# Looking for Therapeutic Antibodies in Next Generation Sequencing Repositories

**DOI:** 10.1101/572958

**Authors:** Konrad Krawczyk, Matthew Raybould, Aleksandr Kovaltsuk, Charlotte M. Deane

## Abstract

Recently it has become possible to query the great diversity of natural antibody repertoires using Next Generation Sequencing (NGS). These methods are capable of producing millions of sequences in a single experiment. Here we compare Clinical Stage Therapeutic antibodies to the ∼1b sequences from 60 independent sequencing studies in the Observed Antibody Space Database. Of the 242 post Phase I antibodies, we find 16 with sequence identity matches of 95% or better for both heavy and light chains. There are also 54 perfect matches to therapeutic CDR-H3 regions in the NGS outputs, suggesting a nontrivial amount of convergence between naturally observed sequences and those developed artificially. This has potential implications for both the discovery of antibody therapeutics and the legal protection of commercial antibodies.

## Introduction

Antibodies are proteins in jawed vertebrates that recognize noxious molecules (antigens) for elimination. An organism expresses millions of diverse antibodies to increase the chances that some of them will be able to bind the foreign antigen, initiating the adaptive immune response. This great diversity can now be queried using Next Generation Sequencing (NGS) of B-cell receptor repertoires, enabling rapid collection of millions of antibody sequences from any given individual ^1-3^.

The sequence of a protein, such as an antibody, is one of the chief vehicles to characterize the molecule in the patent claim^4^. Natural sequences however cannot be patented in the USA^5^. The seminal rulings in the Mayo and Prometheus cases set the precedent to reject patent claims to naturally occurring DNA sequences and it was subsequently extended to include all products of nature^6,7^. The large numbers of antibody sequences now becoming available in the public domain raise the possibility of natural sequences being found that are identical to commercial sequences^5^.

This is especially pertinent in the face of large-scale organized efforts to make naturally sourced antibody NGS data^8^ and analytics^9,10^ more accessible^11^. Specifically, we recently created the Observed Antibody Space (OAS) database that curates the NGS antibody data from public archives and makes them available for easy processing^12^. OAS currently holds ∼1b (∼960m heavy chain and ∼60m light chain) sequences from 60 independent studies. The datasets cover multiple organisms (human, mouse, rabbit, camel etc.), individuals, immune states (non-immunized, immunized etc.) and include the deep sequencing of the totality of adaptive repertoires of mice^13^ (246m sequences) and humans^3^ (318m sequences). Here we quantify how close a sequence match to current Clinical Stage Therapeutic (CST) antibody sequences we can find in OAS.

## Results

We used a set of 242 CST antibody sequences^14^. These are all sequences that have passed Phase I of clinical trials. We separately aligned the CST variable regions (VH or VL), combination of the three Complementarity Determining Regions (CDRs) from VH or VL and CDR-H3s to all the sequences in OAS (see methods). We performed the search across all organisms, individuals and immune states to be comprehensive and to reflect the myriad antibody types (humanized, fully human, chimeric or mouse^15^). The individual identities of the CSTs with respect to the best match from OAS are given in Figure 1 and Table 1 and their distributions are plotted in Figure 2. The aligned sequences are available in the Supplementary Material and on our website http://naturalantibody.com/therapeutics.

**Table 1.** Best sequence identities of Clinical Stage Therapeutic (CST) antibodies to sequences found in public NGS repositories. Sequence identities are given for the best alignment of a sequence from a public repository to a CST heavy or light chain variable region, heavy or light CDR region or CDR-H3 alone (IMGT-defined). The CSTs are identified by their names in the leftmost column. The entries are sorted from top to bottom by the highest heavy chain identity. An interactive version of this table together with aligned sequences are available at http://naturalantibody.com/therapeutics.

| CST Name | Best Heavy Chain Identity (%) | Best Light Identity (%) | Best Heavy CDRs Identity (%) | Best Light CDRs Identity (%) | Best CDR-H3 Identity (%) |
| --- | --- | --- | --- | --- | --- |
| *Enfortumab* | 98 | 98 | 96 | 100 | 100 |
| *Racotumomab* | 97 | 100 | 90 | 100 | 92 |
| *Tabalumab* | 97 | 99 | 96 | 100 | 100 |
| *Emapalumab* | 97 | 99 | 93 | 95 | 87 |
| *Tremelimumab* | 97 | 97 | 94 | 94 | 88 |
| *Ascrinvacumab* | 96 | 100 | 96 | 100 | 100 |
| *Derlotuximab* | 96 | 100 | 89 | 100 | 92 |
| *Zolbetuximab* | 96 | 100 | 88 | 100 | 81 |
| *Ganitumab* | 96 | 99 | 92 | 100 | 91 |
| *Rilotumumab* | 96 | 98 | 93 | 94 | 100 |
| *Durvalumab* | 96 | 98 | 90 | 94 | 92 |
| *Patritumab* | 96 | 97 | 92 | 95 | 90 |
| *Brazikumab* | 96 | 96 | 90 | 95 | 94 |
| *Carotuximab* | 95 | 100 | 85 | 100 | 77 |
| *Varlilumab* | 95 | 98 | 89 | 100 | 91 |
| *Brodalumab* | 95 | 96 | 88 | 100 | 100 |
| *Futuximab* | 95 | 92 | 87 | 88 | 81 |
| *Ramucirumab* | 95 | 87 | 100 | 88 | 100 |
| *Zanolimumab* | 94 | 99 | 100 | 100 | 100 |
| *Foravirumab* | 94 | 98 | 89 | 100 | 100 |
| *Dusigitumab* | 94 | 97 | 100 | 86 | 100 |
| *Rituximab* | 94 | 97 | 90 | 94 | 85 |
| *Muromonab* | 94 | 97 | 82 | 100 | 83 |
| *Ublituximab* | 94 | 96 | 96 | 88 | 100 |
| *Dectrekumab* | 94 | 96 | 93 | 95 | 100 |
| *Necitumumab* | 94 | 95 | 93 | 94 | 92 |
| *Cixutumumab* | 94 | 94 | 89 | 85 | 82 |
| *Fasinumab* | 94 | 93 | 89 | 88 | 83 |
| *Sifalimumab* | 93 | 100 | 88 | 100 | 100 |
| *Modotuximab* | 93 | 100 | 82 | 100 | 91 |
| *Golimumab* | 93 | 99 | 88 | 94 | 94 |
| *Brentuximab* | 93 | 98 | 96 | 100 | 100 |
| *Suvratoxumab* | 93 | 98 | 87 | 94 | 87 |
| *Zalutumumab* | 93 | 98 | 85 | 100 | 88 |
| *Bavituximab* | 93 | 98 | 82 | 94 | 92 |
| *Basiliximab* | 93 | 97 | 88 | 93 | 90 |
| *Radretumab* | 93 | 96 | 80 | 84 | 100 |
| *Ofatumumab* | 92 | 100 | 90 | 100 | 93 |
| *Bezlotoxumab* | 92 | 100 | 89 | 100 | 91 |
| *Daratumumab* | 92 | 100 | 83 | 100 | 86 |
| *Inclacumab* | 92 | 100 | 75 | 100 | 88 |
| *Siltuximab* | 92 | 99 | 89 | 100 | 91 |
| *Canakinumab* | 92 | 99 | 85 | 100 | 100 |
| *Lirilumab* | 92 | 99 | 84 | 100 | 87 |
| *Abrilumab* | 92 | 97 | 85 | 100 | 90 |
| *Tisotumab* | 92 | 97 | 81 | 100 | 81 |
| *Indusatumab* | 92 | 96 | 82 | 100 | 84 |
| *Carlumab* | 92 | 92 | 82 | 70 | 83 |
| *Tovetumab* | 92 | 90 | 86 | 89 | 92 |
| *Utomilumab* | 92 | 89 | 88 | 55 | 100 |
| *Tesidolumab* | 92 | 87 | 92 | 65 | 100 |
| *Glembatumumab* | 91 | 99 | 92 | 100 | 100 |
| *Ipilimumab* | 91 | 99 | 88 | 100 | 90 |
| *Iratumumab* | 91 | 98 | 85 | 100 | 100 |
| *Cetuximab* | 91 | 97 | 82 | 94 | 92 |
| *Burosumab* | 91 | 97 | 80 | 94 | 90 |
| *Anifrolumab* | 91 | 96 | 84 | 89 | 90 |
| *Pritoxaximab* | 91 | 96 | 80 | 100 | 80 |
| *Seribantumab* | 91 | 95 | 78 | 95 | 83 |
| *Girentuximab* | 91 | 95 | 78 | 88 | 91 |
| *Guselkumab* | 91 | 94 | 80 | 82 | 90 |
| *Lenzilumab* | 91 | 91 | 78 | 83 | 83 |
| *Abagovomab* | 91 | 90 | 89 | 94 | 100 |
| *Domagrozumab* | 91 | 89 | 92 | 100 | 88 |
| *Briakinumab* | 91 | 88 | 87 | 65 | 75 |
| *Otelixizumab* | 91 | 71 | 82 | 75 | 83 |
| *Intetumumab* | 90 | 100 | 85 | 100 | 91 |
| *Icrucumab* | 90 | 100 | 82 | 100 | 78 |
| *Foralumab* | 90 | 100 | 81 | 100 | 90 |
| *Fulranumab* | 90 | 100 | 78 | 100 | 93 |
| *Aducanumab* | 90 | 100 | 78 | 100 | 88 |
| *Sarilumab* | 90 | 99 | 88 | 100 | 100 |
| *Bleselumab* | 90 | 98 | 80 | 100 | 84 |
| *Tezepelumab* | 90 | 98 | 80 | 100 | 80 |
| *Opicinumab* | 90 | 98 | 77 | 100 | 90 |
| *Panitumumab* | 90 | 97 | 89 | 94 | 90 |
| *Tomuzotuximab* | 90 | 97 | 82 | 94 | 92 |
| *Timolumab* | 90 | 97 | 80 | 100 | 100 |
| *Adalimumab* | 90 | 97 | 80 | 94 | 71 |
| *Figitumumab* | 90 | 96 | 91 | 100 | 88 |
| *Evolocumab* | 90 | 96 | 91 | 90 | 100 |
| *Berlimatoxumab* | 90 | 95 | 89 | 83 | 90 |
| *Tralokinumab* | 90 | 95 | 80 | 85 | 80 |
| *Ensituximab* | 90 | 94 | 81 | 94 | 85 |
| *Anetumab* | 90 | 92 | 82 | 73 | 84 |
| *Setrusumab* | 90 | 91 | 84 | 78 | 90 |
| *Itolizumab* | 90 | 90 | 82 | 88 | 83 |
| *Ianalumab* | 90 | 88 | 78 | 73 | 71 |
| *Elotuzumab* | 90 | 87 | 96 | 100 | 100 |
| *Emibetuzumab* | 90 | 87 | 87 | 94 | 100 |
| *Evinacumab* | 89 | 100 | 91 | 100 | 94 |
| *Eldelumab* | 89 | 100 | 81 | 100 | 94 |
| *Nivolumab* | 89 | 100 | 77 | 100 | 100 |
| *Avelumab* | 89 | 100 | 75 | 100 | 84 |
| *Denosumab* | 89 | 98 | 87 | 100 | 80 |
| *Atidortoxumab* | 89 | 98 | 67 | 88 | 83 |
| *Setoxaximab* | 89 | 96 | 85 | 100 | 91 |
| *Drozitumab* | 89 | 96 | 80 | 90 | 85 |
| *Indatuximab* | 89 | 95 | 87 | 94 | 100 |
| *Tarextumab* | 89 | 94 | 75 | 89 | 75 |
| *Amatuximab* | 89 | 93 | 82 | 94 | 100 |
| *Infliximab* | 89 | 93 | 75 | 83 | 90 |
| *Lorvatuzumab* | 89 | 92 | 88 | 86 | 100 |
| *Bimagrumab* | 89 | 92 | 87 | 73 | 100 |
| *Solanezumab* | 89 | 92 | 80 | 91 | 100 |
| *Mavrilimumab* | 89 | 91 | 72 | 73 | 61 |
| *Camrelizumab* | 89 | 90 | 92 | 88 | 100 |
| *Tigatuzumab* | 89 | 87 | 89 | 100 | 83 |
| *Anrukinzumab* | 89 | 87 | 85 | 90 | 91 |
| *Urelumab* | 88 | 100 | 80 | 100 | 86 |
| *Secukinumab* | 88 | 100 | 80 | 100 | 80 |
| *Olaratumab* | 88 | 100 | 77 | 100 | 78 |
| *Erenumab* | 88 | 99 | 71 | 100 | 82 |
| *Alirocumab* | 88 | 96 | 85 | 95 | 90 |
| *Gantenerumab* | 88 | 94 | 68 | 89 | 63 |
| *Orticumab* | 88 | 92 | 73 | 77 | 78 |
| *Crenezumab* | 88 | 91 | 95 | 100 | 100 |
| *Concizumab* | 88 | 91 | 80 | 95 | 85 |
| *Bapineuzumab* | 88 | 91 | 75 | 100 | 83 |
| *Actoxumab* | 87 | 100 | 83 | 100 | 86 |
| *Dupilumab* | 87 | 97 | 76 | 95 | 72 |
| *Rafivirumab* | 87 | 95 | 75 | 83 | 70 |
| *Margetuximab* | 87 | 94 | 82 | 94 | 84 |
| *Trevogrumab* | 87 | 94 | 79 | 88 | 69 |
| *Dinutuximab* | 87 | 90 | 86 | 95 | 83 |
| *Mirvetuximab* | 87 | 90 | 77 | 100 | 90 |
| *Olendalizumab* | 87 | 88 | 75 | 100 | 92 |
| *Quilizumab* | 87 | 86 | 88 | 91 | 100 |
| *Obiltoxaximab* | 87 | 85 | 100 | 100 | 100 |
| *Lampalizumab* | 87 | 83 | 79 | 94 | 75 |
| *Pamrevlumab* | 86 | 100 | 82 | 100 | 92 |
| *Fletikumab* | 86 | 100 | 80 | 100 | 85 |
| *Lanadelumab* | 86 | 100 | 67 | 100 | 73 |
| *Ustekinumab* | 86 | 99 | 78 | 100 | 83 |
| *Teprotumumab* | 86 | 98 | 85 | 100 | 90 |
| *Refanezumab* | 86 | 96 | 80 | 100 | 73 |
| *Galiximab* | 86 | 94 | 58 | 90 | 63 |
| *Coltuximab* | 86 | 92 | 96 | 86 | 100 |
| *Ibalizumab* | 86 | 92 | 87 | 95 | 80 |
| *Isatuximab* | 86 | 91 | 89 | 94 | 92 |
| *Otlertuzumab* | 86 | 90 | 92 | 77 | 88 |
| *Rovalpituzumab* | 86 | 90 | 88 | 94 | 90 |
| *Landogrozumab* | 86 | 89 | 81 | 89 | 100 |
| *Daclizumab* | 86 | 87 | 92 | 88 | 100 |
| *Etaracizumab* | 86 | 87 | 84 | 88 | 90 |
| *Enokizumab* | 86 | 87 | 80 | 72 | 86 |
| *Robatumumab* | 86 | 87 | 77 | 100 | 91 |
| *Tislelizumab* | 86 | 86 | 88 | 83 | 91 |
| *Lacnotuzumab* | 86 | 85 | 88 | 94 | 90 |
| *Panobacumab* | 85 | 100 | 84 | 100 | 80 |
| *Fezakinumab* | 85 | 96 | 70 | 95 | 71 |
| *Fresolimumab* | 85 | 95 | 62 | 89 | 84 |
| *Romosozumab* | 85 | 93 | 84 | 100 | 81 |
| *Dalotuzumab* | 85 | 91 | 80 | 100 | 90 |
| *Imgatuzumab* | 85 | 90 | 68 | 76 | 92 |
| *Bococizumab* | 85 | 89 | 77 | 83 | 81 |
| *Atezolizumab* | 85 | 89 | 77 | 77 | 90 |
| *Visilizumab* | 85 | 88 | 89 | 100 | 100 |
| *Lodelcizumab* | 85 | 88 | 70 | 70 | 90 |
| *Lintuzumab* | 85 | 87 | 96 | 100 | 100 |
| *Bimekizumab* | 85 | 84 | 67 | 66 | 66 |
| *Veltuzumab* | 85 | 82 | 90 | 94 | 92 |
| *Rozanolixizumab* | 85 | 82 | 73 | 82 | 80 |
| *Codrituzumab* | 84 | 91 | 83 | 91 | 87 |
| *Plozalizumab* | 84 | 91 | 73 | 100 | 87 |
| *Simtuzumab* | 84 | 90 | 92 | 100 | 100 |
| *Mogamulizumab* | 84 | 88 | 67 | 78 | 75 |
| *Til-drakizumab* | 84 | 87 | 92 | 100 | 100 |
| *Gevokizumab* | 84 | 86 | 79 | 88 | 75 |
| *Sacituzumab* | 84 | 85 | 96 | 94 | 100 |
| *Gedivumab* | 83 | 93 | 67 | 80 | 55 |
| *Obinutuzumab* | 83 | 91 | 78 | 100 | 83 |
| *Ozanezumab* | 83 | 90 | 90 | 100 | 83 |
| *Ixekizumab* | 83 | 90 | 78 | 91 | 75 |
| *Abituzumab* | 83 | 89 | 85 | 100 | 90 |
| *Trastuzumab* | 83 | 89 | 82 | 94 | 84 |
| *Etrolizumab* | 83 | 89 | 76 | 72 | 100 |
| *Ponezumab* | 83 | 89 | 64 | 78 | 77 |
| *Matuzumab* | 83 | 85 | 83 | 88 | 92 |
| *Motavizumab* | 83 | 85 | 75 | 88 | 83 |
| *Inebilizumab* | 83 | 84 | 90 | 90 | 92 |
| *Lifastuzumab* | 83 | 84 | 65 | 78 | 76 |
| *Tanezumab* | 82 | 91 | 80 | 83 | 86 |
| *Olokizumab* | 82 | 90 | 65 | 72 | 81 |
| *Ocrelizumab* | 82 | 88 | 93 | 94 | 93 |
| *Sirukumab* | 82 | 88 | 75 | 82 | 83 |
| *Andecaliximab* | 82 | 85 | 87 | 77 | 100 |
| *Palivizumab* | 82 | 84 | 86 | 94 | 100 |
| *Lumiliximab* | 81 | 94 | 59 | 83 | 88 |
| *Tocilizumab* | 81 | 92 | 82 | 100 | 83 |
| *Galcanezumab* | 81 | 90 | 75 | 83 | 83 |
| *Duligotuzumab* | 81 | 90 | 63 | 77 | 78 |
| *Roledumab* | 81 | 89 | 68 | 94 | 73 |
| *Vadastuximab* | 81 | 88 | 88 | 100 | 100 |
| *Vedolizumab* | 81 | 88 | 86 | 95 | 85 |
| *Mirikizumab* | 81 | 88 | 83 | 77 | 87 |
| *Natalizumab* | 81 | 87 | 90 | 100 | 100 |
| *Eculizumab* | 81 | 87 | 83 | 100 | 86 |
| *Pinatuzumab* | 81 | 86 | 89 | 86 | 100 |
| *Ficlatuzumab* | 81 | 86 | 81 | 88 | 90 |
| *Eptinezumab* | 81 | 80 | 70 | 29 | 100 |
| *Belimumab* | 80 | 98 | 62 | 100 | 62 |
| *Crizanlizumab* | 80 | 91 | 90 | 86 | 93 |
| *Depatuxizumab* | 80 | 88 | 76 | 94 | 88 |
| *Pertuzumab* | 80 | 88 | 75 | 83 | 91 |
| *Ligelizumab* | 80 | 88 | 71 | 88 | 81 |
| *Blosozumab* | 80 | 88 | 66 | 88 | 81 |
| *Ravulizumab* | 80 | 87 | 77 | 100 | 86 |
| *Fremanezumab* | 80 | 87 | 67 | 77 | 53 |
| *Clazakizumab* | 80 | 87 | 65 | 57 | 78 |
| *Pembrolizumab* | 80 | 86 | 86 | 90 | 84 |
| *Inotuzumab* | 80 | 82 | 80 | 95 | 100 |
| *Pidilizumab* | 80 | 82 | 76 | 94 | 90 |
| *Vatelizumab* | 80 | 79 | 82 | 88 | 92 |
| *Benralizumab* | 79 | 89 | 83 | 83 | 71 |
| *Certolizumab* | 79 | 87 | 81 | 100 | 100 |
| *Lebrikizumab* | 79 | 85 | 74 | 95 | 91 |
| *Epratuzumab* | 79 | 84 | 84 | 95 | 88 |
| *Satralizumab* | 79 | 84 | 71 | 72 | 83 |
| *Risankizumab* | 79 | 83 | 82 | 83 | 84 |
| *Reslizumab* | 78 | 89 | 92 | 77 | 100 |
| *Onartuzumab* | 78 | 85 | 78 | 87 | 75 |
| *Farletuzumab* | 78 | 82 | 96 | 90 | 100 |
| *Bevacizumab* | 77 | 93 | 90 | 88 | 93 |
| *Vonlerolizumab* | 77 | 92 | 65 | 94 | 80 |
| *Idarucizumab* | 77 | 91 | 83 | 95 | 87 |
| *Polatuzumab* | 77 | 90 | 80 | 95 | 80 |
| *Rontalizumab* | 77 | 88 | 76 | 95 | 90 |
| *Parsatuzumab* | 77 | 86 | 81 | 82 | 93 |
| *Gemtuzumab* | 77 | 83 | 80 | 86 | 88 |
| *Spartalizumab* | 77 | 83 | 76 | 91 | 90 |
| *Efalizumab* | 76 | 94 | 83 | 100 | 85 |
| *Alemtuzumab* | 76 | 90 | 80 | 66 | 91 |
| *Dacetuzumab* | 76 | 84 | 82 | 91 | 85 |
| *Tregalizumab* | 76 | 84 | 72 | 100 | 93 |
| *Omalizumab* | 75 | 90 | 76 | 100 | 71 |
| *Nimotuzumab* | 75 | 81 | 68 | 95 | 62 |
| *Pateclizumab* | 74 | 91 | 81 | 88 | 81 |
| *Teplizumab* | 74 | 82 | 82 | 100 | 83 |
| *Ranibizumab* | 73 | 92 | 81 | 88 | 93 |
| *Mepolizumab* | 72 | 92 | 78 | 95 | 84 |
| *Ontuxizumab* | 69 | 85 | 78 | 84 | 82 |

**Figure 1.**
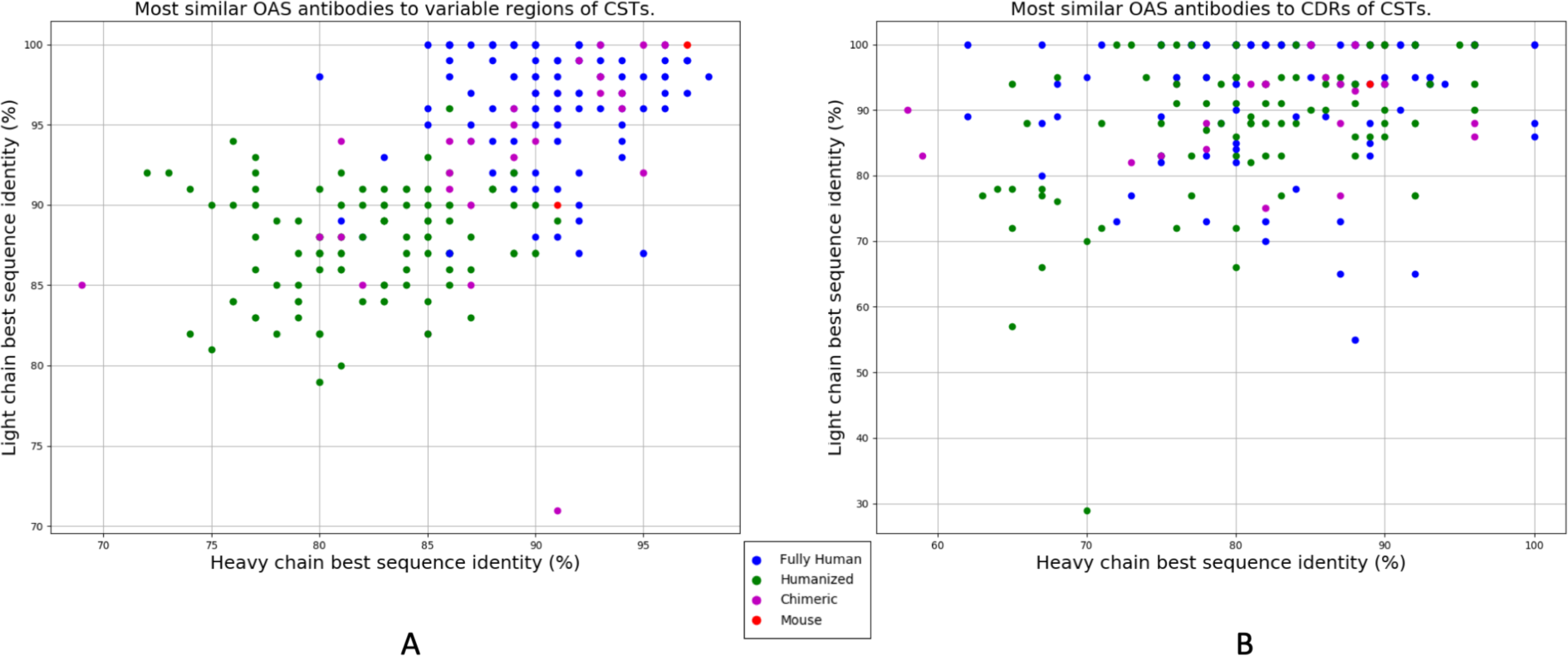
Best sequence identity matches to Clinical Stage Therapeutics (CST) in naturally sourced NGS datasets. **A)** Heavy and light chain variable regions of 242 CST sequences from Raybould et al.^14^ aligned to variable region sequences in OAS^12^. **B)** Heavy and light chain IMGT CDR regions of 242 CS?s aligned to IMGT CDR regions in OAS. Fully human sequences are denoted by blue dots, humanized by green and chimeric by magenta. The two mouse sequences are shown in red. Interactive versions of these charts are available at http://naturalantibody.com/therapeutics.

**Figure 2.**
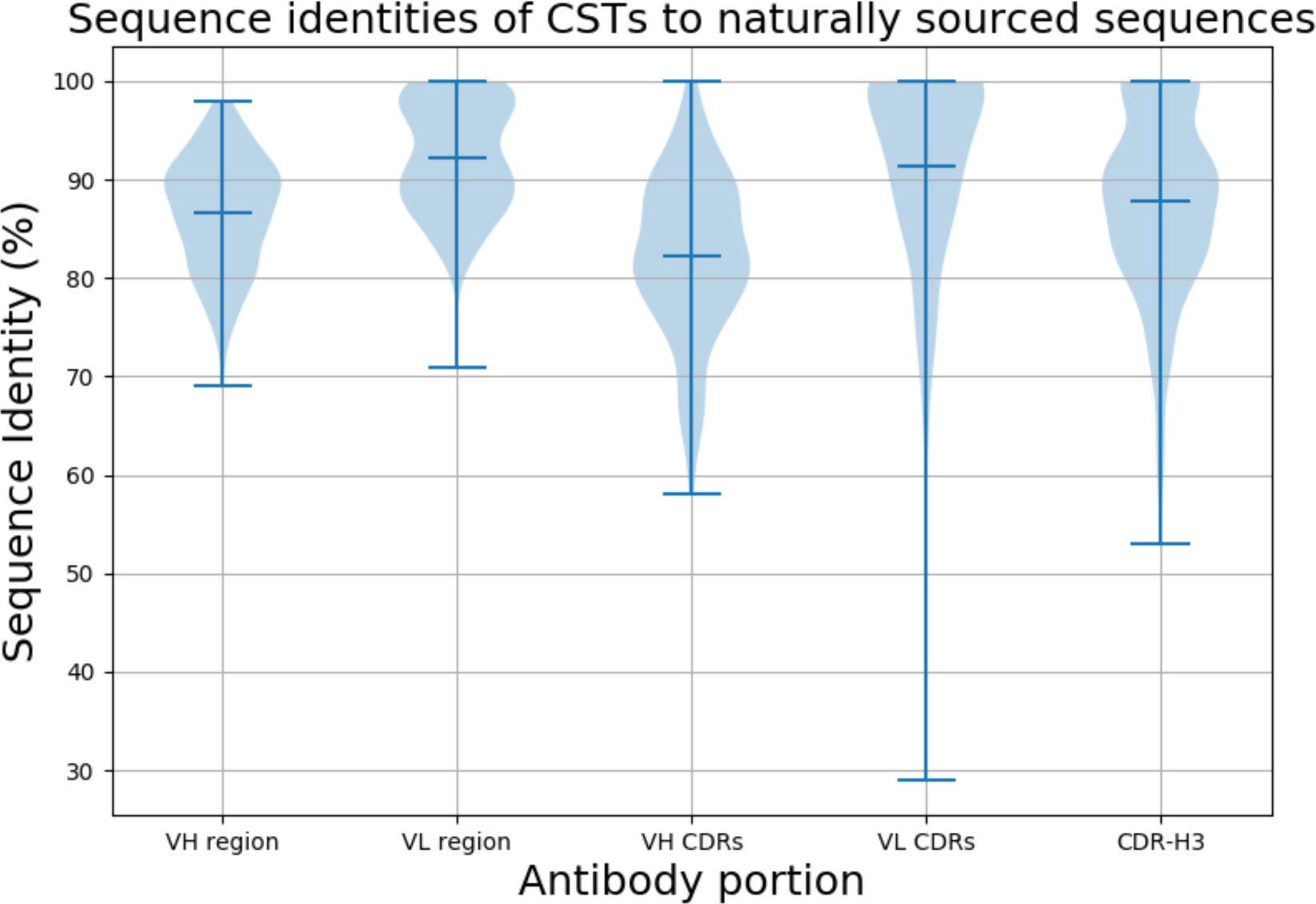
Distribution of sequence identity matches of Clinical Stage Therapeutics (CSTs) to naturally-sourced NGS. The violin plots show the distribution of sequence identities of the variable heavy (VH) and light (VL) chains, heavy and light CDR regions and CDR-H3 of CSTs to best matches in OAS.

### Analysis of Clinical Stage Therapeutic sequence matches to naturally sourced NGS datasets

The best sequence identity matches of CST variable regions to naturally sourced NGS datasets in OAS are given in Figure 1A. Ninety (37.1%) CST heavy chains have matches within OAS of ≥ 90% sequence identity (seqID), with 18 (7.4%) ≥ 95% seqID. We find 158 (65.2%) therapeutic light chains with ≥ 90% seqID to an OAS sequence, with 96 (39.7%) ≥ 95% seqID, and 28 (11.5%) with 100% seqID. For 16 (6.6%) of the CSTs we find both heavy and light chain matches ≥ 95% seqID. In the most extreme case, Enfortumab, we were able to find both heavy and light chain matches of 98% seqID (the differences are H38:N-S, H88:S-Y, L37:G-S, L52:F-L where the first amino acid comes from Enfortumab and the second from an OAS sequence).

The largest discrepancy between the CSTs and any given natively-expressed antibody is typically concentrated in the CDR regions that determine antigen complementarity^16^. It remains unclear, however, the extent to which the highly mutable CDR loops of engineered therapeutics differ from those that are expressed naturally. We searched for the best CST matches to the CDR regions in OAS. The search was performed using IMGT-defined CDR triplets from the heavy or light chain, disregarding the framework region (Table 1, Figure 1B and Figure 2). We find 46 (19.0%) of CST heavy chain CDR regions to have matches to an OAS CDR region with ≥ 90% seqID, 15 (6.1%) with ≥ 95% seqID and 4 (1.6%) with 100% seqID. There were 156 (64.4%) CST light CDR regions with ≥ 90% seqID to an OAS CDR region, with 110 (45.4%) ≥ 95% seqID, and 90 (37.1%) with 100% seqID. We found perfect matches for both light and heavy chain CDRs in two CSTs, Obiltoxaximab and Zanolimumab.

Of the six Complementarity Determining Regions, CDR-H3 is the most sequence and structurally diverse^17,18^. Due to its key role in binding, it is subjected to extensive antibody engineering^19,20^. We checked how likely it is to find CST-derived CDR-H3s in naturally sourced sequences. To assess this, we searched for the best CST CDR-H3 matches in OAS, regardless of the framework region and remaining CDRs (Table 1, Figure 2). Of our 242 CST CDR-H3s, we found 54 perfect matches in OAS. The perfect matches tended to be for shorter CDR-H3s, however some longer loops with perfect matches were also found (see Supplementary Section 1). Twenty-nine perfect matches were found in the recent deep sequencing dataset of Briney et al. (2019)^3^, suggesting that a single comprehensive NGS study can cover a significant amount of CDR-H3 diversity. Forty-seven perfect matches were found in OAS datasets other than that of Briney et al. (2019), showing that certain artificial CDR-H3 sequences can be independently observed in naturally sourced NGS. Twenty-two CDR-H3 matches were found in both Briney et al. data and other OAS datasets. These 22 shared sequences come from 9 humanized and 13 fully human CSTs. The 54 perfect CDR-H3 matches were distributed among all antibody types, with 23 humanized, 22 fully human, 8 chimeric and 1 mouse (21.9%, 22.0%, 22.8% and 50.0% of each category, respectively). These results show that, despite the large theoretical sequence space accessible to the CDR-H3 region^3^, therapeutically-exploitable CDR-H3 loops are found in just ∼960m heavy chain sequences from 60 NGS studies (see Supplementary Section 2). This convergence, coupled with the fact that CDR-H3 loops often mediate antibody specificity^21^ and binding affinity, could suggest intrinsically driven biases in antigen recognition^22^, independent of artificial discovery methods.

### Stratifying the best CST matches in OAS by antibody type

The quality of the variable region match we could find for any given CST sequence appears to be highly dependent on the discovery platform/antibody type. Figure 3 suggests that antibodies produced via more artificial protocols such as humanization have lower variable region sequence identities to sequences in OAS from those of fully human molecules. For the majority of the fully human sequences we find matches of 90% seqID or better whereas matches to the majority of humanized molecules fall below 90% seqID (Figure 3). Chimeric antibodies appear to have seqID values intermediate between the two classes (Figure 3).

**Figure 3.**
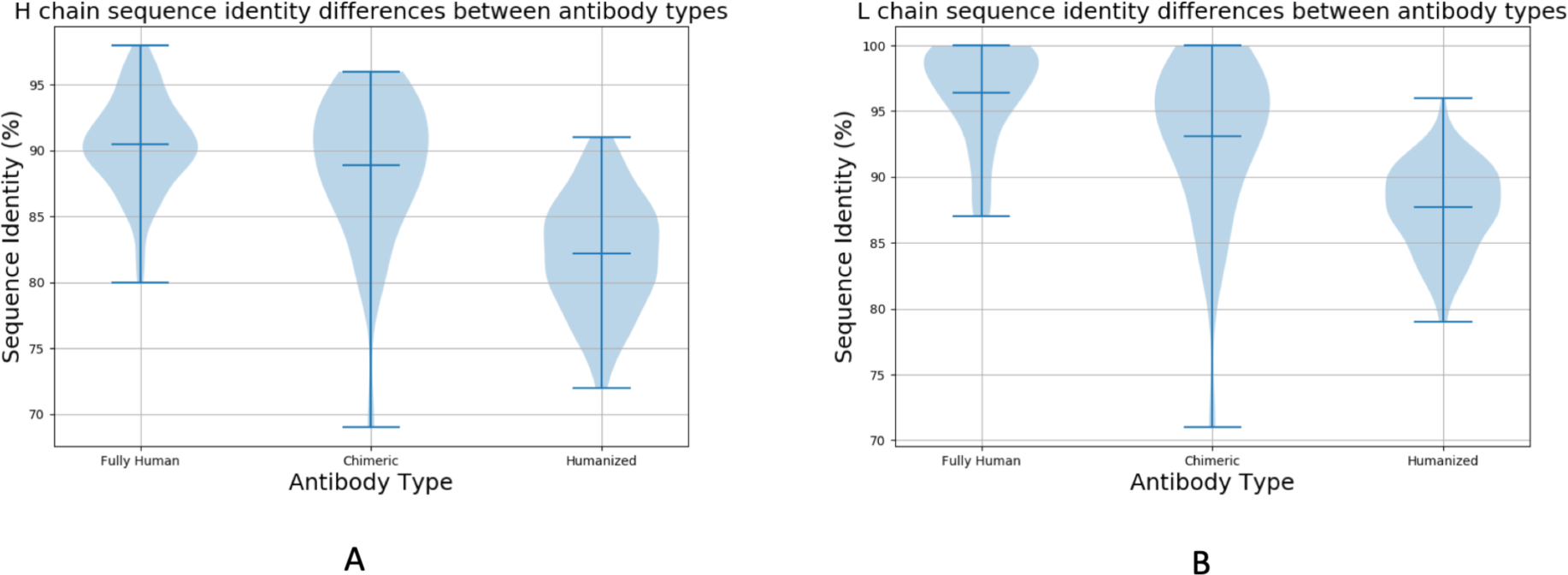
Sequence identity matches of Clinical Stage Therapeutic (CST) variable regions to naturally sourced NGS datasets stratified by CST antibody type. CST **A)** heavy chain and **B)** light chain identities to NGS sequences in OAS stratified by Fully Human, Chimeric and Humanized antibody types. The two mouse molecules were omitted as too small a sample.

The CST antibody type also reflects the organism that produced the best NGS seqID match. Of the 100 fully human CSTs, the 90 (90.0%) most similar heavy chains, 100 (100.0%) most similar light chains, and 55 (55.0%) most similar CDR-H3 loops come from human-sourced NGS. Of the 105 humanized antibodies, 82 (78.0 %) of heavy chains, and 79 (75.2%) of light chains found closest matches in human-sourced NGS, while 71 (67.6%) of the best CDR-H3s matches were identified in mouse-sourced NGS. This further reflects the dominance of CDR-H3 in binding, as therapeutic companies tend to graft this loop from binding mouse antibodies to transfer specificity and binding affinity. It also suggests that data-mining a dataset such as OAS could provide a more accurate measure of antibody ‘humanness’ than our current metrics^23,24^.

## Discussion

Our results demonstrate that despite the theoretically large diversity allowed to antibodies^3,25^, there exists a nontrivial convergence between artificially developed CSTs and naturally sourced NGS sequences. The closest NGS matches to CSTs were sourced from 48 of the 60 (80.0%) independent datasets available in OAS indicating that finding a close match to at least one CST is likely in most NGS datasets.

It was previously suggested that such an overlap could cause issues in claiming patents on therapeutic antibodies^5^. Until a legal opinion on the subject is available, it will remain unclear if NGS data poses potential problems for the patentability of monoclonal antibodies. Firstly, sequence is one of the main ways to characterize a molecule in a patent claim but by itself it does not offer information about its cognate antigen and therapeutic action. Such information however is crucial in patent application to demonstrate novelty of the invention^4^. NGS studies produce copious amounts of sequences but they do not alone relate them to any target molecule. Secondly, the antibody variable region is a product of two polypeptide chains (heavy and light) and its function is intimately related to this combination. Currently the majority of available NGS datasets report heavy and light chains separately. Thirdly, NGS outputs are known to have high error rates^26^ and lastly, it is unclear how close a sequence-identity match to a publicly available sequence or its portion (such as CDR-H3) would cause issues in establishing the inventiveness of a sequence. For instance, there exist only four pairs of CSTs with heavy chain sequence identities of more than 94% (see supplementary section 3). In three of such high sequence-identity pairs, both sequences come from the same company and the fourth is the original patent-expired antibody and its derivative. This is compared to 18 therapeutic heavy chains with matches to OAS better than 95%.

In light of ongoing efforts to further consolidate antibody NGS data and make it more accessible, it follows that finding therapeutic candidate sequences in published NGS datasets will become easier^11,27^.

## Methods

We used the Observed Antibody Space database as the source of NGS sequences. Since its first release, the database has been expanded by four more datasets, most notably the recent deep sequencing of human antibody repertoire by Briney et al. 2019^3^. We employed the processed consensus sequences from Briney et al., removing any sequences with stop codons as these were deemed unproductive.

We used the 242 antibodies from Raybould et al.^14^ as the source of Clinical Stage Therapeutic (CST) antibodies. We numbered the CST sequences according to the IMGT^28^ scheme using ANARCI^29^. The CST sequences were classified into four groups, based on their International Nonproprietary Names^15,30^. Sequences with names containing ‘-xizumab’, ‘-ximab’ or ‘-monab’ were labeled as ‘chimeric’. Sequences not matching this criterion but containing ‘-zumab’ in their name were classified as ‘humanized’. Sequences that contained only ‘-umab’ in their name were labeled as ‘fully human’. Two mouse antibodies (Abagovomab and Racotumomab), were labeled as ‘mouse’.

We separately aligned the heavy chain, light chain, the combination of the three heavy or light chains IMGT-defined CDRs and the IMGT-defined CDR-H3 of CSTs to each of the sequences in OAS^12^. We note a match if an IMGT position in a ‘query’ CST is also found in a ‘template’ sequence from OAS, and they have the same amino acid residue. For the full sequence alignments, the number of matches is divided by the length of the query and by the length of the template, producing two sequence identities. The final sequence identity is the average between these two. Calculating the sequence identity in this way prevents the scenario when one sequence is a substring of another, creating an artificially high sequence identity with a large length discrepancy. The CDR alignments were performed when the IMGT-defined loop lengths matched. The aligned sequences are available in the supplementary section 4 and through an interactive version of Figure 1 and Table 1 accessible at http://naturalantibody.com/therapeutics.

