## Supplementary Information for "Looking for Therapeutic Antibodies in Next Generation Sequencing Repositories"

<u>**Supplementary tables and figures**</u>

### Section 1. Distribution of CDR-H3 lengths in CSTs as compared to perfect matches in OAS.

We found 54 perfect matches to CST CDR-H3s in OAS. We compared the CDR-H3 lengths of the perfect matches to all CDR-H3 lengths of our 242 CSTs (Supplementary Figure 1 and Supplementary Table 1). The mean length of all CST CDR-H3 is 12 whereas this of the 54 perfect matches is 10. As expected, it is easier to find perfect matches for shorter CDR-H3s, however some longer lengths were also covered (Supplementary Figure 1 and Supplementary Table 1).

Of the 54 perfect matches, 22 can be found in the deep sequencing dataset of Birney et al. 2019 and other OAS datasets. The mean length of these shared CDR-H3s is 9, further indicating that finding perfect matches across independent datasets is easier for shorter loop lengths.

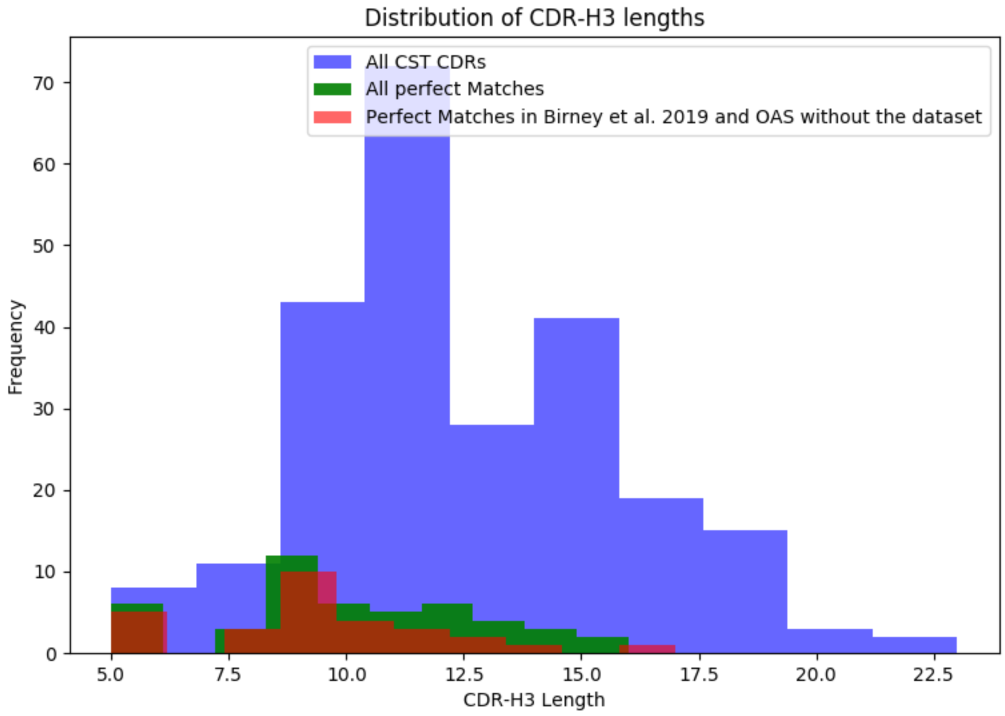


**Supplementary Figure 1.** Distribution of CST CDR-H3 lengths (blue) overlaid on the lengths of the perfect matches (green). The red histogram shows the length distribution of CST CDR-H3 that can be found in the deep sequencing dataset of Birney et al. 2019 and in other datasets in OAS when Birney et al. 2019 data are removed.

### Section 2. Theoretical estimates of probability of finding perfect matches to CDR-H3 region.

We have estimated the number of theoretically allowed CDR-H3s for each length, assuming that each amino acid is allowed at each position. We calculated the number of possible CDR-H3s for a given length as $20^L$, where L is the length of the loop and 20 represents the number of allowed amino acids. We have also estimated how likely it is to find a single sequence for a given length, assuming 960m independent samples (number of our heavy chain sequences, disregarding H3 redundancy and length stratification in our dataset to be deliberately more permissive) as $9.4x10^8/20^L$ for a given length L. The estimates for each loop length are given in Supplementary Table 1. For length 12, which is the mean length of the CSTs, the number of theoretically allowed CDR-H3s is $4.096x10^{15}$ and the probability of finding a single match in 960m independent samples is in the order of $10^{-7}$, whereas we find seven perfect CST matches for this particular length. The longest CDR-H3 we can find a perfect match for is length 17 and the probability of finding a single sequence here is in the order of $10^{-14}$.

| CDR-H3 length | Number of possible sequences | Length frequency in CSTs | Perfect matches | Probability of finding a single sequence in 960m samples |
|---|---|---|---|---|
| **5** | 3,200,000 | 3 | 3 | 1 |
| **6** | 64,000,000 | 5 | 3 | 1 |
| **7** | 1,280,000,000 | 2 | 0 | 0.75 |
| **8** | 25,600,000,000 | 9 | 3 | 0.0375 |
| **9** | 512,000,000,000 | 22 | 14 | 0.00187 |
| **10** | 10,240,000,000,000 | 21 | 7 | 9.375e-05 |
| **11** | 204,800,000,000,000 | 29 | 6 | 4.6875e-06 |
| **12** | 4,096,000,000,000,000 | 43 | 7 | 2.34375e-07 |
| **13** | 81,920,000,000,000,000 | 28 | 5 | 1.171875e-08 |
| **14** | 1,638,400,000,000,000,000 | 23 | 3 | 5.859375e-10 |
| **15** | 32,768,000,000,000,000,000 | 18 | 1 | 2.9296875e-11 |
| **16** | 655,360,000,000,000,000,000 | 14 | 1 | 1.46484375e-12 |
| **17** | 13,107,200,000,000,000,000,000 | 5 | 1 | 7.32421875e-14 |
| **18** | 262,144,000,000,000,000,000,000 | 7 | 0 | 3.662109375e-15 |
| **19** | 5,242,880,000,000,000,000,000,000 | 8 | 0 | 1.8310546875e-16 |
| **20** | 104,857,600,000,000,000,000,000,000 | 3 | 0 | 9.1552734375e-18 |
| **23** | 838,860,800,000,000,000,000,000,000,000 | 2 | 0 | 1.14440917969e-21 |

**Supplementary Table 1.** Estimated theoretical number of CDR-H3 sequences for each IMGT length and probabilities of finding a single sequence given 940m independent samples.

**Section 3. Quantifying pairwise sequence identities of therapeutic sequences.**

To provide context  practically allowed sequence identities in patent claims, we calculated the identity of each pair of therapeutic sequences in our set of 242 CSTs (Supplementary Figure 2 for heavy chains and Supplementary Figure 3 for light chains). In only four cases is it possible to find heavy chains across two different therapeutics that are more than 94% sequence identical and these are given in Supplementary Table 2. These pairs of therapeutics however are by and large produced by the same company as Ravulizumab and Eculizumab are from Alexion Pharmaceuticals, Ranibizumab and Bevacizumab are from Genentech whereas Palivizumab and Motavizumab are from Medimmune. Tomuzotuximab, by Glycotope, is based on Cetuximab by Bristol Myers Squibb, the patent on which expired several years ago (1).

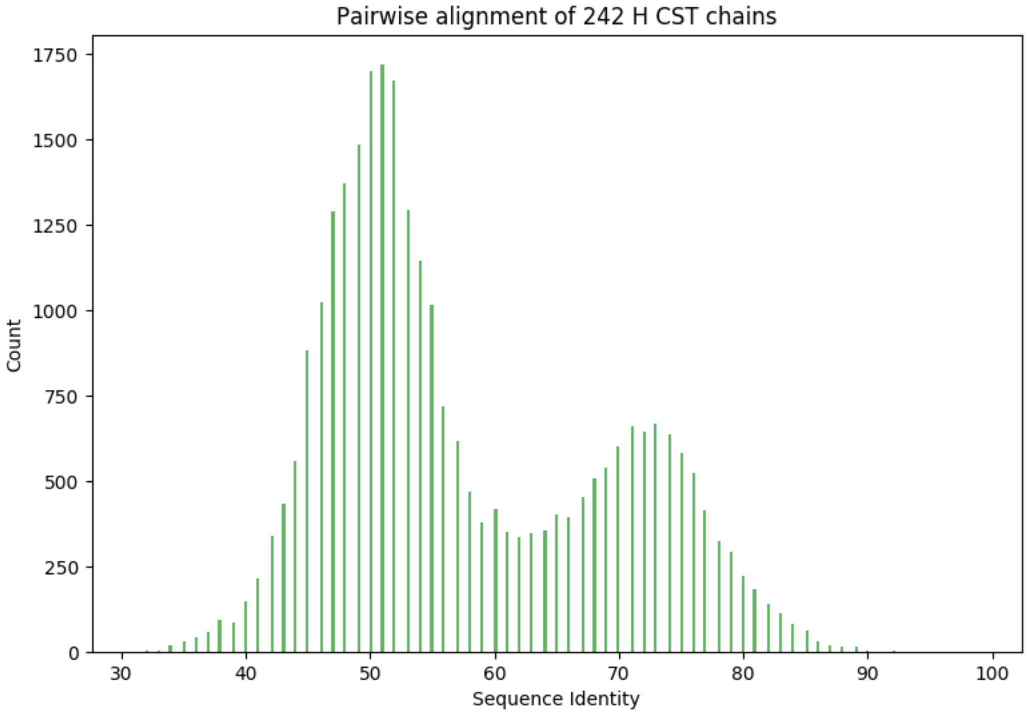


**Supplementary Figure 2.** Histogram of pairwise sequence identities of the 242 CST heavy chains from our dataset.

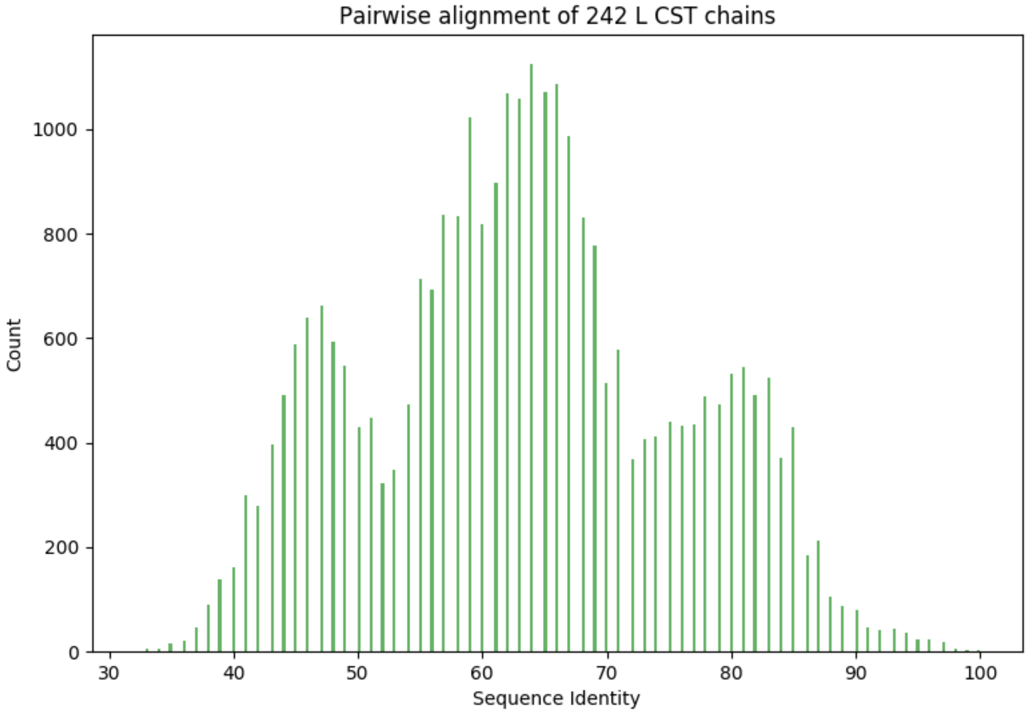


**Supplementary Figure 3**. **.** Histogram of pairwise sequence identities of the 242 CST light chains from our dataset.

| **Therapeutic Antibodies** | **Heavy Chain Sequence Identity** |
|---|---|
| Tomuzotuximab & Cetuximab | 99% |
| Ranibizumab & Bevacizumab | 96% |
| Palivizumab & Motavizumab | 94% |
| Ravulizumab & Eculizumab | 98% |

**Supplementary Table 2.** The pairwise sequence identities of CST heavy that are higher than or equal to 94%.

**Section 4. Alignments between CST antibodies and sequences in OAS.**

Below you can see the precise IMGT-aligned sequences of a CST heavy chain or light chain, heavy and light CDR regions and CDR-H3 alone with the best templates we could find for them in OAS. The alignments are sorted alphabetically by the name of the CST. In each of the alignments, the CST sequence comes first, followed by aligned (|) and unaligned (.) entries of the following sequence in the OAS. IMGT-CDRs are marked by the (^) symbol.

Therapeutic : Abagovomab

Best Alignment of Abagovomab heavy chain to a sequence from OAS
```
QVKLQESGAELARPGASVKLSCKASGYTFTNYWMQWVKQRPGQGLDWIGAIYPGDGNTRYTHKFKGKATLTADKSSSTAYMQLSSLASEDSGVYYCARGEGN-YAWFAYWGQGTTVTVSS
.|||||||||||||||||||||||||||||.||||||||||||||.||||||||||.||||.|||||||||||||||||||||||||||||.||||||||....|||||||||||.||||
-VKLQESGAELARPGASVKLSCKASGYTFTSYWMQWVKQRPGQGLEWIGAIYPGDGDTRYTQKFKGKATLTADKSSSTAYMQLSSLASEDSAVYYCARGEYGNSAWFAYWGQGTTLTVSS
                         ^^^^^^^^                 ^^^^^^^^                                      ^^^^^^^^^^^^^
```

Best Alignment of Abagovomab light chain to a sequence from OAS
```
DIELTQSPASLSASVGETVTITCQASENIYSYLAWHQQKQGKSPQLLVYNAKTLAGGVSSRFSGSGSGTHFSLKIKSLQPEDFGIYYCQHHYGILPTFGGGTKLEIK
||.||||||||||||||||||||.|||||||||||.|||||||||||||||||||.||.||||||||||.|||||.||||||||.||||||||..||||||||||||
DIVLTQSPASLSASVGETVTITCRASENIYSYLAWYQQKQGKSPQLLVYNAKTLAEGVPSRFSGSGSGTQFSLKINSLQPEDFGSYYCQHHYGTPPTFGGGTKLEIK
                          ^^^^^^                 ^^^                                    ^^^^^^^^^
```

Best Alignment of Abagovomab heavy chain CDRs to a sequence from OAS
```
QVKLQESGAELARPGASVKLSCKASGYTFTNYWMQWVKQRPGQGLDWIGAIYPGDGNTRYTHKFKGKATLTADKSSSTAYMQLSSLASEDSGVYYCARGEGNYAWFAYWGQGTTVTVSS
.|.|.||||||.|||.|||.|||||||||||||..|.|||||.||.|||.||||.|.|.|..||||||||||||||||||||.|||.||||..||||||.|||||||||||||.||||.
-VRLVESGAELVRPGTSVKMSCKASGYTFTNYWIGWAKQRPGHGLEWIGDIYPGGGYTNYNEKFKGKATLTADKSSSTAYMQFSSLTSEDSAIYYCARGYGNYAWFAYWGQGTLVTVSA
                         ^^^^^^^^                 ^^^^^^^^                                      ^^^^^^^^^^^^
```

Best Alignment of Abagovomab light CDRs chain to a sequence from OAS
```
DIELTQSPASLSASVGETVTITCQASENIYSYLAWHQQKQGKSPQLLVYNAKTLAGGVSSRFSGSGSGTHFSLKIKSLQPEDFGIYYCQHHYGILPTFGGGTKLEIK
...................||||..||||||||||.|||||||||||||||||||.||.||||||||||.|||||.||||||||.|||||||||.||||.|||||.|
-------------------TITCRPSENIYSYLAWYQQKQGKSPQLLVYNAKTLAEGVPSRFSGSGSGTQFSLKINSLQPEDFGSYYCQHHYGIPPTFGAGTKLELK
                          ^^^^^^                 ^^^                                    ^^^^^^^^^
```

Best Alignment of Abagovomab CDR-H3 chain to a sequence from OAS
```
QVKLQESGAELARPGASVKLSCKASGYTFTNYWMQWVKQRPGQGLDWIGAIYPGDGNTRYTHKFKGKATLTADKSSSTAYMQLSSLASEDSGVYYCARGEGNYAWFAYWGQGTTVTVSS
.|.|.|||..|..||.|.|.||.|||..|..|||.||.|.||.||.|||.|.|......||...|.|.....|....|.|.|.|...|||...||||||||||||||||||||.||||.
EVMLVESGGGLVQPGGSLKPSCAASGFDFSRYWMSWVRQAPGKGLEWIGEINPDSSTINYTPSLKDKFIISRDNAKNTLYLQMSKVRSEDTALYYCARGEGNYAWFAYWGQGTLVTVSA
                         ^^^^^^^^                 ^^^^^^^^                                      ^^^^^^^^^^^^
```

Therapeutic : Abituzumab

Best Alignment of Abituzumab heavy chain to a sequence from OAS
```
QVQLQQSGGELAKPGASVKVSCKASGYTFSSFWMHWVRQAPGQGLEWIGYINPRSGYTEYNEIFRDKATMTTDTSTSTAYMELSSLRSEDTAVYYCASFLGRGAMDYWGQGTTVTVSS
||||||||.||||||||||.|||||||||.|.|||||.|.|||||||||||||.||||.||..|.||||.|.|.|.|||||.||||.|||.||||||.||||||||||||||.|||||
QVQLQQSGAELAKPGASVKLSCKASGYTFTSYWMHWVKQRPGQGLEWIGYINPSSGYTKYNQKFKDKATLTADKSSSTAYMQLSSLTSEDSAVYYCARFLGRGAMDYWGQGTSVTVSS
                         ^^^^^^^^                 ^^^^^^^^                                      ^^^^^^^^^^^
```

Best Alignment of Abituzumab light chain to a sequence from OAS
```
DIQMTQSPSSLSASVGDRVTITCRASQDISNYLAWYQQKPGKAPKLLIYYTSKIHSGVPSRFSGSGSGTDYTFTISSLQPEDIATYYCQQGNTFPYTFGQGTKVEIK
|||||||||||||||||||||||.|||||||||.|||||||||||||||..|....||||||||||||||.|||||||||||||||||||.||||||||||||.|||
DIQMTQSPSSLSASVGDRVTITCQASQDISNYLNWYQQKPGKAPKLLIYDASNLETGVPSRFSGSGSGTDFTFTISSLQPEDIATYYCQQSNTFPYTFGQGTKLEIK
                          ^^^^^^                 ^^^                                    ^^^^^^^^^
```

Best Alignment of Abituzumab heavy chain CDRs to a sequence from OAS
```
QVQLQQSGGELAKPGASVKVSCKASGYTFSSFWMHWVRQAPGQGLEWIGYINPRSGYTEYNEIFRDKATMTTDTSTSTAYMELSSLRSEDTAVYYCASFLGRGAMDYWGQGTTVTVSS
||||||||.||||||||||.|||||||||.|.|||||.|.|||||||||||||.||||.||..|.||||.|.|.|.|||||.||||.|||.||||||.||||||||||||||.|||||
QVQLQQSGAELAKPGASVKLSCKASGYTFTSYWMHWVKQRPGQGLEWIGYINPSSGYTKYNQKFKDKATLTADKSSSTAYMQLSSLTSEDSAVYYCARFLGRGAMDYWGQGTSVTVSS
                         ^^^^^^^^                 ^^^^^^^^                                      ^^^^^^^^^^^
```

Best Alignment of Abituzumab light CDRs chain to a sequence from OAS
```
DIQMTQSPSSLSASVGDRVTITCRASQDISNYLAWYQQKPGKAPKLLIYYTSKIHSGVPSRFSGSGSGTDYTFTISSLQPEDIATYYCQQGNTFPYTFGQGTKVEIK
||.|||..||||||.||||||.|||||||||||.||||||....||||||||..|||||||||||||||||..|||.|..||||||.||||||||||||.|||.|||
DIKMTQTTSSLSASLGDRVTISCRASQDISNYLNWYQQKPDGTVKLLIYYTSRLHSGVPSRFSGSGSGTDYSLTISNLEQEDIATYFCQQGNTFPYTFGGGTKLEIK
                          ^^^^^^                 ^^^                                    ^^^^^^^^^
```

Best Alignment of Abituzumab CDR-H3 chain to a sequence from OAS
```
QVQLQQSGGELAKPGASVKVSCKASGYTFSSFWMHWVRQAPGQGLEWIGYINPRSGYTEYNEIFRDKATMTTDTSTSTAYMELSSLRSEDTAVYYCASFLGRGAMDYWGQGTTVTVSS
.|.|..|||.|.|||.|.|.||.|||.|||...|.||||.|...|||...|.....||.|........|...|......|...|||.|||||.|||||.|||||||||||||.|||||
EVKLMESGGGLVKPGGSLKLSCAASGFTFSDYYMYWVRQTPEKRLEWVATISDGGSYTYYPDSVKGRFTISRDNAKNNLYLQMSSLKSEDTAMYYCASELGRGAMDYWGQGTSVTVSS
                         ^^^^^^^^                 ^^^^^^^^                                      ^^^^^^^^^^^
```

Therapeutic : Abrilumab

Best Alignment of Abrilumab heavy chain to a sequence from OAS
```
QVQLVQSGAEVKKPGASVKVSCKVSGYTLSDLSIHWVRQAPGKGLEWMGGFDPQDGETIYAQKFQGRVTMTEDTSTDTAYMELSSLKSEDTAVYYCATGSSS----SWFDPWGQGTLVTVSS
|||||||||||||||||||||||||||||..||.|||||||||||||||||||.||||||||||||||||||||||||||||||||.|||||||||||.|||.....|||||||||||||||
QVQLVQSGAEVKKPGASVKVSCKVSGYTLTELSMHWVRQAPGKGLEWMGGFDPEDGETIYAQKFQGRVTMTEDTSTDTAYMELSSLRSEDTAVYYCATYSSSWYKKGWFDPWGQGTLVTVSS
                         ^^^^^^^^                 ^^^^^^^^                                      ^^^^^^^^^^^^^^^
```

Best Alignment of Abrilumab light chain to a sequence from OAS
```
DIQMTQSPSSVSASVGDRVTITCRASQGISSWLAWYQQKPGKAPKLLIYGASNLESGVPSRFSGSGSGTDFTLTISSLQPEDFANYYCQQANSFPWTFGQGTKVEIK
||||||||||||||||||||||||||||||||||||||||||||||||||||.|.|||||||||||||||||||||||||||||.||||||||||||||||||||||
DIQMTQSPSSVSASVGDRVTITCRASQGISSWLAWYQQKPGKAPKLLIYGASSLQSGVPSRFSGSGSGTDFTLTISSLQPEDFATYYCQQANSFPWTFGQGTKVEIK
                          ^^^^^^                 ^^^                                    ^^^^^^^^^
```

Best Alignment of Abrilumab heavy chain CDRs to a sequence from OAS
```
QVQLVQSGAEVKKPGASVKVSCKVSGYTLSDLSIHWVRQAPGKGLEWMGGFDPQDGETIYAQKFQGRVTMTEDTSTDTAYMELSSLKSEDTAVYYCATGSSSSWFDPWGQGTLVTVSS
................|||||||||||||..||.|||||||||||||||||||.||||||||||||||||||||||||||||||||.||||||||||||.||||||||||||||||||
----------------SVKVSCKVSGYTLTELSMHWVRQAPGKGLEWMGGFDPEDGETIYAQKFQGRVTMTEDTSTDTAYMELSSLRSEDTAVYYCATGYSSSWFDPWGQGTLVTVSS
                         ^^^^^^^^                 ^^^^^^^^                                      ^^^^^^^^^^^
```

Best Alignment of Abrilumab light CDRs chain to a sequence from OAS

```
DIQMTQSPSSVSASVGDRVTITCRASQGISSWLAWYQQKPGKAPKLLIYGASNLESGVPSRFSGSGSGTDFTLTISSLQPEDFANYYCQQANSFPWTFGQGTKVEIK
.||.||||||||||||||||||||||||||||||||||||||||||||||||.|.|||||||||||||||||||||||||||||.||||||||||||||||||||||
-IQTTQSPSSVSASVGDRVTITCRASQGISSWLAWYQQKPGKAPKLLIYGASSLQSGVPSRFSGSGSGTDFTLTISSLQPEDFATYYCQQANSFPWTFGQGTKVEIK
                          ^^^^^^                 ^^^                                     ^^^^^^^^^
```

Best Alignment of Abrilumab CDR-H3 chain to a sequence from OAS

```
QVQLVQSGAEVKKPGASVKVSCKVSGYTL--SDLSIHWVRQAPGKGLEWMGGFDPQDGETIYAQKFQGRVTMTEDTSTDTAYMELSSLKSEDTAVYYCATGSSSSWFDPWGQGTLVTVSS
....................|.|||.....|.....|.||.||.||||.|......|.|.|......|||...|||.......|||....||||||||.||||||||||||||||||||
-------------SETLSLTCTVSGGSISSSSYYWGWIRQPPGTGLEWIGSIYYS-GSTYYNPSLKSRVTISVDTSKNQFSLKLSSVTAADTAVYYCARGSSSSWFDPWGQGTLVTVSS
                          ^^^^^^^^^^                ^^^^^^^^                                  ^^^^^^^^^^^
```

Therapeutic : Actoxumab

Best Alignment of Actoxumab heavy chain to a sequence from OAS

```
QVQLVESGGGVVQPGRSLRLSCAASGFSFSNYGMHWVRQAPGKGLEWVALIWYDGSNEDYTDSVKGRFTISRDNSKNTLYLQMNSLRAEDTAVYYCARWGMVRG------VIDVFDIWGQGTVVTVSS
|||||||||||||||||||||||||||.||.||||||||||||||||||.|||||||..|.|||||||||||||||||||||||||||||||||||||.|||||......|..||.|||||.|||||
QVQLVESGGGVVQPGRSLRLSCAASGFTFSSYGMHWVRQAPGKGLEWVAVIWYDGSNKYYADSVKGRFTISRDNSKNTLYLQMNSLRAEDTAVYYCARGGMVRGVWSRRNAIRSFDYWGQGTLVTVSS
                          ^^^^^^^^                ^^^^^^^^                                  ^^^^^^^^^^^^^^^^^^^^
```

Best Alignment of Actoxumab light chain to a sequence from OAS

```
DIQMTQSPSSVSASVGDRVTITCRASQGISSWLAWYQHKPGKAPKLLIYAASSLQSGVPSRFSGSGSGTDFTLTISSLQPEDFATYYCQQANSFPWTFGQGTKVEIK
|||||||||||||||||||||||||||||||||||||||||||||||||||||||||||||||||||||||||||||||||||||||||||||||||||||||||||
DIQMTQSPSSVSASVGDRVTITCRASQGISSWLAWYQHKPGKAPKLLIYAASSLQSGVPSRFSGSGSGTDFTLTISSLQPEDFATYYCQQANSFPWTFGQGTKVEIK
                          ^^^^^^                 ^^^                                     ^^^^^^^^^
```

Best Alignment of Actoxumab heavy chain CDRs to a sequence from OAS

```
QVQLVESGGGVVQPGRSLRLSCAASGFSFSNYGMHWVRQAPGKGLEWVALIWYDGSNEDYTDSVKGRFTISRDNSKNTLYLQMNSLRAEDTAVYYCARWGMVRGVIDVFDIWGQGTVVTVSS
................|||||||||||.||.||||||||||||||||||.|||||||..|.|||||||||||||||||||||||||||||||||||||.||||||||.||||||||.|||||
----------------SLRLSCAASGFTFSSYGMHWVRQAPGKGLEWVAVIWYDGSNKYYADSVKGRFTISRDNSKNTLYLQMNSLRAEDTAVYYCARSGMVRGVIDAFDIWGQGTMVTVSS
                          ^^^^^^^^                ^^^^^^^^                                  ^^^^^^^^^^^^^^^
```

Best Alignment of Actoxumab light CDRs chain to a sequence from OAS

```
DIQMTQSPSSVSASVGDRVTITCRASQGISSWLAWYQHKPGKAPKLLIYAASSLQSGVPSRFSGSGSGTDFTLTISSLQPEDFATYYCQQANSFPWTFGQGTKVEIK
|||||||||||||||||||||||||||||||||||||.||||||||||||||||||||||||||||||||||||||||||||.||||||||||||||||||||||||
DIQMTQSPSSVSASVGDRVTITCRASQGISSWLAWYQQKPGKAPKLLIYAASSLQSGVPSRFSGSGSGTDFTLTISSLQPEDLATYYCQQANSFPWTFGQGTKVEIK
                          ^^^^^^                 ^^^                                     ^^^^^^^^^
```

Best Alignment of Actoxumab CDR-H3 chain to a sequence from OAS

```
QVQLVESGGGVVQPGRSLRLSCAASGFSFSNYGMHWVRQAPGKGLEWVALIWYDGSNEDYTDSVKGRFTISRDNSKNTLYLQMNSLRAEDTAVYYCARWGMVRGVIDVFDIWGQGTVVTVSS
................|||||||||||.||.||||||||||||||||||.|||||||..|.|||||||||||||||||||||||||||||||||||||.||||||||.||||||||.|||||
----------------SLRLSCAASGFTFSSYGMHWVRQAPGKGLEWVAVIWYDGSNKYYADSVKGRFTISRDNSKNTLYLQMNSLRAEDTAVYYCARSGMVRGVIDAFDIWGQGTMVTVSS
                          ^^^^^^^^                ^^^^^^^^                                  ^^^^^^^^^^^^^^^
```

Therapeutic : Adalimumab

Best Alignment of Adalimumab heavy chain to a sequence from OAS

```
EVQLVESGGGLVQPGRSLRLSCAASGFTFDDYAMHWVRQAPGKGLEWVSAITWNSGHIDYADSVEGRFTISRDNAKNSLYLQMNSLRAEDTAVYYCAKVSYLS-TASSLDYWGQGTLVTVSS
|||||||||||||||||||||||||||||||||||||||||||||||||.|.||||.|.|||||.||||||||||||||||||||||||||||.|||||.||...||.|||||||||||||||
EVQLVESGGGLVQPGRSLRLSCAASGFTFDDYAMHWVRQAPGKGLEWVSGISWNSGSIGYADSVKGRFTISRDNAKNSLYLQMNSLRAEDTALYYCAKASYTVTTAFSLDYWGQGTLVTVSS
                          ^^^^^^^^                ^^^^^^^^                                  ^^^^^^^^^^^^^^^
```

Best Alignment of Adalimumab light chain to a sequence from OAS

```
DIQMTQSPSSLSASVGDRVTITCRASQGIRNYLAWYQQKPGKAPKLLIYAASTLQSGVPSRFSGSGSGTDFTLTISSLQPEDVATYYCQRYNRAPYTFGQGTKVEIK
||||||||||||||||||||||||||||||||||||||||||.|||||||||||||||||||||||||||||||||||||||||||||||||.||.|||||||||||
DIQMTQSPSSLSASVGDRVTITCRASQGIRNYLAWYQQKPGKVPKLLIYAASTLQSGVPSRFSGSGSGTDFTLTISSLQPEDVATYYCQRYNSAPRTFGQGTKVEIK
                          ^^^^^^                 ^^^                                     ^^^^^^^^^
```

Best Alignment of Adalimumab heavy chain CDRs to a sequence from OAS

```
EVQLVESGGGLVQPGRSLRLSCAASGFTFDDYAMHWVRQAPGKGLEWVSAITWNSGHIDYADSVEGRFTISRDNAKNSLYLQMNSLRAEDTAVYYCAKVSYLSTASSLDYWGQGTLVTVSS
..............|.|||||||||||||||||||||||||||||||||.|.||||.|.|||.|.||||||||||||||||||||||.||||.|||||.||.|..||||||||||||||||
--------------GGSLRLSCAASGFTFDDYAMHWVRQAPGKGLEWVSGISWNSGSIGYADFVKGRFTISRDNAKNSLYLQMNSLRVEDTALYYCAKDSYSSSWSSLDYWGQGTLVTVSS
                          ^^^^^^^^                ^^^^^^^^                                  ^^^^^^^^^^^^^^
```

Best Alignment of Adalimumab light CDRs chain to a sequence from OAS

```
DIQMTQSPSSLSASVGDRVTITCRASQGIRNYLAWYQQKPGKAPKLLIYAASTLQSGVPSRFSGSGSGTDFTLTISSLQPEDVATYYCQRYNRAPYTFGQGTKVEIK
|||||||||||||||||||||||||||||.||||||||||||.||||||||||||||||||||||||||||||||||||||||||||||||||||||||||||.|||
DIQMTQSPSSLSASVGDRVTITCRASQGINNYLAWYQQKPGKVPKLLIYAASTLQSGVPSRFSGSGSGTDFTLTISSLQPEDVATYYCQRYNRAPYTFGQGTKLEIK
                          ^^^^^^                 ^^^                                     ^^^^^^^^^
```

Best Alignment of Adalimumab CDR-H3 chain to a sequence from OAS

```
EVQLVESGGGLVQPGRSLRLSCAASGFTFDDYAMHWVRQAPGKGLEWVSAITWNSGHIDYADSVEGRFTISRDNAKNSLYLQMNSLRAEDTAVYYCAKVSYLSTASSLDYWGQGTLVTVSS
..............|.|||||||||||||..|.|||||||||||||||..|........|||||.|||||||||.||.|||||||||||||||||||||..||..||||||||||||||||
--------------GGSLRLSCAASGFTFSSYGMHWVRQAPGKGLEWVAVISYDGSNKYYADSVKGRFTISRDNSKNTLYLQMNSLRAEDTAVYYCAKVQGLSRRSSLDYWGQGTLVTVSS
                          ^^^^^^^^                ^^^^^^^^                                  ^^^^^^^^^^^^^^
```

Therapeutic : Aducanumab

Best Alignment of Aducanumab heavy chain to a sequence from OAS

```
QVQLVESGGGVVQPGRSLRLSCAASGFAFSSYGMHWVRQAPGKGLEWVAVIWFDGTKKYYTDSVKGRFTISRDNSKNTLYLQMNTLRAEDTAVYYCARDRGIGAR-RGPYYMDVWGKGTTVTVSS
|||||||||||||||||||||||||||.||||||||||||||||||||||||.||..|||.|||||||||||||||||||||||.||||||||||||||.|...|.||.||||||||||||||||
QVQLVESGGGVVQPGRSLRLSCAASGFTFSSYGMHWVRQAPGKGLEWVAVIWYDGSNKYYADSVKGRFTISRDNSKNTLYLQMNSLRAEDTAVYYCARDWGSITRNRGYYYMDVWGKGTTVTVSS
                          ^^^^^^^^                ^^^^^^^^                                  ^^^^^^^^^^^^^^^^^
```

Best Alignment of Aducanumab light chain to a sequence from OAS

```
DIQMTQSPSSLSASVGDRVTITCRASQSISSYLNWYQQKPGKAPKLLIYAASSLQSGVPSRFSGSGSGTDFTLTISSLQPEDFATYYCQQSYSTPLTFGGGTKVEIK
|||||||||||||||||||||||||||||||||||||||||||||||||||||||||||||||||||||||||||||||||||||||||||||||||||||||||||
DIQMTQSPSSLSASVGDRVTITCRASQSISSYLNWYQQKPGKAPKLLIYAASSLQSGVPSRFSGSGSGTDFTLTISSLQPEDFATYYCQQSYSTPLTFGGGTKVEIK
                          ^^^^^^                 ^^^                                     ^^^^^^^^^
```

Best Alignment of Aducanumab heavy chain CDRs to a sequence from OAS

```
QVQLVESGGGVVQPGRSLRLSCAASGFAFSSYGMHWVRQAPGKGLEWVAVIWFDGTKKYYTDSVKGRFTISRDNSKNTLYLQMNTLRAEDTAVYYCARDRGIGARRGPYYMDVWGKGTTVTVSS
..............|.|||||||||||.|||||||||||||||||||||||||||.|||..|||||||||||||||||||||||.||||||||||||||||.|.|...||||||||||||||||
--------------GGSLRLSCAASGFIFSSYGMHWVRQAPGKGLEWVAVIWFDGSKKYFADSVKGRFTISRDNSKNTLYLQMNSLRAEDTAVYYCARDRGYGTRYSYYYMDVWGKGTTVTVSS
                          ^^^^^^^^                ^^^^^^^^                                  ^^^^^^^^^^^^^^^^
```

Best Alignment of Aducanumab light CDRs chain to a sequence from OAS

```
DIQMTQSPSSLSASVGDRVTITCRASQSISSYLNWYQQKPGKAPKLLIYAASSLQSGVPSRFSGSGSGTDFTLTISSLQPEDFATYYCQQSYSTPLTFGGGTKVEIK
|||||||||||||||||||||||||||||||||||||||||||||.|||||||||||||||||||||||||||||||||||||.|||||||||||||||||||||||
DIQMTQSPSSLSASVGDRVTITCRASQSISSYLNWYQQKPGKAPKPLIYAASSLQSGVPSRFSGSGSGTDFTLTISSLQPEDFGTYYCQQSYSTPLTFGGGTKVEIK
                          ^^^^^^                 ^^^                                     ^^^^^^^^^
```

Best Alignment of Aducanumab CDR-H3 chain to a sequence from OAS

```
QVQLVESGGGVVQPGRSLRLSCAASGFAFSSYGMHWVRQAPGKGLEWVAVIWFDGTKKYYTDSVKGRFTISRDNSKNTLYLQMNTLRAEDTAVYYCARDRGIGARRGPYYMDVWGKGTTVTVSS
.............||.|||||||||||.||.|.|.|.|||||||||||..|.......||.|||||||||||||..|.||||||.||||||||||||||||||||.|.|||||||||||||||.
-------------PGGSLRLSCAASGFTFSDYYMSWIRQAPGKGLEWVSYISSSDSTIYYADSVKGRFTISRDNAENSLYLQMNSLRAEDTAVYYCARDRGIGARPGDYYMDVWGKGTTVTVS-
                          ^^^^^^^^                ^^^^^^^^                                  ^^^^^^^^^^^^^^^^
```

Therapeutic : Alemtuzumab

Best Alignment of Alemtuzumab heavy chain to a sequence from OAS
```
QVQLQESGPGLVRPSQTLSLTCTVSGFTF--TDFYMNWVRQPPGRGLEWIGFIRDKAKGYTTEYNPSVKGRVTMLVDTSKNQFSLRLSSVTAADTAVYYCAREGHT----AAPFDYWGQGSLVTVSS
||||||||||||.|||||||||||||||....  ..|||..|.|||||.||||||||.|.....||.||||.|.|||..|||||||||.||||||||||||||||...|......||||||||.||||||
QVQLQESGPGLVKPSQTLSLTCTVSGGSISTADFYWSWIRQPPGKGLEWIGFIHDS---GTTYYNPSLKSRVTISVDTSKNQFSLKLSSVTAADTAVYYCARGLPTRLRTQRPFDYWGQGTLVTVSS
                         ^^^^^^^^^^                 ^^^^^^^^^^                                      ^^^^^^^^^^^^^^^^
```

Best Alignment of Alemtuzumab light chain to a sequence from OAS
```
DIQMTQSPSSLSASVGDRVTITCKASQNIDKYLNWYQQKPGKAPKLLIYNTNNLQTGVPSRFSGSGSGTDFTFTISSLQPEDIATYYCLQHISRPRTFGQGTKVEIK
|||||||||||||||||||||||.|||.|..|||||||||||||||||||...||.|||||||||||||||||||||||||||||||||||||||.|.|||||||||||||
DIQMTQSPSSLSASVGDRVTITCQASQDISNYLNWYQQKPGKAPKLLIYDASNLETGVPSRFSGSGSGTDFTFTISSLQPEDIATYYCLQHNSYPRTFGQGTKVEIK
                          ^^^^^^                 ^^^                                    ^^^^^^^^^
```

Best Alignment of Alemtuzumab heavy chain CDRs to a sequence from OAS
```
QVQLQESGPGLVRPSQTLSLTCTVSGFTFTDFYMNWVRQPPGRGLEWIGFIRDKAKGYTTEYNPSVKGRVTMLVDTSKNQFSLRLSSVTAADTAVYYCAREGHTAAPFDYWGQGSLVTVSS
.|.|.|||.|||.|...|.|.|..|||||||.||.|||||||..|||.||||.||.||||||..|||||.|...|.|.....|......|.|.|.|||||||.|.||.||||||..|||||
EVHLVESGGGLVQPGGSLRLSCATSGFTFTDYYMSWVRQPPGKALEWLGFIRNKANGYTTEYSASVKGRFTISRDNSQSILYLQMNTLRAEDSATYYCAREGITTAPMDYWGQGTSVTVSS
                         ^^^^^^^^                 ^^^^^^^^^^                                      ^^^^^^^^^^^^
```

Best Alignment of Alemtuzumab light CDRs chain to a sequence from OAS
```
DIQMTQSPSSLSASVGDRVTITCKASQNIDKYLNWYQQKPGKAPKLLIYNTNNLQTGVPSRFSGSGSGTDFTFTISSLQPEDIATYYCLQHISRPRTFGQGTKVEIK
|||||||||||||||||..||||.|||.|.|||.|||||||||||||.||..|..||.||||||||||.||.||.|..||.|||.|.||||.|||.|.|||||||||||||
DIQMTQSPSSLSASVGDTGTITCQASQGISKYLAWYQQKPGKAPKPLINDTSTLQSGVPSRFSGIGSWTDCTLPISTLQPADFATYYDLQHNSYPRTFGQGTKVEIK
                          ^^^^^^                 ^^^                                    ^^^^^^^^^
```

Best Alignment of Alemtuzumab CDR-H3 chain to a sequence from OAS
```
QVQLQESGPGLVRPSQTLSLTCTVSGFTF-TDFYMNWVRQPPGRGLEWIGFIRDKAKGYTTEYNPSVKGRVTMLVDTSKNQFSLRLSSVTAADTAVYYCAREGHTAAPFDYWGQGSLVTVSS
.....||||.||.|||.|||||||.|..........|.||.||..|||.|.|........|.||||.|.|.....|||||||.|.|.|||..|||.|||||||.|||||||||||...||||
-----ESGPDLVKPSQSLSLTCTVTGYSITSGYSWHWIRQFPGNKLEWMGYIHYS---GSTNYNPSLKSRISITRDTSKNQFFLQLNSVTTEDTATYYCAREGITAAPFDYWGQGTTLTVSS
                         ^^^^^^^^^                 ^^^^^^^^^^                                      ^^^^^^^^^^^^
```

Therapeutic : Alirocumab

Best Alignment of Alirocumab heavy chain to a sequence from OAS
```
EVQLVESGGGLVQPGGSLRLSCAASGFTFNNYAMNWVRQAPGKGLDWVSTISGSGGTTNYADSVKGRFIISRDSSKHTLYLQMNSLRAEDTAVYYCAKDSNW--GNFDLWGRGTLVTVSS
||||.|||||||||||||||||||||||||..||||||||||||||.||||||||||.|.|||||||||.||||.||.||||||||||||||||||||||||||....|.||||||||||||||
EVQLLESGGGLVQPGGSLRLSCAASGFTFSSYAMNWVRQAPGKGLEWVSTISGSGGSTYYADSVKGRFTISRDNSKNTLYLQMNSLRAEDTAVYYCAKDSSGWGGGFDLWGRGTLVTVSS
                         ^^^^^^^^                 ^^^^^^^^                                      ^^^^^^^^^^^^^
```

Best Alignment of Alirocumab light chain to a sequence from OAS
```
DIVMTQSPDSLAVSLGERATINCKSSQSVLYRSNNRNFLGWYQQKPGQPPNLLIYWASTRESGVPDRFSGSGSGTDFTLTISSLQAEDVAVYYCQQYYTTPYTFGQGTKLEIK
|||||||||||||||||||||||||||||||.|||.|.||||||||||||.|||||||||||||||||||||||||||||||||||||||||||||||||||||||||||||
DIVMTQSPDSLAVSLGERATINCKSSQSVLYSSNNNNYLGWYQQKPGQPPKLLIYWASTRESGVPDRFSGSGSGTDFTLTISSLQAEDVAVYYCQQYYTTPYTFGQGTKLEIK
                          ^^^^^^^^^^^^                 ^^^                                    ^^^^^^^^^
```

Best Alignment of Alirocumab heavy chain CDRs to a sequence from OAS
```
EVQLVESGGGLVQPGGSLRLSCAASGFTFNNYAMNWVRQAPGKGLDWVSTISGSGGTTNYADSVKGRFIISRDSSKHTLYLQMNSLRAEDTAVYYCAKDSNWGNFDLWGRGTLVTVSS
..............||||||||||||||||.|||.||||||||||.||||||||||.|.|||||||||.||||.||.||||||||||||||||||||||||||.||.||.||||||||
--------------GGSLRLSCAASGFTFNSYAMSWVRQAPGKGLEWVSTISGSGGSTYYADSVKGRFTISRDNSKNTLYLQMNSLRAEDTAVYYCAKDSNWGYFDYWGQGTLVTVSS
                         ^^^^^^^^                 ^^^^^^^^                                      ^^^^^^^^^^^
```

Best Alignment of Alirocumab light CDRs chain to a sequence from OAS
```
DIVMTQSPDSLAVSLGERATINCKSSQSVLYRSNNRNFLGWYQQKPGQPPNLLIYWASTRESGVPDRFSGSGSGTDFTLTISSLQAEDVAVYYCQQYYTTPYTFGQGTKLEIK
||||||||||||||||||||.||||||||||||||||||.||||||||||.||||||||||||||||||||||||||||||||||||||||.|||||||.|||||||||||||.||
DIVMTQSPDSLAVSLGERATLNCKSSQSVLYRSNNRNFLAWYQQKPGQPPKLLIYWASTRESGVPDRFSGSGSGTDFTLTISSLQAEDVAGYYCQQYYSTPYTFGQGTKLQIK
                          ^^^^^^^^^^^^                 ^^^                                    ^^^^^^^^^
```

Best Alignment of Alirocumab CDR-H3 chain to a sequence from OAS
```
EVQLVESGGGLVQPGGSLRLSCAASGFTFNNYAMNWVRQAPGKGLDWVSTISGS--GGTTNYADSVKGRFIISRDSSKHTLYLQMNSLRAEDTAVYYCAKDSNWGNFDLWGRGTLVTVSS
..............||||||||||||||..||.|.||||||.|||.|.|.|.|.....|..|..||||||||.||||.|.|..||||||.||..||||||||||||.||||||||||.||||||
--------------GGSLRLSCAASGFTLSNYPMGWVRQAPVKGLEWLSAIGEEKSGSWTKSADSVKGRFTISRDNSENTLYLQMDSLTVEDTAVYYCAKDSSWGNFDLWGRGALVTVSS
                         ^^^^^^^^                 ^^^^^^^^^^                                      ^^^^^^^^^^^
```

Therapeutic : Amatuximab

Best Alignment of Amatuximab heavy chain to a sequence from OAS
```
QVQLQQSGPELEKPGASVKISCKASGYSFTGYTMNWVKQSHGKSLEWIGLITPYNGASSYNQKFRGKATLTVDKSSSTAYMDLLSLTSEDSAVYFCARGGYD-GRGFDYWGSGTPVTVSS
.||||||||||||||||||||||||||||||||||||||||||.||||||||.||||..||||||.|||||||||||||||||.||||||||||||.|||||||..|||.|||.||.||||.
-VQLQQSGPELEKPGASVKISCKASGYSFTGYTMNWVKQSHGKNLEWIGLINPYNGGTSYNQKFKGKATLTVDKSSSTAYMELLSLTSEDSAVYYCARGGYDEDRGFAYWGQGTLVTVSA
                         ^^^^^^^^                 ^^^^^^^^                                      ^^^^^^^^^^^^^
```

Best Alignment of Amatuximab light chain to a sequence from OAS
```
DIELTQSPAIMSASPGEKVTMTCSASSSVSYMHWYQQKSGTSPKRWIYDTSKLASGVPGRFSGSGSGNSYSLTISSVEAEDDATYYCQQWSKHPLTFGSGTKVEIK
||.||||||||||||||||||||||||||||||||||||.||||.|||||||||||||||||||||||||||||||||||||||||||||||..|.|||||||.|||
DIVLTQSPAIMSASPGEKVTMTCSASSSVSYMHWYQQKSSTSPKLWIYDTSKLASGVPGRFSGSGSGNSYSLTISSVEAEDDATYYCQQWSGYPFTFGSGTKLEIK
                          ^^^^^                 ^^^                                    ^^^^^^^^^
```

Best Alignment of Amatuximab heavy chain CDRs to a sequence from OAS
```
QVQLQQSGPELEKPGASVKISCKASGYSFTGYTMNWVKQSHGKSLEWIGLITPYNGASSYNQKFRGKATLTVDKSSSTAYMDLLSLTSEDSAVYFCARGGYDGRGFDYWGSGTPVTVSS
.||||.|||||.|.||||||||||||||||||.|.|||||||||||||||.|..||||.||||||.||||.||||.|||||||...||||||||||.|||||||||||.|||.||.||||.
DVQLQESGPELVKTGASVKISCKASGYSFTGYYMHWVKQSHGKSLEWIGYISCYNGATSYNQKFKGKATFTVDTSSSTAYMQFNSLTSEDSAVYYCARGGYDGRGFAYWGQGTLVTVSA
                         ^^^^^^^^                 ^^^^^^^^                                      ^^^^^^^^^^^^
```

Best Alignment of Amatuximab light CDRs chain to a sequence from OAS
```
DIELTQSPAIMSASPGEKVTMTCSASSSVSYMHWYQQKSGTSPKRWIYDTSKLASGVPGRFSGSGSGNSYSLTISSVEAEDDATYYCQQWSKHPLTFGSGTKVEIK
.|.|||||||||||||||||||||||||||||||||||||||||||||||||||||||.||||||||.||||||||.||||.||||||||||.||||||.|||.|.|
QIVLTQSPAIMSASPGEKVTMTCSASSSVSYMHWYQQKSGTSPKRWIYDTSKLASGVPARFSGSGSGTSYSLTISSMEAEDAATYYCQQWSSHPLTFGAGTKLELK
                          ^^^^^                 ^^^                                    ^^^^^^^^^
```

Best Alignment of Amatuximab CDR-H3 chain to a sequence from OAS
```
QVQLQQSGPELEKPGASVKISCKASGYSF-TGYTMNWVKQSHGKSLEWIGLITPYNGASSYNQKFRGKATLTVDKSSSTAYMDLLSLTSEDSAVYFCARGGYDGRGFDYWGSGTPVTVSS
.||||||||.|.||..|....|...|||...||..||..|..|..|||.|.|....|...||.........|.|.|........|.|.|.|.||.|.|.|||||||||||||||.||.||||.
EVQLQQSGPGLVKPSQSLSLTCSVTGYSITSGYYWNWIRQFPGNKLEWMGYISYD-GSNNYNPSLKNRVSITRDTSKNQFFLKLNSVTTEDTATYYCARGGYDGRGFDYWGQGTLVTVSA
                         ^^^^^^^^^                 ^^^^^^^^                                      ^^^^^^^^^^^^
```

Therapeutic : Andecaliximab

Best Alignment of Andecaliximab heavy chain to a sequence from OAS
```
QVQLQESGPGLVKPSETLSLTCTVSGFSLLSYGVHWVRQPPGKGLEWLGVIWTGGTTNYNSALMSRFTISKDDSKNTVYLKMNSLKTEDTAIYYCARYYY-----GMDYWGQGTLVTVSS
||||.|||||||.||..||.|||||||||.|||||||||||||||||||||||.||.||||||||||..||||.|||.|.||||||.||||||.|||||.||......||||||||.|||||
QVQLKESGPGLVAPSQSLSITCTVSGFSLTSYGVHWVRQPPGKGLEWLGVIWAGGSTNYNSALMSRLSISKDNSKNQVVLKMNSLQTEDTAMYYCARGYYGNPYYAMDYWGQGTSVTVSS
                         ^^^^^^^^                 ^^^^^^^                                       ^^^^^^^^^^^^^^
```

Best Alignment of Andecaliximab light chain to a sequence from OAS

```
DIQMTQSPSSLSASVGDRVTITCKASQDVRNTVAWYQQKPGKAPKLLIYSSSYRNTGVPDRFSGSGSGTDFTLTISSLQAEDVAVYYCQQHYITPYTFGGGTKVEIK
|||||||||||||||||||||||.||||..|...|||||||||||||||..|...||||.||||||||||||||||||||||||||||||.|.||||||.|||.|||
DIQMTQSPSSLSASVGDRVTITCQASQDISNYLNWYQQKPGKAPKLLIYHASNLETGVPSRFSGSGSGTDFTLTISSLQAEDVAVYYCQQYYSTPYTFGQGTKLEIK
                        ^^^^^^          ^^^                   ^^^^^^^^^
```

Best Alignment of Andecaliximab heavy chain CDRs to a sequence from OAS

```
QVQLQESGPGLVKPSETLSLTCTVSGFSLLSYGVHWVRQPPGKGLEWLGVIWTGGTTNYNSALMSRFTISKDDSKNTVYLKMNSLKTEDTAIYYCARYYYGMDYWGQGTLVTVSS
.||..|||||||.||..||.|||||||||.|||||||||.||||||||||||.||.|.||.|..||..||||.||..|..|||||...||||||||||||||||||||||.|||||
-VQRVESGPGLVQPSQSLSITCTVSGFSLTSYGVHWVRQSPGKGLEWLGVIWSGGSTDYNAAFISRLSISKDNSKSQVFFKMNSLQADDTAIYYCARYYYGMDYWGQGTSVTVSS
                        ^^^^^^^^          ^^^^^^^                   ^^^^^^^^^
```

Best Alignment of Andecaliximab light CDRs chain to a sequence from OAS

```
DIQMTQSPSSLSASVGDRVTITCKASQDVRNTVAWYQQKPGKAPKLLIYSSSYRNTGVPDRFSGSGSGTDFTLTISSLQAEDVAVYYCQQHYITPYTFGGGTKVEIK
.|..|.|....|.||.|||.||||||||||.||||||||||..|||||...|.|.|||||||.|||||||.||||||.||||.|.|||||||.||||||||||.|||
-IVLTLSHKFMSTSVVDRVSITCKASQDVRTTVAWYQQKPGQSPKLLIDWASTRDTGVPDRFTGSGSGTDYTLTISSVQAEDLALYYCQQHYSTPYTFGGGTKLEIK
                        ^^^^^^          ^^^                   ^^^^^^^^^
```

Best Alignment of Andecaliximab CDR-H3 chain to a sequence from OAS

```
QVQLQESGPGLVKPSETLSLTCTVSGFSLLSYGVHWVRQPPGKGLEWLGVIWTG-GTTNYNSALMSRFTISKDDSKNTVYLKMNSLKTEDTAIYYCARYYYGMDYWGQGTLVTVSS
.||||.|||.||||.......|..||.|...|...||.|..||.|||.|.|....|.|.||.......|...|.|..|.|.|....||..||.|.|.|||||||||||||||||.|||||
EVQLQQSGPELVKPGASMKIYCKASGYSFTGYTMNWVKQSHGKNLEWIGLINPYNGGTSYNQKFKGKATLTVDKSSSTAYMELLSLTSEDSAVYYCARYYYGMDYWGQGTSVTVSS
                        ^^^^^^^^          ^^^^^^^^                   ^^^^^^^^^
```

Therapeutic : Anetumab

Best Alignment of Anetumab heavy chain to a sequence from OAS

```
QVELVQSGAEVKKPGESLKISCKGSGYSFTSYWIGWVRQAPGKGLEWMGIIDPGDSRTRYSPSFQGQVTISADKSISTAYLQWSSLKASDTAMYYCARGQLYG------GTYMDGWGQGTLVTVSS
.|.||||||||||||||||||||||||||||||||||||.|||||||||||.||||.||||||||||||||||||||||||||||||||||||||||||.||.......|||.|.|||||||||||
EVQLVQSGAEVKKPGESLKISCKGSGYSFTSYWIGWVRQMPGKGLEWMGIIYPGDSDTRYSPSFQGQVTISADKSISTAYLQWSSLKASDTAMYYCARGILYYDSSGYKGTYFDYWGQGTLVTVSS
                        ^^^^^^^^          ^^^^^^^^                   ^^^^^^^^^^^^^^^^^^^
```

Best Alignment of Anetumab light chain to a sequence from OAS

```
DIALTQPASVSGSPGQSITISCTGTSSDIGGYNSVSWYQQHPGKAPKLMIYGVNNRPSGVSNRFSGSKSGNTASLTISGLQAEDEADYYCSSYDIESATPVFGGGTKLTVL
..||||||||||||||||||||||||||.||||.|||||||||||||||||.|||||||||||||||||||||||||||||||||||||||||.|.|.|||||||||||||
QSALTQPASVSGSPGQSITISCTGTSSDVGGYNYVSWYQQHPGKAPKLMIYDVNNRPSGVSNRFSGSKSGNTASLTISGLQAEDEADYYCSSYTI-SSTPVFGGGTKLTVL
                        ^^^^^^^^^          ^^^                   ^^^^^^^^^^^
```

Best Alignment of Anetumab heavy chain CDRs to a sequence from OAS

```
QVELVQSGAEVKKPGESLKISCKGSGYSFTSYWIGWVRQAPGKGLEWMGIIDPGDSRTRYSPSFQGQVTISADKSISTAYLQWSSLKASDTAMYYCARGQLYGGTYMDGWGQGTLVTVSS
.............||||||||||||||||||||||||||.|||||||||||.||||.||||||||||||||||||||||||||||||||||||||||||..|||||.|||||||||||||
-------------PGESLKISCKGSGYSFTSYWIGWVRQMPGKGLEWMGIIYPGDSDTRYSPSFQGQVTISADKSISTAYLQWSSLKASDTAMYYCARGRDYGGTYLDGWGQGTLVTVSS
                        ^^^^^^^^          ^^^^^^^^                   ^^^^^^^^^^^^^
```

Best Alignment of Anetumab light CDRs chain to a sequence from OAS

```
DIALTQPASVSGSPGQSITISCTGTSSDIGGYNSVSWYQQHPGKAPKLMIYGVNNRPSGVSNRFSGSKSGNTASLTISGLQAEDEADYYCSSYDIESATPVFGGGTKLTVL
..|||||.|.|||||||.||||||.|||||||||||||||.||||||||||.||.|||||..||||||||||||||.|||||||||.||||||...||..|||||||||||
QSALTQPPSASGSPGQSVTISCTGSSSDIGGYNSVSWYQQFPGKAPKLMIYEVNKRPSGVPDRFSGSKSGNTASLTVSGLQAEDEAYYYCSSYTSTSAPGVFGGGTKLTVL
                        ^^^^^^^^^          ^^^                   ^^^^^^^^^^^
```

Best Alignment of Anetumab CDR-H3 chain to a sequence from OAS

```
QVELVQSGAEVKKPGESLKISCKGSGYSFTSYWIGWVRQAPGKGLEWMGIIDPGDSRTRYSPSFQGQVTISADKSISTAYLQWSSLKASDTAMYYCARGQLYGGTYMDGWGQGTLVTVSS
...............|.|...|...|.|...|...|.||.|||||||.|.|........||||.....|||.|.|.....|..||..|.|||.|||||||.|||||.|||||||||||||
--------------SETLSLTCAVCGGSLSGYYWSWIRQPPGKGLEWIGEILHS-GSNHYSPSLESRLTISVDTSKNQFSLKLSSVTAADTAVYYCARGQDYGGTYLDGWGQGTLVTVSS
                        ^^^^^^^^          ^^^^^^^^                   ^^^^^^^^^^^^^
```

Therapeutic : Anifrolumab

Best Alignment of Anifrolumab heavy chain to a sequence from OAS

```
EVQLVQSGAEVKKPGESLKISCKGSGYIFTNYWIAWVRQMPGKGLESMGIIYPGDSDIRYSPSFQGQVTISADKSITTAYLQWSSLKASDTAMYYCARHDI----EGFDYWGRGTLVTVSS
||||||||||||||||||||||||||||||.|||.|||||||||||.||||||||||.||||||||||||||||||.||||||||||||||||||||||.|.....||||||.||||||||
EVQLVQSGAEVKKPGESLKISCKGSGYIFTSYWIGWVRQMPGKGLEWMGIIYPGDSDTRYSPSFQGQVTISADKSISTAYLQWSSLKASDTAMYYCARHGIAAAGTGFDYWGQGTLVTVSS
                        ^^^^^^^^          ^^^^^^^^                   ^^^^^^^^^^^^^^
```

Best Alignment of Anifrolumab light chain to a sequence from OAS

```
EIVLTQSPGTLSLSPGERATLSCRASQSVSSSFFAWYQQKPGQAPRLLIYGASSRATGIPDRLSGSGSGTDFTLTITRLEPEDFAVYYCQQYDSSAITFGQGTRLEIK
||||||||||||||||||||||||||||||||..||||||||||||||||||||||||||||.|||||||||||||||||||||||||||||.|||||||||||||||
EIVLTQSPGTLSLSPGERATLSCRASQSVSSSYLAWYQQKPGQAPRLLIYGASSRATGIPDRFSGSGSGTDFTLTITRLEPEDFAVYYCQQYGSSAITFGQGTRLEIK
                        ^^^^^^^          ^^^                   ^^^^^^^^^
```

Best Alignment of Anifrolumab heavy chain CDRs to a sequence from OAS

```
EVQLVQSGAEVKKPGESLKISCKGSGYIFTNYWIAWVRQMPGKGLESMGIIYPGDSDIRYSPSFQGQVTISADKSITTAYLQWSSLKASDTAMYYCARHDIEGFDYWGRGTLVTVSS
..............|||||||||||||.||||||.|||||||||||.|||||||||||||||||||||||||||||.|||||||||||||||||||||..||.|||||.||||||||
--------------GESLKISCKGSGYSFTNYWIGWVRQMPGKGLEWMGIIYPGDSDIRYSPSFQGQVTISADKSISTAYLQWSSLKASDTAMYYCARRPIEEFDYWGQGTLVTVSS
                        ^^^^^^^^          ^^^^^^^^                   ^^^^^^^^^^
```

Best Alignment of Anifrolumab light CDRs chain to a sequence from OAS

```
EIVLTQSPGTLSLSPGERATLSCRASQSVSSSFFAWYQQKPGQAPRLLIYGASSRATGIPDRLSGSGSGTDFTLTITRLEPEDFAVYYCQQYDSSAITFGQGTRLEIK
||||||||.|||.||||||||||||||||||||||||||.||||||||||||.|.||..||.|||||||||||||.||||.|.||||||||.||.|||||||||||.
EIVLTQSPDTLSVSPGERATLSCRASQSVSSSFFAWYQQKGGQAPRLLIYGASNRDTGVSDRFSGSGSGTDFTLTISRLEPDDSAVYYCQQYGSSPITFGQGTRLEIE
                        ^^^^^^^          ^^^                   ^^^^^^^^^
```

Best Alignment of Anifrolumab CDR-H3 chain to a sequence from OAS

```
EVQLVQSGAEVKKPGESLKISCKGSGYIFTNYWIAWVRQMPGKGLESMGIIYPGDSDIRYSPSFQGQVTISADKSITTAYLQWSSLKASDTAMYYCARHDIEGFDYWGRGTLVTVSS
............||.|.|...|..||.....|...|.||.||||||..|.||.......|.||....||||.|.|.....|..||..|.|||.|||||||.|||||||.||||||||
---------GLVKPSETLSLTCTVSGGSISSYYWSWIRQPPGKGLEWIGYIYYS-GSTNYNPSLKSRVTISVDTSKNQFSLKLSSVTAADTAVYYCARHDREGFDYWGQGTLVTVSS
                        ^^^^^^^^          ^^^^^^^^                   ^^^^^^^^^^
```

Therapeutic : Anrukinzumab

Best Alignment of Anrukinzumab heavy chain to a sequence from OAS

```
EVQLVESGGGLVQPGGSLRLSCAASGFTFISYAMSWVRQAPGKGLEWVASISSG-GNTYYPDSVKGRFTISRDNAKNSLYLQMNSLRAEDTAVYYCARLDGYYFGFAYWGQGTLVTVSS
|||||||||||||||||||||||||||||.||||||||||||||||||..||...|.|||.|||||||||||||||||||||||||||||||||||||||..|.|.||.||||||||||||
EVQLVESGGGLVQPGGSLRLSCAASGFTFSSYAMSWVRQAPGKGLEWVSAISGSGGSTYYADSVKGRFTISRDNAKNSLYLQMNSLRAEDTAVYYCARGSGWYAGFDYWGQGTLVTVSS
                        ^^^^^^^^          ^^^^^^^^                   ^^^^^^^^^^^^
```

Best Alignment of Anrukinzumab light chain to a sequence from OAS

```
DIQMTQSPSSLSASVGDRVTITCKASESVDNYGKSLMHWYQQKPGKAPKLLIYRASNLESGVPSRFSGSGSGTDFTLTISSLQPEDFATYYCQQSNEDPWTFGGGTKVEIK
|||||||||||||||||||||||.||||......|...|||||||||||||||.||.|||||||||||||||||||||||||||||||||||||||..|.|||||||||||
DIQMTQSPSSLSASVGDRVTITCRASESI----SSYLNWYQQKPGKAPKLLIYAASSLESGVPSRFSGSGSGTDFTLTISSLQPEDFATYYCQQSNSNPLTFGGGTKVEIK
                        ^^^^^^^^^^          ^^^                   ^^^^^^^^^
```

Best Alignment of Anrukinzumab heavy chain CDRs to a sequence from OAS

```
EVQLVESGGGLVQPGGSLRLSCAASGFTFISYAMSWVRQAPGKGLEWVASISSGGNTYYPDSVKGRFTISRDNAKNSLYLQMNSLRAEDTAVYYCARLDGYYFGFAYWGQGTLVTVSS
||.|||||||||.|||||.||||||||||.|||||||||.|.|.|||||||||||.||||||||||||||||||.|.|||||.|||.||||.|||||.||||.||||||||||||||.
EVKLVESGGGLVKPGGSLKLSCAASGFTFSSYAMSWVRQTPEKRLEWVASISSGGSTYYPDSVKGRFTISRDNARNILYLQMSSLRSEDTAMYYCARSDGYYGGFAYWGQGTLVTVSA
                        ^^^^^^^^          ^^^^^^^                   ^^^^^^^^^^^^
```

Best Alignment of Anrukinzumab light CDRs chain to a sequence from OAS

```
DIQMTQSPSSLSASVGDRVTITCKASESVDNYGKSLMHWYQQKPGKAPKLLIYRASNLESGVPSRFSGSGSGTDFTLTISSLQPEDFATYYCQQSNEDPWTFGGGTKVEIK
||..||||.||..|...|..|.|.||||||.||||.|||||||||..||||||||||||||.|.|||||||.|||||||......|.||||||||||||||||||||.|||
DILLTQSPASLAVSLEQRASIACRASESVDSYGKSFMHWYQQKPGQPPKLLIYRASNLESGIPARFSGSGSRTDFTLTINPVEADDVATYYCQQSNEDPWTFGGGTKLEIK
           ^^^^^^^^^^       ^^^               ^^^^^^^^^
```

Best Alignment of Anrukinzumab CDR-H3 chain to a sequence from OAS

```
EVQLVESGGGLVQPGGSLRLSCAASGFTFISYAMSWVRQAPGKGLEWVASISSG-GNTYYPDSVKGRFTISRDNAKNSLYLQMNSLRAEDTAVYYCARLDGYYFGFAYWGQGTLVTVSS
||||..||..||.||.|..|||.||||....|.|.||.|....||||...|....|.|.|.....|..||..|...|..|||..||..|||||||||||||||.||||||||||||||.
EVQLQQSGAELVKPGASVKLSCTASGFNIKDYYMHWVKQRTEQGLEWIGRIDPEDGETKYAPKFQGKATITADTSSNTAYLQLSSLTSEDTAVYYCARLDGYYEGFAYWGQGTLVTVSA
           ^^^^^^^^       ^^^^^^^^               ^^^^^^^^^^^^
```

Therapeutic : Ascrinvacumab

Best Alignment of Ascrinvacumab heavy chain to a sequence from OAS

```
QVQLQESGPGLVKPSQTLSLTCTVSGGSISSGEYYWNWIRQHPGKGLEWIGYIYYSGSTYYNPSLKSRVTISVDTSKNQFSLKLSSVTAADTAVYYCARESV--AGFDYWGQGTLVTVSS
||||||||||||||||||||||||||||||||.|||.|||||||||||||||||||||||||||||||||||||||||||||||||||||||||||||||||..|.||||||||||||||
QVQLQESGPGLVKPSQTLSLTCTVSGGSISSGGYYWSWIRQHPGKGLEWIGYIYYSGSTYYNPSLKSRVTISVDTSKNQFSLKLSSVTAADTAVYYCARESVDTANFDYWGQGTLVTVSS
           ^^^^^^^^^^       ^^^^^^^               ^^^^^^^^^^^^
```

Best Alignment of Ascrinvacumab light chain to a sequence from OAS

```
EIVLTQSPGTLSLSPGERATLSCRASQSVSSSYLAWYQQKPGQAPRLLIYGTSSRATGIPDRFSGSGSGTDFTLTISRLEPEDFAVYYCQQYGSSPITFGQGTRLEIK
||||||||||||||||||||||||||||||||||||||||||||||||||||||||||||||||||||||||||||||||||||||||||||||||||||||||||||
EIVLTQSPGTLSLSPGERATLSCRASQSVSSSYLAWYQQKPGQAPRLLIYGTSSRATGIPDRFSGSGSGTDFTLTISRLEPEDFAVYYCQQYGSSPITFGQGTRLEIK
           ^^^^^^^       ^^^               ^^^^^^^^^
```

Best Alignment of Ascrinvacumab heavy chain CDRs to a sequence from OAS

```
QVQLQESGPGLVKPSQTLSLTCTVSGGSISSGEYYWNWIRQHPGKGLEWIGYIYYSGSTYYNPSLKSRVTISVDTSKNQFSLKLSSVTAADTAVYYCARESVAGFDYWGQGTLVTVSS
||||||||||||.|||||||||.|||||||||||||.||||.|||||||||||||||||.||||||||.|.|.||||.||||.|.||||||||||||||.||||||||||||||||||
QVQLQESGPGLVNPSQTLSLTCNVSGGSISSGEYYWTWIRQPPGKGLEWIGYIYYSGSTSYNPSLKSRLTLSLDTSKIQFSLRLASVTAADTAVYYCARVSVAGFDYWGQGTLVTVSS
           ^^^^^^^^^^       ^^^^^^^               ^^^^^^^^^^
```

Best Alignment of Ascrinvacumab light CDRs chain to a sequence from OAS

```
EIVLTQSPGTLSLSPGERATLSCRASQSVSSSYLAWYQQKPGQAPRLLIYGTSSRATGIPDRFSGSGSGTDFTLTISRLEPEDFAVYYCQQYGSSPITFGQGTRLEIK
|.|||.||||||.|||||||||||||||||||||||||||||||||||||||||||||||||||||||||||||||||||||||||||||||||||||||||||||||
ELVLTLSPGTLSFSPGERATLSCRASQSVSSSYLAWYQQKPGQAPRLLIYGTSSRATGIPDRFSGSGSGTDFTLTISRLEPEDFAVYYCQQYGSSPITFGQGTRLEIK
           ^^^^^^^       ^^^               ^^^^^^^^^
```

Best Alignment of Ascrinvacumab CDR-H3 chain to a sequence from OAS

```
QVQLQESGPGLVKPSQTLSLTCTVSGGSISSGEYYWNWIRQHPGKGLEWIGYIYYSGSTYYNPSLKSRVTISVDTSKNQFSLKLSSVTAADTAVYYCARESVAGFDYWGQGTLVTVSS
.................|.|.|..||.......|...|.||..||||||...|...|.|||..|.|.|.|||....||...|...|..|.||||||||||||||||||||||||||||
----------------SLRLSCAASGFTF--SSYDMHWVRQATGKGLEWVSAIGTAGDTYYPGSVKGRFTISRENAKNSLYLQMNSLRAGDTAVYYCARESVAGFDYWGQGTLVTVSS
           ^^^^^^^^^^       ^^^^^^^               ^^^^^^^^^^
```

Therapeutic : Atezolizumab

Best Alignment of Atezolizumab heavy chain to a sequence from OAS

```
EVQLVESGGGLVQPGGSLRLSCAASGFTFSDSWIHWVRQAPGKGLEWVAWISPYGGSTYYADSVKGRFTISADTSKNTAYLQMNSLRAEDTAVYYCARRHWP------GGFDYWGQGTLVTVSS
||||||||||||||||||||||||||||||..|.|||||||||||||..||..|||||||||||||||||.|.||||.||||||||||||||||||||.........|.||||||||||||||
EVQLVESGGGLVQPGGSLRLSCAASGFTFSSYWMHWVRQAPGKGLEWVSAISGSGGSTYYADSVKGRFTISRDNSKNTLYLQMNSLRAEDTAVYYCARRYYDSSGYPGGDFDYWGQGTLVTVSS
           ^^^^^^^^       ^^^^^^^^               ^^^^^^^^^^^^^^^^^
```

Best Alignment of Atezolizumab light chain to a sequence from OAS

```
DIQMTQSPSSLSASVGDRVTITCRASQDVSTAVAWYQQKPGKAPKLLIYSASFLYSGVPSRFSGSGSGTDFTLTISSLQPEDFATYYCQQYLYHPATFGQGTKVEIK
|||||||||||||||||||||||||||..||..||||||||||||||||.||.|.||||||||||||||||||||||||||||||||||||...|.|||||||||||
DIQMTQSPSSLSASVGDRVTITCRASQGISTGLAWYQQKPGKAPKLLIYRASSLQSGVPSRFSGSGSGTDFTLTISSLQPEDFATYYCQQYNSTPPTFGQGTKVEIK
           ^^^^^^       ^^^               ^^^^^^^^^
```

Best Alignment of Atezolizumab heavy chain CDRs to a sequence from OAS

```
EVQLVESGGGLVQPGGSLRLSCAASGFTFSDSWIHWVRQAPGKGLEWVAWISPYGGSTYYADSVKGRFTISADTSKNTAYLQMNSLRAEDTAVYYCARRHWPGGFDYWGQGTLVTVSS
||.|||||||||||||||.||||||||||||....||||.|.|.|.|||||.||..||||||.|.||||||||.|..|||.||||..|..||||.||||||.|.||||||||||..||||
EVMLVESGGGLVQPGGSLKLSCAASGFTFSDYYMYWVRQTPEKRLEWVAYISNGGGSTYYPDTVKGRFTISRDNAKNTLYLQMSRLKSEDTAMYYCARRNWDGGFDYWGQGTTLTVSS
           ^^^^^^^^       ^^^^^^^^               ^^^^^^^^^^^
```

Best Alignment of Atezolizumab light CDRs chain to a sequence from OAS

```
DIQMTQSPSSLSASVGDRVTITCRASQDVSTAVAWYQQKPGKAPKLLIYSASFLYSGVPSRFSGSGSGTDFTLTISSLQPEDFATYYCQQYLYHPATFGQGTKVEIK
|...|.........|||||.|||.|||||||||||||||||..|||||||||....|||.||.|||||||||||||..|.||.|.||||||...|.|||.|||.|||
DFVLTRPKNFIPTPVGDRVSITCKASQDVSTAVAWYQQKPGQSPKLLIYSASNRHTGVPDRFTGSGSGTDFTLTISNVQAEDLADYYCQQYYSYPRTFGGGTKLEIK
           ^^^^^^       ^^^               ^^^^^^^^^
```

Best Alignment of Atezolizumab CDR-H3 chain to a sequence from OAS

```
EVQLVESGGGLVQPGGSLRLSCAASGFTFSDSWIHWVRQAPGKGLEWVAWISPYGGSTYYADSVKGRFTISADTSKNTAYLQMNSLRAEDTAVYYCARRHWPGGFDYWGQGTLVTVSS
................|||||||||||..|.....|||||||||||||..|....||.|||||||||||||.|.||||.|||||||||.||||||||||||.||||||||||||||||
----------------SLRLSCAASGFSVSTNYMSWVRQAPGKGLEWVSLIDSG-GSSYYADSVKGRFTISRDNSKNTLYLQMNSLRADDTAVYYCARRHWNGGFDYWGQGTLVTVSS
           ^^^^^^^^       ^^^^^^^^               ^^^^^^^^^^^
```

Therapeutic : Atidortoxumab

Best Alignment of Atidortoxumab heavy chain to a sequence from OAS

```
EVQLQESGPGLVRPSETLSLTCAVSGYSISSGMGWGWIRQPPGKGLEWIGSIDQRGSTYYNPSLKSRVTISVDTSKNQFSLKLSSVTAADTAVYYCARDAGH--AVDMDVWGKGTTVTVSS
.|||||||||||.|||||||||||||||||||..|||||||||||||||||||..|||||||||||||||||||||||||||||||||||||||||||||..||.....||||||||||||||
QVQLQESGPGLVKPSETLSLTCAVSGYSISSGYYWGWIRQPPGKGLEWIGSIDHSGSTYYNPSLKSRVTISVDTSKNQFSLKLSSVTAADTAVYYCARHSGHRRHYYMDVWGKGTTVTVSS
           ^^^^^^^^^       ^^^^^^^               ^^^^^^^^^^^^^^
```

Best Alignment of Atidortoxumab light chain to a sequence from OAS

```
DIQMTQSPSSVSASVGDRVTITCRASQGISRWLAWYQQKPGKAPKLLIYAASSLQSGVPSRFSGSGSGTDFTLTISSLQPEDFATYYCQQGYVFPLTFGGGTKVEIK
||||||||||||||||||||||||||||||||||||||||||||||||||||||||||||||||||||||||||||||||||||||||||.|.||||||||||||||
DIQMTQSPSSVSASVGDRVTITCRASQGISRWLAWYQQKPGKAPKLLIYAASSLQSGVPSRFSGSGSGTDFTLTISSLQPEDFATYYCQQAYSFPLTFGGGTKVEIK
           ^^^^^^       ^^^               ^^^^^^^^^
```

Best Alignment of Atidortoxumab heavy chain CDRs to a sequence from OAS

```
EVQLQESGPGLVRPSETLSLTCAVSGYSISSGMGWGWIRQPPGKGLEWIGSIDQRGSTYYNPSLKSRVTISVDTSKNQFSLKLSSVTAADTAVYYCARDAGHAVDMDVWGKGTTVTVSS
.........|||.|||||||||.|||||||||..||||||||||||||||||...||||||.|||||||||||||||||||||.|||||||||||||||.||...|||||.||||||||
---------GLVKPSETLSLTCTVSGYSISSGYFWGWIRQPPGKGLEWIGSIYHSGSTYYNSSLKSRVTISVDTSKNQFSLKLGSVTAADTAVYYCARDGGHYFGMDVWGQGTTVTVSS
           ^^^^^^^^^       ^^^^^^^               ^^^^^^^^^^^^
```

Best Alignment of Atidortoxumab light CDRs chain to a sequence from OAS

```
DIQMTQSPSSVSASVGDRVTITCRASQGISRWLAWYQQKPGKAPKLLIYAASSLQSGVPSRFSGSGSGTDFTLTISSLQPEDFATYYCQQGYVFPLTFGGGTKVEIK
||||||||||.||||||.||||||||||||||..|||||||||||||.|||||.|||.|||||||||||...||||||||||||||||||||..||||||||||.||
DIQMTQSPSSLSASVGDKVTITCRASQGISRWVDWYQQKPGKAPKLLSYAASSWQSGAPSRFSGSGSGTVYPLTISSLQPEDFATYYCQQGYNTPLTFGGGTKVAIK
           ^^^^^^       ^^^               ^^^^^^^^^
```

Best Alignment of Atidortoxumab CDR-H3 chain to a sequence from OAS

```
EVQLQESGPGLVRPSETLSLTCAVSGYSISSGMGWGWIRQPPGKGLEWIGSIDQR-GSTYYNPSLKSRVTISVDTSKNQFSLKLSSVTAADTAVYYCARDAGHAVDMDVWGKGTTVTVSS
...............|.|...||.||..........|.||.|||||||...|....|...|..|...|.|.|.|.||....|...|....||||||||||.|.|||||||||||||.|||
--------------GESLKISCATSGFIF-RDYAMHWVRQAPGKGLEWVALISFDRGTQHYADSVRGRFTVSSDSSKDTVFLQMNSLRPEDTAVYYCARDRGIAVDMDVWGKGTTVIVSS
           ^^^^^^^^^       ^^^^^^^^               ^^^^^^^^^^^^
```

Therapeutic : Avelumab

Best Alignment of Avelumab heavy chain to a sequence from OAS

```
EVQLLESGGGLVQPGGSLRLSCAASGFTFSSYIMMWVRQAPGKGLEWVSSIYPSGGITFYADTVKGRFTISRDNSKNTLYLQMNSLRAEDTAVYYCARIKLGT-VTTVDYWGQGTLVTVSS
||||||||||||||||||||||||||||||||.|.||||||||||||||.|..|||.|.|||.|||||||||||||||||||||||||||||||||||..||..|||.|||||||||||||
EVQLLESGGGLVQPGGSLRLSCAASGFTFSSYAMSWVRQAPGKGLEWVSAISGSGGSTYYADSVKGRFTISRDNSKNTLYLQMNSLRAEDTAVYYCARDHLGATVTTADYWGQGTLVTVSS
                      ^^^^^^^^                ^^^^^^^^                                  ^^^^^^^^^^^^^^
```

Best Alignment of Avelumab light chain to a sequence from OAS

```
QSALTQPASVSGSPGQSITISCTGTSSDVGGYNYVSWYQQHPGKAPKLMIYDVSNRPSGVSNRFSGSKSGNTASLTISGLQAEDEADYYCSSYTSSTRVFGTGTKVTVL
|||||||||||||||||||||||||||||||||||||||||||||||||||||||||||||||||||||||||||||||||||||||||||||||||||||||||||||
QSALTQPASVSGSPGQSITISCTGTSSDVGGYNYVSWYQQHPGKAPKLMIYDVSNRPSGVSNRFSGSKSGNTASLTISGLQAEDEADYYCSSYTSSTRVFGTGTKVTVL
                      ^^^^^^^^^                ^^^                                 ^^^^^^^^^^
```

Best Alignment of Avelumab heavy chain CDRs to a sequence from OAS

```
EVQLLESGGGLVQPGGSLRLSCAASGFTFSSYIMMWVRQAPGKGLEWVSSIYPSGGITFYADTVKGRFTISRDNSKNTLYLQMNSLRAEDTAVYYCARIKLGTVTTVDYWGQGTLVTVSS
................||||||||||||||||.|.||||||||||||||.|..|||.|.|||.|||||||||||.||.||||||||||||||||||||...|||||||||||||||||||
----------------SLRLSCAASGFTFSSYAMSWVRQAPGKGLEWVSAISGSGGSTYYADSVKGRFTISRDNAKNSLYLQMNSLRAEDTAVYYCARSSGGTVTTVDYWGQGTLVTVSS
                      ^^^^^^^^                ^^^^^^^^                                  ^^^^^^^^^^^^^
```

Best Alignment of Avelumab light CDRs chain to a sequence from OAS

```
QSALTQPASVSGSPGQSITISCTGTSSDVGGYNYVSWYQQHPGKAPKLMIYDVSNRPSGVSNRFSGSKSGNTASLTISGLQAEDEADYYCSSYTSSTRVFGTGTKVTVL
||||||||||||||||||||||||||||||||||||||.||||||||||||||||||||||||||||||||||||||||||.|||||||||||||||||||.|||.|||
QSALTQPASVSGSPGQSITISCTGTSSDVGGYNYVSWCQQHPGKAPKLMIYDVSNRPSGVSNRFSGSKSGNTASLTISGLQTEDEADYYCSSYTSSTRVFGGGTKLTVL
                      ^^^^^^^^^                ^^^                                 ^^^^^^^^^^
```

Best Alignment of Avelumab CDR-H3 chain to a sequence from OAS

```
EVQLLESGGGLVQPGGSLRLSCAASGFTF--SSYIMMWVRQAPGKGLEWVSSIYPSGGITFYADTVKGRFTISRDNSKNTLYLQMNSLRAEDTAVYYCARIKLGTVTTVDYWGQGTLVTVSS
.............|...|.|.|..|||....|.....|.||.|||.|||...|........|....|.|.|||.|.|||...|.|......|||.|||||||..||||||||||||||||||
------------KPTQTLTLTCTFSGFSLSTSGMCVSWIRQPPGKALEWLARIDWD-DDKYYSTSLKTRLTISKDTSKNQVVLTMTNMDPVDTATYYCARIKMTTVTTVDYWGQGTLVTVSS
                      ^^^^^^^^^^                ^^^^^^^^                                  ^^^^^^^^^^^^^
```

Therapeutic : Bapineuzumab

Best Alignment of Bapineuzumab heavy chain to a sequence from OAS

```
EVQLLESGGGLVQPGGSLRLSCAASGFTFSNYGMSWVRQAPGKGLEWVASIRSGGGRTYYSDNVKGRFTISRDNSKNTLYLQMNSLRAEDTAVYYCVRYDHY-SGSSDYWGQGTLVTVSS
||||||||||||||||||||||||||||||.|.|||||||||||||||..|.|||..|||.|.|||||||||||||||||||||||||||||||||...|.|.|||||||||||||||||
EVQLLESGGGLVQPGGSLRLSCAASGFTFSSYAMSWVRQAPGKGLEWVSVIYSGGSSTYYADSVKGRFTISRDNSKNTLYLQMNSLRAEDTAVYYCAKVDYYGSGSSDYWGQGTLVTVSS
                      ^^^^^^^^                ^^^^^^^^                                  ^^^^^^^^^^^^^
```

Best Alignment of Bapineuzumab light chain to a sequence from OAS

```
DVVMTQSPLSLPVTPGEPASISCKSSQSLLDSDGKTYLNWLLQKPGQSPQRLIYLVSKLDSGVPDRFSGSGSGTDFTLKISRVEAEDVGVYYCWQGTHFPRTFGQGTKVEIK
|||||||||.|.||.|.||||||||||||||||||||||||||.|||||.|||||||||||||||||.|||||||||||||||||||.||||||||||||||||.|||.|||
DVVMTQSPLTLSVTIGQPASISCKSSQSLLDSDGKTYLNWLLQRPGQSPKRLIYLVSKLDSGVPDRFTGSGSGTDFTLKISRVEAEDLGVYYCWQGTHFPRTFGGGTKLEIK
                       ^^^^^^^^^^^                ^^^                                 ^^^^^^^^^
```

Best Alignment of Bapineuzumab heavy chain CDRs to a sequence from OAS

```
EVQLLESGGGLVQPGGSLRLSCAASGFTFSNYGMSWVRQAPGKGLEWVASIRSGGGRTYYSDNVKGRFTISRDNSKNTLYLQMNSLRAEDTAVYYCVRYDHYSGSSDYWGQGTLVTVSS
||.|.|||||||||||||.|||||||||||.|.|.||||.|.|.|||||.||.|||||||.|.|||||||||||.||||||||..|..||||.|||.|..||.||||||||||..||||
EVMLVESGGGLVQPGGSLKLSCAASGFTFSDYYMYWVRQTPEKRLEWVAYIRNGGGRTYYPDTVKGRFTISRDNAKNTLYLQMSRLKSEDTAMYYCARGGHYYGSSDYWGQGTTLTVSS
                      ^^^^^^^^                ^^^^^^^^                                  ^^^^^^^^^^^^
```

Best Alignment of Bapineuzumab light CDRs chain to a sequence from OAS

```
DVVMTQSPLSLPVTPGEPASISCKSSQSLLDSDGKTYLNWLLQKPGQSPQRLIYLVSKLDSGVPDRFSGSGSGTDFTLKISRVEAEDVGVYYCWQGTHFPRTFGQGTKVEIK
||||||.||.|.||.|.|||.||||||||||||||||||||||.|||||.|||||||||||||||||.|||||||||||||||||||.||||||||||||||||.|||.|||
DVVMTQTPLTLSVTIGQPASFSCKSSQSLLDSDGKTYLNWLLQRPGQSPKRLIYLVSKLDSGVPDRFTGSGSGTDFTLKISRVEAEDLGVYYCWQGTHFPRTFGGGTKLEIK
                       ^^^^^^^^^^^                ^^^                                 ^^^^^^^^^
```

Best Alignment of Bapineuzumab CDR-H3 chain to a sequence from OAS

```
EVQLLESGGGLVQPGGSLRLSCAASGFTFSNYGMSWVRQAPGKGLEWVASIRSGGGRTYYSDNVKGRFTISRDNSKNTLYLQMNSLRAEDTAVYYCVRYDHYSGSSDYWGQGTLVTVSS
.|||...|..||.||.|..|||.|||.||..|.|.||.|.||.||||...|....|.|.|....|...|...|.|..|.|.|..||..||.|||||.|.||||||||||||||..||||
QVQLQQPGAELVKPGASVKLSCKASGYTFTSYWMHWVKQRPGQGLEWIGMIHPNSGSTNYNEKFKSKATLTVDKSSSTAYMQLSSLTSEDSAVYYCARCDHYSGSSDYWGQGTTLTVSS
                      ^^^^^^^^                ^^^^^^^^                                  ^^^^^^^^^^^^
```

Therapeutic : Basiliximab

Best Alignment of Basiliximab heavy chain to a sequence from OAS

```
QVQLQQSGTVLARPGASVKMSCKASGYSFTRYWMHWIKQRPGQGLEWIGAIYPGNSDTSYNQKFEGKAKLTAVTSASTAYMELSSLTHEDSAVYYCSRDYG--YYFDFWGQGTTLTVSS
.|||||||||||||||||||||||||||||.|||||.|||||||||||||||||||||||||||.||||||||||||||||||||||.||||||||.||||..||||.|||||||||||
EVQLQQSGTVLARPGASVKMSCKASGYSFTSYWMHWVKQRPGQGLEWIGAIYPGNSDTSYNQKFKGKAKLTAVTSASTAYMELSSLTNEDSAVYYCTRDYGSPYYFDYWGQGTTLTVSS
                      ^^^^^^^^                ^^^^^^^^                                  ^^^^^^^^^^^^
```

Best Alignment of Basiliximab light chain to a sequence from OAS

```
QIVSTQSPAIMSASPGEKVTMTCSASSSRSYMQWYQQKPGTSPKRWIYDTSKLASGVPARFSGSGSGTSYSLTISSMEAEDAATYYCHQRSSYTFGGGTKLEIK
|||.||||||||||||||||||||||||.|||.|||||||||||||||||||||||||||||||||||||||||||||||||||||||||||||||||||||||
QIVLTQSPAIMSASPGEKVTMTCSASSSISYMHWYQQKPGTSPKRWIYDTSKLASGVPARFSGSGSGTSYSLTISSMEAEDAATYYCHQRSSYTFGGGTKLEIK
                       ^^^^^                ^^^                                 ^^^^^^^
```

Best Alignment of Basiliximab heavy chain CDRs to a sequence from OAS

```
QVQLQQSGTVLARPGASVKMSCKASGYSFTRYWMHWIKQRPGQGLEWIGAIYPGNSDTSYNQKFEGKAKLTAVTSASTAYMELSSLTHEDSAVYYCSRDYGYYFDFWGQGTTLTVSS
.|.||.||||||||||||||||||||||||.|||||.|||||||||||||||||||||||||||.||||||||||||||||.|||||.||||||||.||||||||.|||||||||||
EVKLQESGTVLARPGASVKMSCKASGYSFTSYWMHWVKQRPGQGLEWIGAIYPGNSDTSYNQKFKGKAKLTAVTSASTAYMQLSSLTSEDSAVYYCARDYGYYFDYWGQGTTLTVSS
                      ^^^^^^^^                ^^^^^^^^                                  ^^^^^^^^^^
```

Best Alignment of Basiliximab light CDRs chain to a sequence from OAS

```
QIVSTQSPAIMSASPGEKVTMTCSASSSRSYMQWYQQKPGTSPKRWIYDTSKLASGVPARFSGSGSGTSYSLTISSMEAEDAATYYCHQRSSYTFGGGTKLEIK
||..||||||||||||||||||||||||.|||.|||||||||||||||||||||.|||||||||||||||.|||||||||||||||.|||||||||||||||||
QILLTQSPAIMSASPGEKVTMTCSASSSISYMHWYQQKPGTSPKRWIYDTSKLAAGVPARFSGSGSGTSYALTISSMEAEDAATYYGHQRSSYTFGGGTKLEIK
                       ^^^^^                ^^^                                 ^^^^^^^
```

Best Alignment of Basiliximab CDR-H3 chain to a sequence from OAS

```
QVQLQQSGTVLARPGASVKMSCKASGYSFTRYWMHWIKQRPGQGLEWIGAIYPGNSDTSYNQKFEGKAKLTAVTSASTAYMELSSLTHEDSAVYYCSRDYGYYFDFWGQGTTLTVSS
||...|.|..|..||||||.|||||||.||.|||||.||||||||||||.|.|....|.||.||..||.||...|.|||||.|||||.||||||.||||||||||.|||||||||||
QVHVKQPGAELVKPGASVKLSCKASGYTFTSYWMHWVKQRPGQGLEWIGMIHPNSGSTNYNEKFKSKATLTVDKSSSTAYMQLSSLTSEDSAVYSCSRDYGYYFDYWGQGTTLTVSS
                      ^^^^^^^^                ^^^^^^^^                                  ^^^^^^^^^^
```

Therapeutic : Bavituximab

Best Alignment of Bavituximab heavy chain to a sequence from OAS

```
EVQLQQSGPELEKPGASVKLSCKASGYSFTGYNMNWVKQSHGKSLEWIGHIDPYYGDTSYNQKFRGKATLTVDKSSSTAYMQLKSLTSEDSAVYYCVKGGYYG--HWYFDVWGAGTTVTVSS
|||||||||||||||||||.||||||||||||||||||||.||||||||.||||||.|||||||.|||||||||||||||||||||||||||||||..|||||..|||||||||||||||||
EVQLQQSGPELEKPGASVKISCKASGYSFTGYNMNWVKQSNGKSLEWIGNIDPYYGGTSYNQKFKGKATLTVDKSSSTAYMQLKSLTSEDSAVYYCARGGYYGSSHWYFDVWGAGTTVTVSS
                      ^^^^^^^^                ^^^^^^^^                                  ^^^^^^^^^^^^^^^
```

Best Alignment of Bavituximab light chain to a sequence from OAS

```
DIQMTQSPSSLSASLGERVSLTCRASQDIGSSLNWLQQGPDGTIKRLIYATSSLDSGVPKRFSGSRSGSDYSLTISSLESEDFVDYYCLQYVSSPPTFGAGTKLELK
||||||||||||||||||||||||||||||||||||||.||||||||||||||||||||||||||||||||||||||||||||||||||||.|||||||||||||||
DIQMTQSPSSLSASLGERVSLTCRASQDIGSSLNWLQQEPDGTIKRLIYATSSLDSGVPKRFSGSRSGSDYSLTISSLESEDFVDYYCLQYASSPPTFGAGTKLELK
                          ^^^^^^                 ^^^                                    ^^^^^^^^^
```

Best Alignment of Bavituximab heavy chain CDRs to a sequence from OAS

```
EVQLQQSGPELEKPGASVKLSCKASGYSFTGYNMNWVKQSHGKSLEWIGHIDPYYGDTSYNQKFRGKATLTVDKSSSTAYMQLKSLTSEDSAVYYCVKGGYYGHWYFDVWGAGTTVTVSS
.|||||.|.||..|||||||||||||||||||||||||||.||||||||.||||||.|||||||.|||||||||||||||||||||||||||||||...||||.||||||||||||||||
-VQLQQPGAELVRPGASVKLSCKASGYSFTGYNMNWVKQSNGKSLEWIGNIDPYYGGTSYNQKFKGKATLTVDKSSSTAYMQLKSLTSEDSAVYYCARFGYYGNWYFDVWGAGTTVTVSS
                         ^^^^^^^^                 ^^^^^^^^                                      ^^^^^^^^^^^^^
```

Best Alignment of Bavituximab light CDRs chain to a sequence from OAS

```
DIQMTQSPSSLSASLGERVSLTCRASQDIGSSLNWLQQGPDGTIKRLIYATSSLDSGVPKRFSGSRSGSDYSLTISSLESEDFVDYYCLQYVSSPPTFGAGTKLELK
|||||||||||||||||||||||||.||||||.|||||.||||.|||||||||||||.||||||||||||||||||||||||.||||||||.|||||||.|||||.|
DIQMTQSPSSLSASLGERVSLTCRARQDIGSSVNWLQQEPDGTNKRLIYATSSLDSGDPKRFSGSRSGSDYSLTISSLESEDCVDYYCLQYASSPPTFGGGTKLESK
                          ^^^^^^                 ^^^                                    ^^^^^^^^^
```

Best Alignment of Bavituximab CDR-H3 chain to a sequence from OAS

```
EVQLQQSGPELEKPGASVKLSCKASGYSFTGYNMNWVKQSHGKSLEWIGHIDPYYGDTSYNQKFRGKATLTVDKSSSTAYMQLKSLTSEDSAVYYCVKGGYYGHWYFDVWGAGTTVTVSS
|||||||||||.|||||.|.||||||||||||.||||||||||.|||||.|.||.|.|||||||.||||||||.|||||||.|.||||||||||||.|||||||||||||||||||||||
EVQLQQSGPELVKPGASMKISCKASGYSFTGYTMNWVKQSHGKNLEWIGLINPYNGGTSYNQKFKGKATLTVDESSSTAYMELLSLTSEDSAVYYCAKGGYYGHWYFDVWGAGTTVTVSS
                         ^^^^^^^^                 ^^^^^^^^                                      ^^^^^^^^^^^^^
```

Therapeutic : Belimumab

Best Alignment of Belimumab heavy chain to a sequence from OAS

```
QVQLQQSGAEVKKPGSSVRVSCKASGGTFNNNAINWVRQAPGQGLEWMGGIIPMFGTAKYSQNFQGRVAITADESTGTASMELSSLRSEDTAVYYCARSRDLLL-----FPHHALSPWGRGTMVTVSS
||||.|||||||||||||.||||||||||...||.||||||||||||||||||.||||.|.|.|.|||||.||||||||||.||||||||||||||||||.|..|........|||...||.||||||||
QVQLVQSGAEVKKPGSSVKVSCKASGGTFSSYAISWVRQAPGQGLEWMGGIIPIFGTANYAQKFQGRVTITADESTGTAYMELSSLRSEDTAVYYCARDRQVLRYFDWLDRHHAFDIWGQGTMVTVSS
                         ^^^^^^^^                 ^^^^^^^^                                      ^^^^^^^^^^^^^^^^^^^^^
```

Best Alignment of Belimumab light chain to a sequence from OAS

```
SSELTQDPAVSVALGQTVRVTCQGDSLRSYYASWYQQKPGQAPVLVIYGKNNRPSGIPDRFSGSSSGNTASLTITGAQAEDEADYYCSSRDSSGNHWVFGGGTELTVL
|||||||||||||||||||||||||||||||||||||||||||||||||||||||||||||||||||||||||||||||||||||||.|||||||||||||||.||||
SSELTQDPAVSVALGQTVRVTCQGDSLRSYYASWYQQKPGQAPVLVIYGKNNRPSGIPDRFSGSSSGNTASLTITGAQAEDEADYYCNSRDSSGNHWVFGGGTKLTVL
                         ^^^^^^                 ^^^                                    ^^^^^^^^^^^
```

Best Alignment of Belimumab heavy chain CDRs to a sequence from OAS

```
QVQLQQSGAEVKKPGSSVRVSCKASGGTFNNNAINWVRQAPGQGLEWMGGIIPMFGTAKYSQNFQGRVAITADESTGTASMELSSLRSEDTAVYYCARSRDLLLFPHHALSPWGRGTMVTVSS
................||.||||||||||...||.||||||||||||||||||.||||.|.|.|.|||||..|.|.|.||.||.|||.|||..|||||||||..|..|.||.||..||.||||||||
----------------SVKVSCKASGGTFSSYAISWVRQAPGQGLEWMGGIIPIFGTANYAQKFQGRVTMTTDTSTSTAYMELRSLRFDDTAVYYCARFVDRALAPHGALDIWGQGTMVTVSS
                         ^^^^^^^^                 ^^^^^^^^                                      ^^^^^^^^^^^^^^^^
```

Best Alignment of Belimumab light CDRs chain to a sequence from OAS

```
SSELTQDPAVSVALGQTVRVTCQGDSLRSYYASWYQQKPGQAPVLVIYGKNNRPSGIPDRFSGSSSGNTASLTITGAQAEDEADYYCSSRDSSGNHWVFGGGTELTVL
|||||||||||||||||||.|||||||||||||||||||||||||||||||||||||||||||||||||||||||||||||||||||||||||||||||.||..|||
SSELTQDPAVSVALGQTVRITCQGDSLRSYYASWYQQKPGQAPVLVIYGKNNRPSGIPDRFSGSSSGNTASLTITGAQAEDEADYYCSSRDSSGNHWVFGTGTKVTVL
                         ^^^^^^                 ^^^                                    ^^^^^^^^^^^
```

Best Alignment of Belimumab CDR-H3 chain to a sequence from OAS

```
QVQLQQSGAEVKKPGSSVRVSCKASGGTFNNNAINWVRQAPGQGLEWMGGIIPMFGTAKYSQNFQGRVAITADESTGTASMELSSLRSEDTAVYYCARSRDLLLFPHHALSPWGRGTMVTVSS
.|||||||.|..|||.||..||||||.||....||||.|.|||||||.|.|.|.|..|..||...|.|....|.|.|..||.|||.||.|||.|||.|||.|.||||||.|...||.||.|||||
-VQLQQSGPELVKPGASVKLSCKASGYTFTSYDINWVKQRPGQGLEWIGWIYPRDGSTKYNEKFKGKATLTVDTSSSTAYMELHSLTSEDSAVYFCARERGLLLFPHYAMDYWGQGTSVTVSS
                         ^^^^^^^^                 ^^^^^^^^                                      ^^^^^^^^^^^^^^^^
```

Therapeutic : Benralizumab

Best Alignment of Benralizumab heavy chain to a sequence from OAS

```
EVQLVQSGAEVKKPGASVKVSCKASGYTFTSYVIHWVRQRPGQGLAWMGYINPYNDGTKYNERFKGKVTITSDRSTSTVYMELSSLRSEDTAVYLCGREGIRYYGLLGDYWGQGTLVTVSS
||||.|||.|..|||||||.|||||||||||||.|||.|.|||||.|.||||||||||||||.||||.|.|||.|.||.|||||||.|||.|||.|.||||.|||...|||||||.|||||
EVQLQQSGPELVKPGASVKMSCKASGYTFTSYVMHWVKQKPGQGLEWIGYINPYNDGTKYNEKFKGKATLTSDKSSSTAYMELSSLTSEDSAVYYCAREGIFYYGGAMDYWGQGTSVTVSS
                         ^^^^^^^^                 ^^^^^^^^                                      ^^^^^^^^^^^^^^
```

Best Alignment of Benralizumab light chain to a sequence from OAS

```
DIQMTQSPSSLSASVGDRVTITCGTSEDIINYLNWYQQKPGKAPKLLIYHTSRLQSGVPSRFSGSGSGTDFTLTISSLQPEDFATYYCQQGYT-LPYTFGQGTKVEIK
|||||||||||||||||||||||..|.||.|||||||||||||||||||..|.|||||||||||||||||||||||||||||||||||||.|...|||||||||||||
DIQMTQSPSSLSASVGDRVTITCQASQDISNYLNWYQQKPGKAPKLLIYAASSLQSGVPSRFSGSGSGTDFTLTISSLQPEDFATYYCQQFYSTPPYTFGQGTKVEIK
                          ^^^^^^                 ^^^                                    ^^^^^^^^^^
```

Best Alignment of Benralizumab heavy chain CDRs to a sequence from OAS

```
EVQLVQSGAEVKKPGASVKVSCKASGYTFTSYVIHWVRQRPGQGLAWMGYINPYNDGTKYNERFKGKVTITSDRSTSTVYMELSSLRSEDTAVYLCGREGIRYYGLLGDYWGQGTLVTVSS
||||.|||.|..|||||||.|||||||||||||.|||.|.|||||.|.||||||||||||||.||||.|.|||.|.||.|||||||.|||.|||.|.||||.|||...|||||||.|||||
EVQLQQSGPELVKPGASVKMSCKASGYTFTSYVMHWVKQKPGQGLEWIGYINPYNDGTKYNEKFKGKATLTSDKSSSTAYMELSSLTSEDSAVYYCAREGIFYYGGAMDYWGQGTSVTVSS
                         ^^^^^^^^                 ^^^^^^^^                                      ^^^^^^^^^^^^^^
```

Best Alignment of Benralizumab light CDRs chain to a sequence from OAS

```
DIQMTQSPSSLSASVGDRVTITCGTSEDIINYLNWYQQKPGKAPKLLIYHTSRLQSGVPSRFSGSGSGTDFTLTISSLQPEDFATYYCQQGYTLPYTFGQGTKVEIK
||||||..||||||.||||||.|..|.|||||||||||||....|||||.||||.|||||||||||||||..||||.|..||.|||.||||.|||||||.|||.|||
DIQMTQTTSSLSASLGDRVTISCRASQDIINYLNWYQQKPDGTVKLLIYYTSRLHSGVPSRFSGSGSGTDYSLTISNLEQEDIATYFCQQGNTLPYTFGGGTKLEIK
                          ^^^^^^                 ^^^                                    ^^^^^^^^^
```

Best Alignment of Benralizumab CDR-H3 chain to a sequence from OAS

```
EVQLVQSGAEVKKPGASVKVSCKASGYTFTSYVIHWVRQRPGQGLAWMGYINPYNDGTKYNERFKGKVTITSDRSTSTVYMELSSLRSEDTAVYLCGREGIRYYGLLGDYWGQGTLVTVSS
.|||.|.|||...||.|||.||||||||||||..|||.|||.|||.|.|.|.|.|....|.||..||.|.|.|.|.|.|.||.||.||||.|||.|||.|.||||.|||..||||||||..||||
-VQLQQPGAELVRPGSSVKLSCKASGYTFTSYWMHWVKQRPIQGLEWIGNIDPSDSETHYNQKFKDKATLTVDKSSSTAYMQLSSLTSEDSAVYYCAREGIYYYGSSGDYWGQGTTLTVSS
                         ^^^^^^^^                 ^^^^^^^^                                      ^^^^^^^^^^^^^^
```

Therapeutic : Berlimatoxumab

Best Alignment of Berlimatoxumab heavy chain to a sequence from OAS

```
ELQLQESGPGLVKPSETLSLTCTVSGGSISSGSYYWDWIRQPPGKGLEWIGNIYKSGSTYYNPSLKSRVTISVDTSKNQFSLKLSSVTAADTAVYYCARERGM----------HYMDVWGKGTTVTVSS
.||||||||||||||||||||||||||||||.||||.||||||||||||||.||.|||||||||||||||||||||||||||||||||||||||||||||||............|||||||||||||||
QLQLQESGPGLVKPSETLSLTCTVSGGSISSSSYYWGWIRQPPGKGLEWIGSIYYSGSTYYNPSLKSRVTISVDTSKNQFSLKLSSVTAADTAVYYCARERGYCSGGSCRYYYYYMDVWGKGTTVTVSS
                         ^^^^^^^^^^                 ^^^^^^^                                      ^^^^^^^^^^^^^^^^^^^^^
```

Best Alignment of Berlimatoxumab light chain to a sequence from OAS

```
DIQMTQSPSSLSASVGDRVTITCRASQSINSYLNWYQQKPGKAPKLLIYAASSLQSGVPSRFSGSGSGTDFTLTISSLQPEDFATYYCQQQFD-PPFTFGGGTKVEIK
|||||||||||||||||||||||||||||.||||||||||||||||||||||||||||||||||||||||||||||||||||||||||||....||||||||||||||
DIQMTQSPSSLSASVGDRVTITCRASQSISSYLNWYQQKPGKAPKLLIYAASSLQSGVPSRFSGSGSGTDFTLTISSLQPEDFATYYCQQSYSTPPFTFGGGTKVEIK
                          ^^^^^^                 ^^^                                    ^^^^^^^^^^
```

Best Alignment of Berlimatoxumab heavy chain CDRs to a sequence from OAS

```
ELQLQESGPGLVKPSETLSLTCTVSGGSISSGSYYWDWIRQPPGKGLEWIGNIYKSGSTYYNPSLKSRVTISVDTSKNQFSLKLSSVTAADTAVYYCARERGMHYMDVWGKGTTVTVSS
.............|||||||||||||||||||||||.||||||||||||||.||.|||||||||||||||||||||||||||||||||||||||||||||||..||||||||||||||.
-------------PSETLSLTCTVSGGSISSGSYYWSWIRQPPGKGLEWIGYIYYSGSTYYNPSLKSRVTISVDTSKNQFSLKLSSVTAADTAVYYCARERGGYYMDVWGKGTTVTVS-
                         ^^^^^^^^^^                 ^^^^^^^                                      ^^^^^^^^^^^
```

Best Alignment of Berlimatoxumab light CDRs chain to a sequence from OAS
DIQMTQSPSSLSASVGDRVTITCRASQSINSYLNWYQQKPGKAPKLLIYAASSLQSGVPSRFSGSGSGTDFTLTISSLQPEDFATYYCQQQFDPPFTFGGGTKVEIK
|||||.||||||||||||||||||||||||||||||||||||..||||||||.||||||||.|||||||||.||||||||.|||.|||..|.|||||.||.|..|
DIQMTPSPSSLSASVGDRVTITCRASQSINSYLNWYQQKPGKAPNVLIYAASSLESGVPSRFSASGSGTDFTLIISSLQPEDSATYFCQQSDDVPFTFGPGTAVDVK
^^^^^^ ^^^ ^^^^^^^^^

Best Alignment of Berlimatoxumab CDR-H3 chain to a sequence from OAS
ELQLQESGPGLVKPSETLSLTCTVSGGSISSGSYYWDWIRQPPGKGLEWIGNIYKS-GSTYYNPSLKSRVTISVDTSKNQFSLKLSSVTAADTAVYYCARERGMHYMDVWGKGTTVTVSS
.............|...|.|.|..||......||...|.||.|||||||...|..|.|||||..|.|.|.|||.|.|||...|...|..|.||||||||||||.|||||||||||||||.
-------------PRGSLRLSCAASGFTF--SSYAMSWVRQAPGKGLEWVSAISGSGGSTYYADSVKGRFTISRDNSKNTLYLQMNSLRAEDTAVYYCARERGEHYMDVWGKGTTVTVS-
^^^^^^^^^^ ^^^^^^^^ ^^^^^^^^^^^

Therapeutic : Bevacizumab

Best Alignment of Bevacizumab heavy chain to a sequence from OAS
EVQLVESGGGLVQPGGSLRLSCAASGYTFTNYGMNWVRQAPGKGLEWVGWINTYTGEPTYAADFKRRFTFSLDTSKSTAYLQMNSLRAEDTAVYYCAKYPHYYGSSHWYFDVWGQGTLVTVSS
.||||||||||||||||||||||||||||.|.||||||||||.||||.|||||.||.||||..|..||.||||||.||||||..||.||||||||||..|.........||.|||||||||||
-VQLVESGGGLVQPGGSLRLSCAASGYTFTSYAMNWVRQAPGQGLEWMGWINTNTGNPTYAQGFTGRFVFSLDTSVSTAYLQISSLKAEDTAVYYCARDPTITI-FEYWFDPWGQGTLVTVSS
^^^^^^^^ ^^^^^^^^ ^^^^^^^^^^^^^^^^

Best Alignment of Bevacizumab light chain to a sequence from OAS
DIQMTQSPSSLSASVGDRVTITCSASQDISNYLNWYQQKPGKAPKVLIYFTSSLHSGVPSRFSGSGSGTDFTLTISSLQPEDFATYYCQQYSTVPWTFGQGTKVEIK
|||||||||||||||||||||||.|||||||||||||||||||||||||.||.|.||||||||||||||||||||||||||||||||||||||.||.|.|||||||||||
DIQMTQSPSSLSASVGDRVTITCQASQDISNYLNWYQQKPGKAPKVLIYGTSILQSGVPSRFSGSGSGTDFTLTISSLQPEDFATYYCQQSSTTPRTFGQGTKVEIK
^^^^^^ ^^^ ^^^^^^^^^

Best Alignment of Bevacizumab heavy chain CDRs to a sequence from OAS
EVQLVESGGGLVQPGGSLRLSCAASGYTFTNYGMNWVRQAPGKGLEWVGWINTYTGEPTYAADFKRRFTFSLDTSKSTAYLQMNSLRAEDTAVYYCAKYPHYYGSSHWYFDVWGQGTLVTVSS
..|||.||..|..||.....||.|||||||||||||||.|||||||.|.|||||||||||||.|||.||.|||.||.||||||.|.|..||.|.|.||.||.|||||.|||||||.||.|||||
QIQLVQSGPELKKPGETVKISCKASGYTFTNYGMNWVKQAPGKGLKWMGWINTYTGEPTYADDFKGRFAFSLETSASTAYLQINNLKNEDMATYFCARYPFYYGSSYWYFDVWGAGTTVTVSS
^^^^^^^^ ^^^^^^^^ ^^^^^^^^^^^^^^^^

Best Alignment of Bevacizumab light CDRs chain to a sequence from OAS
DIQMTQSPSSLSASVGDRVTITCSASQDISNYLNWYQQKPGKAPKVLIYFTSSLHSGVPSRFSGSGSGTDFTLTISSLQPEDFATYYCQQYSTVPWTFGQGTKVEIK
||||||..||||||.||||||.||||||||||||||||||....|.||||||||||||||||||||||||..||||.|.|||.|||||||||..|||||.|||.|||
DIQMTQTTSSLSASLGDRVTISCSASQDISNYLNWYQQKPDGTVKLLIYFTSSLHSGVPSRFSGSGSGTDYFLTISNLEPEDIATYYCQQYSKLPWTFGGGTKLEIK
^^^^^^ ^^^ ^^^^^^^^^

Best Alignment of Bevacizumab CDR-H3 chain to a sequence from OAS
EVQLVESGGGLVQPGGSLRLSCAASGYTFTNYGMNWVRQAPGKGLEWVGWINTYTGEPTYAADFKRRFTFSLDTSKSTAYLQMNSLRAEDTAVYYCAKYPHYYGSSHWYFDVWGQGTLVTVSS
.|||..||..||.||.|...||.||||.|..|.||||.|.|||||||.|.|....|...|...||...|...|.|.||||.|..||..||.|||.||.||||||||||||||||.||.|||||
QVQLQQSGAELVKPGASVKISCKASGYAFSSYWMNWVKQRPGKGLEWIGQIYPGDGDTNYNGKFKGKATLTADKSSSTAYMQLSSLTSEDSAVYFCARYPHYYGSSHWYFDVWGTGTTVTVSS
^^^^^^^^ ^^^^^^^^ ^^^^^^^^^^^^^^^^

Therapeutic : Bezlotoxumab

Best Alignment of Bezlotoxumab heavy chain to a sequence from OAS
EVQLVQSGAEVKKSGESLKISCKGSGYSFTSYWIGWVRQMPGKGLEWMGIFYPGDSSTRYSPSFQGQVTISADKSVNTAYLQWSSLKASDTAMYYCARRRNW---GNAFDIWGQGTMVTVSS
|||||||||||||.||||||||||||||||||||||||||||||||||||.|||||.||||||||||||||||||..||||||||||||||||||||||||||.....|||||||||||||||
EVQLVQSGAEVKKPGESLKISCKGSGYSFTSYWIGWVRQMPGKGLEWMGIIYPGDSDTRYSPSFQGQVTISADKSISTAYLQWSSLKASDTAMYYCARRRNWNYVRGAFDIWGQGTMVTVSS
^^^^^^^^ ^^^^^^^^ ^^^^^^^^^^^^^^^

Best Alignment of Bezlotoxumab light chain to a sequence from OAS
EIVLTQSPGTLSLSPGERATLSCRASQSVSSSYLAWYQQKPGQAPRLLIYGASSRATGIPDRFSGSGSGTDFTLTISRLEPEDFAVYYCQQYGSSTWTFGQGTKVEIK
||||||||||||||||||||||||||||||||||||||||||||||||||||||||||||||||||||||||||||||||||||||||||||||||||||||||||||
EIVLTQSPGTLSLSPGERATLSCRASQSVSSSYLAWYQQKPGQAPRLLIYGASSRATGIPDRFSGSGSGTDFTLTISRLEPEDFAVYYCQQYGSSTWTFGQGTKVEIK
^^^^^^^ ^^^ ^^^^^^^^^

Best Alignment of Bezlotoxumab heavy chain CDRs to a sequence from OAS
EVQLVQSGAEVKKSGESLKISCKGSGYSFTSYWIGWVRQMPGKGLEWMGIFYPGDSSTRYSPSFQGQVTISADKSVNTAYLQWSSLKASDTAMYYCARRRNWGNAFDIWGQGTMVTVSS
..............||||||||||||||||||||||||||||||||||||.|||||.||||||||||||||||||..||||||||||||||||||||||||||.|||||||||||||||||
-------------GESLKISCKGSGYSFTSYWIGWVRQMPGKGLEWMGIIYPGDSDTRYSPSFQGQVTISADKSISTAYLQWSSLKASDTAMYYCARRRNYGNAFDIWGQGTMVTVSS
^^^^^^^^ ^^^^^^^^ ^^^^^^^^^^^^

Best Alignment of Bezlotoxumab light CDRs chain to a sequence from OAS
EIVLTQSPGTLSLSPGERATLSCRASQSVSSSYLAWYQQKPGQAPRLLIYGASSRATGIPDRFSGSGSGTDFTLTISRLEPEDFAVYYCQQYGSSTWTFGQGTKVEIK
||||||||||||||||||||||||||||||||||||||||||||||||||||||||||||||||||||||||||||||||||||||||||||||||||||||||||||
EIVLTQSPGTLSLSPGERATLSCRASQSVSSSYLAWYQQKPGQAPRLLIYGASSRATGIPDRFSGSGSGTDFTLTISRLEPEDFAVYYCQQYGSSTWTFGQGTKVEIK
^^^^^^^ ^^^ ^^^^^^^^^

Best Alignment of Bezlotoxumab CDR-H3 chain to a sequence from OAS
EVQLVQSGAEVKKSGESLKISCKGSGYSFTSYWIGWVRQMPGKGLEWMGIFYPGDSSTRYSPSFQGQVTISADKSVNTAYLQWSSLKASDTAMYYCARRRNWGNAFDIWGQGTMVTVSS
..............|.|...|..||.||......|.||.|||||||.|.......||.|.||....||||.|.|.|...|..||..|.|||.||||||||||.|||||||||||||||
-------------SETLSLTCAVSGGSFSGFYWSWIRQPPGKGLEWIGEINHS-GSTNYNPSLKSRVTISVDTSKNQFSLKLSSVTAADTAVYYCARRRNWGRAFDIWGQGTMVTVSS
^^^^^^^^ ^^^^^^^^ ^^^^^^^^^^^^

Therapeutic : Bimagrumab

Best Alignment of Bimagrumab heavy chain to a sequence from OAS
QVQLVQSGAEVKKPGASVKVSCKASGYTFTSSYINWVRQAPGQGLEWMGTINPVSGSTSYAQKFQGRVTMTRDTSISTAYMELSRLRSDDTAVYYCARGG----------WFDYWGQGTLVTVSS
||||||||||||||||||||||||||||||..|||||||||||||||||.|||.||.|.||||||||||||||||||||||||||||||||||||||||||............||||||||||||
QVQLVQSGAEVKKPGASVKVSCKASGYTFTSYDINWVRQAPGQGLEWMGWINPNSGGTNYAQKFQGRVTMTRDTSISTAYMELSRLRSDDTAVYYCARGGEYYDFWSGYYPTDYWGQGTLVTVSS
^^^^^^^^ ^^^^^^^^ ^^^^^^^^^^^^^^^^^^

Best Alignment of Bimagrumab light chain to a sequence from OAS
QSALTQPASVSGSPGQSITISCTGTSSDVGSYNYVNWYQQHPGKAPKLMIYGVSKRPSGVSNRFSGSKSGNTASLTISGLQAEDEADYYCGTFAGGSYYGVFGGGTKLTVL
||||||||||||||||||||||||||||||||.||||.|||||||||||||||||.||||||||||||||||||||||||||||||||||||||||||||...||.|.|||||||||||||
QSALTQPASVSGSPGQSITISCTGTSSDVGGYNYVSWYQQHPGKAPKLMIYDVSKRPSGVSNRFSGSKSGNTASLTISGLQAEDEADYYCCSYAGSSTYGVFGGGTKLTVL
^^^^^^^^^ ^^^ ^^^^^^^^^^^

Best Alignment of Bimagrumab heavy chain CDRs to a sequence from OAS
QVQLVQSGAEVKKPGASVKVSCKASGYTFTSSYINWVRQAPGQGLEWMGTINPVSGSTSYAQKFQGRVTMTRDTSISTAYMELSRLRSDDTAVYYCARGGWFDYWGQGTLVTVSS
..............|||||||||||||||||.|..||||||||||||||.|||.||||||||||||||||||||||.||.|||||.|||.||||||||||||||||.|||||||||||
-------------GASVKVSCKASGYTFTSYYMHWVRQAPGQGLEWMGIINPSSGSTSYAQKFQGRVTMTRDTSTSTVYMELSSLRSEDTAVYYCARGGWFDPWGQGTLVTVSS
^^^^^^^^ ^^^^^^^^ ^^^^^^^^

Best Alignment of Bimagrumab light CDRs chain to a sequence from OAS
QSALTQPASVSGSPGQSITISCTGTSSDVGSYNYVNWYQQHPGKAPKLMIYGVSKRPSGVSNRFSGSKSGNTASLTISGLQAEDEADYYCGTFAGGSYYGVFGGGTKLTVL
||||||||||||.||.|||..|||||||||||||...||||||||||.||||||||||||||.|||||||||||||.||.|||||||||||||..||.|.|.||||||.|||||
QSALTQPASVSGAPGPSITNPCTGTSSDVGSYNLFSWYQQHPGKAPNLMIYGVSKRPSGVANRFSGSKSGNTASVTIAGLQAEDEADYYCGSEAGISTYWVFGGGIKLTVL
^^^^^^^^^ ^^^ ^^^^^^^^^^^

Best Alignment of Bimagrumab CDR-H3 chain to a sequence from OAS
QVQLVQSGAEVKKPGASVKVSCKASGYTFTSSYINWVRQAPGQGLEWMGTINPVSGSTSYAQKFQGRVTMTRDTSISTAYMELSRLRSDDTAVYYCARGGWFDYWGQGTLVTVSS
.|.||.|......||.|.|.||.|||.||...|..||||.|..||||...||....||.|......|....||......|...|.|.|.|||.||||||||||||||||..||||
EVKLVESEGGLVQPGSSMKLSCTASGFTFSDYYMAWVRQVPEKGLEWVANINYDGSSTYYLDSLKSRFIISRDNAKNILYLQMSSLKSEDTATYYCARGGWFDYWGQGTTLTVSS
^^^^^^^^ ^^^^^^^^ ^^^^^^^^

Therapeutic : Bimekizumab

Best Alignment of Bimekizumab heavy chain to a sequence from OAS
EVQLVESGGGLVQPGGSLRLSCAASGFTFSDYNMAWVRQAPGKGLEWVATITYEGRNTYYRDSVKGRFTISRDNAKNSLYLQMNSLRAEDTAVYYCASPPQYYEGSIYRLWFAHWGQGTLVTVSS
|||||||||||||||.||.||||||||||||||||||||||.||||||||||.|.|..|||||||||||||||||||..|||||||||.||||.|||||.|.||.||.|..|||.|||||||||||
EVQLVESGGGLVQPGRSLKLSCAASGFTFSDYNMAWVRQAPKKGLEWVATISYDGSSTYYRDSVKGRFTISRDNAKSTLYLQMNSLRSEDTATYYCASSPYYYDGSYYYGWFAYWGQGTLVTVSS
^^^^^^^^ ^^^^^^^^ ^^^^^^^^^^^^^^^^^

Best Alignment of Bimekizumab light chain to a sequence from OAS
AIQLTQSPSSLSASVGDRVTITCRADESVRTLMHWYQQKPGKAPKLLIYLVSNSEIGVPDRFSGSGSGTDFRLTISSLQPEDFATYYCQQTWSDPWTFGQGTKVEIK
|||||||||||||||||||||||||.........|||||||||||||||..||.|.|||.|||||||||||.||||||||||||||||||..|||||||||||||||
AIQLTQSPSSLSASVGDRVTITCRASQGISSALAWYQQKPGKAPKLLIYDASNLESGVPSRFSGSGSGTDFTLTISSLQPEDFATYYCQQFNSDPWTFGQGTKVEIK
^^^^^^ ^^^ ^^^^^^^^^

Best Alignment of Bimekizumab heavy chain CDRs to a sequence from OAS
EVQLVESGGGLVQPGGSLRLSCAASGFTFSDYNMAWVRQAPGKGLEWVATITYEGRNTYYRDSVKGRFTISRDNAKNSLYLQMNSLRAEDTAVYYCASPPQYYEGSIYRLWFAHWGQGTLVTVSS
|||||||||||||||.||.|||||||||||||.||||||||.|||||||.|.|||..|||.|||||||||||||||..|||||||||.||||.||||.||.||.||.|..|||.|||||||||||
EVQLVESGGGLVQPGRSLKLSCAASGFTFSDYYMAWVRQAPKKGLEWVASISYEGSSTYYGDSVKGRFTISRDNAKSTLYLQMNSLRSEDTATYYCARPPYYYDGSYYANWFAYWGQGTLVTVSS
^^^^^^^^ ^^^^^^^^ ^^^^^^^^^^^^^^^^^

Best Alignment of Bimekizumab light CDRs chain to a sequence from OAS
AIQLTQSPSSLSASVGDRVTITCRADESVRTLMHWYQQKPGKAPKLLIYLVSNSEIGVPDRFSGSGSGTDFRLTISSLQPEDFATYYCQQTWSDPWTFGQGTKVEIK
.|.|||||...|...|..|||||||..||.|..|||||.|...|||||..||.|..|||.|||||||||...|||.||..||.||||||||.|.|||||||||||||
EIALTQSPDFQSVTPGEKVTITCRASQSVGTSLHWYQQQPEQSPKLLIKYVSQSFSGVPSRFSGSGSGTTYTLTIKSLEAEDGATYYCQQTSSLPWTFGQGTKVEIK
^^^^^^ ^^^ ^^^^^^^^^

Best Alignment of Bimekizumab CDR-H3 chain to a sequence from OAS
EVQLVESGGGLVQPGGSLRLSCAASGFTFSDYNMAWVRQAPGKGLEWVATITYEGRNTYYRDSVKGRFTISRDNAKNSLYLQMNSLRAEDTAVYYCASPPQYYEGSIYRLWFAHWGQGTLVTVSS
.|||...|..||.||.|..|||.|||.||..|.|.||.|.||.||||...|.............|...|..........|.|..||..||.|||||||||.||.||.|..|||.||||||||||.
QVQLQQPGAELVKPGASVKLSCKASGYTFTSYWMHWVKQRPGQGLEWIGMIHPNSGRINHNEKFKSKATLTVVKSSSTAYMQLSSLTSEDSAVYYCASPPIYYYGSSYLSWFAYWGQGTLVTVSA
^^^^^^^^ ^^^^^^^^ ^^^^^^^^^^^^^^^^^

Therapeutic : Bleselumab

Best Alignment of Bleselumab heavy chain to a sequence from OAS
QLQLQESGPGLLKPSETLSLTCTVSGGSISSPGYYGGWIRQPPGKGLEWIGSIYKSGSTYHNPSLKSRVTISVDTSKNQFSLKLSSVTAADTAVYYCTRPVVRY-----FGWFDPWGQGTLVTVSS
|||||||||||.||||||||||||||||||||..||.||||||||||||||||.|||||.||||||||||||||||||||||||||||||||||||||.|||.||......||||||||||||||||
QLQLQESGPGLVKPSETLSLTCTVSGGSISSSSYYWGWIRQPPGKGLEWIGSIYYSGSTYYNPSLKSRVTISVDTSKNQFSLKLSSVTAADTAVYYCARPVIRYSYGSLRGWFDPWGQGTLVTVSS
^^^^^^^^^^ ^^^^^^^ ^^^^^^^^^^^^^^^^^

Best Alignment of Bleselumab light chain to a sequence from OAS
AIQLTQSPSSLSASVGDRVTITCRASQGISSALAWYQQKPGKAPKLLIYDASNLESGVPSRFSGSGSGTDFTLTISSLQPEDFATYYCQQFNSYPTFGQGTKVEIK
||||||||||||||||||||||||||||||||||||||||||||||||||||.|||||||||||||||||||||||||||||||||||||||.|||||||||||||
AIQLTQSPSSLSASVGDRVTITCRASQGISSALAWYQQKPGKAPKLLIYDASSLESGVPSRFSGSGSGTDFTLTISSLQPEDFATYYCQQFNNYPTFGQGTKVEIK
^^^^^^ ^^^ ^^^^^^^^

Best Alignment of Bleselumab heavy chain CDRs to a sequence from OAS
QLQLQESGPGLLKPSETLSLTCTVSGGSISSPGYYGGWIRQPPGKGLEWIGSIYKSGSTYHNPSLKSRVTISVDTSKNQFSLKLSSVTAADTAVYYCTRPVVRYFGWFDPWGQGTLVTVSS
..........|.|||.|||||||||||||||.|||..||||.|||||||||.||.|||||.||||||.||||||||||||||||||||||||||||||||.||||...||||||||||||||||
---------GLVKPSQTLSLTCTVSGGSISSGGYYWSWIRQHPGKGLEWIGYIYYSGSTYYNPSLKSLVTISVDTSKNQFSLKLSSVTAADTAVYYCARPVVTPSGWFDPWGQGTLVTVSS
^^^^^^^^^^ ^^^^^^^ ^^^^^^^^^^^^^

Best Alignment of Bleselumab light CDRs chain to a sequence from OAS
AIQLTQSPSSLSASVGDRVTITCRASQGISSALAWYQQKPGKAPKLLIYDASNLESGVPSRFSGSGSGTDFTLTISSLQPEDFATYYCQQFNSYPTFGQGTKVEIK
|||||||||||||||||||||||||||||||||.||||||||||||||||||.||||||||||||||||||||||||||||||||||||||||||||||||..|||
AIQLTQSPSSLSASVGDRVTITCRASQGISSALVWYQQKPGKAPKLLIYDASSLESGVPSRFSGSGSGTDFTLTISSLQPEDFATYYCQQFNSYPTFGQGTRLEIK
^^^^^^ ^^^ ^^^^^^^^

Best Alignment of Bleselumab CDR-H3 chain to a sequence from OAS
QLQLQESGPGLLKPSETLSLTCTVSGGSISSPGYYGGWIRQPPGKGLEWIGSIYKSGSTYHNPSLKSRVTISVDTSKNQFSLKLSSVTAADTAVYYCTRPVVRYFGWFDPWGQGTLVTVSS
..............|||||||||||||||....||..||||||||||||||.||.||||..|||||||||||||||||||||||||||||||||.||.||||||.||||||||||||||||
--------------SETLSLTCTVSGGSI--SSYYWSWIRQPPGKGLEWIGYIYYSGSTNYNPSLKSRVTISVDTSKNQFSLKLSSVTAADTAVHYCARPVVRYMGWFDPWGQGTLVTVSS
^^^^^^^^^^ ^^^^^^^ ^^^^^^^^^^^^^

Therapeutic : Blosozumab

Best Alignment of Blosozumab heavy chain to a sequence from OAS
QVQLVQSGAEVKKPGASVKVSCKVSGFPIKDTFQHWVRQAPGKGLEWMGWSDPEIGDTEYASKFQGRVTMTEDTSTDTAYMELSSLRSEDTAVYYCATGDTT------------YKFDFWGQGTTVTVSS
||||||||||||||||||||||||||........|||||||||||||||..|||.|.|.||.||||||||||||||||||||||||||||||||||||||..............|..|||||||||||||
QVQLVQSGAEVKKPGASVKVSCKVSGYTLTELSMHWVRQAPGKGLEWMGGFDPEDGETIYAQKFQGRVTMTEDTSTDTAYMELSSLRSEDTAVYYCATGDPLAVAVAPTNYYYYYGMDFWGQGTTVTVSS
^^^^^^^^ ^^^^^^^^ ^^^^^^^^^^^^^^^^^^^^^^

Best Alignment of Blosozumab light chain to a sequence from OAS
DIQMTQSPSSLSASVGDRVTITCKASQDVHTAVAWYQQKPGKAPKLLIYWASTRWTGVPSRFSGSGSGTDFTLTISSLQPEDFATYYCQQYSDYPWTFGGGTKVEIK
|||||||||.|||||||||||||.|||.|.|..||||||||||||||||.|||..||||||||||||||||||||||||||||||||||||..||.|||||||||||
DIQMTQSPSTLSASVGDRVTITCRASQTVNTWLAWYQQKPGKAPKLLIYKASTLETGVPSRFSGSGSGTDFTLTISSLQPEDFATYYCQQYNSYPLTFGGGTKVEIK
^^^^^^ ^^^ ^^^^^^^^^

Best Alignment of Blosozumab heavy chain CDRs to a sequence from OAS
QVQLVQSGAEVKKPGASVKVSCKVSGFPIKDTFQHWVRQAPGKGLEWMGWSDPEIGDTEYASKFQGRVTMTEDTSTDTAYMELSSLRSEDTAVYYCATGDTTYKFDFWGQGTTVTVSS
.|||.||||||...||||||.||..|||.|||...|||.|....||||.|..|||.|||.||.||||..|.|.||...||...||||.|||||||||.||||..||.||.||||||||
EVQLQQSGAELVTPGASVKLSCTASGFNIKDYYMHWVKQRTEQGLEWIGRIDPEDGDTKYAPKFQGKATLTADTASNTADLQLSSLTSEDTAVYYCARGDTTWYFDVWGTGTTVTVSS
^^^^^^^^ ^^^^^^^^ ^^^^^^^^^^^

Best Alignment of Blosozumab light CDRs chain to a sequence from OAS
DIQMTQSPSSLSASVGDRVTITCKASQDVHTAVAWYQQKPGKAPKLLIYWASTRWTGVPSRFSGSGSGTDFTLTISSLQPEDFATYYCQQYSDYPWTFGGGTKVEIK
||.||||....|.||||||.|||||||||.|||||||||||..|||||||||||..||||.||.|||||||||||||..|.||.|.|.|||||.||||||||||.|||
DIVMTQSHKFMSTSVGDRVSITCKASQDVGTAVAWYQQKPGQSPKLLIYWASTLHTGVPDRFTGSGSGTDFTLTISNVQSEDLADYFCQQYSSYPWTFGGGTKLEIK
^^^^^^ ^^^ ^^^^^^^^^

Best Alignment of Blosozumab CDR-H3 chain to a sequence from OAS
QVQLVQSGAEVKKPGASVKVSCKVSGFPIKDTFQHWVRQAPGKGLEWMGWSDPEIGDTEYASKFQGRVTMTEDTSTDTAYMELSSLRSEDTAVYYCATGDTTYKFDFWGQGTTVTVSS
||||.|||||..|||||||.|||.||.........||.|.|||||||.|...|..|||.|...|.|..|.|.|.|..||||.||||.|||.|||.||.|||||.|||||||||.||||
QVQLQQSGAELVKPGASVKISCKASGYAFSNYWMNWVKQRPGKGLEWIGQIYPGDGDTNYNGNFKGKATLTADKSSSTAYMQLSSLTSEDSAVYFCARGDTTYYFDFWGQGTTLTVSS
^^^^^^^^ ^^^^^^^^ ^^^^^^^^^^^

Therapeutic : Bococizumab

Best Alignment of Bococizumab heavy chain to a sequence from OAS
QVQLVQSGAEVKKPGASVKVSCKASGYTFTSYYMHWVRQAPGQGLEWMGEISPFGGRTNYNEKFKSRVTMTRDTSTSTVYMELSSLRSEDTAVYYCARERPL-------YASDLWGQGTTVTVSS
|||||||||||||||||||||||||||||||||||||||||||||||||.|||.||.|.|..||..||||||||||||||||||||||||||||||||.|.........|..|.|||||||||||
QVQLVQSGAEVKKPGASVKVSCKASGYTFTSYYMHWVRQAPGQGLEWMGIISPSGGSTSYAQKFQGRVTMTRDTSTSTVYMELSSLRSEDTAVYYCARDRRVTSLYYYYYGMDVWGQGTTVTVSS
^^^^^^^^ ^^^^^^^^ ^^^^^^^^^^^^^^^^^

Best Alignment of Bococizumab light chain to a sequence from OAS
DIQMTQSPSSLSASVGDRVTITCRASQGISSALAWYQQKPGKAPKLLIYSASYRTGVPSRFSGSGSGTDFTFTISSLQPEDIATYYCQQRYSLWRTFGQGTKLEIK
|||||||||||||||||||||||||||.|||.|.||||||||||||||||||....||||||||||||||||.|||||||||||||||||..||.||||||||||||
DIQMTQSPSSLSASVGDRVTITCRASQSISSYLNWYQQKPGKAPKLLIYSASSLQSGVPSRFSGSGSGTDFTLTISSLQPEDIATYYCQQCDSLPRTFGQGTKLEIK
           ^^^^^^       ^^^                ^^^^^^^^^

Best Alignment of Bococizumab heavy chain CDRs to a sequence from OAS
QVQLVQSGAEVKKPGASVKVSCKASGYTFTSYYMHWVRQAPGQGLEWMGEISPFGGRTNYNEKFKSRVTMTRDTSTSTVYMELSSLRSEDTAVYYCARERPLYASDLWGQGTTVTVSS
................|||||||||||||||||||||||||||||||||.|.|.||.|.|..||..|||||||||||||||||||||||||||||||||||.||.|.|||||.||..|
----------------SVKVSCKASGYTFTSYYMHWVRQAPGQGLEWMGIINPSGGSTSYAQKFQDRVTMTRDTSTSTVYMELSSLRSEDTAVYYCARERPGYACDIWGQGTMVTAPS
           ^^^^^^^^       ^^^^^^^^                ^^^^^^^^^^^

Best Alignment of Bococizumab light CDRs chain to a sequence from OAS
DIQMTQSPSSLSASVGDRVTITCRASQGISSALAWYQQKPGKAPKLLIYSASYRTGVPSRFSGSGSGTDFTFTISSLQPEDIATYYCQQRYSLWRTFGQGTKLEIK
|||.||||||||||||||||||||.||||||.|.||.|||||.|||||||||....||||||||||||||||.|||||||||.|||||||.|||.||||||||.|||
DIQLTQSPSSLSASVGDRVTITCRVSQGISSYLNWYRQKPGKVPKLLIYSASNLQSGVPSRFSGSGSGTDFTLTISSLQPEDFATYYCQQSYSLPRTFGQGTKVEIK
           ^^^^^^       ^^^                ^^^^^^^^^

Best Alignment of Bococizumab CDR-H3 chain to a sequence from OAS
QVQLVQSGAEVKKPGASVKVSCKASGYTFTSYYMHWVRQAPGQGLEWMGEISPFGGRTNYNEKFKSRVTMTRDTSTSTVYMELSSLRSEDTAVYYCARERPLYASDLWGQGTTVTVSS
.|||.|||.|..|||||||.|||.||||||.|.||||.|..|..|||.|.|.|..|.|.||.|||...|.|.|.|.||.||||.||.|||.|||||||||||||.|.|||||.|||||
EVQLQQSGPELVKPGASVKISCKTSGYTFTEYTMHWVKQSHGKSLEWIGGINPNNGGTSYNQKFKGKATLTVDKSSSTAYMELRSLTSEDSAVYYCARERPLYAMDYWGQGTSVTVSS
           ^^^^^^^^       ^^^^^^^^                ^^^^^^^^^^^

Therapeutic : Brazikumab

Best Alignment of Brazikumab heavy chain to a sequence from OAS
QVQLVESGGGVVQPGRSLRLSCAASGFTFSSYGMHWVRQAPGKGLEWVAVIWYDGSNEYYADSVKGRFTISRDNSKNTLYLQMNSLRAEDTAVYYCARDRGYTSS--WYPDAFDIWGQGTMVTVSS
|||||||||||||||||||||||||||||||||||||||||||||||||||||||||.||||||||||||||||||||||||||||||||||||||||||||.||...||||||||||||||||||
QVQLVESGGGVVQPGRSLRLSCAASGFTFSSYGMHWVRQAPGKGLEWVAVIWYDGSNKYYADSVKGRFTISRDNSKNTLYLQMNSLRAEDTAVYYCARDRGYCSSTSCYPDAFDIWGQGTMVTVSS
           ^^^^^^^^       ^^^^^^^^                ^^^^^^^^^^^^^^^^^^

Best Alignment of Brazikumab light chain to a sequence from OAS
QSVLTQPPSVSGAPGQRVTISCTGSSSNTGAGYDVHWYQQVPGTAPKLLIYGSGNRPSGVPDRFSGSKSGTSASLAITGLQAEDEADYYCQSYDSSLSGWVFGGGTRLTVL
||||||||||||||||||||||||||||||||||||||||||||||||||||..|||||||||||||||||||||||||||||||||||||||||||||.||||||.||||
QSVLTQPPSVSGAPGQRVTISCTGSSSNTGAGYDVHWYQQVPGTAPKLLIYGNSNRPSGVPDRFSGSKSGTSASLAITGLQAEDEADYYCQSYDSSLSGVVFGGGTKLTVL
           ^^^^^^^^^       ^^^               ^^^^^^^^^^^

Best Alignment of Brazikumab heavy chain CDRs to a sequence from OAS
QVQLVESGGGVVQPGRSLRLSCAASGFTFSSYGMHWVRQAPGKGLEWVAVIWYDGSNEYYADSVKGRFTISRDNSKNTLYLQMNSLRAEDTAVYYCARDRGYTSSWYPDAFDIWGQGTMVTVSS
................|||||||||||||||||||||||||||||||||.|||||||..|||||||||||||||||||||||||||||||||.||.||||||.||||.||||||||||||||||
---------------GSLRLSCAASGFTFSSYGMHWVRQAPGKGLEWVAFIWYDGSNKFYADSVKGRFTISRDNSKNTLYLQMNSLRAEDTAMYYSARDRGYSSSWYDDAFDIWGQGTMVTVSS
           ^^^^^^^^       ^^^^^^^^                ^^^^^^^^^^^^^^^^

Best Alignment of Brazikumab light CDRs chain to a sequence from OAS
QSVLTQPPSVSGAPGQRVTISCTGSSSNTGAGYDVHWYQQVPGTAPKLLIYGSGNRPSGVPDRFSGSKSGTSASLAITGLQAEDEADYYCQSYDSSLSGWVFGGGTRLTVL
||||||||||||||||||||||||||||||||||||||||.||||||||||||.|||.||||||||||||||||||||||||||||||||||||||||||||||||.||||
QSVLTQPPSVSGAPGQRVTISCTGSSSNTGAGYDVHWYQQLPGTAPKLLIYGSINRPPGVPDRFSGSKSGTSASLAITGLQAEDEADYYCQSYDSSLSGWVFGGGTKLTVL
           ^^^^^^^^^       ^^^               ^^^^^^^^^^^

Best Alignment of Brazikumab CDR-H3 chain to a sequence from OAS
QVQLVESGGGVVQPGRSLRLSCAASGFTF--SSYGMHWVRQAPGKGLEWVAVIWYDGSNEYYADSVKGRFTISRDNSKNTLYLQMNSLRAEDTAVYYCARDRGYTSSWYPDAFDIWGQGTMVTVSS
.|||.|||.|.|.|...|.|.|..||.......|...|.||.|||||||...|.|.....||..|.|.|.|||.|.|||...|...|..|.|||||||||||||.|||||||||||||||.|||||
-VQLQESGPGLVKPSQTLSLTCTVSGGSISSGDYYWSWIRQPPGKGLEWIGYIYYS-GSTYYNPSLKSRVTISVDTSKNQFSLKLSSVTAADTAVYYCARDRGYSSSWYPDAFDIWGQGTTVTVSS
           ^^^^^^^^^^       ^^^^^^^^                ^^^^^^^^^^^^^^^^

Therapeutic : Brentuximab

Best Alignment of Brentuximab heavy chain to a sequence from OAS
QIQLQQSGPEVVKPGASVKISCKASGYTFTDYYITWVKQKPGQGLEWIGWIYPGSGNTKYNEKFKGKATLTVDTSSSTAFMQLSSLTSEDTAVYFCANYGN-YWFAYWGQGTQVTVSA
||||||||||.|||||||||||||||||||||||.||||||||||||||||||||||||||||||||||||||||||||||.|||||||||||||||||.|||...||||||||.|||||
QIQLQQSGPELVKPGASVKISCKASGYTFTDYYINWVKQKPGQGLEWIGWIYPGSGNTKYNEKFKGKATLTVDTSSSTAYMQLSSLTSEDTAVYFCASYGNYVGFAYWGQGTLVTVSA
           ^^^^^^^^       ^^^^^^^^                ^^^^^^^^^^^

Best Alignment of Brentuximab light chain to a sequence from OAS
DIVLTQSPASLAVSLGQRATISCKASQSVDFDGDSYMNWYQQKPGQPPKVLIYAASNLESGIPARFSGSGSGTDFTLNIHPVEEEDAATYYCQQSNEDPWTFGGGTKLEIK
|||||||||||||||||||||||||||||||||||||||||||||||||.|||||||||||||||||||||||||||||||||||||||||||||||||.|||||||||||
DIVLTQSPASLAVSLGQRATISCKASQSVDFDGDSYMNWYQQKPGQPPKLLIYAASNLESGIPARFSGSGSGTDFTLNIHPVEEEDAATYYCQQSNEDPRTFGGGTKLEIK
           ^^^^^^^^^^       ^^^                ^^^^^^^^^

Best Alignment of Brentuximab heavy chain CDRs to a sequence from OAS
QIQLQQSGPEVVKPGASVKISCKASGYTFTDYYITWVKQKPGQGLEWIGWIYPGSGNTKYNEKFKGKATLTVDTSSSTAFMQLSSLTSEDTAVYFCANYGNYWFAYWGQGTQVTVSA
|.||||||.|...||||||.||||||||||||||.||||..||||||||.||||||||.||||||||||||.|.|||||.||||||||||.||||||.|||||||||||||.|||||
QVQLQQSGAELARPGASVKLSCKASGYTFTDYYINWVKQRTGQGLEWIGEIYPGSGNTYYNEKFKGKATLTADKSSSTAYMQLSSLTSEDSAVYFCARYGNYWFAYWGQGTLVTVSA
           ^^^^^^^^       ^^^^^^^^                ^^^^^^^^^^

Best Alignment of Brentuximab light CDRs chain to a sequence from OAS
DIVLTQSPASLAVSLGQRATISCKASQSVDFDGDSYMNWYQQKPGQPPKVLIYAASNLESGIPARFSGSGSGTDFTLNIHPVEEEDAATYYCQQSNEDPWTFGGGTKLEIK
|||.||.|||||||||||||||||||||||||||||||||||||||||||.|||||||||||||||||||||||||||||||||||||||||||||||||||||||||||||
DIVITQTPASLAVSLGQRATISCKASQSVDFDGDSYMNWYQQKPGQPPKLLIYAASNLESGIPARFSGSGSGTDFTLNIHPVEEEDAATYYCQQSNEDPWTFGGGTKLEIK
           ^^^^^^^^^^       ^^^                ^^^^^^^^^

Best Alignment of Brentuximab CDR-H3 chain to a sequence from OAS
QIQLQQSGPEVVKPGASVKISCKASGYTFTDYYITWVKQKPGQGLEWIGWIYPGSGNTKYNEKFKGKATLTVDTSSSTAFMQLSSLTSEDTAVYFCANYGNYWFAYWGQGTQVTVSA
..||||||||.||||||||||||.||||||.|...||||..|..|||||.|.|..|.|.||.|||||||||||.|||||.|.|.||||||.|||.||||||||||||||||.|||||
EVQLQQSGPELVKPGASVKISCKTSGYTFTEYTMHWVKQSHGKSLEWIGGINPNNGGTSYNQKFKGKATLTVDKSSSTAYMELRSLTSEDSAVYYCANYGNYWFAYWGQGTLVTVSA
           ^^^^^^^^       ^^^^^^^^                ^^^^^^^^^^

Therapeutic : Briakinumab

Best Alignment of Briakinumab heavy chain to a sequence from OAS
QVQLVESGGGVVQPGRSLRLSCAASGFTFSSYGMHWVRQAPGKGLEWVAFIRYDGSNKYYADSVKGRFTISRDNSKNTLYLQMNSLRAEDTAVYYCKTHG---------SHDNWGQGTMVTVSS
||||||||||||||||||||||||||||||||||||||||||||||||||||||||||||||||||||||||||||||||||||||||||||||||.||............|.|||||||||||
QVQLVESGGGVVQPGRSLRLSCAASGFTFSSYGMHWVRQAPGKGLEWVAFIRYDGSNKYYADSVKGRFTISRDNSKNTLYLQMNSLRAEDTAVYYCATHPRYSSGWYWDAFDIWGQGTMVTVSS
           ^^^^^^^^       ^^^^^^^^                ^^^^^^^^^^^^^^^^

Best Alignment of Briakinumab light chain to a sequence from OAS
QSVLTQPPSVSGAPGQRVTISCSGSRSNIGSNTVKWYQQLPGTAPKLLIYYNDQRPSGVPDRFSGSKSGTSASLAITGLQAEDEADYYCQSYDRYTHPALLFGTGTKVTVL
||||||||.||.||||||||||||.||||||||.|||||||||||||||.|.||||||||||||||||||||||||||||.|||||||||||.....||.||||||||||
QSVLTQPPSASGTPGQRVTISCSGSSSNIGSNTVNWYQQLPGTAPKLLIYRNNQRPSGVPDRFSGSKSGTSASLAITGLQADDEADYYCQSYDGS-LSALVFGTGTKVTVL
           ^^^^^^^^       ^^^               ^^^^^^^^^^^^
Best Alignment of Briakinumab heavy chain CDRs to a sequence from OAS
QVQLVESGGGVVQPGRSLRLSCAASGFTFSSYGMHWVRQAPGKGLEWVAFIRYDGSNKYYADSVKGRFTISRDNSKNTLYLQMNSLRAEDTAVYYCKTHGSHDNWGQGTMVTVSS
................||||||||||||||||||||||||||||||||||||||||||||||||||||||||||||||||||||||||||||||||.|.||||.|||||.|||||
----------------SLRLSCAASGFTFSSYGMHWVRQAPGKGLEWVAFIRYDGSNKYYADSVKGRFTISRDNSKNTLYLQMNSLRAEDTAVYYCATSGSHDYWGQGTLVTVSS
           ^^^^^^^^       ^^^^^^^^                ^^^^^^^^

Best Alignment of Briakinumab light CDRs chain to a sequence from OAS
QSVLTQPPSVSGAPGQRVTISCSGSRSNIGSNTVKWYQQLPGTAPKLLIYYNDQRPSGVPDRFSGSKSGTSASLAITGLQAEDEADYYCQSYDRYTHPALLFGTGTKVTVL
|||||||||.||.|||.|||||||||||||||||.|||||||||||||||.|||||||||||||||||||||||||||||||||||||||||||.......||.|||.|||
QSVLTQPPSASGTPGQSVTISCSGSRSNIGSNTVTWYQQLPGTAPKLLIYTNDQRPSGVPDRFSGSKSGTSASLAITGLQAEDEADYYCQSYDRRLSASVVFGGGTKLTVL
                    ^^^^^^^^                ^^^                                        ^^^^^^^^^^^^

Best Alignment of Briakinumab CDR-H3 chain to a sequence from OAS
QVQLVESGGGVVQPGRSLRLSCAASGFTFSSYGMHWVRQAPGKGLEWVAFIRYDGSNKYYADSVKGRFTISRDNSKNTLYLQMNSLRAEDTAVYYCKTHGSHDNWGQGTMVTVSS
.|||..||...|.||.|..|||.||||......||||.|.|..||||...|........||....|..||..|.|.||.|||..||..||||||||.||||.||||||...||.|
-VQLQQSGAELVRPGASVKLSCTASGFNIKDDYMHWVKQRPEQGLEWIGWIDPENGDTEYASKFQGKATITADTSSNTAYLQLSSLTSEDTAVYYCTTHGSPDNWGQGDTLTVCS
                    ^^^^^^^^                ^^^^^^^^                                        ^^^^^^^^

Therapeutic : Brodalumab

Best Alignment of Brodalumab heavy chain to a sequence from OAS
QVQLVQSGAEVKKPGASVKVSCKASGYTFTRYGISWVRQAPGQGLEWMGWISTYSGNTNYAQKLQGRVTMTTDTSTSTAYMELRSLRSDDTAVYYCARRQL---YFDYWGQGTLVTVSS
||||||||||||||||||||||||||||||.|||||||||||||||||||||.|||||||||||||||||||||||||||||||||||||||||||||.||...|||||||||||||||
QVQLVQSGAEVKKPGASVKVSCKASGYTFTSYGISWVRQAPGQGLEWMGWISAYSGNTNYAQKLQGRVTMTTDTSTSTAYMELRSLRSDDTAVYYCARTQLLKAYFDYWGQGTLVTVSS
                    ^^^^^^^^                ^^^^^^^^                                        ^^^^^^^^^^^^

Best Alignment of Brodalumab light chain to a sequence from OAS
EIVMTQSPATLSVSPGERATLSCRASQSVSSNLAWFQQKPGQAPRPLIYDASTRATGVPARFSGSGSGTDFTLTISSLQSEDFAVYYCQQYDNWPLTFGGGTKVEIK
|||||||||||||||||||||||||||||||||.|.|||||||||.|||||||||||||||||||||||||||||||||||||||||||||||.|||||||||||||||
EIVMTQSPATLSVSPGERATLSCRASQSVSSNLTWYQQKPGQAPRLLIYDASTRATGVPARFSGSGSGTDFTLTISSLQSEDFAVYYCQQYNNWPLTFGGGTKVEIK
                     ^^^^^^              ^^^                                 ^^^^^^^^^

Best Alignment of Brodalumab heavy chain CDRs to a sequence from OAS
QVQLVQSGAEVKKPGASVKVSCKASGYTFTRYGISWVRQAPGQGLEWMGWISTYSGNTNYAQKLQGRVTMTTDTSTSTAYMELRSLRSDDTAVYYCARRQLYFDYWGQGTLVTVSS
...............|||||||||||||||.|||||||||||||||||||||||.|||||||||||.|||||||.|||||||||||.||||||||||||.||||||||||||||||
---------------ASVKVSCKASGYTFTNYGISWVRQAPGQGLEWMGWISTYTGNTNYAQKLQGSVTMTTDTATSTAYMELRSLTSDDTAVYYCARRALYFDYWGQGTLVTVSS
                    ^^^^^^^^                ^^^^^^^^                                        ^^^^^^^^^

Best Alignment of Brodalumab light CDRs chain to a sequence from OAS
EIVMTQSPATLSVSPGERATLSCRASQSVSSNLAWFQQKPGQAPRPLIYDASTRATGVPARFSGSGSGTDFTLTISSLQSEDFAVYYCQQYDNWPLTFGGGTKVEIK
..||||||||||||.|.||||||||.||||||.||.||||.||||.|||||||||||||||||.|||||.||||||||||||||||||||||||||||||||.||||
--VMTQSPATLSVSLGDRATLSCRAGQSVSSNVAWYQQKPEQAPRILIYDASTRATGVPARFSCSGSGTEFTLTISSLQSEDFAVYYCQQYDNWPLTFGGGTNVEIK
                     ^^^^^^              ^^^                                 ^^^^^^^^^

Best Alignment of Brodalumab CDR-H3 chain to a sequence from OAS
QVQLVQSGAEVKKPGASVKVSCKASGYTFTRYGISWVRQAPGQGLEWMGWISTYSGNTNYAQKLQGRVTMTTDTSTSTAYMELRSLRSDDTAVYYCARRQLYFDYWGQGTLVTVSS
.|||.|||||...||||||.||||||||||.|...||.|.||.||||.|........|||.||..|..|.|.|.|.|||||.|.||.|.|.|||||||||||||||||||..||||
-VQLQQSGAELVMPGASVKLSCKASGYTFTSYWMHWVKQRPGKGLEWIGEFDPSDSYTNYNQKFKGKSTLTVDKSSSTAYMQLSSLTSEDSAVYYCARRQLYFDYWGQGTTLTVSS
                    ^^^^^^^^                ^^^^^^^^                                        ^^^^^^^^^

Therapeutic : Burosumab

Best Alignment of Burosumab heavy chain to a sequence from OAS
QVQLVQSGAEVKKPGASVKVSCKASGYTFTNHYMHWVRQAPGQGLEWMGIINPISGSTSNAQKFQGRVTMTRDTSTSTVYMELSSLRSEDTAVYYCARDIV-------DAFDFWGQGTMVTVSS
||||||||||||||||||||||||||||||..|||||||||||||||||||||..||||.||||||||||||||||||||||||||||||||||||||.||.......||||.|||||||||||
QVQLVQSGAEVKKPGASVKVSCKASGYTFTSYYMHWVRQAPGQGLEWMGIINPSGGSTSYAQKFQGRVTMTRDTSTSTVYMELSSLRSEDTAVYYCARNIVVVPAATGDAFDIWGQGTMVTVSS
                    ^^^^^^^^                ^^^^^^^^                                        ^^^^^^^^^^^^^^^^^

Best Alignment of Burosumab light chain to a sequence from OAS
AIQLTQSPSSLSASVGDRVTITCRASQGISSALVWYQQKPGKAPKLLIYDASSLESGVPSRFSGSGSGTDFTLTISSLQPEDFATYYCQQFN-DYFTFGPGTKVDIK
|||||||||||||||||||||||||||||||||.||||||||||||||||||||||||||||||||||||||||||||||||||||||||||..|||||||||||||
AIQLTQSPSSLSASVGDRVTITCRASQGISSALAWYQQKPGKAPKLLIYDASSLESGVPSRFSGSGSGTDFTLTISSLQPEDFATYYCQQFNNYYFTFGPGTKVDIK
                     ^^^^^^              ^^^                                 ^^^^^^^^^

Best Alignment of Burosumab heavy chain CDRs to a sequence from OAS
QVQLVQSGAEVKKPGASVKVSCKASGYTFTNHYMHWVRQAPGQGLEWMGIINPISGSTSNAQKFQGRVTMTRDTSTSTVYMELSSLRSEDTAVYYCARDIVDAFDFWGQGTMVTVSS
................||||||||||||||..|||||||||||||||||||||.|||||.||||||||||||||||||||||||||||||||||||||||.||||.|||||||||||
----------------SVKVSCKASGYTFTSYYMHWVRQAPGQGLEWMGIINPSSGSTSYAQKFQGRVTMTRDTSTSTVYMELSSLRSEDTAVYYCARDISDAFDIWGQGTMVTVSS
                    ^^^^^^^^                ^^^^^^^^                                        ^^^^^^^^^^

Best Alignment of Burosumab light CDRs chain to a sequence from OAS
AIQLTQSPSSLSASVGDRVTITCRASQGISSALVWYQQKPGKAPKLLIYDASSLESGVPSRFSGSGSGTDFTLTISSLQPEDFATYYCQQFNDYFTFGPGTKVDIK
|||||||||||||||||||||||||||||||||.||||||||||||||||||||||||||||||||||||||||||||||||||||||||||.|||||.||...||
AIQLTQSPSSLSASVGDRVTITCRASQGISSALAWYQQKPGKAPKLLIYDASSLESGVPSRFSGSGSGTDFTLTISSLQPEDFATYYCQQFNSYFTFGQGTRLEIK
                     ^^^^^^              ^^^                                 ^^^^^^^^

Best Alignment of Burosumab CDR-H3 chain to a sequence from OAS
QVQLVQSGAEVKKPGASVKVSCKASGYTFTNHYMHWVRQAPGQGLEWMGIINPISGSTSNAQKFQGRVTMTRDTSTSTVYMELSSLRSEDTAVYYCARDIVDAFDFWGQGTMVTVSS
||||||||.||||||||||||||.|||||..|...||||.|||||||||.|....|.|..||.|||||.||.||||||.||||..|.|.|||||.||||.|||||||||||.|||||
QVQLVQSGTEVKKPGASVKVSCKTSGYTFSGHGISWVRQTPGQGLEWMGWISGSNGKTIYAQRFQGRVSMTTDTSTSTGYMELRNLTSDDTAVYFCARDRVDAFDFWGQGTVVTVSS
                    ^^^^^^^^                ^^^^^^^^                                        ^^^^^^^^^^

Therapeutic : Camrelizumab

Best Alignment of Camrelizumab heavy chain to a sequence from OAS
EVQLVESGGGLVQPGGSLRLSCAASGFTFSSYMMSWVRQAPGKGLEWVATISGGGANTYYPDSVKGRFTISRDNAKNSLYLQMNSLRAEDTAVYYCARQLY--------YFDYWGQGTTVTVSS
||||||||||||||||||||||||||||||||.|||||||||||||||..|||.|..|||.||||||||||||||||||||||||||||||||||||||.|.........||||||||||||||
EVQLVESGGGLVQPGGSLRLSCAASGFTFSSYAMSWVRQAPGKGLEWVSAISGSGGSTYYADSVKGRFTISRDNAKNSLYLQMNSLRAEDTAVYYCARQTYCGGDCYSGIFDYWGQGTTVTVSS
                    ^^^^^^^^                ^^^^^^^^                                        ^^^^^^^^^^^^^^^^^

Best Alignment of Camrelizumab light chain to a sequence from OAS
DIQMTQSPSSLSASVGDRVTITCLASQTIGTWLTWYQQKPGKAPKLLIYTATSLADGVPSRFSGSGSGTDFTLTISSLQPEDFATYYCQQVYSIPWTFGGGTKVEIK
|||||||||||||||||||||||.|||||||.|.|||||||||||||||.|.||..||||||||||||||||||||||||||||||||||.||.|||||.|||||||
DIQMTQSPSSLSASVGDRVTITCRASQTIGTTLNWYQQKPGKAPKLLIYAASSLQSGVPSRFSGSGSGTDFTLTISSLQPEDFATYYCQQSYSTPWTFGQGTKVEIK
                     ^^^^^^              ^^^                                 ^^^^^^^^^

Best Alignment of Camrelizumab heavy chain CDRs to a sequence from OAS
EVQLVESGGGLVQPGGSLRLSCAASGFTFSSYMMSWVRQAPGKGLEWVATISGGGANTYYPDSVKGRFTISRDNAKNSLYLQMNSLRAEDTAVYYCARQLYYFDYWGQGTTVTVSS
||||||||||||.|||||.|||||||||||||.||||||.|.|.|||||||||||.||||||||||||.||||||||.|||||.|||.||||.||||||||||||||.|||.||||
EVQLVESGGGLVKPGGSLKLSCAASGFTFSSYTMSWVRQTPEKRLEWVATISGGGGNTYYPDSVKGRFPISRDNAKNNLYLQMSSLRSEDTALYYCARQLYYFDYWGKGTTLTVSS
                    ^^^^^^^^                ^^^^^^^^                                        ^^^^^^^^^

Best Alignment of Camrelizumab light CDRs chain to a sequence from OAS
DIQMTQSPSSLSASVGDRVTITCLASQTIGTWLTWYQQKPGKAPKLLIYTATSLADGVPSRFSGSGSGTDFTLTISSLQPEDFATYYCQQVYSIPWTFGGGTKVEIK
||||||||.|.|||.|..|||||||||||||||.||||||||.|.||||||||||||||||||||||||.|...|||||.|||..|||||.||.|||||||||.|||
DIQMTQSPASQSASLGESVTITCLASQTIGTWLAWYQQKPGKSPQLLIYTATSLADGVPSRFSGSGSGTKFSFKISSLQAEDFVSYYCQQLYSTPWTFGGGTKLEIK
                     ^^^^^^              ^^^                                 ^^^^^^^^^
Best Alignment of Camrelizumab CDR-H3 chain to a sequence from OAS
EVQLVESGGGLVQPGGSLRLSCAASGFTFSSYMMSWVRQAPGKGLEWVATISGGGANTYYPDSVKGRFTISRDNAKNSLYLQMNSLRAEDTAVYYCARQLYYFDYWGQGTTVTVSS
.|||..||..||.||.|..|||.|||.||..|...||.|.||.||||.|.|..|..||||....||..|..........|.|..||..||.|||.||||||||||||||||.||||
QVQLKQSGAELVRPGASVKLSCKASGYTFTDYYINWVKQRPGQGLEWIARIYPGSGNTYYNEKFKGKATLTAEKSSSTAYMQLSSLTSEDSAVYFCARQLYYFDYWGQGTTLTVSS
                    ^^^^^^^^                ^^^^^^^^                                        ^^^^^^^^^

Therapeutic : Canakinumab

Best Alignment of Canakinumab heavy chain to a sequence from OAS

```
QVQLVESGGGVVQPGRSLRLSCAASGFTFSVYGMNWVRQAPGKGLEWVAIIWYDGDNQYYADSVKGRFTISRDNSKNTLYLQMNGLRAEDTAVYYCARDLRT-----GPFDYWGQGTLVTVSS
||||||||||||||||||||||||||||||.|||.||||||||||||||.|||||.|.||||||||||||||||||||||||||.||||||||||||||||......||||||||||||||||
QVQLVESGGGVVQPGRSLRLSCAASGFTFSSYGMHWVRQAPGKGLEWVAVIWYDGSNKYYADSVKGRFTISRDNSKNTLYLQMNSLRAEDTAVYYCARDLRDHGGNSGPFDYWGQGTLVTVSS
                         ^^^^^^^^                 ^^^^^^^^                                      ^^^^^^^^^^^^^^^^
```

Best Alignment of Canakinumab light chain to a sequence from OAS

```
EIVLTQSPDFQSVTPKEKVTITCRASQSIGSSLHWYQQKPDQSPKLLIKYASQSFSGVPSRFSGSGSGTDFTLTINSLEAEDAAAYYCHQSSSLPFTFGPGTKVDIK
||||||||||||||||||||||||||||||||||||||||||||||||||||||||||||||||||||||||||||||||||||.||||||||||||||||||||||
EIVLTQSPDFQSVTPKEKVTITCRASQSIGSSLHWYQQKPDQSPKLLIKYASQSFSGVPSRFSGSGSGTDFTLTINSLEAEDAATYYCHQSSSLPFTFGPGTKVDIK
                          ^^^^^^                 ^^^                                    ^^^^^^^^^
```

Best Alignment of Canakinumab heavy chain CDRs to a sequence from OAS

```
QVQLVESGGGVVQPGRSLRLSCAASGFTFSVYGMNWVRQAPGKGLEWVAIIWYDGDNQYYADSVKGRFTISRDNSKNTLYLQMNGLRAEDTAVYYCARDLRTGPFDYWGQGTLVTVSS
..............|.||..||||||||||.|||.||||||||||||||.|||||.|.||||||||||||||||||||||||||.||||||||||||||||||.||||||||||||||
--------------GESLKISCAASGFTFSSYGMHWVRQAPGKGLEWVAVIWYDGSNKYYADSVKGRFTISRDNSKNTLYLQMNSLRAEDTAVYYCARDLRTGAFDYWGQGTLVTVSS
                         ^^^^^^^^                 ^^^^^^^^                                      ^^^^^^^^^^^
```

Best Alignment of Canakinumab light CDRs chain to a sequence from OAS

```
EIVLTQSPDFQSVTPKEKVTITCRASQSIGSSLHWYQQKPDQSPKLLIKYASQSFSGVPSRFSGSGSGTDFTLTINSLEAEDAAAYYCHQSSSLPFTFGPGTKVDIK
|.||||||||||||.|||||.|||||||||||.|||.|||||||||||||||||.||||||||||||||||.||||||||||||.||||||||||||||.||...||
EVVLTQSPDFQSVTSKEKVTVTCRASQSIGSSFHWYNQKPDQSPKLLIKYASQSISGVPSRFSGSGSGTDFALTINSLEAEDAATYYCHQSSSLPFTFGQGTRLEIK
                          ^^^^^^                 ^^^                                    ^^^^^^^^^
```

Best Alignment of Canakinumab CDR-H3 chain to a sequence from OAS

```
QVQLVESGGGVVQPGRSLRLSCAASGFTFSVYGMNWVRQAPGKGLEWVAIIWYDGDNQYYADSVKGRFTISRDNSKNTLYLQMNGLRAEDTAVYYCARDLRTGPFDYWGQGTLVTVSS
..............|.||||||||||||||.|.|.||||||||||.||..|..||....|||||||||||||||.|||||||||.|||||||||||||||||||||||||||||||||
--------------GGSLRLSCAASGFTFSSYWMHWVRQAPGKGLVWVSRINTDGSSTSYADSVKGRFTISRDNAKNTLYLQMNSLRAEDTAVYYCARDLRTGPFDYWGQGTLVTVSS
                         ^^^^^^^^                 ^^^^^^^^                                      ^^^^^^^^^^^
```

Therapeutic : Carlumab

Best Alignment of Carlumab heavy chain to a sequence from OAS

```
QVQLVQSGAEVKKPGSSVKVSCKASGGTFSSYGISWVRQAPGQGLEWMGGIIPIFGTANYAQKFQGRVTITADESTSTAYMELSSLRSEDTAVYYCARYDGI-------YGELDFWGQGTLVTVSS
||||||||||||||||||||||||||||||||.|||||||||||||||||||||||||||||||||||||||||||||||||||||||||||||||||..||........||||.|||||||||||
QVQLVQSGAEVKKPGSSVKVSCKASGGTFSSYAISWVRQAPGQGLEWMGGIIPIFGTANYAQKFQGRVTITADESTSTAYMELSSLRSEDTAVYYCARDLGIVVVPAALFGELDYWGQGTLVTVSS
                         ^^^^^^^^                 ^^^^^^^^                                      ^^^^^^^^^^^^^^^^^^
```

Best Alignment of Carlumab light chain to a sequence from OAS

```
EIVLTQSPATLSLSPGERATLSCRASQSVSDAYLAWYQQKPGQAPRLLIYDASSRATGVPARFSGSGSGTDFTLTISSLEPEDFAVYYCHQYIQLHSFTFGQGTKVEIK
|||||||||||||||||||||||||||||.|.|||||||||||||||||||||.|||||||||||||||||||||||||||||||||||.||....|.|||||||||||
EIVLTQSPATLSLSPGERATLSCRASQSV-DSYLAWYQQKPGQAPRLLIYDASYRATGVPARFSGSGSGTDFTLTISSLEPEDFAVYYCQQYG--GSPTFGQGTKVEIK
                          ^^^^^^^                 ^^^                                    ^^^^^^^^^^
```

Best Alignment of Carlumab heavy chain CDRs to a sequence from OAS

```
QVQLVQSGAEVKKPGSSVKVSCKASGGTFSSYGISWVRQAPGQGLEWMGGIIPIFGTANYAQKFQGRVTITADESTSTAYMELSSLRSEDTAVYYCARYDGIYGELDFWGQGTLVTVSS
||||||||||||||||||||||||||||||||.|||||||||||||||||||||||||||||||||||||||||||||||||||||||||||||||||.||.|.|||.|||||||||||
QVQLVQSGAEVKKPGSSVKVSCKASGGTFSSYAISWVRQAPGQGLEWMGGIIPIFGTANYAQKFQGRVTITADESTSTAYMELSSLRSEDTAVYYCAREDGGYYELDYWGQGTLVTVSS
                         ^^^^^^^^                 ^^^^^^^^                                      ^^^^^^^^^^^^
```

Best Alignment of Carlumab light CDRs chain to a sequence from OAS

```
EIVLTQSPATLSLSPGERATLSCRASQSVSDAYLAWYQQKPGQAPRLLIYDASSRATGVPARFSGSGSGTDFTLTISSLEPEDFAVYYCHQYIQLHSFTFGQGTKVEIK
.|||||||||||.||||||||||.||||||..|||.||.|||||||||.||||.|..|...||.||||||..|||.|..||.||||||||||....|||.||||.||||
-IVLTQSPATLSVSPGERATLSCGASQSVSQNYLARYQKKPGQAPRLLMYDASRRDIGIADRFRGSGSGTNVTLTTSRVEPKDFAVYYCHQYGSPPSFTIGQGTQVEIK
                          ^^^^^^^                 ^^^                                    ^^^^^^^^^^
```

Best Alignment of Carlumab CDR-H3 chain to a sequence from OAS

```
QVQLVQSGAEVKKPGSSVKVSCKASGGTFSSYGISWVRQAPGQGLEWMGGIIPIFGTANYAQKFQGRVTITADESTSTAYMELSSLRSEDTAVYYCARYDGIYGELDFWGQGTLVTVSS
||||.|||||..|||.|||.||||.|.||..|.|.||.|.||.||||.|.|.|..|..||..||.|..|.|||.|..||||.||||..||.|.||||||||.|||||.|||||..||||
QVQLQQSGAELMKPGASVKLSCKATGYTFTGYWIEWVKQRPGHGLEWIGEILPGSGSTNYNEKFKGKATFTADTSSNTAYMQLSSLTTEDSAIYYCARYDGYYGELDYWGQGTTLTVSS
                         ^^^^^^^^                 ^^^^^^^^                                      ^^^^^^^^^^^^
```

Therapeutic : Carotuximab

Best Alignment of Carotuximab heavy chain to a sequence from OAS

```
EVKLEESGGGLVQPGGSMKLSCAASGFTFSDAWMDWVRQSPEKGLEWVAEIRSKASNHATYYAESVKGRFTISRDDSKSSVYLQMNSLRAEDTGIYYCTRWRR--FFDSWGQGTTLTVSS
|||||||||||||||||||||||||||||||||||||||||||||||||||||||.||||||||||||||||||||||||||||||||||||||||||||.|...|||.|||||||||||
EVKLEESGGGLVQPGGSMKLSCAASGFTFSDAWMDWVRQSPEKGLEWVAEIRSKANNHATYYAESVKGRFTISRDDSKSSVYLQMNSLRAEDTGIYYCTRRRLGGFFDYWGQGTTLTVSS
                         ^^^^^^^^                 ^^^^^^^^^^                                      ^^^^^^^^^^^
```

Best Alignment of Carotuximab light chain to a sequence from OAS

```
QIVLSQSPAILSASPGEKVTMTCRASSSVSYMHWYQQKPGSSPKPWIYATSNLASGVPVRFSGSGSGTSYSLTISRVEAEDAATYYCQQWSSNPLTFGAGTKLELK
||||||||||||||||||||||||||||||||||||||||||||||||||||||||||||||||||||||||||||||||||||||||||||||||||||||||||
QIVLSQSPAILSASPGEKVTMTCRASSSVSYMHWYQQKPGSSPKPWIYATSNLASGVPVRFSGSGSGTSYSLTISRVEAEDAATYYCQQWSSNPLTFGAGTKLELK
                          ^^^^^                 ^^^                                    ^^^^^^^^^
```

Best Alignment of Carotuximab heavy chain CDRs to a sequence from OAS

```
EVKLEESGGGLVQPGGSMKLSCAASGFTFSDAWMDWVRQSPEKGLEWVAEIRSKASNHATYYAESVKGRFTISRDDSKSSVYLQMNSLRAEDTGIYYCTRWRRFFDSWGQGTTLTVSS
||.|.||||||||||||||||||||||||||||||||||||||||||||||||||.||||||||||||||||||||||||||||||||||||||||||||.||.||.|||||||||||
EVMLVESGGGLVQPGGSMKLSCAASGFTFSDAWMDWVRQSPEKGLEWVAEIRSKANNHATYYAESVKGRFTISRDDSKSSVYLQMNSLRAEDTGIYYCTRGRRYFDYWGQGTTLTVSS
                         ^^^^^^^^                 ^^^^^^^^^^                                      ^^^^^^^^^
```

Best Alignment of Carotuximab light CDRs chain to a sequence from OAS

```
QIVLSQSPAILSASPGEKVTMTCRASSSVSYMHWYQQKPGSSPKPWIYATSNLASGVPVRFSGSGSGTSYSLTISRVEAEDAATYYCQQWSSNPLTFGAGTKLELK
.|||.|||...|.||||||||||.||||||||.||||||||||.|.||||||||||||.|||||||||||||||||.|||||||||||||||||||||||||||.|
HIVLTQSPELMSVSPGEKVTMTCSASSSVSYMYWYQQKPGSSPRPLIYATSNLASGVPARFSGSGSGTSYSLTISRMEAEDAATYYCQQWSSNPLTFGAGTKLEIK
                          ^^^^^                 ^^^                                    ^^^^^^^^^
```

Best Alignment of Carotuximab CDR-H3 chain to a sequence from OAS

```
EVKLEESGGGLVQPGGSMKLSCAASGFTF--SDAWMDWVRQSPEKGLEWVAEIRSKASNHATYYAESVKGRFTISRDDSKSSVYLQMNSLRAEDTGIYYCTRWRRFFDSWGQGTTLTVSS
...................|.|...|...........|.||||..||||......|..|....||.|||.|...|.|.|.....|..||....|...|||||.||.||||||||..||||
--------------SQTLSLTCVIYGDSVSNKNSAWNWIRQSPSRGLEWLGRTYYKS-NWYNEYAVSVKSRISFSADSSENQFSLHLNSVTPDDSAVYYCTRGRREFDSWGQGTLVTVSS
                         ^^^^^^^^^^                 ^^^^^^^^^^                                      ^^^^^^^^^
```

Therapeutic : Certolizumab

Best Alignment of Certolizumab heavy chain to a sequence from OAS

```
EVQLVESGGGLVQPGGSLRLSCAASGYVFTDYGMNWVRQAPGKGLEWMGWINTYIGEPIYADSVKGRFTFSLDTSKSTAYLQMNSLRAEDTAVYYCARGYRS---YAMDYWGQGTLVTVSS
||||||||||||||||||||||||||..|.||||.||||||||||||...||...|...||||||||||.|.|.||.|.|||||||||||||||||||..|....|..|||||||||||||
EVQLVESGGGLVQPGGSLRLSCAASGFTFDDYGMSWVRQAPGKGLEWVSGINWNGGSTGYADSVKGRFTISRDNSKNTLYLQMNSLRAEDTAVYYCAREGRGYGDYVIDYWGQGTLVTVSS
                         ^^^^^^^^                 ^^^^^^^^                                      ^^^^^^^^^^^^^^
```

Best Alignment of Certolizumab light chain to a sequence from OAS
DIQMTQSPSSLSASVGDRVTITCKASQNVGTNVAWYQQKPGKAPKALIYSASFLYSGVPYRFSGSGSGTDFTLTISSLQPEDFATYYCQQYNIYPLTFGQGTKVEIK
|||||||||||||||||||||||.||||..|...||||||||||.|||.||.|.||||.||||||||||||||||||||||||||||||||||.|||||||.|||||||
DIQMTQSPSSLSASVGDRVTITCRASQNIATYLNWYQQKPGKAPKLLIYAASSLQSGVPSRFSGSGSGTDFTLTISSLQPEDFATYYCQQYKIYPLTFGGGTKVEIK
^^^^^^ ^^^ ^^^^^^^^^

Best Alignment of Certolizumab heavy chain CDRs to a sequence from OAS
EVQLVESGGGLVQPGGSLRLSCAASGYVFTDYGMNWVRQAPGKGLEWMGWINTYIGEPIYADSVKGRFTFSLDTSKSTAYLQMNSLRAEDTAVYYCARGYRSYAMDYWGQGTLVTVSS
..|||.||..|..||.....||.||||.||.||||||.|||||||.||||||||.|||.|||..||||.|||.||.||||||.|.|..||||.|.||||..||||||||||||.|||||
QIQLVQSGPELKKPGETVKISCKASGYTFTNYGMNWVKQAPGKGLKWMGWINTYTGEPTYADDFKGRFAFSLETSASTAYLQINNLKNEDTATYFCARGGLSYAMDYWGQGTSVTVSS
^^^^^^^^ ^^^^^^^^ ^^^^^^^^^^^

Best Alignment of Certolizumab light CDRs chain to a sequence from OAS
DIQMTQSPSSLSASVGDRVTITCKASQNVGTNVAWYQQKPGKAPKALIYSASFLYSGVPYRFSGSGSGTDFTLTISSLQPEDFATYYCQQYNIYPLTFGQGTKVEIK
||.||.|....|.||||.|..||||||||||||||||||||..|||||||||..|||||.||.|||||||||||||..|.|..|.|.||||||||||||.|||.|||
DIVMTESQKFMSTSVGDGVSVTCKASQNVGTNVAWYQQKPGESPKALIYSASYRYSGVPGRFTGSGSGTDFTLTISNVQSENLAEYFCQQYNIYPLTFGSGTKLEIK
^^^^^^ ^^^ ^^^^^^^^^

Best Alignment of Certolizumab CDR-H3 chain to a sequence from OAS
EVQLVESGGGLVQPGGSLRLSCAASGYVFTDYGMNWVRQAPGKGLEWMGWINTYIGEPIYADSVKGRFTFSLDTSKSTAYLQMNSLRAEDTAVYYCARGYRSYAMDYWGQGTLVTVSS
.|.|.|||.|||.|..||..||...|.||..||||.|||||||.|.|....|...|......|...|.|.|||...|.||||...|||.|||||||||||||||||||.|||||
-VKLMESGPGLVAPSQSLSITCTVSGFSLTSYGVHWVRQPPGKGLEWLGVIWAG-GSTNYNSALMSRLSISKDNSKSQVFLKMNSLQTDDTAMYYCARGYRSYAMDYWGQGTSVTVSS
^^^^^^^^ ^^^^^^^^ ^^^^^^^^^^^

Therapeutic : Cetuximab

Best Alignment of Cetuximab heavy chain to a sequence from OAS
QVQLKQSGPGLVQPSQSLSITCTVSGFSLTNYGVHWVRQSPGKGLEWLGVIWSGGNTDYNTPFTSRLSINKDNSKSQVFFKMNSLQSNDTAIYYCARALTYYDYEFAYWGQGTLVTVSA
||||||||||||||||||||||||||||||.|||||||||||||||||||||||||.||||..|.|||||.||||||||||||||||.||||||||||.|.|.||||||||||||||||||
QVQLKQSGPGLVQPSQSLSITCTVSGFSLTSYGVHWVRQSPGKGLEWLGVIWSGGSTDYNAAFISRLSISKDNSKSQVFFKMNSLQANDTAIYYCARGLYY-DYEFAYWGQGTLVTVSA
^^^^^^^^ ^^^^^^^ ^^^^^^^^^^^^^

Best Alignment of Cetuximab light chain to a sequence from OAS
DILLTQSPVILSVSPGERVSFSCRASQSIGTNIHWYQQRTNGSPRLLIKYASESISGIPSRFSGSGSGTDFTLSINSVESEDIADYYCQQNNNWPTTFGAGTKLELK
||||||||.||||||||||||||||||||||.|||||||||||||||||||||||||||||||||||||||||||||||||||||||||||||||||.|||||||||||
DILLTQSPAILSVSPGERVSFSCRASQSIGTSIHWYQQRTNGSPRLLIKYASESISGIPSRFSGSGSGTDFTLSINSVESEDIADYYCQQNNNWPLTFGAGTKLELK
^^^^^^ ^^^ ^^^^^^^^^

Best Alignment of Cetuximab heavy chain CDRs to a sequence from OAS
QVQLKQSGPGLVQPSQSLSITCTVSGFSLTNYGVHWVRQSPGKGLEWLGVIWSGGNTDYNTPFTSRLSINKDNSKSQVFFKMNSLQSNDTAIYYCARALTYYDYEFAYWGQGTLVTVSA
.|.|..||||||||||||||||||||||||.|||||||||||||||||||||||||.||||..|.|||||.||||||||||||||||..|||||||||.|.||||.||||||||||||||
-VMLVESGPGLVQPSQSLSITCTVSGFSLTSYGVHWVRQSPGKGLEWLGVIWSGGSTDYNAAFISRLSISKDNSKSQVFFKMNSLQADDTAIYYCARNLDYYDYGFAYWGQGTLVTVSA
^^^^^^^^ ^^^^^^^ ^^^^^^^^^^^^^

Best Alignment of Cetuximab light CDRs chain to a sequence from OAS
DILLTQSPVILSVSPGERVSFSCRASQSIGTNIHWYQQRTNGSPRLLIKYASESISGIPSRFSGSGSGTDFTLSINSVESEDIADYYCQQNNNWPTTFGAGTKLELK
||||||||.||||||||||||||||||||||.||||||||||||||||||||||||||||||||||||||.|||.||||||||||||||||||||||||||||.|||||.|
DILLTQSPAILSVSPGERVSFSCRASQSIGTSIHWYQQRTNGSPRLLIKYASESISGIPSRFSGSGSGTYFTLTINSVESEDIADYYCQQNNNWPTTFGGGTKLEIK
^^^^^^ ^^^ ^^^^^^^^^

Best Alignment of Cetuximab CDR-H3 chain to a sequence from OAS
QVQLKQSGPGLVQPSQSLSITCTVSGFSLTNYGVHWVRQSPGKGLEWLGVIWSG-GNTDYNTPFTSRLSINKDNSKSQVFFKMNSLQSNDTAIYYCARALTYYDYEFAYWGQGTLVTVSA
.|||.|||..||.|..|....|..||...|.|...|..|..||.|||.|........|.||..|..........|.|.|......|.|.|.|.|||||||.|||||||||||||||||||
-VQLQQSGAELVKPGASVKMSCKASGYTFTTYPIEWMKQNHGKSLEWIGNFHPYNDDTKYNEKFKGKATLTVEKSSSTVYLELSRLTSDDSAVYYCARALIYYDYEFAYWGQGTLVTVSA
^^^^^^^^ ^^^^^^^^ ^^^^^^^^^^^^^

Therapeutic : Cixutumumab

Best Alignment of Cixutumumab heavy chain to a sequence from OAS
EVQLVQSGAEVKKPGSSVKVSCKASGGTFSSYAISWVRQAPGQGLEWMGGIIPIFGTANYAQKFQGRVTITADKSTSTAYMELSSLRSEDTAVYYCARAPLRFLEWSTQDHYYYYYMDVWGKGTTVTVSS
..||||||||||||||||||||||||||||||||||||||||||||||||||||||||||||||||||||||||||||||||||||||||||||||||||..|||||.....|||||||||||||||||||
--QLVQSGAEVKKPGSSVKVSCKASGGTFSSYAISWVRQAPGQGLEWMGGIIPIFGTANYAQKFQGRVTITADKSTSTAYMELSSLRSEDTAVYYCARASRRFLEWL-LEDYYYYYMDVWGKGTTVTVSS
^^^^^^^^ ^^^^^^^^ ^^^^^^^^^^^^^^^^^^^^^^

Best Alignment of Cixutumumab light chain to a sequence from OAS
SSELTQDPAVSVALGQTVRITCQGDSLRSYYATWYQQKPGQAPILVIYGENKRPSGIPDRFSGSSSGNTASLTITGAQAEDEADYYCKSRDGSGQHLVFGGGTKLTVL
||||||||||||||||||||||||||||||||.||||||||||.|||||||.|||||||||||||||||||||||||||||||||||||||.|||.||.||||||||||||||
SSELTQDPAVSVALGQTVRITCQGDSLRSYYASWYQQKPGQAPVLVIYGENNRPSGIPDRFSGSSSGNTASLTITGAQAEDEADYYCNSRDSSGNHLVFGGGTKLTVL
^^^^^^ ^^^ ^^^^^^^^^^^

Best Alignment of Cixutumumab heavy chain CDRs to a sequence from OAS
EVQLVQSGAEVKKPGSSVKVSCKASGGTFSSYAISWVRQAPGQGLEWMGGIIPIFGTANYAQKFQGRVTITADKSTSTAYMELSSLRSEDTAVYYCARAPLRFLEWSTQDHYYYYYMDVWGKGTTVTVSS
................||||||||||||||||||||||||||.|||||||||||||||||||||||||||||||||||.|||||||||||||||||||||||||||||.|||||||||...||||||||||||||||||.
---------------ASVKVSCKASGGTFSSYAISWVREAPGQGLEWMGGIIPIFGTANYAQKFQGRVTITADESTSTAYMELSSLRSEDTAVYYCARCPLRFLEWSTVGYYYYYYMDVWGKGTTVTVS-
^^^^^^^^ ^^^^^^^^ ^^^^^^^^^^^^^^^^^^^^^^

Best Alignment of Cixutumumab light CDRs chain to a sequence from OAS
SSELTQDPAVSVALGQTVRITCQGDSLRSYYATWYQQKPGQAPILVIYGENKRPSGIPDRFSGSSSGNTASLTITGAQAEDEADYYCKSRDGSGQHLVFGGGTKLTVL
||||||||||||||||||||||||||||||.|||||||.|||||||||.||||.||||||.|||||||||||||||||.||||||||||.||.|||||||.||||.|||
SSELTQDPAVSVALGQTVRITCQGDSLRSYHATWYQQKQGQAPILVMYGENNRPSGIPERFSGSSSGNTASLTITGTQAEDEADYYCNSRDSSGQHLVFAGGTKVTVL
^^^^^^ ^^^ ^^^^^^^^^^^

Best Alignment of Cixutumumab CDR-H3 chain to a sequence from OAS
EVQLVQSGAEVKKPGSSVKVSCKASGGTFSSYAISWVRQAPGQGLEWMGGIIPIFGTANYAQKFQGRVTITADKSTSTAYMELSSLRSEDTAVYYCARAPLRFLEWSTQDHYYYYYMDVWGKGTTVTVSS
................||||||||||||||||||||||||||.|||||||||||||||||||||||||||||||||||.|||||||||||||||||||||||||||||.|||||||||...||||||||||||||||||.
---------------ASVKVSCKASGGTFSSYAISWVREAPGQGLEWMGGIIPIFGTANYAQKFQGRVTITADESTSTAYMELSSLRSEDTAVYYCARCPLRFLEWSTVGYYYYYYMDVWGKGTTVTVS-
^^^^^^^^ ^^^^^^^^ ^^^^^^^^^^^^^^^^^^^^^^

Therapeutic : Clazakizumab

Best Alignment of Clazakizumab heavy chain to a sequence from OAS
EVQLVESGGGLVQPGGSLRLSCAASGFSLSNYYVTWVRQAPGKGLEWVGIIYGS-DETAYATSAIGRFTISRDNSKNTLYLQMNSLRAEDTAVYYCARDDSSD-WDAKFNLWGQGTLVTVSS
||||.||||||||||||||||||||||..|||..||||||||||||||..|.||...|.||.|..|||||||||||||||||||||||||||||||||||||||.||......|.|||||||||||
EVQLLESGGGLVQPGGSLRLSCAASGFTFSNYAMTWVRQAPGKGLEWVSAISGSGGSTYYADSVKGRFTISRDNSKNTLYLQMNSLRAEDTAVYYCARDPSSGYYGGFFDYWGQGTLVTVSS
^^^^^^^^ ^^^^^^^^ ^^^^^^^^^^^^^^^

Best Alignment of Clazakizumab light chain to a sequence from OAS
AIQMTQSPSSLSASVGDRVTITCQASQSINNELSWYQQKPGKAPKLLIYRASTLASGVPSRFSGSGSGTDFTLTISSLQPDDFATYYCQQGYSLRNIDNAFGGGTKVEIK
.||||||||||||||||||||||.||||||||.||||||||||||||||||.||.|.|||||||||||||||||||||||||||||||.||||||||||||.......||||||||||
DIQMTQSPSSLSASVGDRVTITCRASQSINNYLSWYQQKPGKAPKLLIYAASSLQSGVPSRFSGSGSGTDFTLTISSLQPEDFATYYCQQGYS--TPRITFGGGTKVEIK
^^^^^^ ^^^ ^^^^^^^^^^^^

Best Alignment of Clazakizumab heavy chain CDRs to a sequence from OAS
EVQLVESGGGLVQPGGSLRLSCAASGFSLSNYYVTWVRQAPGKGLEWVGIIYGSDETAYATSAIGRFTISRDNSKNTLYLQMNSLRAEDTAVYYCARDDSSDWDAKFNLWGQGTLVTVSS
..||.||||.||.|.|||.|||.||||.||.||..||||||||||||.|.|..|..|.||....|||.||.||..||..||||||.|.|||.|.||||.||||||...|||||.|||||||.
--QLEESGGDLVKPEGSLTLSCKASGFDLSTYYMNWVRQAPGKGLEWIGMILPSGTTYYASWVNGRFAISSDNAQNTVDLQMNSLTAADTATYFCARDYSSDWGWEFNLWGPGTLVTVS-
^^^^^^^^ ^^^^^^^ ^^^^^^^^^^^^^^

Best Alignment of Clazakizumab light CDRs chain to a sequence from OAS

```
AIQMTQSPSSLSASVGDRVTITCQASQSINNELSWYQQKPGKAPKLLIYRASTLASGVPSRFSGSGSGTDFTLTISSLQPDDFATYYCQQGYSLRNIDNAFGGGTKVEIK
.................|.|||.|||||||.|.||||||||||||||||.||.|.|||||||||||||||||||||||||||.|||||||||.||.....|.||.|||.|||
-----------------VSITCRASQSINNYLNWYQQKPGKAPKLLIYAASSLQSGVPSRFSGSGSGTDFTLTISSLQPEDFATYYCQQSYSDPPKLNTFGQGTKLEIK
                    ^^^^^^          ^^^                    ^^^^^^^^^^^^
```

Best Alignment of Clazakizumab CDR-H3 chain to a sequence from OAS

```
EVQLVESGGGLVQPGGSLRLSCAASGFSL-SNYYVTWVRQAPGKGLEWVGIIYGS-DETAYATSAIGRFTISRDNSKNTLYLQMNSLRAEDTAVYYCARDDSSDWDAKFNLWGQGTLVTVSS
...|.|||||||||||||.|.|.|||||....|...||||||||||||...|..|...|.||..|.|.||||...|..|..||||||.|.|||.|.||||||||||...|||||.|||||||.
Q-SLEESGGGLVQPGGSLTLTCKASGFSFSNRYVMCWVRQAPGKGLEWIACINTSSGNTVYASWAKGPFTISKTSS-TTVTLQMNSLTAADTATYFCARDDSSDWGWEFNLWGPGTLVTVS-
                    ^^^^^^^^^          ^^^^^^^^                    ^^^^^^^^^^^^^^
```

Therapeutic : Codrituzumab

Best Alignment of Codrituzumab heavy chain to a sequence from OAS

```
QVQLVQSGAEVKKPGASVKVSCKASGYTFTDYEMHWVRQAPGQGLEWMGALDPKTGDTAYSQKFKGRVTLTADKSTSTAYMELSSLTSEDTAVYYCTRFY-SYTYWGQGTLVTVSS
||||||||||||||||||||||||||||||.|.||||||||||||||||...|..|.|.|.|||.||||.||||||||||||||||.|||||||||.|.|..|.||||||||||||
QVQLVQSGAEVKKPGASVKVSCKASGYTFTSYYMHWVRQAPGQGLEWMGIINPSGGSTSYAQKFQGRVTITADKSTSTAYMELSSLRSEDTAVYYCARGYSNYDYWGQGTLVTVSS
                    ^^^^^^^^          ^^^^^^^^                    ^^^^^^^^^
```

Best Alignment of Codrituzumab light chain to a sequence from OAS

```
DVVMTQSPLSLPVTPGEPASISCRSSQSLVHSNRNTYLHWYLQKPGQSPQLLIYKVSNRFSGVPDRFSGSGSGTDFTLKISRVEAEDVGVYYCSQNTHVPPTFGQGTKLEIK
||||||.||||||.||.||||||||||||||||.||||||||||||||||.||||||||||||||||||||||||||||||||||||||.|||.|||.||||.|||.|||||||
DVVMTQTPLSLPVSPGDPASISCRSSQSLVHSNGNTYLHWYLQKPGQSPKLLIYKVSNRFSGVPDRFSGSGSGTDFTLKISRVEAEDLGVYFCSQSTHVPWTFGGGTKLEIK
                     ^^^^^^^^^^^          ^^^                    ^^^^^^^^^
```

Best Alignment of Codrituzumab heavy chain CDRs to a sequence from OAS

```
QVQLVQSGAEVKKPGASVKVSCKASGYTFTDYEMHWVRQAPGQGLEWMGALDPKTGDTAYSQKFKGRVTLTADKSTSTAYMELSSLTSEDTAVYYCTRFYSYTYWGQGTLVTVSS
.|||.|||||...|||||..||||||||||||||||.|.|..||||.||.||.||.|||.|||||...||||||.||||.||.||||||.|||||||||.||||||||..||||
-VQLQQSGAELVRPGASVTLSCKASGYTFTDYEMHWVKQTPVHGLEWIGAIDPETGGTAYNQKFKGKAILTADKSSSTAYLELRSLTSEDSAVYYCTRFYGYTYWGQGTTLTVSS
                    ^^^^^^^^          ^^^^^^^^                    ^^^^^^^^
```

Best Alignment of Codrituzumab light CDRs chain to a sequence from OAS

```
DVVMTQSPLSLPVTPGEPASISCRSSQSLVHSNRNTYLHWYLQKPGQSPQLLIYKVSNRFSGVPDRFSGSGSGTDFTLKISRVEAEDVGVYYCSQNTHVPPTFGQGTKLEIK
||||||.||||||..|..|||||||||||||||.||||||||||||||||.||||||||||||||||||||||||||||||||||||||.|||.|||.|||||||||.|||||||
DVVMTQTPLSLPVSLGDQASISCRSSQSLVHSNGNTYLHWYLQKPGQSPKLLIYKVSNRFSGVPDRFSGSGSGTDFTLKISRVEAEDLGVYFCSQSTHVPPTFGGGTKLEIK
                     ^^^^^^^^^^^          ^^^                    ^^^^^^^^^
```

Best Alignment of Codrituzumab CDR-H3 chain to a sequence from OAS

```
QVQLVQSGAEVKKPGASVKVSCKASGYTFTDYEMHWVRQAPGQGLEWMGALDPK--TGDTAYSQKFKGRVTLTADKSTSTAYMELSSLTSEDTAVYYCTRFYSYTYWGQGTLVTVSS
...||.||.....||.|.|.||.|||.||..|.|.||||.|..||||......|.....|.|....|||.|...|.|.|..|.....|..|||..|||||.|||||||.||..||||
DGMLVESGGGLVQPGGSMKLSCVASGFTFSNYWMNWVRQSPEKGLEWVAEIRLKSNNYATHYAESVKGRFTISRDDSKSSVYLQMNNLRAEDTGIYYCTRGYSYTYWGPGTTLTVSS
                    ^^^^^^^^          ^^^^^^^^^^                    ^^^^^^^^
```

Therapeutic : Coltuximab

Best Alignment of Coltuximab heavy chain to a sequence from OAS

```
QVQLVQPGAEVVKPGASVKLSCKTSGYTFTSNWMHWVKQAPGQGLEWIGEIDPSDSYTNYNQNFQGKAKLTVDKSTSTAYMEVSSLRSDDTAVYYCARGSNPYYYAMDYWGQGTSVTVSS
||||.|||||.||||||||||||.|||||||.||.||||.||||||||||||||||||||||.|.|||.||||.|.|||||..|||.|.|.||||||||||||||||||||||||||||||
QVQLQQPGAELVKPGASVKLSCKASGYTFTSYWMQWVKQRPGQGLEWIGEIDPSDSYTNYNQKFKGKATLTVDTSSSTAYMQLSSLTSEDSAVYYCARGSNPYYYAMDYWGQGTSVTVSS
                    ^^^^^^^^          ^^^^^^^^                    ^^^^^^^^^^^^^
```

Best Alignment of Coltuximab light chain to a sequence from OAS

```
EIVLTQSPAIMSASPGERVTMTCSASSGVNYMHWYQQKPGTSPRRWIYDTSKLASGVPARFSGSGSGTDYSLTISSMEPEDAATYYCHQRGSYTFGGGTKLEIK
.|||||||||||||||.||||||||||..||||||||||||||.||||||||||||||||||||||||.|||||||||.||||||||||||.|||||||||||||
QIVLTQSPAIMSASPGEKVTMTCSASSSINYMHWYQQKPGTSPKRWIYDTSKLASGVPARFSGSGSGTSYSLTISSMEAEDAATYYCHQRSSYTFGGGTKLEIK
                     ^^^^^          ^^^                    ^^^^^^^
```

Best Alignment of Coltuximab heavy chain CDRs to a sequence from OAS

```
QVQLVQPGAEVVKPGASVKLSCKTSGYTFTSNWMHWVKQAPGQGLEWIGEIDPSDSYTNYNQNFQGKAKLTVDKSTSTAYMEVSSLRSDDTAVYYCARGSNPYYYAMDYWGQGTSVTVSS
.|||.|||||.|.||||||||||.|||||||.|||||||.||||||||||||||||||||||.|.||..||||||.|||||..|||.|.|.||||||||||||||||||||||||||||||
-VQLQQPGAELVMPGASVKLSCKASGYTFTSYWMHWVKQRPGQGLEWIGEIDPSDSYTNYNQKFKGKSTLTVDKSSSTAYMQLSSLTSEDSAVYYCARGSNPYYYAMDYWGQGTSVTVSS
                    ^^^^^^^^          ^^^^^^^^                    ^^^^^^^^^^^^^
```

Best Alignment of Coltuximab light CDRs chain to a sequence from OAS

```
EIVLTQSPAIMSASPGERVTMTCSASSGVNYMHWYQQKPGTSPRRWIYDTSKLASGVPARFSGSGSGTDYSLTISSMEPEDAATYYCHQRGSYTFGGGTKLEIK
.||.||..||||||.||.|||.|.|||.||||.|||||...||.|||||||||||||||||||||.||.|||||||||.|||||||||||.|||||||||||||
-IVMTQTTAIMSASLGEKVTMSCRASSSVNYMYWYQQKSDASPKRWIYDTSKLASGVPARFSGSGPGTSYSLTISSMEAEDAATYYCHQRSSYTFGGGTKLEIK
                     ^^^^^          ^^^                    ^^^^^^^
```

Best Alignment of Coltuximab CDR-H3 chain to a sequence from OAS

```
QVQLVQPGAEVVKPGASVKLSCKTSGYTFTSNWMHWVKQAPGQGLEWIGEIDPSDSYTNYNQNFQGKAKLTVDKSTSTAYMEVSSLRSDDTAVYYCARGSNPYYYAMDYWGQGTSVTVSS
.|||.|.|.|.||.|||||.|||.|||.||...||||||..|..|||||.|......|.|||.|.|||..|||.|.|||||...||.|.|.||||||||||||||||||||||||||||||
-VQLQQSGPELVKTGASVKISCKASGYSFTGYYMHWVKQSHGKSLEWIGYISCYNGATSYNQKFKGKATFTVDTSSSTAYMQSNSLTSEDSAVYYCARGSNPYYYAMDYWGQGTSVTVSS
                    ^^^^^^^^          ^^^^^^^^                    ^^^^^^^^^^^^^
```

Therapeutic : Concizumab

Best Alignment of Concizumab heavy chain to a sequence from OAS

```
EVQLVESGGGLVKPGGSLRLSCAASGFTFSNYAMSWVRQTPEKRLEWVATISRSGSYSYFPDSVQGRFTISRDNAKNSLYLQMNSLRAEDTAVYYCARLGGYDEGDAMDSWGQGTTVTVSS
|||||||||||||||||.|||||||||||.|||||||||||||||||||||..|||.|.||||.||||||||||||.|||||.|||.||||.|||||||||||.||||.|||||.|||||
EVQLVESGGGLVKPGGSLKLSCAASGFTFSSYAMSWVRQTPEKRLEWVATISSGGSYTYYPDSVKGRFTISRDNAKNTLYLQMSSLRSEDTAMYYCARLGGYDERDAMDYWGQGTSVTVSS
                    ^^^^^^^^          ^^^^^^^^                    ^^^^^^^^^^^^^^
```

Best Alignment of Concizumab light chain to a sequence from OAS

```
DIVMTQTPLSLSVTPGQPASISCKSSQSLLESDGKTYLNWYLQKPGQSPQLLIYLVSILDSGVPDRFSGSGSGTDFTLKISRVEAEDVGVYYCLQATHFPQTFGGGTKVEIK
|||||||||||||||||||||||||||||||.|||||||.|||||||||||||||.||...|||||||||||||||||||||||||||||||||||||.||.|.|||||||||||
DIVMTQTPLSLSVTPGQPASISCKSSQSLLHSDGKTYLCWYLQKPGQSPQLLIYEVSSRFSGVPDRFSGSGSGTDFTLKISRVEAEDVGVYYCLQGTHLPLTFGGGTKVEIK
                     ^^^^^^^^^^^          ^^^                    ^^^^^^^^^
```

Best Alignment of Concizumab heavy chain CDRs to a sequence from OAS

```
EVQLVESGGGLVKPGGSLRLSCAASGFTFSNYAMSWVRQTPEKRLEWVATISRSGSYSYFPDSVQGRFTISRDNAKNSLYLQMNSLRAEDTAVYYCARLGGYDEGDAMDSWGQGTTVTVSS
.|||..||||||||||||.|||||||||||.|||||||||||||||||||||..|||.|.||||.||||||||||||.|||||.|||.||||.|||||||||||.||||.|||||.|||||
-VQLKQSGGGLVKPGGSLKLSCAASGFTFSSYAMSWVRQTPEKRLEWVATISSGGSYTYYPDSVKGRFTISRDNAKNTLYLQMSSLRSEDTAMYYCARLGGYDERDAMDYWGQGTSVTVSS
                    ^^^^^^^^          ^^^^^^^^                    ^^^^^^^^^^^^^^
```

Best Alignment of Concizumab light CDRs chain to a sequence from OAS

```
DIVMTQTPLSLSVTPGQPASISCKSSQSLLESDGKTYLNWYLQKPGQSPQLLIYLVSILDSGVPDRFSGSGSGTDFTLKISRVEAEDVGVYYCLQATHFPQTFGGGTKVEIK
|.|||||||.||||.||||||||||||||||.||||||||||.||.||.||.||||||||.|||||||||.||||||||||||||||||||||.||||||||||||||||||||.|||
DVVMTQTPLTLSVTIGQPASISCKSSQSLLDSDGKTYLNWLLQRPGESPKLLIYLVSKLDSGVPDRFTGSGSGTDFTLKISRVEAEDLGVYYCLQATHFPQTFGGGTKLEIK
                     ^^^^^^^^^^^          ^^^                    ^^^^^^^^^
```

Best Alignment of Concizumab CDR-H3 chain to a sequence from OAS

```
EVQLVESGGGLVKPGGSLRLSCAASGFTFSNYAMSWVRQTPEKRLEWVATISRSGSYSYFPDSVQGRFTISRDNAKNSLYLQMNSLRAEDTAVYYCARLGGYDEGDAMDSWGQGTTVTVSS
.|||..||..|||||.|...||.|||.||..|.|.||.|...|.|||...|..............|..|...|......|...|||..||.|||||||.||||||||||.|||||.|||||
-VQLQQSGPVLVKPGASVKMSCQASGYTFTDYYMNWVKQSHGKSLEWIGVINPYNGGTSYNQKFKGKATLTVDKSSSTAYMELNSLTSEDSAVYYCAREGGYDEGDAMDYWGQGTSVTVSS
                    ^^^^^^^^          ^^^^^^^^                    ^^^^^^^^^^^^^^
```

Therapeutic : Crenezumab

Best Alignment of Crenezumab heavy chain to a sequence from OAS
EVQLVESGGGLVQPGGSLRLSCAASGFTFSSYGMSWVRQAPGKGLELVASINSNGGSTYYPDSVKGRFTISRDNAKNSLYLQMNSLRAEDTAVYYCASG--------DYWGQGTTVTVSS
||||.|||||||||||||||||||||||||||.|||||||||||||.|..|...||||||.|||||||||||||||||||||||||||||||||||||||........|||||||.|||||
EVQLLESGGGLVQPGGSLRLSCAASGFTFSSYAMSWVRQAPGKGLEWVSAISGSGGSTYYADSVKGRFTISRDNAKNSLYLQMNSLRAEDTAVYYCASGYNYGYLYFDYWGQGTLVTVSS
^^^^^^^^ ^^^^^^^^ ^^^^^^^^^^^^^

Best Alignment of Crenezumab light chain to a sequence from OAS
DIVMTQSPLSLPVTPGEPASISCRSSQSLVYSNGDTYLHWYLQKPGQSPQLLIYKVSNRFSGVPDRFSGSGSGTDFTLKISRVEAEDVGVYYCSQSTHVPWTFGQGTKVEIK
||||||||||||||||||||||||||||||.|||..||.|||||||||||||||.||||.||||||||||||||||||||||||||||||||||||.|..|.|||||||||||||
DIVMTQSPLSLPVTPGEPASISCRSSQSLVRSNGNIYLDWYLQKPGQSPQLLIYLVSNRASGVPDRFSGSGSGTDFTLKISRVEAEDVGVYYCMQGIHLPWTFGQGTKVEIK
^^^^^^^^^^^ ^^^ ^^^^^^^^^

Best Alignment of Crenezumab heavy chain CDRs to a sequence from OAS
EVQLVESGGGLVQPGGSLRLSCAASGFTFSSYGMSWVRQAPGKGLELVASINSNGGSTYYPDSVKGRFTISRDNAKNSLYLQMNSLRAEDTAVYYCASGDYWGQGTTVTVSS
.|||..||||||||||||.|||||||||||||||||||..|...|||||.|||||||||||||||||.|||||||||.||||||.||..||||.|||||.||||||||.|||||
-VQLKQSGGGLVQPGGSLKLSCAASGFTFSSYGMSWVRRTPDERLELVATINSNGGSTYYPDSVKGRSTISRDNAKNTLYLQMSSLKSEDTAMYYCASWDYWGQGTTLTVSS
^^^^^^^^ ^^^^^^^^ ^^^^^

Best Alignment of Crenezumab light CDRs chain to a sequence from OAS
DIVMTQSPLSLPVTPGEPASISCRSSQSLVYSNGDTYLHWYLQKPGQSPQLLIYKVSNRFSGVPDRFSGSGSGTDFTLKISRVEAEDVGVYYCSQSTHVPWTFGQGTKVEIK
|||.|||||||||..|..|||||.||||||||||||||||||||||||||.||||||||||||.|||||||||||||||||||||||||||.|||.|||||||||||||.|||.|||
DIVITQSPLSLPVSLGDQASISCSSSQSLVYSNGDTYLHWYLQKPGQSPKLLIYKVSNRFSGAPDRFSGSGSGTDFTLKISRVEAEDLGVYVCSQSTHVPWTFGGGTKLEIK
^^^^^^^^^^^ ^^^ ^^^^^^^^^

Best Alignment of Crenezumab CDR-H3 chain to a sequence from OAS
EVQLVESGGGLVQPGGSLRLSCAASGFTFSSYGMSWVRQAPGKGLELVASINSNGGSTYYPDSVKGRFTISRDNAKNSLYLQMNSLRAEDTAVYYCASGDYWGQGTTVTVSS
||||..|...||.||.|....|.|||.||..|.|.||.|..||.||....||.|.|.|.|....||......|....||..||||||||||||||||||||.|||||
EVQLQQSRPELVKPGASVKIPCKASGYTFTDYNMDWVNQSHGKSLEWIGDINPNNGGTIYNQKFKGKATLTVDKSSSTAYMELRSLTSEDTAVYYCASGDYWGQGTSVTVSS
^^^^^^^^ ^^^^^^^^ ^^^^^

Therapeutic : Crizanlizumab

Best Alignment of Crizanlizumab heavy chain to a sequence from OAS
QVQLVQSGAEVKKPGASVKVSCKVSGYTFTSYDINWVRQAPGKGLEWMGWIYPGDGSIKYNEKFKGRVTMTVDKSTDTAYMELSSLRSEDTAVYYCARRGEYGN--YEGAMDYWGQGTLVTVSS
|||||||||||||||||||||||.||||||||...|||||||.||||||.|.|..||..||..||.|||||.|.||.|||||||||||||||||||||||.|.||.....||.|||||||||||||
QVQLVQSGAEVKKPGASVKVSCKASGYTFTSYYMHWVRQAPGQGLEWMGIINPSGGSTSYAQKFQGRVTMTRDTSTSTAYMELSSLRSEDTAVYYCARAGTYGITGTTGAFDYWGQGTLVTVSS
^^^^^^^^ ^^^^^^^^ ^^^^^^^^^^^^^^^^

Best Alignment of Crizanlizumab light chain to a sequence from OAS
DIQMTQSPSSLSASVGDRVTITCKASQSVDYDGHSYMNWYQQKPGKAPKLLIYAASNLESGVPSRFSGSGSGTDFTLTISSLQPEDFATYYCQQSDENPLTFGGGTKVEIK
|||||||||||||||||||||||.||||......||.|||||||||||||||||||||.|.|||||||||||||||||||||||||||||||||||||||.|||||||||||||||
DIQMTQSPSSLSASVGDRVTITCRASQSI----SSYLNWYQQKPGKAPKLLIYAASSLQSGVPSRFSGSGSGTDFTLTISSLQPEDFATYYCQQSDSNPLTFGGGTKVEIK
^^^^^^^^^^ ^^^ ^^^^^^^^^

Best Alignment of Crizanlizumab heavy chain CDRs to a sequence from OAS
QVQLVQSGAEVKKPGASVKVSCKVSGYTFTSYDINWVRQAPGKGLEWMGWIYPGDGSIKYNEKFKGRVTMTVDKSTDTAYMELSSLRSEDTAVYYCARRGEYGNYEGAMDYWGQGTLVTVSS
||||.|||.|..||||.||.|||.|||||||||||||.|.||.||||.|||||||||.||||||||..|.|.|||..||||.||||.||..|||.|||.||||||.||||||||||.|||||
QVQLQQSGPELVKPGALVKISCKASGYTFTSYDINWVKQRPGQGLEWIGWIYPGDGSTKYNEKFKGKATLTADKSSSTAYMQLSSLTSENSAVYFCARSGEYGNYGGAMDYWGQGTSVTVSS
^^^^^^^^ ^^^^^^^^ ^^^^^^^^^^^^^^^

Best Alignment of Crizanlizumab light CDRs chain to a sequence from OAS
DIQMTQSPSSLSASVGDRVTITCKASQSVDYDGHSYMNWYQQKPGKAPKLLIYAASNLESGVPSRFSGSGSGTDFTLTISSLQPEDFATYYCQQSDENPLTFGGGTKVEIK
||..||||.||..|.|.|.||.|||||||||||.||||||||||...|||||||||||.||.|.|||||||||||||||.|.....||.|||.||||.|||.|||||||.|||
DIVLTQSPASLAVSLGQRATISCKASQSVDYDGDSYMNWYQQKPVQPPKLLIYAASNLGSGIPARFSGSGSGTDFTLNIHPVEEEDAATYCCQQSNENPYTFGGGTKLEIK
^^^^^^^^^^ ^^^ ^^^^^^^^^

Best Alignment of Crizanlizumab CDR-H3 chain to a sequence from OAS
QVQLVQSGAEVKKPGASVKVSCKVSGYTFTSYDINWVRQAPGKGLEWMGWIYPGDGSIKYNEKFKGRVTMTVDKSTDTAYMELSSLRSEDTAVYYCARRGEYGNYEGAMDYWGQGTLVTVSS
||||.|.|||...||.|||.|||.||||||||...||.|.|..||||.|.|.|.|....||.|.|...|.|||||..||||.||||.|||..||||||||||||||.||||||||||.|||||
QVQLKQPGAELVRPGSSVKLSCKASGYTFTSYWMHWVKQRPIQGLEWIGNIDPSDSETYYNQKIKDHATLTVDKSSSTAYMQLSSLTSEDSEVYYCARRGEYGNYGGAMDYWGQGTSVTVSS
^^^^^^^^ ^^^^^^^^ ^^^^^^^^^^^^^^^

Therapeutic : Dacetuzumab

Best Alignment of Dacetuzumab heavy chain to a sequence from OAS
EVQLVESGGGLVQPGGSLRLSCAASGYSFTGYYIHWVRQAPGKGLEWVARVIPNAGGTSYNQKFKGRFTLSVDNSKNTAYLQMNSLRAEDTAVYYCAREG-----IYWWGQGTLVTVSS
||||||||||||||||||||||||||..|..|..|||||||||||.||.|.......|||....|||||.|.||.|.||||||||||||||||||||||||.....|..|||||||||||
EVQLVESGGGLVQPGGSLRLSCAASGFTFSSYWMHWVRQAPGKGLVWVSRINSDGSSTSYADSVKGRFTISRDNAKNTLYLQMNSLRAEDTAVYYCAREGYYDNYIDYWGQGTLVTVSS
^^^^^^^^ ^^^^^^^^ ^^^^^^^^^^^^

Best Alignment of Dacetuzumab light chain to a sequence from OAS
DIQMTQSPSSLSASVGDRVTITCRSSQSLVHSNGNTFLHWYQQKPGKAPKLLIYTVSNRFSGVPSRFSGSGSGTDFTLTISSLQPEDFATYFCSQTTHVPWTFGQGTKVEIK
|||||||||||||||||||||||||||.|||.......||||||||||||||||||||..||..|||||||||||||||||||||||||||||||||.|.|....|.|||||||||||
DIQMTQSPSSLSASVGDRVTITCRASQSI-----STFLHWYQQKPGKAPKLLIYAASNLQSGVPSRFSGSGSGTDFTLTISSLQPEDFATYYCQQIYSTPKTFGQGTKVEIK
^^^^^^^^^^^ ^^^ ^^^^^^^^^

Best Alignment of Dacetuzumab heavy chain CDRs to a sequence from OAS
EVQLVESGGGLVQPGGSLRLSCAASGYSFTGYYIHWVRQAPGKGLEWVARVIPNAGGTSYNQKFKGRFTLSVDNSKNTAYLQMNSLRAEDTAVYYCAREGIYWWGQGTLVTVSS
||.|.|||..||.||.|...||.||||||||||.|||.|..||.|||..||.||.|||||||||||||...|.||.|..|||....||.|||.|||||||||.|.|||||..||||
EVKLMESGPDLVKPGASVKISCKASGYSFTGYYMHWVKQSHGKSLEWIGRVNPNNGGTSYNQKFKGKAILTVDKSSSTAYMELRSLTSEDSAVYYCAREGYYGWGQGTTLTVSS
^^^^^^^^ ^^^^^^^^ ^^^^^^^

Best Alignment of Dacetuzumab light CDRs chain to a sequence from OAS
DIQMTQSPSSLSASVGDRVTITCRSSQSLVHSNGNTFLHWYQQKPGKAPKLLIYTVSNRFSGVPSRFSGSGSGTDFTLTISSLQPEDFATYFCSQTTHVPWTFGQGTKVEIK
|..|||.|.||..|.||...|.||||||||||||||.||||.||||..||||||||||||||||||.|||||||||||||||.||....||...|||||.||||||||.|||.|||
DVVMTQTPLSLPVSLGDQASISCRSSQSLVHSNGNTYLHWYLQKPGQSPKLLIYTVSNRFSGVPDRFSGSGSGTDFTLKISRVEAEDLGVYFCSQSTHVPWTFGGGTKLEIK
^^^^^^^^^^^ ^^^ ^^^^^^^^^

Best Alignment of Dacetuzumab CDR-H3 chain to a sequence from OAS
EVQLVESGGGLVQPGGSLRLSCAASGYSFTGYYIHWVRQAPGKGLEWVARVIPNAGGTSYNQKFKGRFTLSVDNSKNTAYLQMNSLRAEDTAVYYCAREGIYWWGQGTLVTVSS
||.|||||||||.|||||.|||||||..|..|...||||.|.|.|||||......|.|.|....|||||.|.||.|...||||.|||.||||.|||||||||||.|||.|..||||
EVMLVESGGGLVKPGGSLKLSCAASGFTFSSYAMSWVRQTPEKRLEWVASISSG-GSTYYPDSVKGRFTISRDNARNILYLQMSSLRSEDTAMYYCAREGIYYWGQCTTLTVSS
^^^^^^^^ ^^^^^^^^ ^^^^^^^

Therapeutic : Daclizumab

Best Alignment of Daclizumab heavy chain to a sequence from OAS
QVQLVQSGAEVKKPGSSVKVSCKASGYTFTSYRMHWVRQAPGQGLEWIGYINPSTGYTEYNQKFKDKATITADESTNTAYMELSSLRSEDTAVYYCARGGG---VFDYWGQGTLVTVSS
|||||||||||||||.||||||||||||||||.||||||||||||||.|.||||.|.|.|.|||.|..||||||||.||||||||||||||||||||||.||....||||||||||||||
QVQLVQSGAEVKKPGASVKVSCKASGYTFTSYYMHWVRQAPGQGLEWMGIINPSGGSTSYAQKFQDRVTITADESTSTAYMELSSLRSEDTAVYYCARDGGSGWDFDYWGQGTLVTVSS
^^^^^^^^ ^^^^^^^^ ^^^^^^^^^^^^

Best Alignment of Daclizumab light chain to a sequence from OAS

```
DIQMTQSPSTLSASVGDRVTITCSASSSI-SYMHWYQQKPGKAPKLLIYTTSNLASGVPARFSGSGSGTEFTLTISSLQPDDFATYYCHQRSTYPLTFGQGTKVEVK
||||||||||||||||||||||.||.||.|...|||||||||||||||..|||.||||.|||||||||||||||||||||||||||||.|.||||.|||||||||||
DIQMTQSPSTLSASVGDRVTITCRASQSISSWLAWYQQKPGKAPKLLIYKASNLESGVPSRFSGSGSGTEFTLTISSLQPDDFATYYCQQYSTYPWTFGQGTKVEVK
                        ^^^^^^          ^^^                   ^^^^^^^^^
```

Best Alignment of Daclizumab heavy chain CDRs to a sequence from OAS

```
QVQLVQSGAEVKKPGSSVKVSCKASGYTFTSYRMHWVRQAPGQGLEWIGYINPSTGYTEYNQKFKDKATITADESTNTAYMELSSLRSEDTAVYYCARGGGVFDYWGQGTLVTVSS
.|.|..||||..|||.|||.|||||||||||||||||.|.|||||||||||||||||||||||||||||.|||.|..||||.||||.|||.||||||||||.|.|.|||||||||||.
-VKLMESGAELAKPGASVKMSCKASGYTFTSYRMHWVKQRPGQGLEWIGYINPSTGYTEYNQKFKDKATLTADKSSSTAYMQLSSLTSEDSAVYYCARGGGWFAYWGQGTLVTVSA
                       ^^^^^^^^        ^^^^^^^^                   ^^^^^^^^^
```

Best Alignment of Daclizumab light CDRs chain to a sequence from OAS

```
DIQMTQSPSTLSASVGDRVTITCSASSSISYMHWYQQKPGKAPKLLIYTTSNLASGVPARFSGSGSGTEFTLTISSLQPDDFATYYCHQRSTYPLTFGQGTKVEVK
.|..|||....|||.|..||.||.||||||||||||||||..|...||.||.||||||||||||||||...|||||....|.|||||||||.||||||.|||.|.|
QIVLTQSRAIMSASPGEKVTMTCRASSSISYMHWYQQKPGTSPQSWIYYTSKLASGVPARFSGSGSGTAYSLTISSMEAEDAATYYCHQRSNYPLTFGGGTKLEIK
                        ^^^^^          ^^^                   ^^^^^^^^^
```

Best Alignment of Daclizumab CDR-H3 chain to a sequence from OAS

```
QVQLVQSGAEVKKPGSSVKVSCKASGYTFTSYRMHWVRQAPGQGLEWIGYINPSTGYTEYNQKFKDKATITADESTNTAYMELSSLRSEDTAVYYCARGGGVFDYWGQGTLVTVSS
.............|.......|..||....||...|.||.||.||||||||..|.|.|.||...|...||..|.|.|.....|||....|||||||||||||||||||||||||.|
-------------PSETLSLTCTVSGGSISSYYWSWIRQPPGKGLEWIGYIYYS-GSTNYNPSLKSRVTISVDTSKNQFSLKLSSVTAADTAVYYCARGGGVFDYWGQGTLVTVYS
                       ^^^^^^^^        ^^^^^^^^                   ^^^^^^^^^
```

Therapeutic : Dalotuzumab

Best Alignment of Dalotuzumab heavy chain to a sequence from OAS

```
QVQLQESGPGLVKPSETLSLTCTVSGYSITGGYLWNWIRQPPGKGLEWIGYISYDGTNNYKPSLKDRVTISRDTSKNQFSLKLSSVTAADTAVYYCARYGR-------VFFDYWGQGTLVTVSS
||||||||||||||||||||||||||.||...|.|.|||||||||||||||||.|.|..||.||||.|||||.|||||||||||||||||||||||||||.||.......|||||||||||||||
QVQLQESGPGLVKPSETLSLTCTVSGGSI-SSYYWSWIRQPPGKGLEWIGYIYYSGSTNYNPSLKSRVTISVDTSKNQFSLKLSSVTAADTAVYYCARGGRYYDSSSAVFFDYWGQGTLVTVSS
                       ^^^^^^^^^       ^^^^^^^                   ^^^^^^^^^^^^^^^^^
```

Best Alignment of Dalotuzumab light chain to a sequence from OAS

```
DIVMTQSPLSLPVTPGEPASISCRSSQSIVHSNGNTYLQWYLQKPGQSPQLLIYKVSNRLYGVPDRFSGSGSGTDFTLKISRVEAEDVGVYYCFQGSHVPWTFGQGTKVEIK
||||||||||||||||||||||||||||.|.||||.||.|||||||||||||||.||||..||||||||||||||||||||||||||||||||.||.|.|||||||||||||
DIVMTQSPLSLPVTPGEPASISCRSSQSLVRSNGNIYLDWYLQKPGQSPQLLIYLVSNRASGVPDRFSGSGSGTDFTLKISRVEAEDVGVYYCMQGIHLPWTFGQGTKVEIK
                        ^^^^^^^^^^^          ^^^                  ^^^^^^^^^
```

Best Alignment of Dalotuzumab heavy chain CDRs to a sequence from OAS

```
QVQLQESGPGLVKPSETLSLTCTVSGYSITGGYLWNWIRQPPGKGLEWIGYISYDGTNNYKPSLKDRVTISRDTSKNQFSLKLSSVTAADTAVYYCARYGRVFFDYWGQGTLVTVSS
.|||||||||||||..|||||.|.|||||.||.||||||.||..|||.|||||||.|||.||||.|..|.||||||||.|||.|||..|||.||||||.|.|||||||||..||||
EVQLQESGPGLVKPSQSLSLTCSVTGYSITSGYYWNWIRQFPGNKLEWMGYISYDGSNNYNPSLKNRISITRDTSKNQFFLKLNSVTTEDTATYYCARYYRPFFDYWGQGTTLTVSS
                       ^^^^^^^^^       ^^^^^^^                   ^^^^^^^^^^
```

Best Alignment of Dalotuzumab light CDRs chain to a sequence from OAS

```
DIVMTQSPLSLPVTPGEPASISCRSSQSIVHSNGNTYLQWYLQKPGQSPQLLIYKVSNRLYGVPDRFSGSGSGTDFTLKISRVEAEDVGVYYCFQGSHVPWTFGQGTKVEIK
|..|||.||||||..|..|||||||||||||||||||||.||||||||||.||||||||||.||||||||||||||||||||||||||.|||||||||||||||||.|||.|||
DVLMTQTPLSLPVSLGDHASISCRSSQSIVHSNGNTYLEWYLQKPGQSPKLLIYKVSNRLSGVPDRFSGSGSGTDFTLKISRVEAEDLGVYYCFQGSHVPWTFGGGTKLEIK
                        ^^^^^^^^^^^          ^^^                  ^^^^^^^^^
```

Best Alignment of Dalotuzumab CDR-H3 chain to a sequence from OAS

```
QVQLQESGPGLVKPSETLSLTCTVSGYSITGGYLWNWIRQPPGKGLEWIGYISYD-GTNNYKPSLKDRVTISRDTSKNQFSLKLSSVTAADTAVYYCARYGRVFFDYWGQGTLVTVSS
.||||.|||.||||.......|..|||.....|..||..|..||.|||||.|....|...|....|...|...|.|.......|.|.|..|.|||||||||||.||||||||..||||
EVQLQQSGPVLVKPGASVKMSCKASGYTF-TDYYMNWVKQSHGKSLEWIGVINPYNGGTSYNQKFKGKATLTVDKSSSTAYMELNSLTSEDSAVYYCARYGRVYFDYWGQGTTLTVSS
                       ^^^^^^^^^       ^^^^^^^^                   ^^^^^^^^^^
```

Therapeutic : Daratumumab

Best Alignment of Daratumumab heavy chain to a sequence from OAS

```
EVQLLESGGGLVQPGGSLRLSCAVSGFTFNSFAMSWVRQAPGKGLEWVSAISGSGGGTYYADSVKGRFTISRDNSKNTLYLQMNSLRAEDTAVYFCAKDKILWFGEPVFDYWGQGTLVTVSS
||||||||||||||||||||||||.|||||.|.||||||||||||||||||||||||.||||||||||||||||||||||||||||||||||||||||||.||||..|||||..||||||||||||||
EVQLLESGGGLVQPGGSLRLSCAASGFTFSSYAMSWVRQAPGKGLEWVSAISGSGGSTYYADSVKGRFTISRDNSKNTLYLQMNSLRAEDTAVYYCAKDASLWFGEGYFDYWGQGTLVTVSS
                       ^^^^^^^^        ^^^^^^^^                   ^^^^^^^^^^^^^^^
```

Best Alignment of Daratumumab light chain to a sequence from OAS

```
EIVLTQSPATLSLSPGERATLSCRASQSVSSYLAWYQQKPGQAPRLLIYDASNRATGIPARFSGSGSGTDFTLTISSLEPEDFAVYYCQQRSNWPPTFGQGTKVEIK
||||||||||||||||||||||||||||||||||||||||||||||||||||||||||||||||||||||||||||||||||||||||||||||||||||||||||||
EIVLTQSPATLSLSPGERATLSCRASQSVSSYLAWYQQKPGQAPRLLIYDASNRATGIPARFSGSGSGTDFTLTISSLEPEDFAVYYCQQRSNWPPTFGQGTKVEIK
                        ^^^^^^          ^^^                   ^^^^^^^^^
```

Best Alignment of Daratumumab heavy chain CDRs to a sequence from OAS

```
EVQLLESGGGLVQPGGSLRLSCAVSGFTFNSFAMSWVRQAPGKGLEWVSAISGSGGGTYYADSVKGRFTISRDNSKNTLYLQMNSLRAEDTAVYFCAKDKILWFGEPVFDYWGQGTLVTVSS
.............|||||||||.|||||.|.||||||||||||||||||||||||.|||||||||||||||||||||||||||||||||||||||||.|||||.||||||.|||||||||||||
-------------GGSLRLSCAASGFTFSSYAMSWVRQAPGKGLEWVSAISGSGGSTYYADSVKGRFTISRDNSKNTLYLQMNSLRAEDTAVYYCAKDKELWFGEPYFDYWGQGTLVTVSS
                       ^^^^^^^^        ^^^^^^^^                   ^^^^^^^^^^^^^^^
```

Best Alignment of Daratumumab light CDRs chain to a sequence from OAS

```
EIVLTQSPATLSLSPGERATLSCRASQSVSSYLAWYQQKPGQAPRLLIYDASNRATGIPARFSGSGSGTDFTLTISSLEPEDFAVYYCQQRSNWPPTFGQGTKVEIK
||||||||||||||||||||||||||.|||||||||||||||||||||.||||||||||||.|||||||||||||||||.||||||.|.|||||||||||||.||||..|||
EIVLTQSPATLSLSPGERATLSCRARQSVSSYLAWYQQKPGQAPRLLMYDASNRATGIPGRFSGSGSGTDFTLTINSLEPEDSAIYYCQQRSNWPPTLGQGTRLEIK
                        ^^^^^^          ^^^                   ^^^^^^^^^
```

Best Alignment of Daratumumab CDR-H3 chain to a sequence from OAS

```
EVQLLESGGGLVQPGGSLRLSCAVSGFTFNSFAMSWVRQAPGKGLEWVSAISGSGGGTYYADSVKGRFTISRDNSKNTLYLQMNSLRAEDTAVYFCAKDKILWFGEPVFDYWGQGTLVTVSS
.............|||||||||.|||||...|||||||.||||||||||||.|||..|||||||||.||||||||||||||||||||||||||||.|.|||||.||||||.||||||||||||||
-------------GGSLRLSCAASGFTFSNYAMSWVRQNPGKGLEWVSAIRGSGRRTYYADSVKARFTISRDNSKNTLYLQMNSLRAEDTAIYYCAKDKELWFGEPYFDYWGQGTLVTVSS
                       ^^^^^^^^        ^^^^^^^^                   ^^^^^^^^^^^^^^^
```

Therapeutic : Dectrekumab

Best Alignment of Dectrekumab heavy chain to a sequence from OAS

```
EVQLVESGGGVVQPGRSLRLSCAASGFTFSSYGMHWVRQAPGKGLEWVAIIWYDGSNKYYADSVKGRFTISRDNSKNTLYLQMNSLRAEDTAVYYCARLWFGD---LDAFDIWGQGTMVTVSS
.|||||||||||||||||||||||||||||||||||||||||||||||.|||||||||||||||||||||||||||||||||||||||||||||||||.|.|.....|||||||||||||||
-VQLVESGGGVVQPGRSLRLSCAASGFTFSSYGMHWVRQAPGKGLEWVAVIWYDGSNKYYADSVKGRFTISRDNSKNTLYLQMNSLRAEDTAVYYCARDWGGRVATQDAFDIWGQGTMVTVSS
                       ^^^^^^^^        ^^^^^^^^                   ^^^^^^^^^^^^^^^^
```

Best Alignment of Dectrekumab light chain to a sequence from OAS

```
EIVLTQSPATLSLSPGERAILSCRAGQSVSSYLVWYQQKPGQAPRLLIYDASNRATGIPARFSGSGSGTDFTLTISSLEPEDFAVYYCQQRSSWPPVYTFGQGTKLEIK
||||||||||||||||||||.|||||.|||||||.|||||||||||||||||||||||||||||||||||||||||||||||||||||||||||||.||||||||||||||||
EIVLTQSPATLSLSPGERATLSCRASQSVSSYLAWYQQKPGQAPRLLIYDASNRATGIPARFSGSGSGTDFTLTISSLEPEDFAVYYCQQRSNWPPVYTFGQGTKLEIK
                        ^^^^^^          ^^^                   ^^^^^^^^^^^
```

Best Alignment of Dectrekumab heavy chain CDRs to a sequence from OAS

```
EVQLVESGGGVVQPGRSLRLSCAASGFTFSSYGMHWVRQAPGKGLEWVAIIWYDGSNKYYADSVKGRFTISRDNSKNTLYLQMNSLRAEDTAVYYCARLWFGDLDAFDIWGQGTMVTVSS
.............|.|||||||||||||||||||||||||||||||||||.|||||||||||||||||||||||||||||||||||||||||||||||||..||||||||||||||||
-------------GGSLRLSCAASGFTFSSYGMHWVRQAPGKGLEWVAVIWYDGSNKYYADSVKGRFTISRDNSKNTLYLQMNSLRAEDTAVYYCARLWFGEPDAFDIWGQGTMVTVSS
                       ^^^^^^^^        ^^^^^^^^                   ^^^^^^^^^^^^^
```

Best Alignment of Dectrekumab light CDRs chain to a sequence from OAS
EIVLTQSPATLSLSPGERAILSCRAGQSVSSYLVWYQQKPGQAPRLLIYDASNRATGIPARFSGSGSGTDFTLTISSLEPEDFAVYYCQQRSSWPPVYTFGQGTKLEIK
|||||||||||||||||||.|||||.|||||||.||||||||||||||||||||||||||||||||||||||||||||||||||||||||||.||||||||||||||||
EIVLTQSPATLSLSPGERATLSCRASQSVSSYLAWYQQKPGQAPRLLIYDASNRATGIPARFSGSGSGTDFTLTISSLEPEDFAVYYCQQRSNWPPVYTFGQGTKLEIK
                    ^^^^^^               ^^^                              ^^^^^^^^^^^

Best Alignment of Dectrekumab CDR-H3 chain to a sequence from OAS
EVQLVESGGGVVQPGRSLRLSCAASGFTFSSYGMHWVRQAPGKGLEWVAIIWYDGSNKYYADSVKGRFTISRDNSKNTLYLQMNSLRAEDTAVYYCARLWFGDLDAFDIWGQGTMVTVSS
..............|.||||||||||||||||||.|||||||||||||..|...|...||||||||||||||||.||.||||||||||||||||||||||||||||||||||||||||||
------------KSGGSLRLSCAASGFTFSSYGMSWVRQAPGKGLEWVSSISSSGGYIYYADSVKGRFTISRDNAKNSLYLQMNSLRAEDTAVYYCARLWFGDLDAFDIWGQGTMVTVSS
                   ^^^^^^^^               ^^^^^^^^                              ^^^^^^^^^^^^^

Therapeutic : Denosumab

Best Alignment of Denosumab heavy chain to a sequence from OAS
EVQLLESGGGLVQPGGSLRLSCAASGFTFSSYAMSWVRQAPGKGLEWVSGITGSGGSTYYADSVKGRFTISRDNSKNTLYLQMNSLRAEDTAVYYCAKDPGTTV----------------IMSWFDPWGQGTLVTVSS
|||||||||||||||||||||||||||||||||||||||||||||||||.|.||||||||||||||||||||||||||||||||||||||||||||||||.||...................||||||||||||||||
EVQLLESGGGLVQPGGSLRLSCAASGFTFSSYAMSWVRQAPGKGLEWVSAISGSGGSTYYADSVKGRFTISRDNSKNTLYLQMNSLRAEDTAVYYCAKDPFTTPLSRLSTIAVAGMLGDLAESWFDPWGQGTLVTVSS
                   ^^^^^^^^               ^^^^^^^^                              ^^^^^^^^^^^^^^^^^^^^^^^^^^^^^^

Best Alignment of Denosumab light chain to a sequence from OAS
EIVLTQSPGTLSLSPGERATLSCRASQSVRGRYLAWYQQKPGQAPRLLIYGASSRATGIPDRFSGSGSGTDFTLTISRLEPEDFAVFYCQQYGSSPRTFGQGTKVEIK
|||||||||||||||||||||||||||||.||||||||||||||||||||||||||||||||||||||||||||||||||||||||.|||||||||||||||||||||
EIVLTQSPGTLSLSPGERATLSCRASQSV-GRYLAWYQQKPGQAPRLLIYGASSRATGIPDRFSGSGSGTDFTLTISRLEPEDFAVYYCQQYGSSPRTFGQGTKVEIK
                    ^^^^^^^               ^^^                              ^^^^^^^^^

Best Alignment of Denosumab heavy chain CDRs to a sequence from OAS
EVQLLESGGGLVQPGGSLRLSCAASGFTFSSYAMSWVRQAPGKGLEWVSGITGSGGSTYYADSVKGRFTISRDNSKNTLYLQMNSLRAEDTAVYYCAKDPGTTVIMSWFDPWGQGTLVTVSS
..............|.||..|||||||||||||||||||||||||||||.|.||||||||||||||||||||||||||||||||||||||||||||||||||||...|||||||||||||||
--------------GESLKISCAASGFTFSSYAMSWVRQAPGKGLEWVSAISGSGGSTYYADSVKGRFTISRDNSKNTLYLQMNSLRAEDTAVYYCAKDPGTTVTTGWFDPWGQGTLVTVSS
                   ^^^^^^^^               ^^^^^^^^                              ^^^^^^^^^^^^^^^

Best Alignment of Denosumab light CDRs chain to a sequence from OAS
EIVLTQSPGTLSLSPGERATLSCRASQSVRGRYLAWYQQKPGQAPRLLIYGASSRATGIPDRFSGSGSGTDFTLTISRLEPEDFAVFYCQQYGSSPRTFGQGTKVEIK
||||||||||||||||||||.|.|||||||||||||||||||||||||||||||||||||||||||||||||||||||||||||||.|||||||||||||||||||||
EIVLTQSPGTLSLSPGERATFSGRASQSVRGRYLAWYQQKPGQAPRLLIYGASSRATGIPDRFSGSGSGTDFTLTISRLEPEDFAVYYCQQYGSSPRTFGQGTKVEIK
                    ^^^^^^^               ^^^                              ^^^^^^^^^

Best Alignment of Denosumab CDR-H3 chain to a sequence from OAS
EVQLLESGGGLVQPGGSLRLSCAASGFTFSSYAMSWVRQAPGKGLEWVSGITGSGGSTYYADSVKGRFTISRDNSKNTLYLQMNSLRAEDTAVYYCAKDPGTTVIMSWFDPWGQGTLVTVSS
..............|.||..|||||||||||||||||||||||||||||.|.||||||||||||||||||||||||||||||||||||||||||||||||||||...|||||||||||||||
--------------GESLKISCAASGFTFSSYAMSWVRQAPGKGLEWVSAISGSGGSTYYADSVKGRFTISRDNSKNTLYLQMNSLRAEDTAVYYCAKDPGTTVTTGWFDPWGQGTLVTVSS
                   ^^^^^^^^               ^^^^^^^^                              ^^^^^^^^^^^^^^^

Therapeutic : Depatuxizumab

Best Alignment of Depatuxizumab heavy chain to a sequence from OAS
QVQLQESGPGLVKPSQTLSLTCTVSGYSIS-SDFAWNWIRQPPGKGLEWMGYISYSGNTRYQPSLKSRITISRDTSKNQFFLKLNSVTAADTATYYCVTAGR-----------GFPYWGQGTLVTVSS
||||||||||||||||||||||||||.|||..|..|.||||||||||||.|||.|||.|.|.||||||.|||.|||||||.||||||||||||.||||.||.............|.||||||||||||
QVQLQESGPGLVKPSQTLSLTCTVSGGSISSGDYYWSWIRQPPGKGLEWIGYIYYSGSTYYNPSLKSRVTISIDTSKNQFSLKLNSVTAADTAVYYCVRAGVENFYDSSASGGCFDYWGQGTLVTVSS
                   ^^^^^^^^^^               ^^^^^^^                              ^^^^^^^^^^^^^^^^^^^^

Best Alignment of Depatuxizumab light chain to a sequence from OAS
DIQMTQSPSSMSVSVGDRVTITCHSSQDINSNIGWLQQKPGKSFKGLIYHGTNLDDGVPSRFSGSGSGTDYTLTISSLQPEDFATYYCVQYAQFPWTFGGGTKLEIK
||.|||||||||||.||.|.||||.||||.||||||||||||||||||||||||.|||||||||||||.||.||||||..||||.||||||||||||||||||||||
DILMTQSPSSMSVSLGDTVSITCHASQDISSNIGWLQQKPGKSFKGLIYHGTNLEDGVPSRFSGSGSGADYSLTISSLESEDFADYYCVQYAQFPWTFGGGTKLEIK
                    ^^^^^^               ^^^                              ^^^^^^^^^

Best Alignment of Depatuxizumab heavy chain CDRs to a sequence from OAS
QVQLQESGPGLVKPSQTLSLTCTVSGYSISSDFAWNWIRQPPGKGLEWMGYISYSGNTRYQPSLKSRITISRDTSKNQFFLKLNSVTAADTATYYCVTAGRGFPYWGQGTLVTVSS
.||||.||||||||||.|||||||.||||.||.|||||.|.||..|||||||||||.|.|.|.|||||||.|.||||||||||.|||||..|||||||.|.||||.|||||||||||.
EVQLQQSGPGLVKPSQSLSLTCTVTGYSITSDYAWNWIWQFPGNKLEWMGYISYSGSTSYNPSLKSRISITRDTSKNQFFLQLNSVTTEDTATYYCATLGRGFAYWGQGTLVTVSA
                   ^^^^^^^^^               ^^^^^^^                              ^^^^^^^^^

Best Alignment of Depatuxizumab light CDRs chain to a sequence from OAS
DIQMTQSPSSMSVSVGDRVTITCHSSQDINSNIGWLQQKPGKSFKGLIYHGTNLDDGVPSRFSGSGSGTDYTLTISSLQPEDFATYYCVQYAQFPWTFGGGTKLEIK
||.|.|.|||||||..|...||.|.||.||||||||||||||||||||||||||.|||||||||||||.||.||||||..||||.||||||||||||||||||||||
DILMSQPPSSMSVSLEDTASITFHASQGINSNIGWLQQKPGKSFKGLIYHGTNLEDGVPSRFSGSGSGADYSLTISSLESEDFADYYCVQYAQFPWTFGGGTKLEIK
                    ^^^^^^               ^^^                              ^^^^^^^^^

Best Alignment of Depatuxizumab CDR-H3 chain to a sequence from OAS
QVQLQESGPGLVKPSQTLSLTCTVSGYSISSDFAWNWIRQPPGKGLEWMGYISYSGNTRYQPSLKSRITISRDTSKNQFFLKLNSVTAADTATYYCVTAGRGFPYWGQGTLVTVSS
.||||||||.||||||.|||||||.||||.|...|.||||.||..|||||||.|.|||.|.|||||||.|.||||.|||||.|||||..|||||||.|||||||||.||||||||.
EVQLQESGPDLVKPSQSLSLTCTVTGYSITSGYSWHWIRQFPGNKLEWMGYIHYGGNTNYNPSLKSRISITRDTSRNQFFLQLNSVTTEDTATYYCATAGRGFPYWCQGTLVTVSA
                   ^^^^^^^^^               ^^^^^^^                              ^^^^^^^^^

Therapeutic : Derlotuximab

Best Alignment of Derlotuximab heavy chain to a sequence from OAS
QVQLKESGPGLVAPSQSLSITCTVSGFSLTDYGVRWIRQPPGKGLEWLGVIWGGGSTYYNSALKSRLSISKDNSKSQVFLKMNSLQTDDTAMYYCAKEKRRGYYYAMDYWGQGTSVTVSS
||||||||||||||||||||||||||||||||||.||||||||||||||||||||||||||||||||||||||||||||||||||||||||||||||..|||.|||||||||||||||||
QVQLKESGPGLVAPSQSLSITCTVSGFSLTDYGVSWIRQPPGKGLEWLGVIWGGGSTYYNSALKSRLSISKDNSKSQVFLKMNSLQTDDTAMYYCAKHVRRGDYYAMDYWGQGTSVTVSS
                   ^^^^^^^^               ^^^^^^^                              ^^^^^^^^^^^^^^

Best Alignment of Derlotuximab light chain to a sequence from OAS
ENVLTQSPAIMSASPGEKVTMTCRASSSVSSSYLHWYQQKSGASPKLWIYSTSNLASGVPARFSGSGSGTSYSLTISSVEAEDAATYYCQQYSGYPLTFGGGTKLEIK
||||||||||||||||||||||||||||||||||||||||||||||||||||||||||||||||||||||||||||||||||||||||||||||||||||||||||||
ENVLTQSPAIMSASPGEKVTMTCRASSSVSSSYLHWYQQKSGASPKLWIYSTSNLASGVPARFSGSGSGTSYSLTISSVEAEDAATYYCQQYSGYPLTFGGGTKLEIK
                    ^^^^^^^               ^^^                              ^^^^^^^^^

Best Alignment of Derlotuximab heavy chain CDRs to a sequence from OAS
QVQLKESGPGLVAPSQSLSITCTVSGFSLTDYGVRWIRQPPGKGLEWLGVIWGGGSTYYNSALKSRLSISKDNSKSQVFLKMNSLQTDDTAMYYCAKEKRRGYYYAMDYWGQGTSVTVSS
.|||..||||||||||||||||||||||||.|||.|.||||||||||||||||.|||.|.|||.||||.||||||||||||.|||||||||.||||||.|||||||||||||||||||||
-VQLQQSGPGLVAPSQSLSITCTVSGFSLTSYGVSWVRQPPGKGLEWLGVIWGDGSTNYHSALISRLSFSKDNSKSQVFLKLNSLQTDDTATYYCAKEVRRGYYYAMDYWGQGTSVTVSS
                   ^^^^^^^^               ^^^^^^^                              ^^^^^^^^^^^^^^

Best Alignment of Derlotuximab light CDRs chain to a sequence from OAS
ENVLTQSPAIMSASPGEKVTMTCRASSSVSSSYLHWYQQKSGASPKLWIYSTSNLASGVPARFSGSGSGTSYSLTISSVEAEDAATYYCQQYSGYPLTFGGGTKLEIK
|||.|.||||||||||||||||||||||||||||||||..||||||||||||||||||||||||||||||||||||||||||||||||||||||||||||.|||||||
ENVHTPSPAIMSASPGEKVTMTCRASSSVSSSYLHWYQENSGASPKLWIYSTSNLASGVPARFSGSGSGTSYSLTISSVEAEDAATYYCQQYSGYPLTFGSGTKLEIK
                    ^^^^^^^               ^^^                              ^^^^^^^^^
Best Alignment of Derlotuximab CDR-H3 chain to a sequence from OAS
QVQLKESGPGLVAPSQSLSITCTVSGFSLTDYGVRWIRQPPGKGLEWLGVIWGGGSTYYNSALKSRLSISKDNSKSQVFLKMNSLQTDDTAMYYCAKEKRRGYYYAMDYWGQGTSVTVSS
.|||..||||||||||||||||||||||||.|||.|.||||||||||||||||.|||.|.|||.||||.||||||||||||.|||||||||.||||||.|||||||||||||||||||||
-VQLQQSGPGLVAPSQSLSITCTVSGFSLTSYGVSWVRQPPGKGLEWLGVIWGDGSTNYHSALISRLSFSKDNSKSQVFLKLNSLQTDDTATYYCAKEVRRGYYYAMDYWGQGTSVTVSS
                   ^^^^^^^^               ^^^^^^^                              ^^^^^^^^^^^^^^

Therapeutic : Dinutuximab

Best Alignment of Dinutuximab heavy chain to a sequence from OAS
EVQLLQSGPELEKPGASVMISCKASGSSFTGYNMNWVRQNIGKSLEWIGAIDPYYGGTSYNQKFKGRATLTVDKSSSTAYMHLKSLTSEDSAVYYCVSG-------MEYWGQGTSVTVSS
||||.|||||||||||||.|||||||.||||||||||.|..||||||||.||||||||||||||||.||||||||||||||.||||||||||||||.||.......|.||||||||||||
EVQLQQSGPELEKPGASVKISCKASGYSFTGYNMNWVKQSNGKSLEWIGNIDPYYGGTSYNQKFKGKATLTVDKSSSTAYMQLKSLTSEDSAVYYCASGGKPDYYAMDYWGQGTSVTVSS
^^^^^^^^ ^^^^^^^^ ^^^^^^^^^^^^^

Best Alignment of Dinutuximab light chain to a sequence from OAS
EIVMTQSPATLSVSPGERATLSCRSSQSLVHRNGNTYLHWYLQKPGQSPKLLIHKVSNRFSGVPDRFSGSGSGTDFTLKISRVEAEDLGVYFCSQSTHVPPLTFGAGTKLELK
.|||||||..|.||.|..|..||||||||||.|||||||||||||||||||||.|||||||||||||||||||||||||||||||||||||||||||||||||||||||||||||||
DIVMTQSPLSLPVSLGDQASISCRSSQSLVHSNGNTYLHWYLQKPGQSPKLLIYKVSNRFSGVPDRFSGSGSGTDFTLKISRVEAEDLGVYFCSQSTHVPPLTFGAGTKLELK
^^^^^^^^^^^ ^^^ ^^^^^^^^^^

Best Alignment of Dinutuximab heavy chain CDRs to a sequence from OAS
EVQLLQSGPELEKPGASVMISCKASGSSFTGYNMNWVRQNIGKSLEWIGAIDPYYGGTSYNQKFKGRATLTVDKSSSTAYMHLKSLTSEDSAVYYCVSGMEYWGQGTSVTVSS
||||.|||||||||||||.|||||||.||||||||||.|..||||||||.||||||||||||||||.||||||||||||||.|||||||||||||||.||.||||||||||.|
EVQLQQSGPELEKPGASVKISCKASGYSFTGYNMNWVKQSNGKSLEWIGNIDPYYGGTSYNQKFKGKATLTVDKSSSTAYMQLKSLTSEDSAVYYCVMGMDYWGQGTSVTVAS
^^^^^^^^ ^^^^^^^^ ^^^^^^

Best Alignment of Dinutuximab light CDRs chain to a sequence from OAS
EIVMTQSPATLSVSPGERATLSCRSSQSLVHRNGNTYLHWYLQKPGQSPKLLIHKVSNRFSGVPDRFSGSGSGTDFTLKISRVEAEDLGVYFCSQSTHVPPLTFGAGTKLELK
..|||.|..|.||.|..|..||||||||||.|||||||||||||||||||||.|||||||||||||||||||||||||||||||||||||||||||||||||||.|||||.|
DVVMTQTPLSLPVSLGDQASISCRSSQSLVHSNGNTYLHWYLQKPGQSPKLLIYKVSNRFSGVPDRFSGSGSGTDFTLKISRVEAEDLGVYFCSQSTHVPPLTFGSGTKLEIK
^^^^^^^^^^^ ^^^ ^^^^^^^^^^

Best Alignment of Dinutuximab CDR-H3 chain to a sequence from OAS
EVQLLQSGPELEKPGASVMISCKASGSSFTGYNMNWVRQNIGKSLEWIGAIDPYYGGTSYNQKFKGRATLTVDKSSSTAYMHLKSLTSEDSAVYYCVSGMEYWGQGTSVTVSS
||||..||..|.|||.|...||.|||..|..|.|.||||...|.|||...|......|.|....|||.|...|......|.....|.|||.|.||.|.|||||||.||||||.
EVQLVESGGGLVKPGGSLTLSCAASGFTFSSYAMSWVRQTPEKRLEWVVSISDCGNYTYYPDNVKGRFTSSRDNDKNNLYLQMSHLKSEDTAMYYGVRGMEYWGQETSVTVST
^^^^^^^^ ^^^^^^^^ ^^^^^^

Therapeutic : Domagrozumab

Best Alignment of Domagrozumab heavy chain to a sequence from OAS
EVQLLESGGGLVQPGGSLRLSCAASGFTFSSYAMSWVRQAPGKGLEWVSTISSGGSYTSYPDSVKGRFTISRDNSKNTLYLQMNSLRAEDTAVYYCAKQDY----AMNYWGQGTLVTVSS
|||||||||||||||||||||||||||||||||||||||||||||||||.|.||||.|.|.|||||||||||||||||||||||||||||||||||||||.....|..||||||||||||
EVQLLESGGGLVQPGGSLRLSCAASGFTFSSYAMSWVRQAPGKGLEWVSVIYSGGSSTYYADSVKGRFTISRDNSKNTLYLQMNSLRAEDTAVYYCAKQDGYKERAFDYWGQGTLVTVSS
^^^^^^^^ ^^^^^^^^ ^^^^^^^^^^^^^

Best Alignment of Domagrozumab light chain to a sequence from OAS
DIQMTQSPSSLSASVGDRVTITCKASQDVSTAVAWYQQKPGKAPKLLIYSASYRYTGVPSRFSGSGSGTDFTLTISSLQPEDFATYYCQQHYSTPWTFGGGTKVEIK
|||||||||||||||||||||||.||||.||..||||||||||||||||.||....||||||||||||||||||||||||||||||||||.||||||||.|||||||
DIQMTQSPSSLSASVGDRVTITCRASQDISTSLAWYQQKPGKAPKLLIYAASSLQSGVPSRFSGSGSGTDFTLTISSLQPEDFATYYCQQSYSTPWTFGQGTKVEIK
^^^^^^ ^^^ ^^^^^^^^^

Best Alignment of Domagrozumab heavy chain CDRs to a sequence from OAS
EVQLLESGGGLVQPGGSLRLSCAASGFTFSSYAMSWVRQAPGKGLEWVSTISSGGSYTSYPDSVKGRFTISRDNSKNTLYLQMNSLRAEDTAVYYCAKQDYAMNYWGQGTLVTVSS
||.|.|||||||.|||||.||||||||||||||||||||.|.|.||||.|||||||||.|||||||||||||||.||||||||.|||.||||.||||.|||||.||||||.|||||
EVKLVESGGGLVKPGGSLKLSCAASGFTFSSYAMSWVRQTPEKRLEWVATISSGGSYTYYPDSVKGRFTISRDNAKNTLYLQMSSLRSEDTAMYYCARQDYAMDYWGQGTSVTVSS
^^^^^^^^ ^^^^^^^^ ^^^^^^^^^

Best Alignment of Domagrozumab light CDRs chain to a sequence from OAS
DIQMTQSPSSLSASVGDRVTITCKASQDVSTAVAWYQQKPGKAPKLLIYSASYRYTGVPSRFSGSGSGTDFTLTISSLQPEDFATYYCQQHYSTPWTFGGGTKVEIK
||.||||....|.||||||.|||||||||||||||||||||..||||||||||||||||.||.|||||||||.||||.|.||.|.||||||||||||||||||.|||
DIVMTQSHKFMSTSVGDRVSITCKASQDVSTAVAWYQQKPGHSPKLLIYSASYRYTGVPDRFTGSGSGTDFTFTISSVQAEDLAVYYCQQHYSTPWTFGGGTKLEIK
^^^^^^ ^^^ ^^^^^^^^^

Best Alignment of Domagrozumab CDR-H3 chain to a sequence from OAS
EVQLLESGGGLVQPGGSLRLSCAASGFTFSSYAMSWVRQAPGKGLEWVSTISSGGSYTSYPDSVKGRFTISRDNSKNTLYLQMNSLRAEDTAVYYCAKQDYAMNYWGQGTLVTVSS
|||..||||.||.|||||.|||||||||||||.||||||.|.|.||||.|||||||||.|||||||||||||||.||||||||.||..||||.||||||||||.||||||.|||||
EVQRVESGGDLVKPGGSLKLSCAASGFTFSSYGMSWVRQTPDKRLEWVATISSGGSYTYYPDSVKGRFTISRDNAKNTLYLQMSSLKSEDTAMYYCAKQDYAMDYWGQGTSVTVSS
^^^^^^^^ ^^^^^^^^ ^^^^^^^^^

Therapeutic : Drozitumab

Best Alignment of Drozitumab heavy chain to a sequence from OAS
EVQLVQSGGGVERPGGSLRLSCAASGFTFDDYAMSWVRQAPGKGLEWVSGINWQGGSTGYADSVKGRVTISRDNAKNSLYLQMNSLRAEDTAVYYCAKILGAG--RGWYFDYWGKGTTVTVSS
|||||.|||||.||||||||||||||||||||.||||||||||||||||||||.|||||||||||||.||||||||||||||||||||||||.|||||...||..||.||||||.||.|||||
EVQLVESGGGVVRPGGSLRLSCAASGFTFDDYGMSWVRQAPGKGLEWVSGINWNGGSTGYADSVKGRFTISRDNAKNSLYLQMNSLRAEDTALYYCAKYTVAGTSRGGYFDYWGQGTMVTVSS
^^^^^^^^ ^^^^^^^^ ^^^^^^^^^^^^^^^^

Best Alignment of Drozitumab light chain to a sequence from OAS
-SELTQDPAVSVALGQTVRITCSGDSLRSYYASWYQQKPGQAPVLVIYGANNRPSGIPDRFSGSSSGNTASLTITGAQAEDEADYYCNSADSSGNHVVFGGGTKLTVL
.|||||||||||||||||||||.||||||||||||||||||||||||||.|||||||||||||||||||||||||||||||||||||||.||||||||||||||||||
SSELTQDPAVSVALGQTVRITCQGDSLRSYYASWYQQKPGQAPVLVIYGKNNRPSGIPDRFSGSSSGNTASLTITGAQAEDEADYYCNSRDSSGNHVVFGGGTKLTVL
^^^^^^ ^^^ ^^^^^^^^^^^

Best Alignment of Drozitumab heavy chain CDRs to a sequence from OAS
EVQLVQSGGGVERPGGSLRLSCAASGFTFDDYAMSWVRQAPGKGLEWVSGINWQGGSTGYADSVKGRVTISRDNAKNSLYLQMNSLRAEDTAVYYCAKILGAGRGWYFDYWGKGTTVTVSS
................||||||||||||||||.|.||||||||||||||.|.|.||||.||||||||.||||||.||||||||||||.||||.|||||.||||||..|||||.||.|||||
----------------SLRLSCAASGFTFDDYTMHWVRQAPGKGLEWVSLISWDGGSTYYADSVKGRFTISRDNSKNSLYLQMNSLRTEDTALYYCAKDLGAGRGTAFDYWGQGTLVTVSS
^^^^^^^^ ^^^^^^^^ ^^^^^^^^^^^^^^

Best Alignment of Drozitumab light CDRs chain to a sequence from OAS
-SELTQDPAVSVALGQTVRITCSGDSLRSYYASWYQQKPGQAPVLVIYGANNRPSGIPDRFSGSSSGNTASLTITGAQAEDEADYYCNSADSSGNHVVFGGGTKLTVL
.|||||||||||||||||||||.||||||||||||||||||||.|||||.|||||||||||||||||||||||||||||||||||||||.||||||||||||||||||
SSELTQDPAVSVALGQTVRITCQGDSLRSYYASWYQQKPGQAPLLVIYGKNNRPSGIPDRFSGSSSGNTASLTITGAQAEDEADYYCNSRDSSGNHVVFGGGTKLTVL
^^^^^^ ^^^ ^^^^^^^^^^^

Best Alignment of Drozitumab CDR-H3 chain to a sequence from OAS
EVQLVQSGGGVERPGGSLRLSCAASGFTFDDYAMSWVRQAPGKGLEWVSGINWQGGSTGYADSVKGRVTISRDNAKNSLYLQMNSLRAEDTAVYYCAKILGAGRGWYFDYWGKGTTVTVSS
................|||||||||||||..|||.||||||||||||||.|...|..|.||||||||.||||||.|..||||||||||.|||||||||.||||||.||||||.||.|||||
----------------SLRLSCAASGFTFRSYAMNWVRQAPGKGLEWVSAISGSGVRTYYADSVKGRFTISRDNSKKTLYLQMNSLRADDTAVYYCAKDLGAGRGYYFDYWGQGTLVTVSS
^^^^^^^^ ^^^^^^^^ ^^^^^^^^^^^^^^

Therapeutic : Duligotuzumab

Best Alignment of Duligotuzumab heavy chain to a sequence from OAS
EVQLVESGGGLVQPGGSLRLSCAASGFTLSGDWIHWVRQAPGKGLEWVGEISAAGGYTDYADSVKGRFTISADTSKNTAYLQMNSLRAEDTAVYYCARESRVS-----FEAAMDYWGQGTLVTVSS
.|||..|||||||||||||||||||||..|...|||||||||||||||..||...|||.|||||||||||||||||||||||||||||||||||||||..|.|........|||||||||||||||
QVQLLKSGGGLVQPGGSLRLSCAASGFNFSSSSIHWVRQAPGKGLEWVAYISSSYGYTYYADSVKGRFTISADTSKNTAYLQMNSLRAEDTAVYYCARTVRGSKKPYFSGWAMDYWGQGTLVTVSS
^^^^^^^^ ^^^^^^^^ ^^^^^^^^^^^^^^^^^^

Best Alignment of Duligotuzumab light chain to a sequence from OAS
```
DIQMTQSPSSLSASVGDRVTITCRASQNIATDVAWYQQKPGKAPKLLIYSASFLYSGVPSRFSGSGSGTDFTLTISSLQPEDFATYYCQQSEPEPYTFGQGTKVEIK
|||||||||||||||||||||||||||||||...|||||||||||||||.||.|.||||||||||||||||||||||||||||||||||||...|.|||||||||||
DIQMTQSPSSLSASVGDRVTITCRASQNIATFLNWYQQKPGKAPKLLIYAASSLQSGVPSRFSGSGSGTDFTLTISSLQPEDFATYYCQQSYSTPWTFGQGTKVEIK
                          ^^^^^^                 ^^^                                    ^^^^^^^^^
```

Best Alignment of Duligotuzumab heavy chain CDRs to a sequence from OAS
```
EVQLVESGGGLVQPGGSLRLSCAASGFTLSGDWIHWVRQAPGKGLEWVGEISAAGGYTDYADSVKGRFTISADTSKNTAYLQMNSLRAEDTAVYYCARESRVSFEAAMDYWGQGTLVTVSS
|||.||||||||.|||||.|||||||||.|.....||||.|.|.||||..||..|.||.|.||||||||||.|..||..||||.||..||||.|||||||.|.|..|||||||||.|||||
EVQRVESGGGLVKPGGSLKLSCAASGFTFSDYYMYWVRQTPEKRLEWVATISDGGSYTYYPDSVKGRFTISRDNAKNNLYLQMSSLKSEDTAMYYCARESMVTFRYAMDYWGQGTSVTVSS
                         ^^^^^^^^                 ^^^^^^^^                                      ^^^^^^^^^^^^^^
```

Best Alignment of Duligotuzumab light CDRs chain to a sequence from OAS
```
DIQMTQSPSSLSASVGDRVTITCRASQNIATDVAWYQQKPGKAPKLLIYSASFLYSGVPSRFSGSGSGTDFTLTISSLQPEDFATYYCQQSEPEPYTFGQGTKVEIK
||.|||||.||||||||||.|||||||||.||..||||||||||||||||||.|..|||||||||||||||||||||.||..|.|||||...|||||||||.|||
DIRMTQSPPSLSASVGDRVIITCRASQNINTDLNWYQQKPGKAPKLLIYSASNLEGGVPSRFSGSGSGTDFTLTISSLQLEDLSTFYCQQSYSAPYTFGQGTKLEIK
                          ^^^^^^                 ^^^                                    ^^^^^^^^^
```

Best Alignment of Duligotuzumab CDR-H3 chain to a sequence from OAS
```
EVQLVESGGGLVQPGGSLRLSCAASGFTLSGDWIHWVRQAPGKGLEWVGEISAAGGYTDYADSVKGRFTISADTSKNTAYLQMNSLRAEDTAVYYCARESRVSFEAAMDYWGQGTLVTVSS
.|||...|..||.||.|..|||.|||.|....|..||.|.||.||||.|.|......|.|....|...|...|.|..|||.|..||..||.||||||||.||||..|||||||||.|||||
-VQLQQPGAELVRPGSSVKLSCKASGYTFTSYWMDWVKQRPGQGLEWIGNIYPSDSETHYNQKFKDKATLTVDKSSSTAYMQLSSLTSEDSAVYYCARENRVSFFYAMDYWGQGTSVTVSS
                         ^^^^^^^^                 ^^^^^^^^                                      ^^^^^^^^^^^^^^
```

Therapeutic : Dupilumab

Best Alignment of Dupilumab heavy chain to a sequence from OAS
```
EVQLVESGGGLEQPGGSLRLSCAGSGFTFRDYAMTWVRQAPGKGLEWVSSISGSGGNTYYADSVKGRFTISRDNSKNTLYLQMNSLRAEDTAVYYCAKDRLSITI--RPRYYGLDVWGQGTTVTVSS
||||.||||||.||||||||||||.|||||..|||||||||||||||||||.||||||||||||||||||||||||||||||||||||||||||||||||..........|||.|||||||||||||
EVQLLESGGGLVQPGGSLRLSCAASGFTFSSYAMTWVRQAPGKGLEWVSAISGSGGNTYYADSVKGRFTISRDNSKNTLYLQMNSLRAEDTAVYYCAKDRSITMVRGQSGYYGMDVWGQGTTVTVSS
                         ^^^^^^^^                 ^^^^^^^^                                      ^^^^^^^^^^^^^^^^^^^^
```

Best Alignment of Dupilumab light chain to a sequence from OAS
```
DIVMTQSPLSLPVTPGEPASISCRSSQSLLYSIGYNYLDWYLQKSGQSPQLLIYLGSNRASGVPDRFSGSGSGTDFTLKISRVEAEDVGFYYCMQALQTPYTFGQGTKLEIK
||||||||||||||||||||||||||||||.|.||||||||||||||||||||||||||||||||||||||||||||||||||||||||.||||||||||||||||||||||
DIVMTQSPLSLPVTPGEPASISCRSSQSLLHSNGYNYLDWYLQKSGQSPQLLIYLGSNRASGVPDRFSGSGSGTDFTLKISRVEAEDVGVYYCMQALQTPYTFGQGTKLEIK
                          ^^^^^^^^^^^                 ^^^                                    ^^^^^^^^^
```

Best Alignment of Dupilumab heavy chain CDRs to a sequence from OAS
```
EVQLVESGGGLEQPGGSLRLSCAGSGFTFRDYAMTWVRQAPGKGLEWVSSISGSGGNTYYADSVKGRFTISRDNSKNTLYLQMNSLRAEDTAVYYCAKDRLSITIRPRYYGLDVWGQGTTVTVSS
..............|||||||||.|||||..|||.||||||||||||||.||||||.|||||||||||||||||||||||||||||||||||||||||||.||||...|||.||||||||||||.
--------------GGSLRLSCAASGFTFSSYAMSWVRQAPGKGLEWVSAISGSGGSTYYADSVKGRFTISRDNSKNTLYLQMNSLRAEDTAVYYCAKDRGSITIWDYYYGMDVWGQGTTVTVS-
                         ^^^^^^^^                 ^^^^^^^^                                      ^^^^^^^^^^^^^^^^^^
```

Best Alignment of Dupilumab light CDRs chain to a sequence from OAS
```
DIVMTQSPLSLPVTPGEPASISCRSSQSLLYSIGYNYLDWYLQKSGQSPQLLIYLGSNRASGVPDRFSGSGSGTDFTLKISRVEAEDVGFYYCMQALQTPYTFGQGTKLEIK
||||||.|||||||||||||||||||||||||.|||||||||||.||||||||||||.|||||||||||||.||||||.||||||||||.||||||||||||||||||||||
DIVMTQPPLSLPVTPGEPASISCRSSQSLLYSNGYNYLDWYLQKPGQSPQLLIYLGSTRASGVPDRFSGSGTGTDFTLNISRVEAEDVGMYYCMQALQTPYTFGQGTKLEIK
                          ^^^^^^^^^^^                 ^^^                                    ^^^^^^^^^
```

Best Alignment of Dupilumab CDR-H3 chain to a sequence from OAS
```
EVQLVESGGGLEQPGGSLRLSCAGSGFTFRDYAMTWVRQAPGKGLEWVSSISGSGGNTYYADSVKGRFTISRDNSKNTLYLQMNSLRAEDTAVYYCAKDRLSITIRPRYYGLDVWGQGTTVTVSS
..............|||||||||.|||||..|||.||||||||||||||.||||||.|||||||||||||||||||||||||||||||||||||||||||.||||...|||.||||||||||||.
--------------GGSLRLSCAASGFTFSSYAMSWVRQAPGKGLEWVSAISGSGGSTYYADSVKGRFTISRDNSKNTLYLQMNSLRAEDTAVYYCAKDRGSITIWDYYYGMDVWGQGTTVTVS-
                         ^^^^^^^^                 ^^^^^^^^                                      ^^^^^^^^^^^^^^^^^^
```

Therapeutic : Durvalumab

Best Alignment of Durvalumab heavy chain to a sequence from OAS
```
EVQLVESGGGLVQPGGSLRLSCAASGFTFSRYWMSWVRQAPGKGLEWVANIKQDGSEKYYVDSVKGRFTISRDNAKNSLYLQMNSLRAEDTAVYYCAREGGWFGELAFDYWGQGTLVTVSS
||||||||||||||||||||||||||||||.|||||||||||||||||||||||||||||||||||||||||||||||||||||||||||||||||||.|.|||||.||||||||||||||
EVQLVESGGGLVQPGGSLRLSCAASGFTFSSYWMSWVRQAPGKGLEWVANIKQDGSEKYYVDSVKGRFTISRDNAKNSLYLQMNSLRAEDTAVYYCARSGLWFGELPFDYWGQGTLVTVSS
                         ^^^^^^^^                 ^^^^^^^^                                      ^^^^^^^^^^^^^^
```

Best Alignment of Durvalumab light chain to a sequence from OAS
```
EIVLTQSPGTLSLSPGERATLSCRASQRVSSSYLAWYQQKPGQAPRLLIYDASSRATGIPDRFSGSGSGTDFTLTISRLEPEDFAVYYCQQYGS-LPWTFGQGTKVEIK
|||||||||||||||||||||||||||||||.||||||||||||||||||||||||||||||||||||||||||||||||||||||||||||||.||||||||||||||
EIVLTQSPGTLSLSPGERATLSCRASQRVSSNYLAWYQQKPGQAPRLLIYDASSRATGIPDRFSGSGSGTDFTLTISRLEPEDFAVYYCQQYGSSLPWTFGQGTKVEIK
                          ^^^^^^^                 ^^^                                    ^^^^^^^^^^
```

Best Alignment of Durvalumab heavy chain CDRs to a sequence from OAS
```
EVQLVESGGGLVQPGGSLRLSCAASGFTFSRYWMSWVRQAPGKGLEWVANIKQDGSEKYYVDSVKGRFTISRDNAKNSLYLQMNSLRAEDTAVYYCAREGGWFGELAFDYWGQGTLVTVSS
................||||||||||||||.|||||||||||||||||||||||||||||||||||||||||||||||||||||||||||||||||||.|||||||.||||||||||||||
----------------SLRLSCAASGFTFSSYWMSWVRQAPGKGLEWVANIKQDGSEKYYVDSVKGRFTISRDNAKNSLYLQMNSLRAEDTAVYYCARDGGWFGELPFDYWGQGTLVTVSS
                         ^^^^^^^^                 ^^^^^^^^                                      ^^^^^^^^^^^^^^
```

Best Alignment of Durvalumab light CDRs chain to a sequence from OAS
```
EIVLTQSPGTLSLSPGERATLSCRASQRVSSSYLAWYQQKPGQAPRLLIYDASSRATGIPDRFSGSGSGTDFTLTISRLEPEDFAVYYCQQYGSLPWTFGQGTKVEIK
||||||||.||||||||||||||.||||||||||||||||||.|||||||||||||||||||||||||||||||||||||||||||||||||||.|||||||||||||
EIVLTQSPATLSLSPGERATLSCGASQRVSSSYLAWYQQKPGLAPRLLIYDASSRATGIPDRFSGSGSGTDFTLTISRLEPEDFAVYYCQQYGSSPWTFGQGTKVEIK
                          ^^^^^^^                 ^^^                                    ^^^^^^^^^
```

Best Alignment of Durvalumab CDR-H3 chain to a sequence from OAS
```
EVQLVESGGGLVQPGGSLRLSCAASGFTFSRYWMSWVRQAPGKGLEWVANIKQDGSEKYYVDSVKGRFTISRDNAKNSLYLQMNSLRAEDTAVYYCAREGGWFGELAFDYWGQGTLVTVSS
...............|||||||||||||||.|.|.||||||||||||||.|..|||.|||.||||||||||||||||||.||||||||||||.|||||||.|||||||||||||||||||.
---------------GSLRLSCAASGFTFSSYGMHWVRQAPGKGLEWVAVIWYDGSNKYYADSVKGRFTISRDNAKNSLFLQMNSLRAEDTAMYYCAREGLWFGELAFDYWGQGTLVTVS-
                         ^^^^^^^^                 ^^^^^^^^                                      ^^^^^^^^^^^^^^
```

Therapeutic : Dusigitumab

Best Alignment of Dusigitumab heavy chain to a sequence from OAS
```
QVQLVQSGAEVKKPGASVKVSCKASGYTFTSYDINWVRQATGQGLEWMGWMNPNSGNTGYAQKFQGRVTMTRNTSISTAYMELSSLRSEDTAVYYCARDPYYY-------------YYGMDVWGQGTTVTVSS
||||||||||||||||||||||||||||||||||||||||||||||||||||||||||||||||||||||||||||||||||||||||||||||||||||.||.............|||||||||||||||||
QVQLVQSGAEVKKPGASVKVSCKASGYTFTSYDINWVRQATGQGLEWMGWMNPNSGNTGYAQKFQGRVTMTRNTSISTAYMELSSLRSEDTAVYYCARDPSYYYGSGSYRPPLEYYYYGMDVWGQGTTVTVSS
                         ^^^^^^^^                 ^^^^^^^^                                      ^^^^^^^^^^^^^^^^^^^^^^^^^^
```

Best Alignment of Dusigitumab light chain to a sequence from OAS
```
QSVLTQPPSVSAAPGQKVTISCSGSSSNIENNHVSWYQQLPGTAPKLLIYDNNKRPSGIPDRFSGSKSGTSATLGITGLQTGDEADYYCETWDTSLSAGRVFGGGTKLTVL
|||||||||||||||||||||||||||||.||.||||||||||||||||||||||||||||||||||||||||||||||||||||||||.|||||||||||||||||||||
QSVLTQPPSVSAAPGQKVTISCSGSSSNIGNNYVSWYQQLPGTAPKLLIYDNNKRPSGIPDRFSGSKSGTSATLGITGLQTGDEADYYCGTWDTSLSAGRVFGGGTKLTVL
                         ^^^^^^^^                 ^^^                                    ^^^^^^^^^^^^
```

Best Alignment of Dusigitumab heavy chain CDRs to a sequence from OAS
```
QVQLVQSGAEVKKPGASVKVSCKASGYTFTSYDINWVRQATGQGLEWMGWMNPNSGNTGYAQKFQGRVTMTRNTSISTAYMELSSLRSEDTAVYYCARDPYYYYYGMDVWGQGTTVTVSS
..............|||||||||||||||||||||||||||||||||||||||||||||||||||||||||||||||||||||||||||||||||||||||||||||||||||||||||.
--------------GASVKVSCKASGYTFTSYDINWVRQATGQGLEWMGWMNPNSGNTGYAQKFQGRVTMTRNTSISTAYMELSSLRSEDTAVYYCARDPYYYYYGMDVWGQGTTVTVS-
                         ^^^^^^^^                 ^^^^^^^^                                      ^^^^^^^^^^^^^
```

```
Best Alignment of Dusigitumab light CDRs chain to a sequence from OAS
QSVLTQPPSVSAAPGQKVTISCSGSSSNIENNHVSWYQQLPGTAPKLLIYDNNKRPSGIPDRFSGSKSGTSATLGITGLQTGDEADYYCETWDTSLSAGRVFGGGTKLTVL
|||||||||||||||.|.|||||||||||||.|||||.|||||||||||||||||||.||||||||||||||||||||||||||||.|||||||||.|||.|||.|||
QSVLTQPPSVSAAPGQNVNISCSGSSSNIENNFVSWYQKLPGTAPKLLIYDNNKRPSGIRDRFSGSKSGTSATLGITGLQTGDEADYYCGTWDTSLSAGGVFGSGTKVTVL
                 ^^^^^^^^         ^^^                 ^^^^^^^^^^^^
```

```
Best Alignment of Dusigitumab CDR-H3 chain to a sequence from OAS
QVQLVQSGAEVKKPGASVKVSCKASGYTFTSYDINWVRQATGQGLEWMGWMNPNSGNTGYAQKFQGRVTMTRNTSISTAYMELSSLRSEDTAVYYCARDPYYYYYGMDVWGQGTTVTVSS
..............|||||||||||.||.||.|.|||||.||||||||...|..|...||||||||||.|...|.|||||||||||||||||||||||||||||||||||||||||||||
--------------GASVKVSCKASGGTFSSYAISWVRQAPGQGLEWMGGIIPIFGTANYAQKFQGRVTITADESTSTAYMELSSLRSEDTAVYYCARDPYYYYYGMDVWGQGTTVTVSS
                 ^^^^^^^^         ^^^^^^^^                 ^^^^^^^^^^^^^
```

Therapeutic : Eculizumab

```
Best Alignment of Eculizumab heavy chain to a sequence from OAS
QVQLVQSGAEVKKPGASVKVSCKASGYIFSNYWIQWVRQAPGQGLEWMGEILPGSGSTEYTENFKDRVTMTRDTSTSTVYMELSSLRSEDTAVYYCARYFFGSS---PNWYFDVWGQGTLVTVSS
||||||||||||||||||||||||||.|..|...||||||||||||||.|.|..|||.|...|..|||||||||||||||||||||||||||||||||.|.||....||.|||.|||||||||||
QVQLVQSGAEVKKPGASVKVSCKASGYTFTSYYMHWVRQAPGQGLEWMGIINPSGGSTSYAQKFQGRVTMTRDTSTSTVYMELSSLRSEDTAVYYCARGFGGSYQINPNYYFDYWGQGTLVTVSS
                 ^^^^^^^^         ^^^^^^^^                 ^^^^^^^^^^^^^^^^^^
```

```
Best Alignment of Eculizumab light chain to a sequence from OAS
DIQMTQSPSSLSASVGDRVTITCGASENIYGALNWYQQKPGKAPKLLIYGATNLADGVPSRFSGSGSGTDFTLTISSLQPEDFATYYCQNVLNTPLTFGQGTKVEIK
|||||||||||||||||||||||.||.||...|||||||||||||||||.|.||..|||||||||||||||||||||||||||||||||...|||||||.|||||||
DIQMTQSPSSLSASVGDRVTITCRASQNIRNYLNWYQQKPGKAPKLLIYTASNLQSGVPSRFSGSGSGTDFTLTISSLQPEDFATYYCQQSDNTPLTFGGGTKVEIK
                  ^^^^^^         ^^^                 ^^^^^^^^^
```

```
Best Alignment of Eculizumab heavy chain CDRs to a sequence from OAS
QVQLVQSGAEVKKPGASVKVSCKASGYIFSNYWIQWVRQAPGQGLEWMGEILPGSGSTEYTENFKDRVTMTRDTSTSTVYMELSSLRSEDTAVYYCARYFFGSSPNWYFDVWGQGTLVTVSS
.|||.|||||..|||||||.||||.||.||.|||.||.|.||.||||.||||||||||.|.|.||...|.|.|||..|.||.||||.|||.|||||||...||||||||||||.||.|||||
-VQLQQSGAELMKPGASVKMSCKATGYTFSSYWIEWVKQRPGHGLEWIGEILPGSGSTNYNEKFKGKATFTADTSSNTAYMQLSSLTSEDSAVYYCARDYYGSSPNWYFDVWGAGTTVTVSS
                 ^^^^^^^^         ^^^^^^^^                 ^^^^^^^^^^^^^^^
```

```
Best Alignment of Eculizumab light CDRs chain to a sequence from OAS
DIQMTQSPSSLSASVGDRVTITCGASENIYGALNWYQQKPGKAPKLLIYGATNLADGVPSRFSGSGSGTDFTLTISSLQPEDFATYYCQNVLNTPLTFGQGTKVEIK
||||||||.|||||||..||||||||||||||||||||.|.|..|.||||||||||||..|||||||||....|..|||.|.|.||||||||||||||||||.|||.|.|
DIQMTQSPASLSASVGETVTITCGASENIYGALNWYQRKQGISPQLLIYGATNLADGMSSRFSGSGSGRQYSLKVSSLHPDDVATYYCQNVLNTPLTFGAGTKLELK
                  ^^^^^^         ^^^                 ^^^^^^^^^
```

```
Best Alignment of Eculizumab CDR-H3 chain to a sequence from OAS
QVQLVQSGAEVKKPGASVKVSCKASGYIFSNYWIQWVRQAPGQGLEWMGEILPGSGSTEYTENFKDRVTMTRDTSTSTVYMELSSLRSEDTAVYYCARYFFGSSPNWYFDVWGQGTLVTVSS
.|.||.||||..|||||||.||||.||.|..|||.||.|.||.||||.||||||||||.|.|.||...|.|.|||..|.||.||||..||.|.||||||..||||||||||||.||.|||||
-VKLVESGAELMKPGASVKLSCKATGYTFTGYWIEWVKQRPGHGLEWIGEILPGSGSTNYNEKFKGKATFTADTSSNTAYMQLSSLTTEDSAIYYCARYYYGSSPNWYFDVWGTGTTVTVSS
                 ^^^^^^^^         ^^^^^^^^                 ^^^^^^^^^^^^^^^
```

Therapeutic : Efalizumab

```
Best Alignment of Efalizumab heavy chain to a sequence from OAS
EVQLVESGGGLVQPGGSLRLSCAASGYSFTGHWMNWVRQAPGKGLEWVGMIHPSDSETRYNQKFKDRFTISVDKSKNTLYLQMNSLRAEDTAVYYCARGIYFY----------GTTYFDYWGQGTLVTVSS
||||||||||||||||||||||||||..|....||||||||||||||..|..|...|.|....|.|||||.|.||||||||||||||||||||||||.|..||..........||.|||||||||||||||
EVQLVESGGGLVQPGGSLRLSCAASGFTFSSYSMNWVRQAPGKGLEWVSAISGSGGSTYYADSVKGRFTISRDNSKNTLYLQMNSLRAEDTAVYYCAKGSGFYDILTGYYGPTGTHYFDYWGQGTLVTVSS
                 ^^^^^^^^         ^^^^^^^^                 ^^^^^^^^^^^^^^^^^^^^^^^^
```

```
Best Alignment of Efalizumab light chain to a sequence from OAS
DIQMTQSPSSLSASVGDRVTITCRASKTISKYLAWYQQKPGKAPKLLIYSGSTLQSGVPSRFSGSGSGTDFTLTISSLQPEDFATYYCQQHNEYPLTFGQGTKVEIK
||||||||||||||||||||||||||..||.|||||||||||||||||||.||||||||||||||||||||||||||||||||||||||||.||.|||||||||||
DIQMTQSPSSLSASVGDRVTITCRASQGISSYLAWYQQKPGKAPKLLIYSASTLQSGVPSRFSGSGSGTDFTLTISSLQPEDFATYYCQQHNSYPWTFGQGTKVEIK
                  ^^^^^^         ^^^                 ^^^^^^^^^
```

```
Best Alignment of Efalizumab heavy chain CDRs to a sequence from OAS
EVQLVESGGGLVQPGGSLRLSCAASGYSFTGHWMNWVRQAPGKGLEWVGMIHPSDSETRYNQKFKDRFTISVDKSKNTLYLQMNSLRAEDTAVYYCARGIYFYGTTYFDYWGQGTLVTVSS
.|||..||..||.||.|..|||.|||||||..|||||.|.||.||||.|||||||||||||.||||||..|..||||..|.|.|..|...||.||||||||||.||..||||||||||..||||
QVQLQQSGAELVRPGASVKLSCKASGYSFTSYWMNWVKQRPGQGLEWIGMIHPSDSETRLNQKFKDKATLTVDKSSSTAYMQLSSPTSEDSAVYYCARGIYYYGSSYFDYWGQGTTLTVSS
                 ^^^^^^^^         ^^^^^^^^                 ^^^^^^^^^^^^^^
```

```
Best Alignment of Efalizumab light CDRs chain to a sequence from OAS
DIQMTQSPSSLSASVGDRVTITCRASKTISKYLAWYQQKPGKAPKLLIYSGSTLQSGVPSRFSGSGSGTDFTLTISSLQPEDFATYYCQQHNEYPLTFGQGTKVEIK
|.|.|||||.|.||.|...||.||||||||||||||||.||||..|||||||||||||.|||||||||||||||||||||.|||||.|||||||||||||||.|||.|.|
DVQITQSPSYLAASPGETITINCRASKTISKYLAWYQEKPGKTNKLLIYSGSTLQSGIPSRFSGSGSGTDFTLTISSLEPEDFAMYYCQQHNEYPLTFGAGTKLELK
                  ^^^^^^         ^^^                 ^^^^^^^^^
```

```
Best Alignment of Efalizumab CDR-H3 chain to a sequence from OAS
EVQLVESGGGLVQPGGSLRLSCAASGYSFTGHWMNWVRQAPGKGLEWVGMIHPSDSETRYNQKFKDRFTISVDKSKNTLYLQMNSLRAEDTAVYYCARGIYFYGTTYFDYWGQGTLVTVSS
||.|||||||||.|||||.|||.|||..|....|.||||.|.|.||||..|......|.|....|.|||||.|...|.|||||.|.|.||||.||||||.||||.||||||||||...|||
EVELVESGGGLVKPGGSLKLSCEASGFAFSSYVMSWVRQTPEKRLEWVASISSG-GNTYYPDSVKGRFTISRDDARNILYLQMSSQRSEDTAMYYCARGPYFYGITYFDYWGQGTTLSVSS
                 ^^^^^^^^         ^^^^^^^^                 ^^^^^^^^^^^^^^
```

Therapeutic : Eldelumab

```
Best Alignment of Eldelumab heavy chain to a sequence from OAS
QMQLVESGGGVVQPGRSLRLSCTASGFTFSNNGMHWVRQAPGKGLEWVAVIWFDGMNKFYVDSVKGRFTISRDNSKNTLYLEMNSLRAEDTAIYYCAREGDGSGI-YYYYGMDVWGQGTTVTVSS
|.||||||||||||||||||||.|||||||..||||||||||||||||||||.||.||.|.||||||||||||||||||||.||||||||||.||||||||.||..|||||||||||||||||||
QVQLVESGGGVVQPGRSLRLSCAASGFTFSSYGMHWVRQAPGKGLEWVAVIWYDGSNKYYADSVKGRFTISRDNSKNTLYLQMNSLRAEDTAVYYCAREGDSSGYYYYYYGMDVWGQGTTVTVSS
                 ^^^^^^^^         ^^^^^^^^                 ^^^^^^^^^^^^^^^^^^
```

```
Best Alignment of Eldelumab light chain to a sequence from OAS
EIVLTQSPGTLSLSPGERATLSCRASQSVSSSYLAWYQQKPGQAPRLLIYGASSRATGIPDRFSGSGSGTDFTLTISRLEPEDFAVYYCQQYGSSPIFTFGPGTKVDIK
||||||||||||||||||||||||||||||||||||||||||||||||||||||||||||||||||||||||||||||||||||||||||||||||||||||||||||
EIVLTQSPGTLSLSPGERATLSCRASQSVSSSYLAWYQQKPGQAPRLLIYGASSRATGIPDRFSGSGSGTDFTLTISRLEPEDFAVYYCQQYGSSPIFTFGPGTKVDIK
                  ^^^^^^^         ^^^                 ^^^^^^^^^^
```

```
Best Alignment of Eldelumab heavy chain CDRs to a sequence from OAS
QMQLVESGGGVVQPGRSLRLSCTASGFTFSNNGMHWVRQAPGKGLEWVAVIWFDGMNKFYVDSVKGRFTISRDNSKNTLYLEMNSLRAEDTAIYYCAREGDGSGIYYYYGMDVWGQGTTVTVSS
|.||||||||||||||||||||.|||||||..|||||||||||||||.||||.||.||.|.||||||.|||||||||||.|.||||||||||.|.||||||||..||||||||||||||||||
QVQLVESGGGVVQPGRSLRLSCAASGFTFSSYGMHWVRQAPGKGLEWGAVIWYDGSNKYYADSVKGRLTISRDNSKNTLHLQMNSLRAEDTAVYFCAREGDGSYGYYYYGMDVWGQGTTVTVSS
                 ^^^^^^^^         ^^^^^^^^                 ^^^^^^^^^^^^^^^^^
```

```
Best Alignment of Eldelumab light CDRs chain to a sequence from OAS
EIVLTQSPGTLSLSPGERATLSCRASQSVSSSYLAWYQQKPGQAPRLLIYGASSRATGIPDRFSGSGSGTDFTLTISRLEPEDFAVYYCQQYGSSPIFTFGPGTKVDIK
||||||||||||||||||||||||||||||||||||||||||||||||||||||||||||||||||||||||||||||||||||||||||||||||||||||||||||
EIVLTQSPGTLSLSPGERATLSCRASQSVSSSYLAWYQQKPGQAPRLLIYGASSRATGIPDRFSGSGSGTDFTLTISRLEPEDFAVYYCQQYGSSPIFTFGPGTKVDIK
                  ^^^^^^^         ^^^                 ^^^^^^^^^^
```

```
Best Alignment of Eldelumab CDR-H3 chain to a sequence from OAS
QMQLVESGGGVVQPGRSLRLSCTASGFTFSNNGMHWVRQAPGKGLEWVAVIWFDGMNKFYVDSVKGRFTISRDNSKNTLYLEMNSLRAEDTAIYYCAREGDGSGIYYYYGMDVWGQGTTVTVSS
..............|.||||||.|||||.|.|.|.|||||||||||||.||........|.|||||||||||||||||||.||||||||||.||||||||||.|||||||||||||||||||.
--------------GGSLRLSCAASGFTVSSNYMSWVRQAPGKGLEWVSVIYSG-GSTYYADSVKGRFTISRDNSKNTLYLQMNSLRAEDTAVYYCAREGDGSGGYYYYGMDVWGQGTTVTVS-
                 ^^^^^^^^         ^^^^^^^^                 ^^^^^^^^^^^^^^^^^
```

Therapeutic : Elotuzumab

Best Alignment of Elotuzumab heavy chain to a sequence from OAS
EVQLVESGGGLVQPGGSLRLSCAASGFDFSRYWMSWVRQAPGKGLEWIGEINPDSSTINYAPSLKDKFIISRDNAKNSLYLQMNSLRAEDTAVYYCARPDGNYWYFDVWGQGTLVTVSS
||||||||||||||||||.|||||||||||||||||||||||||||||||||||||||||.||||||||||||||||.|||||...|.||||.|||||.|||||||||||.||.|||||
EVQLVESGGGLVQPGGSLKLSCAASGFDFSRYWMSWVRQAPGKGLEWIGEINPDSSTINYTPSLKDKFIISRDNAKNTLYLQMSKVRSEDTALYYCARHDGNYWYFDVWGAGTTVTVSS
                    ^^^^^^^^            ^^^^^^^^                ^^^^^^^^^^^^

Best Alignment of Elotuzumab light chain to a sequence from OAS
DIQMTQSPSSLSASVGDRVTITCKASQDVGIAVAWYQQKPGKVPKLLIYWASTRHTGVPDRFSGSGSGTDFTLTISSLQPEDVATYYCQQYSS-YPYTFGQGTKVEIK
|||||||||||||||||||||||.|||.|.|.|||||||||||||||||.|||...|||.|||||||||||||||||||||||||||||||.|..|.|||.|||||||
DIQMTQSPSSLSASVGDRVTITCRASQGVSIYVAWYQQKPGKVPKLLIYAASTLQSGVPSRFSGSGSGTDFTLTISSLQPEDVATYYCQQYNSAPPLTFGGGTKVEIK
                    ^^^^^^            ^^^                ^^^^^^^^^^

Best Alignment of Elotuzumab heavy chain CDRs to a sequence from OAS
EVQLVESGGGLVQPGGSLRLSCAASGFDFSRYWMSWVRQAPGKGLEWIGEINPDSSTINYAPSLKDKFIISRDNAKNSLYLQMNSLRAEDTAVYYCARPDGNYWYFDVWGQGTLVTVSS
||.|||||||||||||||.|||||||||||||||||||||||||||||||||||||||||.||||||||||||||||.|||||.....||||.||||||||.||||||||.||.|||||
EVKLVESGGGLVQPGGSLKLSCAASGFDFSRYWMSWVRQAPGKGLEWIGEINPDSSTINYTPSLKDKFIISRDNAKNTLYLQMSKVISEDTALYYCARPDGYYWYFDVWGAGTTVTVSS
                    ^^^^^^^^            ^^^^^^^^                ^^^^^^^^^^^^

Best Alignment of Elotuzumab light CDRs chain to a sequence from OAS
DIQMTQSPSSLSASVGDRVTITCKASQDVGIAVAWYQQKPGKVPKLLIYWASTRHTGVPDRFSGSGSGTDFTLTISSLQPEDVATYYCQQYSSYPYTFGQGTKVEIK
||.||||....|.||||||.|||||||||||||||||||||..|||||||||||||||||||.|||||||||||||..|.||.|.|.||||||||||||.|||.|||
DIVMTQSHKFMSTSVGDRVSITCKASQDVGIAVAWYQQKPGQSPKLLIYWASTRHTGVPDRFTGSGSGTDFTLTISNVQSEDLADYFCQQYSSYPYTFGGGTKLEIK
                    ^^^^^^            ^^^                ^^^^^^^^^

Best Alignment of Elotuzumab CDR-H3 chain to a sequence from OAS
EVQLVESGGGLVQPGGSLRLSCAASGFDFSRYWMSWVRQAPGKGLEWIGEINPDSSTINYAPSLKDKFIISRDNAKNSLYLQMNSLRAEDTAVYYCARPDGNYWYFDVWGQGTLVTVSS
||.|||||||||||||||.||||||||.||.|.|.||||.|.|.|||...|........|....|....||||.|||.|||||..|..||||.|||||||||||||||||.||.|||||
EVHLVESGGGLVQPGGSLKLSCAASGFTFSDYYMYWVRQTPEKRLEWVAYISNGGGSTYYPDTVKGRSTISRDTAKNTLYLQMSRLKSEDTAMYYCARPDGNYWYFDVWGTGTTVTVSS
                    ^^^^^^^^            ^^^^^^^^                ^^^^^^^^^^^^

Therapeutic : Emapalumab

Best Alignment of Emapalumab heavy chain to a sequence from OAS
EVQLLESGGGLVQPGGSLRLSCAASGFTFSSYAMSWVRQAPGKGLEWVSAISGSGGSTYYADSVKGRFTISRDNSKNTLYLQMNSLRAEDTAVYYCAKDGSSGWYVPHWFDPWGQGTLVTVSS
||||||||||||||||||||||||||||||||||||||||||||||||||||||||||||||||||||||||||||||||||||||||||||||||||||.||.|.|||||||||||||||
EVQLLESGGGLVQPGGSLRLSCAASGFTFSSYAMSWVRQAPGKGLEWVSAISGSGGSTYYADSVKGRFTISRDNSKNTLYLQMNSLRAEDTAVYYCAKDGSSSWYGPNWFDPWGQGTLVTVSS
                    ^^^^^^^^            ^^^^^^^^                ^^^^^^^^^^^^^^^^

Best Alignment of Emapalumab light chain to a sequence from OAS
NFMLTQPHSVSESPGKTVTISCTRSSGSIASNYVQWYQQRPGSSPTTVIYEDNQRPSGVPDRFSGSIDSSSNSASLTISGLKTEDEADYYCQSYDGSNRWMFGGGTKLTVL
||||||||||||||||||||||||||||||||||||||||||||||||||||||||||||||||||||||||||||||||||||||||||||||||||||.||||||||||
NFMLTQPHSVSESPGKTVTISCTRSSGSIASNYVQWYQQRPGSSPTTVIYEDNQRPSGVPDRFSGSIDSSSNSASLTISGLKTEDEADYYCQSYDGSNRWVFGGGTKLTVL
                    ^^^^^^^^            ^^^                ^^^^^^^^^^

Best Alignment of Emapalumab heavy chain CDRs to a sequence from OAS
EVQLLESGGGLVQPGGSLRLSCAASGFTFSSYAMSWVRQAPGKGLEWVSAISGSGGSTYYADSVKGRFTISRDNSKNTLYLQMNSLRAEDTAVYYCAKDGSSGWYVPHWFDPWGQGTLVTVSS
.............||||||||||||||||||||||||||||||||||||||||||||||||||||||||||||||||||||||||||||||||||||||||||.|.|||||||||||||||
-------------PGGSLRLSCAASGFTFSSYAMSWVRQAPGKGLEWVSAISGSGGSTYYADSVKGRFTISRDNSKNTLYLQMNSLRAEDTAVYYCAKDGSSGWYTPGWFDPWGQGTLVTVSS
                    ^^^^^^^^            ^^^^^^^^                ^^^^^^^^^^^^^^^^

Best Alignment of Emapalumab light CDRs chain to a sequence from OAS
NFMLTQPHSVSESPGKTVTISCTRSSGSIASNYVQWYQQRPGSSPTTVIYEDNQRPSGVPDRFSGSIDSSSNSASLTISGLKTEDEADYYCQSYDGSNRWMFGGGTKLTVL
|.|||||||||||||||||||||||||||||||||||||||||||||||||||||||||||||||.||.||||||||||||||.||||||||||||||||.||||||||||
NLMLTQPHSVSESPGKTVTISCTRSSGSIASNYVQWYQQRPGSSPTTVIYEDNQRPSGVPDRFSGPIDGSSNSASLTISGLKTGDEADYYCQSYDGSNRWVFGGGTKLTVL
                    ^^^^^^^^            ^^^                ^^^^^^^^^^

Best Alignment of Emapalumab CDR-H3 chain to a sequence from OAS
EVQLLESGGGLVQPGGSLRLSCAASGFTFSSYAMSWVRQAPGKGLEWVSAISGSGGSTYYADSVKGRFTISRDNSKNTLYLQMNSLRAEDTAVYYCAKDGSSGWYVPHWFDPWGQGTLVTVSS
..............||||||||||||||||||.|.||||||||||||||.||.|....||||||||||||||||.||.|||||||||||||||||||.|.|||||||||||||||||||||||
--------------GGSLRLSCAASGFTFSSYSMNWVRQAPGKGLEWVSSISSSSSYIYYADSVKGRFTISRDNAKNSLYLQMNSLRAEDTAVYYCARDPSSGWYVPHWFDPWGQGTLVTVSS
                    ^^^^^^^^            ^^^^^^^^                ^^^^^^^^^^^^^^^^

Therapeutic : Emibetuzumab

Best Alignment of Emibetuzumab heavy chain to a sequence from OAS
QVQLVQSGAEVKKPGASVKVSCKASGYTFTDYYMHWVRQAPGQGLEWMGRVNPNRRGTTYNQKFEGRVTMTTDTSTSTAYMELRSLRSDDTAVYYCARAN--WLDYWGQGTTVTVSS
||||||||||||||||||||||||||||||.|||||||||||||||||||.|||..||.|.|||.||||||||||||||||||||||||||||||||||...|.|||||||.|||||
QVQLVQSGAEVKKPGASVKVSCKASGYTFTGYYMHWVRQAPGQGLEWMGRINPNSGGTNYAQKFQGRVTMTTDTSTSTAYMELRSLRSDDTAVYYCARAWNYWGDYWGQGTLVTVSS
                    ^^^^^^^^            ^^^^^^^^                ^^^^^^^^^^

Best Alignment of Emibetuzumab light chain to a sequence from OAS
DIQMTQSPSSLSASVGDRVTITCSVSSSVSSIYLHWYQQKPGKAPKLLIYSTSNLASGVPSRFSGSGSGTDFTLTISSLQPEDFATYYCQVYSGYPLTFGGGTKVEIK
|||||||||||||||||||||||..|.|..|.||.|||||||||||||||..|.|.||||||||||||||||||||||||||||||||||||||.|..||||||||||||||
DIQMTQSPSSLSASVGDRVTITCRASQSI-SSYLNWYQQKPGKAPKLLIYAASSLQSGVPSRFSGSGSGTDFTLTISSLQPEDFATYYCQQYNSYPLTFGGGTKVEIK
                    ^^^^^^^            ^^^                ^^^^^^^^^

Best Alignment of Emibetuzumab heavy chain CDRs to a sequence from OAS
QVQLVQSGAEVKKPGASVKVSCKASGYTFTDYYMHWVRQAPGQGLEWMGRVNPNRRGTTYNQKFEGRVTMTTDTSTSTAYMELRSLRSDDTAVYYCARANWLDYWGQGTTVTVSS
.|||.|||.|..|||||||.||||||||||||||.||.|..|..|||.|..|||..||.|||||.|..|.|.|.|.||||||||||.|.|.|||||||||||||||||||.||||
EVQLQQSGPELVKPGASVKISCKASGYTFTDYYMNWVKQSHGKSLEWIGDINPNNGGTSYNQKFKGKATLTVDKSSSTAYMELRSLTSEDSAVYYCARANWLDYWGQGTTLTVSS
                    ^^^^^^^^            ^^^^^^^^                ^^^^^^^^

Best Alignment of Emibetuzumab light CDRs chain to a sequence from OAS
DIQMTQSPSSLSASVGDRVTITCSVSSSVSSIYLHWYQQKPGKAPKLLIYSTSNLASGVPSRFSGSGSGTDFTLTISSLQPEDFATYYCQVYSGYPLTFGGGTKVEIK
....||||...|||.|..||.||..|||||||||||||||.|..|||.||||||||||||.|||||||||...|||||...||.||||||.|||||||||||||.|||
ENVLTQSPAIMSASPGEKVTMTCRASSSVSSIYLHWYQQKSGASPKLWIYSTSNLASGVPARFSGSGSGTSYSLTISSVEAEDAATYYCQQYSGYPLTFGGGTKLEIK
                    ^^^^^^^            ^^^                ^^^^^^^^^

Best Alignment of Emibetuzumab CDR-H3 chain to a sequence from OAS
QVQLVQSGAEVKKPGASVKVSCKASGYTFTDYYMHWVRQAPGQGLEWMGRVNPNRRGTTYNQKFEGRVTMTTDTSTSTAYMELRSLRSDDTAVYYCARANWLDYWGQGTTVTVSS
.|||.|||.|..|||||||.||||||||||||||.||.|..|..|||.|..|||..||.|||||.|..|.|.|.|.||||||||||.|.|.|||||||||||||||||||.||||
EVQLQQSGPELVKPGASVKISCKASGYTFTDYYMNWVKQSHGKSLEWIGDINPNNGGTSYNQKFKGKATLTVDKSSSTAYMELRSLTSEDSAVYYCARANWLDYWGQGTTLTVSS
                    ^^^^^^^^            ^^^^^^^^                ^^^^^^^^

Therapeutic : Enfortumab

Best Alignment of Enfortumab heavy chain to a sequence from OAS
EVQLVESGGGLVQPGGSLRLSCAASGFTFSSYNMNWVRQAPGKGLEWVSYISSSSSTIYYADSVKGRFTISRDNAKNSLSLQMNSLRDEDTAVYYCARAYYYGMDVWGQGTTVTVSS
||||||||||||||||||||||||||||||||.||||||||||||||||||||||||||||||||||||||||||||||.|||||||||||||||||||||||||||||||||||||
EVQLVESGGGLVQPGGSLRLSCAASGFTFSSYSMNWVRQAPGKGLEWVSYISSSSSTIYYADSVKGRFTISRDNAKNSLYLQMNSLRDEDTAVYYCARAYYYGMDVWGQGTTVTVSS
                    ^^^^^^^^            ^^^^^^^^                ^^^^^^^^^^

Best Alignment of Enfortumab light chain to a sequence from OAS

```
DIQMTQSPSSVSASVGDRVTITCRASQGISGWLAWYQQKPGKAPKFLIYAASTLQSGVPSRFSGSGSGTDFTLTISSLQPEDFATYYCQQANSFPPTFGGGTKVEIK
||||||||||||||||||||||||||||||.||||||||||||||.|||||||||||||||||||||||||||||||||||||||||||||||||||||||||||||
DIQMTQSPSSVSASVGDRVTITCRASQGISSWLAWYQQKPGKAPKLLIYAASTLQSGVPSRFSGSGSGTDFTLTISSLQPEDFATYYCQQANSFPPTFGGGTKVEIK
                        ^^^^^^          ^^^                ^^^^^^^^^
```

Best Alignment of Enfortumab heavy chain CDRs to a sequence from OAS

```
EVQLVESGGGLVQPGGSLRLSCAASGFTFSSYNMNWVRQAPGKGLEWVSYISSSSSTIYYADSVKGRFTISRDNAKNSLSLQMNSLRDEDTAVYYCARAYYYGMDVWGQGTTVTVSS
||||||||||||||||||||||||||||||||.||||||||||||||||||||||||||||||||||||||||||||||.|||||||||||||||||||||||||||||||||||||
EVQLVESGGGLVQPGGSLRLSCAASGFTFSSYSMNWVRQAPGKGLEWVSYISSSSSTIYYADSVKGRFTISRDNAKNSLYLQMNSLRDEDTAVYYCARAYYYGMDVWGQGTTVTVSS
                       ^^^^^^^^        ^^^^^^^^                  ^^^^^^^^^^
```

Best Alignment of Enfortumab light CDRs chain to a sequence from OAS

```
DIQMTQSPSSVSASVGDRVTITCRASQGISGWLAWYQQKPGKAPKFLIYAASTLQSGVPSRFSGSGSGTDFTLTISSLQPEDFATYYCQQANSFPPTFGGGTKVEIK
|||||||||||||||||||||||||||||||||||||||||||||.||||||.||||||||||||||||||||||||||||||||||||||||||||||.||..|||
DIQMTQSPSSVSASVGDRVTITCRASQGISGWLAWYQQKPGKAPKLLIYAASSLQSGVPSRFSGSGSGTDFTLTISSLQPEDFATYYCQQANSFPPTFGQGTRLEIK
                        ^^^^^^          ^^^                ^^^^^^^^^
```

Best Alignment of Enfortumab CDR-H3 chain to a sequence from OAS

```
EVQLVESGGGLVQPGGSLRLSCAASGFTF--SSYNMNWVRQAPGKGLEWVSYISSSSSTIYYADSVKGRFTISRDNAKNSLSLQMNSLRDEDTAVYYCARAYYYGMDVWGQGTTVTVSS
.................|.|.|..||.......|...|.||.||.||||..||..|....||..|.|.|.|||.|..||..||...|....||||||||||||||||||||||||||||
--------------SETLSLTCTVSGGSISSGDYYWSWIRQPPGTGLEWIGYIYYS-GSTYYNPSLKSRVTISVDTSKNQFSLKLSSVTAADTAVYYCARAYYYGMDVWGQGTTVTVSS
                       ^^^^^^^^^^        ^^^^^^^^                  ^^^^^^^^^^
```

Therapeutic : Enokizumab

Best Alignment of Enokizumab heavy chain to a sequence from OAS

```
QVQLVQSGAEVKKPGSSVKVSCKASGGTFSYYWIEWVRQAPGQGLEWMGEILPGSGTTNPNEKFKGRVTITADESTSTAYMELSSLRSEDTAVYYCARADYYGS-DYVKFDYWGQGTLVTVSS
||||||||||||||||||||||||||||||.|.|.||||||||||||||.|.|..||.|...||.|||||||||||||||||||||||||||||||||||||.|..||.||||||||||||||
QVQLVQSGAEVKKPGSSVKVSCKASGGTFSSYAISWVRQAPGQGLEWMGRIIPIFGTANYAQKFQGRVTITADESTSTAYMELSSLRSEDTAVYYCARADYYDSSGYVFFDYWGQGTLVTVSS
                       ^^^^^^^^        ^^^^^^^^                  ^^^^^^^^^^^^^^^^
```

Best Alignment of Enokizumab light chain to a sequence from OAS

```
DIQMTQSPSSLSASVGDRVTITCKASQHVITHVTWYQQKPGKAPKLLIYGTSYSYSGVPSRFSGSGSGTDFTLTISSLQPEDFATYYCQQFYEYPLTFGGGTKVEIK
|||||||||||||||||||||||.|||.|.|.|.|||.|||||||||||..|...||||||||||||||||||||||||||||||||||||..||||||||||||||
DIQMTQSPSSLSASVGDRVTITCQASQSVSTWVAWYQHKPGKAPKLLIYKASSLQSGVPSRFSGSGSGTDFTLTISSLQPEDFATYYCQQFNSYPLTFGGGTKVEIK
                        ^^^^^^          ^^^                ^^^^^^^^^
```

Best Alignment of Enokizumab heavy chain CDRs to a sequence from OAS

```
QVQLVQSGAEVKKPGSSVKVSCKASGGTFSYYWIEWVRQAPGQGLEWMGEILPGSGTTNPNEKFKGRVTITADESTSTAYMELSSLRSEDTAVYYCARADYYGSDYVKFDYWGQGTLVTVSS
.|||.|||||..|||.|||.||||.|.|||.||||||.|.||.||||.||||||||.||.||||||..|.|||.|..||||.||||.|||.|||||||.|||||.||.||||||||..||||
-VQLQQSGAELMKPGASVKISCKATGYTFSSYWIEWVKQRPGHGLEWIGEILPGSGSTNYNEKFKGKATFTADTSSNTAYMQLSSLTSEDSAVYYCARPDYYGSSYVSFDYWGQGTTLTVSS
                       ^^^^^^^^        ^^^^^^^^                  ^^^^^^^^^^^^^^^
```

Best Alignment of Enokizumab light CDRs chain to a sequence from OAS

```
DIQMTQSPSSLSASVGDRVTITCKASQHVITHVTWYQQKPGKAPKLLIYGTSYSYSGVPSRFSGSGSGTDFTLTISSLQPEDFATYYCQQFYEYPLTFGGGTKVEIK
|||||.|.|||.|||||||||.|.|||.|||...||||.||||||||..|.|....|||||||||||..||||||.|.||||||||.|||.|.||||.||||.||||
DIQMTHSSSSLAASVGDRVTIACRASQNVITYLAWYQQTPGKAPKLLLSGASTLQGGVPSRFSGSGSESDFTLTITSVQPEDFATYFCQQTYAYPLTCGGGTRVEIK
                        ^^^^^^          ^^^                ^^^^^^^^^
```

Best Alignment of Enokizumab CDR-H3 chain to a sequence from OAS

```
QVQLVQSGAEVKKPGSSVKVSCKASGGTFSYYWIEWVRQAPGQGLEWMGEILPGSGTTNPNEKFKGRVTITADESTSTAYMELSSLRSEDTAVYYCARADYYGSDYVKFDYWGQGTLVTVSS
.......|||..|||.|||.||||||.||..||..||.|.|||||||.|||.|....||.|.||||..|.|.|.|.|.|||||.||||.|||.|||||||||||||.||.||||||||..||||
-------GAELVKPGASVKLSCKASGYTFTSYWMQWVKQRPGQGLEWIGEIDPSDSYTNYNQKFKGKATLTVDTSSSTAYMQLSSLTSEDSAVYYCARADYYGSSYVGFDYWGQGTTLTVSS
                       ^^^^^^^^        ^^^^^^^^                  ^^^^^^^^^^^^^^^
```

Therapeutic : Ensituximab

Best Alignment of Ensituximab heavy chain to a sequence from OAS

```
QVQLKESGPDLVAPSQSLSITCTVSGFSLSKFGVNWVRQPPGKGLEWLGVIWGDGSTSYNSGLISRLSISKENSKSQVFLKLNSLQADDTATYYCVKPGGDYWGHGTSVTVSS
|||||||||.|||||||||||||||||||...||.||||||||||||||||||||||.|.|.||||||||||||||||||||||||.||||||||.||||||||.||||||||
QVQLKESGPGLVAPSQSLSITCTVSGFSLTSYGVSWVRQPPGKGLEWLGVIWGDGSTNYHSALISRLSISKENSKSQVFLKLNSLQTDDTATYYCAKPGGDYWGQGTSVTVSS
                       ^^^^^^^^        ^^^^^^^                 ^^^^^^^
```

Best Alignment of Ensituximab light chain to a sequence from OAS

```
QVVLTQSPVIMSASPGEKVTMTCSASSSISYMYWYQQKPGTSPKRWIYDTSKLASGVPARFSGSGSGTSYSLTISNMEAGDAATYYCHQRDSYPWTFGGGTNLEIK
|.||||||.||||||||||||||||||||||||||||||||||||||||||||||||||||||||||.|||.||||||||||.||||||||||.||||
QIVLTQSPAIMSASPGEKVTMTCSASSSISYMYWYQQKPGTSPKRWIYDTSKLASGVPARFSGSGSGTSYSLTISSMEAEDAATYYCHQRSSYPWTFGGGTKLEIK
                        ^^^^^          ^^^                ^^^^^^^^^
```

Best Alignment of Ensituximab heavy chain CDRs to a sequence from OAS

```
QVQLKESGPDLVAPSQSLSITCTVSGFSLSKFGVNWVRQPPGKGLEWLGVIWGDGSTSYNSGLISRLSISKENSKSQVFLKLNSLQADDTATYYCVKPGGDYWGHGTSVTVSS
.|.|.||||.|||||||||||||||||||...||.||||||||||||||||||||||.|.|.|||||||||.||||||||||||||.|.||||||.||||||||.||||||||
-VKLVESGPGLVAPSQSLSITCTVSGFSLTSYGVSWVRQPPGKGLEWLGVIWGDGSTNYHSALISRLSISKDNSKSQVFLKLNSLQTDVTATYYCAKPGGDYWGQGTSVTVSS
                       ^^^^^^^^        ^^^^^^^                 ^^^^^^^
```

Best Alignment of Ensituximab light CDRs chain to a sequence from OAS

```
QVVLTQSPVIMSASPGEKVTMTCSASSSISYMYWYQQKPGTSPKRWIYDTSKLASGVPARFSGSGSGTSYSLTISNMEAGDAATYYCHQRDSYPWTFGGGTNLEIK
|.||||||.|||||.|||||||||||||||||.||||||||||||||||||||||||||||||||||||||||||.|||.||||||||||.||||||||||.||||
QIVLTQSPAIMSASTGEKVTMTCSASSSISYMHWYQQKPGTSPKRWIYDTSKLASGVPARFSGSGSGTSYSLTISSMEAEDAATYYCHQRSSYPWTFGGGTKLEIK
                        ^^^^^          ^^^                ^^^^^^^^^
```

Best Alignment of Ensituximab CDR-H3 chain to a sequence from OAS

```
QVQLKESGPDLVAPSQSLSITCTVSGFSLSKFGVNWVRQPPGKGLEWLGVIWGD---GSTSYNSGLISRLSISKENSKSQVFLKLNSLQADDTATYYCVKPGGDYWGHGTSVTVSS
.|||.|||..||.|..||..||..||||......|||||.|||||||...|........|.|......|..|....|.|...|..|.|...|||.||||.|||||||.||||||||
EVQLVESGGGLVQPKGSLKLSCAASGFSFNTYAMNWVRQAPGKGLEWVARIRSKSNKYATYYADSVKDRFTIYRDDSESMLYLQMNNLKTEDTAMYYCVSPGGDYWGQGTSVTVSS
                       ^^^^^^^^        ^^^^^^^^^^                  ^^^^^^^
```

Therapeutic : Epratuzumab

Best Alignment of Epratuzumab heavy chain to a sequence from OAS

```
QVQLVQSGAEVKKPGSSVKVSCKASGYTFTSYWLHWVRQAPGQGLEWIGYINPRNDYTEYNQNFKDKATITADESTNTAYMELSSLRSEDTAFYFCARRDITTFYWGQGTTVTVSS
||||.|||||..|||.|||.|||||||||||||.|||.|.|||||||||||||...||||||.||||||.|||.|..||||.||||.|||.|.|.|.|||.||||.|||||||.||||
QVQLKQSGAELAKPGASVKMSCKASGYTFTSYWMHWVKQRPGQGLEWIGYINPSTGYTEYNQKFKDKATLTADKSSSTAYMQLSSLTSEDSAVYYCARSDITTHYWGQGTTLTVSS
                       ^^^^^^^^        ^^^^^^^^                  ^^^^^^^^^
```

Best Alignment of Epratuzumab light chain to a sequence from OAS

```
DIQLTQSPSSLSASVGDRVTMSCKSSQSVLYSANHKNYLAWYQQKPGKAPKLLIYWASTRESGVPSRFSGSGSGTDFTFTISSLQPEDIATYYCHQYLSSWTFGGGTKLEIK
.|..|||||||..|.||.||||||||||||||.|.||||||||||||.|||||||||||||||||.||.|||||||||.||||.|.||.|.|||||||||||||||||||||
NIMMTQSPSSLAVSAGDKVTMSCKSSQSVLYSSNQKNYLAWYQQKPGQAPKLLIYWASTRESGVPDRFTGSGSGTDFTLTISSVQAEDLAVYYCHQYLSSWTFGGGTKLEIK
                        ^^^^^^^^^^^^          ^^^                  ^^^^^^^^
```

Best Alignment of Epratuzumab heavy chain CDRs to a sequence from OAS

```
QVQLVQSGAEVKKPGSSVKVSCKASGYTFTSYWLHWVRQAPGQGLEWIGYINPRNDYTEYNQNFKDKATITADESTNTAYMELSSLRSEDTAFYFCARRDITTFYWGQGTTVTVSS
.|||.|||||...||.|||.|||||||||||||.|||.|.|||||||||.|||...||.|.|.||.|||.|.|.|.|..||||.||||.|||.|.|.|.|||||.||||||||||.||||
-VQLQQSGAELVRPGTSVKLSCKASGYTFTSYWMHWVKQRPGQGLEWIGVINPSDSYTNYKQKFKGKATLTVDTSSSTAYMQLSSLTSEDSAVYYCARRDTTTFYWGQGTTLTVSS
                       ^^^^^^^^        ^^^^^^^^                  ^^^^^^^^^
```

Best Alignment of Epratuzumab light CDRs chain to a sequence from OAS
DIQLTQSPSSLSASVGDRVTMSCKSSQSVLYSANHKNYLAWYQQKPGKAPKLLIYWASTRESGVPSRFSGSGSGTDFTFTISSLQPEDIATYYCHQYLSSWTFGGGTKLEIK
.|..|||||||..|.|..||||||||||||||||.|||.|||||||...||||||||||||||||.||.|||||||||.||||.|.||.|.|||||||||||||||||||||
NIMMTQSPSSLAVSAGEKVTMSCKSSQSVLYSANQKNYVAWYQQKPAQSPKLLIYWASTRESGVPDRFTGSGSGTDFTLTISSVQAEDLAVYYCHQYLSSWTFGGGTKLEIK
                    ^^^^^^^^^^^^          ^^^                    ^^^^^^^^

Best Alignment of Epratuzumab CDR-H3 chain to a sequence from OAS
QVQLVQSGAEVKKPGSSVKVSCKASGYTFTSYWLHWVRQAPGQGLEWIGYINPRNDYTEYNQNFKDKATITADESTNTAYMELSSLRSEDTAFYFCARRDITTFYWGQGTTVTVSS
||||.|.|||..|||.|||.|||||||||||||..||.|.||||||||||.|.|....|.||..||.|||.|.|.|..||||.||||.|||.|.|.||||||||.|||||||.||||
QVQLKQPGAELVKPGASVKMSCKASGYTFTSYWITWVKQRPGQGLEWIGDIYPGSGSTNYNEKFKSKATLTVDTSSSTAYMQLSSLTSEDSAVYYCARRDITTDYWGQGTTLTVSS
                    ^^^^^^^^          ^^^^^^^^                    ^^^^^^^^^

Therapeutic : Eptinezumab

Best Alignment of Eptinezumab heavy chain to a sequence from OAS
EVQLVESGGGLVQPGGSLRLSCAVSGIDLSGYYMNWVRQAPGKGLEWVGVIGINGATYYASWAKGRFTISRDNSKTTVYLQMNSLRAEDTAVYFCARG---------DIWGQGTLVTVSS
|||||||||||||||||||||||||...|..||||||||||||||||.||...|.||||...||||||||||||.|.||||||||||||||||.||||.........|.|||||||||||
EVQLVESGGGLVQPGGSLRLSCAVSGFSVSSNYMNWVRQAPGKGLEWVSVIYSGGSTYYADSVKGRFTISRDNSKNTLYLQMNSLRAEDTAVYYCARGAGIVVVIHFDYWGQGTLVTVSS
                    ^^^^^^^^          ^^^^^^^                    ^^^^^^^^^^^^^^

Best Alignment of Eptinezumab light chain to a sequence from OAS
-QVLTQSPSSLSASVGDRVTINCQASQSVYHNTYLAWYQQKPGKVPKQLIYDASTLASGVPSRFSGSGSGTDFTLTISSLQPEDVATYYCLGSYDCTNGDCFVFGGGTKVEIK
....||||||||||||||||.|.||||.....||||||||||||||.|||.||||.||||||||||||||||||||||||||||.|||||..|||........||||||||||
DIQMTQSPSSLSASVGDRVTITCRASQSI--SNYLAWYQQKPGKVPKLLIYAASTLQSGVPSRFSGSGSGTDFTLTISSLQPEDFATYYCQQSYD----TPLTFGGGTKVEIK
                    ^^^^^^^^          ^^^                    ^^^^^^^^^^^^^

Best Alignment of Eptinezumab heavy chain CDRs to a sequence from OAS
EVQLVESGGGLVQPGGSLRLSCAVSGIDLSGYYMNWVRQAPGKGLEWVGVIGINGATYYASWAKGRFTISRDNSKTTVYLQMNSLRAEDTAVYFCARGDIWGQGTLVTVSS
..||...|.||..|...|.|.|.|.|..|||||..|.||.|||||||.|.|...|.|.|......|.|||.|.||....|...|..|.|||||.|||||||||||.|||||
QMQLQQWGAGLLKPSETLSLTCGVHGGSLSGYYWSWIRQPPGKGLEWIGEINHSGSTNYNPSLESRVTISVDTSKNQFSLKLSSVTAADTAVYYCARGDIWGQGTMVTVSS
                    ^^^^^^^^          ^^^^^^^                    ^^^^^

Best Alignment of Eptinezumab light CDRs chain to a sequence from OAS
-QVLTQSPSSLSASVGDRVTINCQASQSVYHNTYLAWYQQKPGKVPKQLIYDASTLASGVPSRFSGSGSGTDFTLTISSLQPEDVATYYCLGSYDCTNGDCFVFGGGTKVEIK
..||||||..||.|.|.|.|..|.||||.....||||||..||..|..|||.||..|.|.|.|||||||||||||||||||||||.||||||..||.........||.|||.|||
EIVLTQSPGTLSLSPGERATLSCRASQSLETSNYLAWYQKRPGQAPRLLIYNASSRATGIPDRFSGSGSGTDFTLTISSLQPEDFATYYCQQSYSTPPSIGYTFGQGTKLEIK
                    ^^^^^^^^          ^^^                    ^^^^^^^^^^^^^

Best Alignment of Eptinezumab CDR-H3 chain to a sequence from OAS
EVQLVESGGGLVQPGGSLRLSCAVSGIDLSGYYMNWVRQAPGKGLEWVGVIGIN-GATYYASWAKGRFTISRDNSKTTVYLQMNSLRAEDTAVYFCARGDIWGQGTLVTVSS
..............|.||..|||.||...|.|.|.|||||||||||||..|.......||....||||||||||.|...||||||||||||||.|||||||||||.|||||
--------------GESLKISCAASGFTFSSYWMSWVRQAPGKGLEWVANIKQDGSEKYYVDSVKGRFTISRDNAKNSLYLQMNSLRAEDTAVYYCARGDIWGQGTMVTVSS
                    ^^^^^^^^          ^^^^^^^^                    ^^^^^

Therapeutic : Erenumab

Best Alignment of Erenumab heavy chain to a sequence from OAS
QVQLVESGGGVVQPGRSLRLSCAASGFTFSSFGMHWVRQAPGKGLEWVAVISFD-GSIKYSVDSVKGRFTISRDNSKNTLFLQMNSLRAEDTAVYYCARDRLNYYDSSGYYHYKYYGMAVWGQGTTVTVSS
|||||||||||||||||||||||||||||||.|||||||||||||||||||||.|.....|..||||||||||||||||||.|||||||||||||||||||||...||||||||.|.||||.||||||||||||
QVQLVESGGGVVQPGRSLRLSCAASGFTFSSYGMHWVRQAPGKGLEWVAVISYDGSNKYY-ADSVKGRFTISRDNSKNTLYLQMNSLRAEDTAVYYCARDLPYYYDSSGYYLYYYYGMDVWGQGTTVTVSS
                    ^^^^^^^^          ^^^^^^^^                    ^^^^^^^^^^^^^^^^^^^^^^^

Best Alignment of Erenumab light chain to a sequence from OAS
QSVLTQPPSVSAAPGQKVTISCSGSSSNIGNNYVSWYQQLPGTAPKLLIYDNNKRPSGIPDRFSGSKSGTSTTLGITGLQTGDEADYYCGTWDSRLSAVVFGGGTKLTVL
|||||||||||||||||||||||||||||||||||||||||||||||||||||||||||||||||||||||||||||||||||||||||||||.|||||||||||||||
QSVLTQPPSVSAAPGQKVTISCSGSSSNIGNNYVSWYQQLPGTAPKLLIYDNNKRPSGIPDRFSGSKSGTSTTLGITGLQTGDEADYYCGTWDSSLSAVVFGGGTKLTVL
                    ^^^^^^^^          ^^^                    ^^^^^^^^^^^

Best Alignment of Erenumab heavy chain CDRs to a sequence from OAS
QVQLVESGGGVVQPGRSLRLSCAASGFTFSSFGMHWVRQAPGKGLEWVAVISFDGSIKYSVDSVKGRFTISRDNSKNTLFLQMNSLRAEDTAVYYCARDRLNYYDSSGYYHYKYYGMAVWGQGTTVTVSS
..............|.||||||||||||.||..|.||||||||||||||.||...||..|..||||||||||||||||||.|||||||||||||||||||||||.|||||||||.|.||||.||||||||||||
--------------GGSLRLSCAASGFTVSSNYMSWVRQAPGKGLEWVSVIYSGGSTYY-ADSVKGRFTISRDNSKNTLYLQMNSLRAEDTAVYYCARDRLYYYDSSGYYYYYYYGMDVWGQGTTVTVSS
                    ^^^^^^^^          ^^^^^^^                    ^^^^^^^^^^^^^^^^^^^^^^^

Best Alignment of Erenumab light CDRs chain to a sequence from OAS
QSVLTQPPSVSAAPGQKVTISCSGSSSNIGNNYVSWYQQLPGTAPKLLIYDNNKRPSGIPDRFSGSKSGTSTTLGITGLQTGDEADYYCGTWDSRLSAVVFGGGTKLTVL
|||||||||||||||||||||||||||||||||||||||||||||||||||||||||||||||||||||||.||||.||||||||||||||||||||||||||||||||||.
QSVLTQPPSVSAAPGQKVTISCSGSSSNIGNNYVSWYQQLPGTAPKLLIYDNNKRPSGIPDRFSGSKSGTSATLGIAGLQTGDEADYYCGTWDSRLSAVVFGGGTKLTVV
                    ^^^^^^^^          ^^^                    ^^^^^^^^^^^

Best Alignment of Erenumab CDR-H3 chain to a sequence from OAS
QVQLVESGGGVVQPGRSLRLSCAASGFTFSSFGMHWVRQAPGKGLEWVAVISFDGSIKYSVDSVKGRFTISRDNSKNTLFLQMNSLRAEDTAVYYCARDRLNYYDSSGYYHYKYYGMAVWGQGTTVTVSS
..............|.||||||||||||.||..|.||||||||||||||.||...||..|..||||||||||||||||||.|||||||||||||||||||||||.|||||||||.|.||||.||||||||||||
--------------GGSLRLSCAASGFTVSSNYMSWVRQAPGKGLEWVSVIYSGGSTYY-ADSVKGRFTISRDNSKNTLYLQMNSLRAEDTAVYYCARDRLYYYDSSGYYYYYYYGMDVWGQGTTVTVSS
                    ^^^^^^^^          ^^^^^^^                    ^^^^^^^^^^^^^^^^^^^^^^^

Therapeutic : Etaracizumab

Best Alignment of Etaracizumab heavy chain to a sequence from OAS
QVQLVESGGGVVQPGRSLRLSCAASGFTFSSYDMSWVRQAPGKGLEWVAKVSSGGGSTYYLDTVQGRFTISRDNSKNTLYLQMNSLRAEDTAVYYCARHLH-----GSFASWGQGTTVTVSS
|||||||||||||||||||||||||||||||.|.|||||||||||||||...||.|||||.|.|.|||||||||||||||||||||||||||||||||||||||......|.|..|||||.|||||
QVQLVESGGGVVQPGRSLRLSCAASGFTFSSYGMHWVRQAPGKGLEWVAVIYSG-GSTYYADSVKGRFTISRDNSKNTLYLQMNSLRAEDTAVYYCARHLPAAIGMGAFDIWGQGTMVTVSS
                    ^^^^^^^^          ^^^^^^^^                    ^^^^^^^^^^^^^^^

Best Alignment of Etaracizumab light chain to a sequence from OAS
EIVLTQSPATLSLSPGERATLSCQASQSISNFLHWYQQRPGQAPRLLIRYRSQSISGIPARFSGSGSGTDFTLTISSLEPEDFAVYYCQQSGS-WPLTFGGGTKVEIK
|||||||||||||||||||||||.|||||.|||.||||||||||||||...|....|||||||||||||||||||||||||||||||||||||.||..|||||||||||||
EIVLTQSPATLSLSPGERATLSCRASQSIINFLAWYQQRPGQAPRLLIYDASNRAIGIPARFSGSGSGTDFTLTISSLEPEDFAVYYCQQYGSSPPLTFGGGTKVEIK
                    ^^^^^^          ^^^                    ^^^^^^^^^^

Best Alignment of Etaracizumab heavy chain CDRs to a sequence from OAS
QVQLVESGGGVVQPGRSLRLSCAASGFTFSSYDMSWVRQAPGKGLEWVAKVSSGGGSTYYLDTVQGRFTISRDNSKNTLYLQMNSLRAEDTAVYYCARHLHGSFASWGQGTTVTVSS
.||.||||||.|.||.||.||||||||.|||||||||||.|.|.|||||.||||||||||.|||.||||||||||.||||||||.||..||||.||||||..||||.|||||.||||.
EVQRVESGGGLVKPGGSLKLSCAASGFAFSSYDMSWVRQTPEKRLEWVAYVSSGGGSTYYPDTVKGRFTISRDNAKNTLYLQMSSLKSEDTAMYYCARHEDGSFAYWGQGTLVTVSA
                    ^^^^^^^^          ^^^^^^^^                    ^^^^^^^^^^

Best Alignment of Etaracizumab light CDRs chain to a sequence from OAS
EIVLTQSPATLSLSPGERATLSCQASQSISNFLHWYQQRPGQAPRLLIRYRSQSISGIPARFSGSGSGTDFTLTISSLEPEDFAVYYCQQSGSWPLTFGGGTKVEIK
.||.||||||||..||.|..|||.|||||||||||||||.....||.||.|.||||.|.|.||||||||||||||.|.|.|.|||..|.||||.||||||||.|||.|.|
DIVMTQSPATLSVTPGDRVSLSCRASQSISNFLHWYQQKSHESPRFLIKYASQSIPGTPSRFSGSGSGTDFTLSINSVETEDFGIYFCQQSNSWPLTFGAGTKLELK
                    ^^^^^^          ^^^                    ^^^^^^^^^
Best Alignment of Etaracizumab CDR-H3 chain to a sequence from OAS
QVQLVESGGGVVQPGRSLRLSCAASGFTFSSYDMSWVRQAPGKGLEWVAKVSSGGGSTYYLDTVQGRFTISRDNSKNTLYLQMNSLRAEDTAVYYCARHLHGSFASWGQGTTVTVSS
.|.||||||..|.||.||.|||.|||||||.|.||||||.|.|.|||||..||.|......|.|.||||||||||..||||||||.|||.|||..||||||||.|||||||||.|.||.|.
EVELVESGGDLVRPGGSLKLSCSASGFTFSTYSMSWVRQTPDKRLEWVATISSAGDYIFFPDSVKGRFTISRDNARNTLYLQMSSLRSEDTTMYYCARHLTGSFASWGQATLVTGSA
                    ^^^^^^^^          ^^^^^^^^                    ^^^^^^^^^^

Therapeutic : Etrolizumab

Best Alignment of Etrolizumab heavy chain to a sequence from OAS
```
EVQLVESGGGLVQPGGSLRLSCAASGFFITNNYWGWVRQAPGKGLEWVGYISYS-GSTSYNPSLKSRFTISRDTSKNTFYLQMNSLRAEDTAVYYCARTGSSGYFDFWGQGTLVTVSS
|||||||||||||||||||||||||||...|||..|||||||.|||||..||.|.|||.|..|.|.|||||||.||||.|||||||||||||||||||.|||.|||.|||||||||||
EVQLVESGGGLVQPGGSLRLSCAASGFTVSNNYMSWVRQAPGRGLEWVSAISGSGGSTYYADSVKGRFTISRDNSKNTLYLQMNSLRAEDTAVYYCAREGSSWYFDYWGQGTLVTVSS
          ^^^^^^^^              ^^^^^^^^                       ^^^^^^^^^^^
```

Best Alignment of Etrolizumab light chain to a sequence from OAS
```
DIQMTQSPSSLSASVGDRVTITCRASESVDDLLHWYQQKPGKAPKLLIKYASQSISGVPSRFSGSGSGTDFTLTISSLQPEDFATYYCQQGNSLPNTFGQGTKVEIK
|||||||||||||||||||||||||||.|...||||||||||||||||..||...|||||||||||||||||||||||||||||||||||.||||.|||||||||||
DIQMTQSPSSLSASVGDRVTITCRASENVNNYLHWYQQKPGKAPKLLIYKASTLQSGVPSRFSGSGSGTDFTLTISSLQPEDFATYYCQQYNSLPRTFGQGTKVEIK
           ^^^^^^             ^^^                 ^^^^^^^^^
```

Best Alignment of Etrolizumab heavy chain CDRs to a sequence from OAS
```
EVQLVESGGGLVQPGGSLRLSCAASGFFITNNYWGWVRQAPGKGLEWVGYISYSGSTSYNPSLKSRFTISRDTSKNTFYLQMNSLRAEDTAVYYCARTGSSGYFDFWGQGTLVTVSS
||||.|||.||..|...|.|.|...|..||..||.|.|..||..||..|||||||||.||||||||..|.||||||..|||.||...||||.|||||.|||||||.|||||..||||
EVQLQESGPGLAKPSQTLSLTCSVTGYSITSDYWNWIRKFPGNKLEYMGYISYSGSTYYNPSLKSRISITRDTSKNQYYLQLNSVTTEDTATYYCARWGSSGYFDYWGQGTTLTVSS
          ^^^^^^^^              ^^^^^^^                      ^^^^^^^^^^^
```

Best Alignment of Etrolizumab light CDRs chain to a sequence from OAS
```
DIQMTQSPSSLSASVGDRVTITCRASESVDDLLHWYQQKPGKAPKLLIKYASQSISGVPSRFSGSGSGTDFTLTISSLQPEDFATYYCQQGNSLPNTFGQGTKVEIK
|||||||||.||.|||||||||||||.|||.||.|||.|||.||.|||..||...||||||||||||||.||||||||||.|.|||||||.||.|||||||||..||
DIQMTQSPSTLSVSVGDRVTITCRASQSVDSLLAWYQHKPGRAPTLLIYKASSLESGVPSRFSGSGSGTEFTLTISSLQPDDVATYYCQQYNSYPNTFGQGTKLDIK
           ^^^^^^             ^^^                 ^^^^^^^^^
```

Best Alignment of Etrolizumab CDR-H3 chain to a sequence from OAS
```
EVQLVESGGGLVQPGGSLRLSCAASGFFITNNYWGWVRQAPGKGLEWVGYISYS-GSTSYNPSLKSRFTISRDTSKNTFYLQMNSLRAEDTAVYYCARTGSSGYFDFWGQGTLVTVSS
||||||||||.||||.||.|||.||||...|....|.|||||||||||..||.....|.|..|.|.||..|||..|.|.|||.||||.||||.||||||||||||||||.||.|||||
EVQLVESGGGPVQPGRSLKLSCVASGFIFSNYWMTWIRQAPGKGLEWVASISNTGDNTYYSDSVKGRFSLSRDNAKSTLYLQVNSLRSEDTATYYCARTGSSGYFDFWGPGTMVTVSS
          ^^^^^^^^              ^^^^^^^^                       ^^^^^^^^^^^
```

Therapeutic : Evinacumab

Best Alignment of Evinacumab heavy chain to a sequence from OAS
```
EVQLVESGGGVIQPGGSLRLSCAASGFTFDDYAMNWVRQGPGKGLEWVSAISGDGGSTYYADSVKGRFTISRDNSKNSLYLQMNSLRAEDTAFFYCAKDLRNTIFGVVIPDAFDIWGQGTMVTVSS
|||||||||||.||||||||||||||||||||||.||||.|||||||||.|||||||||||||||||||||||||||||||||||||.||||..||||||..|....|.|.|||||||||||||||
EVQLVESGGGVVQPGGSLRLSCAASGFTFDDYAMHWVRQAPGKGLEWVSLISGDGGSTYYADSVKGRFTISRDNSKNSLYLQMNSLRTEDTALYYCAKDLLTTV---VTPYAFDIWGQGTMVTVSS
          ^^^^^^^^              ^^^^^^^^                       ^^^^^^^^^^^^^^^^^^
```

Best Alignment of Evinacumab light chain to a sequence from OAS
```
DIQMTQSPSTLSASVGDRVTITCRASQSIRSWLAWYQQKPGKAPKLLIYKASSLESGVPSRFSGSGSGTEFTLTISSLQPDDFATYYCQQYNSYSYTFGQGTKLEIK
|||||||||||||||||||||||||||||||||||||||||||||||||||||||||||||||||||||||||||||||||||||||||||||||||||||||||||
DIQMTQSPSTLSASVGDRVTITCRASQSIRSWLAWYQQKPGKAPKLLIYKASSLESGVPSRFSGSGSGTEFTLTISSLQPDDFATYYCQQYNSYSYTFGQGTKLEIK
           ^^^^^^             ^^^                 ^^^^^^^^^
```

Best Alignment of Evinacumab heavy chain CDRs to a sequence from OAS
```
EVQLVESGGGVIQPGGSLRLSCAASGFTFDDYAMNWVRQGPGKGLEWVSAISGDGGSTYYADSVKGRFTISRDNSKNSLYLQMNSLRAEDTAFFYCAKDLRNTIFGVVIPDAFDIWGQGTMVTVSS
............||||||||||||||||||||||.||||.|||||||||.|||||||||||||||||||||||||||||||||||||.||||..|||||.|.|||||||.||||||||||||||||
------------QPGGSLRLSCAASGFTFDDYAMHWVRQAPGKGLEWVSLISGDGGSTYYADSVKGRFTISRDNSKNSLYLQMNSLRTEDTALYYCAKDYRGTIFGVVIMDAFDIWGQGTMVTVSS
          ^^^^^^^^              ^^^^^^^^                       ^^^^^^^^^^^^^^^^^^
```

Best Alignment of Evinacumab light CDRs chain to a sequence from OAS
```
DIQMTQSPSTLSASVGDRVTITCRASQSIRSWLAWYQQKPGKAPKLLIYKASSLESGVPSRFSGSGSGTEFTLTISSLQPDDFATYYCQQYNSYSYTFGQGTKLEIK
|||||.|.|||||||||||||||||||||||||||||||||||||||||||||||||||||||||||||||||||||||||||||||||||||||||||||||||||
DIQMTPSTSTLSASVGDRVTITCRASQSIRSWLAWYQQKPGKAPKLLIYKASSLESGVPSRFSGSGSGTEFTLTISSLQPDDFATYYCQQYNSYSYTFGQGTKLEIK
           ^^^^^^             ^^^                 ^^^^^^^^^
```

Best Alignment of Evinacumab CDR-H3 chain to a sequence from OAS
```
EVQLVESGGGVIQPGGSLRLSCAASGFTFDDYAMNWVRQGPGKGLEWVSAISGDGGSTYYADSVKGRFTISRDNSKNSLYLQMNSLRAEDTAFFYCAKDLRNTIFGVVIPDAFDIWGQGTMVTVSS
..............|||||.|||||||||..|.|.||||.||||||||..||.||...|||||||||||||||||||.||||||||||||||..|||||||.|||||||||||||||||||||||||
--------------GGSLRPSCAASGFTFSSYGMHWVRQAPGKGLEWVAVISYDGSNKYYADSVKGRFTISRDNSKNTLYLQMNSLRAEDTAVYYCAKDLRITIFGVVIPDAFDIWGQGTMVTVSS
          ^^^^^^^^              ^^^^^^^^                       ^^^^^^^^^^^^^^^^^^
```

Therapeutic : Evolocumab

Best Alignment of Evolocumab heavy chain to a sequence from OAS
```
EVQLVQSGAEVKKPGASVKVSCKASGYTLTSYGISWVRQAPGQGLEWMGWVSFYNGNTNYAQKLQGRGTMTTDPSTSTAYMELRSLRSDDTAVYYCARGY------------GMDVWGQGTTVTVSS
.|||||||||||||||||||||||||||.|||||||||||||||||||||.|.||||||||||||||.||||||||||||||||||||||||||||||||............|||||||||||||||
QVQLVQSGAEVKKPGASVKVSCKASGYTFTSYGISWVRQAPGQGLEWMGWISAYNGNTNYAQKLQGRVTMTTDPSTSTAYMELRSLRSDDTAVYYCARGYYGSGSYSRYYYYGMDVWGQGTTVTVSS
          ^^^^^^^^              ^^^^^^^^                       ^^^^^^^^^^^^^^^^^^^^
```

Best Alignment of Evolocumab light chain to a sequence from OAS
```
ESALTQPASVSGSPGQSITISCTGTSSDVGGYNSVSWYQQHPGKAPKLMIYEVSNRPSGVSNRFSGSKSGNTASLTISGLQAEDEADYYCNSYTS--TSMVFGGGTKLTVL
.||||||||||||||||||||||||||||||||.||||||||||||||||||||||||||||||||||||||||||||||||||||||||||.||||..||||||||||||||
QSALTQPASVSGSPGQSITISCTGTSSDVGGYNYVSWYQQHPGKAPKLMIYEVSNRPSGVSNRFSGSKSGNTASLTISGLQAEDEADYYCSSYTSSSTSMVFGGGTKLTVL
          ^^^^^^^^^              ^^^                   ^^^^^^^^^^^
```

Best Alignment of Evolocumab heavy chain CDRs to a sequence from OAS
```
EVQLVQSGAEVKKPGASVKVSCKASGYTLTSYGISWVRQAPGQGLEWMGWVSFYNGNTNYAQKLQGRGTMTTDPSTSTAYMELRSLRSDDTAVYYCARGYGMDVWGQGTTVTVSS
..............||||||||||||||||||||||||||||||||||||.|.||||||||||||||.|||||.|||||||||||||||||||||||||||||||||||.||||.
--------------GASVKVSCKASGYTLTSYGISWVRQAPGQGLEWMGWISAYNGNTNYAQKLQGRVTMTTDTSTSTAYMELRSLRSDDTAVYYCARGYGMDVWGQGTLVTVS-
          ^^^^^^^^              ^^^^^^^^                       ^^^^^^^^
```

Best Alignment of Evolocumab light CDRs chain to a sequence from OAS
```
ESALTQPASVSGSPGQSITISCTGTSSDVGGYNSVSWYQQHPGKAPKLMIYEVSNRPSGVSNRFSGSKSGNTASLTISGLQAEDEADYYCNSYTSTSMVFGGGTKLTVL
..||||||.|||||.|.||.|..|.||||||||.||||||||||||.|||||||||||||||||.||||||||||||||||||||||||||||||.|||||||||||||
QPALTQPATVSGSPAQAITTSFPGSSSDVGGYNYVSWYQQHPGKAPILMIYEVSNRPSGVSNRFSASKSGNTASLTISGLQAEDEADYYCNSYTSSSMVFGGGTKLTVL
          ^^^^^^^^^              ^^^                   ^^^^^^^^^
```

Best Alignment of Evolocumab CDR-H3 chain to a sequence from OAS
```
EVQLVQSGAEVKKPGASVKVSCKASGYTLTSYGISWVRQAPGQGLEWMGWVSFYNGNTNYAQKLQGRGTMTTDPSTSTAYMELRSLRSDDTAVYYCARGYGMDVWGQGTTVTVSS
..............|.|.|.||.|||.|..||...|||||.|.||||........|.|.|.....||.|..........|....|||..||||||||||||||||||||||||||
--------------GESLKISCAASGFTFSSYDMHWVRQATGKGLEWVSAIGTA-GDTYYPGSVKGRFTISRENAKNSLYLQMNSLRAGDTAVYYCARGYGMDVWGQGTTVTVSS
          ^^^^^^^^              ^^^^^^^^                       ^^^^^^^^
```

Therapeutic : Farletuzumab

Best Alignment of Farletuzumab heavy chain to a sequence from OAS
```
EVQLVESGGGVVQPGRSLRLSCSASGFTFSGYGLSWVRQAPGKGLEWVAMISSGGSYTYYADSVKGRFAISRDNAKNTLFLQMDSLRPEDTGVYFCARHGDD---------PAWFAYWGQGTPVTVSS
.|||||||||||||||||||||.|||||||.|||.||||||||||||||.||..||..||||||||||.|||||.||||.|||.|||.|||.||.|||.||..........|..|.||||||.|||||
QVQLVESGGGVVQPGRSLRLSCVASGFTFSSYGLHWVRQAPGKGLEWVAVISYDGSNKYYADSVKGRFTISRDNSKNTLYLQMNSLRAEDTAVYYCARDGDNDILTGYYRPPYYFDYWGQGTLVTVSS
          ^^^^^^^^              ^^^^^^^^                       ^^^^^^^^^^^^^^^^^^^^
```

Best Alignment of Farletuzumab light chain to a sequence from OAS
DIQLTQSPSSLSASVGDRVTITCSVSSSISSNNLHWYQQKPGKAPKPWIYGTSNLASGVPSRFSGSGSGTDYTFTISSLQPEDIATYYCQQWSSYPYMYTFGQGTKVEIK
|||.|||||||||||||||||||..|..|.||.|.|||||||||||..||..|||..||||||||||||||.|||||||||||||||||||..|||.|||||||||.|||
DIQMTQSPSSLSASVGDRVTITCQASQDI-SNYLNWYQQKPGKAPKLLIYDASNLETGVPSRFSGSGSGTDFTFTISSLQPEDIATYYCQQFNSYPPMYTFGQGTKLEIK
                       ^^^^^^^            ^^^                        ^^^^^^^^^^^

Best Alignment of Farletuzumab heavy chain CDRs to a sequence from OAS
EVQLVESGGGVVQPGRSLRLSCSASGFTFSGYGLSWVRQAPGKGLEWVAMISSGGSYTYYADSVKGRFAISRDNAKNTLFLQMDSLRPEDTGVYFCARHGDDPAWFAYWGQGTPVTVSS
||..|||||..|.||.||.|||.|||||||.||.|||||.|.|.|||||.||||||||||.|||||||.||||||||||.|||.||..|||..|.|||||||||||||||||||.||||.
EVKVVESGGDLVKPGGSLKLSCAASGFTFSSYGMSWVRQTPDKRLEWVATISSGGSYTYYPDSVKGRFTISRDNAKNTLYLQMSSLKSEDTAMYYCARHGDDPAWFAYWGQGTLVTVSA
                      ^^^^^^^^           ^^^^^^^^                         ^^^^^^^^^^^^

Best Alignment of Farletuzumab light CDRs chain to a sequence from OAS
DIQLTQSPSSLSASVGDRVTITCSVSSSISSNNLHWYQQKPGKAPKPWIYGTSNLASGVPSRFSGSGSGTDYTFTISSLQPEDIATYYCQQWSSYPYMYTFGQGTKVEIK
.|.|||||....||.|..||.||||||||||.||||||||....|||||||||||||||.|||||||||.|..||||...||.|||||||||||||.|||||.|||.|||
EIVLTQSPALMAASPGEKVTFTCSVSSSISSSNLHWYQQKSETSPKPWIYGTSNLASGVPVRFSGSGSGTSYSLTISSMEAEDAATYYCQQWSSYPLMYTFGGGTKLEIK
                       ^^^^^^^            ^^^                        ^^^^^^^^^^^

Best Alignment of Farletuzumab CDR-H3 chain to a sequence from OAS
EVQLVESGGGVVQPGRSLRLSCSASGFTFSGYGLSWVRQAPGKGLEWVAMISSGGSYTYYADSVKGRFAISRDNAKNTLFLQMDSLRPEDTGVYFCARHGDDPAWFAYWGQGTPVTVSS
|||.||||||.|.||.||.|||.|||||||.|..|||||.|.|.|||||.||.||..|||.|||||||.||||||||||.|||.|||.|||..|.|||||||||||||||||||.||||.
EVQRVESGGGLVKPGGSLKLSCAASGFTFSSYTMSWVRQTPEKRLEWVATISGGGGNTYYPDSVKGRFTISRDNAKNTLYLQMSSLRSEDTALYYCARHGDDPAWFAYWGQGTLVTVSA
                      ^^^^^^^^           ^^^^^^^^                         ^^^^^^^^^^^^

Therapeutic : Fasinumab

Best Alignment of Fasinumab heavy chain to a sequence from OAS
QVQLVQSGAEVKKPGASVKVSCKVSGFTLTELSIHWVRQAPGKGLEWMGGFDPEDGETIYAQKFQGRVTMTEDTSTDTAYMELTSLRSEDTAVYYCSTIFGVVTNFDNWGQGTLVTVSS
||||||||||||||||||||||||||.||||||.||||||||||||||||||||||||||||||||||||||||||||||||||.||||||||||||.|.|||||.||.|||||||||||
QVQLVQSGAEVKKPGASVKVSCKVSGYTLTELSMHWVRQAPGKGLEWMGGFDPEDGETIYAQKFQGRVTMTEDTSTDTAYMELSSLRSEDTAVYYCATAFGVVTLFDYWGQGTLVTVSS
                      ^^^^^^^^           ^^^^^^^^                         ^^^^^^^^^^^^

Best Alignment of Fasinumab light chain to a sequence from OAS
DIQMTQSPSSLSASAGDRVTITCRASQAIRNDLGWYQQKPGKAPKRLIYAAFNLQSGVPSRFSGSGSGTEFTLTISSLQPEDLASYYCQQYNRYPWTFGQGTKVEIK
|||||||||||||.||||||||||||.|||||||||||.||||||||||.||||||||||||||||||||||||||||||||.|.|||||||.||||||||||||||
DIQMTQSPSSLSASVGDRVTITCRASQGIRNDLGWYQQKSGKAPKRLIYAASNLQSGVPSRFSGSGSGTEFTLTISSLQPEDFATYYCQQYNSYPWTFGQGTKVEIK
                       ^^^^^^           ^^^                       ^^^^^^^^^

Best Alignment of Fasinumab heavy chain CDRs to a sequence from OAS
QVQLVQSGAEVKKPGASVKVSCKVSGFTLTELSIHWVRQAPGKGLEWMGGFDPEDGETIYAQKFQGRVTMTEDTSTDTAYMELTSLRSEDTAVYYCSTIFGVVTNFDNWGQGTLVTVSS
................|||.||||||.||||||.|||||||||||||||||||||||||||||||||||||||||||||||||.||||||||||||.||||||||||.|||||||||||
----------------SVKASCKVSGYTLTELSMHWVRQAPGKGLEWMGGFDPEDGETIYAQKFQGRVTMTEDTSTDTAYMELSSLRSEDTAVYYCVTIFGVVTNFDYWGQGTLVTVSS
                      ^^^^^^^^           ^^^^^^^^                         ^^^^^^^^^^^^

Best Alignment of Fasinumab light CDRs chain to a sequence from OAS
DIQMTQSPSSLSASAGDRVTITCRASQAIRNDLGWYQQKPGKAPKRLIYAAFNLQSGVPSRFSGSGSGTEFTLTISSLQPEDLASYYCQQYNRYPWTFGQGTKVEIK
|||||||||||||.|||||||||||||||||.|.||||||||.|||.||..|||||||||||.|||||||||||||||.|.|.|||||||.||||||||||||||
DIQMTQSPSSLSASIGDRVTITCRASQAIRNDLDWFQQKPGKAPQRLISAASSLQSGVPSRFSGRGSGTEFTLTISSLQPDDFATYYCQQYNSYPWTFGQGTKVEIK
                       ^^^^^^           ^^^                       ^^^^^^^^^

Best Alignment of Fasinumab CDR-H3 chain to a sequence from OAS
QVQLVQSGAEVKKPGASVKVSCKVSGFTLTELSIHWVRQAPGKGLEWMGGFDPEDGETIYAQKFQGRVTMTEDTSTDTAYMELTSLRSEDTAVYYCSTIFGVVTNFDNWGQGTLVTVSS
................|...||..||||.......|.||||||||||............||....||.|...|......|....||..|||||||||||.||||.||||||||||||||
----------------SLRLSCAASGFTFSDHYMSWIRQAPGKGLEWISYISGSGRTIYYADSVKGRFTISRDNAKNSLYLQMNSLGAEDTAVYYCSTIVGVVTIFDNWGQGTLVTVSS
                      ^^^^^^^^           ^^^^^^^^                         ^^^^^^^^^^^^

Therapeutic : Fezakinumab

Best Alignment of Fezakinumab heavy chain to a sequence from OAS
QVQLVQSGAEVKKPGASVKVSCKASGYTFTNYYMHWVRQAPGQGLEWVGWINPYTGSAFYAQKFRGRVTMTRDTSISTAYMELSRLRSDDTAVYYCAREPEKF---DSDDSDVWGRGTLVTVSS
||||||||||||||||||||||||||||||.||||||||||||||||.|||||..|...|||||.||||||||||||||||||||||||||||||||||||||.....|....||||.||.|||||
QVQLVQSGAEVKKPGASVKVSCKASGYTFTGYYMHWVRQAPGQGLEWMGWINPNSGGTNYAQKFQGRVTMTRDTSISTAYMELSRLRSDDTAVYYCAREPELEYPSDYYGMDVWGQGTTVTVSS
                      ^^^^^^^^           ^^^^^^^^                         ^^^^^^^^^^^^^^^^^

Best Alignment of Fezakinumab light chain to a sequence from OAS
QAVLTQPPSVSGAPGQRVTISCTGSSSNIGAGYGVHWYQQLPGTAPKLLIYGDSNRPSGVPDRFSGSKSGTSASLAITGLQAEDEADYYCQSYDNSLSGYVFGGGTQLTVL
|.||||||||||||||||||||||||||||||||.||||||||||||||||||||.|||||||||||||||||||||||||||||||||||||||||||||||||||||.|||||||||||
QSVLTQPPSVSGAPGQRVTISCTGSSSNIGAGYDVHWYQQLPGTAPKLLIYGNSNRPSGVPDRFSGSKSGTSASLAITGLQAEDEADYYCQSYDNSLSGWVFGGGTQLTVL
                      ^^^^^^^^^            ^^^                        ^^^^^^^^^^^

Best Alignment of Fezakinumab heavy chain CDRs to a sequence from OAS
QVQLVQSGAEVKKPGASVKVSCKASGYTFTNYYMHWVRQAPGQGLEWVGWINPYTGSAFYAQKFRGRVTMTRDTSISTAYMELSRLRSDDTAVYYCAREPEKFDSDDSDVWGRGTLVTVSS
................||||||||||||||.|||||||||||||||||.|.|||..||..|||||.|||||||||||.||.|||||.|||..|||||||||...|.|||.||||.||.|||||
----------------SVKVSCKASGYTFTGYYMHWVRQAPGQGLEWMGIINPSGGSTSYAQKFQGRVTMTRDTSTSTVYMELSSLRSEETAVYYCARERDDFWSDDMDVWGKGTTVTVSS
                      ^^^^^^^^           ^^^^^^^^                         ^^^^^^^^^^^^^^

Best Alignment of Fezakinumab light CDRs chain to a sequence from OAS
QAVLTQPPSVSGAPGQRVTISCTGSSSNIGAGYGVHWYQQLPGTAPKLLIYGDSNRPSGVPDRFSGSKSGTSASLAITGLQAEDEADYYCQSYDNSLSGYVFGGGTQLTVL
|.||||.||||||||||||||||||||||||..|||||||||||||||||||||||||||||||||||||||||||||||||||||||||||||||||||||||||||.||..|||
QSVLTQSPSVSGAPGQRVTISCTGSSSNIGAGYDIHWYQQLPGTAPKLLIYGDSNRPSGVPDRFSGSKSGTSASLAITGLQAEDEADYYCQSYDNSLSGYVFGTGTKVTVL
                      ^^^^^^^^^            ^^^                        ^^^^^^^^^^^

Best Alignment of Fezakinumab CDR-H3 chain to a sequence from OAS
QVQLVQSGAEVKKPGASVKVSCKASGYTF--TNYYMHWVRQAPGQGLEWVGWINPYTGSAFYAQKFRGRVTMTRDTSISTAYMELSRLRSDDTAVYYCAREPEKFDSDDSDVWGRGTLVTVSS
....................|..||..|..|.||..||||.||.||||...|....||..|....|.||.|..||........||......||||||||.||.|.|||.||||.|.||||||
--------------SETLSLTCTVSGGSFASTTYYWGWVRQSPGKGLEWIAHIHYR-GSTYYKSSLRSRVIMSVDTFKNQFSLKLSSVTAAATAVYYCARDPEPFFSDDGDVWGQGILVTVSS
                      ^^^^^^^^^^           ^^^^^^^^                         ^^^^^^^^^^^^^^

Therapeutic : Ficlatuzumab

Best Alignment of Ficlatuzumab heavy chain to a sequence from OAS
QVQLVQPGAEVKKPGTSVKLSCKASGYTFTTYWMHWVRQAPGQGLEWIGEINPTNGHTNYNQKFQGRATLTVDKSTSTAYMELSSLRSEDTAVYYCARNYVG---SIFDYWGQGTLLTVSS
||||||.||||||||.|||.||||||||||.|.||||||||||||||.|.|||..|.|||.||||||.|.|.|.|||||||||||||||||||||||||..|.....||||||||||.||||
QVQLVQSGAEVKKPGASVKVSCKASGYTFTGYYMHWVRQAPGQGLEWMGWINPNSGGTNYAQKFQGRVTITADESTSTAYMELSSLRSEDTAVYYCARSGVPYGDYIFDYWGQGTLVTVSS
                      ^^^^^^^^           ^^^^^^^^                         ^^^^^^^^^^^^^^

Best Alignment of Ficlatuzumab light chain to a sequence from OAS
DIVMTQSPDSLAMSLGERVTLNCKASENVVSYVSWYQQKPGQSPKLLIYGASNRESGVPDRFSGSGSATDFTLTISSVQAEDVADYHCGQSYNYPYTFGQGTKLEIK
.|||||||.|..||.||||||.||||||||..||||||||||||||||||||||||..||||||.|||||||||||||||||||.||||||||||.|||||||.||||||||
NIVMTQSPKSMSMSVGERVTLSCKASENVGTYVSWYQQKPGQSPKLLIYGASNRYTGVPDRFTGSGSATDFTLTISSVQAEDLADYHCGQSYSYPYTFGGGTKLEIK
                       ^^^^^^           ^^^                       ^^^^^^^^^

Best Alignment of Ficlatuzumab heavy chain CDRs to a sequence from OAS
QVQLVQPGAEVKKPGTSVKLSCKASGYTFTTYWMHWVRQAPGQGLEWIGEINPTNGHTNYNQKFQGRATLTVDKSTSTAYMELSSLRSEDTAVYYCARNYVGSIFDYWGQGTLLTVSS
||||.|||.|..|||.||||||||||||||.||||||.|.||||||||||.|||.||.||||.||...||||||||.|||||.||||.|||.|||||||.|.||||||||||||.|||||

QVQLQQPGTELVKPGASVKLSCKASGYTFTSYWMHWVKQRPGQGLEWIGNINPSNGGTNYNEKFKSKATLTVDKSSSTAYMQLSSLTSEDSAVYYCARAYYGSIFDYWGQGTTLTVSS
^^^^^^^^ ^^^^^^^^ ^^^^^^^^^^^

Best Alignment of Ficlatuzumab light CDRs chain to a sequence from OAS
DIVMTQSPDSLAMSLGERVTLNCKASENVVSYVSWYQQKPGQSPKLLIYGASNRESGVPDRFSGSGSATDFTLTISSVQAEDVADYHCGQSYNYPYTFGQGTKLEIK
.|||||||.|..||.|||||||.|||||||.|||||||||..|||||.|||||||..||||||||.||||||||||||||||||||.|||||||||.||||||.|||||||
NIVMTQSPKSMSMSVGERVTLSCKASENVGSYVSWYQQKTEQSPKLVIYGASNRYTGVPDRFTGSGSATDFTLTISSVQAEDLADYHCGQSYSYPYTFGGGTKLEIK
^^^^^^ ^^^ ^^^^^^^^^

Best Alignment of Ficlatuzumab CDR-H3 chain to a sequence from OAS
QVQLVQPGAEVKKPGTSVKLSCKASGYTFTTYWMHWVRQAPGQGLEWIGEINPTNGHTNYNQKFQGRATLTVDKSTSTAYMELSSLRSEDTAVYYCARNYVGSIFDYWGQGTLLTVSS
||...|.|.....|..|....|..||...|.|..|||||.||.||||.|.|....|.|.||..|..|.....|.|.|.......||...|||.|||||||.|||||||||||.|||||
QVHVKQSGPGLVQPSQSLSITCTVSGFSLTSYGVHWVRQSPGKGLEWLGVIWSG-GSTDYNAAFISRLSISKDNSKSQVFFKMNSLQADDTAIYYCARNYYGSIFDYWGQGTTLTVSS
^^^^^^^^ ^^^^^^^^ ^^^^^^^^^^^

Therapeutic : Figitumumab

Best Alignment of Figitumumab heavy chain to a sequence from OAS
EVQLLESGGGLVQPGGSLRLSCTASGFTFSSYAMNWVRQAPGKGLEWVSAISGSGGTTFYADSVKGRFTISRDNSRTTLYLQMNSLRAEDTAVYYCAKDLGWSDS---------YYYYYGMDVWGQGTTVTVSS
||||||||||||||||||||||.|||||||||||.|||||||||||||||||||||.|.||||||||||||||||..||||||||||||||||||||||||||..|.........||||||||||||||||||||
EVQLLESGGGLVQPGGSLRLSCAASGFTFSSYAMSWVRQAPGKGLEWVSAISGSGGSTYYADSVKGRFTISRDNSKNTLYLQMNSLRAEDTAVYYCAKDLGWYSSSWFSGAIGPYYYYYGMDVWGQGTTVTVSS
^^^^^^^^ ^^^^^^^^ ^^^^^^^^^^^^^^^^^^^^^^^^^^

Best Alignment of Figitumumab light chain to a sequence from OAS
DIQMTQFPSSLSASVGDRVTITCRASQGIRNDLGWYQQKPGKAPKRLIYAASRLHRGVPSRFSGSGSGTEFTLTISSLQPEDFATYYCLQHNSYPCSFGQGTKLEIK
||||||.|||||||||||||||||||||||||||||||||||||||||||||.|..|||||||||||||||||||||||||||||||||||||||||||||||||||
DIQMTQSPSSLSASVGDRVTITCRASQGIRNDLGWYQQKPGKAPKRLIYAASSLQSGVPSRFSGSGSGTEFTLTISSLQPEDFATYYCLQHNSYPCSFGQGTKLEIK
^^^^^^ ^^^ ^^^^^^^^^

Best Alignment of Figitumumab heavy chain CDRs to a sequence from OAS
EVQLLESGGGLVQPGGSLRLSCTASGFTFSSYAMNWVRQAPGKGLEWVSAISGSGGTTFYADSVKGRFTISRDNSRTTLYLQMNSLRAEDTAVYYCAKDLGWSDSYYYYYGMDVWGQGTTVTVSS
..............||||||||.|||||||||||.|||||||||||||||||||||.|.||||||||||||||||..||||||||||||||||||||||||.|.|||||||||||||||||||||
--------------GGSLRLSCAASGFTFSSYAMSWVRQAPGKGLEWVSAISGSGGSTYYADSVKGRFTISRDNSKNTLYLQMNSLRAEDTAVYYCAKDLGSSSSYYYYYGMDVWGQGTTVTVSS
^^^^^^^^ ^^^^^^^^ ^^^^^^^^^^^^^^^^^

Best Alignment of Figitumumab light CDRs chain to a sequence from OAS
DIQMTQFPSSLSASVGDRVTITCRASQGIRNDLGWYQQKPGKAPKRLIYAASRLHRGVPSRFSGSGSGTEFTLTISSLQPEDFATYYCLQHNSYPCSFGQGTKLEIK
||.|||.|||||||||||||||||||||||||||||||||||.||||||||||.||.||||||||.|||||||||||||||||||.||||||||||||||||||||.|||
DIRMTQSPSSLSASVGDRVTITCRASQGIRNDLGWYQQKPGQAPKRLIYAASSLHSGVPSRFSGGGSGTEFTLTISSLQTEDFATYYCLQHNSYPCSFGQGTKVEIK
^^^^^^ ^^^ ^^^^^^^^^

Best Alignment of Figitumumab CDR-H3 chain to a sequence from OAS
EVQLLESGGGLVQPGGSLRLSCTASGFTFSSYAMNWVRQAPGKGLEWVSAISGSGGTTFYADSVKGRFTISRDNSRTTLYLQMNSLRAEDTAVYYCAKDLGWSDSYYYYYGMDVWGQGTTVTVSS
..............||||||||.|||||||||||.|||||||||||||||||||||.|.||||||||||||||||..||||||||||||||||||||||||.|.|||||||||||||||||||||
--------------GGSLRLSCAASGFTFSSYAMSWVRQAPGKGLEWVSAISGSGGSTYYADSVKGRFTISRDNSKNTLYLQMNSLRAEDTAVYYCAKDLGSSSSYYYYYGMDVWGQGTTVTVSS
^^^^^^^^ ^^^^^^^^ ^^^^^^^^^^^^^^^^^

Therapeutic : Fletikumab

Best Alignment of Fletikumab heavy chain to a sequence from OAS
QVQLVQSGAEVKRPGASVKVSCKASGYTFTNDIIHWVRQAPGQRLEWMGWINAGYGNTQYSQNFQDRVSITRDTSASTAYMELISLRSEDTAVYYCAREPLWFGESSPHDYYGMDVWGQGTTVTVSS
||||||||||||.|||||||||||||||||....||||||||||||||||||||||.|||.|||.||.||.||||||||||||||||||.||||||||||||||..||||||....|||||||||||||||||
QVQLVQSGAEVKKPGASVKVSCKASGYTFTSYAMHWVRQAPGQRLEWMGWINAGNGNTKYSQKFQGRVTITRDTSASTAYMELSSLRSEDTAVYYCARDLLWFGES-NYYYYGMDVWGQGTTVTVSS
^^^^^^^^ ^^^^^^^^ ^^^^^^^^^^^^^^^^^^^

Best Alignment of Fletikumab light chain to a sequence from OAS
AIQLTQSPSSLSASVGDRVTITCRASQGISSALAWYQQKPGKAPKLLIYDASSLESGVPSRFSGSGSGTDFTLTISSLQPEDFATYYCQQFNSYPLTFGGGTKVEIK
|||||||||||||||||||||||||||||||||||||||||||||||||||||||||||||||||||||||||||||||||||||||||||||||||||||||||||
AIQLTQSPSSLSASVGDRVTITCRASQGISSALAWYQQKPGKAPKLLIYDASSLESGVPSRFSGSGSGTDFTLTISSLQPEDFATYYCQQFNSYPLTFGGGTKVEIK
^^^^^^ ^^^ ^^^^^^^^^

Best Alignment of Fletikumab heavy chain CDRs to a sequence from OAS
QVQLVQSGAEVKRPGASVKVSCKASGYTFTNDIIHWVRQAPGQRLEWMGWINAGYGNTQYSQNFQDRVSITRDTSASTAYMELISLRSEDTAVYYCAREPLWFGESSPHDYYGMDVWGQGTTVTVSS
................||||||||||||||....||||||||||||||||||||.|||.|||.||.||.||||||||||||||.|||||||||||||||.|||||.|||.||||||||||||||||.
----------------SVKVSCKASGYTFTSYAMHWVRQAPGQRLEWMGWINAGNGNTKYSQKFQGRVTITRDTSASTAYMELSSLRSEDTAVYYCAREGLWFGELSPHYYYGMDVWGQGTTVTVS-
^^^^^^^^ ^^^^^^^^ ^^^^^^^^^^^^^^^^^^^

Best Alignment of Fletikumab light CDRs chain to a sequence from OAS
AIQLTQSPSSLSASVGDRVTITCRASQGISSALAWYQQKPGKAPKLLIYDASSLESGVPSRFSGSGSGTDFTLTISSLQPEDFATYYCQQFNSYPLTFGGGTKVEIK
|||||||||||||||||||||||||||||||||||||||||||||||||||||||||||||||||||||||||||||||||||||||||||||||||||.||||.||
AIQLTQSPSSLSASVGDRVTITCRASQGISSALAWYQQKPGKAPKLLIYDASSLESGVPSRFSGSGSGTDFTLTISSLQPEDFATYYCQQFNSYPLTFGPGTKVDIK
^^^^^^ ^^^ ^^^^^^^^^

Best Alignment of Fletikumab CDR-H3 chain to a sequence from OAS
QVQLVQSGAEVKRPGASVKVSCKASGYTFTNDIIHWVRQAPGQRLEWMGWINAGYGNTQYSQNFQDRVSITRDTSASTAYMELISLRSEDTAVYYCAREPLWFGESSPHDYYGMDVWGQGTTVTVSS
.....|||||.|.||.||||||||||.||||....||||||||.||.||.|...|....|...|..||.||.|.|...||.||||.||.||||||.|||||..|||||||||.||||||||||||.|||
-----QSGAEAKKPGSSVKVSCKASGGTFTNYAFSWVRQAPGQGLEGMGGIIPTYNSAHYARRFVGRVTITADEATSTVYMELSSLTSEDTAVCYCAREQRWFGESSPHDDYGMDVWGQGTTVIVSS
^^^^^^^^ ^^^^^^^^ ^^^^^^^^^^^^^^^^^^^

Therapeutic : Foralumab

Best Alignment of Foralumab heavy chain to a sequence from OAS
QVQLVESGGGVVQPGRSLRLSCAASGFKFSGYGMHWVRQAPGKGLEWVAVIWYDGSKKYYVDSVKGRFTISRDNSKNTLYLQMNSLRAEDTAVYYCARQMGY-----------WHFDLWGRGTLVTVSS
|||||||||||||||||||||||||||.||.|||||||||||||||||||||||||.|||.|||||||||||||||||||||||||||||||||||||||..||...........|.||||||||||||||
QVQLVESGGGVVQPGRSLRLSCAASGFTFSSYGMHWVRQAPGKGLEWVAVIWYDGSNKYYADSVKGRFTISRDNSKNTLYLQMNSLRAEDTAVYYCARESGYDFWSGYYGSSYWYFDLWGRGTLVTVSS
^^^^^^^^ ^^^^^^^^ ^^^^^^^^^^^^^^^^^^^^^

Best Alignment of Foralumab light chain to a sequence from OAS
EIVLTQSPATLSLSPGERATLSCRASQSVSSYLAWYQQKPGQAPRLLIYDASNRATGIPARFSGSGSGTDFTLTISSLEPEDFAVYYCQQRSNWPPLTFGGGTKVEIK
||||||||||||||||||||||||||||||||||||||||||||||||||||||||||||||||||||||||||||||||||||||||||||||||||||||||||||
EIVLTQSPATLSLSPGERATLSCRASQSVSSYLAWYQQKPGQAPRLLIYDASNRATGIPARFSGSGSGTDFTLTISSLEPEDFAVYYCQQRSNWPPLTFGGGTKVEIK
^^^^^^ ^^^ ^^^^^^^^^^

Best Alignment of Foralumab heavy chain CDRs to a sequence from OAS
QVQLVESGGGVVQPGRSLRLSCAASGFKFSGYGMHWVRQAPGKGLEWVAVIWYDGSKKYYVDSVKGRFTISRDNSKNTLYLQMNSLRAEDTAVYYCARQMGYWHFDLWGRGTLVTVSS
.............||||||||||||||.||.|||||||||||||||||||||||||||||.||||||||||||||||||||||||||.||||||||||..|||.|||||||.||||||
-------------PGRSLRLSCAASGFTFSIYGMHWVRQAPGKGLEWVAVIWYDGSKKYYADSVKGRFTISRDNSKNTLYLQMNSLRVEDTAVYYCARGNGYWYFDLWGRGALVTVSS
^^^^^^^^ ^^^^^^^^ ^^^^^^^^^^^

Best Alignment of Foralumab light CDRs chain to a sequence from OAS
EIVLTQSPATLSLSPGERATLSCRASQSVSSYLAWYQQKPGQAPRLLIYDASNRATGIPARFSGSGSGTDFTLTISSLEPEDFAVYYCQQRSNWPPLTFGGGTKVEIK
|||||||||||||||||||||||||||||||||||||||||||||||||||||||||||||||||||||||||||||.||||||.|||||||||||||||||||||||
EIVLTQSPATLSLSPGERATLSCRASQSVSSYLAWYQQKPGQAPRLLIYDASNRATGIPARFSGSGSGTDFTLTISSIEPEDFAIYYCQQRSNWPPLTFGGGTKVEIK
^^^^^^ ^^^ ^^^^^^^^^^

Best Alignment of Foralumab CDR-H3 chain to a sequence from OAS
QVQLVESGGGVVQPGRSLRLSCAASGFKFSGYGMHWVRQAPGKGLEWVAVIWYDGSKKYYVDSVKGRFTISRDNSKNTLYLQMNSLRAEDTAVYYCARQMGYWHFDLWGRGTLVTVSS
................|.|.|..||...||....|.||.|||||||...|.|......|..|.|.|.|.|.|.|||...|...|..|.||||||||||||||.||||||||||||||

--------------SETLSLTCTVSGGSISGHSWSWIRQPPGKGLEWIGYIYYS-GSANYGPSLKSRVTLSVDTSKNQFSLNLASVTAADTAVYYCARQMGYWYFDLWGRGTLVTVSS
                    ^^^^^^^^                ^^^^^^^^                                    ^^^^^^^^^^^

Therapeutic : Foravirumab

Best Alignment of Foravirumab heavy chain to a sequence from OAS
QVQLVESGGGAVQPGRSLRLSCAASGFTFSSYGMHWVRQAPGKGLEWVAVILYDGSDKFYADSVKGRFTISRDNSKNTLYLQMNSLRAEDTAVYYCAKVAVAGTHFDYWGQGTLVTVSS
||||||||||.||||||||||||||||||||||||||||||||||||||||.||||.|.|||||||||||||||||||||||||||||||||||||||||..|.|||||||||||||||
QVQLVESGGGVVQPGRSLRLSCAASGFTFSSYGMHWVRQAPGKGLEWVAVIWYDGSNKYYADSVKGRFTISRDNSKNTLYLQMNSLRAEDTAVYYCAKVAPSGYHFDYWGQGTLVTVSS
                    ^^^^^^^^                ^^^^^^^^                                    ^^^^^^^^^^^^

Best Alignment of Foravirumab light chain to a sequence from OAS
DIQMTQSPSSLSASVGDRVTITCRASQGIRNDLGWYQQKPGKAPKLLIYAASSLQSGVPSRFSGSGSGTDFTLTISSLQPEDFATYYCQQLNSYPPTFGGGTKVEIK
.||||||||||||||||||||||||||||||||||||||||||||||||||||||||||||||||||||||||||||||||||||||||||||||.|||||||||||
AIQMTQSPSSLSASVGDRVTITCRASQGIRNDLGWYQQKPGKAPKLLIYAASSLQSGVPSRFSGSGSGTDFTLTISSLQPEDFATYYCQQLNSYPLTFGGGTKVEIK
                     ^^^^^^                ^^^                                 ^^^^^^^^^

Best Alignment of Foravirumab heavy chain CDRs to a sequence from OAS
QVQLVESGGGAVQPGRSLRLSCAASGFTFSSYGMHWVRQAPGKGLEWVAVILYDGSDKFYADSVKGRFTISRDNSKNTLYLQMNSLRAEDTAVYYCAKVAVAGTHFDYWGQGTLVTVSS
................|||||||||||||||||.|||||||||||||||||.||||.|.|||||||||||||||||||||||||||||||||||||||||||||.||||||||||||||
----------------SLRLSCAASGFTFSSYGTHWVRQAPGKGLEWVAVISYDGSNKYYADSVKGRFTISRDNSKNTLYLQMNSLRAEDTAVYYCAKVAVAGTDFDYWGQGTLVTVSS
                    ^^^^^^^^                ^^^^^^^^                                    ^^^^^^^^^^^^

Best Alignment of Foravirumab light CDRs chain to a sequence from OAS
DIQMTQSPSSLSASVGDRVTITCRASQGIRNDLGWYQQKPGKAPKLLIYAASSLQSGVPSRFSGSGSGTDFTLTISSLQPEDFATYYCQQLNSYPPTFGGGTKVEIK
.|||||||||||.||||||||||||||||||||||.||||.|||..||||||||.||||||||.|||||.|||||.|||||||||||||||||||||||||||||||
-IQMTQSPSSLSPSVGDRVTITCRASQGIRNDLGWSQQKPVKAPNVLIYAASSLESGVPSRFSASGSGTEFTLTIGSLQPEDFATYYCQQLNSYPPTFGGGTKVEIK
                     ^^^^^^                ^^^                                 ^^^^^^^^^

Best Alignment of Foravirumab CDR-H3 chain to a sequence from OAS
QVQLVESGGGAVQPGRSLRLSCAASGFTFSSYGMHWVRQAPGKGLEWVAVILYDGSDKFYADSVKGRFTISRDNSKNTLYLQMNSLRAEDTAVYYCAKVAVAGTHFDYWGQGTLVTVSS
..............|.||..||..||..|.||...||||.|||||||...|....||..|..|..|..|||.|.|..|.|||..||.|.|||.||||||||||||||||||||||||||
--------------GESLKISCKGSGYSFTSYWIGWVRQMPGKGLEWMGIIYPGDSDTRYSPSFQGQVTISADKSISTAYLQWSSLKASDTAMYYCAKVAVAGTHFDYWGQGTLVTVSS
                    ^^^^^^^^                ^^^^^^^^                                    ^^^^^^^^^^^^

Therapeutic : Fremanezumab

Best Alignment of Fremanezumab heavy chain to a sequence from OAS
EVQLVESGGGLVQPGGSLRLSCAASGFTFSNYWISWVRQAPGKGLEWVAEIRSESDASATHYAEAVKGRFTISRDNAKNSLYLQMNSLRAEDTAVYYCLAYFDYG---LAIQNYWGQGTLVTVSS
||||||||||||||||||||||||||||||.||.||||||||||||||.|......|...|...|||||||||||||||||||||||||||||||||...||||........||||||||||||
EVQLVESGGGLVQPGGSLRLSCAASGFTFSSYWMSWVRQAPGKGLEWVANIKQD--GSEKYYVDSVKGRFTISRDNAKNSLYLQMNSLRAEDTAVYYCARDFDYGGNSPSYFDYWGQGTLVTVSS
                    ^^^^^^^^                ^^^^^^^^^^                                    ^^^^^^^^^^^^^^^^

Best Alignment of Fremanezumab light chain to a sequence from OAS
EIVLTQSPATLSLSPGERATLSCKASKRVTTYVSWYQQKPGQAPRLLIYGASNRYLGIPARFSGSGSGTDFTLTISSLEPEDFAVYYCSQSYN-YPYTFGQGTKLEIK
|||||||||||||||||||||||.||..||||..|||||||||||||||.||||..||||||||||||||||||||||||||||||||.|..|..|||||||||||||
EIVLTQSPATLSLSPGERATLSCRASQSVTTYLAWYQQKPGQAPRLLIYDASNRATGIPARFSGSGSGTDFTLTISSLEPEDFAVYYCQQRSNWPPYTFGQGTKLEIK
                     ^^^^^^                ^^^                                 ^^^^^^^^^^

Best Alignment of Fremanezumab heavy chain CDRs to a sequence from OAS
EVQLVESGGGLVQPGGSLRLSCAASGFTFSNYWISWVRQAPGKGLEWVAEIRSESDASATHYAEAVKGRFTISRDNAKNSLYLQMNSLRAEDTAVYYCLAYFDYGLAIQNYWGQGTLVTVSS
..|||||||||||||||..|||.||||||||||..||||.|.|||||||.||..||..||||||.||||||||||..|.|.|||||.||||||..|||.||.||.||...|||||||||||.
--QLVESGGGLVQPGGSMKLSCVASGFTFSNYWMNWVRQSPAKGLEWVAQIRLKSDNYATHYAESVKGRFTISRDDSKSSVYLQMNNLRAEDTGIYYCTAYYDYDLAWFAYWGQGTLVTVSA
                    ^^^^^^^^                ^^^^^^^^^^                                    ^^^^^^^^^^^^^

Best Alignment of Fremanezumab light CDRs chain to a sequence from OAS
EIVLTQSPATLSLSPGERATLSCKASKRVTTYVSWYQQKPGQAPRLLIYGASNRYLGIPARFSGSGSGTDFTLTISSLEPEDFAVYYCSQSYNYPYTFGQGTKLEIK
.||.||||...|.|.|||..|.|..||.|.|||.||||||.|.|.||||||||||.|.|.||.||||.|.|||||||...||.|.|.|.|||.||||||.|||||||
-IVMTQSPKSMSMSVGERDSLICQSSKNVRTYVHWYQQKPEQSPKLLIYGASNRYTGVPDRFTGSGSATYFTLTISSVQAEDLADYHCRQSYSYPYTFGAGTKLEIK
                     ^^^^^^                ^^^                                 ^^^^^^^^^

Best Alignment of Fremanezumab CDR-H3 chain to a sequence from OAS
EVQLVESGGGLVQPGGSLRLSCAASGFTFSNYWISWVRQAPGKGLEWVAEIRSESDASATHYAEAVKGRFTISRDNAKNSLYLQMNSLRAEDTAVYYCLAYFDYGLAIQNYWGQGTLVTVSS
................||||||.||||||..|..||||||||.|.|||..|.......|..||..|||||||.||.||||.||..||||||||||.||||..|||...|.||||||||||||
----------------SLRLSCKASGFTFRTYAMSWVRQAPGQGPEWVSDISTS--GTAKYYAHTVKGRFTITRDDAKNSPYLRINSLRAEDTAVFYCLAGPDYGDYGQDYWGQGTLVTVSS
                    ^^^^^^^^                ^^^^^^^^^^                                    ^^^^^^^^^^^^^

Therapeutic : Fresolimumab

Best Alignment of Fresolimumab heavy chain to a sequence from OAS
QVQLVQSGAEVKKPGSSVKVSCKASGYTFSSNVISWVRQAPGQGLEWMGGVIPIVDIANYAQRFKGRVTITADESTSTTYMELSSLRSEDTAVYYCASTLGLV-----LDAMDYWGQGTLVTVSS
||||||||||||||||||||||||||.||||..|||||||||||||||||.|||...|||||.|.|||||||||||||.||||||||||||||||||.|||.......||..|||||||||||||
QVQLVQSGAEVKKPGSSVKVSCKASGGTFSSYAISWVRQAPGQGLEWMGGIIPIFGTANYAQKFQGRVTITADESTSTAYMELSSLRSEDTAVYYCARTLGYSGYDPDLDNFDYWGQGTLVTVSS
                    ^^^^^^^^                ^^^^^^^^                                    ^^^^^^^^^^^^^^^^^^

Best Alignment of Fresolimumab light chain to a sequence from OAS
ETVLTQSPGTLSLSPGERATLSCRASQSLGSSYLAWYQQKPGQAPRLLIYGASSRAPGIPDRFSGSGSGTDFTLTISRLEPEDFAVYYCQQYADSPITFGQGTRLEIK
||||||||||||||||||||||||||||..||||||||||||||||||||||||||.|||||||||||||||||||||||||||||||||||..||||||||||||||
ETVLTQSPGTLSLSPGERATLSCRASQSVSSSYLAWYQQKPGQAPRLLIYGASSRATGIPDRFSGSGSGTDFTLTISRLEPEDFAVYYCQQYGSSPITFGQGTRLEIK
                     ^^^^^^^                ^^^                                 ^^^^^^^^^

Best Alignment of Fresolimumab heavy chain CDRs to a sequence from OAS
QVQLVQSGAEVKKPGSSVKVSCKASGYTFSSNVISWVRQAPGQGLEWMGGVIPIVDIANYAQRFKGRVTITADESTSTTYMELSSLRSEDTAVYYCASTLGLVLDAMDYWGQGTLVTVSS
................||||||||||.||||..|||||||||||||||||.|||..||||||.|.||||||||.||||.||||||||.|||||||||||.|..|||.|.|||||.|||||
----------------SVKVSCKASGGTFSSYAISWVRQAPGQGLEWMGGIIPILGIANYAQKFQGRVTITADKSTSTAYMELSSLRAEDTAVYYCASTYGDYLDAFDIWGQGTMVTVSS
                    ^^^^^^^^                ^^^^^^^^                                    ^^^^^^^^^^^^^

Best Alignment of Fresolimumab light CDRs chain to a sequence from OAS
ETVLTQSPGTLSLSPGERATLSCRASQSLGSSYLAWYQQKPGQAPRLLIYGASSRAPGIPDRFSGSGSGTDFTLTISRLEPEDFAVYYCQQYADSPITFGQGTRLEIK
|.||||||||||||||||||||||||||||||||||||||||||||||||||||||.||||||||||||||||||||.|||||||||||||||.||.|||||||..||
EIVLTQSPGTLSLSPGERATLSCRASQSLGSSYLAWYQQKPGQAPRLLIYGASSRATGIPDRFSGSGSGTDFTLTINRLEPEDFAVYYCQQYASSPWTFGQGTRVDIK
                     ^^^^^^^                ^^^                                 ^^^^^^^^^

Best Alignment of Fresolimumab CDR-H3 chain to a sequence from OAS
QVQLVQSGAEVKKPGSSVKVSCKASGYTFSSNVISWVRQAPGQGLEWMGGVIPIVDIANYAQRFKGRVTITADESTSTTYMELSSLRSEDTAVYYCASTLGLVLDAMDYWGQGTLVTVSS
.|.||.||.|..|||.|||.|||||||.|.|....||.|.||.||||.|...|.....||...|||..|.|||....|.||.||.|.||...||.|||||.|||.|||||||||.|||||
-VKLVESGPELVKPGASVKISCKASGYAFTSSWVNWVKQRPGKGLEWIGLIYPGGGATNYNGKFKGQATLTADKYSRTAYMQLSRLTSEASVVYFCASTLRLVLYAMDYWGQGTSVTVSS
                    ^^^^^^^^                ^^^^^^^^                                    ^^^^^^^^^^^^^

Therapeutic : Fulranumab

Best Alignment of Fulranumab heavy chain to a sequence from OAS
EVQLVESGGGLVQPGGSLRLSCAASGFTLRSYSMNWVRQAPGKGLEWVSYISRSSHTIFYADSVKGRFTISRDNAKNSLYLQMDSLRDEDTAMYYCARVYSSGWHVSDYFDYWGQGILVTVSS
||||||||||||||||||||||||||||..||||||||||||||||||||||.||.||.||||||||||||||||||||||||.||||||||.|||||.|||||.|..||||||||.||||||
EVQLVESGGGLVQPGGSLRLSCAASGFTFSSYSMNWVRQAPGKGLEWVSYISSSSSTIYYADSVKGRFTISRDNAKNSLYLQMNSLRDEDTAVYYCARGYSSGWYVVYYFDYWGQGTLVTVSS
                    ^^^^^^^^                ^^^^^^^^                                    ^^^^^^^^^^^^^^^^

Best Alignment of Fulranumab light chain to a sequence from OAS

```
AIQLTQSPSSLSASVGDRVTITCRASQGISSALAWYQQKPGKAPKLLIYDASSLESGVPSRFSGSGSGTDFTLTISSLQPEDFATYYCQQFNSYPLTFGGGTKVEIK
|||||||||||||||||||||||||||||||||||||||||||||||||||||||||||||||||||||||||||||||||||||||||||||||||||||||||||
AIQLTQSPSSLSASVGDRVTITCRASQGISSALAWYQQKPGKAPKLLIYDASSLESGVPSRFSGSGSGTDFTLTISSLQPEDFATYYCQQFNSYPLTFGGGTKVEIK
                         ^^^^^^          ^^^                      ^^^^^^^^^
```

Best Alignment of Fulranumab heavy chain CDRs to a sequence from OAS

```
EVQLVESGGGLVQPGGSLRLSCAASGFTLRSYSMNWVRQAPGKGLEWVSYISRSSHTIFYADSVKGRFTISRDNAKNSLYLQMDSLRDEDTAMYYCARVYSSGWHVSDYFDYWGQGILVTVSS
..............||||||||||||||..||||||||||||||||||||||.||.||.||||||||||||||||||||||||.|||.||||.|||||.|||||..||||||||||.||||||
--------------GGSLRLSCAASGFTFSSYSMNWVRQAPGKGLEWVSYISSSSSTIYYADSVKGRFTISRDNAKNSLYLQMNSLRAEDTAVYYCARAYSSGWYPSDYFDYWGQGTLVTVSS
                         ^^^^^^^^        ^^^^^^^^                  ^^^^^^^^^^^^^^^^
```

Best Alignment of Fulranumab light CDRs chain to a sequence from OAS

```
AIQLTQSPSSLSASVGDRVTITCRASQGISSALAWYQQKPGKAPKLLIYDASSLESGVPSRFSGSGSGTDFTLTISSLQPEDFATYYCQQFNSYPLTFGGGTKVEIK
|||||||||||||||||||||||||||||||||||||||||||||||||||||||||||||||||||||||||||||||||||||||||||||||||||.||||.||
AIQLTQSPSSLSASVGDRVTITCRASQGISSALAWYQQKPGKAPKLLIYDASSLESGVPSRFSGSGSGTDFTLTISSLQPEDFATYYCQQFNSYPLTFGPGTKVDIK
                         ^^^^^^          ^^^                      ^^^^^^^^^
```

Best Alignment of Fulranumab CDR-H3 chain to a sequence from OAS

```
EVQLVESGGGLVQPGGSLRLSCAASGFTL--RSYSMNWVRQAPGKGLEWVSYISRSSHTIFYADSVKGRFTISRDNAKNSLYLQMDSLRDEDTAMYYCARVYSSGWHVSDYFDYWGQGILVTVSS
.................|.|.|..||......||...|.||.|||||||...|..|.....|..|.|.|.|||.|..||...|...|....|||.|||||||||||.|||||||||||.||||||
--------------SETLSLTCTVSGGSISSSSYYWGWIRQPPGKGLEWIGSIYYS-GSTYYNPSLKSRVTISVDTSKNQFSLKLSSVTAADTAVYYCARVYSSGWYVSDYFDYWGQGTLVTVSS
                         ^^^^^^^^^^        ^^^^^^^^                  ^^^^^^^^^^^^^^^^
```

Therapeutic : Futuximab

Best Alignment of Futuximab heavy chain to a sequence from OAS

```
EVQLQQPGSELVRPGASVKLSCKASGYTFTSYWMHWVKQRPGQGLEWIGNIYPGSRSTNYDEKFKSKATLTVDTSSSTAYMQLSSLTSEDSAVYYCTRNGDYYVSSGDAMDYWGQGTSVTVSS
.||||||||||||||||||||||||||||||||||||||||||||||||||||||.||||||||||||||||||||||||||||||||||||||||||.|.||.||.||||||||||||||||
QVQLQQPGSELVRPGASVKLSCKASGYTFTSYWMHWVKQRPGQGLEWIGNIYPGSGSTNYDEKFKSKATLTVDTSSSTAYMQLSSLTSEDSAVYYCTRRGSYYGSSYDAMDYWGQGTSVTVSS
                         ^^^^^^^^        ^^^^^^^^                  ^^^^^^^^^^^^^^^^
```

Best Alignment of Futuximab light chain to a sequence from OAS

```
DIQMTQTTSSLSASLGDRVTISCRTSQDIGNYLNWYQQKPDGTVKLLIYYTSRLHSGVPSRFSGSGSGTDFSLTINNVEQEDVATYFCQHYNTVPPTFGGGTKLEIK
||||||||||||||||||||||||.||||.||||||||||||||||||||||||||||||||||||||||.||||.|.|||||||||||..||.|||||||||||||
DIQMTQTTSSLSASLGDRVTISCRASQDISNYLNWYQQKPDGTVKLLIYYTSRLHSGVPSRFSGSGSGTDYSLTISNLEQEDVATYFCQQGNTLPPTFGGGTKLEIK
                         ^^^^^^          ^^^                      ^^^^^^^^^
```

Best Alignment of Futuximab heavy chain CDRs to a sequence from OAS

```
EVQLQQPGSELVRPGASVKLSCKASGYTFTSYWMHWVKQRPGQGLEWIGNIYPGSRSTNYDEKFKSKATLTVDTSSSTAYMQLSSLTSEDSAVYYCTRNGDYYVSSGDAMDYWGQGTSVTVSS
.||||||||||||||||||||||||||||||||||||||||||||||||||||||.||||||||||||||||||||||||||||||||||||||||||.|.||.|||||||||||||||||||
-VQLQQPGSELVRPGASVKLSCKASGYTFTSYWMHWVKQRPGQGLEWIGNIYPGSGSTNYDEKFKSKATLTVDTSSSTAYMQLSSLTSEDSAVYYCTRWGYYYGSSGDAMDYWGQGTSVTVSS
                         ^^^^^^^^        ^^^^^^^^                  ^^^^^^^^^^^^^^^^
```

Best Alignment of Futuximab light CDRs chain to a sequence from OAS

```
DIQMTQTTSSLSASLGDRVTISCRTSQDIGNYLNWYQQKPDGTVKLLIYYTSRLHSGVPSRFSGSGSGTDFSLTINNVEQEDVATYFCQHYNTVPPTFGGGTKLEIK
||||||..||||||.|||.||.||.||||.|||.||||||....|.||||||.|..|||||||||.||||..|||.....||.|||.||||||.|||||||||.|.|
DIQMTQSPSSLSASVGDRITITCRASQDINNYLSWYQQKPGKAPKPLIYYTSSLETGVPSRFSGSRSGTDYTLTISSLQPEDIATYYCQHYNTLPPTFGGGTKGELK
                         ^^^^^^          ^^^                      ^^^^^^^^^
```

Best Alignment of Futuximab CDR-H3 chain to a sequence from OAS

```
EVQLQQPGSELVRPGASVKLSCKASGYTFTSYWMHWVKQRPGQGLEWIGNIYPGSRSTNYDEKFKSKATLTVDTSSSTAYMQLSSLTSEDSAVYYCTRNGDYYVSSGDAMDYWGQGTSVTVSS
.|||||.|..|.|||||||.|||.||||||||||||||||||.||||||.||||...|.|..|||.||.||..|.|.||||.|||||.||||||||||.||||.|||.|||||||||||||||
-VQLQQSGTVLARPGASVKMSCKTSGYTFTSYWMHWVKQRPGPGLEWIGAIYPGNSDTSYNQKFKGKAKLTAVTSASTAYMELSSLTNEDSAVYYCTRSGDYYGSSGGAMDYWGQGTSVTVSS
                         ^^^^^^^^        ^^^^^^^^                  ^^^^^^^^^^^^^^^^
```

Therapeutic : Galcanezumab

Best Alignment of Galcanezumab heavy chain to a sequence from OAS

```
QVQLVQSGAEVKKPGSSVKVSCKASGYTFGNYWMQWVRQAPGQGLEWMGAIYEGTGKTVYIQKFADRVTITADKSTSTAYMELSSLRSEDTAVYYCARLSDY-VSGFGYWGQGTTVTVSS
|||||||||||||||.|||||||||||||..|.|.||||||||||||||.|....|.|.|.|||..||||||||||||||||||||||||||||||||||.|....|.||||||.|||||
QVQLVQSGAEVKKPGASVKVSCKASGYTFTSYYMHWVRQAPGQGLEWMGIINPSGGSTSYAQKFQGRVTITADKSTSTAYMELSSLRSEDTAVYYCARLSGYNKYYFDYWGQGTLVTVSS
                         ^^^^^^^^        ^^^^^^^^                  ^^^^^^^^^^^^^
```

Best Alignment of Galcanezumab light chain to a sequence from OAS

```
DIQMTQSPSSLSASVGDRVTITCRASKDISKYLNWYQQKPGKAPKLLIYYTSGYHSGVPSRFSGSGSGTDFTLTISSLQPEDFATYYCQQGDALPPTFGGGTKVEIK
||||||||||||||||||||||||||..||.||||||||||||||||||..|...|||||||||||||||||||||||||||||||||||.|.||||||||||||||
DIQMTQSPSSLSASVGDRVTITCRASQSISSYLNWYQQKPGKAPKLLIYAASSLQSGVPSRFSGSGSGTDFTLTISSLQPEDFATYYCQQYDNLPPTFGGGTKVEIK
                         ^^^^^^          ^^^                      ^^^^^^^^^
```

Best Alignment of Galcanezumab heavy chain CDRs to a sequence from OAS

```
QVQLVQSGAEVKKPGSSVKVSCKASGYTFGNYWMQWVRQAPGQGLEWMGAIYEGTGKTVYIQKFADRVTITADKSTSTAYMELSSLRSEDTAVYYCARLSDYVSGFGYWGQGTTVTVSS
||||.|||||...||.|||.|||||||||.|||..|..|.||.||||.|.|.||.|.|.|.|.|..||....|.|||||.|||||..|||.|||||.|||||||||..||.|||||||.||||
QVQLQQSGAELVRPGTSVKMSCKASGYTFTNYWIGWAKQRPGHGLEWIGDIYPGGGYTNYNEKFKGKATLTADKSSSTAYMQFSSLTSEDSAIYYCARLSDYDGGFDYWGQGTTLTVSS
                         ^^^^^^^^        ^^^^^^^^                  ^^^^^^^^^^^^
```

Best Alignment of Galcanezumab light CDRs chain to a sequence from OAS

```
DIQMTQSPSSLSASVGDRVTITCRASKDISKYLNWYQQKPGKAPKLLIYYTSGYHSGVPSRFSGSGSGTDFTLTISSLQPEDFATYYCQQGDALPPTFGGGTKVEIK
||||||..||||||.||||||.||||.|||||||||||||....||||||||..||||||||||||||||..||||.|..||.||.||||.|||.|||.|||.|||
DIQMTQTTSSLSASLGDRVTISCRASQDISKYLNWYQQKPDGTVKLLIYYTSRLHSGVPSRFSGSGSGTDYSLTISNLEQEDIATYFCQQGNALPLTFGAGTKLEIK
                         ^^^^^^          ^^^                      ^^^^^^^^^
```

Best Alignment of Galcanezumab CDR-H3 chain to a sequence from OAS

```
QVQLVQSGAEVKKPGSSVKVSCKASGYTFGNYWMQWVRQAPGQGLEWMGAIYEGTGKTVYIQKFADRVTITADKSTSTAYMELSSLRSEDTAVYYCARLSDYVSGFGYWGQGTTVTVSS
................|...||.|||.|...|.|.|||||||.||||...|..|.....|......|.||..|......|....|.|.||||||||||..|||||||||||||.|||||
----------------SLRLSCAASGFTLSSYSMNWVRQAPGKGLEWVASITTGGSYKYYGDSVKGRFTISRDNAKNSLYLQMNSMRAEDTAVYYCARSGDYVSGFGYWGQGTLVTVSS
                         ^^^^^^^^        ^^^^^^^^                  ^^^^^^^^^^^^
```

Therapeutic : Galiximab

Best Alignment of Galiximab heavy chain to a sequence from OAS

```
QVQLQESGPGLVKPSETLSLTCAVSGGSISGGYGWGWIRQPPGKGLEWIGSFYSSSGNTYYNPSLKSQVTISTDTSKNQFSLKLNSMTAADTAVYYCVRDRLFSVVGMVYNNWFDVWGPGVLVTVSS
||||||||||||||||||||||||||||.|||||||||||||||||||||..||.||||||||||||.||||||||||||||||.|.||||||||||.|...........|||||||||||||||||
QVQLQESGPGLVKPSETLSLTCAVSGGSVSGGYGWGWIRQPPGKGLEWIGTIYSGSGNTYYNPSLKSRVTISTDTSKNQFSLKLTSVTAADTAVYYCAREPFSRY---SWNNWFDVWGPGVLVTVSS
                         ^^^^^^^^^       ^^^^^^^^                  ^^^^^^^^^^^^^^^^^^^
```

Best Alignment of Galiximab light chain to a sequence from OAS

```
ESALTQPPSVSGAPGQKVTISCTGSTSNIGGYDLHWYQQLPGTAPKLLIYDINKRPSGISDRFSGSKSGTAASLAITGLQTEDEADYYCQSYDSSLNAQVFGGGTRLTVL
.|.||||||||||||||||||||||.|||||||.|||||||||||||||||.||||||||||||||||||.|||||||||||||||||||||||||||||||||||||||
QSVLTQPPSVSGAPGQKVTISCTGSSSNIGGYDVHWYQQLPGTAPKLLIYDNNKRPSGISDRFSGSKSGTSASLAITGLQTEDEADYYCQSYDSSLNAQVFGGGTRLTVL
                         ^^^^^^^^        ^^^                      ^^^^^^^^^^^
```

Best Alignment of Galiximab heavy chain CDRs to a sequence from OAS

```
QVQLQESGPGLVKPSETLSLTCAVSGGSISGGYGWGWIRQPPGKGLEWIGSFYSSSGNTYYNPSLKSQVTISTDTSKNQFSLKLNSMTAADTAVYYCVRDRLFSVVGMVYNNWFDVWGPGVLVTVSS
||||||||||.||||||||||||||||||||||.|.||||||||||||||.....||||.|.|||||.||||.|||||||||.|.|.||||||||||.|||.....|..|.||.||||||||.||||
```

QVQLQESGPGLLKPSETLSLTCAVSGGSISGGYAWSWIRQPPGKGLEWIGFIGGYSGNTDYDPSLKSRVTISKDTSKNQFSLNLSSVTAADTAVYYCARDRGYCSGGVCYGNWVDVWGPGVLATVSS
^^^^^^^^^ ^^^^^^^^ ^^^^^^^^^^^^^^^^^^^

Best Alignment of Galiximab light CDRs chain to a sequence from OAS
ESALTQPPSVSGAPGQKVTISCTGSTSNIGGYDLHWYQQLPGTAPKLLIYDINKRPSGISDRFSGSKSGTAASLAITGLQTEDEADYYCQSYDSSLNAQVFGGGTRLTVL
.|.||||||||||||||||||||||.|||||||.||||||||||||||||||||||||.|||||||||||.||||||||||||||||||||||||||||.|||.||.||||
QSVLTQPPSVSGAPGQKVTISCTGSSSNIGGYDVHWYQQLPGTAPKLLIYDINKRPSGVSDRFSGSKSGTSASLAITGLQTEDEADYYCQSYDSSLNAHVFGSGTKLTVL
^^^^^^^^ ^^^ ^^^^^^^^^^^

Best Alignment of Galiximab CDR-H3 chain to a sequence from OAS
QVQLQESGPGLVKPSETLSLTCAVSGGSISGGYGWGWIRQPPGKGLEWIGSFYSSSGNTYYNPSLKSQVTISTDTSKNQFSLKLNSMTAADTAVYYCVRDRLFSVVGMVYNNWFDVWGPGVLVTVSS
....................|..||......|...|.||.||.||||.|........|.|.......||..|||..|.....|.|....|||||||||..|..|||.|.|||||.||.|.||||||
----------------SVKVSCKASGYRF-STYSITWVRQAPGQGLEWMGWISPYNRDTKYARKFQGRVTMTTDTITNTAYMELRSLRSDDTAVYYCVRGALVGVVGGVQNNWFDPWGQGTLVTVSS
^^^^^^^^^ ^^^^^^^^ ^^^^^^^^^^^^^^^^^^^

Therapeutic : Ganitumab

Best Alignment of Ganitumab heavy chain to a sequence from OAS
QVQLQESGPGLVKPSGTLSLTCAVSGGSISSSNWWSWVRQPPGKGLEWIGEIYHSGSTNYNPSLKSRVTISVDKSKNQFSLKLSSVTAADTAVYYCARWTGR-TDAFDIWGQGTMVTVSS
||||||||||||||||||||||||||||||||||||||||||||||||||||||||||||||||||||||||||||||||||||||||||||||||||.|.|..||||||||||||||||
QVQLQESGPGLVKPSGTLSLTCAVSGGSISSSNWWSWVRQPPGKGLEWIGEIYHSGSTNYNPSLKSRVTISVDKSKNQFSLKLSSVTAADTAVYYCARDTEREGDAFDIWGQGTMVTVSS
^^^^^^^^^ ^^^^^^^ ^^^^^^^^^^^^^

Best Alignment of Ganitumab light chain to a sequence from OAS
DVVMTQSPLSLPVTPGEPASISCRSSQSLLHSNGYNYLDWYLQKPGQSPQLLIYLGSNRASGVPDRFSGSGSGTDFTLKISRVEAEDVGVYYCMQGTHWPLTFGQGTKVEIK
|.|||||||||||||||||||||||||||||||||||||||||||||||||||||||||||||||||||||||||||||||||||||||||||||||||||||||||||||
DIVMTQSPLSLPVTPGEPASISCRSSQSLLHSNGYNYLDWYLQKPGQSPQLLIYLGSNRASGVPDRFSGSGSGTDFTLKISRVEAEDVGVYYCMQGTHWPLTFGQGTKVEIK
^^^^^^^^^^^ ^^^ ^^^^^^^^^

Best Alignment of Ganitumab heavy chain CDRs to a sequence from OAS
QVQLQESGPGLVKPSGTLSLTCAVSGGSISSSNWWSWVRQPPGKGLEWIGEIYHSGSTNYNPSLKSRVTISVDKSKNQFSLKLSSVTAADTAVYYCARWTGRTDAFDIWGQGTMVTVSS
..............|.||||||||||||||||||||||||||||||||||||||||||||||||||||||||||||||||||||||||||.||.||||||||||||||||
--------------SETLSLTCAVSGGSISSSNWWSWVRQPPGKGLEWIGEIYHSGSTNYNPSLKSRVTISVDKSKNQFSLKLSSVTAADTAVYYCARWPGRYDAFDIWGQGTMVTVSS
^^^^^^^^^ ^^^^^^^ ^^^^^^^^^^^^

Best Alignment of Ganitumab light CDRs chain to a sequence from OAS
DVVMTQSPLSLPVTPGEPASISCRSSQSLLHSNGYNYLDWYLQKPGQSPQLLIYLGSNRASGVPDRFSGSGSGTDFTLKISRVEAEDVGVYYCMQGTHWPLTFGQGTKVEIK
|.||||||||||||||||||||||||||||||||||||||||||||||||||||||||||||||||||||||||||||||..||||||||||||||||||||||.|||||||
DIVMTQSPLSLPVTPGEPASISCRSSQSLLHSNGYNYLDWYLQKPGQSPQLLIYLGSNRASGVPDRFSGSGSGTDFTLKITSVEAEDVGVYYCMQGTHWPLTFGGGTKVEIK
^^^^^^^^^^^ ^^^ ^^^^^^^^^

Best Alignment of Ganitumab CDR-H3 chain to a sequence from OAS
QVQLQESGPGLVKPSGTLSLTCAVSGGSIS-SSNWWSWVRQPPGKGLEWIGEIYHSGSTNYNPSLKSRVTISVDKSKNQFSLKLSSVTAADTAVYYCARWTGRTDAFDIWGQGTMVTVSS
..............|.||||||.|||||||.||..|.|.|||||||||.||.||.||||.||||||||||||||.||.||.|||||||||||||||||||||.||||||||||||||||
--------------SETLSLTCTVSGGSISSSSYYWGWLRQPPGKGLEGIGSIYYSGSTYYNPSLKSRVTISVDTSKYQFYLKLSSVTAADTAVYYCARWTGGTDAFDIWGQGTMVTVSS
^^^^^^^^^^ ^^^^^^^ ^^^^^^^^^^^^

Therapeutic : Gantenerumab

Best Alignment of Gantenerumab heavy chain to a sequence from OAS
QVELVESGGGLVQPGGSLRLSCAASGFTFSSYAMSWVRQAPGKGLEWVSAINASGTRTYYADSVKGRFTISRDNSKNTLYLQMNSLRAEDTAVYYCARGKGNTHKPYGYVRYFDVWGQGTLVTVSS
.|.|||||||||||||||||||||||||||||||||||||||||||||||..||..|||||||||||||||||||||||||||||||||||||||||||||..|.....|..|||.|||||||||||
EVQLVESGGGLVQPGGSLRLSCAASGFTFSSYAMSWVRQAPGKGLEWVSAISGSGGSTYYADSVKGRFTISRDNSKNTLYLQMNSLRAEDTAVYYCARGKPDTR---TYKYYFDYWGQGTLVTVSS
^^^^^^^^ ^^^^^^^^ ^^^^^^^^^^^^^^^^^^^

Best Alignment of Gantenerumab light chain to a sequence from OAS
DIVLTQSPATLSLSPGERATLSCRASQSVSSSYLAWYQQKPGQAPRLLIYGASSRATGVPARFSGSGSGTDFTLTISSLEPEDFATYYCLQIYNMPITFGQGTKVEIK
.|||||||||||||||||||||||||||||||||||||||||||||||||||||||||.|||||||||||||||||||||||||||||||.|.||.|.|||||||||||
EIVLTQSPATLSLSPGERATLSCRASQSVSSSYLAWYQQKPGQAPRLLIYGASSRATGIPARFSGSGSGTDFTLTISSLEPEDFATYYCQQNYNAPWTFGQGTKVEIK
^^^^^^^ ^^^ ^^^^^^^^^

Best Alignment of Gantenerumab heavy chain CDRs to a sequence from OAS
QVELVESGGGLVQPGGSLRLSCAASGFTFSSYAMSWVRQAPGKGLEWVSAINASGTRTYYADSVKGRFTISRDNSKNTLYLQMNSLRAEDTAVYYCARGKGNTHKPYGYVRYFDVWGQGTLVTVSS
............|||||||||||||||||||||||||||||||||||.|||.||..||||.||||||||||.||||||||||||||||.||||......|.|.||.|||||||.||||.|||
------------PGGSLRLSCAASGFTFSSYAMSWVRQAPGKGLEWVSAISASGSRTHFADSVAGRFTISRDNSRNTLYLQMNSLRAEDTAIYYCAKDRVYSHDPSGYPRYFDVWGRGTLVAVSS
^^^^^^^^ ^^^^^^^^ ^^^^^^^^^^^^^^^^^^^

Best Alignment of Gantenerumab light CDRs chain to a sequence from OAS
DIVLTQSPATLSLSPGERATLSCRASQSVSSSYLAWYQQKPGQAPRLLIYGASSRATGVPARFSGSGSGTDFTLTISSLEPEDFATYYCLQIYNMPITFGQGTKVEIK
.||||||.||||||||||||||||||||||||||||||.||||||||||.||||.||||.|||||||||||||.||||||||..|||||||||||.||.||||||||..|||
EIVLTQSPGTLSLSPGERATLSCRASQSVSSSYLAWYRQKPGQAPRLLMYGASTRATGIPARFSGSGSGTEFTLTISSLQSEDFATYYCLQDYNYPITFGQGTRLEIK
^^^^^^^ ^^^ ^^^^^^^^^

Best Alignment of Gantenerumab CDR-H3 chain to a sequence from OAS
QVELVESGGGLVQPGGSLRLSCAASGFTFSSYAMSWVRQAPGKGLEWVSAINASGTRTYYADSVKGRFTISRDNSKNTLYLQMNSLRAEDTAVYYCARGKGNTHKPYGYVRYFDVWGQGTLVTVSS
................|.|.||..|...|.|..||.||.|||||||...||.|...|.|..|.|.|.|||.|.|||...|...|..|.||||||||||.|.....||||||||.||.||||||||
--------------SETLSLTCAVYGGSYSGYYWSWIRQPPGKGLEWIGEINHS-GSTNYNPSLKSRVTISVDTSKNQFSLKLSSVTAADTAVYYCARGRGRSGYSYGYVRYFDLWGRGTLVTVSS
^^^^^^^^ ^^^^^^^^ ^^^^^^^^^^^^^^^^^^^

Therapeutic : Gedivumab

Best Alignment of Gedivumab heavy chain to a sequence from OAS
EVQLVESGGGVVQPGKSLRLSCAASGLTFSSYAVHWVRQAPGKGLEWVTLISYDGANQYYADSVKGRFTISRDNSKNTVYLQMNSLRPEDTAVYYCAVPGPVFGI--FPPWSYFDNWGQGILVTVSS
.|||||||||||||.||||||||||.||||||.||||||||||||||..|||||.|.|||||||||||||||||||||.|||||||||.|||||||||......|...|||.||||.||||.||||||
QVQLVESGGGVVQPGRSLRLSCAASGFTFSSYAMHWVRQAPGKGLEWVAVISYDGSNKYYADSVKGRFTISRDNSKNTLYLQMNSLRAEDTAVYYCARDAYGWGGYDFPPRSYFDYWGQGTLVTVSS
^^^^^^^^ ^^^^^^^^ ^^^^^^^^^^^^^^^^^^^^

Best Alignment of Gedivumab light chain to a sequence from OAS
EIVLTQSPATLSVSPGERATLSCRASQVISHNLAWYQQKPGQAPRLLIYGASTRASGIPARFSGSGSGTDYTLTITSLQSEDFAVYYCQHYSNWPPRLTFGGGTKVEIK
|||.||||||||||||||||||||||||..|.||||||||||||||||||||||||||.|||||||||||||||.|||||||||||||||||||.||||||||||||||||||||
EIVMTQSPATLSVSPGERATLSCRASQSVSSNLAWYQQKPGQAPRLLIYGASTRATGIPARFSGSGSGTDFTLTITSLQSEDFAVYYCQQYSNWPPRLTFGGGTKVEIK
^^^^^^ ^^^ ^^^^^^^^^^^

Best Alignment of Gedivumab heavy chain CDRs to a sequence from OAS
EVQLVESGGGVVQPGKSLRLSCAASGLTFSSYAVHWVRQAPGKGLEWVTLISYDGANQYYADSVKGRFTISRDNSKNTVYLQMNSLRPEDTAVYYCAVPGPVFGIFPPWSYFDNWGQGILVTVSS
..............|.||||||||||.||||||.||||||||||||||.||||||.|.||||||||||||||||||||.|||||||||.|||||||||.||..||.|.|..|||.||||.||||||
--------------GRSLRLSCAASGFTFSSYAMHWVRQAPGKGLEWVALISYDGSNKYYADSVKGRFTISRDNSKNTLYLQMNSLRAEDTAVYYCARPGSWFGEFIPGYYFDYWGQGTLVTVSS
^^^^^^^^ ^^^^^^^^ ^^^^^^^^^^^^^^^^^^

Best Alignment of Gedivumab light CDRs chain to a sequence from OAS
EIVLTQSPATLSVSPGERATLSCRASQVISHNLAWYQQKPGQAPRLLIYGASTRASGIPARFSGSGSGTDYTLTITSLQSEDFAVYYCQHYSNWPPRLTFGGGTKVEIK
|||.||||.||||||||.|||||||||.||.|||||||||||.||||.|||||||.||||||.||||||..||||.|.||.|.|||||||||.|||||.|||.||...||
EIVMTQSPDTLSVSPGETATLSCRASQSISNNLAWYQQKPGQSPRLLFYGASTRATGIPARFRGSGSGTEFTLTISSIQSGDCAVYYCQHYNNWPPRITFGQGTRLAIK
^^^^^^ ^^^ ^^^^^^^^^^^
Best Alignment of Gedivumab CDR-H3 chain to a sequence from OAS
EVQLVESGGGVVQPGKSLRLSCAASGLTFSSYAVHWVRQAPGKGLEWVTLISYDGANQYYADSVKGRFTISRDNSKNTVYLQMNSLRPEDTAVYYCAVPGPVFGIFPPWSYFDNWGQGILVTVSS
............||.||..||||||.||..||.|||||||||.||||||.|||.|...|||||||||||||||||.|.||||.|||.||.||||||..||.|||.|..|.||.||||..|.|||

-------------PGESLKISCAASGFTFNNYAMHWVRQAPGQGLEWVTVISYNGNDKYYADSVKGRFTISRDNSKDTLYLQMRSLRAEDSAVYYCATGGPIFGIRPITSAFDIWGQGTMVSVSS
^^^^^^^^ ^^^^^^^^ ^^^^^^^^^^^^^^^^^

Therapeutic : Gemtuzumab

Best Alignment of Gemtuzumab heavy chain to a sequence from OAS
EVQLVQSGAEVKKPGSSVKVSCKASGYTITDSNIHWVRQAPGQSLEWIGYIYPYNGGTDYNQKFKNRATLTVDNPTNTAYMELSSLRSEDTAFYYCVNGNPWLAYWGQGTLVTVSS
||||..||.|..|||.|||.||||||||.||.||||..|.|||||||||||||||||.|||||||||||||||...|||||.|.||.|||.|.|||.||..|.||||||||||||.
EVQLQESGPELVKPGASVKISCKASGYTFTDNNIHWLKQSPGQSLEWIGYIYPYNGGTGYNQKFKNRATLTVDDSSNTAYMDLRSLTSEDSAVYYCANGYAWFAYWGQGTLVTVSA
^^^^^^^^ ^^^^^^^^ ^^^^^^^^^

Best Alignment of Gemtuzumab light chain to a sequence from OAS
DIQLTQSPSTLSASVGDRVTITCRASESLDNYGIRFLTWFQQKPGKAPKLLMYAASNQGSGVPSRFSGSGSGTEFTLTISSLQPDDFATYYCQQTKEVPWSFGQGTKVEVK
|||.||||||||||||||||||||||.|........|||.||||||||||.|.|||..||||||||||||||||||||||||||||||||||||.|..||.||||||||||
DIQMTQSPSTLSASVGDRVTITCRASQSI----SSWLTWYQQKPGKAPKLLIYKASNLESGVPSRFSGSGSGTEFTLTISSLQPDDFATYYCQQYKSYPWTFGQGTKVEVK
^^^^^^^^^^ ^^^ ^^^^^^^^^

Best Alignment of Gemtuzumab heavy chain CDRs to a sequence from OAS
EVQLVQSGAEVKKPGSSVKVSCKASGYTITDSNIHWVRQAPGQSLEWIGYIYPYNGGTDYNQKFKNRATLTVDNPTNTAYMELSSLRSEDTAFYYCVNGNPWLAYWGQGTLVTVSS
.|||.|||.|..|||.|||.||||||||.||.|.|||.|..|.|||||||||||||||.||||||..|||||||...||||||.||.|||.|.|||.|||.|.|||||||||||||.
-VQLQQSGPELVKPGASVKISCKASGYTFTDYNMHWVKQSHGKSLEWIGYIYPYNGGTGYNQKFKSKATLTVDNSSSTAYMELRSLTSEDSAVYYCANGNAWFAYWGQGTLVTVSA
^^^^^^^^ ^^^^^^^^ ^^^^^^^^^

Best Alignment of Gemtuzumab light CDRs chain to a sequence from OAS
DIQLTQSPSTLSASVGDRVTITCRASESLDNYGIRFLTWFQQKPGKAPKLLMYAASNQGSGVPSRFSGSGSGTEFTLTISSLQPDDFATYYCQQTKEVPWSFGQGTKVEVK
||.|||||..|..|.|.|.||.||||||.|||||||..|||||.|..||||.|||||||||||.|||||||||.|.|.|.....||.|.|.|||.|||||.||.|||.|.|
DIVLTQSPASLAVSLGQRATISCRASESVDNYGIRFMNWFQQKTGQPPKLLIYAASNQGSGVPARFSGSGSGTDFSLNIHPMEEDDTAMYFCQQSKEVPWTFGGGTKLEIK
^^^^^^^^^^ ^^^ ^^^^^^^^^

Best Alignment of Gemtuzumab CDR-H3 chain to a sequence from OAS
EVQLVQSGAEVKKPGSSVKVSCKASGYTITDSNIHWVRQAPGQSLEWIGYIYPYNGGTDYNQKFKNRATLTVDNPTNTAYMELSSLRSEDTAFYYCVNGNPWLAYWGQGTLVTVSS
.|||.|||||..|||.|||.||.|||..|.|...|||.|.|.|.|||||.|.|.||.|.|..||...||.|.|...||||..||||.|||||.|||||||||.||||||||||||.
-VQLQQSGAELVKPGASVKLSCTASGFNIKDTYMHWVKQRPEQGLEWIGRIDPANGNTKYDPKFQGKATITADTSSNTAYLQLSSLTSEDTAVYYCVNGNPWFAYWGQGTLVTVSA
^^^^^^^^ ^^^^^^^^ ^^^^^^^^^

Therapeutic : Gevokizumab

Best Alignment of Gevokizumab heavy chain to a sequence from OAS
QVQLQESGPGLVKPSQTLSLTCSFSGFSLSTSGMGVGWIRQPSGKGLEWLAHIWWDGDESYNPSLKSRLTISKDTSKNQVSLKITSVTAADTAVYFCARNRYDPPWFVDWGQGTLVTVSS
||.|.|||||..|||||||||||||||||||||||||||||||||||||||||||||.|..|||||||.||||||||.|||.||||||..||||.|.|||.|||.|||..||||||||||.
QVTLKESGPGILKPSQTLSLTCSFSGFSLSTSGMGVGWIRQPSGKGLEWLAHIWWDDDKYYNPSLKSQLTISKDTSRNQVFLKITSVDTADTATYYCARRRYDGPWFAYWGQGTLVTVSA
^^^^^^^^^^ ^^^^^^^ ^^^^^^^^^^^^

Best Alignment of Gevokizumab light chain to a sequence from OAS
DIQMTQSTSSLSASVGDRVTITCRASQDISNYLSWYQQKPGKAVKLLIYYTSKLHSGVPSRFSGSGSGTDYTLTISSLQQEDFATYFCLQGKMLPWTFGQGTKLEIK
|||||||.||||||||||||||||||||.|||||||||||||||.|||||..|.|.|||||||||||||||||.||||||||.||||||.|||.|..||||||||.|||
DIQMTQSPSSLSASVGDRVTITCRASQGISNYLSWYQQKPGKAPKLLIYDASTLQSGVPSRFSGSGSGTDFTLTISSLQPEDFATYYCLQYKSDPWTFGQGTKVEIK
^^^^^^ ^^^ ^^^^^^^^^

Best Alignment of Gevokizumab heavy chain CDRs to a sequence from OAS
QVQLQESGPGLVKPSQTLSLTCSFSGFSLSTSGMGVGWIRQPSGKGLEWLAHIWWDGDESYNPSLKSRLTISKDTSKNQVSLKITSVTAADTAVYFCARNRYDPPWFVDWGQGTLVTVSS
||.|.|||||..|||||||||||||||||||||||||||||||||||||||||||||.|..|||||||.||||||||.|||.||||||..||||.|.|||.|||.|||..||||||||||.
QVTLKESGPGILKPSQTLSLTCSFSGFSLSTSGMGVGWIRQPSGKGLEWLAHIWWDDDKYYNPSLKSQLTISKDTSRNQVFLKITSVDTADTATYYCARRRYDGPWFAYWGQGTLVTVSA
^^^^^^^^^^ ^^^^^^^ ^^^^^^^^^^^^

Best Alignment of Gevokizumab light CDRs chain to a sequence from OAS
DIQMTQSTSSLSASVGDRVTITCRASQDISNYLSWYQQKPGKAVKLLIYYTSKLHSGVPSRFSGSGSGTDYTLTISSLQQEDFATYFCLQGKMLPWTFGQGTKLEIK
|||||..||.|||..|||||||.|||||||||||.||||||...|||||||||..||||||.|||||||||||.||||.|.|||.|||||.||.|||||||.|||||||
DIQMTWPTSPLSAPLGDRVTISCRASQDISNYLNWYQQKPDGTVKLLIYYTSRIHSGVPSTFSGSGSGTDYSLTISNLEQEDIATYFCQQGNMLPWTFGGGTKLEIK
^^^^^^ ^^^ ^^^^^^^^^

Best Alignment of Gevokizumab CDR-H3 chain to a sequence from OAS
QVQLQESGPGLVKPSQTLSLTCSFSGFSLSTSGMGVGWIRQPSGKGLEWLAHIWWDGDESYNPSLKSRLTISKDTSKNQVSLKITSVTAADTAVYFCARNRYDPPWFVDWGQGTLVTVSS
.|||.||||.||||||||||||..|.|.........|||...|..||....|...|...||||||||..|..||||||..|...||||.|||.|.||||||..|||.|||||||||.
EVQLVESGPSLVKPSQTLSLTCSVTGDSI--TSGYWNWIRKFPGNKLEYMGYINYSGNTYYNPSLKSRISITRDTSKNQFYLQLNSVTADDTATYYCARNRYDEAWFVYWGQGTLVTVSP
^^^^^^^^^^ ^^^^^^^ ^^^^^^^^^^^^

Therapeutic : Girentuximab

Best Alignment of Girentuximab heavy chain to a sequence from OAS
DVKLVESGGGLVKLGGSLKLSCAASGFTFSNYYMSWVRQTPEKRLELVAAINSDGGITYYLDTVKGRFTISRDNAKNTLYLQMSSLKSEDTALFYCARHRSG---YFSMDYWGQGTSVTVSS
||||||||||||||||||||||||||||||.||||||||||||||||||||||.||.|||.||||||||||||||||||||||||||||||||.||||||.|...|..||||||||||||||
DVKLVESGGGLVKLGGSLKLSCAASGFTFSSYYMSWVRQTPEKRLELVAAINSNGGSTYYPDTVKGRFTISRDNAKNTLYLQMSSLKSEDTALYYCARHRDGNYDYYAMDYWGQGTSVTVSS
^^^^^^^^ ^^^^^^^^ ^^^^^^^^^^^^^^^

Best Alignment of Girentuximab light chain to a sequence from OAS
DIVMTQSQRFMSTTVGDRVSITCKASQNVVSAVAWYQQKPGQSPKLLIYSASNRYTGVPDRFTGSGSGTDFTLTISNMQSEDLADFFCQQYSNYPWTFGGGTKLEIK
||||||||.||||||||||||||||||||..||||||||||||||||||||||||||||||||||||||||||||||||||||||.||||||.||||||||||||||
DIVMTQSQKFMSTTVGDRVSITCKASQNVGTAVAWYQQKPGQSPKLLIYSASNRYTGVPDRFTGSGSGTDFTLTISNMQSEDLADYFCQQYSSYPWTFGGGTKLEIK
^^^^^^ ^^^ ^^^^^^^^^

Best Alignment of Girentuximab heavy chain CDRs to a sequence from OAS
DVKLVESGGGLVKLGGSLKLSCAASGFTFSNYYMSWVRQTPEKRLELVAAINSDGGITYYLDTVKGRFTISRDNAKNTLYLQMSSLKSEDTALFYCARHRSGYFSMDYWGQGTSVTVSS
.|.||||.||||..|.|.||||.|||||||.|||.||||.|||.||.||.||.||..|||||..|.|||.||||||||.||||||.|.|||..||||.|.||||||||||||||||||
EVHLVESEGGLVQPGSSMKLSCTASGFTFSDYYMAWVRQVPEKGLEWVANINYDGSSTYYLDSLKSRFIISRDNAKNILYLQMSSLQSDDTATYYCARERGGYFSMDYWGQGTSVTVSS
^^^^^^^^ ^^^^^^^^ ^^^^^^^^^^^^

Best Alignment of Girentuximab light CDRs chain to a sequence from OAS
DIVMTQSQRFMSTTVGDRVSITCKASQNVVSAVAWYQQKPGQSPKLLIYSASNRYTGVPDRFTGSGSGTDFTLTISNMQSEDLADFFCQQYSNYPWTFGGGTKLEIK
||||||||.||||.|||||..||||||||.|.||||.|||||||..||.|||.||.|||||||||||||||||.|.||||||..|||||||||||||||||||||
DIVMTQSQKFMSTSVGDRVTVTCKASQNVGSNVAWYEQKPGQSPTALIHSASYRYSGVPDRFTGSGSGTDFTLTINNVQSEDLAEYFCQQYSNYPWTFGGGTKLEIK
^^^^^^ ^^^ ^^^^^^^^^

Best Alignment of Girentuximab CDR-H3 chain to a sequence from OAS
DVKLVESGGGLVKLGGSLKLSCAASGFTFSNYYMSWVRQTPEKRLELVAAINSDGGITYYLDTVKGRFTISRDNAKNTLYLQMSSLKSEDTALFYCARHRSGYFSMDYWGQGTSVTVSS
.||||||||.|||.||||||||||||||||.|.||||||||.||||.||.|.|.|..|||.|.|||||||||||||||||||||||||||||||..||||||.||||||||||||||||||
EVKLVESGGDLVKPGGSLKLSCAASGFTFSSYGMSWVRQTPDKRLEWVATISSGGSYTYYPDSVKGRFTISRDNAKNTLYLQMSSLKSEDTAMYYCARHRDGYFSMDYWGQGTSVTVSS
^^^^^^^^ ^^^^^^^^ ^^^^^^^^^^^^

Therapeutic : Glembatumumab

Best Alignment of Glembatumumab heavy chain to a sequence from OAS
QVQLQESGPGLVKPSQTLSLTCTVSGGSISSFNYYWSWIRHHPGKGLEWIGYIYYSGSTYSNPSLKSRVTISVDTSKNQFSLTLSSVTAADTAVYYCARGYNW------NYFDYWGQGTLVTVSS
|||||||||||||||||||||||||||||||..|||||||.|||||||||||||||||||.|||||||||||||||||||||.||||||||||||||||||........||||||||||||||||
QVQLQESGPGLVKPSQTLSLTCTVSGGSISSGGYYWSWIRQHPGKGLEWIGYIYYSGSTYYNPSLKSRVTISVDTSKNQFSLKLSSVTAADTAVYYCARGYLSYYTRIRNYFDYWGQGTLVTVSS
^^^^^^^^^^ ^^^^^^^ ^^^^^^^^^^^^^^^^

Best Alignment of Glembatumumab light chain to a sequence from OAS

```
EIVMTQSPATLSVSPGERATLSCRASQSVDNNLVWYQQKPGQAPRLLIYGASTRATGIPARFSGSGSGTEFTLTISSLQSEDFAVYYCQQYNNWPPWTFGQGTKVEIK
|||||||||||||||||||||||||||||.||||||||||||||||||||||||||||||||||||||||||||||||||||||||||||||||||||||||||||||
EIVMTQSPATLSVSPGERATLSCRASQSVSNNLVWYQQKPGQAPRLLIYGASTRATGIPARFSGSGSGTEFTLTISSLQSEDFAVYYCQQYNNWPPWTFGQGTKVEIK
                          ^^^^^^                 ^^^                                    ^^^^^^^^^^
```

Best Alignment of Glembatumumab heavy chain CDRs to a sequence from OAS

```
QVQLQESGPGLVKPSQTLSLTCTVSGGSISSFNYYWSWIRHHPGKGLEWIGYIYYSGSTYSNPSLKSRVTISVDTSKNQFSLTLSSVTAADTAVYYCARGYNWNYFDYWGQGTLVTVSS
..............|.|||||||||||||||..|||.|||..|||||||||||||||||..|||||||||||||||||||||.||||||||||||||||||||||||||||||||||||
--------------SETLSLTCTVSGGSISSSSYYWGWIRQPPGKGLEWIGYIYYSGSTNYNPSLKSRVTISVDTSKNQFSLKLSSVTAADTAVYYCARGYNWNYFDYWGQGTLVTVSS
                         ^^^^^^^^^^                ^^^^^^^                                       ^^^^^^^^^^^
```

Best Alignment of Glembatumumab light CDRs chain to a sequence from OAS

```
EIVMTQSPATLSVSPGERATLSCRASQSVDNNLVWYQQKPGQAPRLLIYGASTRATGIPARFSGSGSGTEFTLTISSLQSEDFAVYYCQQYNNWPPWTFGQGTKVEIK
||||||||||||||||||||||||||||||||..|||||||||||||||||||||||||||||||||||||||||||.||||||||||||||||||||||||||||||
EIVMTQSPATLSVSPGERATLSCRASQSVDNNFAWYQQKPGQAPRLLIYGASTRATGIPARFSGSGSGTEFTLTISSRQSEDFAVYYCQQYNNWPPWTFGQGTKVEIK
                          ^^^^^^                 ^^^                                    ^^^^^^^^^^
```

Best Alignment of Glembatumumab CDR-H3 chain to a sequence from OAS

```
QVQLQESGPGLVKPSQTLSLTCTVSGGSISSFNYYWSWIRHHPGKGLEWIGYIYYS-GSTYSNPSLKSRVTISVDTSKNQFSLTLSSVTAADTAVYYCARGYNWNYFDYWGQGTLVTVSS
.....................|..||.......|...|.|..||..|||.|.|....|.|........||||..|||.......|.|....|||||||||||||||||||.|||||||||
----------------SVKVSCKASGYTF--TTYAIHWVRQAPGQRLEWMGWINAGNGNTKYAQKFQGRVTITRDTSASTAYMELRSLRSEDTAVYYCARGYNWNYFDYCGQGTLVTVSS
                         ^^^^^^^^^^                ^^^^^^^^                                       ^^^^^^^^^^^
```

Therapeutic : Golimumab

Best Alignment of Golimumab heavy chain to a sequence from OAS

```
QVQLVESGGGVVQPGRSLRLSCAASGFIFSSYAMHWVRQAPGNGLEWVAFMSYDGSNKKYADSVKGRFTISRDNSKNTLYLQMNSLRAEDTAVYYCARDRGIAAGGNYYYYGMDVWGQGTTVTVSS
|||||||||||||||||||||||||||.||||||||||||||.||||||..|||||||.|||||||||||||||||||||||||||||||||||||||||||||.|..||||||||||||||||||
QVQLVESGGGVVQPGRSLRLSCAASGFTFSSYAMHWVRQAPGKGLEWVAVISYDGSNKYYADSVKGRFTISRDNSKNTLYLQMNSLRAEDTAVYYCARDRGIAAAGLRYYYGMDVWGQGTTVTVSS
                         ^^^^^^^^                 ^^^^^^^^                                      ^^^^^^^^^^^^^^^^^^^
```

Best Alignment of Golimumab light chain to a sequence from OAS

```
EIVLTQSPATLSLSPGERATLSCRASQSVYSYLAWYQQKPGQAPRLLIYDASNRATGIPARFSGSGSGTDFTLTISSLEPEDFAVYYCQQRSNWPPFTFGPGTKVDIK
|||||||||||||||||||||||||||||.||||||||||||||||||||||||||||||||||||||||||||||||||||||||||||||||||||||||||||||
EIVLTQSPATLSLSPGERATLSCRASQSVSSYLAWYQQKPGQAPRLLIYDASNRATGIPARFSGSGSGTDFTLTISSLEPEDFAVYYCQQRSNWPPFTFGPGTKVDIK
                          ^^^^^^                 ^^^                                    ^^^^^^^^^^
```

Best Alignment of Golimumab heavy chain CDRs to a sequence from OAS

```
QVQLVESGGGVVQPGRSLRLSCAASGFIFSSYAMHWVRQAPGNGLEWVAFMSYDGSNKKYADSVKGRFTISRDNSKNTLYLQMNSLRAEDTAVYYCARDRGIAAGGNYYYYGMDVWGQGTTVTVSS
................|||||||||||.||||||||||||||.||||||..|||||||.|||||||||||||||||||||||||||||.|||||||||||||||.|.|||||||||||||||||||
----------------SLRLSCAASGFTFSSYAMHWVRQAPGKGLEWVAVISYDGSNKYYADSVKGRFTISRDNSKNTLYLQMNSLRAADTAVYYCARDRGIAAAGYYYYYGMDVWGQGTTVTVSS
                         ^^^^^^^^                 ^^^^^^^^                                      ^^^^^^^^^^^^^^^^^^^
```

Best Alignment of Golimumab light CDRs chain to a sequence from OAS

```
EIVLTQSPATLSLSPGERATLSCRASQSVYSYLAWYQQKPGQAPRLLIYDASNRATGIPARFSGSGSGTDFTLTISSLEPEDFAVYYCQQRSNWPPFTFGPGTKVDIK
|||||||||||||||||||||||||||||.||||||||||||||||||||||||||||||||||||||||||||||||||||||||||||||||||||||.||||.||
EIVLTQSPATLSLSPGERATLSCRASQSVSSYLAWYQQKPGQAPRLLIYDASNRATGIPARFSGSGSGTDFTLTISSLEPEDFAVYYCQQRSNWPPFTFGGGTKVEIK
                          ^^^^^^                 ^^^                                    ^^^^^^^^^^
```

Best Alignment of Golimumab CDR-H3 chain to a sequence from OAS

```
QVQLVESGGGVVQPGRSLRLSCAASGFIFSSYAMHWVRQAPGNGLEWVAFMSYDGSNKKYADSVKGRFTISRDNSKNTLYLQMNSLRAEDTAVYYCARDRGIAAGGNYYYYGMDVWGQGTTVTVSS
.............|...|.|.|.||...|||...|.||.||.||||.....|......|..|.|.|.|||.|.|||...|...|..|.|||||||||||||||.||||||||||||||||||||.
-------------PSETLSLTCTVSGGSISSYYWSWIRQPPGKGLEWIGYIYYS-GSTNYNPSLKSRVTISVDTSKNQFSLKLSSVTAADTAVYYCARDRGIAAAGNYYYYGMDVWGQGTTVTVS-
                         ^^^^^^^^                 ^^^^^^^^                                      ^^^^^^^^^^^^^^^^^^^
```

Therapeutic : Guselkumab

Best Alignment of Guselkumab heavy chain to a sequence from OAS

```
EVQLVQSGAEVKKPGESLKISCKGSGYSFSNYWIGWVRQMPGKGLEWMGIIDPSNSYTRYSPSFQGQVTISADKSISTAYLQWSSLKASDTAMYYCARWYY----KPFDVWGQGTLVTVSS
|||||||||||||||||||||||||||||.|||||||||||||||||||||||..|.|||||||||||||||||||||||||||||||||||||||||||||..|.....|||.|||||||||||
EVQLVQSGAEVKKPGESLKISCKGSGYSFTNYWIGWVRQMPGKGLEWMGIIDPGDSDTRYSPSFQGQVTISADKSISTAYLQWSSLKASDTAMYYCARHSYSNYAPPFDYWGQGTLVTVSS
                         ^^^^^^^^                 ^^^^^^^^                                      ^^^^^^^^^^^^^^
```

Best Alignment of Guselkumab light chain to a sequence from OAS

```
QSVLTQPPSVSGAPGQRVTISCTGSSSNIGSGYDVHWYQQLPGTAPKLLIYGNSKRPSGVPDRFSGSKSGTSASLAITGLQSEDEADYYCASWTDGLSLVVFGGGTKLTVL
||||||||||||||||||||||||||||||.|||||||||||||||||||||||.||||||||||||||||||||||||||.|||||||||||.|.||.||||||||||||
QSVLTQPPSVSGAPGQRVTISCTGSSSNIGAGYDVHWYQQLPGTAPKLLIYGNSNRPSGVPDRFSGSKSGTSASLAITGLQAEDEADYYCASWDDSLSGVVFGGGTKLTVL
                        ^^^^^^^^^                  ^^^                                    ^^^^^^^^^^^
```

Best Alignment of Guselkumab heavy chain CDRs to a sequence from OAS

```
EVQLVQSGAEVKKPGESLKISCKGSGYSFSNYWIGWVRQMPGKGLEWMGIIDPSNSYTRYSPSFQGQVTISADKSISTAYLQWSSLKASDTAMYYCARWYYKPFDVWGQGTLVTVSS
..............||||||||||||||||||||.||||||||||||||.||||.|||.|||||||.|||||||||||||||||||||||||||||||.||...|||||||.|||||
--------------GESLKISCKGSGYSFSNYWISWVRQMPGKGLEWMGRIDPSDSYTNYSPSFQGHVTISADKSISTAYLQWSSLKASDTAMYYCARLYYYAMDVWGQGTTVTVSS
                         ^^^^^^^^                 ^^^^^^^^                                      ^^^^^^^^^^
```

Best Alignment of Guselkumab light CDRs chain to a sequence from OAS

```
QSVLTQPPSVSGAPGQRVTISCTGSSSNIGSGYDVHWYQQLPGTAPKLLIYGNSKRPSGVPDRFSGSKSGTSASLAITGLQSEDEADYYCASWTDGLSLVVFGGGTKLTVL
||||||||||||||||||||||||||||||.||||||||||||.||||||||||.||||||||||||||||||||||||||.|||||||||||.|.||.||||||||||||
QSVLTQPPSVSGAPGQRVTISCTGSSSNIGAGYDVHWYQQLPGAAPKLLIYGNSNRPSGVPDRFSGSKSGTSASLAITGLQAEDEADYYCASWDDSLSGVVFGGGTKLTVL
                        ^^^^^^^^^                  ^^^                                    ^^^^^^^^^^^
```

Best Alignment of Guselkumab CDR-H3 chain to a sequence from OAS

```
EVQLVQSGAEVKKPGESLKISCKGSGYSFSNYWIGWVRQMPGKGLEWMGIIDPSNSYTRYSPSFQGQVTISADKSISTAYLQWSSLKASDTAMYYCARWYYKPFDVWGQGTLVTVSS
.|||.|.|||...||.|.|.|||.|||.|..||..||.|.||.||||.|.|.||.|.|.|...|....|...|||.||||.|.|||...|.|.||||||||.||||||.||.|||||
QVQLQQPGAELVRPGSSVKLSCKASGYTFTSYWMDWVKQRPGQGLEWIGNIYPSDSETHYNQKFKDKATLTVDKSSSTAYMQLSSLTSEDSAVYYCARWYYGPFDVWGTGTTVTVSS
                         ^^^^^^^^                 ^^^^^^^^                                      ^^^^^^^^^^
```

Therapeutic : Ianalumab

Best Alignment of Ianalumab heavy chain to a sequence from OAS

```
QVQLQQSGPGLVKPSQTLSLTCAISGDSVSSNSAAWGWIRQSPGRGLEWLGRIYYRSKWYNSYAVSVKSRITINPDTSKNQFSLQLNSVTPEDTAVYYCARYQWVP--KIGVFDSWGQGTLVTVSS
||||||||||||||||||||||||||||||||||||.||||||.||||||||.||||||||.|||||||||||||||||||||||||||||||||||||||..|......||||.|||||||||||
QVQLQQSGPGLVKPSQTLSLTCAISGDSVSSNSAAWNWIRQSPSRGLEWLGRTYYRSKWYNDYAVSVKSRITINPDTSKNQFSLQLNSVTPEDTAVYYCARESWGGGWYKGVFDYWGQGTLVTVSS
                         ^^^^^^^^^^                 ^^^^^^^^^                                      ^^^^^^^^^^^^^^^^
```

Best Alignment of Ianalumab light chain to a sequence from OAS

```
DIVLTQSPATLSLSPGERATLSCRASQFILPEYLSWYQQKPGQAPRLLIYGSSSRATGVPARFSGSGSGTDFTLTISSLEPEDFAVYYCQQFYSSPLTFGQGTKVEIK
|||||||||||||||||||||||||||.....||.||||||||||||||||.||||||||.|||||||||||||||||||||||||||||..|||.||||||.||||
DIVLTQSPATLSLSPGERATLSCRASQSV-SSYLAWYQQKPGQAPRLLIYGASSRATGVPDRFSGSGSGTDFTLTISSLEPEDFAVYYCQQYGSSPYTFGQGTRVEIK
                          ^^^^^^^                 ^^^                                   ^^^^^^^^^
```

Best Alignment of Ianalumab heavy chain CDRs to a sequence from OAS

```
QVQLQQSGPGLVKPSQTLSLTCAISGDSVSSNSAAWGWIRQSPGRGLEWLGRIYYRSKWYNSYAVSVKSRITINPDTSKNQFSLQLNSVTPEDTAVYYCARYQWVPKIGVFDSWGQGTLVTVSS
....................|||||||||||||||.||||||.||||||||.||||||||.|||||.|||||||||||.||||||||||||||||||||.|..||.|.||.|||||||||||
```

```
---------------------CAISGDSVSSNSAAWNWIRQSPSRGLEWLGRTYYRSKWYNDYAVSVESRITINPDTSKIQFSLQLNSVTPEDTAVYYCARSQITPKLGWFDPWGQGTLVTVSS
                         ^^^^^^^^^^          ^^^^^^^^^                            ^^^^^^^^^^^^^^
```

Best Alignment of Ianalumab light CDRs chain to a sequence from OAS

```
DIVLTQSPATLSLSPGERATLSCRASQFILPEYLSWYQQKPGQAPRLLIYGSSSRATGVPARFSGSGSGTDFTLTISSLEPEDFAVYYCQQFYSSPLTFGQGTKVEIK
.||||||.||||||||||||||||||.||..||.|||||||||||||||||.||||||.|||||||||.||||||.|||||||||||||||..||||||.|||.|||
EIVLTQSPGTLSLSPGERATLSCRASQTILRSYLAWYQQKPGQAPRLLIYGSSNRATGVPDRFSGSGSGTEFTLTISRLEPEDFAVYYCQQFETSPLTFGGGTKLEIK
                          ^^^^^^^          ^^^                          ^^^^^^^^^
```

Best Alignment of Ianalumab CDR-H3 chain to a sequence from OAS

```
QVQLQQSGPGLVKPSQTLSLTCAISGDSVSSNSAAWGWIRQSPGRGLEWLGRIYYRSKWYNSYAVSVKSRITINPDTSKNQFSLQLNSVTPEDTAVYYCARYQWVPKIGVFDSWGQGTLVTVSS
.............|.||||||..||.|.||.|..||||||.||.||||.|.|||.......|..|.|||.||..|||||||||.|.|||..|||||||||.|||..|||||.|||||||||||
--------------SETLSLTCTVSGGSISSSSYYWGWIRQPPGKGLEWIGSIYYS--GSTYYNPSLKSRVTISVDTSKNQFSLKLSSVTAADTAVYYCARDQWVATIGVFDYWGQGTLVTVSS
                         ^^^^^^^^^^          ^^^^^^^^^                            ^^^^^^^^^^^^^^
```

Therapeutic : Ibalizumab

Best Alignment of Ibalizumab heavy chain to a sequence from OAS

```
QVQLQQSGPEVVKPGASVKMSCKASGYTFTSYVIHWVRQKPGQGLDWIGYINPYNDGTDYDEKFKGKATLTSDTSTSTAYMELSSLRSEDTAVYYCAREKDNYATGAWFAYWGQGTLVTVSS
.||||||||.|||||||||||||||||||||||.|||.|||||||.||||||||||||.|.|||||||||||||.|.|||||||||||.|||.||||||||.|....||||||||||||||||.
EVQLQQSGPELVKPGASVKMSCKASGYTFTSYVMHWVKQKPGQGLEWIGYINPYNDGTKYNEKFKGKATLTSDKSSSTAYMELSSLTSEDSAVYYCAREWDYD--GAWFAYWGQGTLVTVSA
                         ^^^^^^^^        ^^^^^^^^                         ^^^^^^^^^^^^^^^
```

Best Alignment of Ibalizumab light chain to a sequence from OAS

```
DIVMTQSPDSLAVSLGERVTMNCKSSQSLLYSTNQKNYLAWYQQKPGQSPKLLIYWASTRESGVPDRFSGSGSGTDFTLTISSVQAEDVAVYYCQQYYSYRTFGGGTKLEIK
||||.|||.|||||.||.|||.|||||||||||.|||||||||||||||||||||||||||||||||||||||||.||||||||||||||||||||.||||||||||||||||||||||
DIVMSQSPSSLAVSVGEKVTMSCKSSQSLLYSSNQKNYLAWYQQKPGQSPKLLIYWASTRESGVPDRFTGSGSGTDFTLTISSVQAEDLAVYYCQQYYSYRTFGGGTKLEIK
                          ^^^^^^^^^^^^          ^^^                  ^^^^^^^^
```

Best Alignment of Ibalizumab heavy chain CDRs to a sequence from OAS

```
QVQLQQSGPEVVKPGASVKMSCKASGYTFTSYVIHWVRQKPGQGLDWIGYINPYNDGTDYDEKFKGKATLTSDTSTSTAYMELSSLRSEDTAVYYCAREKDNYATGAWFAYWGQGTLVTVSS
.||||||||.|||||||||||||||||||||||.|||.|||||||.||||||||||||.|.|||||||||||||.|.|||||||||||.|||.|||||||..|.|.|||||||||||||||||.
EVQLQQSGPELVKPGASVKMSCKASGYTFTSYVMHWVKQKPGQGLEWIGYINPYNDGTKYNEKFKGKATLTSDKSSSTAYMELSSLTSEDSAVYYCARGGDYYGTGAWFAYWGQGTLVTVSA
                         ^^^^^^^^        ^^^^^^^^                         ^^^^^^^^^^^^^^^
```

Best Alignment of Ibalizumab light CDRs chain to a sequence from OAS

```
DIVMTQSPDSLAVSLGERVTMNCKSSQSLLYSTNQKNYLAWYQQKPGQSPKLLIYWASTRESGVPDRFSGSGSGTDFTLTISSVQAEDVAVYYCQQYYSYRTFGGGTKLEIK
||.|.|||.|||||.||..||.|.|||||||||.||||||||||||||||||||||||||||||.||||.||||||||||||||||.|||.|||||||||||||||||.|||||.|
DIAMLQSPSSLAVSVGEKATMSCNSSQSLLYSSNQKNYLAWYQQKPGQSPKLLIYWASTRESGIPDRFTGSGSGTDFTLTISSVKAEDLAVYYCQQYYSYRTFGAGTKLELK
                          ^^^^^^^^^^^^          ^^^                  ^^^^^^^^
```

Best Alignment of Ibalizumab CDR-H3 chain to a sequence from OAS

```
QVQLQQSGPEVVKPGASVKMSCKASGYTFTSYVIHWVRQKPGQGLDWIGYINPYNDGTDYDEKFKGKATLTSDTSTSTAYMELSSLRSEDTAVYYCAREKDNYATGAWFAYWGQGTLVTVSS
.|||||.|||.||||||||.|||||||.||.|..|||.|.....|.|||.|||||..|.|...||.||.||.|.|.||||||||.||.|||.||||||||.|.||.|||||||||||||||||.
-VQLQQFGPELVKPGASVKISCKASGYSFTGYYMHWVKQSHVKSLEWIGRINPYNGATSYNQNFKDKASLTVDKSSSTAYMELHSLTSEDSAVYYCAREEDYYASGAWFAYWGQGTLVTVSA
                         ^^^^^^^^        ^^^^^^^^                         ^^^^^^^^^^^^^^^
```

Therapeutic : Icrucumab

Best Alignment of Icrucumab heavy chain to a sequence from OAS

```
QAQVVESGGGVVQSGRSLRLSCAASGFAFSSYGMHWVRQAPGKGLEWVAVIWYDGSNKYYADSVRGRFTISRDNSENTLYLQMNSLRAEDTAVYYCARDHYGSGVH---HYFYYGLDVWGQGTTVTVSS
|.|.|||||||||.|||||||||||||.|||||||||||||||||||||||||||||||||||||||.||||||||||.||||||||||||||||||||||||||.||||.....||.|||.|||||||||||||
QVQLVESGGGVVQPGRSLRLSCAASGFTFSSYGMHWVRQAPGKGLEWVAVIWYDGSNKYYADSVKGRFTISRDNSKNTLYLQMNSLRAEDTAVYYCARDYYGSGSYYSFHYYYYGVDVWGQGTTVTVSS
                         ^^^^^^^^        ^^^^^^^^                         ^^^^^^^^^^^^^^^^^^^^^^
```

Best Alignment of Icrucumab light chain to a sequence from OAS

```
EIVLTQSPGTLSLSPGERATLSCRASQSVSSSYLAWYQQKPGQAPRLLIYGASSRATGIPDRFSGSGSGTDFTLTISRLEPEDFAVYYCQQYGSSPLTFGGGTKVEIK
||||||||||||||||||||||||||||||||||||||||||||||||||||||||||||||||||||||||||||||||||||||||||||||||||||||||||||
EIVLTQSPGTLSLSPGERATLSCRASQSVSSSYLAWYQQKPGQAPRLLIYGASSRATGIPDRFSGSGSGTDFTLTISRLEPEDFAVYYCQQYGSSPLTFGGGTKVEIK
                          ^^^^^^^          ^^^                  ^^^^^^^^^
```

Best Alignment of Icrucumab heavy chain CDRs to a sequence from OAS

```
QAQVVESGGGVVQSGRSLRLSCAASGFAFSSYGMHWVRQAPGKGLEWVAVIWYDGSNKYYADSVRGRFTISRDNSENTLYLQMNSLRAEDTAVYYCARDHYGSGVHHYFYYGLDVWGQGTTVTVSS
|.|.|||||||||.|||||||||||||.|||||||||||||||||||||||||||||||||||||||.||||||||||.||||||||||||||||||||||||||||||||...|.|||.|||||||||||||
QVQLVESGGGVVQPGRSLRLSCAASGFTFSSYGMHWVRQAPGKGLEWVAVIWYDGSNKYYADSVKGRFTISRDNSKNTLYLQMNSLRAEDTAVYYCARDHYGSGHALYYYYGMDVWGQGTTVTVSS
                         ^^^^^^^^        ^^^^^^^^                         ^^^^^^^^^^^^^^^^^^
```

Best Alignment of Icrucumab light CDRs chain to a sequence from OAS

```
EIVLTQSPGTLSLSPGERATLSCRASQSVSSSYLAWYQQKPGQAPRLLIYGASSRATGIPDRFSGSGSGTDFTLTISRLEPEDFAVYYCQQYGSSPLTFGGGTKVEIK
|||||||||||||||||||||||||||||||||||||||||||||||||||||||||||||||||||||.|||||||||||||||||||||||||||||||.||..|||
EIVLTQSPGTLSLSPGERATLSCRASQSVSSSYLAWYQQKPGQAPRLLIYGASSRATGIPDRFSGSGSGADFTLTISRLEPEDFAVYYCQQYGSSPLTFGQGTRLEIK
                          ^^^^^^^          ^^^                  ^^^^^^^^^
```

Best Alignment of Icrucumab CDR-H3 chain to a sequence from OAS

```
QAQVVESGGGVVQSGRSLRLSCAASGFAFSSYGMHWVRQAPGKGLEWVAVIWYDGSNKYYADSVRGRFTISRDNSENTLYLQMNSLRAEDTAVYYCARDHYGSGVHHYFYYGLDVWGQGTTVTVSS
.............|.||..|||||||.||||.|.|||||||||||||..|...||..||||||.|||||||||..|.||||||||||||||||||||||||||||||..||.|||.|||||||||||||
--------------GESLKISCAASGFTFSSYSMNWVRQAPGKGLEWVSYISSSGSTIYYADSVKGRFTISRDNAKNSLYLQMNSLRAEDTAVYYCARDHYGSGSYHYYYYGMDVWGQGTTVTVSS
                         ^^^^^^^^        ^^^^^^^^                         ^^^^^^^^^^^^^^^^^^
```

Therapeutic : Idarucizumab

Best Alignment of Idarucizumab heavy chain to a sequence from OAS

```
QVQLQESGPGLVKPSETLSLTCTVSGFSLTSYIVDWIRQPPGKGLEWIGVIWAGGSTGYNSALRSRVSITKDTSKNQFSLKLSSVTAADTAVYYCASAAYYSY-------YNYDGFAYWGQGTLVTVSS
||||||||||||||||||||||||||.|..||...||||||||||||||.|...|||.||..|.|||.|..|||||||||||||||||||||||||||||||..|.|........|.||.|.||||||||||||
QVQLQESGPGLVKPSETLSLTCTVSGGSISSYYWSWIRQPPGKGLEWIGYIYYSGSTNYNPSLKSRVTISVDTSKNQFSLKLSSVTAADTAVYYCASSGYCSGGSLCGYSYGYDYFDYWGQGTLVTVSS
                         ^^^^^^^^        ^^^^^^^                         ^^^^^^^^^^^^^^^^^^^^^^^
```

Best Alignment of Idarucizumab light chain to a sequence from OAS

```
DVVMTQSPLSLPVTLGQPASISCKSSQSLLYTDGKTYLYWFLQRPGQSPRRLIYLVSKLDSGVPDRFSGSGSGTDFTLKISRVEAEDVGVYYCLQSTHFPHTFGGGTKVEIK
||||||.||.|.||.||||||||||||||||||.|||||.|.||||||||.|||||||||||||||||||||||||||||||||||||.||||||||||||||||||||.|||
DVVMTQTPLTLSVTIGQPASISCKSSQSLLYTNGKTYLNWLLQRPGQSPKRLIYLVSKLDSGVPDRFSGSGSGTDFTLKISRVEAEDLGVYYCLQSTHFPHTFGGGTKLEIK
                          ^^^^^^^^^^^          ^^^                  ^^^^^^^^^
```

Best Alignment of Idarucizumab heavy chain CDRs to a sequence from OAS

```
QVQLQESGPGLVKPSETLSLTCTVSGFSLTSYIVDWIRQPPGKGLEWIGVIWAGGSTGYNSALRSRVSITKDTSKNQFSLKLSSVTAADTAVYYCASAAYYSYYNYDGFAYWGQGTLVTVSS
|||||.||||||.||..||.|||||||||||||.|||.||.|||||||.||||.||||.|||||.||.||.||.||.|..||..|....|||.|||||.||||||.||.|||||||||||||.
QVQLQQSGPGLVAPSQSLSITCTVSGFSLTSYGVDWVRQSPGKGLEWLGVIWGGGSTNYNSALKSRLSISKDNSKSQVFLKMNSLQTDDTAMYYCASSAYYSYYSYDPFAYWGQGTLVTVSA
                         ^^^^^^^^        ^^^^^^^                         ^^^^^^^^^^^^^^^
```

Best Alignment of Idarucizumab light CDRs chain to a sequence from OAS

```
DVVMTQSPLSLPVTLGQPASISCKSSQSLLYTDGKTYLYWFLQRPGQSPRRLIYLVSKLDSGVPDRFSGSGSGTDFTLKISRVEAEDVGVYYCLQSTHFPHTFGGGTKVEIK
||||||.||.|.||.||||||||||||||||||.|||||.|.||||||||.|||||||||||||||||||||||||||||||||||||.|||||||||||||||||.|||.|||
DVVMTQTPLTLSVTIGQPASISCKSSQSLLYTNGKTYLNWLLQRPGQSPKRLIYLVSKLDSGVPDRFSGSGSGTDFTLKISRVEAEDLGVYYCLQSTHFPHTFGSGTKLEIK
                          ^^^^^^^^^^^          ^^^                  ^^^^^^^^^
```

Best Alignment of Idarucizumab CDR-H3 chain to a sequence from OAS

```
QVQLQESGPGLVKPSETLSLTCTVSGFSLTSYIVDWIRQPPGKGLEWIGVIWAG-GSTGYNSALRSRVSITKDTSKNQFSLKLSSVTAADTAVYYCASAAYYSYYNYDGFAYWGQGTLVTVSS
|||||..|..||||.....|.|..||...|||...|..|.||.||||||.|....|||.||....|....|.|.|.......|||.|..|.|||||||.||||||.||||||||||||||||.
```

QVQLQQPGAELVKPGASVKLSCKASGYTFTSYWMHWVKQRPGQGLEWIGMIHPNSGSTNYNEKFKSKATLTVDKSSSTAYMQLSSLTSEDSAVYYCASEAYYSYSYSYDGFAYWGQGTLVTVSA
                    ^^^^^^^^               ^^^^^^^^                                    ^^^^^^^^^^^^^^^^

Therapeutic : Imgatuzumab

Best Alignment of Imgatuzumab heavy chain to a sequence from OAS
QVQLVQSGAEVKKPGSSVKVSCKASGFTFTDYKIHWVRQAPGQGLEWMGYFNPNSGYSTYAQKFQGRVTITADKSTSTAYMELSSLRSEDTAVYYCARLSPGG----YYVMDAWGQGTTVTVSS
|||||||||||||.||||||||||.|||.|..|||||||||||||||..|||||...|||||||||||||||||||||||||||||||||||||||.|..|....||.|.|||||||||||
QVQLVQSGAEVKKPGASVKVSCKASGYTFTGYYMHWVRQAPGQGLEWMGWINPNSGGTNYAQKFQGRVTITADKSTSTAYMELSSLRSEDTAVYYCARGSVRGVIDYYYGMDVWGQGTTVTVSS
                    ^^^^^^^^               ^^^^^^^^                                    ^^^^^^^^^^^^^^^^

Best Alignment of Imgatuzumab light chain to a sequence from OAS
DIQMTQSPSSLSASVGDRVTITCRASQGINNYLNWYQQKPGKAPKRLIYNTNNLQTGVPSRFSGSGSGTEFTLTISSLQPEDFATYYCLQHN-SFPTFGQGTKLEIK
|||||||||||||||||||||||||||||.|.|.||||||||||||||.|.|||.||||||||||||||||||||||||||||||||||||.|..|||||||.|||
DIQMTQSPSSLSASVGDRVTITCRASQGIRNGLGWYQQKPGKAPKRLIYATSNLQSGVPSRFSGSGSGTEFTLTISSLQPEDFATYYCLQHNSSPRTFGQGTKVEIK
                     ^^^^^^                ^^^                                ^^^^^^^^^

Best Alignment of Imgatuzumab heavy chain CDRs to a sequence from OAS
QVQLVQSGAEVKKPGSSVKVSCKASGFTFTDYKIHWVRQAPGQGLEWMGYFNPNSGYSTYAQKFQGRVTITADKSTSTAYMELSSLRSEDTAVYYCARLSPGGYYVMDAWGQGTTVTVSS
|||||||||||.|||.|||||||||.|||||.|..||||||||||||||..|||||...|||||||||.|.|.|.||||||||.|||.|||||||||||..|||.||.|.|||||||||
QVQLVQSGAEVQKPGASVKVSCKASGYTFTDYYMHWVRQAPGQGLEWMGWINPNSGGTNYAQKFQGRVTMTRDTSISTAYMELSRLRSDDTAVYYCARLSGDGYYYMDVWGKGTTVTVSS
                    ^^^^^^^^               ^^^^^^^^                                    ^^^^^^^^^^^^^

Best Alignment of Imgatuzumab light CDRs chain to a sequence from OAS
DIQMTQSPSSLSASVGDRVTITCRASQGINNYLNWYQQKPGKAPKRLIYNTNNLQTGVPSRFSGSGSGTEFTLTISSLQPEDFATYYCLQHNSFPTFGQGTKLEIK
|||||||||||||||||||||||||||||||||||||||||.|||||.|.|.||.|..||||||||||||||||||||.||||||||||||.|.||.||||.|||.|||
DIQMTQSPSSLSASVGDRVTITCRASQGINNYLNWYQQKQGKAPKLLNYYTNRLERGVPSRFSGSGSGTEFTLTIGSLQPEDFATYYCQQYNSLPTFGGGTKVEIK
                     ^^^^^^                ^^^                                ^^^^^^^^

Best Alignment of Imgatuzumab CDR-H3 chain to a sequence from OAS
QVQLVQSGAEVKKPGSSVKVSCKASGFTFTDYKIHWVRQAPGQGLEWMGYFNPNSGYSTYAQKFQGRVTITADKSTSTAYMELSSLRSEDTAVYYCARLSPGGYYVMDAWGQGTTVTVSS
.||||.|......||.|.|.||.||||||..|...||||||..||||.....|..|...|.....||.||.|...||.|....||||||||.||.|||||.|||||||||||..|||||
EVQLVESDGGLVQPGRSLKLSCAASGFTFSNYDMAWVRQAPTKGLEWVASISPSGGSTYYRDSVKGRFTISRDNAKSTLYLQMDSLRSEDTATYYGARLSPTGYYVMDAWGQGASVTVSS
                    ^^^^^^^^               ^^^^^^^^                                    ^^^^^^^^^^^^^

Therapeutic : Inclacumab

Best Alignment of Inclacumab heavy chain to a sequence from OAS
EVQLVESGGGLVRPGGSLRLSCAASGFTFSNYDMHWVRQATGKGLEWVSAITAAGDIYYPGSVKGRFTISRENAKNSLYLQMNSLRAGDTAVYYCARGRYSGSGSYYNDWFDPWGQGTLVTVSS
||||||||||||.|||||||||||||||||.|||||||||||||||||||||..|||.||||||||||||||||||||||||||||||||||||||||||||.||||...||..|||||||||||||||
EVQLVESGGGLVQPGGSLRLSCAASGFTFSSYDMHWVRQATGKGLEWVSAIGTAGDTYYPGSVKGRFTISRENAKNSLYLQMNSLRAGDTAVYYCARGSYSGS---YRNWFDPWGQGTLVTVSS
                    ^^^^^^^^               ^^^^^^^                                   ^^^^^^^^^^^^^^^^^^

Best Alignment of Inclacumab light chain to a sequence from OAS
EIVLTQSPATLSLSPGERATLSCRASQSVSSYLAWYQQKPGQAPRLLIYDASNRATGIPARFSGSGSGTDFTLTISSLEPEDFAVYYCQQRSNWPLTFGGGTKVEIK
||||||||||||||||||||||||||||||||||||||||||||||||||||||||||||||||||||||||||||||||||||||||||||||||||||||||||||
EIVLTQSPATLSLSPGERATLSCRASQSVSSYLAWYQQKPGQAPRLLIYDASNRATGIPARFSGSGSGTDFTLTISSLEPEDFAVYYCQQRSNWPLTFGGGTKVEIK
                     ^^^^^^                ^^^                                ^^^^^^^^^

Best Alignment of Inclacumab heavy chain CDRs to a sequence from OAS
EVQLVESGGGLVRPGGSLRLSCAASGFTFSNYDMHWVRQATGKGLEWVSAITAAGDIYYPGSVKGRFTISRENAKNSLYLQMNSLRAGDTAVYYCARGRYSGSGSYYNDWFDPWGQGTLVTVSS
.............|||||||||||||||||.||||||||||||||||||||..|||.|||||||||||||||.||||||||||||||||.|||||||||.||.|||||||..|||||||||||||||
--------------GGSLRLSCAASGFTFSSYDMHWVRQATGKGLEWVSAIGTAGDTYYPGSVKGRFTISRDNAKNSLYLQMNSLRAEDTAVYYCARDRYYGSGSYYNVPFDPWGQGTLVTVSS
                    ^^^^^^^^               ^^^^^^^                                   ^^^^^^^^^^^^^^^^^^

Best Alignment of Inclacumab light CDRs chain to a sequence from OAS
EIVLTQSPATLSLSPGERATLSCRASQSVSSYLAWYQQKPGQAPRLLIYDASNRATGIPARFSGSGSGTDFTLTISSLEPEDFAVYYCQQRSNWPLTFGGGTKVEIK
|||||||||||||||||||||||||||||||||||||||||||||||||||||||||||||||||.||||||||||||||||||||||||||||||||||||||||||
EIVLTQSPATLSLSPGERATLSCRASQSVSSYLAWYQQKPGQAPRLLIYDASNRATGIPARFSGSGPGTDFTLTISSLEPEDFAVYYCQQRSNWPLTFGGGTKVEIK
                     ^^^^^^                ^^^                                ^^^^^^^^^

Best Alignment of Inclacumab CDR-H3 chain to a sequence from OAS
EVQLVESGGGLVRPGGSLRLSCAASGFTFSNYDMHWVRQATGKGLEWVSAITAA-GDIYYPGSVKGRFTISRENAKNSLYLQMNSLRAGDTAVYYCARGRYSGSGSYYNDWFDPWGQGTLVTVSS
............|||||||||||||||||.|.|||||||.|||||||..|.......||..|||||||||||.|.||.||||||||||.||||||||||||.|||||||.|||||||||||||||
-------------PGGSLRLSCAASGFTFSSYGMHWVRQAPGKGLEWVAFIRYDGSNKYYADSVKGRFTISRDNSKNTLYLQMNSLRAEDTAVYYCARGRYYGSGSYYNFWFDPWGQGTLVTVSS
                    ^^^^^^^^               ^^^^^^^^                                    ^^^^^^^^^^^^^^^^^^

Therapeutic : Indatuximab

Best Alignment of Indatuximab heavy chain to a sequence from OAS
QVQLQQSGSELMMPGASVKISCKATGYTFSNYWIEWVKQRPGHGLEWIGEILPGTGRTIYNEKFKGKATFTADISSNTVQMQLSSLTSEDSAVYYCARRDYYGN-FYYAMDYWGQGTSVTVSS
||||||||.|||.||||||||||||||||||.|||||||||||||||||||||||||.|.|.|||||||||||||||.||||..||||||||||||||||||||.|||...|||||||||||||||||
QVQLQQSGAELMKPGASVKISCKATGYTFSSYWIEWVKQRPGHGLEWIGEILPGSGSTNYNEKFKGKATFTADTSSNTAYMQLSSLTSEDSAVYYCARRGYYGSTHYYAMDYWGQGTSVTVSS
                    ^^^^^^^^               ^^^^^^^^                                    ^^^^^^^^^^^^^^^^

Best Alignment of Indatuximab light chain to a sequence from OAS
DIQMTQSTSSLSASLGDRVTISCSASQGINNYLNWYQQKPDGTVELLIYYTSTLQSGVPSRFSGSGSGTDYSLTISNLEPEDIGTYYCQQYSKLPRTFGGGTKLEIK
||||||.|||||||||||||||||||||||.||||||||||||||||.|||||||.||||||||||||||||||||||||||||||||||||.|||||||||||||||||||||||
DIQMTQTTSSLSASLGDRVTISCSASQGISNYLNWYQQKPDGTVKLLIYYTSSLQSGVPSRFSGSGSGTDYSLTISNLEPEDIATYYCQQYSKLPRTFGGGTKLEIK
                     ^^^^^^                ^^^                                ^^^^^^^^^

Best Alignment of Indatuximab heavy chain CDRs to a sequence from OAS
QVQLQQSGSELMMPGASVKISCKATGYTFSNYWIEWVKQRPGHGLEWIGEILPGTGRTIYNEKFKGKATFTADISSNTVQMQLSSLTSEDSAVYYCARRDYYGNFYYAMDYWGQGTSVTVSS
.||||||.|.|.||||||.|||||||||..|||||||||||||||||||||||||||.|.||.|||.|||.|.|.|.|..|||.|||||||||||||||.||||||||||||||||||||||
-VQLQQSGDEVMKPGASVKLSCKATGYTFTDYWIEWVKQRPGHGLEWIGEILPGTGSTTYNQKFKAKATLTVDKSSSTAYMQLKSLTSEDSAVYYCARYDYYGNFYYAMDYWGQGTSVTVSS
                    ^^^^^^^^               ^^^^^^^^                                    ^^^^^^^^^^^^^^^

Best Alignment of Indatuximab light CDRs chain to a sequence from OAS
DIQMTQSTSSLSASLGDRVTISCSASQGINNYLNWYQQKPDGTVELLIYYTSTLQSGVPSRFSGSGSGTDYSLTISNLEPEDIGTYYCQQYSKLPRTFGGGTKLEIK
||||||.|||||.||||||||||||||||||||||||||.|||||.|||||||.|.||||||||||||||||||||||||||||||.|||||||||||.|||.|||||||
DIQMTQTTSSLSVSLGDRVTISCSASQGINNYLNWYQQNPDGTVKLLIYYTSSLHSGVPSRFSGSGSGTDYSLTISNLEPEDIATYYCQQYSKLPFTFGSGTKLEIK
                     ^^^^^^                ^^^                                ^^^^^^^^^

Best Alignment of Indatuximab CDR-H3 chain to a sequence from OAS
QVQLQQSGSELMMPGASVKISCKATGYTFSNYWIEWVKQRPGHGLEWIGEILPGTGRTIYNEKFKGKATFTADISSNTVQMQLSSLTSEDSAVYYCARRDYYGNFYYAMDYWGQGTSVTVSS
||||..||.||..||||||.||||.||||..|.|.||.|||..||||||.|.|.|.|.||||||||||.|.|.||.|..||||.|||||||||.||||||||||||||||||||||||||||||
QVQLKESGAELVKPGASVKLSCKASGYTFTSYDINWVRQRPEQGLEWIGWIFPGDGSTKYNEKFKGKATLTTDKSSSTAYMQLSRLTSEDSAVYFCARRDYYGNFYYAMDYWGQGTSVTVSS
                    ^^^^^^^^               ^^^^^^^^                                    ^^^^^^^^^^^^^^^

Therapeutic : Indusatumab

Best Alignment of Indusatumab heavy chain to a sequence from OAS
QVQLQQWGAGLLKPSETLSLTCAVFGGSFSGYYWSWIRQPPGKGLEWIGEINHRGNTNDNPSLKSRVTISVDTSKNQFALKLSSVTAADTAVYYCARERGYT--YGNFDHWGQGTLVTVSS
||||||||||||||||||||||.||||||||||||||||||||||||||||||.|.||.|||||||||||||||||||.||||||||||||||||||||||||.|..|.|||.|||||||||||
QVQLQQWGAGLLKPSETLSLTCAVYGGSFSGYYWSWIRQPPGKGLEWIGEINHSGSTNYNPSLKSRVTISVDTSKNQFSLKLSSVTAADTAVYYCARERGITGYYTNFDYWGQGTLVTVSS
                    ^^^^^^^^               ^^^^^^^                                   ^^^^^^^^^^^^^^^

Best Alignment of Indusatumab light chain to a sequence from OAS
```
EIVMTQSPATLSVSPGERATLSCRASQSVSRNLAWYQQKPGQAPRLLIYGASTRATGIPARFSGSGSGTEFTLTIGSLQSEDFAVYYCQQYKTWPRTFGQGTNVEIK
||||||||||||||||||||||||||||||.||||||||||||||||||||||||||||||||||||||||||||||||||||||||||||..|||||||||.||||
EIVMTQSPATLSVSPGERATLSCRASQSVSSNLAWYQQKPGQAPRLLIYGASTRATGIPARFSGSGSGTEFTLTIGSLQSEDFAVYYCQQYNNWPRTFGQGTKVEIK
                          ^^^^^^                 ^^^                                    ^^^^^^^^^
```

Best Alignment of Indusatumab heavy chain CDRs to a sequence from OAS
```
QVQLQQWGAGLLKPSETLSLTCAVFGGSFSGYYWSWIRQPPGKGLEWIGEINHRGNTNDNPSLKSRVTISVDTSKNQFALKLSSVTAADTAVYYCARERGYTYGNFDHWGQGTLVTVSS
..............||||||||||.||||||||||||||||||||||||||||.|.||.|||||||||.|||||||||.||||||||||||||||||||||.||.||.|||||||||||
--------------SETLSLTCAVYGGSFSGYYWSWIRQPPGKGLEWIGEINHSGSTNYNPSLKSRVTVSVDTSKNQFSLKLSSVTAADTAVYYCARERGYSYGYFDYWGQGTLVTVSS
                         ^^^^^^^^                 ^^^^^^^                                      ^^^^^^^^^^^^^
```

Best Alignment of Indusatumab light CDRs chain to a sequence from OAS
```
EIVMTQSPATLSVSPGERATLSCRASQSVSRNLAWYQQKPGQAPRLLIYGASTRATGIPARFSGSGSGTEFTLTIGSLQSEDFAVYYCQQYKTWPRTFGQGTNVEIK
||||||||||||||||||||||||||||||||||||||||||||||||||||.|||||||.|||.||||||||||..|..||.||||.|||||||||.|.||.||.|
EIVMTQSPATLSVSPGERATLSCRASQSVSRNLAWYQQKPGQAPRLLIYGASNRATGIPAGFSGGGSGTEFTLTITRLEPEDLAVYYWQQYKTWPRTLGHGTKVEVK
                          ^^^^^^                 ^^^                                    ^^^^^^^^^
```

Best Alignment of Indusatumab CDR-H3 chain to a sequence from OAS
```
QVQLQQWGAGLLKPSETLSLTCAVFGGSFSGYYWSWIRQPPGKGLEWIGEINHR-GNTNDNPSLKSRVTISVDTSKNQFALKLSSVTAADTAVYYCARERGYTYGNFDHWGQGTLVTVSS
.................|.|.||..|..||.|..||.||.|||||||...|...........|.|.|.|||.|..||...|...|..|.|||||||||||||.|||||.|||||||||||
--------------GGSLRLSCAASGFTFSSYWMSWVRQAPGKGLEWVANIKQDGSEKYYVDSVKGRFTISRDNAKNSLYLQMNSLRAEDTAVYYCARERGYSYGNFDYWGQGTLVTVSS
                         ^^^^^^^^                 ^^^^^^^^                                      ^^^^^^^^^^^^^
```

Therapeutic : Inebilizumab

Best Alignment of Inebilizumab heavy chain to a sequence from OAS
```
EVQLVESGGGLVQPGGSLRLSCAASGFTFSSSWMNWVRQAPGKGLEWVGRIYPG--DGDTNYNVKFKGRFTISRDDSKNSLYLQMNSLKTEDTAVYYCARSGFITTVRDFDYWGQGTLVTVSS
||||||||||||||||||||||||||||||..|||||||||||||||||||.....||.|.|....|||||||||||||.||||||||||||||||||||.......|.||||||||||||||
EVQLVESGGGLVQPGGSLRLSCAASGFTFSNAWMNWVRQAPGKGLEWVGRIKSKTDDGTTDYATPVKGRFTISRDDSKNTLYLQMNSLKTEDTAVYYCAREAA----RRFDYWGQGTLVTVSS
                         ^^^^^^^^                 ^^^^^^^^^^                                      ^^^^^^^^^^^^^^
```

Best Alignment of Inebilizumab light chain to a sequence from OAS
```
EIVLTQSPDFQSVTPKEKVTITCRASESVDTFGISFMNWFQQKPDQSPKLLIHEASNQGSGVPSRFSGSGSGTDFTLTINSLEAEDAATYYCQQSKEVPFTFGGGTKVEIK
||||||||||||||||||||||||||.|......|..||.||||||||||||..||..|||||||||||||||||||||||||||||||||||||||...|.|||||||||||
EIVLTQSPDFQSVTPKEKVTITCRASQSI----GSSLNWYQQKPDQSPKLLIKYASQYGSGVPSRFSGSGSGTDFTLTINSLEAEDAATYYCQQSSSLPITFGGGTKVEIK
                          ^^^^^^^^^^                 ^^^                                    ^^^^^^^^^
```

Best Alignment of Inebilizumab heavy chain CDRs to a sequence from OAS
```
EVQLVESGGGLVQPGGSLRLSCAASGFTFSSSWMNWVRQAPGKGLEWVGRIYPGDGDTNYNVKFKGRFTISRDDSKNSLYLQMNSLKTEDTAVYYCARSGFITTVRDFDYWGQGTLVTVSS
.|||.|||..||.||.|...||.|||..|||||||||.|.|||||||.|||||||||||||.||||..|...|.|....|.|..||..||.|||.||||||||||.|||||||||..||||
QVQLKESGPELVKPGASVKISCKASGYAFSSSWMNWVKQRPGKGLEWIGRIYPGDGDTNYNGKFKGKATLTADKSSSTAYMQLSSLTSEDSAVYFCARSGFITTVVDFDYWGQGTTLTVSS
                         ^^^^^^^^                 ^^^^^^^^                                      ^^^^^^^^^^^^^^
```

Best Alignment of Inebilizumab light CDRs chain to a sequence from OAS
```
EIVLTQSPDFQSVTPKEKVTITCRASESVDTFGISFMNWFQQKPDQSPKLLIHEASNQGSGVPSRFSGSGSGTDFTLTINSLEAEDAATYYCQQSKEVPFTFGGGTKVEIK
.|||||||....|......||.|||||||||.||||||||||||.|.|||||..|||||||||.|||||||||||.|.|...|..|.|.|.||||||||||||.|||.|||
DIVLTQSPASLAVSLWQRATIPCRASESVDTYGISFMNWFQQKPGQPPKLLIYAASNQGSGVPARFSGSGSGTDFSLNIHPMEEDDTAMYFCQQSKEVPFTFGSGTKLEIK
                          ^^^^^^^^^^                 ^^^                                    ^^^^^^^^^
```

Best Alignment of Inebilizumab CDR-H3 chain to a sequence from OAS
```
EVQLVESGGGLVQPGGSLRLSCAASGFTFSSSWMNWVRQAPGKGLEWVGRIYPGDGDTNYNVKFKGRFTISRDDSKNSLYLQMNSLKTEDTAVYYCARSGFITTVRDFDYWGQGTLVTVSS
.|||...|..||.||.|..|||.|||.||.|.||.||.|.||.||||.|.|.|.|..||||.||||..|...|......|.|..||..||.|||||||.|||||||||||||.||..||||
QVQLQQPGAELVKPGASVKLSCKASGYTFTSYWMQWVKQRPGQGLEWIGEIDPSDSYTNYNQKFKGKATLTGDTASSTAYMQLSSLTSEDSAVYYCARGGFITTVRDFDYWGKGTTLTVSS
                         ^^^^^^^^                 ^^^^^^^^                                      ^^^^^^^^^^^^^^
```

Therapeutic : Infliximab

Best Alignment of Infliximab heavy chain to a sequence from OAS
```
EVKLEESGGGLVQPGGSMKLSCVASGFIFSNHWMNWVRQSPEKGLEWVAEIRSKSINSATHYAESVKGRFTISRDDSKSAVYLQMTDLRTEDTGVYYCSRNYYGSTYDYWGQGTTLTVSS
|||||||||||||||||||||||||||.|||.||||||||||||||||||||.||.|.|||||||||||||||||||||.|||||..||.||||||||.|.||||..|||||||||||||
EVKLEESGGGLVQPGGSMKLSCVASGFTFSNYWMNWVRQSPEKGLEWVAEIRLKSNNYATHYAESVKGRFTISRDDSKSSVYLQMNNLRAEDTGVYYCTRHYYGSSFDYWGQGTTLTVSS
                         ^^^^^^^^                 ^^^^^^^^^^                                      ^^^^^^^^^^^
```

Best Alignment of Infliximab light chain to a sequence from OAS
```
DILLTQSPAILSVSPGERVSFSCRASQFVGSSIHWYQQRTNGSPRLLIKYASESMSGIPSRFSGSGSGTDFTLSINTVESEDIADYYCQQSHSWPFTFGSGTNLEVK
|||||||||||||||||||||||||||..|.||||||||||||||||||||||||.|||||||||||||||||||||.|||||||||||||||||||||||||.||.|
DILLTQSPAILSVSPGERVSFSCRASQSIGTSIHWYQQRTNGSPRLLIKYASESISGIPSRFSGSGSGTDFTLSINSVESEDIADYYCQQSHSWPFTFGSGTKLEIK
                          ^^^^^^                 ^^^                                    ^^^^^^^^^
```

Best Alignment of Infliximab heavy chain CDRs to a sequence from OAS
```
EVKLEESGGGLVQPGGSMKLSCVASGFIFSNHWMNWVRQSPEKGLEWVAEIRSKSINSATHYAESVKGRFTISRDDSKSAVYLQMTDLRTEDTGVYYCSRNYYGSTYDYWGQGTTLTVSS
|||||||||||||||||||||||||||.|||.|||||||||||||||||.||.||.||||.|||||.||||||||||||.|||||..||.||||.|||.|.|||||.|||||||||||||
EVKLEESGGGLVQPGGSMKLSCVASGFTFSNYWMNWVRQSPEKGLEWVAQIRLKSDNSATPYAESVQGRFTISRDDSKSSVYLQMNNLRAEDTGIYYCTRSYYGSTFDYWGQGTTLTVSS
                         ^^^^^^^^                 ^^^^^^^^^^                                      ^^^^^^^^^^^
```

Best Alignment of Infliximab light CDRs chain to a sequence from OAS
```
DILLTQSPAILSVSPGERVSFSCRASQFVGSSIHWYQQRTNGSPRLLIKYASESMSGIPSRFSGSGSGTDFTLSINTVESEDIADYYCQQSHSWPFTFGSGTNLEVK
...........|||.||.....|.|.|.||||....||.....|.|||..||....|||.||.||||||.|||.|.....||.|.||||||||||.|||.||..|.|
-----------SVSAGESSTPVCGAGQSVGSSVALFQQKPGQAPSLLIYGASTRATGIPARFGGSGSGTEFTLTISSLQAEDFAVYYCQQSHSWPVTFGQGTRVEIK
                          ^^^^^^                 ^^^                                    ^^^^^^^^^
```

Best Alignment of Infliximab CDR-H3 chain to a sequence from OAS
```
EVKLEESGGGLVQPGGSMKLSCVASGFIFSNHWMNWVRQSPEKGLEWVAEIRSKSINSATHYAESVKGRFTISRDDSKSAVYLQMTDLRTEDTGVYYCSRNYYGSTYDYWGQGTTLTVSS
...|...|..|..||...|.||.|||..|....|.||.|.|.|||.|...|..........||...||||..|...|.|..|||...|..|||..|.||||||||.||||||||||||||
QIQLVQYGPELKKPGETVKISCKASGYTFTTYGMSWVKQAPGKGLKWMGWINTY--AGVPTYADDFKGRFAFSLETSASTAYLQINNLKNEDTATYFCSRNYYGSSYDYWGQGTTLTVSS
                         ^^^^^^^^                 ^^^^^^^^^^                                      ^^^^^^^^^^^
```

Therapeutic : Inotuzumab

Best Alignment of Inotuzumab heavy chain to a sequence from OAS
```
EVQLVQSGAEVKKPGASVKVSCKASGYRFTNYWIHWVRQAPGQGLEWIGGINPGNNYATYRRKFQGRVTMTADTSTSTVYMELSSLRSEDTAVYYCTREGYGN------YGAWFAYWGQGTLVTVSS
.||||||||||||||||||||||||||.||.|..|||||||||||||.|.|||.....||..|||||||||.||||||||||||||||||||||||.|||.|.......|.|.|.||||||||||||
QVQLVQSGAEVKKPGASVKVSCKASGYTFTSYYMHWVRQAPGQGLEWMGIINPSGGSTTYAQKFQGRVTMTRDTSTSTVYMELSSLRSEDTAVYYCAREGDGYYDFSGYYYAPFQYWGQGTLVTVSS
                         ^^^^^^^^                 ^^^^^^^^                                      ^^^^^^^^^^^^^^^^^^^^
```

Best Alignment of Inotuzumab light chain to a sequence from OAS
```
DVQVTQSPSSLSASVGDRVTITCRSSQSLANSYGNTFLSWYLHKPGKAPQLLIYGISNRFSGVPDRFSGSGSGTDFTLTISSLQPEDFATYYCLQGTHQPYTFGQGTKVEIK
||.|||.|.||..|.||.|.|.||||||||||||||.||||||||||.||||||||||||||||||||||||||||||.||...|||...||||||||||||||.|||.|||
DVVVTQTPLSLPVSFGDQVSISCRSSQSLANSYGNTYLSWYLHKPGKSPQLLIYGISNRFSGVPDRFSGSGSGTDFTLKISTIKPEDLGMYYCLQGTHQPYTFGGGTKLEIK
                          ^^^^^^^^^^^                 ^^^                                    ^^^^^^^^^
```

Best Alignment of Inotuzumab heavy chain CDRs to a sequence from OAS
```
EVQLVQSGAEVKKPGASVKVSCKASGYRFTNYWIHWVRQAPGQGLEWIGGINPGNNYATYRRKFQGRVTMTADTSTSTVYMELSSLRSEDTAVYYCTREGYGNYGAWFAYWGQGTLVTVSS
||||.|||.....||||||.|||||||.||.||.|||.|.|||||||||.|.|||....|..||.|....||.|||||.|||||||..||.|||||||||||||||||||||||||||||.
```

EVQLQQSGTVLARPGASVKMSCKASGYTFTSYWMHWVKQRPGQGLEWIGAIYPGNSDTSYNQKFKGKAKLTAVTSTSTAYMELSSLTNEDSAVYYCTREGYGNYGAWFAYWGQGTLVTVSA
^^^^^^^^ ^^^^^^^^ ^^^^^^^^^^^^^^

Best Alignment of Inotuzumab light CDRs chain to a sequence from OAS
DVQVTQSPSSLSASVGDRVTITCRSSQSLANSYGNTFLSWYLHKPGKAPQLLIYGISNRFSGVPDRFSGSGSGTDFTLTISSLQPEDFATYYCLQGTHQPYTFGQGTKVEIK
||.|||.|.||..|.||.|.|.||||||||||||||||||||||||..||.|||||||||||||||||||||||||||.||...|||...||||||||||.|||.|||.|||
DVVVTQTPLSLPVSFGDQVSISCRSSQSLANSYGNTFLSWYLHKPGQSPQFLIYGISNRFSGVPDRFSGSGSGTDFTLKISTIKPEDLGMYYCLQGTHQPFTFGSGTKLEIK
^^^^^^^^^^^ ^^^ ^^^^^^^^^

Best Alignment of Inotuzumab CDR-H3 chain to a sequence from OAS
EVQLVQSGAEVKKPGASVKVSCKASGYRFTNYWIHWVRQAPGQGLEWIGGINPGNNYATYRRKFQGRVTMTADTSTSTVYMELSSLRSEDTAVYYCTREGYGNYGAWFAYWGQGTLVTVSS
||||.|||.....||||||.|||||||.||.||.|||.|.|||||||||.|.|||....|..||.|....||.|||||.|||||||..||.|||||||||||||||||||||||||||||.
EVQLQQSGTVLARPGASVKMSCKASGYTFTSYWMHWVKQRPGQGLEWIGAIYPGNSDTSYNQKFKGKAKLTAVTSTSTAYMELSSLTNEDSAVYYCTREGYGNYGAWFAYWGQGTLVTVSA
^^^^^^^^ ^^^^^^^^ ^^^^^^^^^^^^^^

Therapeutic : Intetumumab

Best Alignment of Intetumumab heavy chain to a sequence from OAS
QVQLVESGGGVVQPGRSRRLSCAASGFTFSRYTMHWVRQAPGKGLEWVAVISFDGSNKYYVDSVKGRFTISRDNSENTLYLQVNILRAEDTAVYYCAREARG----SYAFDIWGQGTMVTVSS
||||||||||||||||.||||||||||||.|.||||||||||||||||||||.|||||||.||||||||||||||.||||||.|.|||||||||||||||||....|.|||||||||||||||
QVQLVESGGGVVQPGRSLRLSCAASGFTFSSYAMHWVRQAPGKGLEWVAVISYDGSNKYYADSVKGRFTISRDNSKNTLYLQMNSLRAEDTAVYYCAREARGYGDYSDAFDIWGQGTMVTVSS
^^^^^^^^ ^^^^^^^^ ^^^^^^^^^^^^^^^^

Best Alignment of Intetumumab light chain to a sequence from OAS
EIVLTQSPATLSLSPGERATLSCRASQSVSSYLAWYQQKPGQAPRLLIYDASNRATGIPARFSGSGSGTDFTLTISSLEPEDFAVYYCQQRSNWPPFTFGPGTKVDIK
|||||||||||||||||||||||||||||||||||||||||||||||||||||||||||||||||||||||||||||||||||||||||||||||||||||||||||||
EIVLTQSPATLSLSPGERATLSCRASQSVSSYLAWYQQKPGQAPRLLIYDASNRATGIPARFSGSGSGTDFTLTISSLEPEDFAVYYCQQRSNWPPFTFGPGTKVDIK
^^^^^^ ^^^ ^^^^^^^^^^

Best Alignment of Intetumumab heavy chain CDRs to a sequence from OAS
QVQLVESGGGVVQPGRSRRLSCAASGFTFSRYTMHWVRQAPGKGLEWVAVISFDGSNKYYVDSVKGRFTISRDNSENTLYLQVNILRAEDTAVYYCAREARGSYAFDIWGQGTMVTVSS
..............|.|.||||||||||||.|.||||||||||||||||||||.|||||||.||||||||||||||.||||||.|.|||||||||||||||.||||||||||||||||||
--------------GGSLRLSCAASGFTFSSYAMHWVRQAPGKGLEWVAVISYDGSNKYYADSVKGRFTISRDNSKNTLYLQMNSLRAEDTAVYYCARETRGSYAFDIWGQGTMVTVSS
^^^^^^^^ ^^^^^^^^ ^^^^^^^^^^^^

Best Alignment of Intetumumab light CDRs chain to a sequence from OAS
EIVLTQSPATLSLSPGERATLSCRASQSVSSYLAWYQQKPGQAPRLLIYDASNRATGIPARFSGSGSGTDFTLTISSLEPEDFAVYYCQQRSNWPPFTFGPGTKVDIK
||||||||||||||||||||||||||||||||||||||||||||||||||||||||||||||||||||||||||||||||||||||||||||||||||||.||||.||
EIVLTQSPATLSLSPGERATLSCRASQSVSSYLAWYQQKPGQAPRLLIYDASNRATGIPARFSGSGSGTDFTLTISSLEPEDFAVYYCQQRSNWPPFTFGGGTKVEIK
^^^^^^ ^^^ ^^^^^^^^^^

Best Alignment of Intetumumab CDR-H3 chain to a sequence from OAS
QVQLVESGGGVVQPGRSRRLSCAASGFTFSRYTMHWVRQAPGKGLEWVAVISFDGSNKYYVDSVKGRFTISRDNSENTLYLQVNILRAEDTAVYYCAREARGSYAFDIWGQGTMVTVSS
................|.||||||||||||.|.|||||||.|||||||..|.......||..||||||||||.|..|.||||.|.|||.||||||||||||||.||||||||||||||||
----------------SLRLSCAASGFTFSSYDMHWVRQATGKGLEWVSAIGTA-GDTYYPGSVKGRFTISRENAKNSLYLQMNSLRAGDTAVYYCAREARGGYAFDIWGQGTMVTVSS
^^^^^^^^ ^^^^^^^^ ^^^^^^^^^^^^

Therapeutic : Ipilimumab

Best Alignment of Ipilimumab heavy chain to a sequence from OAS
QVQLVESGGGVVQPGRSLRLSCAASGFTFSSYTMHWVRQAPGKGLEWVTFISYDGNNKYYADSVKGRFTISRDNSKNTLYLQMNSLRAEDTAIYYCARTGWL--------GPFDYWGQGTLVTVSS
||||||||||||||||||||||||||||||||.|||||||||||||||||.|||||.||||||||||||||||||||||||||||||||||||||||||||.|||||.|..........|||||||||||||||
QVQLVESGGGVVQPGRSLRLSCAASGFTFSSYAMHWVRQAPGKGLEWVTVISYDGSNKYYADSVKGRFTISRDNSKNTLYLQMNSLRAEDTAVYYCARAGLGDYGDYRRPGPFDYWGQGTLVTVSS
^^^^^^^^ ^^^^^^^^ ^^^^^^^^^^^^^^^^^^^

Best Alignment of Ipilimumab light chain to a sequence from OAS
EIVLTQSPGTLSLSPGERATLSCRASQSVGSSYLAWYQQKPGQAPRLLIYGAFSRATGIPDRFSGSGSGTDFTLTISRLEPEDFAVYYCQQYGSSPWTFGQGTKVEIK
||||||||||||||||||||||||||||||||||||||||||||||||||||||||||||||||||||||||||||||||||||||||.||||||||||||||||||||||
EIVLTQSPGTLSLSPGERATLSCRASQSVGSSYLAWYQQKPGQAPRLLIYGAFSRATGIPDRFSGSGSGTDFTLTISRLEPEDFALYYCQQYGSSPWTFGQGTKVEIK
^^^^^^^ ^^^ ^^^^^^^^^

Best Alignment of Ipilimumab heavy chain CDRs to a sequence from OAS
QVQLVESGGGVVQPGRSLRLSCAASGFTFSSYTMHWVRQAPGKGLEWVTFISYDGNNKYYADSVKGRFTISRDNSKNTLYLQMNSLRAEDTAIYYCARTGWLGPFDYWGQGTLVTVSS
..............|.||||||||||||||||.||||||||||||||||..|||||.||||||||||||||||||||||||||||||||||||.||||||.|||||||||||||||||||
--------------GGSLRLSCAASGFTFSSYAMHWVRQAPGKGLEWVAVISYDGSNKYYADSVKGRFTISRDNSKNTLYLQMNSLRAEDTAVYYCARTLWLGPFDYWGQGTLVTVSS
^^^^^^^^ ^^^^^^^^ ^^^^^^^^^^^

Best Alignment of Ipilimumab light CDRs chain to a sequence from OAS
EIVLTQSPGTLSLSPGERATLSCRASQSVGSSYLAWYQQKPGQAPRLLIYGAFSRATGIPDRFSGSGSGTDFTLTISRLEPEDFAVYYCQQYGSSPWTFGQGTKVEIK
||||||||||||||||||||||||||||||||||||||||||||||||||||||||||||||||||||||||||||||||||||||||.||||||||||||||||||||||
EIVLTQSPGTLSLSPGERATLSCRASQSVGSSYLAWYQQKPGQAPRLLIYGAFSRATGIPDRFSGSGSGTDFTLTISRLEPEDFALYYCQQYGSSPWTFGQGTKVEIK
^^^^^^^ ^^^ ^^^^^^^^^

Best Alignment of Ipilimumab CDR-H3 chain to a sequence from OAS
QVQLVESGGGVVQPGRSLRLSCAASGFTFSSYTMHWVRQAPGKGLEWVTFISYDGNNKYYADSVKGRFTISRDNSKNTLYLQMNSLRAEDTAIYYCARTGWLGPFDYWGQGTLVTVSS
..................||.|||.|||||...|||||||.||||...|........||....||.||..|.|..|.|....|||.||||.||||||.||||||||||||||||||
------------------KISCKASGGTFSSYAISWVRQAPGQGLEWMGGIIPIFGTANYAQKFQGRVTITADESTSTAYMELSSLRSEDTAVYYCARTQWLGPFDYWGQGTLVTVSS
^^^^^^^^ ^^^^^^^^ ^^^^^^^^^^^

Therapeutic : Iratumumab

Best Alignment of Iratumumab heavy chain to a sequence from OAS
QVQLQQWGAGLLKPSETLSLTCAVYGGSFSAYYWSWIRQPPGKGLEWIGDINHGGGTNYNPSLKSRVTISVDTSKNQFSLKLNSVTAADTAVYYCASL------TAYWGQGSLVTVSS
||||||||||||||||||||||||||||||.||||||||||||||||||.|||.|.||||||||||||||||||||||||||||||||||||||||||||.......||||||.||||||
QVQLQQWGAGLLKPSETLSLTCAVYGGSFSGYYWSWIRQPPGKGLEWIGEINHSGSTNYNPSLKSRVTISVDTSKNQFSLKLNSVTAADTAVYYCASLFMIVEDMAYWGQGTLVTVSS
^^^^^^^^ ^^^^^^^ ^^^^^^^^^^^^

Best Alignment of Iratumumab light chain to a sequence from OAS
DIQMTQSPTSLSASVGDRVTITCRASQGISSWLTWYQQKPEKAPKSLIYAASSLQSGVPSRFSGSGSGTDFTLTISSLQPEDFATYYCQQYDSYPITFGQGTRLEIK
||||||||.||||||||||||||||||||||||.|||||||||||||||||||||||||||||||||||||||||||||||||||||||||||||||||||||||||
DIQMTQSPSSLSASVGDRVTITCRASQGISSWLAWYQQKPEKAPKSLIYAASSLQSGVPSRFSGSGSGTDFTLTISSLQPEDFATYYCQQYDSYPITFGQGTRLEIK
^^^^^^ ^^^ ^^^^^^^^^

Best Alignment of Iratumumab heavy chain CDRs to a sequence from OAS
QVQLQQWGAGLLKPSETLSLTCAVYGGSFSAYYWSWIRQPPGKGLEWIGDINHGGGTNYNPSLKSRVTISVDTSKNQFSLKLNSVTAADTAVYYCASLTAYWGQGSLVTVSS
.............|||||||||||||||||.||||||||||||||||||.|||||.|||||||||||||||||||||||||||.|||||||||||||.||||||||.||||||
-------------PSETLSLTCAVYGGSFSGYYWSWIRQPPGKGLEWIGEINHGGSTNYNPSLKSRVTISVDTSKNQFSLKLSSVTAADTAVYYCARLTAYWGQGTLVTVSS
^^^^^^^^ ^^^^^^^ ^^^^^^

Best Alignment of Iratumumab light CDRs chain to a sequence from OAS
DIQMTQSPTSLSASVGDRVTITCRASQGISSWLTWYQQKPEKAPKSLIYAASSLQSGVPSRFSGSGSGTDFTLTISSLQPEDFATYYCQQYDSYPITFGQGTRLEIK
|.|.||||.|.||||||||||||||||||||||.||||||||||||||||||||||||||||||||||||||||||||||||||||||||||||||||||||||||
DSQLTQSPSSPSASVGDRVTITCRASQGISSWLAWYQQKPEKAPKSLIYAASSLQSGVPSRFSGSGSGTDFTLTISSLQPEDFATYYCQQYDSYPITFGQGTRLEIK
^^^^^^ ^^^ ^^^^^^^^^
Best Alignment of Iratumumab CDR-H3 chain to a sequence from OAS
QVQLQQWGAGLLKPSETLSLTCAVYGGSFSAYYWSWIRQPPGKGLEWIGDINHGGGTNYNPSLKSRVTISVDTSKNQFSLKLNSVTAADTAVYYCASLTAYWGQGSLVTVSS
.............|||||||||.|.|||.|.||||||||||||||||||.|...|.||||||||||||||||||||||||||.||||||||||||||||||||||.||||||

-------------PSETLSLTCTVSGGSISSYYWSWIRQPPGKGLEWIGYIYYSGSTNYNPSLKSRVTISVDTSKNQFSLKLSSVTAADTAVYYCASLTAYWGQGTLVTVSS
                         ^^^^^^^^                 ^^^^^^^                                      ^^^^^^

Therapeutic : Isatuximab

Best Alignment of Isatuximab heavy chain to a sequence from OAS
QVQLVQSGAEVAKPGTSVKLSCKASGYTFTDYWMQWVKQRPGQGLEWIGTIYPGDGDTGYAQKFQGKATLTADKSSKTVYMHLSSLASEDSAVYYCARGDYYG---SNSLDYWGQGTSVTVSS
||||.|||||.|.||.||||||||||||||.||||||||||||||||||.||||||||.|.|||.|||||||||||.|.||.|||||||||||||||||||||...|...|||||||||||||
QVQLQQSGAELARPGASVKLSCKASGYTFTSYWMQWVKQRPGQGLEWIGAIYPGDGDTRYTQKFKGKATLTADKSSSTAYMQLSSLASEDSAVYYCARGDYYGSSYSYAMDYWGQGTSVTVSS
                         ^^^^^^^^                 ^^^^^^^^                                      ^^^^^^^^^^^^^^^^

Best Alignment of Isatuximab light chain to a sequence from OAS
DIVMTQSHLSMSTSLGDPVSITCKASQDVSTVVAWYQQKPGQSPRRLIYSASYRYIGVPDRFTGSGAGTDFTFTISSVQAEDLAVYYCQQHYSPPYTFGGGTKLEIK
||||||||..||||.||.|||||||||||||.||||||||||||.||||||||||.||||||||||.||||||||||||||||||||||||||.|||||||||||||
DIVMTQSHKFMSTSVGDRVSITCKASQDVSTAVAWYQQKPGQSPKRLIYSASYRYTGVPDRFTGSGSGTDFTFTISSVQAEDLAVYYCQQHYSTPYTFGGGTKLEIK
                          ^^^^^^                 ^^^                                    ^^^^^^^^^

Best Alignment of Isatuximab heavy chain CDRs to a sequence from OAS
QVQLVQSGAEVAKPGTSVKLSCKASGYTFTDYWMQWVKQRPGQGLEWIGTIYPGDGDTGYAQKFQGKATLTADKSSKTVYMHLSSLASEDSAVYYCARGDYYGSNSLDYWGQGTSVTVSS
.|||..||||.|.||.||||||||||||||.||||||||||||||||||.||||||||.|.|||.|||||||||||.|.||.|||||||||||||||||||||||..|||||||..||||
-VQLQESGAELARPGASVKLSCKASGYTFTSYWMQWVKQRPGQGLEWIGAIYPGDGDTRYTQKFKGKATLTADKSSSTAYMQLSSLASEDSAVYYCARGDYYGSNYFDYWGQGTTLTVSS
                         ^^^^^^^^                 ^^^^^^^^                                      ^^^^^^^^^^^^^

Best Alignment of Isatuximab light CDRs chain to a sequence from OAS
DIVMTQSHLSMSTSLGDPVSITCKASQDVSTVVAWYQQKPGQSPRRLIYSASYRYIGVPDRFTGSGAGTDFTFTISSVQAEDLAVYYCQQHYSPPYTFGGGTKLEIK
||||||||..||||.||.||||||||||||||||||||||||||..|||||||||.|||.||||||.||||||||||||||||||||||||||.|||||||||||||
DIVMTQSHKFMSTSVGDRVSITCKASQDVSTVVAWYQQKPGQSPKLLIYSASYRYTGVPGRFTGSGSGTDFTFTISSVQAEDLAVYYCQQHYSTPYTFGGGTKLEIK
                          ^^^^^^                 ^^^                                    ^^^^^^^^^

Best Alignment of Isatuximab CDR-H3 chain to a sequence from OAS
QVQLVQSGAEVAKPGTSVKLSCKASGYTF--TDYWMQWVKQRPGQGLEWIGTIYPGDGDTGYAQKFQGKATLTADKSSKTVYMHLSSLASEDSAVYYCARGDYYGSNSLDYWGQGTSVTVSS
...................|.|..||.......|...|..|.||.||||||.||...|.|.|........|...|.|.......|||....|.|||||||||||||.|||||||||.|||||
----------------TLSLTCTVSGGSISSGGYYWSWIRQHPGKGLEWIGYIYYS-GSTYYNPSLKSRVTISVDTSKNQFSLKLSSVTAADTAVYYCARGDYYGSGSLDYWGQGTLVTVSS
                         ^^^^^^^^^^                 ^^^^^^^^                                      ^^^^^^^^^^^^^

Therapeutic : Itolizumab

Best Alignment of Itolizumab heavy chain to a sequence from OAS
EVQLVESGGGLVKPGGSLKLSCAASGFKFSRYAMSWVRQAPGKRLEWVATISSGGSYIYYPDSVKGRFTISRDNVKNTLYLQMSSLRSEDTAMYYCARRDYD--LDYFDSWGQGTLVTVSS
|||||||||||||||||||||||||||.||.||||||||.|.|||||||||||||||.||||||||||||||||.|||||||||||||||||||||||||||...||||.|||||..||||
EVQLVESGGGLVKPGGSLKLSCAASGFTFSSYAMSWVRQTPEKRLEWVATISSGGSYTYYPDSVKGRFTISRDNAKNTLYLQMSSLRSEDTAMYYCARRDYDGSSDYFDYWGQGTTLTVSS
                         ^^^^^^^^                 ^^^^^^^^                                      ^^^^^^^^^^^^^^

Best Alignment of Itolizumab light chain to a sequence from OAS
DIQMTQSPSSLSASVGDRVTITCKASRDIRSYLTWYQQKPGKAPKTLIYYATSLADGVPSRFSGSGSGQDYSLTISSLESDDTATYYCLQHGESPFTLGSGTKLEIK
||.||||||||.||.|.|||||||||.||.|||.||||||.|.||||||||||||||||||||||||||||||||||||||||||||||||||||||.|||||||||
DIKMTQSPSSLYASLGERVTITCKASQDIKSYLSWYQQKPWKSPKTLIYYATSLADGVPSRFSGSGSGQDYSLTISSLESDDTATYYCLQHGESPFTFGSGTKLEIK
                          ^^^^^^                 ^^^                                    ^^^^^^^^^

Best Alignment of Itolizumab heavy chain CDRs to a sequence from OAS
EVQLVESGGGLVKPGGSLKLSCAASGFKFSRYAMSWVRQAPGKRLEWVATISSGGSYIYYPDSVKGRFTISRDNVKNTLYLQMSSLRSEDTAMYYCARRDYDLDYFDSWGQGTLVTVSS
||||.||||||||||||||||||||||.||.||||||||.|.|||||||||||||||.||||||||||||||||.||||||||||||||||||||||||||||.|||.|||||..||||
EVQLQESGGGLVKPGGSLKLSCAASGFTFSSYAMSWVRQTPEKRLEWVATISSGGSYTYYPDSVKGRFTISRDNAKNTLYLQMSSLRSEDTAMYYCARRDYDLYYFDYWGQGTTLTVSS
                         ^^^^^^^^                 ^^^^^^^^                                      ^^^^^^^^^^^^

Best Alignment of Itolizumab light CDRs chain to a sequence from OAS
DIQMTQSPSSLSASVGDRVTITCKASRDIRSYLTWYQQKPGKAPKTLIYYATSLADGVPSRFSGSGSGQDYSLTISSLESDDTATYYCLQHGESPFTLGSGTKLEIK
||.|||||||..||.|.||||||||||||.|||.||||||.|.||||||||||||||||||||||||||||||||||||||||||||||||||||.|.|.|.|||||||
DIKMTQSPSSMYASLGERVTITCKASRDIKSYLSWYQQKPWKSPKTLIYYATSLADGVPSRFSGSGSGQDYSLTISSLESDDTATYYCLQHGESPPTFGGGTKLEIK
                          ^^^^^^                 ^^^                                    ^^^^^^^^^

Best Alignment of Itolizumab CDR-H3 chain to a sequence from OAS
EVQLVESGGGLVKPGGSLKLSCAASGFKF-SRYAMSWVRQAPGKRLEWVATISSGGSYIYYPDSVKGRFTISRDNVKNTLYLQMSSLRSEDTAMYYCARRDYDLDYFDSWGQGTLVTVSS
||||||||.|||||..||.|.|........|.||..|.||.||..|||...||.......|..|.|.|..|.||..||...||..|...||||.|||||||||.||||.|||||..||||
EVQLVESGPGLVKPSQSLSLTCTVTSYSITSDYAWNWIRQFPGNKLEWMGYISYS-GSTSYNPSLKSRISITRDTSKNQFFLQLNSVTTEDTATYYCARRDYDYDYFDYWGQGTTLTVSS
                         ^^^^^^^^^                 ^^^^^^^^                                      ^^^^^^^^^^^^

Therapeutic : Ixekizumab

Best Alignment of Ixekizumab heavy chain to a sequence from OAS
QVQLVQSGAEVKKPGSSVKVSCKASGYSFTDYHIHWVRQAPGQGLEWMGVINPMYGTTDYNQRFKGRVTITADESTSTAYMELSSLRSEDTAVYYCARYDYF-----TGTGVYWGQGTLVTVSS
|||||||||||||||.|||||||||||.||.|.||||||||||||||||.|||..|||.|.|.|.|||||||||||||||||||||||||||||||||..||......|...||||||||||||
QVQLVQSGAEVKKPGASVKVSCKASGYTFTGYYIHWVRQAPGQGLEWMGWINPIFGTTNYAQKFQGRVTITADESTSTAYMELSSLRSEDTAVYYCARDQYFGSGSLIGRFDYWGQGTLVTVSS
                         ^^^^^^^^                 ^^^^^^^^                                      ^^^^^^^^^^^^^^^^^

Best Alignment of Ixekizumab light chain to a sequence from OAS
DIVMTQTPLSLSVTPGQPASISCRSSRSLVHSRGNTYLHWYLQKPGQSPQLLIYKVSNRFIGVPDRFSGSGSGTDFTLKISRVEAEDVGVYYCSQSTHLPFTFGQGTKLEIK
|||||||||||||||||||||.||||.|||||.|.|||.|||||||||||||||.||.||.||||||||||||||||||||||||||||||||.|.||||.|||||||||||
DIVMTQTPLSLSVTPGQPASIFCRSSQSLVHSDGSTYLYWYLQKPGQSPQLLIYEVSKRFSGVPDRFSGSGSGTDFTLKISRVEAEDVGVYYCMQGTHLPDTFGQGTKLEIK
                          ^^^^^^^^^^^                 ^^^                                    ^^^^^^^^^

Best Alignment of Ixekizumab heavy chain CDRs to a sequence from OAS
QVQLVQSGAEVKKPGSSVKVSCKASGYSFTDYHIHWVRQAPGQGLEWMGVINPMYGTTDYNQRFKGRVTITADESTSTAYMELSSLRSEDTAVYYCARYDYFTGTGVYWGQGTLVTVSS
.|||.|||.|..|||.|||.||||||||||||...||.|..|..|||.|||||.||||.|||.|||..|.|.|.|.|||||.|.||.|||.||||||||||..||..||||||..||||
EVQLQQSGPELVKPGASVKISCKASGYSFTDYNMNWVKQSNGKSLEWIGVINPNYGTTSYNQKFKGKATLTVDQSSSTAYMQLNSLTSEDSAVYYCARYDYDRGTFDYWGQGTTLTVSS
                         ^^^^^^^^                 ^^^^^^^^                                      ^^^^^^^^^^^^

Best Alignment of Ixekizumab light CDRs chain to a sequence from OAS
DIVMTQTPLSLSVTPGQPASISCRSSRSLVHSRGNTYLHWYLQKPGQSPQLLIYKVSNRFIGVPDRFSGSGSGTDFTLKISRVEAEDVGVYYCSQSTHLPFTFGQGTKLEIK
|||.||..|||.|..|..||||||||.|||||.||||||||||||||||.||||||||||.||||||||||||||||||||||||||.|||.||||||||||||.|||||||
DIVLTQSTLSLPVSLGDQASISCRSSQSLVHSNGNTYLHWYLQKPGQSPKLLIYKVSNRFSGVPDRFSGSGSGTDFTLKISRVEAEDLGVYFCSQSTHLPFTFGSGTKLEIK
                          ^^^^^^^^^^^                 ^^^                                    ^^^^^^^^^

Best Alignment of Ixekizumab CDR-H3 chain to a sequence from OAS
QVQLVQSGAEVKKPGSSVKVSCKASGYSFTDYHIHWVRQAPGQGLEWMGVINPMYGTTDYNQRFKGRVTITADESTSTAYMELSSLRSEDTAVYYCARYDYFTGTGVYWGQGTLVTVSS
.....||||||||||.|.|.|||.||||||.|.|.||||.||.||||||.|.|....|.|...|.|.|||.||.|.||||...|||...|||.||||||||..|.||||||||||||||
-----QSGAEVKKPGESLKISCKGSGYSFTNYWIAWVRQMPGKGLEWMGIIYPGDSDTRYSPSFQGQVTISADKSISTAYLQWSSLKASDTAMYYCARYDYDSGGGVYWGQGTLVTVSS
                         ^^^^^^^^                 ^^^^^^^^                                      ^^^^^^^^^^^^

Therapeutic : Lacnotuzumab

Best Alignment of Lacnotuzumab heavy chain to a sequence from OAS
QVQLQESGPGLVKPSQTLSLTCTVSDYSITSDYAWNWIRQFPGKGLEWMGYISYSGSTSYNPSLKSRITISRDTSKNQFSLQLNSVTAADTAVYYCASFDYA-HAMDYWGQGTTVTVSS
.|||||||||||||||.|||||||..|||||||||||||||||..|||||||||||||||||||||||.|.||||||||.|||||||..|||.|||||.|||..|||||||||.|||||
DVQLQESGPGLVKPSQSLSLTCTVTGYSITSDYAWNWIRQFPGNKLEWMGYISYSGSTSYNPSLKSRISITRDTSKNQFFLQLNSVTTEDTATYYCASYDYAYYAMDYWGQGTSVTVSS
                         ^^^^^^^^^                 ^^^^^^^                                      ^^^^^^^^^^^^

Best Alignment of Lacnotuzumab light chain to a sequence from OAS
DIVLTQSPAFLSVTPGEKVTFTCQASQSIGTSIHWYQQKTDQAPKLLIKYASESISGIPSRFSGSGSGTDFTLTISSVEAEDAADYYCQQINSWPTTFGGGTKLEIK
||.||||||.|||.|||.|.|.|.||||||||||||||.|...|.||||||||||||||||||||||||||||||.|||.||.|||||||.||||||||||||||||
DILLTQSPAILSVSPGERVSFSCRASQSIGTSIHWYQQRTNDSPRLLIKYASESISGIPSRFSGSGSGTDFTLTINSVESEDIADYYCQQSNSWPTTFGGGTKLEIK
^^^^^^ ^^^ ^^^^^^^^^

Best Alignment of Lacnotuzumab heavy chain CDRs to a sequence from OAS
QVQLQESGPGLVKPSQTLSLTCTVSDYSITSDYAWNWIRQFPGKGLEWMGYISYSGSTSYNPSLKSRITISRDTSKNQFSLQLNSVTAADTAVYYCASFDYAHAMDYWGQGTTVTVSS
.|.|||||||||||||.|||||||..|||||||||||||||||..|||||||||||||||||||||||.|.||||||||.|||||||..|||.|||||.|||.|||||||||.|||||
EVKLQESGPGLVKPSQSLSLTCTVTGYSITSDYAWNWIRQFPGNKLEWMGYISYSGSTSYNPSLKSRISITRDTSKNQFFLQLNSVTTEDTATYYCASYDYAYAMDYWGQGTSVTVSS
^^^^^^^^^ ^^^^^^^ ^^^^^^^^^^^

Best Alignment of Lacnotuzumab light CDRs chain to a sequence from OAS
DIVLTQSPAFLSVTPGEKVTFTCQASQSIGTSIHWYQQKTDQAPKLLIKYASESISGIPSRFSGSGSGTDFTLTISSVEAEDAADYYCQQINSWPTTFGGGTKLEIK
||.||||||.|||.|||.|.|.|.||||||||||||||.|...|.||||||||||||||||||||||||||||.|.|||.||.|.|||||.||||||||.|||||||
DILLTQSPAILSVSPGERVSFSCRASQSIGTSIHWYQQRTNGSPRLLIKYASESISGIPSRFSGSGSGTDFTLSINSVESEDIAEYYCQQSNSWPTTFGSGTKLEIK
^^^^^^ ^^^ ^^^^^^^^^

Best Alignment of Lacnotuzumab CDR-H3 chain to a sequence from OAS
QVQLQESGPGLVKPSQTLSLTCTVSDYSITSDYAWNWIRQFPGKGLEWMGYISYS-GSTSYNPSLKSRITISRDTSKNQFSLQLNSVTAADTAVYYCASFDYAHAMDYWGQGTTVTVSS
|||||..|..||||.......|..|.|.....|...|..|.||.||||.|.|....|||.||...||..|...|||......||.|.|..|.||||||||||||.|||||||||.|||||
QVQLQQPGAELVKPGASVKMSCKASGYTF-TSYWITWVKQRPGQGLEWIGDIYPGSGSTNYNEKFKSKATLTVDTSSSTAYNQLSSLTTGDSAVYYCASFDYAYAMDYWGQGTSVTVSS
^^^^^^^^^ ^^^^^^^^ ^^^^^^^^^^^

Therapeutic : Lampalizumab

Best Alignment of Lampalizumab heavy chain to a sequence from OAS
EVQLVQSGPELKKPGASVKVSCKASGYTFTNYGMNWVRQAPGQGLEWMGWINTYTGETTYADDFKGRFVFSLDTSVSTAYLQISSLKAEDTAVYYCEREG---GVNNWGQGTLVTVSS
.|||||||.|||||||||||||||||||||.|.||||||||||||||||||||.||.||||..|.|||||||||||||||||||||||||||||||||.|||....|..|||||||||||
QVQLVQSGSELKKPGASVKVSCKASGYTFTSYAMNWVRQAPGQGLEWMGWINTNTGNTTYAQGFTGRFVFSLDTSVSTAYLQISSLKAEDTAVYYCAREGGFRSVDYWGQGTLVTVSS
^^^^^^^^ ^^^^^^^^ ^^^^^^^^^^^

Best Alignment of Lampalizumab light chain to a sequence from OAS
DIQVTQSPSSLSASVGDRVTITCITSTDIDDDMNWYQQKPGKVPKLLISGGNTLRPGVPSRFSGSGSGTDFTLTISSLQPEDVATYYCLQSDSLPYTFGQGTKVEIK
|||.|||||||||||||||||||..|.||.....||||||||||||||.|..||..||||||||||||||||||||||||||||||||.|.|.||||||||||.|||
DIQMTQSPSSLSASVGDRVTITCRASQDISNYLAWYQQKPGKVPKLLIYGASTLQSGVPSRFSGSGSGTDFTLTISSLQPEDVATYYCQQYDNLPYTFGQGTKLEIK
^^^^^^ ^^^ ^^^^^^^^^

Best Alignment of Lampalizumab heavy chain CDRs to a sequence from OAS
EVQLVQSGPELKKPGASVKVSCKASGYTFTNYGMNWVRQAPGQGLEWMGWINTYTGETTYADDFKGRFVFSLDTSVSTAYLQISSLKAEDTAVYYCEREGGVNNWGQGTLVTVSS
..|||||||||||||..||.|||||||||||||||||.||||.||.|||||||||||.||||||||||.|||.||.|||||||..||.||||.|...||||...|||||..||||
QIQLVQSGPELKKPGETVKISCKASGYTFTNYGMNWVKQAPGKGLKWMGWINTYTGEPTYADDFKGRFAFSLETSASTAYLQINNLKNEDTATYFWAREGGTDYWGQGTTLTVSS
^^^^^^^^ ^^^^^^^^ ^^^^^^^^

Best Alignment of Lampalizumab light CDRs chain to a sequence from OAS
DIQVTQSPSSLSASVGDRVTITCITSTDIDDDMNWYQQKPGKVPKLLISGGNTLRPGVPSRFSGSGSGTDFTLTISSLQPEDVATYYCLQSDSLPYTFGQGTKVEIK
...|||||.|||...|..|||.|||.||||||||||||||||..|||||||||||||||||||||||.||.||||..||.....||||.|||||||||.|||||.|||.|||
ETTVTQSPASLSVATGEKVTIRCITGTDIDDDMNWYQQKPGEPPKLLISGGNTLRPGVPSRFSSSGYGTDFVFTIDNTLSEDVADYYCLQSDSMPYTFGGGTKLEIK
^^^^^^ ^^^ ^^^^^^^^^

Best Alignment of Lampalizumab CDR-H3 chain to a sequence from OAS
EVQLVQSGPELKKPGASVKVSCKASGYTFTNYGMNWVRQAPGQGLEWMGWINTYTGETTYADDFKGRFVFSLDTSVSTAYLQISSLKAEDTAVYYCEREGGVNNWGQGTLVTVSS
.....................|...|..|..|...|.||.||.||||.|.||...|.|......|.|...|.|||.....|..||..|.|||||||.||||||.||.||.|||||
----------------TLSLTCAVYGGSFSGYYWSWIRQPPGKGLEWIGEINHS-GSTNSKPSLKSRVTISVDTSKNQFSLILSSVTAADTAVYYCAREGGVNVWGKGTTVTVSS
^^^^^^^^ ^^^^^^^^ ^^^^^^^^

Therapeutic : Lanadelumab

Best Alignment of Lanadelumab heavy chain to a sequence from OAS
EVQLLESGGGLVQPGGSLRLSCAASGFTFSHYIMMWVRQAPGKGLEWVSGIYSSGGITVYADSVKGRFTISRDNSKNTLYLQMNSLRAEDTAVYYCAYRRIGVP-----RRDEFDIWGQGTMVTVSS
||||||||||||||||||||||||||||||.|.|.||||||||||||||||..|||.|.|||||||||||||||||||||||||||||||||||||||||...|||.......|..||||||||||||||
EVQLLESGGGLVQPGGSLRLSCAASGFTFSSYAMSWVRQAPGKGLEWVSGISGSGGSTYYADSVKGRFTISRDNSKNTLYLQMNSLRAEDTAVYYCAKDSIGVITFGGVNRGSFDIWGQGTMVTVSS
^^^^^^^^ ^^^^^^^^ ^^^^^^^^^^^^^^^^^^^

Best Alignment of Lanadelumab light chain to a sequence from OAS
DIQMTQSPSTLSASVGDRVTITCRASQSISSWLAWYQQKPGKAPKLLIYKASTLESGVPSRFSGSGSGTEFTLTISSLQPDDFATYYCQQYNTYWTFGQGTKVEIK
||||||||||||||||||||||||||||||||||||||||||||||||||||||||||||||||||||||||||||||||||||||||||||||||||||||||||
DIQMTQSPSTLSASVGDRVTITCRASQSISSWLAWYQQKPGKAPKLLIYKASTLESGVPSRFSGSGSGTEFTLTISSLQPDDFATYYCQQYNTYWTFGQGTKVEIK
^^^^^^ ^^^ ^^^^^^^^

Best Alignment of Lanadelumab heavy chain CDRs to a sequence from OAS
EVQLLESGGGLVQPGGSLRLSCAASGFTFSHYIMMWVRQAPGKGLEWVSGIYSSGGITVYADSVKGRFTISRDNSKNTLYLQMNSLRAEDTAVYYCAYRRIGVPRRDEFDIWGQGTMVTVSS
................||||||||||||||.|.|.||||||||||||||.|..|||.|.||||||||||||||||||||||||||||||||||||||.||||...||.||||||||||||||
----------------SLRLSCAASGFTFSSYAMSWVRQAPGKGLEWVSVISGSGGTTYYADSVKGRFTISRDNSKNTLYLQMNSLRAEDTAVYYCAKRRIGGVVRDAFDIWGQGTMVTVSS
^^^^^^^^ ^^^^^^^^ ^^^^^^^^^^^^^^^

Best Alignment of Lanadelumab light CDRs chain to a sequence from OAS
DIQMTQSPSTLSASVGDRVTITCRASQSISSWLAWYQQKPGKAPKLLIYKASTLESGVPSRFSGSGSGTEFTLTISSLQPDDFATYYCQQYNTYWTFGQGTKVEIK
|||||||||||||||||||||.||||||||||||||||||||||||||||||.|||||||||||||||||||||||||||||.|||||||||||||||||||||||
DIQMTQSPSTLSASVGDRVTIACRASQSISSWLAWYQQKPGKAPKLLIYKASSLESGVPSRFSGSGSGTEFTLTISSLQPDDLATYYCQQYNTYWTFGQGTKVEIK
^^^^^^ ^^^ ^^^^^^^^

Best Alignment of Lanadelumab CDR-H3 chain to a sequence from OAS
EVQLLESGGGLVQPGGSLRLSCAASGFTFSHYIMMWVRQAPGKGLEWVSGIYSSGGITVYADSVKGRFTISRDNSKNTLYLQMNSLRAEDTAVYYCAYRRIGVPRRDEFDIWGQGTMVTVSS
.............|||||||||||||||||.|.|.|||||||||||||..|...|....|.||||||||||||||.||.||||||||||||||||||||||..|||||||.|.||||||||||||||
------------RPGGSLRLSCAASGFTFSSYWMSWVRQAPGKGLEWVANIKQDGSEKYYVDSVKGRFTISRDNAKNSLYLQMNSLRAEDTAVYYCARVRIGVPRWDAFDIWGQGTMVTVSS
^^^^^^^^ ^^^^^^^^ ^^^^^^^^^^^^^^^

Therapeutic : Landogrozumab

Best Alignment of Landogrozumab heavy chain to a sequence from OAS
EVQLVESGGGLVQPGGSLRLSCAASGLTFSRYPMSWVRQAPGKGLVWVSAITSSGGSTYYSDTVKGRFTISRDNAKNTLYLQMNSLRAEDTAVYYCARL---------------PDYWGQGTLVTVSS
||||||||||||||||||||||||||.|||.|.||||||||||||.|||||..||||||||.|.|||||||||||||.|||||||||||||||||||||||...............||||||||||||||
EVQLVESGGGLVQPGGSLRLSCAASGFTFSSYAMSWVRQAPGKGLEWVSAISGSGGSTYYADSVKGRFTISRDNSKNTLYLQMNSLRAEDTAVYYCARLEFFIAAAGTLVPPSVPDYWGQGTLVTVSS
^^^^^^^^ ^^^^^^^^ ^^^^^^^^^^^^^^^^^^^^^

Best Alignment of Landogrozumab light chain to a sequence from OAS
EIVLTQSPGTLSLSPGERATLSCRASSSVSSSYLHWYQQKPGQAPRLLIYSTSNLVAGIPDRFSGSGSGTDFTLTISRLEPEDFAVYYCQHHSGYHFTFGGGTKVEIK
||||||||||||||||||||||||||.|||||||.|||||||||||||||.|||..||||||||||||||||||||||||||||||||||...|...|||||||||||
EIVLTQSPGTLSLSPGERATLSCRASQSVSSSYLAWYQQKPGQAPRLLIYGTSNRAAGIPDRFSGSGSGTDFTLTISRLEPEDFAVYYCQQYDGSPLTFGGGTKVEIK
^^^^^^^ ^^^ ^^^^^^^^^

Best Alignment of Landogrozumab heavy chain CDRs to a sequence from OAS
EVQLVESGGGLVQPGGSLRLSCAASGLTFSRYPMSWVRQAPGKGLVWVSAITSSGGSTYYSDTVKGRFTISRDNAKNTLYLQMNSLRAEDTAVYYCARLPDYWGQGTLVTVSS
|||||||||.|||||.||.|||.|||.||..|.|.|.|..|||||.||..||||||||.|.|.||||.||||||||.||||||||||.||.|.||||||||||||||..|||||

```
EVQLVESGGDLVQPGRSLKLSCVASGFTFNNYWMTWIRHVPGKGLEWVASITSSGGSTNYRDSVKGRLTISRDNAKSTLYLQMNSLRSEDMATYYCARLPDYWGQGVMVTVSS
                      ^^^^^^^^               ^^^^^^^^                        ^^^^^^
```

Best Alignment of Landogrozumab light CDRs chain to a sequence from OAS

```
EIVLTQSPGTLSLSPGERATLSCRASSSVSSSYLHWYQQKPGQAPRLLIYSTSNLVAGIPDRFSGSGSGTDFTLTISRLEPEDFAVYYCQHHSGYHFTFGGGTKVEIK
|.||||||...|.|||...|..||||||||||||||||||.|..|.|.||||||||..|.|.|||||||||...||||..|.||.|.||||..||||||||.|||.|||
ENVLTQSPAIMSASPGDKVTMTCRASSSVSSSYLHWYQQKSGASPKLWIYSTSNLASGVPARFSGSGSGTSYSLTISSVEAEDAATYYCQQYSGYHFTFGSGTKLEIK
                       ^^^^^^^            ^^^                    ^^^^^^^^^
```

Best Alignment of Landogrozumab CDR-H3 chain to a sequence from OAS

```
EVQLVESGGGLVQPGGSLRLSCAASGLTFSRYPMSWVRQAPGKGLVWVSAITSSGGSTYYSDTVKGRFTISRDNAKNTLYLQMNSLRAEDTAVYYCARLPDYWGQGTLVTVSS
.............|.||..||.|||.||..|...|||||.|.||.|........|.|.|.....||.|..|.....|.|....|||.|||||||||||||||||||||||||
--------------GESLKISCKASGYTFTSYDINWVRQATGQGLEWMGWMNPNSGNTGYAQKFQGRVTMTRNTSISTAYMELSSLRSEDTAVYYCARLPDYWGQGTLVTVSS
                      ^^^^^^^^               ^^^^^^^^                        ^^^^^^
```

Therapeutic : Lebrikizumab

Best Alignment of Lebrikizumab heavy chain to a sequence from OAS

```
QVTLRESGPALVKPTQTLTLTCTVSGFSL--SAYSVNWIRQPPGKALEWLAMIWGDGKIVYNSALKSRLTISKDTSKNQVVLTMTNMDPVDTATYYCAGDGYY----PYAMDNWGQGSLVTVSS
||||||||||||||||||||||||.|||||..|...|.||||||||||||||.|..||...||..||.||||||||||||||||||||||||||||||||...||....|.|.|.||||..|||||
QVTLRESGPALVKPTQTLTLTCTFSGFSLSTSGMCVSWIRQPPGKALEWLARIDWDGDKYYNTSLKTRLTISKDTSKNQVVLTMTNMDPVDTATYYCARIHYYDSSGPRAFDIWGQGTMVTVSS
                      ^^^^^^^^^^               ^^^^^^^                        ^^^^^^^^^^^^^^^^
```

Best Alignment of Lebrikizumab light chain to a sequence from OAS

```
DIVMTQSPDSLSVSLGERATINCRASKSVDSYGNSFMHWYQQKPGQPPKLLIYLASNLESGVPDRFSGSGSGTDFTLTISSLQAEDVAVYYCQQNNEDPRTFGGGTKVEIK
|||.||||.||.||||.||||.||||.||||||||||||||||||||||||||.|||||||||.|||||||.||||||||...|.|||.|||||.|||||||||||||.|||
DIVLTQSPASLAVSLGQRATISCRASESVDSYGNSFMHWYQQKPGQPPKLLIYRASNLESGVPARFSGSGSRTDFTLTISPVEADDVATYYCQQSNEDPRTFGGGTKLEIK
                       ^^^^^^^^^^              ^^^                    ^^^^^^^^^
```

Best Alignment of Lebrikizumab heavy chain CDRs to a sequence from OAS

```
QVTLRESGPALVKPTQTLTLTCTVSGFSLSAYSVNWIRQPPGKALEWLAMIWGDGKIVYNSALKSRLTISKDTSKNQVVLTMTNMDPVDTATYYCAGDGYYPYAMDNWGQGSLVTVSS
||.|.||||.||.|.|.|..|||||||||..|.|.|.||||||.||||..|||||...|.|||.|||.||||.||.||.|........||||||||.|||||||||.||||..|||||
QVQLKESGPGLVAPSQSLSITCTVSGFSLTSYGVSWVRQPPGKGLEWLGVIWGDGSTNYHSALISRLSISKDNSKSQVFLKLNSLQTDDTATYYCAKDGYYPYAMDYWGQGTSVTVSS
                      ^^^^^^^^               ^^^^^^^                        ^^^^^^^^^^^^
```

Best Alignment of Lebrikizumab light CDRs chain to a sequence from OAS

```
DIVMTQSPDSLSVSLGERATINCRASKSVDSYGNSFMHWYQQKPGQPPKLLIYLASNLESGVPDRFSGSGSGTDFTLTISSLQAEDVAVYYCQQNNEDPRTFGGGTKVEIK
.||.||||.||.||||.||||.||||.||||||||||||||||||||||||||||||||||||.|||||||.|||||||....|.|.|.||||||||||||||||||||.|||
NIVPTQSPASLAVSLGQRATISCRASESVDSYGNSFMHWYQQKPGQPPKLLIYLASNLESGVPARFSGSGSRTDFTLTIDPVEADDAATYYCQQNNEDPRTFGGGTKLEIK
                       ^^^^^^^^^^              ^^^                    ^^^^^^^^^
```

Best Alignment of Lebrikizumab CDR-H3 chain to a sequence from OAS

```
QVTLRESGPALVKPTQTLTLTCTVSGFSLSAYSVNWIRQPPGKALEWLAMIWGD---GKIVYNSALKSRLTISKDTSKNQVVLTMTNMDPVDTATYYCAGDGYYPYAMDNWGQGSLVTVSS
.|.|.|||..||.|...|.|.|..|||....|...|.|||||||||||..|..........|....|.|...|.|||.......|......|.|.||.||||||||||||.||||..|||||
EVELVESGGDLVQPGGSLRLSCATSGFTFTDYYMSWVRQPPGKALEWLGVIRNKANGYTTEYSASVKGRFPCSRDTSQSILYRQMDTLRAEDGASYYRAGDGYYPYAMDYWGQGASVTVSS
                      ^^^^^^^^               ^^^^^^^^^^                        ^^^^^^^^^^^^
```

Therapeutic : Lenzilumab

Best Alignment of Lenzilumab heavy chain to a sequence from OAS

```
QVQLVQSGAEVKKPGASVKVSCKASGYSFTNYYIHWVRQAPGQRLEWMGWINAGNGNTKYSQKFQGRVTITRDTSASTAYMELSSLRSEDTAVYYCVRRQRF--------PYYFDYWGQGTLVTVSS
|||||||||||||||||||||||||||.||.||.||||||||||||||||||||||||||||||||||||||||||||||||||||||||||||||||||.|.||..........|||||||||||||||
QVQLVQSGAEVKKPGASVKVSCKASGYTFTGYYMHWVRQAPGQRLEWMGWINAGNGNTKYSQKFQGRVTITRDTSASTAYMELSSLRSEDTAVYYCARDQRKGTAAGSPIAYYFDYWGQGTLVTVSS
                      ^^^^^^^^               ^^^^^^^^                        ^^^^^^^^^^^^^^^^^^^
```

Best Alignment of Lenzilumab light chain to a sequence from OAS

```
EIVLTQSPATLSVSPGERATLSCRASQSVGTNVAWYQQKPGQAPRVLIYSTSSRATGITDRFSGSGSGTDFTLTISRLEPEDFAVYYCQQFNKSPLTFGGGTKVEIK
||||||||||||||||||||||||||||||.|.||||||||||||.|||..|||||||.||||||||||||||||||||||||||||||||...|||||||||||||
EIVLTQSPATLSVSPGERATLSCRASQSVGNNLAWYQQKPGQAPRLLIYGASSRATGIPDRFSGSGSGTDFTLTISRLEPEDFAVYYCQQYSSSPLTFGGGTKVEIK
                       ^^^^^^             ^^^                    ^^^^^^^^^
```

Best Alignment of Lenzilumab heavy chain CDRs to a sequence from OAS

```
QVQLVQSGAEVKKPGASVKVSCKASGYSFTNYYIHWVRQAPGQRLEWMGWINAGNGNTKYSQKFQGRVTITRDTSASTAYMELSSLRSEDTAVYYCVRRQRFPYYFDYWGQGTLVTVSS
................|||||||||||.||||..|||||||||.|||||||||||||||||||||||||||||||||.||||||||||||||||||||.|...|||||||||||||||||
----------------SVKVSCKASGYTFTNYAMHWVRQAPGQWLEWMGWINAGNGNTKYSQKFQGRVTITRDTSARTAYMELSSLRSEDTAVYYCARGFDFPYYFDYWGQGTLVTVSS
                      ^^^^^^^^               ^^^^^^^^                        ^^^^^^^^^^^^
```

Best Alignment of Lenzilumab light CDRs chain to a sequence from OAS

```
EIVLTQSPATLSVSPGERATLSCRASQSVGTNVAWYQQKPGQAPRVLIYSTSSRATGITDRFSGSGSGTDFTLTISRLEPEDFAVYYCQQFNKSPLTFGGGTKVEIK
||||||||||||..|||.||||||||||.||.||||||||||||.|.||.|.|||||..|||||||||.||||||.|..||.||||||||||.|||||||||||||
EIVLTQSPATLSVAEGERTTLSCRASQSVNTNLAWYQQKPGQAPRLLLYSASTRATGIPARFSGSGSGTEFTLTISSLQSEDVAVYYCQQFNKWPLTFGGGTKVEIK
                       ^^^^^^             ^^^                    ^^^^^^^^^
```

Best Alignment of Lenzilumab CDR-H3 chain to a sequence from OAS

```
QVQLVQSGAEVKKPGASVKVSCKASGYSFTNYYIHWVRQAPGQRLEWMGWINAGNGNTKYSQKFQGRVTITRDTSASTAYMELSSLRSEDTAVYYCVRRQRFPYYFDYWGQGTLVTVSS
...................|..||.|.......|.||.||..|||.|.|....|...|.......|||..|||........||....||||||||||.|.|||||||||||||||||
----------------TLSLTCTVSGVSISDHSWGWIRQPPGKGLEWIGYISYS-GRSNYNPPLKSQVTISLDTSKNQVSLKVSSVTAADTAVYYCVRRFRDPYYFDYWGQGTLVTVSS
                      ^^^^^^^^               ^^^^^^^^                        ^^^^^^^^^^^^
```

Therapeutic : Lifastuzumab

Best Alignment of Lifastuzumab heavy chain to a sequence from OAS

```
EVQLVESGGGLVQPGGSLRLSCAASGFSFSDFAMSWVRQAPGKGLEWVATIGRVAFHTYYPDSMKGRFTISRDNSKNTLYLQMNSLRAEDTAVYYCARHRGFD---VGHFDFWGQGTLVTVSS
||||||||||||||||||||||||||||.||..|||||||||||||||||..|......|||.||.||||||||||||||||||||||||||||||||||||.||.|....|.||.|||||||||||
EVQLVESGGGLVQPGGSLRLSCAASGFTFSSYAMSWVRQAPGKGLEWVSAISGSGGSTYYADSVKGRFTISRDNSKNTLYLQMNSLRAEDTAVYYCARDRGPDYGDYGWFDYWGQGTLVTVSS
                      ^^^^^^^^               ^^^^^^^^                        ^^^^^^^^^^^^^^^^
```

Best Alignment of Lifastuzumab light chain to a sequence from OAS

```
DIQMTQSPSSLSASVGDRVTITCRSSETLVHSSGNTYLEWYQQKPGKAPKLLIYRVSNRFSGVPSRFSGSGSGTDFTLTISSLQPEDFATYYCFQGSFNPLTFGQGTKVEIK
||||||||||||||||||||||||.|.|......||||.|||||||||||||||..|...||||||||||||||||||||||||||||||||||||.|.|..|.|||||||||||
DIQMTQSPSSLSASVGDRVTITCRASQTI-----NTYLNWYQQKPGKAPKLLIYAASSLQSGVPSRFSGSGSGTDFTLTISSLQPEDFATYYCQQSSSTPWTFGQGTKVEIK
                       ^^^^^^^^^^^              ^^^                    ^^^^^^^^^
```

Best Alignment of Lifastuzumab heavy chain CDRs to a sequence from OAS

```
EVQLVESGGGLVQPGGSLRLSCAASGFSFSDFAMSWVRQAPGKGLEWVATIGRVAFHTYYPDSMKGRFTISRDNSKNTLYLQMNSLRAEDTAVYYCARHRGFDVGHFDFWGQGTLVTVSS
................||||||||||.||||.|.||||||||||||...|...|.||.|.||..||||.||||..|.|||||||||.||||||||||.||...|||||||||.||.|||
----------------SLRLSCAASGFTFSDFYMTWVRQAPGKGLEWISDISDTAAHTNYADSVRGRFTVSRDNAHNSLYLQMNSLRGEDTAVYYCARGRGDGFGHFDFWGQGRLVVVSS
                      ^^^^^^^^               ^^^^^^^^                        ^^^^^^^^^^^^^
```

Best Alignment of Lifastuzumab light CDRs chain to a sequence from OAS

```
DIQMTQSPSSLSASVGDRVTITCRSSETLVHSSGNTYLEWYQQKPGKAPKLLIYRVSNRFSGVPSRFSGSGSGTDFTLTISSLQPEDFATYYCFQGSFNPLTFGQGTKVEIK
.....|.|.||..|.||...|.||||..||||||||||.||.||||..||||||.|||||||||.||||||||||||||.||....||...|||||||..|||||.|||.|.|
-----QTPLSLPVSLGDQAAISCRSSLSLVHSSGNTYLHWYLQKPGQSPKLLIYKVSNRFSGVPDRFSGSGSGTDFTLKISRVEAEDLGVYYCFQGSHVPLTFGAGTKLELK
                       ^^^^^^^^^^^              ^^^                    ^^^^^^^^^
```

Best Alignment of Lifastuzumab CDR-H3 chain to a sequence from OAS

```
EVQLVESGGGLVQPGGSLRLSCAASGFSF-SDFAMSWVRQAPGKGLEWVATIGRVAFHTYYPDSMKGRFTISRDNSKNTLYLQMNSLRAEDTAVYYCARHRGFDVGHFDFWGQGTLVTVSS
.................|.|.|..||.|.......|||||||||||||.........|.|..|.|.|.|.|.|.|.|.|...|...||.|.||.||.||||||..||.||||||||||||||
```

--------------SQTLSLTCTVSGVSVTTTNWWSWVRQAPGKGLEWVGEVYQD-GSTNYNPSLKSRVTVSLDKSNNKFSLKLTSLTAADTGVYFCARHRGATVGPFDFWGQGTLVTVSS
^^^^^^^^^ ^^^^^^^^ ^^^^^^^^^^^^^

Therapeutic : Ligelizumab

Best Alignment of Ligelizumab heavy chain to a sequence from OAS
QVQLVQSGAEVMKPGSSVKVSCKASGYTFSWYWLEWVRQAPGHGLEWMGEIDPGTFTTNYNEKFKARVTFTADTSTSTAYMELSSLRSEDTAVYYCARFSHFSGSNYDFDYWGQGTLVTVSS
|||||||||||.||||||||||||||.|||.|.|.|||||||.||||||.|.|...|.||..||..|||.|||.||||||||||||||||||||||||....|||.|.|||||||||||||||
QVQLVQSGAEVKKPGSSVKVSCKASGGTFSSYALSWVRQAPGQGLEWMGGIIPIFGTANYAQKFQGRVTITADESTSTAYMELSSLRSEDTAVYYCARVGIYSGSYYHYFDYWGQGTLVTVSS
^^^^^^^^ ^^^^^^^^ ^^^^^^^^^^^^^^^^

Best Alignment of Ligelizumab light chain to a sequence from OAS
EIVMTQSPATLSVSPGERATLSCRASQSIGTNIHWYQQKPGQAPRLLIYYASESISGIPARFSGSGSGTEFTLTISSLQSEDFAVYYCQQSWSWPTTFGGGTKVEIK
|||||||||||||||||||||||||||||..|..|||||||||||||||.||....||||||||||||||||||||||||||||||||||..|||.|||||||||||
EIVMTQSPATLSVSPGERATLSCRASQSISSNLAWYQQKPGQAPRLLIYGASTRATGIPARFSGSGSGTEFTLTISSLQSEDFAVYYCQQYNSWPLTFGGGTKVEIK
^^^^^^ ^^^ ^^^^^^^^^

Best Alignment of Ligelizumab heavy chain CDRs to a sequence from OAS
QVQLVQSGAEVMKPGSSVKVSCKASGYTFSWYWLEWVRQAPGHGLEWMGEIDPGTFTTNYNEKFKARVTFTADTSTSTAYMELSSLRSEDTAVYYCARFSHFSGSNYDFDYWGQGTLVTVSS
.|||.|.|||...||||||.|||||||||..|||.||.|.|..|.||.|.|||....|.||.|||...|.|.|.|.|||||.||||.|||.||||||||||..||.||||||||||..||||
-VQLQQPGAELVRPGSSVKLSCKASGYTFTSYWLHWVKQRPIQGHEWIGNIDPSDSETHYNQKFKDKATLTVDKSSSTAYMQLSSLTSEDSAVYYCARFSHYYGSSYDFDYWGQGTTLTVSS
^^^^^^^^ ^^^^^^^^ ^^^^^^^^^^^^^^^^

Best Alignment of Ligelizumab light CDRs chain to a sequence from OAS
EIVMTQSPATLSVSPGERATLSCRASQSIGTNIHWYQQKPGQAPRLLIYYASESISGIPARFSGSGSGTEFTLTISSLQSEDFAVYYCQQSWSWPTTFGGGTKVEIK
.|..|||||.||||||||...|||||||||||||||||.....|||||.|||||.||||.|||||||||.|||.|.|..|||.|.||||||.|||.|||||||.|||
DILLTQSPAILSVSPGERVSFSCRASQSIGTNIHWYQQRTNGSPRLLIKYASESSSGIPSRFSGSGSGTDFTLSINSVESEDIADYYCQQSNSWPYTFGGGTKLEIK
^^^^^^ ^^^ ^^^^^^^^^

Best Alignment of Ligelizumab CDR-H3 chain to a sequence from OAS
QVQLVQSGAEVMKPGSSVKVSCKASGYTFSWYWLEWVRQAPGHGLEWMGEIDPGTFTTNYNEKFKARVTFTADTSTSTAYMELSSLRSEDTAVYYCARFSHFSGSNYDFDYWGQGTLVTVSS
.|||.|.|||...||||||.|||||||||..|||.||.|.|..|.||.|.|||....|.||.|||...|.|.|.|.|||||.||||.|||.||||||||||..||.||||||||||..||||
-VQLQQPGAELVRPGSSVKLSCKASGYTFTSYWLHWVKQRPIQGHEWIGNIDPSDSETHYNQKFKDKATLTVDKSSSTAYMQLSSLTSEDSAVYYCARFSHYYGSSYDFDYWGQGTTLTVSS
^^^^^^^^ ^^^^^^^^ ^^^^^^^^^^^^^^^^

Therapeutic : Lintuzumab

Best Alignment of Lintuzumab heavy chain to a sequence from OAS
QVQLVQSGAEVKKPGSSVKVSCKASGYTFTDYNMHWVRQAPGQGLEWIGYIYPYNGGTGYNQKFKSKATITADESTNTAYMELSSLRSEDTAVYYCARGRP--AMDYWGQGTLVTVSS
|||||||||||||||||||||||||||||.||.||||||||||||||.|.|.|.|.||||.|.|||....||||||||.|||||||||||||||||||||.|...|.|||||||||||||
QVQLVQSGAEVKKPGSSVKVSCKASGYTFIDYYMHWVRQAPGQGLEWMGWINPNNGGTNYAQKFQGRVTITADESTSTAYMELSSLRSEDTAVYYCARLRSTSALDYWGQGTLVTVSS
^^^^^^^^ ^^^^^^^^ ^^^^^^^^^^^

Best Alignment of Lintuzumab light chain to a sequence from OAS
DIQMTQSPSSLSASVGDRVTITCRASESVDNYGISFMNWFQQKPGKAPKLLIYAASNQGSGVPSRFSGSGSGTDFTLTISSLQPDDFATYYCQQSKEVPWTFGQGTKVEIK
||||||||||||||||||||||||||.||.....||.||.||||||||||||||||...|||||||||||||||||||||||||||||||.||||...|||||||||||||
DIQMTQSPSSLSASVGDRVTITCRASQSV----SSFLNWYQQKPGKAPKLLIYAASRLQSGVPSRFSGSGSGTDFTLTISSLQPDDFATYSCQQSLVLPWTFGQGTKVEIK
^^^^^^^^^^ ^^^ ^^^^^^^^^

Best Alignment of Lintuzumab heavy chain CDRs to a sequence from OAS
QVQLVQSGAEVKKPGSSVKVSCKASGYTFTDYNMHWVRQAPGQGLEWIGYIYPYNGGTGYNQKFKSKATITADESTNTAYMELSSLRSEDTAVYYCARGRPAMDYWGQGTLVTVSS
.|||.|||.|..|||.|||.|||||||||||||||||.|..|..|||||||||||||||||||||||||.|.|.|.|..||||||.||.|||.|||||||||.|||||||||.|||||
-VQLQQSGPELVKPGASVKISCKASGYTFTDYNMHWVKQSHGKSLEWIGYIYPYNGGTGYNQKFKSKATLTVDNSSSTAYMELRSLTSEDSAVYYCARGRDAMDYWGQGTSVTVSS
^^^^^^^^ ^^^^^^^^ ^^^^^^^^^

Best Alignment of Lintuzumab light CDRs chain to a sequence from OAS
DIQMTQSPSSLSASVGDRVTITCRASESVDNYGISFMNWFQQKPGKAPKLLIYAASNQGSGVPSRFSGSGSGTDFTLTISSLQPDDFATYYCQQSKEVPWTFGQGTKVEIK
||..||||.||..|.|.|.||.|||||||||||||||||||||||..||||||||||||||||.|||||||||||.|.|.|.....||.|.|.||||||||||||.|||....
DIVLTQSPASLAVSLGQRATISCRASESVDNYGISFMNWFQQKPGQPPKLLIYAASNQGSGVPARFSGSGSGTDFSLNIHPMEEDDTAMYFCQQSKEVPWTFGGGTKLKSN
^^^^^^^^^^ ^^^ ^^^^^^^^^

Best Alignment of Lintuzumab CDR-H3 chain to a sequence from OAS
QVQLVQSGAEVKKPGSSVKVSCKASGYTFTDYNMHWVRQAPGQGLEWIGYIYPYNGGTGYNQKFKSKATITADESTNTAYMELSSLRSEDTAVYYCARGRPAMDYWGQGTLVTVSS
.|||.|.|||...||||||.||||||||||.|.|.||.|.|||||||||.|||....|.||||.|.|||.|.|.|.|..||||.||||.|||.||||||||||||||.||||.|||||
-VQLQQPGAELVRPGSSVKLSCKASGYTFTSYWMDWVKQRPGQGLEWIGNIYPSDSETHYNQKVKDKATLTVDKSSSTAYMQLSSLTSEDSAVYYCARGRPAMDYRGQGTSVTVSS
^^^^^^^^ ^^^^^^^^ ^^^^^^^^^

Therapeutic : Lirilumab

Best Alignment of Lirilumab heavy chain to a sequence from OAS
QVQLVQSGAEVKKPGSSVKVSCKASGGTFSFYAISWVRQAPGQGLEWMGGFIPIFGAANYAQKFQGRVTITADESTSTAYMELSSLRSDDTAVYYCARIPSGSY-----YYDYDMDVWGQGTTVTVSS
||||||||||||||||||||||||||||||.|||||||||||||||||||.|||||.|||||||||||||||||||||||||||||||.||||||||||.||||.....||.|.||||||||||||||
QVQLVQSGAEVKKPGSSVKVSCKASGGTFSSYAISWVRQAPGQGLEWMGGIIPIFGTANYAQKFQGRVTITADESTSTAYMELSSLRSEDTAVYYCARIRSGSYYNENYYYYGMDVWGQGTTVTVSS
^^^^^^^^ ^^^^^^^^ ^^^^^^^^^^^^^^^^^^^^^

Best Alignment of Lirilumab light chain to a sequence from OAS
EIVLTQSPVTLSLSPGERATLSCRASQSVSSYLAWYQQKPGQAPRLLIYDASNRATGIPARFSGSGSGTDFTLTISSLEPEDFAVYYCQQRSNWMYTFGQGTKLEIK
||||||||.||||||||||||||||||||||||||||||||||||||||||||||||||||||||||||||||||||||||||||||||||||||||||||||||||
EIVLTQSPATLSLSPGERATLSCRASQSVSSYLAWYQQKPGQAPRLLIYDASNRATGIPARFSGSGSGTDFTLTISSLEPEDFAVYYCQQRSNWMYTFGQGTKLEIK
^^^^^^ ^^^ ^^^^^^^^^

Best Alignment of Lirilumab heavy chain CDRs to a sequence from OAS
QVQLVQSGAEVKKPGSSVKVSCKASGGTFSFYAISWVRQAPGQGLEWMGGFIPIFGAANYAQKFQGRVTITADESTSTAYMELSSLRSDDTAVYYCARIPSGSYYYDYDMDVWGQGTTVTVSS
..............|.||||||||||||||.|||||||||||||||||||.||||||||||||||||||||||||||||||||.||||.|||||||||..||||||.|||||||.||||||||
--------------GASVKVSCKASGGTFSSYAISWVRQAPGQGLEWMGGIIPIFGAANYAQKFQGRVTITADESTSTAYMELRSLRSEDTAVYYCARAHSGSYYYYYDMDVWGKGTTVTVSS
^^^^^^^^ ^^^^^^^^ ^^^^^^^^^^^^^^^^

Best Alignment of Lirilumab light CDRs chain to a sequence from OAS
EIVLTQSPVTLSLSPGERATLSCRASQSVSSYLAWYQQKPGQAPRLLIYDASNRATGIPARFSGSGSGTDFTLTISSLEPEDFAVYYCQQRSNWMYTFGQGTKLEIK
||||||||.||||||||||||||||||||||||||||||||||||||||||||||||||||||||||||||||||||||||||||||||||||||||||||||||||
EIVLTQSPATLSLSPGERATLSCRASQSVSSYLAWYQQKPGQAPRLLIYDASNRATGIPARFSGSGSGTDFTLTISSLEPEDFAVYYCQQRSNWMYTFGQGTKLEIK
^^^^^^ ^^^ ^^^^^^^^^

Best Alignment of Lirilumab CDR-H3 chain to a sequence from OAS
QVQLVQSGAEVKKPGSSVKVSCKASGGTF--SFYAISWVRQAPGQGLEWMGGFIPIFGAANYAQKFQGRVTITADESTSTAYMELSSLRSDDTAVYYCARIPSGSYYYDYDMDVWGQGTTVTVSS
.....................|..|||......|...|.||.||.||||.|..........|......||||..|.|.......|.|....|||||||||..||||||||||||||.||.|||||
----------------TLSLTCTVSGGSISSIGYFWTWIRQHPGKGLEWIGYKYCS-ESTDYNPSLKSRVTISVDTSENQFSLKLRSVTAADTAVYYCARGYSGSYYYDYDMDVWGKGTSVTVSS
^^^^^^^^^^ ^^^^^^^^ ^^^^^^^^^^^^^^^^

Therapeutic : Lodelcizumab

Best Alignment of Lodelcizumab heavy chain to a sequence from OAS
QVQLVQSGAEVKKPGASVKVSCKASGYTFSTMYMSWVRQAPGQGLEWMGRIDPANEHTNYAQKFQGRVTMTRDTSISTAYMELSRLTSDDTAVYYCARSYYY------YNMDYWGQGTLVTVSS
|||||||||||||||||||||||||||||...|..||||||||||||||||.|....|||||||||||||||||||||||||||||.|||||||||||.|||......|..|||||||||||||
QVQLVQSGAEVKKPGASVKVSCKASGYTFTGYYIQWVRQAPGQGLEWMGRINPNSGGTNYAQKFQGRVTMTRDTSISTAYMELSRLRSDDTAVYYCARVYYYDSSGYYYYFDYWGQGTLVTVSS
^^^^^^^^ ^^^^^^^^ ^^^^^^^^^^^^^^^^^

Best Alignment of Lodelcizumab light chain to a sequence from OAS
QIVLTQSPATLSVSPGERATLSCRASQSV--SYMHWYQQKPGQAPRLLIYGVFRRATGIPDRFSGSGSGTDFTLTIGRLEPEDFAVYYCLQWSS-DPPTFGQGTKLEIK
.||||||||||||||||||||||||||||..||..|||||||||||||||||..||||||||||||||||||||||.||||||||||||.|..|..|.|||||||||||
EIVLTQSPATLSVSPGERATLSCRASQSVSSSYLAWYQQKPGQAPRLLIYGVSTRATGIPDRFSGSGSGTDFTLTISRLEPEDFAVYYCQQYGSSPPYTFGQGTKLEIK
^^^^^^^ ^^^ ^^^^^^^^^^

Best Alignment of Lodelcizumab heavy chain CDRs to a sequence from OAS
QVQLVQSGAEVKKPGASVKVSCKASGYTFSTMYMSWVRQAPGQGLEWMGRIDPANEHTNYAQKFQGRVTMTRDTSISTAYMELSRLTSDDTAVYYCARSYYYYNMDYWGQGTLVTVSS
.|||.|.|||..|||||||.|||||||||....|.||.|.|.|||||.|||||||..|.|..||||..|.|.|||..|||..||.|||.||||||||||.|||.||||||||.|||||
-VQLQQPGAELVKPGASVKLSCKASGYTFTSYWMHWVKQRPEQGLEWIGRIDPANGNTKYDPKFQGKATITADTSSNTAYLQLSSLTSEDTAVYYCARSGYYYAMDYWGQGTSVTVSS
^^^^^^^^ ^^^^^^^^ ^^^^^^^^^^^

Best Alignment of Lodelcizumab light CDRs chain to a sequence from OAS
QIVLTQSPATLSVSPGERATLSCRASQSVSYMHWYQQKPGQAPRLLIYGVFRRATGIPDRFSGSGSGTDFTLTIGRLEPEDFAVYYCLQWSSDPPTFGQGTKLEIK
|||||||||..|.||||..|..|.||.|||||.|||||||..|||||||....|.|.|.|||||||||...|||.|.|.||.|.|||.||||.|||||.|||||||
QIVLTQSPAIMSASPGEKVTMTCSASSSVSYMYWYQQKPGSSPRLLIYGTSNLASGVPVRFSGSGSGTSYSLTISRMEAEDAATYYCQQWSSYPPTFGGGTKLEIK
^^^^^ ^^^ ^^^^^^^^^

Best Alignment of Lodelcizumab CDR-H3 chain to a sequence from OAS
QVQLVQSGAEVKKPGASVKVSCKASGYTF-STMYMSWVRQAPGQGLEWMGRIDPANEHTNYAQKFQGRVTMTRDTSISTAYMELSRLTSDDTAVYYCARSYYYYNMDYWGQGTLVTVSS
.|||..||....||..|....|...||...|.....|.||.||..|||||.|......|.|......|...|||.|.....|.|...|.|..|||.||||||||||.||||||||.|||||
EVQLQESGPGLVKPSQSLSLTCTVTGYSITSDYAWNWIRQFPGNKLEWMGYISYS-GNTSYNPSLKSRISITRDQSKNQFFMQLNSVTTEDTATYYCARSYYYYAMDYWGQGTSVTVSS
^^^^^^^^^ ^^^^^^^^ ^^^^^^^^^^^

Therapeutic : Lorvatuzumab

Best Alignment of Lorvatuzumab heavy chain to a sequence from OAS
QVQLVESGGGVVQPGRSLRLSCAASGFTFSSFGMHWVRQAPGKGLEWVAYISSGSFTIYYADSVKGRFTISRDNSKNTLYLQMNSLRAEDTAVYYCARMRKG--YAMDYWGQGTLVTVSS
.||||||||||||||||||||||||||||||.||||||||||||||||.||||.|.||||||||||||||||||.||.||||||||||||||||||||.|.|...|.||||||||||||
-VQLVESGGGVVQPGRSLRLSCAASGFTFSSYGMHWVRQAPGKGLEWVSYISSSSSTIYYADSVKGRFTISRDNAKNSLYLQMNSLRAEDTAVYYCARDRVGAIGAFDYWGQGTLVTVSS
^^^^^^^^ ^^^^^^^^ ^^^^^^^^^^^^^

Best Alignment of Lorvatuzumab light chain to a sequence from OAS
DVVMTQSPLSLPVTLGQPASISCRSSQIIIHSDGNTYLEWFQQRPGQSPRRLIYKVSNRFSGVPDRFSGSGSGTDFTLKISRVEAEDVGVYYCFQGSH-VPHTFGQGTKVEIK
|||||||||||||||||||||||||||...||||||||.||||||||||||||||||||.|||||||||||||||||||||||||||||||||.||||..|.|||||||||||
DVVMTQSPLSLPVTLGQPASISCRSSQSLVHSDGNTYLNWFQQRPGQSPRRLIYKVSNRDSGVPDRFSGSGSGTDFTLKISRVEAEDVGVYYCMQGSHFPPWTFGQGTKVEIK
^^^^^^^^^^^ ^^^ ^^^^^^^^^^

Best Alignment of Lorvatuzumab heavy chain CDRs to a sequence from OAS
QVQLVESGGGVVQPGRSLRLSCAASGFTFSSFGMHWVRQAPGKGLEWVAYISSGSFTIYYADSVKGRFTISRDNSKNTLYLQMNSLRAEDTAVYYCARMRKGYAMDYWGQGTLVTVSS
.|||||||||.||||.|..||||||||||||||||||||||.|||||||||||||.||||||.|||||||||||.||||.||..|||.||||.|.|||.|.|||||||||||..||||
EVQLVESGGGLVQPGGSRKLSCAASGFTFSSFGMHWVRQAPEKGLEWVAYISSGSSTIYYADTVKGRFTISRDNPKNTLFLQKTSLRSEDTAMYDCARWRYGYAMDYWGQGTSLTVSS
^^^^^^^^ ^^^^^^^^ ^^^^^^^^^^^

Best Alignment of Lorvatuzumab light CDRs chain to a sequence from OAS
DVVMTQSPLSLPVTLGQPASISCRSSQIIIHSDGNTYLEWFQQRPGQSPRRLIYKVSNRFSGVPDRFSGSGSGTDFTLKISRVEAEDVGVYYCFQGSHVPHTFGQGTKVEIK
||||||...||||..|...|..|||.|.||||.|||||||..|.||..|..|||||||||.||||||||||||||||||||||||||.||||||||||||.|||.|||.|||
DVVMTQTAHSLPVSPGDQVSMPCRSGQSIIHSNGNTYLEWYLQKPGEYPKLLIYKVSNRFPGVPDRFSGSGSGTDFTLKISRVEAEDLGVYYCFQGSHVPWTFGRGTKLEIK
^^^^^^^^^^^ ^^^ ^^^^^^^^^

Best Alignment of Lorvatuzumab CDR-H3 chain to a sequence from OAS
QVQLVESGGGVVQPGRSLRLSCAASGFTF--SSFGMHWVRQAPGKGLEWVAYISSGSFTIYYADSVKGRFTISRDNSKNTLYLQMNSLRAEDTAVYYCARMRKGYAMDYWGQGTLVTVSS
||.|.|||.|..||...|.|.|..|||....|..|..|.||..||||||.|.|........|....|.|.|||.|.|.|...|...|....|||.|||||||||||||||||||.|||||
QVTLKESGPGILQPSQTLSLTCSFSGFSLSTSGMGVGWIRQPSGKGLEWLAHIWWD-DDKRYNPALKSRLTISKDTSSNQVFLKIASVDTADTATYYCARMRKGYAMDYWGQGTSVTVSS
^^^^^^^^^^ ^^^^^^^^ ^^^^^^^^^^^

Therapeutic : Lumiliximab

Best Alignment of Lumiliximab heavy chain to a sequence from OAS
EVQLVESGGGLAKPGGSLRLSCAASGFRFTFNNYYMDWVRQAPGQGLEWVSRISSSGDPTWYADSVKGRFTISRENANNTLFLQMNSLRAEDTAVYYCASLTT----GSDSWGQGVLVTVSS
.||||||||||.|||||||||||||||.|....|||.|.|||||.||||.|.|||||..|.|||||||||||||.||.|.|||||||||||||||||||||||....|.|.||||.||||||
QVQLVESGGGLVKPGGSLRLSCAASGFSF--SDYYMSWIRQAPGKGLEWLSYISSSGTYTTYADSVKGRFTISRDNAKNSLFLQMNSLRAEDTAVYYCASLTTVVTPGLDYWGQGTLVTVSS
^^^^^^^^^^ ^^^^^^^^ ^^^^^^^^^^^^^

Best Alignment of Lumiliximab light chain to a sequence from OAS
DIQMTQSPSSLSASVGDRVTITCRASQDIRYYLNWYQQKPGKAPKLLIYVASSLQSGVPSRFSGSGSGTEFTLTVSSLQPEDFATYYCLQVYSTPRTFGQGTKVEIK
|||||||||||||||||||||||||||.|..||||||||||||||||||.||||||||||||||||||||||||.|||||||||||||||.||||||||||||||||
DIQMTQSPSSLSASVGDRVTITCRASQSISSYLNWYQQKPGKAPKLLIYAASSLQSGVPSRFSGSGSGTEFTLTISSLQPEDFATYYCLQSYSTPRTFGQGTKVEIK
^^^^^^ ^^^ ^^^^^^^^^

Best Alignment of Lumiliximab heavy chain CDRs to a sequence from OAS
EVQLVESGGGLAKPGGSLRLSCAASGFRFTFNNYYMDWVRQAPGQGLEWVSRISSSGDPTWYADSVKGRFTISRENANNTLFLQMNSLRAEDTAVYYCASLTTGSDSWGQGVLVTVSS
...............|||.||||.|||.|....||..|.|||||.||||||.|||||....|||||||||||||.||.|.|.|||||||||||||||||.||||.|.||||.||||||
--------------WGSLCLSCAVSGFSFSSSGYYLGWIRQAPGKGLEWVSYISSSGSTIYYADSVKGRFTISRDNAKNSLYLQMNSLRAEDTAVYYCARLTTGFDYWGQGTLVTVSS
^^^^^^^^^^ ^^^^^^^^ ^^^^^^^^^

Best Alignment of Lumiliximab light CDRs chain to a sequence from OAS
DIQMTQSPSSLSASVGDRVTITCRASQDIRYYLNWYQQKPGKAPKLLIYVASSLQSGVPSRFSGSGSGTEFTLTVSSLQPEDFATYYCLQVYSTPRTFGQGTKVEIK
||||||||||||||.||||||||||||.||.||||||.||||||||||.|||.||||||||||||||||.||||.|||||||||||||||.||||||||||||||||
DIQMTQSPSSLSASEGDRVTITCRASQSIRNYLNWYQEKPGKAPKLLINVASILQSGVPSRFSGSGSGTDFTLTISSLQPEDFATYYCLQSYSTPRTFGQGTKVEIK
^^^^^^ ^^^ ^^^^^^^^^

Best Alignment of Lumiliximab CDR-H3 chain to a sequence from OAS
EVQLVESGGGLAKPGGSLRLSCAASGFRFTFNNYYMDWVRQAPGQGLEWVSRISSSGDPTWYADSVKGRFTISRENANNTLFLQMNSLRAEDTAVYYCASLTTGSDSWGQGVLVTVSS
||||.|||..|.|||.|...||.|||..|....|.|.||.|..|..|||...|......|.|....||..|........|...|.|||..||.||||||||||||.|||||...||||
EVQLQESGPELVKPGASVKISCKASGYSF--TDYNMNWVKQSNGKSLEWIGVINPNYGTTSYNQKFKGKATLTVDQSSSTAYMQLNSLTSEDSAVYYCASLTTGSSSWGQGTTHTVSS
^^^^^^^^^^ ^^^^^^^^ ^^^^^^^^^

Therapeutic : Margetuximab

Best Alignment of Margetuximab heavy chain to a sequence from OAS
QVQLQQSGPELVKPGASLKLSCTASGFNIKDTYIHWVKQRPEQGLEWIGRIYPTNGYTRYDPKFQDKATITADTSSNTAYLQVSRLTSEDTAVYYCSRWGGDG--FYAMDYWGQGASVTVSS
.|||||||.||||||||.|||||||||||||||.|||||||||||||||||.|.||.|.||||||.||||||||||||||||.|.|||||||||||.|.||||..||||||||||.||||||
EVQLQQSGAELVKPGASVKLSCTASGFNIKDTYMHWVKQRPEQGLEWIGRIDPANGNTKYDPKFQGKATITADTSSNTAYLQLSSLTSEDTAVYYCARGGGDGYPFYAMDYWGQGTSVTVSS
^^^^^^^^ ^^^^^^^^ ^^^^^^^^^^^^^^^

Best Alignment of Margetuximab light chain to a sequence from OAS
DIVMTQSHKFMSTSVGDRVSITCKASQDVNTAVAWYQQKPGHSPKLLIYSASFRYTGVPDRFTGSRSGTDFTFTISSVQAEDLAVYYCQQHYTTPPTFGGGTKVEIK
|||||||||||||||||||||||||||||.|||||||||||.||||||||||.||||||||||||.||||||||||||||||||||||||||.||||||||||.|||
DIVMTQSHKFMSTSVGDRVSITCKASQDVSTAVAWYQQKPGQSPKLLIYSASYRYTGVPDRFTGSGSGTDFTFTISSVQAEDLAVYYCQQHYSTPPTFGGGTKLEIK
^^^^^^ ^^^ ^^^^^^^^^

Best Alignment of Margetuximab heavy chain CDRs to a sequence from OAS
QVQLQQSGPELVKPGASLKLSCTASGFNIKDTYIHWVKQRPEQGLEWIGRIYPTNGYTRYDPKFQDKATITADTSSNTAYLQVSRLTSEDTAVYYCSRWGGDGFYAMDYWGQGASVTVSS
.|.|..||.||||||||.|||||||||||||||.|||||||||||||||||.|.||.|.||||||.||||||||||||||||.|.||||||||||..||||||.|||||||||.||||||

```
EVMLVESGAELVKPGASVKLSCTASGFNIKDTYMHWVKQRPEQGLEWIGRIDPANGNTKYDPKFQGKATITADTSSNTAYLQLSSLTSEDTAVYYGARWGGDGVYAMDYWGQGPSVTVSS
                    ^^^^^^^^          ^^^^^^^^                ^^^^^^^^^^^^^
```

Best Alignment of Margetuximab light CDRs chain to a sequence from OAS

```
DIVMTQSHKFMSTSVGDRVSITCKASQDVNTAVAWYQQKPGHSPKLLIYSASFRYTGVPDRFTGSRSGTDFTFTISSVQAEDLAVYYCQQHYTTPPTFGGGTKVEIK
||.|||||||||||||||||||||||||||||||||||||||||||||||.||||||||||.|||||||||.||||||||||||||||.||||||||||.|||
DIMMTQSHKFMSTSVGDRVSITCKASQDVNTAVAWYQQKPGHSPKLLIYSASFRCTGVPDRFTGSGSGTDFTFTINSVQAEDLAVYYCQQHYSTPPTFGGGTKLEIK
                     ^^^^^^          ^^^              ^^^^^^^^^
```

Best Alignment of Margetuximab CDR-H3 chain to a sequence from OAS

```
QVQLQQSGPELVKPGASLKLSCTASGFNIKDTYIHWVKQRPEQGLEWIGRIYPTNGYTRYDPKFQDKATITADTSSNTAYLQVSRLTSEDTAVYYCSRWGGDGFYAMDYWGQGASVTVSS
||||||.|.|||.||||.||||.|||.........|||||||||||||||.|.....||..||.|||..|.|.||.|||.|.|.|||||.||||||||||..||||||||||.||||||
QVQLQQPGAELVRPGASVKLSCKASGYTFTNYWMNWVKQRPEQGLEWIGRIDPYDSEIRYNQKFKDKAILTVDKSSRTAYMQLSSLTSEDSAVYYCSRWGGITFYAMDYWGQGTSVTVSS
                    ^^^^^^^^          ^^^^^^^^                ^^^^^^^^^^^^^
```

Therapeutic : Matuzumab

Best Alignment of Matuzumab heavy chain to a sequence from OAS

```
QVQLVQSGAEVKKPGASVKVSCKASGYTFTSHWMHWVRQAPGQGLEWIGEFNPSNGRTNYNEKFKSKATMTVDTSTNTAYMELSSLRSEDTAVYYCASRDYDYDGRYFDYWGQGTLVTVSS
||||.|||||..|||||||.|||||||||||.|||||.|.||||||||||.||||||||||||||||||.|||.|..||||.||||.|||.|||||||.||||||.|||.||.||.|||||
QVQLQQSGAELVKPGASVKLSCKASGYTFTSYWMHWVKQRPGQGLEWIGEINPSNGRTNYNEKFKSKATLTVDKSSSTAYMQLSSLTSEDSAVYYCASVDYDYDGGYFDVWGAGTTVTVSS
                    ^^^^^^^^          ^^^^^^^^                ^^^^^^^^^^^^^^
```

Best Alignment of Matuzumab light chain to a sequence from OAS

```
DIQMTQSPSSLSASVGDRVTITCSASSSV-TYMYWYQQKPGKAPKLLIYDTSNLASGVPSRFSGSGSGTDYTFTISSLQPEDIATYYCQQWSSHIFTFGQGTKVEIK
|||||||||||||||||||||||.||..|..|..|||||||||||||||..|||.||||||||||||||||||||||||||||||.|||||||..|...|||||||||||
DIQMTQSPSSLSASVGDRVTITCRASENVNNYLNWYQQKPGKAPKLLIYKASNLQSGVPSRFSGSGSGTDYTFTISSLQPEDVATYYCQQHNSYPRTFGQGTKVEIK
                     ^^^^^^          ^^^              ^^^^^^^^^
```

Best Alignment of Matuzumab heavy chain CDRs to a sequence from OAS

```
QVQLVQSGAEVKKPGASVKVSCKASGYTFTSHWMHWVRQAPGQGLEWIGEFNPSNGRTNYNEKFKSKATMTVDTSTNTAYMELSSLRSEDTAVYYCASRDYDYDGRYFDYWGQGTLVTVSS
||||.|.|||..|||||||.|||||||||||.|||||.|.||||||||||.||||||||||||||||||.|||.|..||||.||||.|||.|||||||.||||||.|||.||.||.|||||
QVQLQQPGAELVKPGASVKLSCKASGYTFTSYWMHWVKQRPGQGLEWIGEINPSNGRTNYNEKFKSKATLTVDKSSSTAYMQLSSLTSEDSAVYYCASVDYDYDGGYFDVWGAGTTVTVSS
                    ^^^^^^^^          ^^^^^^^^                ^^^^^^^^^^^^^^
```

Best Alignment of Matuzumab light CDRs chain to a sequence from OAS

```
DIQMTQSPSSLSASVGDRVTITCSASSSVTYMYWYQQKPGKAPKLLIYDTSNLASGVPSRFSGSGSGTDYTFTISSLQPEDIATYYCQQWSSHIFTFGQGTKVEIK
.|..||||....||.|..||.||||||||||||.|||||.|..||..|||||.||||||.|||||||||.|..||||...||.|||||||||||..||||.|||.|||
QIVLTQSPAIMPASPGEKVTMTCSASSSVTYMHWYQQKSGTSPKRWIYDTSKLASGVPARFSGSGSGTSYSLTISSMEAEDAATYYCQQWSSNPFTFGSGTKLEIK
                     ^^^^^          ^^^              ^^^^^^^^^
```

Best Alignment of Matuzumab CDR-H3 chain to a sequence from OAS

```
QVQLVQSGAEVKKPGASVKVSCKASGYTFTSHWMHWVRQAPGQGLEWIGEFNPSNGRTNYNEKFKSKATMTVDTSTNTAYMELSSLRSEDTAVYYCASRDYDYDGRYFDYWGQGTLVTVSS
.||.|.||....|||.|.|.||.|||.||....||||||||..||||............|....|...|...|...||......||||||||.|||||||||||||.|||||||||..||||
EVQRVESGGGLVKPGGSLKLSCAASGFTFSDYGMHWVRQAPEKGLEWVAYISSGSSTIYYADTVKGRFTISRDNAKNTLFLQMTSLRSEDTAMYYCASRDYDYDGGYFDYWGQGTTLTVSS
                    ^^^^^^^^          ^^^^^^^^                ^^^^^^^^^^^^^^
```

Therapeutic : Mavrilimumab

Best Alignment of Mavrilimumab heavy chain to a sequence from OAS

```
QVQLVQSGAEVKKPGASVKVSCKVSGYTLTELSIHWVRQAPGKGLEWMGGFDPEENEIVYAQRFQGRVTMTEDTSTDTAYMELSSLRSEDTAVYYCAIVGSFS-PLTLGLWGQGTMVTVSS
|||||||||||||||||||||||||||||||||.||||||||||||||||||||..|..|||.|||||||||||||||||||||||||||||||||||||.|||.|..|.||.|||||.|||||
QVQLVQSGAEVKKPGASVKVSCKVSGYTLTELSMHWVRQAPGKGLEWMGGFDPEDGETIYAQKFQGRVTMTEDTSTDTAYMELSSLRSEDTAVYYCARVGSSSWYLGLGYWGQGTLVTVSS
                    ^^^^^^^^          ^^^^^^^^                ^^^^^^^^^^^^^^
```

Best Alignment of Mavrilimumab light chain to a sequence from OAS

```
QSVLTQPPSVSGAPGQRVTISCTGSGSNIGAPYDVSWYQQLPGTAPKLLIYHNNKRPSGVPDRFSGSKSGTSASLAITGLQAEDEADYYCATVEAGLSGSVFGGGTKLTVL
|||||||||||||||||||||||||||.||||||||||.|||||||||||||||.||.||||||||||||||||||||||||||||||||||||||.|....||||||||||||||
QSVLTQPPSVSGAPGQRVTISCTGSSSNIGAPYDVHWYQQLPGTAPKLLIYGNNNRPSGVPDRFSGSKSGTSASLAITGLQAEDEADYYCQTYDNSLSGSVFGGGTKLTVL
                    ^^^^^^^^^          ^^^              ^^^^^^^^^^^
```

Best Alignment of Mavrilimumab heavy chain CDRs to a sequence from OAS

```
................|||||||||||||||||.||||||||||||||||||||..|..|||.|||||||||||||||||||.||||||||||||||||..||.|.||||.|||||.|||||
----------------SVKVSCKVSGYTLTELSMHWVRQAPGKGLEWMGGFDPEDGETIYAQKFQGRVTMTEDTSTDTAYMVLSSLRSEDTAVYYCATDGSGSYLTLGYWGQGTLVTVSS
                    ^^^^^^^^          ^^^^^^^^                ^^^^^^^^^^^^^
```

Best Alignment of Mavrilimumab light CDRs chain to a sequence from OAS

```
QSVLTQPPSVSGAPGQRVTISCTGSGSNIGAPYDVSWYQQLPGTAPKLLIYHNNKRPSGVPDRFSGSKSGTSASLAITGLQAEDEADYYCATVEAGLSGSVFGGGTKLTVL
|||||||||||||||||||||||||||.||||||||||.||.||||.||||||||.||.||||.||||||||||||||.|.||||||..||||||||||...||||.|||||||||||
QSVLTQPPSVSGAPGQRVTISCTGSRSNIGAPYDVHWYKQLPGLAPKLLIYENNERPSGIPDRFSGSKSGTSATLDITGLQTGDEADYYCATWDDGLSGRVFGGGTKLTVL
                    ^^^^^^^^^          ^^^              ^^^^^^^^^^^
```

Best Alignment of Mavrilimumab CDR-H3 chain to a sequence from OAS

```
QVQLVQSGAEVKKPGASVKVSCKVSGYTLTELSIHWVRQAPGKGLEWMGGFDPEENEIVYAQRFQGRVTMTEDTSTDTAYMELSSLRSEDTAVYYCAIVGSFSPLTLGLWGQGTMVTVSS
................|||||||.||||.|...|..||||||.||||||...|......|||.||.|||.|.|.|..||||||||||||||||.||||...|.|||||||.||||||.|||||
----------------SVKVSCKASGYTFTSYGISRVRQAPGQGLEWMGWITPFNGNTNYAQKFQDRVTITRDRSMSTAYMELSSLRSEDTAMYYCAGGDSSSPLTLGYWGQGTLVTVSS
                    ^^^^^^^^          ^^^^^^^^                ^^^^^^^^^^^^^
```

Therapeutic : Mepolizumab

Best Alignment of Mepolizumab heavy chain to a sequence from OAS

```
QVTLRESGPALVKPTQTLTLTCTVSGFSL--TSYSVHWVRQPPGKGLEWLGVIWASGGTDYNSALMSRLSISKDTSRNQVVLTMTNMDPVDTATYYCARDPPSS----LLRLDYWGRGTPVTVSS
||||||||||||||||||||||||.|||||......|.|.||||||.||||.||.......||..|..||||||||||.||||||||||||||||||||||||....|.......|||||.||.|||||
QVTLRESGPALVKPTQTLTLTCTFSGFSLNTDEMCVSWIRQPPGKALEWLAVIHRDDNKFYNTSLKTRLSISKDTSKNQVVLTMTNMDPVDTATYYCARIRNYSGYNGYGGLDYWGQGTLVTVSS
                    ^^^^^^^^^^          ^^^^^^^                ^^^^^^^^^^^^^^^^^
```

Best Alignment of Mepolizumab light chain to a sequence from OAS

```
DIVMTQSPDSLAVSLGERATINCKSSQSLLNSGNQKNYLAWYQQKPGQPPKLLIYGASTRESGVPDRFSGSGSGTDFTLTISSLQAEDVAVYYCQNVHSFPFTFGGGTKLEIK
||||||||||||||||||||||||||||||.|.|.|.||||||||||||||||||||.||||||||||||||||||||||||||||||||||||||||..||.|||||.|||||||
DIVMTQSPDSLAVSLGERATINCKSSQSVLYSSNNKNYLAWYQQKPGQPPKLLIYWASTRESGVPDRFSGSGSGTDFTLTISSLQAEDVAVYYCQQDHSPPFTFGQGTKLEIK
                     ^^^^^^^^^^^^          ^^^              ^^^^^^^^^
```

Best Alignment of Mepolizumab heavy chain CDRs to a sequence from OAS

```
QVTLRESGPALVKPTQTLTLTCTVSGFSLTSYSVHWVRQPPGKGLEWLGVIWASGGTDYNSALMSRLSISKDTSRNQVVLTMTNMDPVDTATYYCARDPPSSLLRLDYWGRGTPVTVSS
||.|.||||.||.|.|.|..||||||||||||.|||||||||||||||||||||.|.|.|||||||||||||||.|..||.|.|......|.|.||||||.|..||||||||.||.|||||
QVQLKESGPGLVAPSQSLSITCTVSGFSLTSYGVHWVRQPPGKGLEWLGVIWAGGSTNYNSALMSRLSISKDNSKSQVCLKMNSLQIDDPAMYYCARDAPLLLLRLDYWGQGTSVTVSS
                    ^^^^^^^^          ^^^^^^^                ^^^^^^^^^^^^^
```

Best Alignment of Mepolizumab light CDRs chain to a sequence from OAS

```
DIVMTQSPDSLAVSLGERATINCKSSQSLLNSGNQKNYLAWYQQKPGQPPKLLIYGASTRESGVPDRFSGSGSGTDFTLTISSLQAEDVAVYYCQNVHSFPFTFGGGTKLEIK
||||||||.||.||.||..|..||||||||||||||||||||||||||||||||||||||||||||||.||||||||||||||.||||.|||||||||||.|||||.|||||||
DIVMTQSPSSLSVSAGEKVTMSCKSSQSLLNSGNQKNYLAWYQQKPGQPPKLLIYGASTRESGVPDRFTGSGSGTDFTLTISSVQAEDLAVYYCQNVHSYPFTFGSGTKLEIK
                     ^^^^^^^^^^^^          ^^^              ^^^^^^^^^
```

Best Alignment of Mepolizumab CDR-H3 chain to a sequence from OAS

```
QVTLRESGPALVKPTQTLTLTCTVSGFSLTSYSVHWVRQPPGKGLEWLGVIWAS-GGTDYNSALMSRLSISKDTSRNQVVLTMTNMDPVDTATYYCARDPPSSLLRLDYWGRGTPVTVSS
.............|...|.|.|..||||...|..|||||.|||||||...||.......|......|..||.|.|.|...|.|......|||.||||||||||.|.|||||.||||||||
```

```
-------------PGGSLRLSCAASGFSFSDYGMHWVRQAPGKGLEWVAFIWYDGNNIKYADSVKGRFTISRDNSKNTMYLQMNTLRVEDTAVYYCARDPPSSGLHLDYWGQGTPVTVSS
            ^^^^^^^^          ^^^^^^^^                          ^^^^^^^^^^^^^
```

Therapeutic : Mirikizumab

Best Alignment of Mirikizumab heavy chain to a sequence from OAS

```
QVQLVQSGAEVKKPGSSVKVSCKASGYKFTRYVMHWVRQAPGQGLEWMGYINPYNDGTNYNEKFKGRVTITADKSTSTAYMELSSLRSEDTAVYYCARNW---------------DTGLWGQGTTVTVSS
|||||||||||||||||||||||||||.||.|.|||||||||||||||||.|||...||||..||.||||||||||||||||||||||||||||||||||.....................||||||||||
QVQLVQSGAEVKKPGSSVKVSCKASGYTFTGYYMHWVRQAPGQGLEWMGWINPNSGGTNYAQKFQGRVTITADKSTSTAYMELSSLRSEDTAVYYCARERIAAAAKSSDHHYYYYGMDVWGQGTTVTVSS
            ^^^^^^^^          ^^^^^^^^                          ^^^^^^^^^^^^^^^^^^^^^^
```

Best Alignment of Mirikizumab light chain to a sequence from OAS

```
DIQMTQSPSSLSASVGDRVTITCKASDHILKFLTWYQQKPGKAPKLLIYGATSLETGVPSRFSGSGSGTDFTLTISSLQPEDFATYYCQMYWSTPFTFGGGTKVEIK
||||||||||||||||||||||||.||..|...|.|||||||||||||||.|.||||||||||||||||||||||||||||||||||||||||.|.|||.||||||||||
DIQMTQSPSSLSASVGDRVTITCRASQGISSYLNWYQQKPGKAPKLLIYAASSLETGVPSRFSGSGSGTDFTLTISSLQPEDFATYYCQQYDSTPITFGGGTKVEIK
             ^^^^^^         ^^^                     ^^^^^^^^^
```

Best Alignment of Mirikizumab heavy chain CDRs to a sequence from OAS

```
QVQLVQSGAEVKKPGSSVKVSCKASGYKFTRYVMHWVRQAPGQGLEWMGYINPYNDGTNYNEKFKGRVTITADKSTSTAYMELSSLRSEDTAVYYCARNWDTGLWGQGTTVTVSS
.|||.|||.|..|||.|||.|||||||||||||||||.|.|||||||.||||||||||.|||||||..|.|.|||.||||||||||.|||.||||||.|||...|||.|.||||.
EVQLQQSGPELVKPGASVKMSCKASGYKFTRYVMHWVKQKPGQGLEWIGYINPYNDGTQYNEKFKGKATLTSDKSSSTAYMELSSLTSEDSAVYYCAPNWDSAYWGQWTLVTVSA
            ^^^^^^^^          ^^^^^^^^                          ^^^^^^^^
```

Best Alignment of Mirikizumab light CDRs chain to a sequence from OAS

```
DIQMTQSPSSLSASVGDRVTITCKASDHILKFLTWYQQKPGKAPKLLIYGATSLETGVPSRFSGSGSGTDFTLTISSLQPEDFATYYCQMYWSTPFTFGGGTKVEIK
|||||||.|.||.|.|.|||||||||||||.|.|.||||||||.||.|||.||||||||||||||||||||...||.|.|||.||.||||||.|||||.||||||||.|||
DIQMTQSSSYLSVSLGGRVTITCKASDHINKWLAWYQQKPGNAPRLLISGATSLETGVPSRFSGSGSGKNYTLNITSLQTEDVATYYCQQYWSTPYTFGGGTKLEIK
             ^^^^^^         ^^^                     ^^^^^^^^^
```

Best Alignment of Mirikizumab CDR-H3 chain to a sequence from OAS

```
QVQLVQSGAEVKKPGSSVKVSCKASGYKFTRYVMHWVRQAPGQGLEWMGYINPYNDGTNYNEKFKGRVTITADKSTSTAYMELSSLRSEDTAVYYCARNWDTGLWGQGTTVTVSS
................|||||||||||.||.|..|||||||||...||||||.....|.|......|||||.|.|.||.||.||.|.||||||||||||.||||||||||.|||||
---------------ASVKVSCKASGYIFTNYAIHWVRQAPGQRPQWMGYINTAIGNTKYSQSLQDRVTITSDTSASTVYMHLSRLTSEDTAVYYCARDWDTGLWGQGTLVTVSS
            ^^^^^^^^          ^^^^^^^^                          ^^^^^^^^
```

Therapeutic : Mirvetuximab

Best Alignment of Mirvetuximab heavy chain to a sequence from OAS

```
QVQLVQSGAEVVKPGASVKISCKASGYTFTGYFMNWVKQSPGQSLEWIGRIHPYDGDTFYNQKFQGKATLTVDKSSNTAHMELLSLTSEDFAVYYCTRYDGS---RAMDYWGQGTTVTVSS
.|||.|||.|.|||||||||||||||||.|||||||||||||.||||||||||.||.||||||||||.|||||||||||||.|||||||||||||||||||||.||.||....||||||||||.|||||
-VQLQQSGPELVKPGASVKISCKASGYSFTGYFMNWVKQSHGQSLEWIGRINPYNGDTFYNQKFKGKATLTVDKSSSTAHMELLSLTSEDFAVYYCARYYGSSYDYAMDYWGQGTSVTVSS
            ^^^^^^^^          ^^^^^^^^                          ^^^^^^^^^^^^^^
```

Best Alignment of Mirvetuximab light chain to a sequence from OAS

```
DIVLTQSPLSLAVSLGQPAIISCKASQSVSFAGTSLMHWYHQKPGQQPRLLIYRASNLEAGVPDRFSGSGSKTDFTLTISPVEAEDAATYYCQQSREYPYTFGGGTKLEIK
||||||||.||||||||.||||||||||||||||||||||||.||||||||.|||||||||||||||.||||||||.||||||.|.|||..||||||||||||||.||||||||||||
DIVLTQSPASLAVSLGQRAIISCKASQSVSFAGTSLMHWYQQKPGQQPKLLIYRASNLEAGVPTRFSGSGSRTDFTLNIHPVEEDDAATYYCQQSREYPRTFGGGTKLEIK
             ^^^^^^^^^^         ^^^                     ^^^^^^^^^
```

Best Alignment of Mirvetuximab heavy chain CDRs to a sequence from OAS

```
QVQLVQSGAEVVKPGASVKISCKASGYTFTGYFMNWVKQSPGQSLEWIGRIHPYDGDTFYNQKFQGKATLTVDKSSNTAHMELLSLTSEDFAVYYCTRYDGSRAMDYWGQGTTVTVSS
.|||.|.|.|.|||||||||||||||||.||||||||||.||.|.|||||||||.|||||||||||||.||||||||||||.||||||.||.|||.|||||.||.|..||||||||||.|||||
-VQLQQPGPELVKPGASVKISCKASGYSFTGYFMNWVMQSHGKSLEWIGRINPYDGDTFYNQKFKGKATLTVDKSSSTAHMELRSLASEDSAVYYCARYYGQGAMDYWGQGTSVTVSS
            ^^^^^^^^          ^^^^^^^^                          ^^^^^^^^^^^
```

Best Alignment of Mirvetuximab light CDRs chain to a sequence from OAS

```
DIVLTQSPLSLAVSLGQPAIISCKASQSVSFAGTSLMHWYHQKPGQQPRLLIYRASNLEAGVP-DRFSGSGSKTDFTLTISPVEAEDAATYYCQQSREYPYTFGGGTKLEIK
||||||||.||||||||.||||||||||||||||||||||||.||||||||.|||||||||||||.........|........|.|||..||||||||||||||||||||||||||||
DIVLTQSPASLAVSLGQRAIISCKASQSVSFAGTSLMHWYQQKPGQQPKLLIYRASNLEAGGSYQGLVAVGLGQTSPSNIHPVEEDDAATYYCQQSREYPYTFGGGTKLEIK
             ^^^^^^^^^^         ^^^                     ^^^^^^^^^
```

Best Alignment of Mirvetuximab CDR-H3 chain to a sequence from OAS

```
QVQLVQSGAEVVKPGASVKISCKASGYTFTGYFMNWVKQSPGQSLEWIGRIHPYDGDTFYNQKFQGKATLTVDKSSNTAHMELLSLTSEDFAVYYCTRYDGSRAMDYWGQGTTVTVSS
||||.|||||.|.|||||..||||||||||.|.|.||||.|...|||||.|.|..|.|.|||||.|||.||.||||.||.|||.||||||.|||||||||.|||||||||||.|||||
QVQLQQSGAELVRPGASVALSCKASGYTFTDYEMHWVKQTPVHGLEWIGAIDPETGGTAYNQKFKGKAILTADKSSSTAYMELRSLTSEDSAVYYCTRYDSSRAMDYWGQGTSVTVSS
            ^^^^^^^^          ^^^^^^^^                          ^^^^^^^^^^^
```

Therapeutic : Modotuximab

Best Alignment of Modotuximab heavy chain to a sequence from OAS

```
QVQLQQPGAELVEPGGSVKLSCKASGYTFTSHWMHWVKQRPGQGLEWIGEINPSSGRNNYNEKFKSKATLTVDKSSSTAYMQFSSLTSEDSAVYYCVRYYGYDEAMDYWGQGTSVTVSS
||||||||||||.||.||||||||||||||||.||||||||||||||||||||||.||.||||||||||||||||||||||||.|||||||||||||.|||||.|||||||||||||||||
QVQLQQPGAELVKPGASVKLSCKASGYTFTSYWMHWVKQRPGQGLEWIGEINPSNGRTNYNEKFKSKATLTVDKSSSTAYMQLSSLTSEDSAVYYCARYYGY-EAMDYWGQGTSVTVSS
            ^^^^^^^^          ^^^^^^^^                          ^^^^^^^^^^^^
```

Best Alignment of Modotuximab light chain to a sequence from OAS

```
DIVMTQAAFSNPVTLGTSASISCRSSKSLLHSNGITYLYWYLQKPGQSPQLLIYQMSNLASGVPDRFSSSGSGTDFTLRISRVEAEDVGVYYCAQNLELPYTFGGGTKLEIK
||||||||||||||||||||||||||||||||||||||||||||||||||||||||||||||||||||||||||||||||||||||||||||||||||||||||||||||||||
DIVMTQAAFSNPVTLGTSASISCRSSKSLLHSNGITYLYWYLQKPGQSPQLLIYQMSNLASGVPDRFSSSGSGTDFTLRISRVEAEDVGVYYCAQNLELPYTFGGGTKLEIK
             ^^^^^^^^^^^         ^^^                     ^^^^^^^^^
```

Best Alignment of Modotuximab heavy chain CDRs to a sequence from OAS

```
QVQLQQPGAELVEPGGSVKLSCKASGYTFTSHWMHWVKQRPGQGLEWIGEINPSSGRNNYNEKFKSKATLTVDKSSSTAYMQFSSLTSEDSAVYYCVRYYGYDEAMDYWGQGTSVTVSS
.|||||.||||..||.||||||||||||||||.||||||||||||||||||.||||||...||.|||.||||||.|||||||.|.||||.|||||..|.|.||||||||||||||||.|||
-VQLQQSGAELAKPGASVKLSCKASGYTFTSYWMHWVKQRPGQGLEWIGYINPSSGYTKYNQKFKDKATLTVEKSSSTAYIQLSSLTYEDSAVSSCARVYGYDEAMDYWGQGTSVAVSS
            ^^^^^^^^          ^^^^^^^^                          ^^^^^^^^^^^^
```

Best Alignment of Modotuximab light CDRs chain to a sequence from OAS

```
DIVMTQAAFSNPVTLGTSASISCRSSKSLLHSNGITYLYWYLQKPGQSPQLLIYQMSNLASGVPDRFSSSGSGTDFTLRISRVEAEDVGVYYCAQNLELPYTFGGGTKLEIK
.|.||.||...||.|..||||||.||||||||||||||||||||||||||||||||||||||||||||||||||||||||||||||||||||||||||||||||||||||||
-IAMTPAACCSPVSLLPSASISCGSSKSLLHSNGITYLYWYLQKPGQSPQLLIYQMSNLASGVPDRFSSSGSGTDFTLRISRVEAEDVGVYYCAQNLELPYTFGGGTKLEIK
             ^^^^^^^^^^^         ^^^                     ^^^^^^^^^
```

Best Alignment of Modotuximab CDR-H3 chain to a sequence from OAS

```
QVQLQQPGAELVEPGGSVKLSCKASGYTFTSHWMHWVKQRPGQGLEWIGEINPS--SGRNNYNEKFKSKATLTVDKSSSTAYMQFSSLTSEDSAVYYCVRYYGYDEAMDYWGQGTSVTVSS
.|.|...|..||.||||..|||.|||.|||...|.||.|.||..|||.|.|..........|....|...|...|.|.|..|.|...|...|||.|||.||||||||||||||||||||||
EVKLVESGGGLVQPGGSLSLSCAASGFTFTDYYMSWVRQPPGKALEWLGFIRNKANGYTTEYSASVKGRFTISRDNSQSILYLQMNALRADDSATYYCARYYGYDEAMDYWGQGTSVTVSS
            ^^^^^^^^          ^^^^^^^^^^                          ^^^^^^^^^^^^
```

Therapeutic : Mogamulizumab

Best Alignment of Mogamulizumab heavy chain to a sequence from OAS

```
EVQLVESGGDLVQPGRSLRLSCAASGFIFSNYGMSWVRQAPGKGLEWVATISSASTYSYYPDSVKGRFTISRDNAKNSLYLQMNSLRVEDTALYYCGRHSDG-NFAFGYWGQGTLVTVSS
||||.||||.|||||.|||||||||||.||.|.|||||||||||||||||||||.|.|.||.|||||||||||||||||||||||||||||.||||.|||.|..||.||...|||||||||||||
EVQLLESGGGLVQPGGSLRLSCAASGFTFSSYAMSWVRQAPGKGLEWVATISSSSSYIYYTDSVKGRFTISRDNAKNSLYLQMNSLRAEDTAVYYCARSRDGYNFDIDYWGQGTLVTVSS
            ^^^^^^^^          ^^^^^^^^                          ^^^^^^^^^^^^^
```

Best Alignment of Mogamulizumab light chain to a sequence from OAS
DVLMTQSPLSLPVTPGEPASISCRSSRNIVHINGDTYLEWYLQKPGQSPQLLIYKVSNRFSGVPDRFSGSGSGTDFTLKISRVEAEDVGVYYCFQGSLLPWTFGQGTKVEIK
||.|||||||||||||||||||||||.....|||.|||||||||||||||||||.||||.|||||||||||||||||||||||||||||||||.||...|||||||||||||
DVVMTQSPLSLPVTPGEPASISCRSSQSLMPINGYTYLEWYLQKPGQSPQLLIYLVSNRASGVPDRFSGSGSGTDFTLKISRVEAEDVGVYYCMQGLQIPWTFGQGTKVEIK
^^^^^^^^^^^ ^^^ ^^^^^^^^^

Best Alignment of Mogamulizumab heavy chain CDRs to a sequence from OAS
EVQLVESGGDLVQPGRSLRLSCAASGFIFSNYGMSWVRQAPGKGLEWVATISSASTYSYYPDSVKGRFTISRDNAKNSLYLQMNSLRVEDTALYYCGRHSDGNFAFGYWGQGTLVTVSS
||||||||||||.||.||.||||||||.||.||||||||.|.|.|||||||||...|||||||||||||||||||||.|||||.||..||||.|||.||.||||.|.||||||..||||
EVQLVESGGDLVKPGGSLKLSCAASGFTFSSYGMSWVRQTPDKRLEWVATISSGGSYSYYPDSVKGRFTISRDNAKNTLYLQMSSLKSEDTAMYYCARHEDGNFHFDYWGQGTTLTVSS
^^^^^^^^ ^^^^^^^^ ^^^^^^^^^^^^

Best Alignment of Mogamulizumab light CDRs chain to a sequence from OAS
DVLMTQSPLSLPVTPGEPASISCRSSRNIVHINGDTYLEWYLQKPGQSPQLLIYKVSNRFSGVPDRFSGSGSGTDFTLKISRVEAEDVGVYYCFQGSLLPWTFGQGTKVEIK
||.|||.||.|||..|..||||||||..|||.|||||||||||||||||.|||||||||||||||||||.|||||||||||||||||.|||||||||..|||||.|||.|||
DVVMTQTPLFLPVRLGDHASISCRSSQSIVHSNGDTYLEWYLQKPGQSPKLLIYKVSNRFSGVPDRFSGGGSGTDFTLKISRVEAEDLGVYYCFQGSHVPWTFGGGTKLEIK
^^^^^^^^^^^ ^^^ ^^^^^^^^^

Best Alignment of Mogamulizumab CDR-H3 chain to a sequence from OAS
EVQLVESGGDLVQPGRSLRLSCAASGFIFSNYGMSWVRQAPGKGLEWVATISSASTYSYYPDSVKGRFTISRDNAKNSLYLQMNSLRVEDTALYYCGRHSDGNFAFGYWGQGTLVTVSS
|||.|||||.||.||.||.||||||||.||.|.||||||.|.|.|||||||||...|.||||||.||||||||||||.|.|||..||.||.|.|||.|.|||||.|||||||||||||.
EVQGVESGGGLVKPGGSLKLSCAASGFTFSSYAMSWVRQTPEKRLEWVATISSGGSYTYYPDSVEGRFTISRDNAKNTLCLQMSGLRSEDAAMYYCARGSDGNFPFGYWGQGTLVTVSA
^^^^^^^^ ^^^^^^^^ ^^^^^^^^^^^^

Therapeutic : Motavizumab

Best Alignment of Motavizumab heavy chain to a sequence from OAS
QVTLRESGPALVKPTQTLTLTCTFSGFSLSTAGMSVGWIRQPPGKALEWLADIWWDDKKHYNPSLKDRLTISKDTSKNQVVLKVTNMDPADTATYYCARDMIF-NFYFDVWGQGTTVTVSS
|||||||||||||||||||||||||||||||.||||.||||||||||||||.|.|||.|.||.|||.|||||||||||||||..|||||.|||||||||.......|||.|||||.|||||
QVTLRESGPALVKPTQTLTLTCTFSGFSLSTSGMSVSWIRQPPGKALEWLALIDWDDAKYYNTSLKTRLTISKDTSKNQVVLTMTNMDPVDTATYYCARSYGGRTYYFDYWGQGTLVTVSS
^^^^^^^^^^ ^^^^^^^ ^^^^^^^^^^^^^

Best Alignment of Motavizumab light chain to a sequence from OAS
DIQMTQSPSTLSASVGDRVTITCSASSRV-GYMHWYQQKPGKAPKLLIYDTSKLASGVPSRFSGSGSGTEFTLTISSLQPDDFATYYCFQGSGYPFTFGGGTKVEIK
|||||||||||||||||||||||.||.|......||||||||||||||||.|.|||||||||||||||||||||||||||||||||||.|..|||.|||.|||||||
DIQMTQSPSTLSASVGDRVTITCRASQRFTTWLAWYQQKPGKAPKLLIYDVSSLASGVPSRFSGSGSGTEFTLTISSLQPDDFATYYCQQYDGYPWTFGQGTKVEIK
^^^^^^ ^^^ ^^^^^^^^^

Best Alignment of Motavizumab heavy chain CDRs to a sequence from OAS
QVTLRESGPALVKPTQTLTLTCTFSGFSLSTAGMSVGWIRQPPGKALEWLADIWWDDKKHYNPSLKDRLTISKDTSKNQVVLKVTNMDPADTATYYCARDMIFNFYFDVWGQGTTVTVSS
||||.||||..|.|.|||.|||..|||||||.||.|||||||.||.|||||.|||||.|.||..||..||||||||||||.||....|.||||||||||...|.|||||||.||||||||
QVTLKESGPGIVQPSQTLGLTCSCSGFSLSTSGMGVGWIRQPSGKGLEWLAHIWWDDDKYYNTALKSGLTISKDTSKNQVFLKIASVDTADTATYYCARISLFIFYFDVWGAGTTVTVSS
^^^^^^^^^^ ^^^^^^^ ^^^^^^^^^^^^

Best Alignment of Motavizumab light CDRs chain to a sequence from OAS
DIQMTQSPSTLSASVGDRVTITCSASSRVGYMHWYQQKPGKAPKLLIYDTSKLASGVPSRFSGSGSGTEFTLTISSLQPDDFATYYCFQGSGYPFTFGGGTKVEIK
....||||...|||.|..||.||||||||.||||||||....|||.|||||.|||||..||||||||....|||||....|.||||||||||||.|||||||.|||
ENVLTQSPAIMSASPGEKVTMTCSASSRVSYMHWYQQKSSTSPKLWIYDTSILASGVTGRFSGSGSGNSYSLTISSMEAEDVATYYCFQGSGYPYTFGGGTKLEIK
^^^^^ ^^^ ^^^^^^^^^

Best Alignment of Motavizumab CDR-H3 chain to a sequence from OAS
QVTLRESGPALVKPTQTLTLTCTFSGFSLSTAGMSVGWIRQPPGKALEWLADIWWD-DKKHYNPSLKDRLTISKDTSKNQVVLKVTNMDPADTATYYCARDMIFNFYFDVWGQGTTVTVSS
||.|..||..||.|.......|..||...........|..|.|...|||...|........||...|...|...|.|.|..............|.|.|.|.||||.|||.||||||.||||||||
QVQLQQSGAELVRPGASVKMSCKASGYTF--TSYNMHWVKQTPRQGLEWIGAIYPGNGDTSYNQKFKGKATLTVDKSSSTAYMQLSSLTSEDSAVYFCARDGIFNWYFDVWGTGTTVTVSS
^^^^^^^^^^ ^^^^^^^^ ^^^^^^^^^^^^

Therapeutic : Muromonab

Best Alignment of Muromonab heavy chain to a sequence from OAS
QVQLQQSGAELARPGASVKMSCKASGYTFTRYTMHWVKQRPGQGLEWIGYINPSRGYTNYNQKFKDKATLTTDKSSSTAYMQLSSLTSEDSAVYYCARYYDDHYCLDYWGQGTTLTVSS
||||||||||||||||||||||||||||||.|||||||||||||||||||||||.||||||||||||||||.||||||||||||||||||||||||||.||..|.||||||||||||||
QVQLQQSGAELARPGASVKMSCKASGYTFTSYTMHWVKQRPGQGLEWIGYINPSSGYTNYNQKFKDKATLTADKSSSTAYMQLSSLTSEDSAVYYCARPYDG-YYLDYWGQGTTLTVSS
^^^^^^^^ ^^^^^^^^ ^^^^^^^^^^^^

Best Alignment of Muromonab light chain to a sequence from OAS
QIVLTQSPAIMSASPGEKVTMTCSASSSVSYMNWYQQKSGTSPKRWIYDTSKLASGVPAHFRGSGSGTSYSLTISGMEAEDAATYYCQQWSSNPFTFGSGTKLEIK
|||||||||||||||||||||||||||||||||||||||||||||||||||||||||||.|.|||||||||||||.||||||||||||||||||||||||||||||
QIVLTQSPAIMSASPGEKVTMTCSASSSVSYMNWYQQKSGTSPKRWIYDTSKLASGVPARFSGSGSGTSYSLTISSMEAEDAATYYCQQWSSNPFTFGSGTKLEIK
^^^^^ ^^^ ^^^^^^^^^

Best Alignment of Muromonab heavy chain CDRs to a sequence from OAS
QVQLQQSGAELARPGASVKMSCKASGYTFTRYTMHWVKQRPGQGLEWIGYINPSRGYTNYNQKFKDKATLTTDKSSSTAYMQLSSLTSEDSAVYYCARYYDDHYCLDYWGQGTTLTVSS
|.|||.|..|||||||||.||||||||||||||||||.||||||||||||||||.|||.||||.|||.|||.||||||||||||||||||.||||||||||..|..||||||||.|..|
QLQLQHSASELARPGASVQMSCKASGYTFTRYTMHWVQQRPGQGLEWIGYINPSSGYTEYNQKLKDKTTLTADKSSSTAYMQLSSLTSEDYAVYYCARYYDYDYAMDYWGQGTTDTDTS
^^^^^^^^ ^^^^^^^^ ^^^^^^^^^^^^

Best Alignment of Muromonab light CDRs chain to a sequence from OAS
QIVLTQSPAIMSASPGEKVTMTCSASSSVSYMNWYQQKSGTSPKRWIYDTSKLASGVPAHFRGSGSGTSYSLTISGMEAEDAATYYCQQWSSNPFTFGSGTKLEIK
||||||||||||||||||||||||||||||||.||||||||||||||||||||||||||.|.|||||||||||||.||||||||||||||||||||||.|||||.|
QIVLTQSPAIMSASPGEKVTMTCSASSSVSYMHWYQQKSGTSPKRWIYDTSKLASGVPARFSGSGSGTSYSLTISSMEAEDAATYYCQQWSSNPFTFGAGTKLELK
^^^^^ ^^^ ^^^^^^^^^

Best Alignment of Muromonab CDR-H3 chain to a sequence from OAS
QVQLQQSGAELARPGASVKMSCKASGYTFTRYTMHWVKQRPGQGLEWIGYINPS--RGYTNYNQKFKDKATLTTDKSSSTAYMQLSSLTSEDSAVYYCARYYDDHYCLDYWGQGTTLTVSS
.|....||..|..||.|...||.|||.|||.|.|.||.|.||..|||.|.|........|.|....|...|...|.|.|..|.|...|..||||.||||||.||.||||||||||||||||
EVKVVESGGGLVQPGGSLSLSCAASGFTFTDYYMSWVRQPPGKALEWLGFIRNKANGYTTEYSASVKGRFTISRDNSQSILYLQMNALRAEDSATYYCARYLDDGYCLDYWGQGTTLTVSS
^^^^^^^^ ^^^^^^^^^^ ^^^^^^^^^^^^

Therapeutic : Natalizumab

Best Alignment of Natalizumab heavy chain to a sequence from OAS
QVQLVQSGAEVKKPGASVKVSCKASGFNIKDTYIHWVRQAPGQRLEWMGRIDPANGYTKYDPKFQGRVTITADTSASTAYMELSSLRSEDTAVYYCAREGYYGN-YGVYAMDYWGQGTLVTVSS
||||||||||||||||||||||||||........|||||||||||||||.|...||.|||..|||||||||.||||||||||||||||||||||||||||||....|.|..|||||||||||||
QVQLVQSGAEVKKPGASVKVSCKASGYTFTSYAMHWVRQAPGQRLEWMGWINAGNGNTKYSQKFQGRVTITRDTSASTAYMELSSLRSEDTAVYYCAREGYYYDSSGYYPLDYWGQGTLVTVSS
^^^^^^^^ ^^^^^^^^ ^^^^^^^^^^^^^^^^^

Best Alignment of Natalizumab light chain to a sequence from OAS
DIQMTQSPSSLSASVGDRVTITCKTSQDINKYMAWYQQTPGKAPRLLIHYTSALQPGIPSRFSGSGSGRDYTFTISSLQPEDIATYYCLQYDNLWTFGQGTKVEIK
||||||||||||||.|..|||||||||||||||||||..|||.|||||||||.||||||||||||||||||.|.||.|.||||||||||||||||||||.|||.|||
DIQMTQSPSSLSASLGGKVTITCKTSQDINKYMAWYQHKPGKGPRLLIHYTSTLQPGIPSRFSGSGSGRDYSFSISNLEPEDIATYYCLQYDNLWTFGGGTKLEIK
^^^^^^ ^^^ ^^^^^^^^

Best Alignment of Natalizumab heavy chain CDRs to a sequence from OAS
QVQLVQSGAEVKKPGASVKVSCKASGFNIKDTYIHWVRQAPGQRLEWMGRIDPANGYTKYDPKFQGRVTITADTSASTAYMELSSLRSEDTAVYYCAREGYYGNYGVYAMDYWGQGTLVTVSS
.|||.|||||..|||||||.||.|||||||||||.|||.|.|.|.|||.||||||||.|||.|||||..|||||||..|||..||||.|||||.|||||.|||||||.||||||||||.|||||

```
-VQLQQSGAELVKPGASVKLSCTASGFNIKDTYMHWVKQRPEQGLEWIGRIDPANGNTKYAPKFQGKATITADTSSNTAYLQLSSLTSEDTAIYYCARTGYYGNYGGYAMDYWGQGTSVTVSS
                   ^^^^^^^^        ^^^^^^^^                  ^^^^^^^^^^^^^^^^
```

Best Alignment of Natalizumab light CDRs chain to a sequence from OAS

```
DIQMTQSPSSLSASVGDRVTITCKTSQDINKYMAWYQQTPGKAPRLLIHYTSALQPGIPSRFSGSGSGRDYTFTISSLQPEDIATYYCLQYDNLWTFGQGTKVEIK
.|||.||||||.||.|..||||||.|||||||.||||..|||.|||||||||.||||||||||||||||||.|.||.|.||||||||||||||||||||.|||.|||
-IQMIQSPSSLPASLGGKVTITCKASQDINKYIAWYQHKPGKGPRLLIHYTSTLQPGIPSRFSGSGSGRDYSFSISNLEPEDIATYYCLQYDNLWTFGGGTKLEIK
                   ^^^^^^          ^^^                ^^^^^^^^
```

Best Alignment of Natalizumab CDR-H3 chain to a sequence from OAS

```
QVQLVQSGAEVKKPGASVKVSCKASGFNIKDTYIHWVRQAPGQRLEWMGRIDPANGYTKYDPKFQGRVTITADTSASTAYMELSSLRSEDTAVYYCAREGYYGNYGVYAMDYWGQGTLVTVSS
||||..||.....|..|....|..|||........||||.||..|||.|.|....|.|.|......|..|..|.|.|.......||...|||.|||||||||||||||||||||||||.|||||
QVQLKESGPGLVAPSQSLSITCTVSGFSLTGYGVNWVRQPPGKGLEWLGMIWGD-GSTDYNSALKSRLSISKDNSKSQVFLKMNSLQTDDTARYYCAREGYYGNYGVYAMDYWGQGTSVTVSS
                   ^^^^^^^^        ^^^^^^^^                  ^^^^^^^^^^^^^^^^
```

Therapeutic : Necitumumab

Best Alignment of Necitumumab heavy chain to a sequence from OAS

```
QVQLQESGPGLVKPSQTLSLTCTVSGGSISSGDYYWSWIRQPPGKGLEWIGYIYYSGSTDYNPSLKSRVTMSVDTSKNQFSLKVNSVTAADTAVYYCARVSIFGVGTFDYWGQGTLVTVSS
|||||||||||||||||||||||||||||||||||||||||||||||||||||||.||||||||||.||||||||||||..|||||||||||||||..||||.||||||||||||||||
QVQLQESGPGLVKPSQTLSLTCTVSGGSISSGDYYWSWIRQPPGKGLEWIGYIYYSGSTYYNPSLKSRVTISVDTSKNQFSLKLSSVTAADTAVYYCARATIFGVDTFDYWGQGTLVTVSS
                   ^^^^^^^^^^        ^^^^^^^                  ^^^^^^^^^^^^^
```

Best Alignment of Necitumumab light chain to a sequence from OAS

```
EIVMTQSPATLSLSPGERATLSCRASQSVSSYLAWYQQKPGQAPRLLIYDASNRATGIPARFSGSGSGTDFTLTISSLEPEDFAVYYCHQYGS-TPLTFGGGTKAEIK
|||.||||||||||||||||||||||||||||||||||||||||||||||||||||||||||||||||||||||||||||||||||||||.||||..||||||||||.|||
EIVLTQSPATLSLSPGERATLSCRASQSVSSYLAWYQQKPGQAPRLLIYDASNRATGIPARFSGSGSGTDFTLTISSLEPEDFAVYYCQQYGSSPPLTFGGGTKVEIK
                   ^^^^^^          ^^^                ^^^^^^^^^^
```

Best Alignment of Necitumumab heavy chain CDRs to a sequence from OAS

```
QVQLQESGPGLVKPSQTLSLTCTVSGGSISSGDYYWSWIRQPPGKGLEWIGYIYYSGSTDYNPSLKSRVTMSVDTSKNQFSLKVNSVTAADTAVYYCARVSIFGVGTFDYWGQGTLVTVSS
.............|.||||||||||||||||.||||||||.|||||||||||||||||.||||||||||.||||||||||||..|||||||||||||||.||||||||||||||||||||
-------------SETLSLTCTVSGGSISSGGYYWSWIRQHPGKGLEWIGYIYYSGSTYYNPSLKSRVTISVDTSKNQFSLKLSSVTAADTAVYYCARVTIFGVGTFDYWGQGTLVTVSS
                   ^^^^^^^^^^        ^^^^^^^                  ^^^^^^^^^^^^^
```

Best Alignment of Necitumumab light CDRs chain to a sequence from OAS

```
EIVMTQSPATLSLSPGERATLSCRASQSVSSYLAWYQQKPGQAPRLLIYDASNRATGIPARFSGSGSGTDFTLTISSLEPEDFAVYYCHQYGSTPLTFGGGTKAEIK
|||.||||||||||||||||||||||||||||||||||||||||||||||||||||.||||.||||||||||||.|||||||||||.|||||||||||||||.|||
EIVLTQSPATLSLSPGERATLSCRASQSVSSYLAWYQQKPGQAPRLLIYDASNRATGIPDRFSGGGSGTDFTLTISRLEPEDFAVYYCQQYGSTPLTFGGGTKVEIK
                   ^^^^^^          ^^^                ^^^^^^^^^
```

Best Alignment of Necitumumab CDR-H3 chain to a sequence from OAS

```
QVQLQESGPGLVKPSQTLSLTCTVSGGSISSGDYYWSWIRQPPGKGLEWIGYIYYSGSTDYNPSLKSRVTMSVDTSKNQFSLKVNSVTAADTAVYYCARVSIFGVGTFDYWGQGTLVTVSS
............||.|||||||||||||....||||||||||||||||||||||||||.||||||||||.||||||||||||..||||||||||||||||||||.||||||||||||||||
------------PSETLSLTCTVSGGSI--SSYYWSWIRQPPGKGLEWIGYIYYSGSTNYNPSLKSRVTISVDTSKNQFSLKLSSVTAADTAVYYCARVSIFGVVTFDYWGQGTLVTVSS
                   ^^^^^^^^^^        ^^^^^^^                  ^^^^^^^^^^^^^
```

Therapeutic : Nimotuzumab

Best Alignment of Nimotuzumab heavy chain to a sequence from OAS

```
QVQLQQSGAEVKKPGSSVKVSCKASGYTFTNYYIYWVRQAPGQGLEWIGGINPTSGGSNFNEKFKTRVTITADESSTTAYMELSSLRSEDTAFYFCTRQGLWFD----SDGRGFDFWGQGTTVTVSS
||||.||||||||||.||||||||||||||.|||.|||||||||||||.|.|||.|||.|...||..|||||||||..||||||||||||||||.|.|.|...|......|..|.||.|||||.|||||
QVQLVQSGAEVKKPGASVKVSCKASGYTFTGYYIHWVRQAPGQGLEWMGRINPNSGGTNYAQKFQGRVTITADESTSTAYMELSSLRSEDTAVYYCARGRCWGRYYDSSGYRAFDIWGQGTMVTVSS
                   ^^^^^^^^        ^^^^^^^^                  ^^^^^^^^^^^^^^^^^^^^
```

Best Alignment of Nimotuzumab light chain to a sequence from OAS

```
DIQMTQSPSSLSASVGDRVTITCRSSQNIVHSNGNTYLDWYQQTPGKAPKLLIYKVSNRFSGVPSRFSGSGSGTDFTFTISSLQPEDIATYYCFQYSH---VPWTFGQGTKLQIT
||||||||||||||||||||||||.||||......|||.||||.||||||||||..||..|||||||||||||||||||||||||||||||||||.||......|.||||||.|.|.
DIQMTQSPSSLSASVGDRVTITCRASQNI-----ATYLNWYQQKPGKAPKLLIYVASNLHSGVPSRFSGSGSGTDFTFTISSLQPEDIATYYCQQYDNLPRPPITFGQGTRLEIK
                   ^^^^^^^^^^^          ^^^                ^^^^^^^^^^^^
```

Best Alignment of Nimotuzumab heavy chain CDRs to a sequence from OAS

```
QVQLQQSGAEVKKPGSSVKVSCKASGYTFTNYYIYWVRQAPGQGLEWIGGINPTSGGSNFNEKFKTRVTITADESSTTAYMELSSLRSEDTAFYFCTRQGLWFDSDGRGFDFWGQGTTVTVSS
||||.||||||||||.||||||||||||||.||..||||||||||||.|.|||.|||.|...||..|||.|.|.|..|||||||.|||.|||.|.|.|....|||.|||||.|||||.|||||
QVQLVQSGAEVKKPGASVKVSCKASGYTFTGYYMHWVRQAPGQGLEWMGWINPNSGGTNYAQKFQGRVTMTRDTSISTAYMELSRLRSDDTAVYYCARDTPPFDSSGRGFDYWGQGTLVTVSS
                   ^^^^^^^^        ^^^^^^^^                  ^^^^^^^^^^^^^^^^
```

Best Alignment of Nimotuzumab light CDRs chain to a sequence from OAS

```
DIQMTQSPSSLSASVGDRVTITCRSSQNIVHSNGNTYLDWYQQTPGKAPKLLIYKVSNRFSGVPSRFSGSGSGTDFTFTISSLQPEDIATYYCFQYSHVPWTFGQGTKLQIT
|..|||.|.||..|.||...|.|||||||||||||||||.||.|.||..||||||||||||||||.||||||||||||..||....||...|||||.||||||||.||||.|.
DVLMTQTPLSLPVSLGDQASISCRSSQNIVHSNGNTYLEWYLQKPGQSPKLLIYKVSNRFSGVPDRFSGSGSGTDFTLKISRVEAEDLGVYYCFQGSHVPWTFGGGTKLEIK
                   ^^^^^^^^^^^          ^^^                ^^^^^^^^^
```

Best Alignment of Nimotuzumab CDR-H3 chain to a sequence from OAS

```
QVQLQQSGAEVKKPGSSVKVSCKASGYTFTNYYIYWVRQAPGQGLEWIGGINPTSGGSNFNEKFKTRVTITADESSTTAYMELSSLRSEDTAFYFCTRQGLWFDSDGRGFDFWGQGTTVTVSS
.|||||.||...||.......|..||..|.|....|.||.||.||||||.......|.........|.||..|.........|.|....|||.|.|||.|||...|.||||.|||||.|||||
-VQLQQWGAGLLKPSETLSLTCAVSGGSFSNSLWTWIRQPPGKGLEWIGDVSLD-RGTKNVPSLQSRITISMDRTKNQFSLRLKSVTAADTATYYCTRLGLWGYYDARGFDYWGQGTLVTVSS
                   ^^^^^^^^        ^^^^^^^^                  ^^^^^^^^^^^^^^^^
```

Therapeutic : Nivolumab

Best Alignment of Nivolumab heavy chain to a sequence from OAS

```
QVQLVESGGGVVQPGRSLRLDCKASGITFSNSGMHWVRQAPGKGLEWVAVIWYDGSKRYYADSVKGRFTISRDNSKNTLFLQMNSLRAEDTAVYYCATN-------DDYWGQGTLVTVSS
|||||||||||||||||||.|.|||.||.|.||||||||||||||||||||||||||.||||||.|||||||||||||||||||||||||||||||||||.........||||||||||||
QVQLVESGGGVVQPGRSLRLSCVASGFTFRNYGMHWVRQAPGKGLEWVAVIWYDGSKKYYADSVGGRFTISRDNSKNTLFLQMNSLRAEDTAVYYCATVSKSSSWYTDYWGQGTLVTVSS
                   ^^^^^^^^        ^^^^^^^^                  ^^^^^^^^^^^^^
```

Best Alignment of Nivolumab light chain to a sequence from OAS

```
EIVLTQSPATLSLSPGERATLSCRASQSVSSYLAWYQQKPGQAPRLLIYDASNRATGIPARFSGSGSGTDFTLTISSLEPEDFAVYYCQQSSNWPRTFGQGTKVEIK
|||||||||||||||||||||||||||||||||||||||||||||||||||||||||||||||||||||||||||||||||||||||||||||||||||||||||||
EIVLTQSPATLSLSPGERATLSCRASQSVSSYLAWYQQKPGQAPRLLIYDASNRATGIPARFSGSGSGTDFTLTISSLEPEDFAVYYCQQSSNWPRTFGQGTKVEIK
                   ^^^^^^          ^^^                ^^^^^^^^^
```

Best Alignment of Nivolumab heavy chain CDRs to a sequence from OAS

```
QVQLVESGGGVVQPGRSLRLDCKASGITFSNSGMHWVRQAPGKGLEWVAVIWYDGSKRYYADSVKGRFTISRDNSKNTLFLQMNSLRAEDTAVYYCATNDDYWGQGTLVTVSS
...............||||.|.||||||||.|||||||||||||||||||||||||.||||||||||||||||||||||||.||||.||||||||||||..|.||||||||||||||
--------------GSLRLSCAASGITFSNYGMHWVRQAPGKGLEWVAVIWYDGSNRYYADSVKGRFTISRDNSKNTLYLQMNGLRAEDTAVYYCTSNFDYWGQGTLVTVSS
                   ^^^^^^^^        ^^^^^^^^                  ^^^^^^
```

Best Alignment of Nivolumab light CDRs chain to a sequence from OAS

```
EIVLTQSPATLSLSPGERATLSCRASQSVSSYLAWYQQKPGQAPRLLIYDASNRATGIPARFSGSGSGTDFTLTISSLEPEDFAVYYCQQSSNWPRTFGQGTKVEIK
||||.||||||||||||||||||||||||||||||||||||||||||||||||||||||.|||||||.||||||||||||||||||||||||||||||||||.||||
EIVLAQSPATLSLSPGERATLSCRASQSVSSYLAWYQQKPGQAPRLLIYDASNRATGIPARVSGSGSGTEFTLTISSLEPEDFAVYYCQQSSNWPRTFGQGTRVEIK
                   ^^^^^^          ^^^                ^^^^^^^^^
```

Best Alignment of Nivolumab CDR-H3 chain to a sequence from OAS

```
QVQLVESGGGVVQPGRSLRLDCKASGITFSNSGMHWVRQAPGKGLEWVAVIWYDGSKRYYADSVKGRFTISRDNSKNTLFLQMNSLRAEDTAVYYCATNDDYWGQGTLVTVSS
.|||..||...|.||.|....|||||.||....|.||.|..||.|||...|........|....||..|...|.|..|......||..||.||||||||||||||||..||||
```

-VQLQQSGPELVKPGASVKISCKASGYTFTDYYMNWVKQSHGKSLEWIGDINPNNGGTSYNQKFKGKATLTVDKSSSTAYMELRSLTSEDSAVYYCATNDDYWGQGTTLTVSS
                   ^^^^^^^^            ^^^^^^^^                              ^^^^^^

Therapeutic : Obiltoxaximab

Best Alignment of Obiltoxaximab heavy chain to a sequence from OAS
QVQLQQSGPELKKPGASVKVSCKDSGYAFSSSWMNWVRQAPGQGLEWIGRIYPGDGDTNYNGKFQGRVTITADKSSSTAYMELSSLRSEDTAVYFCARSGLL---RYAMDYWGQGTLVTVSS
|||||||||||.|||||||.|||.|||||||||||||.|.||.|||||||||||||||||||||.|..|.|||||||||||.||||.|||.|||||||||||...|||||||||||.|||||
QVQLQQSGPELVKPGASVKISCKASGYAFSSSWMNWVKQRPGKGLEWIGRIYPGDGDTNYNGKFKGKATLTADKSSSTAYMQLSSLTSEDSAVYFCARSGLLLRSRYAMDYWGQGTSVTVSS
                   ^^^^^^^^            ^^^^^^^^                              ^^^^^^^^^^^^^^^

Best Alignment of Obiltoxaximab light chain to a sequence from OAS
DIQMTQSPSSLSASVGDRVTITCRASQDIRNYLNWYQQKPGKAVKLLIYYTSRLLPGVPSRFSGSGSGTDYSLTISSQEQEDIGTYFCQQGNTLPWTFGQGTKVEIR
||||||..||||||.||||||.|||||||.||||||||||..||||||||||||..||||||||||||||||||||..|||||.|||||||||||||||.|||.||.
DIQMTQITSSLSASLGDRVTISCRASQDISNYLNWYQQKPDGAVKLLIYYTSRLHSGVPSRFSGSGSGTDYSLTISNLEQEDIATYFCQQGNTLPWTFGGGTKLEIK
                    ^^^^^^          ^^^                        ^^^^^^^^^

Best Alignment of Obiltoxaximab heavy chain CDRs to a sequence from OAS
QVQLQQSGPELKKPGASVKVSCKDSGYAFSSSWMNWVRQAPGQGLEWIGRIYPGDGDTNYNGKFQGRVTITADKSSSTAYMELSSLRSEDTAVYFCARSGLLRYAMDYWGQGTLVTVSS
||||..|||||.|||||||.|||.||||||||||.||.|.||||||||||||||||||||||||.|..|.||||||.||||.||||.|||.||||.|||||||||||||||||.|||||
QVQLKESGPELVKPGASVKISCKASGYAFSSSWMHWVKQRPGQGLEWIGRIYPGDGDTNYNGKFKGKATLTADKSSNTAYMPLSSLTSEDSAVYFFARSGLLRYAMDYWGQGTSVTVSS
                   ^^^^^^^^            ^^^^^^^^                              ^^^^^^^^^^^^

Best Alignment of Obiltoxaximab light CDRs chain to a sequence from OAS
DIQMTQSPSSLSASVGDRVTITCRASQDIRNYLNWYQQKPGKAVKLLIYYTSRLLPGVPSRFSGSGSGTDYSLTISSQEQEDIGTYFCQQGNTLPWTFGQGTKVEIR
|||.||..||||||.||||||.||||||||||||||||||...|||||||||||..||||||||||||||||||||..|||||.||||||||||||||..|||.|..
DIQLTQTTSSLSASLGDRVTISCRASQDIRNYLNWYQQKPDGTVKLLIYYTSRLHSGVPSRFSGSGSGTDYSLTISNLEQEDIATYFCQQGNTLPWTFSGGTKLEVK
                    ^^^^^^          ^^^                        ^^^^^^^^^

Best Alignment of Obiltoxaximab CDR-H3 chain to a sequence from OAS
QVQLQQSGPELKKPGASVKVSCKDSGYAFSSSWMNWVRQAPGQGLEWIGRIYPGDGDTNYNGKFQGRVTITADKSSSTAYMELSSLRSEDTAVYFCARSGLLRYAMDYWGQGTLVTVSS
.|||..||..|..||.|.|.||..||..|||..|.||||||..||||......|.....|.....||.||..|....|......||||||||.|.||||||||||||||||||.|||||
EVQLVESGGGLVQPGGSRKLSCAASGFTFSSFGMHWVRQAPEKGLEWVAYVSSGSSTIYYADTVKGRFTISRDNPKNTLFLQMTSLRSEDTAMYYCARSGLLRYAMDYWGQGTSVTVSS
                   ^^^^^^^^            ^^^^^^^^                              ^^^^^^^^^^^^

Therapeutic : Obinutuzumab

Best Alignment of Obinutuzumab heavy chain to a sequence from OAS
QVQLVQSGAEVKKPGSSVKVSCKASGYAFSYSWINWVRQAPGQGLEWMGRIFPGDGDTDYNGKFKGRVTITADKSTSTAYMELSSLRSEDTAVYYCARNVFD--GYWLVYWGQGTLVTVSS
||||||||||||||||||||||||||..|....||||||||||||||||||.|..|.|.|..||.|||||||||||||||||||||||||||||||||..||..||...||||||||||||
QVQLVQSGAEVKKPGSSVKVSCKASGGTF-RYAINWVRQAPGQGLEWMGRIIPIFGTTNYAQKFQGRVTITADKSTSTAYMELSSLRSEDTAVYYCARGPFDNSGYYSFYWGQGTLVTVSS
                   ^^^^^^^^            ^^^^^^^^                              ^^^^^^^^^^^^^^

Best Alignment of Obinutuzumab light chain to a sequence from OAS
DIVMTQTPLSLPVTPGEPASISCRSSKSLLHSNGITYLYWYLQKPGQSPQLLIYQMSNLVSGVPDRFSGSGSGTDFTLKISRVEAEDVGVYYCAQNLELPYTFGGGTKVEIK
||||||...|.|||.|..|||||||||||||||||||||||||||||||||||||||||.||||||||||||||||||.|||||||||||||||||||||||||||||.|||
DIVMTQAAFSNPVTLGTSASISCRSSKSLLHSNGITYLYWYLQKPGQSPQLLIYQMSNLASGVPDRFSGSGSGTDFTLRISRVEAEDVGVYYCAQNLELPYTFGGGTKLEIK
                    ^^^^^^^^^^^          ^^^                        ^^^^^^^^^

Best Alignment of Obinutuzumab heavy chain CDRs to a sequence from OAS
QVQLVQSGAEVKKPGSSVKVSCKASGYAFSYSWINWVRQAPGQGLEWMGRIFPGDGDTDYNGKFKGRVTITADKSTSTAYMELSSLRSEDTAVYYCARNVFDGYWLVYWGQGTLVTVSS
||||.|||.|..|||.|||.||||||||||.||.|||.|.|||||||.|||.||||||.|||||||..|.|||||.|||||.||||.|.|.|||.|||.|.||.|.||||||||||||.
QVQLQQSGPELVKPGASVKISCKASGYAFSSSWMNWVKQRPGQGLEWIGRIYPGDGDTNYNGKFKGKATLTADKSSSTAYMHLSSLTSVDSAVYFCARCVYDGSWFVYWGQGTLVTVSA
                   ^^^^^^^^            ^^^^^^^^                              ^^^^^^^^^^^^

Best Alignment of Obinutuzumab light CDRs chain to a sequence from OAS
DIVMTQTPLSLPVTPGEPASISCRSSKSLLHSNGITYLYWYLQKPGQSPQLLIYQMSNLVSGVPDRFSGSGSGTDFTLKISRVEAEDVGVYYCAQNLELPYTFGGGTKVEIK
.|.||......||.....|||||.|||||||||||||||||||||||||||||||||||.||||||||.|||||||||.|||||||||||||||||||||||||||||.|||
-IAMTPAACCSPVSLLPSASISCGSSKSLLHSNGITYLYWYLQKPGQSPQLLIYQMSNLASGVPDRFSSSGSGTDFTLRISRVEAEDVGVYYCAQNLELPYTFGGGTKLEIK
                    ^^^^^^^^^^^          ^^^                        ^^^^^^^^^

Best Alignment of Obinutuzumab CDR-H3 chain to a sequence from OAS
QVQLVQSGAEVKKPGSSVKVSCKASGYAFSYSWINWVRQAPGQGLEWMGRIFPGDGDTDYNGKFKGRVTITADKSTSTAYMELSSLRSEDTAVYYCARNVFDGYWLVYWGQGTLVTVSS
.|||.|||....|||.|||.|||||||.|.....|||.|..|..|||.|.|.|..|.|..|.||||..|.|.|||.|||||||.||.|||.||.||||||||||.|.||||||..||||
EVQLQQSGPVLVKPGASVKMSCKASGYTFTDYYMNWVKQSHGKSLEWIGVINPYNGGTSSNQKFKGKATLTVDKSSSTAYMELNSLKSEDSAVFYCARNVFDGYYLDYWGQGTTLTVSS
                   ^^^^^^^^            ^^^^^^^^                              ^^^^^^^^^^^^

Therapeutic : Ocrelizumab

Best Alignment of Ocrelizumab heavy chain to a sequence from OAS
EVQLVESGGGLVQPGGSLRLSCAASGYTFTSYNMHWVRQAPGKGLEWVGAIYPGNGDTSYNQKFKGRFTISVDKSKNTLYLQMNSLRAEDTAVYYCARVVYYSN--SYWYFDVWGQGTLVTVSS
||||||||||||||||||||||||||.||.||.|||||||||||||||..||.|.|.|.|....|||||||.|.|||||||||||||||||||||||||||..|.....|||.|||||||||||
EVQLVESGGGLVQPGGSLRLSCAASGFTFSSYAMHWVRQAPGKGLEWVSVIYSG-GSTYYADSVKGRFTISRDNSKNTLYLQMNSLRAEDTAVYYCARVVYGGNVFPSYYFDYWGQGTLVTVSS
                   ^^^^^^^^            ^^^^^^^^                              ^^^^^^^^^^^^^^^^^

Best Alignment of Ocrelizumab light chain to a sequence from OAS
DIQMTQSPSSLSASVGDRVTITCRASSSV-SYMHWYQQKPGKAPKPLIYAPSNLASGVPSRFSGSGSGTDFTLTISSLQPEDFATYYCQQWSFNPPTFGQGTKVEIK
||||||||||||||||||||.|||||.|..||..|||||||||||.||||.|||.|||||||||||||||||||||||||||||||||||...||||||||||||||
DIQMTQSPSSLSASVGDRVTFTCRASQSIISYLNWYQQKPGKAPKLLIYAASNLQSGVPSRFSGSGSGTDFTLTISSLQPEDFATYYCQQSYSNPPTFGQGTKVEIK
                    ^^^^^^          ^^^                        ^^^^^^^^^

Best Alignment of Ocrelizumab heavy chain CDRs to a sequence from OAS
EVQLVESGGGLVQPGGSLRLSCAASGYTFTSYNMHWVRQAPGKGLEWVGAIYPGNGDTSYNQKFKGRFTISVDKSKNTLYLQMNSLRAEDTAVYYCARVVYYSNSYWYFDVWGQGTLVTVSS
...|..||..||.||.|...||.||||||||||||||.|.|..||||.||||||||||||||||||..|..||||..|.|.|..||..||.|||.|||..|||||||||||||.||.|||||
---LQQSGAELVRPGASVKMSCKASGYTFTSYNMHWVKQTPRQGLEWIGAIYPGNGDTSYNQKFKGKATLTVDKSSSTAYMQLSSLTSEDSAVYFCARSDYYSNSYWYFDVWGTGTTVTVSS
                   ^^^^^^^^            ^^^^^^^^                              ^^^^^^^^^^^^^^^

Best Alignment of Ocrelizumab light CDRs chain to a sequence from OAS
DIQMTQSPSSLSASVGDRVTITCRASSSVSYMHWYQQKPGKAPKPLIYAPSNLASGVPSRFSGSGSGTDFTLTISSLQPEDFATYYCQQWSFNPPTFGQGTKVEIK
.|..||||...|||.|..||.||.||||||||||||||.|..|...|||||..|.|||.|||||||||..||||||...||.|||||||||.||||||.|||.|.|
QIVLTQSPAIMSASPGEKVTMTCSASSSVSYMHWYQQKSGPSPTSWIYAPSQRACGVPARFSGSGSGTSYSLTISSMEAEDAATYYCQQWSSNPPTFGAGTKLELK
                    ^^^^^          ^^^                        ^^^^^^^^^

Best Alignment of Ocrelizumab CDR-H3 chain to a sequence from OAS
EVQLVESGGGLVQPGGSLRLSCAASGYTFTSYNMHWVRQAPGKGLEWVGAIYPGNGDTSYNQKFKGRFTISVDKSKNTLYLQMNSLRAEDTAVYYCARVVYYSNSYWYFDVWGQGTLVTVSS
.|||..||..||.||.|...||.||||.|.|.|..|.||.|.||.||||.|.||||.|||.||.|||...|...|||..|.|.|..||..||.|||.|||.||||||||||||||.||.||||.
QVQLQQSGPELVKPGASVKISCKASGYAFSSSWMNWVKQRPGQGLEWIGRIYPGDGDTNYNGKFKCKATLTADKSSSTAYMQRSSLTSEDSAVYFCARSVYYSNSYWYFDVWGTGTTVTVSP
                   ^^^^^^^^            ^^^^^^^^                              ^^^^^^^^^^^^^^^

Therapeutic : Ofatumumab

Best Alignment of Ofatumumab heavy chain to a sequence from OAS
EVQLVESGGGLVQPGRSLRLSCAASGFTFNDYAMHWVRQAPGKGLEWVSTISWNSGSIGYADSVKGRFTISRDNAKKSLYLQMNSLRAEDTALYYCAKDIQYGN-------YYYGMDVWGQGTTVTVSS
|||||||||||||||||||||||||||||.|||||||||||||||||||.||||||||||||||||||||||||||.|||||||||||||||||||||||.|.........||||||||||||||||||
EVQLVESGGGLVQPGRSLRLSCAASGFTFDDYAMHWVRQAPGKGLEWVSGISWNSGSIGYADSVKGRFTISRDNAKNSLYLQMNSLRAEDTALYYCAKDIGYCSSTSCPRPYYYGMDVWGQGTTVTVSS
                   ^^^^^^^^            ^^^^^^^^                              ^^^^^^^^^^^^^^^^^^^^^^

Best Alignment of Ofatumumab light chain to a sequence from OAS

```
EIVLTQSPATLSLSPGERATLSCRASQSVSSYLAWYQQKPGQAPRLLIYDASNRATGIPARFSGSGSGTDFTLTISSLEPEDFAVYYCQQRSNWPITFGQGTRLEIK
|||||||||||||||||||||||||||||||||||||||||||||||||||||||||||||||||||||||||||||||||||||||||||||||||||||||||||
EIVLTQSPATLSLSPGERATLSCRASQSVSSYLAWYQQKPGQAPRLLIYDASNRATGIPARFSGSGSGTDFTLTISSLEPEDFAVYYCQQRSNWPITFGQGTRLEIK
                         ^^^^^^           ^^^                       ^^^^^^^^^
```

Best Alignment of Ofatumumab heavy chain CDRs to a sequence from OAS

```
EVQLVESGGGLVQPGRSLRLSCAASGFTFNDYAMHWVRQAPGKGLEWVSTISWNSGSIGYADSVKGRFTISRDNAKKSLYLQMNSLRAEDTALYYCAKDIQYGNYYYGMDVWGQGTTVTVSS
................|||||||||||||.||||||||||||||||||||.||||||||||||||||||||||||||||.|||||||||||||.||||||||||||.|.|.||||||||||||||||||
----------------SLRLSCAASGFTFDDYAMHWVRQAPGKGLEWVSGISWNSGSIGYADSVKGRFTISRDNAKNSLYLQMNSLRAEGTALYYCAKDIQAGYYYYGMDVWGQGTTVTVSS
                        ^^^^^^^^          ^^^^^^^^                        ^^^^^^^^^^^^^^^
```

Best Alignment of Ofatumumab light CDRs chain to a sequence from OAS

```
EIVLTQSPATLSLSPGERATLSCRASQSVSSYLAWYQQKPGQAPRLLIYDASNRATGIPARFSGSGSGTDFTLTISSLEPEDFAVYYCQQRSNWPITFGQGTRLEIK
|||||||||||||||||||||||||||||||||||||||||||||||||||||||||||||||||||||||||||||||||||||||||||||||||||||||||||
EIVLTQSPATLSLSPGERATLSCRASQSVSSYLAWYQQKPGQAPRLLIYDASNRATGIPARFSGSGSGTDFTLTISSLEPEDFAVYYCQQRSNWPITFGQGTRLEIK
                         ^^^^^^           ^^^                       ^^^^^^^^^
```

Best Alignment of Ofatumumab CDR-H3 chain to a sequence from OAS

```
EVQLVESGGGLVQPGRSLRLSCAASGFTFNDYAMHWVRQAPGKGLEWVSTISWNSGSIGYADSVKGRFTISRDNAKKSLYLQMNSLRAEDTALYYCAKDIQYGNYYYGMDVWGQGTTVTVSS
..............|.|||||||||||||..|.||||||||||||||.|..||.......|.||||||||||||||.|..|||||||||||||||.|||||||.||||||||||||||.|||||||||
--------------GGSLRLSCAASGFTFSSYGMHWVRQAPGKGLESVADISYDGSNKYYVDSVKGRFTISRDNSKNTLYLQMNSLRAEDTAVYYCAKDIAYGNYYYGMDVWGKGTTVTVSS
                        ^^^^^^^^          ^^^^^^^^                        ^^^^^^^^^^^^^^^
```

Therapeutic : Olaratumab

Best Alignment of Olaratumab heavy chain to a sequence from OAS

```
QLQLQESGPGLVKPSETLSLTCTVSGGSINSSSYYWGWLRQSPGKGLEWIGSFFYTGSTYYNPSLRSRLTISVDTSKNQFSLMLSSVTAADTAVYYCARQSTYYYGSGNYYGWFDRWDQGTLVTVSS
|||||||||||||||||||||||||||||||.||||||||.||.||||||||||||..|.||||||||||.||.||||||||||||||||.||||||||||||||||||..||||||||.|||.||.|.||||||||||
QLQLQESGPGLVKPSETLSLTCTVSGGSISSSSYYWGWIRQPPGKGLEWIGSIYYSGSTYYNPSLKSRVTISVDTSKNQFSLKLSSVTAADTAVYYCARGRTYYYGSGSYYGLFDYWGQGTLVTVSS
                        ^^^^^^^^^^          ^^^^^^^                        ^^^^^^^^^^^^^^^^^^^
```

Best Alignment of Olaratumab light chain to a sequence from OAS

```
EIVLTQSPATLSLSPGERATLSCRASQSVSSYLAWYQQKPGQAPRLLIYDASNRATGIPARFSGSGSGTDFTLTISSLEPEDFAVYYCQQRSNWPPAFGQGTKVEIK
|||||||||||||||||||||||||||||||||||||||||||||||||||||||||||||||||||||||||||||||||||||||||||||||||||||||||||
EIVLTQSPATLSLSPGERATLSCRASQSVSSYLAWYQQKPGQAPRLLIYDASNRATGIPARFSGSGSGTDFTLTISSLEPEDFAVYYCQQRSNWPPAFGQGTKVEIK
                         ^^^^^^           ^^^                       ^^^^^^^^^
```

Best Alignment of Olaratumab heavy chain CDRs to a sequence from OAS

```
QLQLQESGPGLVKPSETLSLTCTVSGGSINSSSYYWGWLRQSPGKGLEWIGSFFYTGSTYYNPSLRSRLTISVDTSKNQFSLMLSSVTAADTAVYYCARQSTYYYGSGNYYGWFDRWDQGTLVTVSS
.............||||||||.||||||.|||||||||.||.|||||||||||...|.|||||||||||.||.||||||||||||||||.|||||||||||||||||.|.|||||||.|||||||.|.||||||||||
--------------SETLSLTCAVSGGSISSSSYYWGWIRQPPGKGLEWIGGIYYSGSTYYNPSLKSRVTISVDTSKNQFSLKLSSVTAADTAVYYCARHSPYYYGSGSYYGWFDPWGQGTLVTVSS
                        ^^^^^^^^^^          ^^^^^^^                        ^^^^^^^^^^^^^^^^^^^
```

Best Alignment of Olaratumab light CDRs chain to a sequence from OAS

```
EIVLTQSPATLSLSPGERATLSCRASQSVSSYLAWYQQKPGQAPRLLIYDASNRATGIPARFSGSGSGTDFTLTISSLEPEDFAVYYCQQRSNWPPAFGQGTKVEIK
||||||||||||||||||||||||||||||||||||||||||||||||||||||||||||||||||||||||||||||||||||||||||||||||||.||||||||
EIVLTQSPATLSLSPGERATLSCRASQSVSSYLAWYQQKPGQAPRLLIYDASNRATGIPARFSGSGSGTDFTLTISSLEPEDFAVYYCQQRSNWPPAFGGGTKVEIK
                         ^^^^^^           ^^^                       ^^^^^^^^^
```

Best Alignment of Olaratumab CDR-H3 chain to a sequence from OAS

```
QLQLQESGPGLVKPSETLSLTCTVSGGSINSSSYYWGWLRQSPGKGLEWIGSFFYTGSTYYNPSLRSRLTISVDTSKNQFSLMLSSVTAADTAVYYCARQSTYYYGSGNYYGWFDRWDQGTLVTVSS
............||||||||||||||||..|||||.|.||.||||||||||...|.|||.|||||.||.||||||||||||||.||||||||||||||||.||||||||.|||.||.|.|||.|||||
-------------PSETLSLTCTVSGGSI--SSYYWSWIRQPPGKGLEWIGYIYYSGSTNYNPSLKSRVTISVDTSKNQFSLKLSSVTAADTAVYYCARDSTYYYGSGSYYGAFDIWGQGTMVTVSS
                        ^^^^^^^^^^          ^^^^^^^                        ^^^^^^^^^^^^^^^^^^^
```

Therapeutic : Olendalizumab

Best Alignment of Olendalizumab heavy chain to a sequence from OAS

```
QVQLVQSGAEVKKPGASVKVSCKASGYTFTDYSMDWVRQAPGQGLEWMGAIHLNTGYTNYNQKFKGRVTMTRDTSTSTVYMELSSLRSEDTAVYYCARGFYDGYSPMDYWGQGTTVTVSS
||||||||||||||||||||||||||||||.|.|.|||||||||||||||.|..|.|.|||.|||.|||||||||||||||||||||||||||||||||||.|.||||..||||||||.|||||
QVQLVQSGAEVKKPGASVKVSCKASGYTFTGYYMHWVRQAPGQGLEWMGWINPNSGGTNYAQKFQGRVTMTRDTSTSTVYMELSSLRSEDTAVYYCARDFLDGYSGFDYWGQGTLVTVSS
                        ^^^^^^^^          ^^^^^^^^                        ^^^^^^^^^^^^^
```

Best Alignment of Olendalizumab light chain to a sequence from OAS

```
DIQMTQSPSSLSASVGDRVTITCRASESVDSYGNSFMHWYQQKPGKAPKLLIYRASNLESGVPSRFSGSGSGTDFTLTISSLQPEDFATYYCQQSNEDPYTFGGGTKVEIK
||||||||||||||||||||||||||||.||....||..|||||||||||||||.|.||||.||||||||||||||||||||||||||||||||||||||||...|||||.|||||||
DIQMTQSPSSLSASVGDRVTITCRASQSV----NSYLHWYQQKPGKAPKLLMYAASNLQSGVPSRFSGSGSGTDFTLTISSLQPEDFATYYCQQSYSTPYTFGQGTKVEIK
                         ^^^^^^^^^^          ^^^                      ^^^^^^^^^
```

Best Alignment of Olendalizumab heavy chain CDRs to a sequence from OAS

```
QVQLVQSGAEVKKPGASVKVSCKASGYTFTDYSMDWVRQAPGQGLEWMGAIHLNTGYTNYNQKFKGRVTMTRDTSTSTVYMELSSLRSEDTAVYYCARGFYDGYSPMDYWGQGTTVTVSS
.|||.|.|||..|||||||.||||||||||.|.|.||.|.||||||||.|.||.|.|.||||.|||...|.|.|.|.||.||.||||.|||.||||||||.|||||.||||||||.|||||
-VQLQQPGAELVKPGASVKLSCKASGYTFTSYWMHWVKQRPGQGLEWIGMIHPNSGSTNYNEKFKSKATLTVDKSSSTAYMQLSSLTSEDSAVYYCARGDYDGYSFMDYWGQGTSVTVSS
                        ^^^^^^^^          ^^^^^^^^                        ^^^^^^^^^^^^^
```

Best Alignment of Olendalizumab light CDRs chain to a sequence from OAS

```
DIQMTQSPSSLSASVGDRVTITCRASESVDSYGNSFMHWYQQKPGKAPKLLIYRASNLESGVPSRFSGSGSGTDFTLTISSLQPEDFATYYCQQSNEDPYTFGGGTKVEIK
||..||||.||..|.|.|.||.||||||||||||||||||||||||||..||||||||||||||.|.|||||||.|||||||......|.|||||||||||||||||||||.|||
DIVLTQSPVSLAVSLGQRATISCRASESVDSYGNSFMHWYQQKPGQPPKLLIYRASNLESGIPARFSGSGSRTDFTLTINPVEADDVATYYCQQSNEDPYTFGGGTKLEIK
                         ^^^^^^^^^^          ^^^                      ^^^^^^^^^
```

Best Alignment of Olendalizumab CDR-H3 chain to a sequence from OAS

```
QVQLVQSGAEVKKPGASVKVSCKASGYTFTDYSMDWVRQAPGQGLEWMGAIHLNTGYTNYNQKFKGRVTMTRDTSTSTVYMELSSLRSEDTAVYYCARGFYDGYSPMDYWGQGTTVTVSS
.|||.|||....|||||||.|||||||||||||.|.||.|..||.|||.|.|....|.|.||.||||..|.|.|.|.||.||||.||.|||.|||||||||||||||.||||||||.|||||
EVQLQQSGPVLVKPGASVKMSCKASGYTFTDYYMNWVKQSHGQSLEWIGVINPYNGVTSYNPKFKGKATLTVDKSSSTAYMELNSLTSEDSAVYYCARGFYDGYSFMDYWGQGTSVTVSS
                        ^^^^^^^^          ^^^^^^^^                        ^^^^^^^^^^^^^
```

Therapeutic : Olokizumab

Best Alignment of Olokizumab heavy chain to a sequence from OAS

```
EVQLVESGGGLVQPGGSLRLSCAASGFNFNDYFMNWVRQAPGKGLEWVAQMRNKNYQYGTYYAESLEGRFTISRDDSKNSLYLQMNSLKTEDTAVYYCARESYY---GFTSYWGQGTLVTVSS
|||||||||||||||||||||||||||.|..|.||||||||||||||||...|||...|.|.||.|..||||||||||||||||||||||||||||||||||..||...|...|||||||||||
EVQLVESGGGLVQPGGSLRLSCAASGFTFSSYEMNWVRQAPGKGLEWVGRTRNKANSYTTEYAASVKGRFTISRDDSKNSLYLQMNSLKTEDTAVYYCARVDYYDSSGWYDYWGQGTLVTVSS
                        ^^^^^^^^          ^^^^^^^^^^                        ^^^^^^^^^^^^^^
```

Best Alignment of Olokizumab light chain to a sequence from OAS

```
DIQMTQSPSSLSASVGDRVTITCQASQDIGISLSWYQQKPGKAPKLLIYNANNLADGVPSRFSGSGSGTDFTLTISSLQPEDFATYYCLQHNSAPYTFGQGTKLEIK
||||||||||||||||||||||||||||||...|.||||||||||||||||.|.||..|||||||||||||.||||||||||||||||||||||||.||||||||||||||
DIQMTQSPSSLSASVGDRVTITCQASQDISNNLHWYQQKPGKAPKLLIYDASNLETGVPSRFSGSGSGTEFTLTISSLQPEDFATYYCLQHNSYPYTFGQGTKLEIK
                         ^^^^^^           ^^^                       ^^^^^^^^^
```

Best Alignment of Olokizumab heavy chain CDRs to a sequence from OAS

```
EVQLVESGGGLVQPGGSLRLSCAASGFNFNDYFMNWVRQAPGKGLEWVAQMRNKNYQYGTYYAESLEGRFTISRDDSKNSLYLQMNSLKTEDTAVYYCARESYYGFTSYWGQGTLVTVSS
||.|.|||||||||||||||||.|||.|.||.|.||||.|||.|||....|||...|.|.|..|..||||||||.|...||||||.|..||.|.||||||||||..||||||||||||.
```

EVKLMESGGGLVQPGGSLRLSCATSGFTFTDYYMSWVRQPPGKALEWLGFIRNKANGYTTEYSASVKGRFTISRDNSQSILYLQMNTLRAEDSATYYCARESYYGSSSYWGQGTLVTVSA
^^^^^^^^ ^^^^^^^^^^ ^^^^^^^^^^^

Best Alignment of Olokizumab light CDRs chain to a sequence from OAS
DIQMTQSPSSLSASVGDRVTITCQASQDIGISLSWYQQKPGKAPKLLIYNANNLADGVPSRFSGSGSGTDFTLTISSLQPEDFATYYCLQHNSAPYTFGQGTKLEIK
||||||||||||||||.|||||.|||||.|||.|.|||||||||||||.|.....||||||||.||||||||.||||||||||.||||||||.||.||||||.|||
DIQMTQSPSSLSASVGDTVTITCRASQDISISLNWFQQKPGKAPKLLIYAASSVESGVPSRFSGGGSGTDFTLSISSLQPEDFAVYYCLQHNSYPYSFGQGTKVEIK
^^^^^^ ^^^ ^^^^^^^^^

Best Alignment of Olokizumab CDR-H3 chain to a sequence from OAS
EVQLVESGGGLVQPGGSLRLSCAASGFNFNDYFMNWVRQAPGKGLEWVAQMRNKNYQYGTYYAESLEGRFTISRDDSKNSLYLQMNSLKTEDTAVYYCARESYYGFTSYWGQGTLVTVSS
.............|||||||||||||.|..|.|.||||||||||||||..|........|||.|..||||||||..|||||||||||..|||||||||||||||..|||||||||||||
-------------PGGSLRLSCAASGFTFSSYGMHWVRQAPGKGLEWVAFIRYD--GSNKYYADSVKGRFTISRDNAKNSLYLQMNSLRAEDTAVYYCARESYYGSGSYWGQGTLVTVSS
^^^^^^^^ ^^^^^^^^^^ ^^^^^^^^^^^

Therapeutic : Omalizumab

Best Alignment of Omalizumab heavy chain to a sequence from OAS
EVQLVESGGGLVQPGGSLRLSCAVSGYSITSGYSWNWIRQAPGKGLEWVASITYD-GSTNYNPSLKGRITISRDDSKNTFYLQMNSLRAEDTAVYYCARGSHYF-GHWHFAVWGQGTLVTVSS
||||||||||||||||||||||||.||....|.|..||.|||||||||||||..|....|||.|..|.|||.|||||.||||.|||||||||||||||||||.||....|.|.|..|||||||||||
EVQLVESGGGLVQPGGSLRLSCAASGFTF-SSYAMNWVRQAPGKGLEWVSAISGSGGSTYYADSVKGRFTISRDNSKNTLYLQMNSLRAEDTAVYYCAKGSDFWSGYWWFDPWGQGTLVTVSS
^^^^^^^^^ ^^^^^^^^ ^^^^^^^^^^^^^^^

Best Alignment of Omalizumab light chain to a sequence from OAS
DIQLTQSPSSLSASVGDRVTITCRASQSVDYDGDSYMNWYQQKPGKAPKLLIYAASYLESGVPSRFSGSGSGTDFTLTISSLQPEDFATYYCQQSHEDPYTFGQGTKVEIK
|||||||||||||||||||||||||||||......||.||||||||||||||||||||.|.|||||||||||||||||||||||||||||||||||||||||..|.||||||||||||
DIQLTQSPSSLSASVGDRVTITCRASQSI----SSYLNWYQQKPGKAPKLLIYAASSLQSGVPSRFSGSGSGTDFTLTISSLQPEDFATYYCQQSHSTPRTFGQGTKVEIK
^^^^^^^^^^ ^^^ ^^^^^^^^^

Best Alignment of Omalizumab heavy chain CDRs to a sequence from OAS
EVQLVESGGGLVQPGGSLRLSCAVSGYSITSGYSWNWIRQAPGKGLEWVASITYDGSTNYNPSLKGRITISRDDSKNTFYLQMNSLRAEDTAVYYCARGSHYFGHWHFAVWGQGTLVTVSS
.|||.|||.|||.|..||.|.|.|.||||||||.||||||.||..|||...|.||||.|||||||.||.|.||.|||.|.|..||...||||.||||||||.|.|||.|.|||.||.|||||
-VQLKESGPGLVKPSQSLSLTCSVTGYSITSGYYWNWIRQFPGNKLEWMGYISYDGSNNYNPSLKNRISITRDTSKNQFFLKLNSVTTEDTATYYCARGSDYYGHWYFDVWGAGTTVTVSS
^^^^^^^^^ ^^^^^^^ ^^^^^^^^^^^^^^

Best Alignment of Omalizumab light CDRs chain to a sequence from OAS
DIQLTQSPSSLSASVGDRVTITCRASQSVDYDGDSYMNWYQQKPGKAPKLLIYAASYLESGVPSRFSGSGSGTDFTLTISSLQPEDFATYYCQQSHEDPYTFGQGTKVEIK
||.|||||.||..|.|.|.||.|.|||||||||||||||||||||||.|||||||||||.|.||.|.|||||||||||||||.|.....||.||||||||||||||||||.|||.|||
DIVLTQSPASLAVSLGQRATISCKASQSVDYDGDSYMNWYQQKPGQAPKLLIYAASNLDSGIPARFSGSGSGTDFTLNIHPVEEEDAATYYCQQSHEDPYTFGGGTKLEIK
^^^^^^^^^^ ^^^ ^^^^^^^^^

Best Alignment of Omalizumab CDR-H3 chain to a sequence from OAS
EVQLVESGGGLVQPGGSLRLSCAVSGYSITSGYSWNWIRQAPGKGLEWVASITYD-GSTNYNPSLKGRITISRDDSKNTFYLQMNSLRAEDTAVYYCARGSHYFGHWHFAVWGQGTLVTVSS
................|||||||.||......|...|.|||||||||||..|.....|..|..|..||..||||.||||.|||||.||.||||.||.||||.|..||.||||..|.||||||
----------------SLRLSCAASGFTF-NNYAMAWVRQAPGKGLEWVSVISGSATSMYYAESVRGRFIISRDKSKNTLYLQMNRLRVEDTAEYYWARGSGYYFHWYFAVWASGALVTVSS
^^^^^^^^^ ^^^^^^^^ ^^^^^^^^^^^^^^

Therapeutic : Onartuzumab

Best Alignment of Onartuzumab heavy chain to a sequence from OAS
EVQLVESGGGLVQPGGSLRLSCAASGYTFTSYWLHWVRQAPGKGLEWVGMIDPSNSDTRFNPNFKDRFTISADTSKNTAYLQMNSLRAEDTAVYYCATYRSYVTPLDYWGQGTLVTVSS
||||||||||||||||||||||||||.||.|||.||||||||||||||..|..|...|......|.|||||.|.||||.|||||||||||||||||||..|..|.||||||||||||||||
EVQLVESGGGLVQPGGSLRLSCAASGFTFSSYWMHWVRQAPGKGLEWVSAISGSGGSTYYADSVKGRFTISRDNSKNTLYLQMNSLRAEDTAVYYCARGRIAVAPLDYWGQGTLVTVSS
^^^^^^^^ ^^^^^^^^ ^^^^^^^^^^^^

Best Alignment of Onartuzumab light chain to a sequence from OAS
DIQMTQSPSSLSASVGDRVTITCKSSQSLLYTSSQKNYLAWYQQKPGKAPKLLIYWASTRESGVPSRFSGSGSGTDFTLTISSLQPEDFATYYCQQYYAYPWTFGQGTKVEIK
||.|||||.||..|.|.||||.|||||||||||..||||||||||||.||||||||||||||||||.||||||||||||||||.|||||||||||||...||.|||||||||||
DIVMTQSPDSLAVSLGERVTINCKSSQSLLYTSNNKNYLAWYQQKPGQAPKLLIYWASTRESGVPNRFSGSGSGTDFTLTISGLQPEDFATYYCQQHNSYPRTFGQGTKVEIK
^^^^^^^^^^^^ ^^^ ^^^^^^^^^

Best Alignment of Onartuzumab heavy chain CDRs to a sequence from OAS
EVQLVESGGGLVQPGGSLRLSCAASGYTFTSYWLHWVRQAPGKGLEWVGMIDPSNSDTRFNPNFKDRFTISADTSKNTAYLQMNSLRAEDTAVYYCATYRSYVTPLDYWGQGTLVTVSS
.|||..||..||.||.|...||.|||||||||||.|||.|.||.||||.||||||||.||.|..|||..|...|.|.||||.|..||..||.||||||||||..|...|||||||.|||||
QVQLKQSGPELVRPGASVKMSCKASGYTFTSYWMHWVKQRPGQGLEWIGMIDPSNSETRLNQKFKDKATLNVDKSSNTAYMQLSSLTSEDSAVYYCATYRYDVYAMDYWGQGTSVTVSS
^^^^^^^^ ^^^^^^^^ ^^^^^^^^^^^^

Best Alignment of Onartuzumab light CDRs chain to a sequence from OAS
DIQMTQSPSSLSASVGDRVTITCKSSQSLLYTSSQKNYLAWYQQKPGKAPKLLIYWASTRESGVPSRFSGSGSGTDFTLTISSLQPEDFATYYCQQYYAYPWTFGQGTKVEIK
.|.|.||||||..|||..||..||||||||||||.||||||.||||||||..||.||||||.||||||.||.|||.|||||||||||...||.|.||||||||.||.|||.|||.|||
-IVMSQSPSSLAVSVGEKVTMSCKSSQSLLYTSNQKNYLPWYQQKPGHSPKPLIYWASPRESGVPGRFTGSGTGTDFTLTISSVKAEDLAVYYCQQYYSYPPTFGGGTKLEIK
^^^^^^^^^^^^ ^^^ ^^^^^^^^^

Best Alignment of Onartuzumab CDR-H3 chain to a sequence from OAS
EVQLVESGGGLVQPGGSLRLSCAASGYTFTSYWLHWVRQAPGKGLEWVGMIDPSNSDTRFNPNFKDRFTISADTSKNTAYLQMNSLRAEDTAVYYCATYRSYVTPLDYWGQGTLVTVSS
..............|||||||||||.||.|....|||||||||||||..|..|...|......|.|||||.|.|.||.|||||||||||||||||||||||||||..|.|||||||||||||
--------------GGSLRLSCAASGFTFSSNAMSWVRQAPGKGLEWVSAISGSADSTYYADSVKGRFTISRDNSENTVYLQMNSLRAEDTAVYYCATYRSYYHPFDYWGQGTLVTVSS
^^^^^^^^ ^^^^^^^^ ^^^^^^^^^^^^

Therapeutic : Ontuxizumab

Best Alignment of Ontuxizumab heavy chain to a sequence from OAS
QVQLQESGPGLVRPSQTLSLTCTASGYTFTDYVIHWVKQPPGRGLEWIGYINPYDDDTTYNQKFKGRVTMLVDTSSNTAYLRLSSVTAEDTAVYYCARRGNSYDGYFDYSMDYWGSGTPVTVSS
.||||.|||.||.|.......|.|||||||.||.|||||.||.||||||||||||.|.|.||.||||..|...|.||.|||..|||.|.||.|||||||.|..|||||.|||||||.||.|||||
EVQLQQSGPELVKPGASGKMSCKASGYTFTSYVMHWVKQKPGQGLEWIGYINPYNDGTKYNEKFKGKATLTSDKSSSTAYMELSSLTSEDSAVYYCARGGTPYDGYFYYSMDYWGQGTSVTVSS
^^^^^^^^ ^^^^^^^^ ^^^^^^^^^^^^^^^^^

Best Alignment of Ontuxizumab light chain to a sequence from OAS
DIQMTQSPSSLSASVGDRVTITCRASQNVGTAVAWLQQTPGKAPKLLIYSASNRYTGVPSRFSGSGSGTDYTFTISSLQPEDIATYYCQQYTNYPMYTFGQGTKVQIK
||||||||||||||||||||||||||||......||.||.|||||||||||.|||..||||||||||||||||.|||||||||||||||||||||.|.||||||||||||..||
DIQMTQSPSSLSASVGDRVTITCRASQGISNYLAWYQQKPGKAPKLLIYDASNLETGVPSRFSGSGSGTDFTFTISSLQPEDIATYYCQQYDNLPMYTFGQGTKLEIK
^^^^^^ ^^^ ^^^^^^^^^^

Best Alignment of Ontuxizumab heavy chain CDRs to a sequence from OAS
QVQLQESGPGLVRPSQTLSLTCTASGYTFTDYVIHWVKQPPGRGLEWIGYINPYDDDTTYNQKFKGRVTMLVDTSSNTAYLRLSSVTAEDTAVYYCARRGNSYDGYFDYSMDYWGSGTPVTVSS
.||||.|||.||.|.......|.|||||||.||.|||||.||.||||||||||||.|.|.||.||||..|...|.||.|||..|||.|.||.|||||||.|..|||||.|||||||.||.|||||
EVQLQQSGPELVKPGASGKMSCKASGYTFTSYVMHWVKQKPGQGLEWIGYINPYNDGTKYNEKFKGKATLTSDKSSSTAYMELSSLTSEDSAVYYCARGGTPYDGYFYYSMDYWGQGTSVTVSS
^^^^^^^^ ^^^^^^^^ ^^^^^^^^^^^^^^^^^

Best Alignment of Ontuxizumab light CDRs chain to a sequence from OAS
DIQMTQSPSSLSASVGDRVTITCRASQNVGTAVAWLQQTPGKAPKLLIYSASNRYTGVPSRFSGSGSGTDYTFTISSLQPEDIATYYCQQYTNYPMYTFGQGTKVQIK
.|..|||....|.|||||...||.|||||||.|||.||.||..||.||||||.||.|||.||.||||||||.|.|.|..|.||.|.|.||||.|||.||||.|||..||
-IVLTQSQKFVSPSVGDRDSVTCKASQNVGTNVAWYQQKPGQSPKPLIYSASYRYSGVPDRFTGSGSGTDFTLTVSNVQSEDLAEYFCQQYNNYPPYTFGGGTKLEIK
^^^^^^ ^^^ ^^^^^^^^^^

Best Alignment of Ontuxizumab CDR-H3 chain to a sequence from OAS
QVQLQESGPGLVRPSQTLSLTCTASGYTFTDYVIHWVKQPPGRGLEWIGYINPYDDDTTYNQKFKGRVTMLVDTSSNTAYLRLSSVTAEDTAVYYCARRGNSYDGYFDYSMDYWGSGTPVTVSS
.||||.||..||||.....|.|.|||||||||||.|.||||.||.|||||..|.|....|.||.||||..|.....||.|||..|||.|.||.|||.|||||.|||||.||.|||||.||.|||||

-VQLQQSGAELVRPGASVKLSCKASGYTFTDYYINWVKQRPGQGLEWIARIYPGSGNTYYNEKFKGKATLTAEKSSSTAYMQLSSLTSEDSAVYVCARRGVSYDGYYDYAMDYWGQGTSVTVSS
                         ^^^^^^^^            ^^^^^^^^                    ^^^^^^^^^^^^^^^^^

Therapeutic : Opicinumab

Best Alignment of Opicinumab heavy chain to a sequence from OAS
EVQLLESGGGLVQPGGSLRLSCAASGFTFSAYEMKWVRQAPGKGLEWVSVIGPSGGFTFYADSVKGRFTISRDNSKNTLYLQMNSLRAEDTAVYYCATEGDN---DAFDIWGQGTTVTVSS
||||||||||||||||||||||||||||||.|.|.||||||||||||||.|..|||.|.||||||||||||||||||||||||||||||||||||||.||||...||||||||||.|||||
EVQLLESGGGLVQPGGSLRLSCAASGFTFSSYAMSWVRQAPGKGLEWVSTISGSGGSTYYADSVKGRFTISRDNSKNTLYLQMNSLRAEDTAVYYCAREGDNWNEDAFDIWGQGTMVTVSS
                         ^^^^^^^^            ^^^^^^^^                    ^^^^^^^^^^^^^^

Best Alignment of Opicinumab light chain to a sequence from OAS
DIQMTQSPATLSLSPGERATLSCRASQSVSSYLAWYQQKPGQAPRLLIYDASNRATGIPARFSGSGSGTDFTLTISSLEPEDFAVYYCQQRSNWPMYTFGQGTKLEIK
.|.|||||||||||||||||||||||||||||||||||||||||||||||||||||||||||||||||||||||||||||||||||||||||||||||||||||||||
EIVMTQSPATLSLSPGERATLSCRASQSVSSYLAWYQQKPGQAPRLLIYDASNRATGIPARFSGSGSGTDFTLTISSLEPEDFAVYYCQQRSNWPMYTFGQGTKLEIK
                          ^^^^^^            ^^^                    ^^^^^^^^^^

Best Alignment of Opicinumab heavy chain CDRs to a sequence from OAS
EVQLLESGGGLVQPGGSLRLSCAASGFTFSAYEMKWVRQAPGKGLEWVSVIGPSGGFTFYADSVKGRFTISRDNSKNTLYLQMNSLRAEDTAVYYCATEGDNDAFDIWGQGTTVTVSS
..............|.||..||||||||||.|||.||||||||||||||.|..||..|..||||||||||||||.||.|||||||||||||||||.||||||||||||||.|||||
--------------GESLKISCAASGFTFSNYEMIWVRQAPGKGLEWVSYISSSGSTTYSADSVKGRFTISRDNAKNSLYLQMNSLRAEDTAVYYCAREGDNDAFDIWGQGTMVTVSS
                         ^^^^^^^^            ^^^^^^^^                    ^^^^^^^^^^^

Best Alignment of Opicinumab light CDRs chain to a sequence from OAS
DIQMTQSPATLSLSPGERATLSCRASQSVSSYLAWYQQKPGQAPRLLIYDASNRATGIPARFSGSGSGTDFTLTISSLEPEDFAVYYCQQRSNWPMYTFGQGTKLEIK
.|..||||.|||||||||||||||||||||||||||||||||||||||||||||||||||||||||||||||||||||||||||||||||||||||||||||||||||
EIVLTQSPVTLSLSPGERATLSCRASQSVSSYLAWYQQKPGQAPRLLIYDASNRATGIPARFSGSGSGTDFTLTISSLEPEDFAVYYCQQRSNWPMYTFGQGTKLEIK
                          ^^^^^^            ^^^                    ^^^^^^^^^^

Best Alignment of Opicinumab CDR-H3 chain to a sequence from OAS
EVQLLESGGGLVQPGGSLRLSCAASGFTFSAYEMKWVRQAPGKGLEWVSVIGPSGGFTFYADSVKGRFTISRDNSKNTLYLQMNSLRAEDTAVYYCATEGDNDAFDIWGQGTTVTVSS
.................|.|.|..||...|.|...|.||.|||||||...|..|.|.|.|..|.|.|.|||.|.|||...|...|..|.|||||||||||.||||||||||.|||||
----------------TLSLTCTVSGGSISSYYWSWIRQPPGKGLEWIGYIYYS-GSTNYNPSLKSRVTISVDTSKNQFSLKLSSVTAADTAVYYCATEGDRDAFDIWGQGTMVTVSS
                         ^^^^^^^^            ^^^^^^^^                    ^^^^^^^^^^^

Therapeutic : Orticumab

Best Alignment of Orticumab heavy chain to a sequence from OAS
EVQLLESGGGLVQPGGSLRLSCAASGFTFSNAWMSWVRQAPGKGLEWVSSISVGGHRTYYADSVKGRSTISRDNSKNTLYLQMNSLRAEDTAVYYCARIRVGP-SGGAFDYWGQGTLVTVSS
||||||||||||||||||||||||||||||...||||||||||||||||.||.||..||||||||||.|||||||||||||||||||||||||||||..||..|.|.||||||||||||||
EVQLLESGGGLVQPGGSLRLSCAASGFTFSSYAMSWVRQAPGKGLEWVSAISGGGGSTYYADSVKGRFTISRDNSKNTLYLQMNSLRAEDTAVYYCARGLVGATSRGYFDYWGQGTLVTVSS
                         ^^^^^^^^            ^^^^^^^^                    ^^^^^^^^^^^^^^^

Best Alignment of Orticumab light chain to a sequence from OAS
QSVLTQPPSASGTPGQRVTISCSGSNTNIGKNYVSWYQQLPGTAPKLLIYANSNRPSGVPDRFSGSKSGTSASLAISGLRSEDEADYYCASWDASLNGWVFGGGTKLTVL
|||||||||||||||||||||||||..|||||||.|||||||||||||||.||.||||||||||||||||||||||||||||||||||||.||.||.|||||||||||||
QSVLTQPPSASGTPGQRVTISCSGSSSNIGKNYVYWYQQLPGTAPKLLIYRNSQRPSGVPDRFSGSKSGTSASLAISGLRSEDEADYYCAVWDDSLGGWVFGGGTKLTVL
                         ^^^^^^^^            ^^^                    ^^^^^^^^^^^

Best Alignment of Orticumab heavy chain CDRs to a sequence from OAS
EVQLLESGGGLVQPGGSLRLSCAASGFTFSNAWMSWVRQAPGKGLEWVSSISVGGHRTYYADSVKGRSTISRDNSKNTLYLQMNSLRAEDTAVYYCARIRVGPSGGAFDYWGQGTLVTVSS
||||.||||||||||.||.||||||||||||..|.||||.|.||||||.|||.||..|||.||||||.||||||.|||.||||.|||.||||.|||||..||||||.|||||||..|||||
EVQLVESGGGLVQPGRSLKLSCAASGFTFSNYGMAWVRQTPTKGLEWVASISTGGGNTYYRDSVKGRFTISRDNAKNTQYLQMDSLRSEDTATYYCARHEVGPSGGYFDYWGQGVMVTVSS
                         ^^^^^^^^            ^^^^^^^^                    ^^^^^^^^^^^^^^

Best Alignment of Orticumab light CDRs chain to a sequence from OAS
QSVLTQPPSASGTPGQRVTISCSGSNTNIGKNYVSWYQQLPGTAPKLLIYANSNRPSGVPDRFSGSKSGTSASLAISGLRSEDEADYYCASWDASLNGWVFGGGTKLTVL
|||||||||.|..|||.|||||||||.|||.|||.||||.||||||.|||.||.||||.|.||||||||||||||||||..|||||||||.||.||||||||||||||||
QSVLTQPPSVSAAPGQKVTISCSGSNSNIGNNYVTWYQQFPGTAPKVLIYDNSRRPSGIPARFSGSKSGTSASLAISGLQAEDEADYYCAAWDDSLNGWVFGGGTKLTVL
                         ^^^^^^^^            ^^^                    ^^^^^^^^^^^

Best Alignment of Orticumab CDR-H3 chain to a sequence from OAS
EVQLLESGGGLVQPGGSLRLSCAASGFTFSNAWMSWVRQAPGKGLEWVSSISVGGHRTYYADSVKGRSTISRDNSKNTLYLQMNSLRAEDTAVYYCARIRVGPSGGAFDYWGQGTLVTVSS
.|||.|||.|||.....|.|.|..||...|....||.||..|||||....|......|.|..|.|.|.|.|.|.|||...|...|..|.|||||||||.||..||||||||||||||||||
QVQLQESGPGLVETSETLSLTCNVSGGSISSYYWSWIRQPAGKGLEYIGRIHTS-GSTNYNPSLKSRVTMSVDMSKNQFSLMLTSVTAADTAVYYCARLRVPASGGAFDYWGQGTLVTVSS
                         ^^^^^^^^            ^^^^^^^^                    ^^^^^^^^^^^^^^

Therapeutic : Otelixizumab

Best Alignment of Otelixizumab heavy chain to a sequence from OAS
EVQLLESGGGLVQPGGSLRLSCAASGFTFSSFPMAWVRQAPGKGLEWVSTISTSGGRTYYRDSVKGRFTISRDNSKNTLYLQMNSLRAEDTAVYYCAKFRQY---SGGFDYWGQGTLVTVSS
|||||||||||||||||||||||||||||||..|.||||||||||||||.||.|||.|||.|||||||||||||||||||||||||||||||||||||.|||...|||.|||||||||||||
EVQLLESGGGLVQPGGSLRLSCAASGFTFSSYAMSWVRQAPGKGLEWVSAISGSGGSTYYADSVKGRFTISRDNSKNTLYLQMNSLRAEDTAVYYCAKDRQYYDSSGGVDYWGQGTLVTVSS
                         ^^^^^^^^            ^^^^^^^^                    ^^^^^^^^^^^^^^^

Best Alignment of Otelixizumab light chain to a sequence from OAS
DIQLTQPNSVSTSLGSTVKLSCTLSSGNIENNYVHWYQLYEGRSPTTMIYDDDKRPDGVPDRFSGSIDRSSNSAFLTIHNVAIEDEAIYFCHSYVS-SFNVFGGGTKLTVL
...||||.|||.|.|.||..|||.|||||.||||.|||...|..|||.||||||||.|||||||||||||||||.|||.....||||.|.|.||.|.|..|||||||||||
NFMLTQPHSVSESPGKTVIISCTRSSGNIGNNYVQWYQQRPGSAPTTVIYDDDKRPSGVPDRFSGSIDRSSNSASLTISGLKTEDEADYYCQSYDSSSDVVFGGGTKLTVL
                         ^^^^^^^^            ^^^                    ^^^^^^^^^^

Best Alignment of Otelixizumab heavy chain CDRs to a sequence from OAS
EVQLLESGGGLVQPGGSLRLSCAASGFTFSSFPMAWVRQAPGKGLEWVSTISTSGGRTYYRDSVKGRFTISRDNSKNTLYLQMNSLRAEDTAVYYCAKFRQYSGGFDYWGQGTLVTVSS
||||.||||||||||.|..||||||||||||||||||||||.||||||.|||||||.|||||||||||||||||.|.||||||||||.||||.|||...|.|||||||||||..|||||
EVQLVESGGGLVQPGRSMKLSCAASGFTFSSFPMAWVRQAPTKGLEWVATISTSGGSTYYRDSVKGRFTISRDNAKSTLYLQMNSLRSEDTATYYCTRDRYYSGGFDYWGQGVMVTVSS
                         ^^^^^^^^            ^^^^^^^^                    ^^^^^^^^^^^^

Best Alignment of Otelixizumab light CDRs chain to a sequence from OAS
DIQLTQPNSVSTSLGSTVKLSCTLSSGNIENNYVHWYQLYEGRSPTTMIYDDDKRPDGVPDRFSGSIDRSSNSAFLTIHNVAIEDEAIYFCHSYVSSFNVFGGGTKLTVL
...||||.|||.|.|.||..|||.|||||.||||.|||...|..|||.||||||||.|||||||||||.|||||.|||.....||||.|.|.||.||..|||||||||||
NFMLTQPHSVSESPGKTVIISCTRSSGNIGNNYVQWYQQRPGSAPTTVIYDDDKRPSGVPDRFSGSIDSSSNSASLTISGLKTEDEADYYCQSYDSSNQVFGGGTKLTVL
                         ^^^^^^^^            ^^^                    ^^^^^^^^^

Best Alignment of Otelixizumab CDR-H3 chain to a sequence from OAS
EVQLLESGGGLVQPGGSLRLSCAASGFTFSSFPMAWVRQAPGKGLEWVSTISTSGGRTYYRDSVKGRFTISRDNSKNTLYLQMNSLRAEDTAVYYCAKFRQYSGGFDYWGQGTLVTVSS
.|||...|..||.||.|..|||.|||.||.|..|.||.|.||.||||...|....|.|.|....|...|...|....|.|....||..||.|||||||||.|.||||||||||..||||
QVQLQQPGAELVKPGASVKLSCKASGYTFTSYWMHWVKQRPGRGLEWIGRIDPNSGGTKYNEKFKSKATLTVDKPTSTAYMKLSSLTSEDSAVYYCAKFRDYDGGFDYWGQGTTLTVSS
                         ^^^^^^^^            ^^^^^^^^                    ^^^^^^^^^^^^

Therapeutic : Otlertuzumab

Best Alignment of Otlertuzumab heavy chain to a sequence from OAS
EVQLVQSGAEVKKPGESLKISCKGSGYSFTGYNMNWVRQMPGKGLEWMGNIDPYYGGTTYNRKFKGQVTISADKSISTAYLQWSSLKASDTAMYYCARSVG--PFDSWGQGTLVTVSS
||||||||||||||||||.|||||||||||.|..|||||||||||||||||||....|.|...|.|||||||||||||||||||||||||||||||||.||..|||.|||||||||||
EVQLVQSGAEVKKPGESLRISCKGSGYSFTSYWINWVRQMPGKGLEWMGNIDPGDSDTRYSPSFQGQVTISADKSISTAYLQWSSLKASDTAMYYCARHVGATPFDYWGQGTLVTVSS
                         ^^^^^^^^            ^^^^^^^^                    ^^^^^^^^^^^

Best Alignment of Otlertuzumab light chain to a sequence from OAS

```
EIVLTQSPATLSLSPGERATLSCRASENVYSYLAWYQQKPGQAPRLLIYFAKTLAEGIPARFSGSGSGTDFTLTISSLEPEDFAVYYCQHHSDNPWTFGQGTKVEIK
||||||||||||||||||||||||||.||.||||||||||||||||||||.|.|.|.||||||||||||||||||||||||||||||||||..|..|||||||||||||
EIVLTQSPATLSLSPGERATLSCRASQNVSSYLAWYQQKPGQAPRLLIYDASTRATGIPARFSGSGSGTDFTLTISSLEPEDFAVYYCQQRSIWPWTFGQGTKVEIK
                         ^^^^^^          ^^^                   ^^^^^^^^^
```

Best Alignment of Otlertuzumab heavy chain CDRs to a sequence from OAS

```
EVQLVQSGAEVKKPGESLKISCKGSGYSFTGYNMNWVRQMPGKGLEWMGNIDPYYGGTTYNRKFKGQVTISADKSISTAYLQWSSLKASDTAMYYCARSVGPFDSWGQGTLVTVSS
||||.|||.|..|||.|.|||||.|||||||||||||.|..||.|||.||||||||||.||.||||..|...|||.||||.|..||...|.|.||||||.||||.|||||..||||
EVQLQQSGPELEKPGASVKISCKASGYSFTGYNMNWVKQSNGKSLEWIGNIDPYYGGTSYNQKFKGKATLTVDKSSSTAYMQLKSLTSEDSAVYYCARSSGPFDYWGQGTTLTVSS
                        ^^^^^^^^         ^^^^^^^^                  ^^^^^^^^^
```

Best Alignment of Otlertuzumab light CDRs chain to a sequence from OAS

```
EIVLTQSPATLSLSPGERATLSCRASENVYSYLAWYQQKPGQAPRLLIYFAKTLAEGIPARFSGSGSGTDFTLTISSLEPEDFAVYYCQHHSDNPWTFGQGTKVEIK
.|..|||||.||.|.||..|...|||||.||||||||||.|..|..|.|.|||||||.|.|||||||||.|.|.|.||.||||..||||||.|.|||||.|||.|||
DIQITQSPASLSASVGEAVTITYRASENIYSYLAWYQQKQGKSPQVLVYNAKTLAEGVPSRFSGSGSGTQFSLKINSLQPEDFGNYYCQHHYDTPWTFGGGTKLEIK
                         ^^^^^^          ^^^                   ^^^^^^^^^
```

Best Alignment of Otlertuzumab CDR-H3 chain to a sequence from OAS

```
EVQLVQSGAEVKKPGESLKISCKGSGYSF--TGYNMNWVRQMPGKGLEWMGNIDPY-YGGTTYNRKFKGQVTISADKSISTAYLQWSSLKASDTAMYYCARSVGPFDSWGQGTLVTVSS
................|...|..||.|...|....||.||.|..||||.|...........|.....|..||....|.....||..|....|||.|||||||||.||||||||||||
-------------SQTLSLTCAISGDSVSSTNAAWNWIRQSPSRGLEWLGRTYYRSRWYNEYAVSLRGRITINPVTSKNQFSLQLNSVSPEDTAVYYCARSVGPIDSWGQGTLVTVSS
                        ^^^^^^^^^^         ^^^^^^^^^                    ^^^^^^^^^
```

Therapeutic : Ozanezumab

Best Alignment of Ozanezumab heavy chain to a sequence from OAS

```
QVQLVQSGAEVKKPGASVKVSCKASGYTFTSYWMHWVRQAPGQGLEWIGNINPSNGGTNYNEKFKSKATMTRDTSTSTAYMELSSLRSEDTAVYYCELM--------QGYWGQGTLVTVSS
||||||||||||||||||||||||||||||||.||||||||||||||.|.||||.|||.|..|||.|.||||||||||.||||||||||||||||||............|||||||||||
QVQLVQSGAEVKKPGASVKVSCKASGYTFTSYYMHWVRQAPGQGLEWMGIINPSGGGTSYAQKFKGKVTMTRDTSTSTVYMELSSLRSEDTAVYYCARGYGSGSYYLFDYWGQGTLVTVSS
                        ^^^^^^^^         ^^^^^^^^                  ^^^^^^^^^^^^^^
```

Best Alignment of Ozanezumab light chain to a sequence from OAS

```
DIVMTQSPLSNPVTLGQPVSISCRSSKSLLYKDGKTYLNWFLQRPGQSPQLLIYLMSTRASGVPDRFSGGGSGTDFTLKISRVEAEDVGVYYCQQLVEYPLTFGQGTKLEIK
|||.||..||||||.|..|||||||||||||||||||||||||||||||||||||||||||.|||||.||||||||.||||.|||||||||||||||||||||.|||||||
DIVITQDELSNPVTSGESVSISCRSSKSLLYKDGKTYLNWFLQRPGQSPQLLIYLMSTRASGVSDRFSGSGSGTDFTLEISRVKAEDVGVYYCQQLVEYPLTFGSGTKLEIK
                         ^^^^^^^^^^^          ^^^                   ^^^^^^^^^
```

Best Alignment of Ozanezumab heavy chain CDRs to a sequence from OAS

```
QVQLVQSGAEVKKPGASVKVSCKASGYTFTSYWMHWVRQAPGQGLEWIGNINPSNGGTNYNEKFKSKATMTRDTSTSTAYMELSSLRSEDTAVYYCELMQGYWGQGTLVTVSS
.|||.|.|.|..|||||||.||||||||||||||||.|.||||||||||||||||||||||||||||||.|.|.|.|||||.||||.|||.|||||.|.||||||||..||||
-VQLQQPGTELVKPGASVKLSCKASGYTFTSYWMHWVKQRPGQGLEWIGNINPSNGGTNYNEKFKSKATLTVDKSSSTAYMQLSSLTSEDSAVYYCGLGQGYWGQGTTLTVSS
                        ^^^^^^^^         ^^^^^^^^                  ^^^^^^
```

Best Alignment of Ozanezumab light CDRs chain to a sequence from OAS

```
DIVMTQSPLSNPVTLGQPVSISCRSSKSLLYKDGKTYLNWFLQRPGQSPQLLIYLMSTRASGVPDRFSGGGSGTDFTLKISRVEAEDVGVYYCQQLVEYPLTFGQGTKLEIK
|||.||..||||||.|..|||||||||||||||||||||||||||||||||||||||||||.|||||.||||||||.||||.|||||||||||||||||||||.|||||||
DIVITQDELSNPVTSGESVSISCRSSKSLLYKDGKTYLNWFLQRPGQSPQLLIYLMSTRASGVSDRFSGSGSGTDFTLEISRVKAEDVGVYYCQQLVEYPLTFGSGTKLEIK
                         ^^^^^^^^^^^          ^^^                   ^^^^^^^^^
```

Best Alignment of Ozanezumab CDR-H3 chain to a sequence from OAS

```
QVQLVQSGAEVKKPGASVKVSCKASGYTFTSYWMHWVRQAPGQGLEWIGNINPSNGGTNYNEKFKSKATMTRDTSTSTAYMELSSLRSEDTAVYYCELMQGYWGQGTLVTVSS
||||...|||..|||||||.||||||||||||||.||.|.||||||||||.|.||...||||.|||..||.|.|||.|||||.||||..|.|.||||||.|||||.||..||||
QVQLQPPGAELVKPGASVKLSCKASGYTFTSYWMQWVKQRPGQGLEWIGVIDPSDSYTNYNQKFKGQATLTVDTSSSTAYMQLSSLTYEETGVYYCELGQGYWGKGTTRTVSS
                        ^^^^^^^^         ^^^^^^^^                  ^^^^^^
```

Therapeutic : Palivizumab

Best Alignment of Palivizumab heavy chain to a sequence from OAS

```
QVTLRESGPALVKPTQTLTLTCTFSGFSLSTSGMSVGWIRQPPGKALEWLADIWWDDKKDYNPSLKSRLTISKDTSKNQVVLKVTNMDPADTATYYCARSMIT--------NWYFDVWGAGTTVTVSS
|||||||||||||||||||||||||||||||||||.|.||||||||||||||.|.|||.|.|..|||.|||||||||||||||..|||||.||||||||||..|........|...||||.||||||||
QVTLRESGPALVKPTQTLTLTCTFSGFSLSTSGMCVSWIRQPPGKALEWLALIDWDDDKYYSTSLKTRLTISKDTSKNQVVLTMTNMDPVDTATYYCARSTVTAPIRRRSKNYGMDVWGQGTTVTVSS
                        ^^^^^^^^^^         ^^^^^^^                  ^^^^^^^^^^^^^^^^^^^^
```

Best Alignment of Palivizumab light chain to a sequence from OAS

```
DIQMTQSPSTLSASVGDRVTITCKCQLSV-GYMHWYQQKPGKAPKLLIYDTSKLASGVPSRFSGSGSGTEFTLTISSLQPDDFATYYCFQGSGYPFTFGGGTKLEIK
||||||||||||||||||||||||....|...|..||||||||||||||||..|.||||||||||||||||||||||||||||||||||||.|...||||||.|||||||
DIQMTQSPSTLSASVGDRVTITCRASQSISSYLAWYQQKPGKAPKLLIYKASSLASGVPSRFSGSGSGTEFTLTISSLQPDDFATYYCQQYNSYPFTFGQGTKLEIK
                         ^^^^^^          ^^^                   ^^^^^^^^^
```

Best Alignment of Palivizumab heavy chain CDRs to a sequence from OAS

```
QVTLRESGPALVKPTQTLTLTCTFSGFSLSTSGMSVGWIRQPPGKALEWLADIWWDDKKDYNPSLKSRLTISKDTSKNQVVLKVTNMDPADTATYYCARSMITNWYFDVWGAGTTVTVSS
||||..|||......|||.|||.|.|||||||||.|.|||||.||.|||||.|.|||.|.|||||||||||||||||.|||.||.|..|.|||||||||||||||.|||||||.||||||||
QVTLKVSGPGILQSSQTLSLTCSFPGFSLSTSGMGVSWIRQPSGKGLEWLAHIYWDDDKRYNPSLKSRLTISKDTSRNQVFLKITSVDTADTATYYCARSMITTWYFDVWGTGTTVTVSS
                        ^^^^^^^^^^         ^^^^^^^                  ^^^^^^^^^^^^
```

Best Alignment of Palivizumab light CDRs chain to a sequence from OAS

```
DIQMTQSPSTLSASVGDRVTITCKCQLSVGYMHWYQQKPGKAPKLLIYDTSKLASGVPSRFSGSGSGTEFTLTISSLQPDDFATYYCFQGSGYPFTFGGGTKLEIK
||..||.|...|||.|..||.||...|||.||||||||....|||.||||||||||||.||||||||....|||||....|.||||||||||||||||||.|||||||
DIVLTQTPAIMSASPGEKVTMTCSASLSVSYMHWYQQKSSTSPKLWIYDTSKLASGVPGRFSGSGSGNSYSLTISSMEAEDVATYYCFQGSGYPFTFGSGTKLEIK
                         ^^^^^          ^^^                   ^^^^^^^^^
```

Best Alignment of Palivizumab CDR-H3 chain to a sequence from OAS

```
QVTLRESGPALVKPTQTLTLTCTFSGFSLSTSGMSVGWIRQPPGKALEWLADIWWD-DKKDYNPSLKSRLTISKDTSKNQVVLKVTNMDPADTATYYCARSMITNWYFDVWGAGTTVTVSS
||.|......||.|.....|.|..||...........|..|.||..|||...|........||...|...|...|.|..............|.|.||||||||||||||||||.||||||||
QVQLQQPVAELVMPGASVKLSCKASGYTF--TSYWMHWVKQRPGQGLEWIGEIDPSDSYTNYNQKFKGKSTLTVDKSSSTAYMQLSSLTSEDSAVYYCARSMITNWYFDVWGTGTTVTVSS
                        ^^^^^^^^^^         ^^^^^^^^                   ^^^^^^^^^^^^
```

Therapeutic : Pamrevlumab

Best Alignment of Pamrevlumab heavy chain to a sequence from OAS

```
EGQLVQSGGGLVHPGGSLRLSCAGSGFTFSSYGMHWVRQAPGKGLEWVSGIGTGGGTYSTDSVKGRFTISRDNAKNSLYLQMNSLRAEDMAVYYCARGDYYG--SGSFFDCWGQGTLVTVSS
|.|||||||||||||||||||||.|||||||||||.|||||||||||||||||.||||||||..|||||||||||||||||||||||||||||||||.|||||||..||...||...|.||.||||||||
EVQLVQSGGGLVHPGGSLRLSCSGSGFTFSSYAMHWVRQAPGKGLEWVSAIGTGGGTYYADSVKGRFTISRDNAKNSLYLQMNSLRAEDTAVYYCARDGYYYESSGYTLDYWGRGTLVTVSS
                        ^^^^^^^^         ^^^^^^^                  ^^^^^^^^^^^^^^^^
```

Best Alignment of Pamrevlumab light chain to a sequence from OAS

```
DIQMTQSPSSLSASVGDRVTITCRASQGISSWLAWYQQKPEKAPKSLIYAASSLQSGVPSRFSGSGSGTDFTLTISSLQPEDFATYYCQQYNSYPPTFGQGTKLEIK
|||||||||||||||||||||||||||||||||||||||||||||||||||||||||||||||||||||||||||||||||||||||||||||||||||||||||||
DIQMTQSPSSLSASVGDRVTITCRASQGISSWLAWYQQKPEKAPKSLIYAASSLQSGVPSRFSGSGSGTDFTLTISSLQPEDFATYYCQQYNSYPPTFGQGTKLEIK
                         ^^^^^^          ^^^                   ^^^^^^^^^
```

Best Alignment of Pamrevlumab heavy chain CDRs to a sequence from OAS

```
EGQLVQSGGGLVHPGGSLRLSCAGSGFTFSSYGMHWVRQAPGKGLEWVSGIGTGGGTYSTDSVKGRFTISRDNAKNSLYLQMNSLRAEDMAVYYCARGDYYGSGSFFDCWGQGTLVTVSS
................|||||||.|||||||||.||||||||.|||||||||.|||.|.||...||||||||||.|||||||||||||||.|.|||||||||||||||.||.||||.||||||
```

```
----------------SLRLSCAASGFTFSSYDMHWVRQATGKGLEWVSAIGTAGDTYYPGSVKGRFTISRENAKNSLYLQMNSLRAGDTAVYYCARGDYYGSGSYFDYWGQGALVTVSS
                 ^^^^^^^^        ^^^^^^^                        ^^^^^^^^^^^^^^
```

Best Alignment of Pamrevlumab light CDRs chain to a sequence from OAS

```
DIQMTQSPSSLSASVGDRVTITCRASQGISSWLAWYQQKPEKAPKSLIYAASSLQSGVPSRFSGSGSGTDFTLTISSLQPEDFATYYCQQYNSYPPTFGQGTKLEIK
|.||..|..|...||||||||||||||||||||||||||.||||.|||||||||||||||||||||||||||||||||||||||||||||||||||||.|||.|||
DSQMRKSTASVPPSVGDRVTITCRASQGISSWLAWYQQKPGKAPKLLIYAASSLQSGVPSRFSGSGSGTDFTLTISSLQPEDFATYYCQQYNSYPPTFGEGTKVEIK
                  ^^^^^^        ^^^                    ^^^^^^^^^
```

Best Alignment of Pamrevlumab CDR-H3 chain to a sequence from OAS

```
EGQLVQSGGGLVHPGGSLRLSCAGSGFTFSSYGMHWVRQAPGKGLEWVSGIGTGGGTYSTDSVKGRFTISRDNAKNSLYLQMNSLRAEDMAVYYCARGDYYGSGSFFDCWGQGTLVTVSS
................|.|.|..|....|.|...|.||.|||||||...|...|.|....|.|.|.|||.|...|...|...|..|.|.||||||||||||||||||.|||||||||||
-------------SETLSLTCTVSAGSISGYFWSWIRQPPGKGLEWIGYIYYSGSTNYNPSLKSRVTISVDTSNNQFSLKLSSVTAADKAVYYCARGDYYGSGSFFDYWGQGTLVTVSS
                 ^^^^^^^^        ^^^^^^^                        ^^^^^^^^^^^^^^
```

Therapeutic : Panitumumab

Best Alignment of Panitumumab heavy chain to a sequence from OAS

```
QVQLQESGPGLVKPSETLSLTCTVSGGSVSSGDYYWTWIRQSPGKGLEWIGHIYYSGNTNYNPSLKSRLTISIDTSKTQFSLKLSSVTAADTAIYYCVRDRVT-GAFDIWGQGTMVTVSS
||||||||||||||||||||||||||||||||.|||.||||.|||||||||.|||||.||||||||||.|||.||||.|||||||||||||||.|||.||.||.||||||||||||||||
QVQLQESGPGLVKPSETLSLTCTVSGGSVSSGSYYWSWIRQPPGKGLEWIGYIYYSGSTNYNPSLKSRVTISVDTSKNQFSLKLSSVTAADTAVYYCARDLVTQGAFDIWGQGTMVTVSS
                 ^^^^^^^^^^        ^^^^^^^                        ^^^^^^^^^^^^
```

Best Alignment of Panitumumab light chain to a sequence from OAS

```
DIQMTQSPSSLSASVGDRVTITCQASQDISNYLNWYQQKPGKAPKLLIYDASNLETGVPSRFSGSGSGTDFTFTISSLQPEDIATYFCQHFDHLPLAFGGGTKVEIK
||||||||||||||||||||||||||||||||||||||||||||||||||||||||||||||||||||||||||||||||||||||||||.|.|||.||||||||||
DIQMTQSPSSLSASVGDRVTITCQASQDISNYLNWYQQKPGKAPKLLIYDASNLETGVPSRFSGSGSGTDFTFTISSLQPEDIATYFCQHYDNLPLTFGGGTKVEIK
                  ^^^^^^        ^^^                    ^^^^^^^^^
```

Best Alignment of Panitumumab heavy chain CDRs to a sequence from OAS

```
QVQLQESGPGLVKPSETLSLTCTVSGGSVSSGDYYWTWIRQSPGKGLEWIGHIYYSGNTNYNPSLKSRLTISIDTSKTQFSLKLSSVTAADTAIYYCVRDRVTGAFDIWGQGTMVTVSS
|||||||||||||||.||||||||||||||||||||.||||.||||||.||.|||||||.||||||||.|||.||||.||||||.||||||||.|||.||..|||||||||||||||||
QVQLQESGPGLVKPSQTLSLTCTVSGGSVSSGDYYWSWIRQPPGKGLECIGYIYYSGNTYYNPSLKSRVTISVDTSKNQFSLKLNSVTAADTAVYYCARDGETGAFDIWGQGTMVTVSS
                 ^^^^^^^^^^        ^^^^^^^                        ^^^^^^^^^^^
```

Best Alignment of Panitumumab light CDRs chain to a sequence from OAS

```
DIQMTQSPSSLSASVGDRVTITCQASQDISNYLNWYQQKPGKAPKLLIYDASNLETGVPSRFSGSGSGTDFTFTISSLQPEDIATYFCQHFDHLPLAFGGGTKVEIK
||||||||||||||||||||||||||||||||||||||.||||||||||||||||.||||||||.||||.|.||||.|||||||||.|||||||||.|||||..|||
DIQMTQSPSSLSASVGDRVTITCQASQDISNYLNWYQQRPGKAPKLLIYDASNLESGVPSRFSGRGSGTHFSFTISGLQPEDIATYYCQHFDHLPLTFGGGTRLEIK
                  ^^^^^^        ^^^                    ^^^^^^^^^
```

Best Alignment of Panitumumab CDR-H3 chain to a sequence from OAS

```
QVQLQESGPGLVKPSETLSLTCTVSGGSVSSGDYYWTWIRQSPGKGLEWIGHIYYS-GNTNYNPSLKSRLTISIDTSKTQFSLKLSSVTAADTAIYYCVRDRVTGAFDIWGQGTMVTVSS
................|.|.|.||........|...|.||.||||||||..|....|...|..|...|.|||.|.......|...|..|.|||.|.||||||.||||||||||||||||
--------------GGSLRLSCAVSEFTF--SSYDMNWVRQAPGKGLEWISFITTRSGTIYYADSVRGRFTISRDNAQNSLYLQMNSLRAEDTAVYHCVRDRVGGAFDIWGQGTMVTVSS
                 ^^^^^^^^^^        ^^^^^^^^                        ^^^^^^^^^^^
```

Therapeutic : Panobacumab

Best Alignment of Panobacumab heavy chain to a sequence from OAS

```
EEQVVESGGGFVQPGGSLRLSCAASGFTFSPYWMHWVRQAPGKGLVWVSRINSD-GSTYYADSVKGRFTISRDNARNTLYLQMNSLRAEDTAVYYCARDRY-------------YGPEMWGQGTMVTVSS
|.|.||||||||||||||||||||||||||.|||||||||||||||||||||||..||.||||||||||||||||.|||||||||||||||||||||||||.............||...|||||.|||||
EVQLVESGGGFVQPGGSLRLSCAASGFTFSSYWMHWVRQAPGKGLVWVSRINSDGSSTSYADSVKGRFTISRDNAKNTLYLQMNSLRAEDTAVYYCARDRYYDFWSGYYPFDYYYYGMDVWGQGTTVTVSS
                 ^^^^^^^^        ^^^^^^^^                        ^^^^^^^^^^^^^^^^^^^^^^^^
```

Best Alignment of Panobacumab light chain to a sequence from OAS

```
DVVMTQSPLSLPVTLGQPASISCRSSQSLVYSDGNTYLNWFQQRPGQSPRRLIYKVSNRDSGVPDRFSGSGSGTDFTLKISRVEAEDVGVYYCMQGTHWPLTFGGGTKVEIK
||||||||||||||||||||||||||||||||||||||||||||||||||||||||||||||||||||||||||||||||||||||||||||||||||||||||||||||||
DVVMTQSPLSLPVTLGQPASISCRSSQSLVYSDGNTYLNWFQQRPGQSPRRLIYKVSNRDSGVPDRFSGSGSGTDFTLKISRVEAEDVGVYYCMQGTHWPLTFGGGTKVEIK
                  ^^^^^^^^^^^        ^^^                    ^^^^^^^^^
```

Best Alignment of Panobacumab heavy chain CDRs to a sequence from OAS

```
EEQVVESGGGFVQPGGSLRLSCAASGFTFSPYWMHWVRQAPGKGLVWVSRINSDGSTYYADSVKGRFTISRDNARNTLYLQMNSLRAEDTAVYYCARDRYYGPEMWGQGTMVTVSS
..............||||||||||||||||.|||||||.||||||||||||||||||.||||||||||||||||.|||||||||||||||||||||||||||...|||||.||||.
--------------GGSLRLSCAASGFTFSTYWMHWVRHAPGKGLVWVSRINSDGSTTYADSVKGRFTISRDNAKNTLYLQMNSLRAEDTAVYYCARDRYYGMDVWGQGTTVTVS-
                 ^^^^^^^^        ^^^^^^^                        ^^^^^^^^^^
```

Best Alignment of Panobacumab light CDRs chain to a sequence from OAS

```
DVVMTQSPLSLPVTLGQPASISCRSSQSLVYSDGNTYLNWFQQRPGQSPRRLIYKVSNRDSGVPDRFSGSGSGTDFTLKISRVEAEDVGVYYCMQGTHWPLTFGGGTKVEIK
|||||||||||||||||||||||||||||||||||||||||||||||||||||||||||||||||||||||||||||||||||||||||.||||||||||||||.|||.|||
DVVMTQSPLSLPVTLGQPASISCRSSQSLVYSDGNTYLNWFQQRPGQSPRRLIYKVSNRDSGVPDRFSGSGSGTDFTLKISRVEAEDVGIYYCMQGTHWPLTFGQGTKPEIK
                  ^^^^^^^^^^^        ^^^                    ^^^^^^^^^
```

Best Alignment of Panobacumab CDR-H3 chain to a sequence from OAS

```
EEQVVESGGGFVQPGGSLRLSCAASGFTF--SPYWMHWVRQAPGKGLVWVSRINSDGSTYYADSVKGRFTISRDNARNTLYLQMNSLRAEDTAVYYCARDRYYGPEMWGQGTMVTVSS
..|..|||.|.|.|...|.|.|..||.......|...|.||..||||.|..||...|||.|..|.|.|.|||.|...|...|...|..|.||||||||||||||||..|||||.|||||
QVQLQESGPGLVKPSQTLSLTCTVSGGSISSGSYYWSWIRQPAGKGLEWIGRIYTSGSTNYNPSLKSRVTISVDTSKNQFSLKLSSVTAADTAVYYCARDRYYGPLDWGQGTLVTVSS
                 ^^^^^^^^^^        ^^^^^^^                        ^^^^^^^^^^
```

Therapeutic : Parsatuzumab

Best Alignment of Parsatuzumab heavy chain to a sequence from OAS

```
EVQLVESGGGLVQPGGSLRLSCAASGYTFIDYYMNWVRQAPGKGLEWVGDINLDNSGTHYNQKFKGRFTISRDKSKNTAYLQMNSLRAEDTAVYYCAREGVYHD-----YDDYAMDYWGQGTLVTVSS
||||||||||||||||||||||||||.||..|.|.||||||||||.||..||.|.|.|.|....|||||||||..|||.|||||||||||||||||||||.|.|.....||....|||||||||||||
EVQLVESGGGLVQPGGSLRLSCAASGFTFSSYWMHWVRQAPGKGLVWVSRINSDGSSTSYADSVKGRFTISRDNAKNTLYLQMNSLRAEDTAVYYCAREGTYYDFWSGYYDRGCSDYWGQGTLVTVSS
                 ^^^^^^^^        ^^^^^^^^                        ^^^^^^^^^^^^^^^^^^^^^
```

Best Alignment of Parsatuzumab light chain to a sequence from OAS

```
DIQMTQSPSSLSASVGDRVTITCRTSQSLVHINAITYLHWYQQKPGKAPKLLIYRVSNRFSGVPSRFSGSGSGTDFTLTISSLQPEDFATYYCGQSTHVPLTFGQGTKVEIK
||||||||||||||||||||||||.||||......|||.|||||||||||||||..|...||||||||||||||||||||||||||||||||.||...|||||||||||||
DIQMTQSPSSLSASVGDRVTITCRASQSL-----KTYLNWYQQKPGKAPKLLIYAASTLQSGVPSRFSGSGSGTDFTLTISSLQPEDFATYYCQQSYSTPLTFGQGTKVEIK
                  ^^^^^^^^^^^        ^^^                    ^^^^^^^^^
```

Best Alignment of Parsatuzumab heavy chain CDRs to a sequence from OAS

```
EVQLVESGGGLVQPGGSLRLSCAASGYTFIDYYMNWVRQAPGKGLEWVGDINLDNSGTHYNQKFKGRFTISRDKSKNTAYLQMNSLRAEDTAVYYCAREGVYHDYDDYAMDYWGQGTLVTVSS
||||..||..||.||.|...||.||||||.|||||||.|..||.|||.||||..|.||.|||||||..|...|||..|||....||..||.|||||||||.|.||||||||||||||.|||||
EVQLQQSGPELVKPGASVKISCKASGYTFTDYYMNWVKQSHGKSLEWIGDINPNNGGTSYNQKFKGKATLTVDKSSSTAYMELRSLTSEDSAVYYCAREGGYYDYDDYAMDYWGQGTSVTVSS
                 ^^^^^^^^        ^^^^^^^^                        ^^^^^^^^^^^^^^^^
```

Best Alignment of Parsatuzumab light CDRs chain to a sequence from OAS

```
DIQMTQSPSSLSASVGDRVTITCRTSQSLVHINAITYLHWYQQKPGKAPKLLIYRVSNRFSGVPSRFSGSGSGTDFTLTISSLQPEDFATYYCGQSTHVPLTFGQGTKVEIK
|..|||||.||....|....|.||.||||||.|..|||.||||||...|..|||.||||.||.|.||||||.||||||.||....||...||||||||||||||.|||||||
DVVMTQSPLSLPITPGQPASISCRSSQSLVHSNGNTYLNWYQQKPCQPPRRLIYQVSNRDSGGPDRFSGSGAGTDFTLNISRVESEDVGVYYCGQSTHVPLTFGGGTKVEIK
                  ^^^^^^^^^^^        ^^^                    ^^^^^^^^^
```

Best Alignment of Parsatuzumab CDR-H3 chain to a sequence from OAS

```
EVQLVESGGGLVQPGGSLRLSCAASGYTFIDYYMNWVRQAPGKGLEWVGDINLDNSGTHYNQKFKGRFTISRDKSKNTAYLQMNSLRAEDTAVYYCAREGVYHDYDDYAMDYWGQGTLVTVSS
||.|||||||||||||..|||||||.||....|.||||||.||||||..|....|...|....|||||||||..|||..|||.|||.||||.|||||||||.||||||||||||||.|||||
```

```
EVMLVESGGGLVQPGGSRKLSCAASGFTFSSFGMHWVRQAPEKGLEWVAYISSGSSTIYYADTVKGRFTISRDNPKNTLFLQMTSLRSEDTAMYYCAREGVYYDYDDYAMDYWGQGTSVTVSS
                      ^^^^^^^^              ^^^^^^^^                                    ^^^^^^^^^^^^^^^^
```

Therapeutic : Pateclizumab

Best Alignment of Pateclizumab heavy chain to a sequence from OAS

```
EVQLVESGGGLVQPGGSLRLSCAASGYTFTSYVIHWVRQAPGKGLEWVGYNNPYNAGTNYNEKFKGRFTISSDKSKNTAYLQMNSLRAEDTAVYYCSRPTML---------PWFAYWGQGTLVTVSS
|||||||||||||||||||||||||.||.||..|||||||||||.||...|.....|.|....||||||.|.||||.||||||||||||||||.|...|.........|.|.||||||||||||
EVQLVESGGGLVQPGGSLRLSCAASGFTFSSYWMHWVRQAPGKGLVWVSRINSDGSSTSYADSVKGRFTISRDNSKNTLYLQMNSLRAEDTAVYYCARYGPLEGVAAATQVPYFDYWGQGTLVTVSS
                      ^^^^^^^^              ^^^^^^^^                                    ^^^^^^^^^^^^^^^^^^^^
```

Best Alignment of Pateclizumab light chain to a sequence from OAS

```
DIQMTQSPSSLSASVGDRVTITCRASQAVSSAVAWYQQKPGKAPKLLIYSASHRYTGVPSRFSGSGSGTDFTLTISSLQPEDFATYYCQESYSTPWTFGQGTKVEIK
||||||||||||||||||||||||||..|||.||||||||||||||||.||....||||||||||||||||||||||||||||||||.|||||||||||||||||
DIQMTQSPSSLSASVGDRVTITCRASQNISSALAWYQQKPGKAPKLLIYDASSLESGVPSRFSGSGSGTDFTLTISSLQPEDFATYYCQQSYSTPWTFGQGTKVEIK
                       ^^^^^^             ^^^                                  ^^^^^^^^^
```

Best Alignment of Pateclizumab heavy chain CDRs to a sequence from OAS

```
EVQLVESGGGLVQPGGSLRLSCAASGYTFTSYVIHWVRQAPGKGLEWVGYNNPYNAGTNYNEKFKGRFTISSDKSKNTAYLQMNSLRAEDTAVYYCSRPTMLPWFAYWGQGTLVTVSS
||||..||..||.||.|...||.||||||||||.|||.|.||.||||.||.||||.||.|||||||..|..||||..|||....||..||.|||||.||||..|||||||||||||.
EVQLQQSGPELVKPGASVKMSCKASGYTFTSYVMHWVKQKPGQGLEWIGYINPYNDGTKYNEKFKGKATLTSDKSSSTAYMELSSLTSEDSAVYYCARPTMTTWFAYWGQGTLVTVSA
                      ^^^^^^^^              ^^^^^^^^                                    ^^^^^^^^^^^
```

Best Alignment of Pateclizumab light CDRs chain to a sequence from OAS

```
DIQMTQSPSSLSASVGDRVTITCRASQAVSSAVAWYQQKPGKAPKLLIYSASHRYTGVPSRFSGSGSGTDFTLTISSLQPEDFATYYCQESYSTPWTFGQGTKVEIK
|||||||||||||||||||.||||||.||||.|||.|||||||||||||||....||.||||||||||||||||||||||||||||||||||||.|||.|||||||
DIQMTQSPSSLSASVGDRVTVTCRASQGVSSALAWYHQKPGKAPKLLIYSASTLESGVASRFSGSGSGTDFTLTISSLQPEDFATYYCQESYSTPHTFGGGTKVEIK
                       ^^^^^^             ^^^                                  ^^^^^^^^^
```

Best Alignment of Pateclizumab CDR-H3 chain to a sequence from OAS

```
EVQLVESGGGLVQPGGSLRLSCAASGYTFTSYVIHWVRQAPGKGLEWVGYNNPYNAGTNYNEKFKGRFTISSDKSKNTAYLQMNSLRAEDTAVYYCSRPTMLPWFAYWGQGTLVTVSS
.|||..||..|..||.|...||.|.||||.||.|.||.|.||.||||.|...|....|||||||||..|...|.|.||||.|..||..||.|||||||.||.||||||||||||||.
-VQLQQSGAELMKPGASVKISCKATGYTFSSYWIEWVKQRPGHGLEWIGEILPGSGSTNYNEKFKGKATFTADTSSNTAYMQLSSLTSEDSAVYYCSRSTMIPWFAYWGQGTLVTVSA
                      ^^^^^^^^              ^^^^^^^^                                    ^^^^^^^^^^^
```

Therapeutic : Patritumab

Best Alignment of Patritumab heavy chain to a sequence from OAS

```
QVQLQQWGAGLLKPSETLSLTCAVYGGSFSGYYWSWIRQPPGKGLEWIGEINHSGSTNYNPSLKSRVTISVETSKNQFSLKLSSVTAADTAVYYCARDKWT-------WYFDLWGRGTLVTVSS
||||||||||||||||||||||||||||||||||||||||||||||||||||||||||||||||||||||||.||||||||||||||||||||||||||||.......||||||||||||||||
QVQLQQWGAGLLKPSETLSLTCAVYGGSFSGYYWSWIRQPPGKGLEWIGEINHSGSTNYNPSLKSRVTISVDTSKNQFSLKLSSVTAADTAVYYCARDKWTSNWGPRYWYFDLWGRGTLVTVSS
                      ^^^^^^^^              ^^^^^^^                                    ^^^^^^^^^^^^^^^^^^
```

Best Alignment of Patritumab light chain to a sequence from OAS

```
DIEMTQSPDSLAVSLGERATINCRSSQSVLYSSSNRNYLAWYQQNPGQPPKLLIYWASTRESGVPDRFSGSGSGTDFTLTISSLQAEDVAVYYCQQYYSTPRTFGQGTKVEIK
||.||||||||||||||||||||||||||||||.||||||||||.||||||||||||||||||||||||||||||||||||||||||||||||||||||||||||||||||||
DIVMTQSPDSLAVSLGERATINCRSSQSVLYSSDNRNYLAWYQQKPGQPPKLLIYWASTRESGVPDRFSGSGSGTDFTLTISSLQAEDVAVYYCQQYYSTPRTFGQGTKVEIK
                       ^^^^^^^^^^^^             ^^^                                  ^^^^^^^^^
```

Best Alignment of Patritumab heavy chain CDRs to a sequence from OAS

```
QVQLQQWGAGLLKPSETLSLTCAVYGGSFSGYYWSWIRQPPGKGLEWIGEINHSGSTNYNPSLKSRVTISVETSKNQFSLKLSSVTAADTAVYYCARDKWTWYFDLWGRGTLVTVSS
..............||||||||||||||||||.||||||||||||||||||||||||||||||||||||||.||||||||||||||||||||||||||||.|.||||||||||||||||
--------------ETLSLTCAVYGGSFSGYYWRWIRQPPGKGLEWIGEINHSGSTNYNPSLKSRVTISVDTSKNQFSLKLSSVTAADTAVYYCARDSWYWYFDLWGRGTLVTVSS
                      ^^^^^^^^              ^^^^^^^                                    ^^^^^^^^^^^
```

Best Alignment of Patritumab light CDRs chain to a sequence from OAS

```
DIEMTQSPDSLAVSLGERATINCRSSQSVLYSSSNRNYLAWYQQNPGQPPKLLIYWASTRESGVPDRFSGSGSGTDFTLTISSLQAEDVAVYYCQQYYSTPRTFGQGTKVEIK
||.|||||||||||||||||.||.|||||||||.||||||||||.|||.|||||.||||||||||||||||||||||||||||||||||||||||||||||||||||||.|||
DIVMTQSPDSLAVSLGERATVNCKSSQSVLYSSNNRNYLAWYQQRPGQSPKLLISWASTRESGVPDRFSGSGSGTDFTLTISSLQAEDVAVYYCQQYYSTPRTFGQGTKLEIK
                       ^^^^^^^^^^^^             ^^^                                  ^^^^^^^^^
```

Best Alignment of Patritumab CDR-H3 chain to a sequence from OAS

```
QVQLQQWGAGLLKPSETLSLTCAVYGGSFSGYYWSWIRQPPGKGLEWIGEINHS-GSTNYNPSLKSRVTISVETSKNQFSLKLSSVTAADTAVYYCARDKWTWYFDLWGRGTLVTVSS
||||.|.||...||.......|...|..|..||..|.||..|.||||.|.||...|.|.|......|||....||.......||.....|||||||||||||.||||||||||||||||
QVQLVQSGAEVKKPGASVKVSCKASGYTFTAYYLHWVRQASGQGLEWMGWINPNNGDTIYAQNFQGRVTMTRDTSISTAYMELSNLRSDDTAVYYCARDKWFWYFDLWGRGTLVTVSS
                      ^^^^^^^^              ^^^^^^^^                                    ^^^^^^^^^^^
```

Therapeutic : Pembrolizumab

Best Alignment of Pembrolizumab heavy chain to a sequence from OAS

```
QVQLVQSGVEVKKPGASVKVSCKASGYTFTNYYMYWVRQAPGQGLEWMGGINPSNGGTNFNEKFKNRVTLTTDSSTTTAYMELKSLQFDDTAVYYCARRDYRFDMGFDYWGQGTTVTVSS
||||||||.||||||||||||||||||||||.|||.|||||||||||||||.|||..||||...|...||||.|||.||.||||||.||..|||||||||||.||..|.|.||||||||.|||||
QVQLVQSGAEVKKPGASVKVSCKASGYTFTGYYMHWVRQAPGQGLEWMGRINPNSGGTNYAQKLQGRVTMTTDTSTSTAYMELRSLRSDDTAVYYCARDDYG-DYGGDYWGQGTLVTVSS
                      ^^^^^^^^              ^^^^^^^^                                    ^^^^^^^^^^^^^
```

Best Alignment of Pembrolizumab light chain to a sequence from OAS

```
EIVLTQSPATLSLSPGERATLSCRASKGVSTSGYSYLHWYQQKPGQAPRLLIYLASYLESGVPARFSGSGSGTDFTLTISSLEPEDFAVYYCQHSRDLPLTFGGGTKVEIK
|||||||||||||||||||||||||||.||.....|||.|||||||||||||||||.||....|.||||||||||||||||||||||||||||||||||..|.||||||||||||||
EIVLTQSPATLSLSPGERATLSCRASQGV----RSYLAWYQQKPGQAPRLLIYDASNRATGIPARFSGSGSGTDFTLTISSLEPEDFAVYYCQHRDDWPLTFGGGTKVEIK
                       ^^^^^^^^^^             ^^^                                ^^^^^^^^^
```

Best Alignment of Pembrolizumab heavy chain CDRs to a sequence from OAS

```
QVQLVQSGVEVKKPGASVKVSCKASGYTFTNYYMYWVRQAPGQGLEWMGGINPSNGGTNFNEKFKNRVTLTTDSSTTTAYMELKSLQFDDTAVYYCARRDYRFDMGFDYWGQGTTVTVSS
.|||.|||....|||||||.||||||||||.|||.||.|..|.||||.|.|||.||||..|.|||...|||.|.|..||||||.||...|.|||.||||||.||.||||||||||.||||.
EVQLQQSGPVLVKPGASVKMSCKASGYTFTDYYMNWVKQSHGKGLEWIGVINPYNGGTSSNQKFKGKATLTVDKSSSTAYMELNSLTSEDAAVYSCARRDYGFDGGFDYWGQGTLVTVSA
                      ^^^^^^^^              ^^^^^^^^                                    ^^^^^^^^^^^^^
```

Best Alignment of Pembrolizumab light CDRs chain to a sequence from OAS

```
EIVLTQSPATLSLSPGERATLSCRASKGVSTSGYSYLHWYQQKPGQAPRLLIYLASYLESGVPARFSGSGSGTDFTLTISSLEPEDFAVYYCQHSRDLPLTFGGGTKVEIK
.||||||||.|..|.|.|||.||||||.||||||||.|||||||||.|.|||||||.||||||||||||||||||||.|...|.||.|.|||||||.||||||.|||.|||
DIVLTQSPASLAVSLGQRATISCRASKSVSTSGYSYMHWYQQKPGQPPKLLIYLASNLESGVPARFSGSGSGTDFTLNIHPVEEEDAATYYCQHSRELPLTFGSGTKLEIK
                       ^^^^^^^^^^             ^^^                                ^^^^^^^^^
```

Best Alignment of Pembrolizumab CDR-H3 chain to a sequence from OAS

```
QVQLVQSGVEVKKPGASVKVSCKASGYTFTNYYMYWVRQAPGQGLEWMGGINPSNGGTNFNEKFKNRVTLTTDSSTTTAYMELKSLQFDDTAVYYCARRDYRFDMGFDYWGQGTTVTVSS
.|.||.||.....||.|.|.||.|||.||....|.||||||..||||...|............|.|.|...|....|......||...|||.|||||||||.|.||||||||||.||||
EVKLVESGGGLVQPGGSRKLSCAASGFTFSSFGMHWVRQAPEKGLEWVAYISSGSSTIYYADTVKGRFTISRDNPKNTLFLQMTSLRSEDTAMYYCARRDYRYDVGFDYWGQGTTLTVSS
                      ^^^^^^^^              ^^^^^^^^                                    ^^^^^^^^^^^^^
```

Therapeutic : Pertuzumab

Best Alignment of Pertuzumab heavy chain to a sequence from OAS

```
EVQLVESGGGLVQPGGSLRLSCAASGFTFTDYTMDWVRQAPGKGLEWVADVNPNSGGSIYNQRFKGRFTLSVDRSKNTLYLQMNSLRAEDTAVYYCARNLGP---SFYFDYWGQGTLVTVSS
||||||||||||||.||||||||||||.||.|.|||||||||||||.....|||...|....|||||.|.|.||||||||||||||||||||||||.||.....|||||||||||||||
EVQLVESGGGLVQPGRSLRLSCAASGFTFDDYAMHWVRQAPGKGLEWVSGISWNSGSIGYADSVKGRFTISRDNSKNTLYLQMNSLRAEDTAVYYCARDLGELELLFYFDYWGQGTLVTVSS
                      ^^^^^^^^              ^^^^^^^^                                    ^^^^^^^^^^^^^^^
```

```
Best Alignment of Pertuzumab light chain to a sequence from OAS
DIQMTQSPSSLSASVGDRVTITCKASQDVSIGVAWYQQKPGKAPKLLIYSASYRYTGVPSRFSGSGSGTDFTLTISSLQPEDFATYYCQQYYIYPYTFGQGTKVEIK
|||||||||||||||||||||||.|||...|...|||||||||||.||||||...||||||||||||||||||||||||||||||||||||||||||||||||||.|||
DIQMTQSPSSLSASVGDRVTITCRASQVIRIDLGWYQQKPGKAPKRLIYSASSLQTGVPSRFSGSGSGTDFTLTISSLQPEDFATYYCQQYYIYPYTFGQGTKLEIK
                       ^^^^^^             ^^^                        ^^^^^^^^^
```

```
Best Alignment of Pertuzumab heavy chain CDRs to a sequence from OAS
EVQLVESGGGLVQPGGSLRLSCAASGFTFTDYTMDWVRQAPGKGLEWVADVNPNSGGSIYNQRFKGRFTLSVDRSKNTLYLQMNSLRAEDTAVYYCARNLGPSFYFDYWGQGTLVTVSS
||||..||..||.||.|...||.|||.|||||.|.||.|..||.|||....|||.||..|||.|||..||.|..|..|.|....||..||.||||||||.|.|||||||||||..||||
EVQLQQSGPELVKPGASVKMSCKASGYTFTDYNMHWVKQSHGKSLEWIGYINPNNGGTSYNQKFKGKATLTVNKSSSTAYMELRSLTSEDSAVYYCARNYGSSFYFDYWGQGTTLTVSS
                      ^^^^^^^^           ^^^^^^^^                         ^^^^^^^^^^^^
```

```
Best Alignment of Pertuzumab light CDRs chain to a sequence from OAS
DIQMTQSPSSLSASVGDRVTITCKASQDVSIGVAWYQQKPGKAPKLLIYSASYRYTGVPSRFSGSGSGTDFTLTISSLQPEDFATYYCQQYYIYPYTFGQGTKVEIK
.........|||||||||||.|..|.||||...|..|||||||||....|||....||||.|||.||||||.|||..|||||||||||||||.|||||||||||.|||
---------SLSASVGDRVTLTWRAGQDVSNYLALFQQKPGKAPKSVSSSASTLRSGVPSKFSGGGSGTDFALTITGLQPEDFATYYCQQYYTYPYTFGQGTKLEIK
                       ^^^^^^             ^^^                        ^^^^^^^^^
```

```
Best Alignment of Pertuzumab CDR-H3 chain to a sequence from OAS
EVQLVESGGGLVQPGGSLRLSCAASGFTFTDYTMDWVRQAPGKGLEWVADVNPNSGGSIYNQRFKGRFTLSVDRSKNTLYLQMNSLRAEDTAVYYCARNLGPSFYFDYWGQGTLVTVSS
.............|||||||||.||||||....|.||||||||||||||..........|....|||||.|.|..||.|||||||||||||||||||||||||||.|||||||||||||||
-------------PGGSLRLSCEASGFTFSNDFMTWVRQAPGKGLEWVANMRVDGSDIHYVDSVKGRFTISRDNAKNSLYLQMNSLRAEDTAVYYCARNLGPSYYFDYWGQGTLVTVSS
                      ^^^^^^^^           ^^^^^^^^                         ^^^^^^^^^^^^
```

Therapeutic : Pidilizumab

```
Best Alignment of Pidilizumab heavy chain to a sequence from OAS
QVQLVQSGSELKKPGASVKISCKASGYTFTNYGMNWVRQAPGQGLQWMGWINTDSGESTYAEEFKGRFVFSLDTSVNTAYLQITSLTAEDTGMYFCVRVGY---------DALDYWGQGTLVTVSS
||||||||||||||||||||||||||||||||.|||||||||||||.|||||||..|..|||..|.||||||||||||||||||.||.||||..|.|.|||.........||.|.|||||.|||||
QVQLVQSGSELKKPGASVKISCKASGYTFTNYAMNWVRQAPGQGLEWMGWINTNTGNPTYAQGFTGRFVFSLDTSVNTAYLQISSLKAEDTAFYYCARVGRFLEWFQTKDDAFDIWGQGTMVTVSS
                      ^^^^^^^^           ^^^^^^^^                         ^^^^^^^^^^^^^^^^^^^
```

```
Best Alignment of Pidilizumab light chain to a sequence from OAS
EIVLTQSPSSLSASVGDRVTITCSARSSVSYMHWFQQKPGKAPKLWIYRTSNLASGVPSRFSGSGSGTSYCLTINSLQPEDFATYYCQQRSSFPLTFGGGTKLEIK
.|||||||...|||.|..|||||||.||||||||||||||..||||||.||||||||||||||||||||||||.|||||...||.|||||||||||.|||||.|||||.|
-IVLTQSPAIMSASPGEKVTITCSASSSVSYMHWFQQKPGTSPKLWIYSTSNLASGVPSRFSGSGSGTSYSLTINSMEAEDAATYYCQQRSSYPLTFGAGTKLELK
                       ^^^^^             ^^^                        ^^^^^^^^^
```

```
Best Alignment of Pidilizumab heavy chain CDRs to a sequence from OAS
QVQLVQSGSELKKPGASVKISCKASGYTFTNYGMNWVRQAPGQGLQWMGWINTDSGESTYAEEFKGRFVFSLDTSVNTAYLQITSLTAEDTGMYFCVRVGYDALDYWGQGTLVTVSS
|.||||||.||||||..|||||||||||||||||||||.||||.||.|||||||..||.|||..|||||.|||.||..||||||..|..||...|||.|.||||.||||||||.|||||
QIQLVQSGPELKKPGETVKISCKASGYTFTNYGMNWVKQAPGKGLKWMGWINTYTGEPTYADDFKGRFAFSLETSASTAYLQINNLKNEDMATYFCARRGYDAMDYWGQGTSVTVSS
                      ^^^^^^^^           ^^^^^^^^                         ^^^^^^^^^^
```

```
Best Alignment of Pidilizumab light CDRs chain to a sequence from OAS
EIVLTQSPSSLSASVGDRVTITCSARSSVSYMHWFQQKPGKAPKLWIYRTSNLASGVPSRFSGSGSGTSYCLTINSLQPEDFATYYCQQRSSFPLTFGGGTKLEIK
..||||.|...|||.|..|||||||.||||||||||.||||||..||||||||||||||||||.||||||||||||.|||.....||.|||||||||||.|||||.|||||||
QMVLTQPPQIMSASPGEKVTITCSASSSVSYMHWYQQKPGTSPKLWIYRTSNLASGVPARFSGSGSGTSYSLTISRMEAEDAATYYCQQRSSYPLTFGAGTKLEIK
                       ^^^^^             ^^^                        ^^^^^^^^^
```

```
Best Alignment of Pidilizumab CDR-H3 chain to a sequence from OAS
QVQLVQSGSELKKPGASVKISCKASGYTF-TNYGMNWVRQAPGQGLQWMGWINTDSGESTYAEEFKGRFVFSLDTSVNTAYLQITSLTAEDTGMYFCVRVGYDALDYWGQGTLVTVSS
......................|..||..........||||.||.||.|.|.|....|...|....|.|...|.|.|.|...|...|.||.||..|.|||.||||||||||||||||||
-------------SETLSLTCAVSGGFMSSGDWWSWVRQPPGKGLEWFGEIHHR-GSTNYHPSPKRRVTISVDQSKNQFPLTVNSVTAADTASYDCVRNGYDALDYWGQGTLVTVSS
                      ^^^^^^^^^           ^^^^^^^^                         ^^^^^^^^^^
```

Therapeutic : Pinatuzumab

```
Best Alignment of Pinatuzumab heavy chain to a sequence from OAS
EVQLVESGGGLVQPGGSLRLSCAASGYEFSRSWMNWVRQAPGKGLEWVGRIYPGDGDTNYSGKFKGRFTISADTSKNTAYLQMNSLRAEDTAVYYCARDGSSW--DWYFDVWGQGTLVTVSS
|||||||||||||||||||||||||||..|||..|||||||||||||||..||.|.|||||.|..||||||||.|.||||.|||||||||||||||||.|||||........||||||||||||||
EVQLVESGGGLVQPGGSLRLSCAASGFTFSRNAMNWVRQAPGKGLEWVSLIYSG-GDTNYAGSVKGRFTISRDGSKNTLYLQMNSLRAEDTAVYFCARDGGRDGYNLAFDVWGQGTLVTVSS
                      ^^^^^^^^           ^^^^^^^^                         ^^^^^^^^^^^^^^^
```

```
Best Alignment of Pinatuzumab light chain to a sequence from OAS
DIQMTQSPSSLSASVGDRVTITCRSSQSIVHSVGNTFLEWYQQKPGKAPKLLIYKVSNRFSGVPSRFSGSGSGTDFTLTISSLQPEDFATYYCFQGSQFPYTFGQGTKVEIK
||||||||||||||||||||||||.||||.....|..|.||||||||||||||||||||...|||||||||||||||||||||||||||||||||||||.|...||.||||||||||||
DIQMTQSPSSLSASVGDRVTITCRASQSI-----NNYLNWYQQKPGKAPKLLIYKVSNLQNGVPSRFSGSGSGTDFTLTISSLQPEDFATYYCQQCYNFPWTFGQGTKVEIK
                       ^^^^^^^^^^^           ^^^                        ^^^^^^^^^
```

```
Best Alignment of Pinatuzumab heavy chain CDRs to a sequence from OAS
EVQLVESGGGLVQPGGSLRLSCAASGYEFSRSWMNWVRQAPGKGLEWVGRIYPGDGDTNYSGKFKGRFTISADTSKNTAYLQMNSLRAEDTAVYYCARDGSSWDWYFDVWGQGTLVTVSS
.|||..||..||.||.|...||.||||.||.||||||.|.|||||||.||||||||||||||.|||||..|..||.|..|||.|..||..||.|||.|||||||.||||||||.||.|||||
QVQLQQSGPELVKPGASVKISCKASGYAFSSSWMNWVKQRPGKGLEWIGRIYPGDGDTNYNGKFKGKATLTADKSSSTAYMQLSSLTSEDSAVYFCARDGSSYDWYFDVWGTGTTVTVSS
                      ^^^^^^^^           ^^^^^^^^                         ^^^^^^^^^^^^^
```

```
Best Alignment of Pinatuzumab light CDRs chain to a sequence from OAS
DIQMTQSPSSLSASVGDRVTITCRSSQSIVHSVGNTFLEWYQQKPGKAPKLLIYKVSNRFSGVPSRFSGSGSGTDFTLTISSLQPEDFATYYCFQGSQFPYTFGQGTKVEIK
|..|||.|.||..|.||...|.||||||||||.|||.||||.||||..||||||||||||||||.|||||||||||||.||....||...||||||||.|||||.|||.|||
DVLMTQTPLSLPVSLGDQASISCRSSQSIVHSNGNTYLEWYLQKPGQSPKLLIYKVSNRFSGVPDRFSGSGSGTDFTLKISRVEAEDLGVYYCFQGSQVPYTFGGGTKLEIK
                       ^^^^^^^^^^^           ^^^                        ^^^^^^^^^
```

```
Best Alignment of Pinatuzumab CDR-H3 chain to a sequence from OAS
EVQLVESGGGLVQPGGSLRLSCAASGYEFSRSWMNWVRQAPGKGLEWVGRIYPGDGDTNYSGKFKGRFTISADTSKNTAYLQMNSLRAEDTAVYYCARDGSSWDWYFDVWGQGTLVTVSS
||.|||||||||.|||||.|||||||..||...|.||||||.||||||..|..|.....|....|||||||.|..|||..|||.|||.||||.||||||||||||||||||||.||.|||||
EVMLVESGGGLVKPGGSLKLSCAASGFTFSDYGMHWVRQAPEKGLEWVAYISSGSSTIYYADTVKGRFTISRDNAKNTLFLQMTSLRSEDTAMYYCARDGSSWDWYFDVWGTGTTVTVSS
                      ^^^^^^^^           ^^^^^^^^                         ^^^^^^^^^^^^^
```

Therapeutic : Plozalizumab

```
Best Alignment of Plozalizumab heavy chain to a sequence from OAS
EVQLVESGGGLVKPGGSLRLSCAASGFTFSAYAMNWVRQAPGKGLEWVGRIRTKNNNYATYYADSVKDRFTISRDDSKNTLYLQMNSLKTEDTAVYYCTTFY----------GNGVWGQGTLVTVSS
|||||||||||||||||||||||||||||||.|.|||||||.|||||||||||.|.|.|||.||.|||.||||||||||||||||||||||||||||||||||||||............|..||||||.|||||
EVQLVESGGGLVKPGGSLRLSCAASGFTFSSYSMNWVRQASGKGLEWVGRIRSKANSYATAYAASVKGRFTISRDDSKNTLYLQMNSLKTEDTAVYYCTTDPPYSSDYYYYYGMDVWGQGTTVTVSS
                      ^^^^^^^^           ^^^^^^^^^^                         ^^^^^^^^^^^^^^^^^^
```

```
Best Alignment of Plozalizumab light chain to a sequence from OAS
DVVMTQSPLSLPVTLGQPASISCKSSQSLLDSDGKTFLNWFQQRPGQSPRRLIYLVSKLDSGVPDRFSGSGSGTDFTLKISRVEAEDVGVYYCWQGTHFPYTFGQGTRLEIK
|||||||||||||||||||||||.|||||..|||.|.|||||||||||||||||.|||.|||||||||||||||||||||||||||||||||||||.||||.|.||||||||||||
DVVMTQSPLSLPVTLGQPASISCRSSQSLVYSDGNTYLNWFQQRPGQSPRRLIYKVSKWDSGVPDRFSGSGSGTDFTLKISRVEAEDVGVYYCMQGTHWPPTFGQGTRLEIK
                       ^^^^^^^^^^^           ^^^                        ^^^^^^^^^
```

```
Best Alignment of Plozalizumab heavy chain CDRs to a sequence from OAS
EVQLVESGGGLVKPGGSLRLSCAASGFTFSAYAMNWVRQAPGKGLEWVGRIRTKNNNYATYYADSVKDRFTISRDDSKNTLYLQMNSLKTEDTAVYYCTTFYGNGVWGQGTLVTVSS
.|.|||||||||.||||..|||.|||||||.|.||||||.|.||||||..||.|.|||||.||.|||.||||||||||...|||||.|..|||..||||..|||.|||||||||||.
```

```
DVKLVESGGGLVQPGGSMKLSCVASGFTFSNYWMNWVRQSPEKGLEWVAEIRLKSNNYATHYAESVKGRFTISRDDSKSSVYLQMNNLRAEDTGIYYCTRTYGNYVWGQGTLVTVSA
                    ^^^^^^^^          ^^^^^^^^^^                    ^^^^^^^^
```

Best Alignment of Plozalizumab light CDRs chain to a sequence from OAS

```
DVVMTQSPLSLPVTLGQPASISCKSSQSLLDSDGKTFLNWFQQRPGQSPRRLIYLVSKLDSGVPDRFSGSGSGTDFTLKISRVEAEDVGVYYCWQGTHFPYTFGQGTRLEIK
||||.|.||.|.||.||||||||||||||||||||||||||..|||||||.||||||||||||||||.||||||||||||||||||.||||||||||||||||.||.||||
DVVMPQTPLTLSVTIGQPASISCKSSQSLLDSDGKTFLNWLLQRPGQSPKRLIYLVSKLDSGVPDRFTGSGSGTDFTLKISRVEAEDLGVYYCWQGTHFPYTFGGGTKLEIK
                     ^^^^^^^^^^^          ^^^                    ^^^^^^^^^
```

Best Alignment of Plozalizumab CDR-H3 chain to a sequence from OAS

```
EVQLVESGGGLVKPGGSLRLSCAASGFTFSAYAMNWVRQAPGKGLEWVGRIRTKNNNYATYYADSVKDRFTISRDDSKNTLYLQMNSLKTEDTAVYYCTTFYGNGVWGQGTLVTVSS
.|||.|||..||.||.|..|||.|||||.....|.||.|.|.|||||.|.|.....|..|.||.......||..|.|.||.|||..||..|||||||||||||||.|||||..||||
DVQLQESGAELVRPGASVKLSCTASGFTIKDDYMHWVKQRPEKGLEWIGWIDPE--NGDTEYASKFQGKATITADTSSNTAYLQLSSLTSEDTAVYYCTTFYGNGYWGQGTTLTVSS
                    ^^^^^^^^          ^^^^^^^^^^                    ^^^^^^^^
```

Therapeutic : Polatuzumab

Best Alignment of Polatuzumab heavy chain to a sequence from OAS

```
EVQLVESGGGLVQPGGSLRLSCAASGYTFSSYWIEWVRQAPGKGLEWIGEILPGGGDTNYNEIFKGRATFSADTSKNTAYLQMNSLRAEDTAVYYCTRRVP----IRLDYWGQGTLVTVSS
|||||||||||||||||||||||||||.||||||..|||||||||||||...|...||.|.|....|||.|.|.|.||||.||||||||||||||||||||..|.....|.|||||||||||||
EVQLVESGGGLVQPGGSLRLSCAASGFTFSSYWMSWVRQAPGKGLEWVSAISGSGGSTYYADSVKGRFTISRDNSKNTLYLQMNSLRAEDTAVYYCTRQGPTGTHQRDDYWGQGTLVTVSS
                    ^^^^^^^^          ^^^^^^^^                   ^^^^^^^^^^^^^^
```

Best Alignment of Polatuzumab light chain to a sequence from OAS

```
DIQLTQSPSSLSASVGDRVTITCKASQSVDYEGDSFLNWYQQKPGKAPKLLIYAASNLESGVPSRFSGSGSGTDFTLTISSLQPEDFATYYCQQSNEDPLTFGQGTKVEIK
|||.||||||||||||||||||||.||||......|.||||||||||||||||||||||.|||||||||||||||||||||||||||||||||||||||||||||..|||||||||||||
DIQMTQSPSSLSASVGDRVTITCRASQSI----SSYLNWYQQKPGKAPKLLIYAASSLESGVPSRFSGSGSGTDFTLTISSLQPEDFATYYCQQSNSTPLTFGQGTKVEIK
                     ^^^^^^^^^^          ^^^                   ^^^^^^^^^
```

Best Alignment of Polatuzumab heavy chain CDRs to a sequence from OAS

```
EVQLVESGGGLVQPGGSLRLSCAASGYTFSSYWIEWVRQAPGKGLEWIGEILPGGGDTNYNEIFKGRATFSADTSKNTAYLQMNSLRAEDTAVYYCTRRVPIRLDYWGQGTLVTVSS
.|||...|..|..||.|...||.|.|||||||||||||.|.||.||||||||||||.|.|||||.|||.|||.||||.||||.|..||..||.|||||.|.|.||||||||||||||||.
QVQLQQPGAELMKPGASVKISCKATGYTFSSYWIEWVKQRPGHGLEWIGEILPGSGSTNYNEKFKGEATFTADTSSNTAYMQLSSLTSEDSAVYYCAREVGIRLDYWGQGTLVTVSA
                    ^^^^^^^^          ^^^^^^^^                   ^^^^^^^^^^
```

Best Alignment of Polatuzumab light CDRs chain to a sequence from OAS

```
DIQLTQSPSSLSASVGDRVTITCKASQSVDYEGDSFLNWYQQKPGKAPKLLIYAASNLESGVPSRFSGSGSGTDFTLTISSLQPEDFATYYCQQSNEDPLTFGQGTKVEIK
||.||.||..|..|.|....|.||||||||||||||..|||||||||..|||||||||||||.||.|.|||||||||||||||.|.....|..||||||||||||||||||.|||.|.|
DIVLTPSPVTLAVSQGQWAPISCKASQSVDYEGDSNMNWYQQKPGQPPKLLIYAASNLGSGIPARFSGSGSGTDFTLNIHPVEEENAATYYCQQSNEDPLTFGAGTKLELK
                     ^^^^^^^^^^          ^^^                   ^^^^^^^^^
```

Best Alignment of Polatuzumab CDR-H3 chain to a sequence from OAS

```
EVQLVESGGGLVQPGGSLRLSCAASGYTFSSYWIEWVRQAPGKGLEWIGEILPGGGDTNYNEIFKGRATFSADTSKNTAYLQMNSLRAEDTAVYYCTRRVPIRLDYWGQGTLVTVSS
.|||..||..||.||.|...||.||||||..|||.|..|.||.||||||.|.||||.|||||.|||.||..||.|..|||.|..||..||.|.||||||||..||||||||||..||||
QVQLKQSGAELVRPGTSVKMSCKASGYTFTNYWIGWAKQRPGHGLEWIGDIYPGGGYTNYNEKFKGKATLTADKSSSTAYMQFSSLTSEDSAIYYCTRRVLLRLDYWGQGTTLTVSS
                    ^^^^^^^^          ^^^^^^^^                   ^^^^^^^^^^
```

Therapeutic : Ponezumab

Best Alignment of Ponezumab heavy chain to a sequence from OAS

```
QVQLVQSGAEVKKPGASVKVSCKASGYYTEAYYIHWVRQAPGQGLEWMGRIDPATGNTKYAPRLQDRVTMTRDTSTSTVYMELSSLRSEDTAVYYCASLYS---LPVYWGQGTTVTVSS
|||||||||||||||||||||||||||...||||||||||||||||||||||.|.||.|.||...|.||||||||||||||||||||||||||||||||||..........|||||||||||
QVQLVQSGAEVKKPGASVKVSCKASGYTFTAYYIHWVRQAPGQGLEWMGRIIPITGITNYAQKFQGRVTMTRDTSTSTVYMELSSLRSEDTAVYYCASALTTWGGMDVWGQGTTVTVSS
                    ^^^^^^^^          ^^^^^^^^                   ^^^^^^^^^^^^
```

Best Alignment of Ponezumab light chain to a sequence from OAS

```
DVVMTQSPLSLPVTLGQPASISCKSSQSLLYSDAKTYLNWFQQRPGQSPRRLIYQISRLDPGVPDRFSGSGSGTDFTLKISRVEAEDVGVYYCLQGTH-YPVLFGQGTRLEIK
||||||||||||||||||||||||.||||||||||..||||||.|||||||||||||||.|..|.||||||||||||||||||||||||||||||||||||.||||..||.||||||||||
DVVMTQSPLSLPVTLGQPASISCRSSQSLLYSDGNTYLNWFHQRPGQSPRRLIYQVSKRDSGVPDRFSGSGSGTDFTLKISRVEAEDVGVYYCMQGTHWPPVAFGQGTRLEIK
                     ^^^^^^^^^^^          ^^^                    ^^^^^^^^^^
```

Best Alignment of Ponezumab heavy chain CDRs to a sequence from OAS

```
QVQLVQSGAEVKKPGASVKVSCKASGYYTEAYYIHWVRQAPGQGLEWMGRIDPATGNTKYAPRLQDRVTMTRDTSTSTVYMELSSLRSEDTAVYYCASLYSLPVYWGQGTTVTVSS
.|||.||.||...||||||.||.|||......|.|||.|.|.|||||.||||||.|||||||..|...|.|.|||..|.|..||||.|||||.|||||||..|||||||||.||||
-VQLQQSVAELVRPGASVKLSCTASGFNIKNTYMHWVKQRPEQGLEWIGRIDPANGNTKYAPKFQGKATITADTSSNTAYLQLSSLTSEDTAIYYCASLYDYPVYWGQGTTLTVSS
                    ^^^^^^^^          ^^^^^^^^                   ^^^^^^^^^
```

Best Alignment of Ponezumab light CDRs chain to a sequence from OAS

```
DVVMTQSPLSLPVTLGQPASISCKSSQSLLYSDAKTYLNWFQQRPGQSPRRLIYQISRLDPGVPDRFSGSGSGTDFTLKISRVEAEDVGVYYCLQGTHYPVLFGQGTRLEIK
||||||.||||.||.|.||||||||||||||||.|||||||.||||||.|..|.||.|.||||.||||||||||.|||||||||||||||.|||||||||.||..||.||.||.|
DVVMTQTPLSLSVTIGKPASISCKSSQSLLYSNAKTYLNWLQQRPGQAPKHLMYQVSKLDPGIPDRFSGSGSETDFTLKISRVEAEDLGVYYCLQGTYYPLTFGAGTKLELK
                     ^^^^^^^^^^^          ^^^                    ^^^^^^^^^
```

Best Alignment of Ponezumab CDR-H3 chain to a sequence from OAS

```
QVQLVQSGAEVKKPGASVKVSCKASGYYTEAYYIHWVRQAPGQGLEWMGRIDPATGNTKYAPRLQDRVTMTRDTSTSTVYMELSSLRSEDTAVYYCASLYSLPVYWGQGTTVTVSS
.....|||||||||||||||||||||....||.|||||||||||||||.|.|..|.|.||...|.|||||||||||||||||||||||||||||||||||||.||.||||||.|||||
-----QSGAEVKKPGASVKVSCKASGYTFTSYYMHWVRQAPGQGLEWMGIINPSGGSTSYAQKFQGRVTMTRDTSTSTVYMELSSLRSEDTAVYYCASLYQLPDYWGQGTLVTVSS
                    ^^^^^^^^          ^^^^^^^^                   ^^^^^^^^^
```

Therapeutic : Pritoxaximab

Best Alignment of Pritoxaximab heavy chain to a sequence from OAS

```
QVQLQESGAELVRSGASVRMSCKASGYTFTSYNMHWVKQTPGQGLEWIGYIYPGNGGTNYIQKFKGKAILTADTSSSTAYMQISSLTSEDSAVYFCTRSPSHYSSDPYFDYWGQGTTLTVSS
|||||.||||||||||||.||||||||||||||||||||||||||||||||||||||||||.|||||||.|||||||||||||||||||||||||||||.|||.......||||||||||||||
QVQLQQSGAELVRSGASVKMSCKASGYTFTSYNMHWVKQTPGQGLEWIGYIYPGNGGTNYNQKFKGKATLTADTSSSTAYMQISSLTSEDSAVYFCARSPPS---WYYFDYWGQGTTLTVSS
                    ^^^^^^^^          ^^^^^^^^                   ^^^^^^^^^^^^^^^
```

Best Alignment of Pritoxaximab light chain to a sequence from OAS

```
DIVMSQSHKFMSTSVGDRVSITCKASQDVGTAVAWYQQNPGQSPKFLIYWASTRHTGVPDRFTGSGSGTDFTLTITNVQSEDLADYFCQQYSSYPLTFGAGTSLELK
||||.|||||||||||||||||||||||||||||||||||||.||||||||||||||||||||||||||||.||||||||||||||||||||||||||||.||||
DIVMTQSHKFMSTSVGDRVSITCKASQDVGTAVAWYQQNPGQSPKLLIYWASTRHTGVPDRFTGSGSGTDFTLTISNVQSEDLADYFCQQYSSYPLTFGAGTKLELK
                     ^^^^^^          ^^^                   ^^^^^^^^^
```

Best Alignment of Pritoxaximab heavy chain CDRs to a sequence from OAS

```
QVQLQESGAELVRSGASVRMSCKASGYTFTSYNMHWVKQTPGQGLEWIGYIYPGNGGTNYIQKFKGKAILTADTSSSTAYMQISSLTSEDSAVYFCTRSPSHYSSDPYFDYWGQGTTLTVSS
|||||.|||||||.||||.||||||||||||||||||||||||||.|||||||.||||||.|.|.||||||||.||.|.|||||||||.||||||||||||||.|||..|.|.||||||||||||||||
QVQLQQSGAELVRPGASVKMSCKASGYTFTSYNMHWVKQTPRQGLEWIGAIYPGNGDTSYNQKFKGKATLTVDKSSSTAYMQLSSLTSEDSAVYFCARSPYYYGSSPYFDYWGQGTTLTVSS
                    ^^^^^^^^          ^^^^^^^^                   ^^^^^^^^^^^^^^^
```

Best Alignment of Pritoxaximab light CDRs chain to a sequence from OAS

```
DIVMSQSHKFMSTSVGDRVSITCKASQDVGTAVAWYQQNPGQSPKFLIYWASTRHTGVPDRFTGSGSGTDFTLTITNVQSEDLADYFCQQYSSYPLTFGAGTSLELK
||||.|||||||||||||||||||||||||||||||||.||||||.||.|||||.|||||||||||||||.||||.||||||||||||||||||||||||.||.||.|
DIVMTQSHKFMSTSVGDRVSITCKASQDVGTAVAWYQQKPGQSPKLLIHWASTRNTGVPDRFTGSGSGTDYTLTISNVQSEDLADYFCQQYSSYPLTFGGGTKLEIK
                     ^^^^^^          ^^^                   ^^^^^^^^^
```

Best Alignment of Pritoxaximab CDR-H3 chain to a sequence from OAS

```
QVQLQESGAELVRSGASVRMSCKASGYTFTSYNMHWVKQTPGQGLEWIGYIYPGNGGTNYIQKFKGKAILTADTSSSTAYMQISSLTSEDSAVYFCTRSPSHYSSDPYFDYWGQGTTLTVSS
...............|.|.||.|||....|..|.|..|.||.||||||.|....|.|||....|.......|||........||.|..|.|||.|.||||.||||.||||||||||..||||
```

```
----------------SLRLSCVASGFSISSHWMRWLRQPPGKGLEWIGEINHS-GSTNYNPSLKSRVTISVDTSKNQFSLKLSSVTAADTAVYYCARSPSPYSSDWYFDYWGQGTLVTVSS
                  ^^^^^^^^          ^^^^^^^^                   ^^^^^^^^^^^^^^^
```

Therapeutic : Quilizumab

Best Alignment of Quilizumab heavy chain to a sequence from OAS

```
EVQLVESGGGLVQPGGSLRLSCAASGFTFSDYGIAWVRQAPGKGLEWVAFISDLAYTIYYADTVTGRFTISRDNSKNTLYLQMNSLRAEDTAVYYCARDNW-DAMDYWGQGTLVTVSS
|||||||||||||||||.||||||||||||||.|||||||.||.|||||||.|||.|||||||||||||||.|.||||||.|.|||.||||.|||||.||.||||||||||.|||||
EVQLVESGGGLVQPGGSLKLSCAASGFTFSDYGMAWVRQAPRKGPEWVAFISNLAYSIYYADTVTGRFTISRENAKNTLYLEMSSLRSEDTAMYYCARRNWDDAMDYWGQGTSVTVSS
                  ^^^^^^^^          ^^^^^^^^                   ^^^^^^^^^^^
```

Best Alignment of Quilizumab light chain to a sequence from OAS

```
DIQMTQSPSSLSASVGDRVTITCRSSQSLVHNNANTYLHWYQQKPGKAPKLLIYKVSNRFSGVPSRFSGSGSGTDFTLTISSLQPEDFATYYCSQNTLVPWTFGQGTKVEIK
||||||||||||||||||||||||.|||......|.||.||||||||||||||||||...|||||||||||||||||||||||||||||||||.|....|||||||||||||
DIQMTQSPSSLSASVGDRVTITCRASQSI-----NNYLNWYQQKPGKAPKLLIYKVSNLQNGVPSRFSGSGSGTDFTLTISSLQPEDFATYYCQQCYNFPWTFGQGTKVEIK
                   ^^^^^^^^^^^         ^^^                   ^^^^^^^^^
```

Best Alignment of Quilizumab heavy chain CDRs to a sequence from OAS

```
EVQLVESGGGLVQPGGSLRLSCAASGFTFSDYGIAWVRQAPGKGLEWVAFISDLAYTIYYADTVTGRFTISRDNSKNTLYLQMNSLRAEDTAVYYCARDNWDAMDYWGQGTLVTVSS
||||||||||||||||..||||||||||||||.||||||||||.|||||||.|||.|||||||||||||||.|.||||||.|.|||.||||.|||||||.|||||.||||.|||||
EVQLVESGGGLVQPGGSRKLSCAASGFTFSDYGMAWVRQAPGKGPEWVAFISNLAYSIYYADTVTGRFTISRENAKNTLYLEMSSLRSEDTAMYYCARDNEDAMDYGGQGTSVTVSS
                  ^^^^^^^^          ^^^^^^^^                   ^^^^^^^^^^
```

Best Alignment of Quilizumab light CDRs chain to a sequence from OAS

```
DIQMTQSPSSLSASVGDRVTITCRSSQSLVHNNANTYLHWYQQKPGKAPKLLIYKVSNRFSGVPSRFSGSGSGTDFTLTISSLQPEDFATYYCSQNTLVPWTFGQGTKVEIK
|..|||.|.||..|.||...|.|||||||||||.|||||||.||||..||||||||||||||||.|||||||||||||.||....||...|||||||.||||||.|||.|||
DVVMTQTPLSLPVSLGDQASISCRSSQSLVHNNGNTYLHWYLQKPGQSPKLLIYKVSNRFSGVPDRFSGSGSGTDFTLKISRVEAEDLGVYYCSQNTYVPWTFGGGTKLEIK
                   ^^^^^^^^^^^         ^^^                   ^^^^^^^^^
```

Best Alignment of Quilizumab CDR-H3 chain to a sequence from OAS

```
EVQLVESGGGLVQPGGSLRLSCAASGFTFSDYGIAWVRQAPGKGLEWVAFISDL--AYTIYYADTVTGRFTISRDNSKNTLYLQMNSLRAEDTAVYYCARDNWDAMDYWGQGTLVTVSS
||.|||||||||||||||||||.|||||||....||||.|||.|||.|........||..|...|.|||..|||.|...||||||.|||||||.||||||||||||||||||.|||||
EVMLVESGGGLVQPGGSLRLSCATSGFTFSDFYMEWVRQPPGKRLEWIAASRNKANDYTTEYSASVKGRFIVSRDTSQSILYLQMNALRAEDTAIYYCARDNWDAMDYWGQGTSVTVSS
                  ^^^^^^^^          ^^^^^^^^^^                   ^^^^^^^^^^
```

Therapeutic : Racotumomab

Best Alignment of Racotumomab heavy chain to a sequence from OAS

```
QVQLQQSGAELVKPGASVKLSCKASGYTFTSYDINWVRQRPEQGLEWIGWIFPGDGSTKYNEKFKGKATLTTDKSSSTAYMQLSRLTSEDSAVYFCAREDYYD-NSYYFDYWGQGTTLTVSS
||||||||||||||||||||||||||||||||||||||||||||||||||||||||||||||||||||||||||||||||||||||||||||||||||||||...||||||||||||||||||
QVQLQQSGAELVKPGASVKLSCKASGYTFTSYDINWVRQRPEQGLEWIGWIFPGDGSTKYNEKFKGKATLTTDKSSSTAYMQLSRLTSEDSAVYFCAREDYYYGSSYYFDYWGQGTTLTVSS
                  ^^^^^^^^          ^^^^^^^^                   ^^^^^^^^^^^^^^^
```

Best Alignment of Racotumomab light chain to a sequence from OAS

```
DIQMTQTTSSLSASLGDRVTISCRASQDISNYLNWYQQKPDGTVKLLIYYTSRLHSGVPSRFSGSGSGTDYSLTISNLEQEDIATYFCQQGNTLPWTFGGGTKLEIK
|||||||||||||||||||||||||||||||||||||||||||||||||||||||||||||||||||||||||||||||||||||||||||||||||||||||||||
DIQMTQTTSSLSASLGDRVTISCRASQDISNYLNWYQQKPDGTVKLLIYYTSRLHSGVPSRFSGSGSGTDYSLTISNLEQEDIATYFCQQGNTLPWTFGGGTKLEIK
                   ^^^^^^          ^^^                 ^^^^^^^^^
```

Best Alignment of Racotumomab heavy chain CDRs to a sequence from OAS

```
QVQLQQSGAELVKPGASVKLSCKASGYTFTSYDINWVRQRPEQGLEWIGWIFPGDGSTKYNEKFKGKATLTTDKSSSTAYMQLSRLTSEDSAVYFCAREDYYDNSYYFDYWGQGTTLTVSS
.|||||||||||||||||||||||||||||||||||||||||||||||||||||||.||||||||||||||||.||.|||||.|||||||||||||||.||||.|||||||||||||||||||
-VQLQQSGAELVKPGASVKLSCKASGYTFTSYDINWVRQRPEQGLEWIGWIFPGDGNTKYNEKFKGKATLTTDESSNTAYMQVSRLTSEDSAVYFCAGEDYYGNSYYFDYWGQGTTLTVSS
                  ^^^^^^^^          ^^^^^^^^                   ^^^^^^^^^^^^^^
```

Best Alignment of Racotumomab light CDRs chain to a sequence from OAS

```
DIQMTQTTSSLSASLGDRVTISCRASQDISNYLNWYQQKPDGTVKLLIYYTSRLHSGVPSRFSGSGSGTDYSLTISNLEQEDIATYFCQQGNTLPWTFGGGTKLEIK
|.|..||||||...|||||||||||||||||||||||||||||||||.||||||||||||||||||||.||||||||||||||||.||||||||||||||||||||||||
DMQLAQTTSSLPGALGDRVTISCRASQDISNYLNWYQQKPDGTVKPLIYYTSRLHSGVPSRFSGSASGTDYSLTISNLEQEAIATYFCQQGNTLPWTFGGGTKLEIK
                   ^^^^^^          ^^^                 ^^^^^^^^^
```

Best Alignment of Racotumomab CDR-H3 chain to a sequence from OAS

```
QVQLQQSGAELVKPGASVKLSCKASGYTFTSYDINWVRQRPEQGLEWIGWIFPGDGSTKYNEKFKGKATLTTDKSSSTAYMQLSRLTSEDSAVYFCAREDYYDNSYYFDYWGQGTTLTVSS
................|..|||.||..||..|..|||||.|..||||...|........|....||..|...|......|.|...|..||.|||.||||||||||.|||||||||..||||
----------------SLRLSCAASALTFSTYSMNWVRQTPGRGLEWVSYISSSSSTIYYADSVKGRFTISRDNAKNSVYLQMNSLRAEDTAVYYCAREDYYDNSGYFDYWGQGTLVTVSS
                  ^^^^^^^^          ^^^^^^^^                   ^^^^^^^^^^^^^^
```

Therapeutic : Radretumab

Best Alignment of Radretumab heavy chain to a sequence from OAS

```
EVQLLESGGGLVQPGGSLRLSCAASGFTFSSFSMSWVRQAPGKGLEWVSSISGSSGTTYYADSVKGRFTISRDNSKNTLYLQMNSLRAEDTAVYYCAKPFP----YFDYWGQGTLVTVSS
||||||||||||||||||||||||||||||..||||||||||||||||.||||.|.|||||||||||||||||||||||||||||||||||||||||||||||....|||||||||||||||
EVQLLESGGGLVQPGGSLRLSCAASGFTFSSYAMSWVRQAPGKGLEWVSAISGSGGSTYYADSVKGRFTISRDNSKNTLYLQMNSLRAEDTAVYYCAKPFPRLMYYFDYWGQGTLVTVSS
                  ^^^^^^^^          ^^^^^^^^                   ^^^^^^^^^^^^^
```

Best Alignment of Radretumab light chain to a sequence from OAS

```
EIVLTQSPGTLSLSPGERATLSCRASQSVSSSFLAWYQQKPGQAPRLLIYYASSRATGIPDRFSGSGSGTDFTLTISRLEPEDFAVYYCQQTGRIPPTFGQGTKVEIK
||||||||||||||||||||||||||||||||.|||||||||||||||||.|||||||||||||||||||||||||||||||||||||||||||||.||.||||||||||||||
EIVLTQSPGTLSLSPGERATLSCRASQSVSSSYLAWYQQKPGQAPRLLIYGASSRATGIPDRFSGSGSGTDFTLTISRLEPEDFAVYYCQQYGRSPPTFGQGTKVEIK
                   ^^^^^^^         ^^^                 ^^^^^^^^^
```

Best Alignment of Radretumab heavy chain CDRs to a sequence from OAS

```
EVQLLESGGGLVQPGGSLRLSCAASGFTFSSFSMSWVRQAPGKGLEWVSSISGSSGTTYYADSVKGRFTISRDNSKNTLYLQMNSLRAEDTAVYYCAKPFPYFDYWGQGTLVTVSS
................||||||||||||||..|||||||||.||||||.||||.|.||||||||||||||||||||||||||||||||||||||||||||||.||||||||||||||||
----------------SLRLSCAASGFTFSSYAMSWVRQAPGRGLEWVSAISGSGGSTYYADSVKGRFTISRDNSKNTLYLQMNSLRAEDTAVYYCAKPSPYFDYWGQGTLVTVSS
                  ^^^^^^^^          ^^^^^^^^                   ^^^^^^^^^
```

Best Alignment of Radretumab light CDRs chain to a sequence from OAS

```
EIVLTQSPGTLSLSPGERATLSCRASQSVSSSFLAWYQQKPGQAPRLLIYYASSRATGIPDRFSGSGSGTDFTLTISRLEPEDFAVYYCQQTGRIPPTFGQGTKVEIK
|||||||||||||||||.||||||||||||||||||||||||||||||..||||||||||||||||||||||.||.|||||||||||||||||.||.||||||||||.|||
EIVLTQSPGTLSLSPGERGTLSCRASQSVSSSFLAWYQQKPGQAPRLLIFGASSRATGIPDRFSGSGSGTDFALTLSRLEPEDFAVYYCQQYGRSPPTFGQGTKLEIK
                   ^^^^^^^         ^^^                 ^^^^^^^^^
```

Best Alignment of Radretumab CDR-H3 chain to a sequence from OAS

```
EVQLLESGGGLVQPGGSLRLSCAASGFTFSSFSMSWVRQAPGKGLEWVSSISGSSGTTYYADSVKGRFTISRDNSKNTLYLQMNSLRAEDTAVYYCAKPFPYFDYWGQGTLVTVSS
.|.|.|||.|||.|..||...|..|||...|...|||||.|||||||...|.|..|.|.|......|..||...||....||..||...|||.||||||||||||||||||..||||
-VKLVESGPGLVAPSQSLSITCIVSGFSLTSYGVSWVRQPPGKGLEWLGVIWGD-GSTSYHSALISRLSISKEKSKSQVFLQLSSLQTDDTATYYCAKPFPYFDYWGQGTTLTVSS
                  ^^^^^^^^          ^^^^^^^^                   ^^^^^^^^^
```

Therapeutic : Rafivirumab

Best Alignment of Rafivirumab heavy chain to a sequence from OAS

```
QVQLVQSGAEVKKPGSSVKVSCKASGGTFNRYTVNWVRQAPGQGLEWMGGIIPIFGTANYAQRFQGRLTITADESTSTAYMELSSLRSDDTAVYFCARENLDNSGTYYYFSGWFDPWGQGTLVTVSS
||||||||||||||||||||||||||||||..|...|||||||||||||||||||||||||||.||||.|||||||||||||||||||||.|||||.||||..|.||..||..|||||||||||||||
QVQLVQSGAEVKKPGSSVKVSCKASGGTFSSYAISWVRQAPGQGLEWMGGIIPIFGTANYAQKFQGRVTITADESTSTAYMELSSLRSEDTAVYYCAREYYDSSG--YYYEGWFDPWGQGTLVTVSS
                  ^^^^^^^^          ^^^^^^^^                   ^^^^^^^^^^^^^^^^^^^
```

Best Alignment of Rafivirumab light chain to a sequence from OAS
QSALTQPRSVSGSPGQSVTISCTGTSSDIGGYNFVSWYQQHPGKAPKLMIYDATKRPSGVPDRFSGSKSGNTASLTISGLQAEDEADYYCCSYAGDYTPGVVFGGGTKLTVL
||||||||||||||||||||||||||||.||||.||||||||||||||||||..|||||||||||||||||||||||||||||||||||||||||.||||||||||||||||
QSALTQPRSVSGSPGQSVTISCTGTSSDVGGYNYVSWYQQHPGKAPKLMIYDVSKRPSGVPDRFSGSKSGNTASLTISGLQAEDEADYYCCSYAGSYTPGVVFGGGTKLTVL
^^^^^^^^^ ^^^ ^^^^^^^^^^^^

Best Alignment of Rafivirumab heavy chain CDRs to a sequence from OAS
QVQLVQSGAEVKKPGSSVKVSCKASGGTFNRYTVNWVRQAPGQGLEWMGGIIPIFGTANYAQRFQGRLTITADESTSTAYMELSSLRSDDTAVYFCARENLDNSGTYYYFSGWFDPWGQGTLVTVSS
................||||||||||||||.|...|||||||||||||||||||||||||||.||||.||||||||||||||||||||.|||||.||||..|.||...|.|||||||||||||||||
----------------SVKVSCKASGGTFNSYAISWVRQAPGQGLEWMGGIIPIFGTANYAQKFQGRVTITADESTSTAYMELSSLRSEDTAVYYCARESPDCSGGSCYSSGWFDPWGQGTLVTVSS
^^^^^^^^ ^^^^^^^^ ^^^^^^^^^^^^^^^^^^^^

Best Alignment of Rafivirumab light CDRs chain to a sequence from OAS
QSALTQPRSVSGSPGQSVTISCTGTSSDIGGYNFVSWYQQHPGKAPKLMIYDATKRPSGVPDRFSGSKSGNTASLTISGLQAEDEADYYCCSYAGDYTPGVVFGGGTKLTVL
|||||||||||||||||||.||||||||.|||||||||||..||||||.|||.||||||||||||||||.||||||||||||.|||.||||||||.||.|||||||||||||
QSALTQPRSVSGSPGQSVTFSCTGTSSDVGGYNFVSWYQQKSGKAPKLIIYDVTKRPSGVPDRFSGSKSANTASLTISGLQADDEAHYYCCSYAGSYTVGVVFGGGTKLTVL
^^^^^^^^^ ^^^ ^^^^^^^^^^^^

Best Alignment of Rafivirumab CDR-H3 chain to a sequence from OAS
QVQLVQSGAEVKKPGSSVKVSCKASGGTFNRYTVNWVRQAPGQGLEWMGGIIPIFGTANYAQRFQGRLTITADESTSTAYMELSSLRSDDTAVYFCARENLDNSGTYYYFSGWFDPWGQGTLVTVSS
.....||||||||||.|...|||.||..|..|...||||.||.||||||.|.|.....||...|||..||.||.|.||||...|.|...|||.|.|||.||..||.|.|.||||||||||.||||||
-----QSGAEVKKPGESLRISCKGSGYNFTNYWISWVRQMPGKGLEWMGRIDPSDSYTNYSPSFQGHVTISADKSISTAYLQWSTLKASDTAMYYCARPNLYCSGGYCYVSGWFDPWGQGALVTVSS
^^^^^^^^ ^^^^^^^^ ^^^^^^^^^^^^^^^^^^^^

Therapeutic : Ramucirumab

Best Alignment of Ramucirumab heavy chain to a sequence from OAS
EVQLVQSGGGLVKPGGSLRLSCAASGFTFSSYSMNWVRQAPGKGLEWVSSISSSSSYIYYADSVKGRFTISRDNAKNSLYLQMNSLRAEDTAVYYCARVTD--------AFDIWGQGTMVTVSS
|||||.|||||||||||||||||||||||||||||||||||||||||||||||||||||||||||||||||||||||||||||||||||||||||||||........|||||||||||||||
EVQLVESGGGLVKPGGSLRLSCAASGFTFSSYSMNWVRQAPGKGLEWVSSISSSSSYIYYADSVKGRFTISRDNAKNSLYLQMNSLRAEDTAVYYCARVTDIVVVPAAHAFDIWGQGTMVTVSS
^^^^^^^^ ^^^^^^^^ ^^^^^^^^^^^^^^^^^

Best Alignment of Ramucirumab light chain to a sequence from OAS
DIQMTQSPSSVSASIGDRVTITCRASQGIDNWLGWYQQKPGKAPKLLIYDASNLDTGVPSRFSGSGSGTYFTLTISSLQAEDFAVYFCQQAKAFPPTFGGGTKVDIK
||||||||||||||.||||||||||||||.|||.|||||||||||||||.||.|..|||||||||||||.|||||||||.||||.|||||||.||.||||||||.||
DIQMTQSPSSVSASVGDRVTITCRASQGISNWLAWYQQKPGKAPKLLIYAASSLQSGVPSRFSGSGSGTDFTLTISSLQPEDFATYFCQQAKSFPLTFGGGTKVEIK
^^^^^^ ^^^ ^^^^^^^^^

Best Alignment of Ramucirumab heavy chain CDRs to a sequence from OAS
EVQLVQSGGGLVKPGGSLRLSCAASGFTFSSYSMNWVRQAPGKGLEWVSSISSSSSYIYYADSVKGRFTISRDNAKNSLYLQMNSLRAEDTAVYYCARVTDAFDIWGQGTMVTVSS
..............||||||||||||||||||||||||||||||||||||||||||||||||||||||||||||||||||||||||||||||||||||||||||
--------------GGSLRLSCAASGFTFSSYSMNWVRQAPGKGLEWVSSISSSSSYIYYADSVKGRFTISRDNAKNSLYLQMNSLRAEDTAVYYCARVTDAFDIWGQGTMVTVSS
^^^^^^^^ ^^^^^^^^ ^^^^^^^^^

Best Alignment of Ramucirumab light CDRs chain to a sequence from OAS
DIQMTQSPSSVSASIGDRVTITCRASQGIDNWLGWYQQKPGKAPKLLIYDASNLDTGVPSRFSGSGSGTYFTLTISSLQAEDFAVYFCQQAKAFPPTFGGGTKVDIK
||||||||||||||.||||||||||||||.|||.||||||||||.|||..||||..|||||||||.|||.|||||||||.||||.|.|||||||||||||||||.||
DIQMTQSPSSVSASVGDRVTITCRASQGISNWLAWYQQKPGKAPSLLIDTASNLQSGVPSRFSGSASGTDFTLTISSLQPEDFATYYCQQAKAFPPTFGGGTKVEIK
^^^^^^ ^^^ ^^^^^^^^^

Best Alignment of Ramucirumab CDR-H3 chain to a sequence from OAS
EVQLVQSGGGLVKPGGSLRLSCAASGFTFSSYSMNWVRQAPGKGLEWVSSISSSSSYIYYADSVKGRFTISRDNAKNSLYLQMNSLRAEDTAVYYCARVTDAFDIWGQGTMVTVSS
.................|.|.|..||...|||...|.||.|||||||...|..|.....|..|.|.|.|||.|..||...|...|..|.||||||||||||||||||||||||||
--------------SQILSLTCTVSGGSISSYYWSWIRQPPGKGLEWIGYIYYS-GSTNYNPSLKSRVTISVDTSKNQFSLKLSSVTAADTAVYYCARVTDAFDIWGQGTMVTVSS
^^^^^^^^ ^^^^^^^^ ^^^^^^^^^

Therapeutic : Ranibizumab

Best Alignment of Ranibizumab heavy chain to a sequence from OAS
EVQLVESGGGLVQPGGSLRLSCAASGYDFTHYGMNWVRQAPGKGLEWVGWINTYTGEPTYAADFKRRFTFSLDTSKSTAYLQMNSLRAEDTAVYYCAKYPYYYG---------TSHWYFDVWGQGTLVTVSS
.|||||||||.||||.|||||||||||.||.|.|||||||||.||||.|||||.||.||||..|..||.||||||.||||||..||.||||||||||...|.|.............|.||.|||||||||||
QVQLVESGGGVVQPGRSLRLSCAASGYTFTSYAMNWVRQAPGQGLEWMGWINTNTGNPTYAQGFTGRFVFSLDTSVSTAYLQISSLKAEDTAVYYCARDAYGYRVLRFLEWSQGGSWAFDYWGQGTLVTVSS
^^^^^^^^ ^^^^^^^^ ^^^^^^^^^^^^^^^^^^^^^^^^^

Best Alignment of Ranibizumab light chain to a sequence from OAS
DIQLTQSPSSLSASVGDRVTITCSASQDISNYLNWYQQKPGKAPKVLIYFTSSLHSGVPSRFSGSGSGTDFTLTISSLQPEDFATYYCQQYSTVPWTFGQGTKVEIK
|||.|||||||||||||||||||.|||||||||||||||||||||||||.||.|.|||||||||||||||||||||||||||||||||||.||.|.|||||||||||
DIQMTQSPSSLSASVGDRVTITCQASQDISNYLNWYQQKPGKAPKVLIYGTSILQSGVPSRFSGSGSGTDFTLTISSLQPEDFATYYCQQSSTTPRTFGQGTKVEIK
^^^^^^ ^^^ ^^^^^^^^^

Best Alignment of Ranibizumab heavy chain CDRs to a sequence from OAS
EVQLVESGGGLVQPGGSLRLSCAASGYDFTHYGMNWVRQAPGKGLEWVGWINTYTGEPTYAADFKRRFTFSLDTSKSTAYLQMNSLRAEDTAVYYCAKYPYYYGTSHWYFDVWGQGTLVTVSS
..|||.||..|..||.....||.||||.||.||||||.|||||||.|.|||||||||||||.|||.||.|||.||.||||||||.|.|..||.|.|.||...||||.|||||||||.||.|||||
QIQLVQSGPELKKPGETVKISCKASGYTFTNYGMNWVKQAPGKGLKWMGWINTYTGEPTYADDFKGRFAFSLETSASTAYLQINNLKNEDMATYFCAREAYYYGSSHWYFDVWGAGTTVTVSS
^^^^^^^^ ^^^^^^^^ ^^^^^^^^^^^^^^^^

Best Alignment of Ranibizumab light CDRs chain to a sequence from OAS
DIQLTQSPSSLSASVGDRVTITCSASQDISNYLNWYQQKPGKAPKVLIYFTSSLHSGVPSRFSGSGSGTDFTLTISSLQPEDFATYYCQQYSTVPWTFGQGTKVEIK
|||.||..||||||.||||||.||||||||||||||||||....|.||||||||||||||||||||||||..||||.|.|||.|||||||||..|||||.|||.|||
DIQMTQTTSSLSASLGDRVTISCSASQDISNYLNWYQQKPDGTVKLLIYFTSSLHSGVPSRFSGSGSGTDYFLTISNLEPEDIATYYCQQYSKLPWTFGGGTKLEIK
^^^^^^ ^^^ ^^^^^^^^^

Best Alignment of Ranibizumab CDR-H3 chain to a sequence from OAS
EVQLVESGGGLVQPGGSLRLSCAASGYDFTHYGMNWVRQAPGKGLEWVGWINTYTGEPTYAADFKRRFTFSLDTSKSTAYLQMNSLRAEDTAVYYCAKYPYYYGTSHWYFDVWGQGTLVTVSS
.|||..||..|..||.|..|||.|.||.||.|...||.|.||.||||.|.|....|...|...||...||..|||..|||.|..||..||.|.|||||||||||.||||||||.||.|||||
QVQLQQSGAELMKPGASVKLSCKATGYTFTGYWIEWVKQRPGHGLEWIGEILPGSGSTNYNEKFKGKATFTADTSSNTAYMQLSSLTTEDSAIYYCAKYPYYYGSSHWYFDVWGTGTTVTVSS
^^^^^^^^ ^^^^^^^^ ^^^^^^^^^^^^^^^^

Therapeutic : Ravulizumab

Best Alignment of Ravulizumab heavy chain to a sequence from OAS
QVQLVQSGAEVKKPGASVKVSCKASGHIFSNYWIQWVRQAPGQGLEWMGEILPGSGHTEYTENFKDRVTMTRDTSTSTVYMELSSLRSEDTAVYYCARYFFGSS---PNWYFDVWGQGTLVTVSS
||||||||||||||||||||||||||..|..|...||||||||||||||.|.|..|.|.|...|..||||||||||||||||||||||||||||||||.|.||....||.|||.|||||||||||
QVQLVQSGAEVKKPGASVKVSCKASGYTFTSYYMHWVRQAPGQGLEWMGIINPSGGSTSYAQKFQGRVTMTRDTSTSTVYMELSSLRSEDTAVYYCARGFGGSYQINPNYYFDYWGQGTLVTVSS
^^^^^^^^ ^^^^^^^^ ^^^^^^^^^^^^^^^^^^

Best Alignment of Ravulizumab light chain to a sequence from OAS
DIQMTQSPSSLSASVGDRVTITCGASENIYGALNWYQQKPGKAPKLLIYGATNLADGVPSRFSGSGSGTDFTLTISSLQPEDFATYYCQNVLNTPLTFGQGTKVEIK
|||||||||||||||||||||||.||.||...|||||||||||||||||.|.||..|||||||||||||||||||||||||||||||||...|||||||.|||||||
DIQMTQSPSSLSASVGDRVTITCRASQNIRNYLNWYQQKPGKAPKLLIYTASNLQSGVPSRFSGSGSGTDFTLTISSLQPEDFATYYCQQSDNTPLTFGGGTKVEIK
^^^^^^ ^^^ ^^^^^^^^^

Best Alignment of Ravulizumab heavy chain CDRs to a sequence from OAS
QVQLVQSGAEVKKPGASVKVSCKASGHIFSNYWIQWVRQAPGQGLEWMGEILPGSGHTEYTENFKDRVTMTRDTSTSTVYMELSSLRSEDTAVYYCARYFFGSSPNWYFDVWGQGTLVTVSS
.|||.|||||..|||||||.||||.|..||.|||.||.|.||.||||.||||||||.|.|.|.||...|.|.|||..|.||.||||.|||.|||||||...||||||||||||.||.|||||

```
-VQLQQSGAELMKPGASVKMSCKATGYTFSSYWIEWVKQRPGHGLEWIGEILPGSGSTNYNEKFKGKATFTADTSSNTAYMQLSSLTSEDSAVYYCARDYYGSSPNWYFDVWGAGTTVTVSS
                         ^^^^^^^^          ^^^^^^^^                         ^^^^^^^^^^^^^^^
```

Best Alignment of Ravulizumab light CDRs chain to a sequence from OAS

```
DIQMTQSPSSLSASVGDRVTITCGASENIYGALNWYQQKPGKAPKLLIYGATNLADGVPSRFSGSGSGTDFTLTISSLQPEDFATYYCQNVLNTPLTFGQGTKVEIK
||||||||.|||||||..|||||||||||||||||||.|.|..|.||||||||||||..||||||||||....|..|||.|.|.||||||||||||||||||.|||.|.|
DIQMTQSPASLSASVGETVTITCGASENIYGALNWYQRKQGISPQLLIYGATNLADGMSSRFSGSGSGRQYSLKVSSLHPDDVATYYCQNVLNTPLTFGAGTKLELK
                          ^^^^^^          ^^^                    ^^^^^^^^^
```

Best Alignment of Ravulizumab CDR-H3 chain to a sequence from OAS

```
QVQLVQSGAEVKKPGASVKVSCKASGHIFSNYWIQWVRQAPGQGLEWMGEILPGSGHTEYTENFKDRVTMTRDTSTSTVYMELSSLRSEDTAVYYCARYFFGSSPNWYFDVWGQGTLVTVSS
.|.||.||||..|||||||.||||.|..|..|||.||.|.||.||||.||||||||.|.|.|.||...|.|.|||..|.||.||||..||.|.||||||..|||||||||||||.||.|||||
-VKLVESGAELMKPGASVKLSCKATGYTFTGYWIEWVKQRPGHGLEWIGEILPGSGSTNYNEKFKGKATFTADTSSNTAYMQLSSLTTEDSAIYYCARYYYGSSPNWYFDVWGTGTTVTVSS
                         ^^^^^^^^          ^^^^^^^^                         ^^^^^^^^^^^^^^^
```

Therapeutic : Refanezumab

Best Alignment of Refanezumab heavy chain to a sequence from OAS

```
QVQLVQSGSELKKPGASVKVSCKASGYTFTNYGMNWVRQAPGQGLEWMGWINTYTGEPTYADDFTGRFVFSLDTSVSTAYLQISSLKAEDTAVYYCARNPINYYGI----NYEGYVMDYWGQGTLVTVSS
||||||||||||||||||||||||||||||.|.||||||||||||||||||||.||.||||..|||||||||||||||||||||||||||||||||||||||.|..||.|.......||..|||||||||||||
QVQLVQSGSELKKPGASVKVSCKASGYTFTSYAMNWVRQAPGQGLEWMGWINTNTGNPTYAQGFTGRFVFSLDTSVSTAYLQISSLKAEDTAVYYCARDPPGYYDILTGSVDGGYYFDYWGQGTLVTVSS
                         ^^^^^^^^          ^^^^^^^^                         ^^^^^^^^^^^^^^^^^^^^^^^
```

Best Alignment of Refanezumab light chain to a sequence from OAS

```
DIVMTQSPDSLAVSLGERATINCKSSHSVLYSSNQKNYLAWYQQKPGQPPKLLIYWASTRESGVPDRFSGSGSGTDFTLTISSLQAEDVAVYYCHQYLSSLTFGQGTKLEIK
||||||||||||||||||||||||||.|||||||.||||||||||||||||||||||||||||||||||||||||||||||||||||||||||||||.||.|||||||||||
DIVMTQSPDSLAVSLGERATINCKSSQSVLYSSNNKNYLAWYQQKPGQPPKLLIYWASTRESGVPDRFSGSGSGTDFTLTISSLQAEDVAVYYCHQYYSSYTFGQGTKLEIK
                          ^^^^^^^^^^^^          ^^^                    ^^^^^^^^
```

Best Alignment of Refanezumab heavy chain CDRs to a sequence from OAS

```
QVQLVQSGSELKKPGASVKVSCKASGYTFTNYGMNWVRQAPGQGLEWMGWINTYTGEPTYADDFTGRFVFSLDTSVSTAYLQISSLKAEDTAVYYCARNPINYYGINYEGYVMDYWGQGTLVTVSS
|.||||||.||||||..||.|||||||||||||||||.||||.||.||||||||||||||||||||.|||.|||.||.|||||||..||.||||.|.|||..|.|||..||.|.|||||||||.|||||
QIQLVQSGPELKKPGETVKISCKASGYTFTNYGMNWVKQAPGKGLKWMGWINTYTGEPTYADDFKGRFAFSLETSASTAYLQINNLKNEDTATYFCARHLIYYYGSSYEDYAMDYWGQGTSVTVSS
                         ^^^^^^^^          ^^^^^^^^                         ^^^^^^^^^^^^^^^^^^^
```

Best Alignment of Refanezumab light CDRs chain to a sequence from OAS

```
DIVMTQSPDSLAVSLGERATINCKSSHSVLYSSNQKNYLAWYQQKPGQPPKLLIYWASTRESGVPDRFSGSGSGTDFTLTISSLQAEDVAVYYCHQYLSSLTFGQGTKLEIK
.|.|||||.|||||.||..||.|||||||||||||||||||||||||||||.|||||||||||||||||||||.||||||||||||||.||||.|||||||||||||||||.|||||.|
NIMMTQSPSSLAVSAGEKVTISCKSSHSVLYSSNQKNYLAWYQQKPGQSPKLLIYWASTRESGVPDRFTGSGSGTDFTLTISSVQAEDLAVYYCHQYLSSLTFGAGTKLELK
                          ^^^^^^^^^^^^          ^^^                    ^^^^^^^^
```

Best Alignment of Refanezumab CDR-H3 chain to a sequence from OAS

```
QVQLVQSGSELKKPGASVKVSCKASGYTF-TNYGMNWVRQAPGQGLEWMGWINTYTGEPTYADDFTGRFVFSLDTSVSTAYLQISSLKAEDTAVYYCARNPINYYGINYEGYVMDYWGQGTLVTVSS
.|.|..||..|.||..|....|...||.....|..||.||.||..|||||.|....|...|......|.....|||.....|...|...||||.|||||.|.||||..||||.||||||||.|||||
EVKLMESGPGLVKPSQSLSLTCSVTGYSITSGYYWNWIRQFPGNKLEWMGYISYD-GSNNYNPSLKNRISITRDTSKNQFFLKLNSVTTEDTATYYCARDPSNYYGSSYEGYAMDYWGQGTSVTVSS
                         ^^^^^^^^^          ^^^^^^^^                         ^^^^^^^^^^^^^^^^^^^
```

Therapeutic : Reslizumab

Best Alignment of Reslizumab heavy chain to a sequence from OAS

```
EVQLVESGGGLVQPGGSLRLSCAVSGLSLTSNSVNWIRQAPGKGLEWVGLIWSNGDTDYNSAIKSRFTISRDTSKSTVYLQMNSLRAEDTAVYYCAREYY----GYFDYWGQGTLVTVSS
||||||||.|||||||||||||.||....||...|.|||||||||||..|.|.|.|.|....|.|||||||.||.|||||||||||||||||||||||||||||....||||||||||||||||
EVQLVESGGRLVQPGGSLRLSCAASGFTVSSNYMSWVRQAPGKGLEWVSVIYSGGSTYYADSVKGRFTISRDNSKNTVYLQMNSLRAEDTAVYYCAREYYDILTGYFDYWGQGTLVTVSS
                         ^^^^^^^^          ^^^^^^^                         ^^^^^^^^^^^^^^
```

Best Alignment of Reslizumab light chain to a sequence from OAS

```
DIQMTQSPSSLSASVGDRVTITCLASEGISSYLAWYQQKPGKAPKLLIYGANSLQTGVPSRFSGSGSATDYTLTISSLQPEDFATYYCQQSYKFPNTFGQGTKVEVK
|||||||||||||||||||||||.||.||||||||||||||||||||||||.|.|||.||||||||||||.||.||||||||||||||||||||||||..|.|||||||||.|
DIQMTQSPSSLSASVGDRVTITCRASQGISSYLAWYQQKPGKAPKLLIYAASSLQSGVPSRFSGSGSGTDFTLTISSLQPEDFATYYCQQSYSTPWTFGQGTKVEIK
                          ^^^^^^          ^^^                    ^^^^^^^^^
```

Best Alignment of Reslizumab heavy chain CDRs to a sequence from OAS

```
EVQLVESGGGLVQPGGSLRLSCAVSGLSLTSNSVNWIRQAPGKGLEWVGLIWSNGDTDYNSAIKSRFTISRDTSKSTVYLQMNSLRAEDTAVYYCAREYYGYFDYWGQGTLVTVSS
.|||.|||.|||||...|.|.|.||||||||||||.||||.||||.||.|.|||||.||||||||||..||||||||.|.|.||||..||||.|.|||||.|||||||||..|||||
QVQLKESGPGLVQPSQTLSLTCTVSGLSLTSNSVSWIRQPPGKGMEWLGVIWSNGGTDYNSAIKSRLSISRDTSKSQVFLKMNSLQTEDTAMYFCAREYEGYFDYWGQGVMVTVSS
                         ^^^^^^^^          ^^^^^^^                         ^^^^^^^^^^
```

Best Alignment of Reslizumab light CDRs chain to a sequence from OAS

```
DIQMTQSPSSLSASVGDRVTITCLASEGISSYLAWYQQKPGKAPKLLIYGANSLQTGVPSRFSGSGSATDYTLTISSLQPEDFATYYCQQSYKFPNTFGQGTKVEVK
|||.|||||.||||||||||.||.||.|||||.|||||.|||||.||||||.||..|||||||||.||.|..||||..|||||.|||||||||.||.|||.|||||.|
DIQLTQSPSLLSASVGDRVTLTCRASQGISSYVAWYQQRPGKAPNLLIYGASSLRSGVPSRFSGGGSRTQFTLTINNLQPEDSATYYCQQSYSFPLTFGGGTKVEIK
                          ^^^^^^          ^^^                    ^^^^^^^^^
```

Best Alignment of Reslizumab CDR-H3 chain to a sequence from OAS

```
EVQLVESGGGLVQPGGSLRLSCAVSGLSL-TSNSVNWIRQAPGKGLEWVGLIWSNGDTDYNSAIKSRFTISRDTSKSTVYLQMNSLRAEDTAVYYCAREYYGYFDYWGQGTLVTVSS
.|||.|||..||.|..||.|.|.|.|.|.......|||||.||..|||.|.|...|...||...|.|..|.|||||....|..||...||||.||||||||||||||||||||..||||
DVQLQESGPVLVKPSQSLSLTCSVTGYSITSGYYWNWIRQFPGNKLEWMGYISYDGSNKYNPSLKNRISITRDTSKNQFFLKLNSVTTEDTATYYCAREYYGYFDYWGQGTTLTVSS
                         ^^^^^^^^^          ^^^^^^^                         ^^^^^^^^^^
```

Therapeutic : Rilotumumab

Best Alignment of Rilotumumab heavy chain to a sequence from OAS

```
QVQLQESGPGLVKPSETLSLTCTVSGGSISIYYWSWIRQPPGKGLEWIGYVYYSGSTNYNPSLKSRVTISVDTSKNQFSLKLNSVTAADTAVYYCARGGYDFWSGYFDYWGQGTLVTVSS
||||||||||||||||||||||||||||||.|||||||||||||||||||.|||||||||||||||||||||||||||||||||||.|||||||||||||||.|||||||||||||||||||||
QVQLQESGPGLVKPSETLSLTCTVSGGSISSYYWSWIRQPPGKGLEWIGYIYYSGSTNYNPSLKSRVTISVDTSKNQFSLKLSSVTAADTAVYYCARGDYDFWSGYFDYWGQGTLVTVSS
                         ^^^^^^^^          ^^^^^^^                         ^^^^^^^^^^^^^^
```

Best Alignment of Rilotumumab light chain to a sequence from OAS

```
EIVMTQSPATLSVSPGERATLSCRASQSVDSNLAWYRQKPGQAPRLLIYGASTRATGIPARFSGSGSGTEFTLTISSLQSEDFAVYYCQQYINWPPITFGQGTRLEIK
||||||||||||||||||||||||||||||.|||||||||||||||||||||||||||||||||||||||||||||||||||||||||||||||.||||||||||||||||
EIVMTQSPATLSVSPGERATLSCRASQSVSSNLAWYRQKPGQAPRLLIYGASTRATGIPARFSGSGSGTEFTLTISSLQSEDFAVYYCQQYNNWPPITFGQGTRLEIK
                          ^^^^^^          ^^^                    ^^^^^^^^^^
```

Best Alignment of Rilotumumab heavy chain CDRs to a sequence from OAS

```
QVQLQESGPGLVKPSETLSLTCTVSGGSISIYYWSWIRQPPGKGLEWIGYVYYSGSTNYNPSLKSRVTISVDTSKNQFSLKLNSVTAADTAVYYCARGGYDFWSGYFDYWGQGTLVTVSS
..............||||||||||||||||.|||||||||||||||||||.|||||||||||||||||||||||||||||||||||.|||||||||||||||||||||||||||||||||
--------------SETLSLTCTVSGGSISSYYWSWIRQPPGKGLEWIGYIYYSGSTNYNPSLKSRVTISVDTSKNQFSLKLSSVTAADTAVYYCARGGYDFWSGYFDYWGQGTLVTVSS
                         ^^^^^^^^          ^^^^^^^                         ^^^^^^^^^^^^^^
```

Best Alignment of Rilotumumab light CDRs chain to a sequence from OAS

```
EIVMTQSPATLSVSPGERATLSCRASQSVDSNLAWYRQKPGQAPRLLIYGASTRATGIPARFSGSGSGTEFTLTISSLQSEDFAVYYCQQYINWPPITFGQGTRLEIK
||||||.|||||||||||||||||||||||||||||.||.|||||||||||||||||||.||||||||||||||||||||||||||||||||.||||||||||||||||
EIVMTQPPATLSVSPGERATLSCRASQSVDSNLAWYQQKAGQAPRLLIYGASTRATGIPGRFSGSGSGTEFTLTISSLQSEDFAVYYCQQYNNWPPITFGQGTRLEIK
                          ^^^^^^          ^^^                    ^^^^^^^^^^
```

Best Alignment of Rilotumumab CDR-H3 chain to a sequence from OAS

```
QVQLQESGPGLVKPSETLSLTCTVSGGSISIYYWSWIRQPPGKGLEWIGYVYYS-GSTNYNPSLKSRVTISVDTSKNQFSLKLNSVTAADTAVYYCARGGYDFWSGYFDYWGQGTLVTVSS
.....................|..|||..|.|..||.||.||.||||.|......|..||......||||..|.|.......|.|....|||||||||||||||||||||||||||||||
```

----------------SVKVSCKASGGTFSSYAISWVRQAPGQGLEWMGGIIPIFGTANYAQKFQGRVTITADESTSTAYMELSSLRSEDTAVYYCARGGYDFWSGYFDYWGQGTLVTVSS
                    ^^^^^^^^           ^^^^^^^^                        ^^^^^^^^^^^^^^

Therapeutic : Risankizumab

Best Alignment of Risankizumab heavy chain to a sequence from OAS
QVQLVQSGAEVKKPGSSVKVSCKASGYTFTDQTIHWMRQAPGQGLEWIGYIYPRDDSPKYNENFKGKVTITADKSTSTAYMELSSLRSEDTAVYYCAIPDRSGYAWFIYWGQGTLVTVSS
||||||||||||||||||||||||||.||...||.|.||||||||||.|.|.|......|...|.|.|||||||||||||||||||||||||||||||.|.|||..|.||||||||||||
QVQLVQSGAEVKKPGSSVKVSCKASGGTFSSYTISWVRQAPGQGLEWMGRIIPILGIANYAQKFQGRVTITADKSTSTAYMELSSLRSEDTAVYYCAILDSSGYPPFDYWGQGTLVTVSS
                    ^^^^^^^^           ^^^^^^^^                        ^^^^^^^^^^^^^

Best Alignment of Risankizumab light chain to a sequence from OAS
DIQMTQSPSSLSASVGDRVTITCKASRDVAIAVAWYQQKPGKVPKLLIYWASTRHTGVPSRFSGSGSRTDFTLTISSLQPEDVADYFCHQYSSYPFTFGSGTKLEIK
|||||||||||||||||||||||.||..|....|||||||||||.||||.|||.|.|||||||||||.||||||||||||||||.|.||||..||.|||.|||||||
DIQMTQSPSSLSASVGDRVTITCRASQAVDNHLAWYQQKPGKVPNLLIYAASTLHSGVPSRFSGSGSGTDFTLTISSLQPEDVATYYCHQYNTYPYTFGQGTKLEIK
                     ^^^^^^         ^^^                   ^^^^^^^^^

Best Alignment of Risankizumab heavy chain CDRs to a sequence from OAS
QVQLVQSGAEVKKPGSSVKVSCKASGYTFTDQTIHWMRQAPGQGLEWIGYIYPRDDSPKYNENFKGKVTITADKSTSTAYMELSSLRSEDTAVYYCAIPDRSGYAWFIYWGQGTLVTVSS
.|||..|.||..|||.|||.|||.|||||||.|||||.|.|.|||||||||||||.|.||||.||||.|.|||||.|||||.|.||.|||.|||.|||||.||||||.|||||||||||.
-VQLQESDAELVKPGASVKISCKVSGYTFTDHTIHWMKQRPEQGLEWIGYIYPRDGSTKYNEKFKGKATLTADKSSSTAYMQLNSLTSEDSAVYFCAIPDSSGYAWFAYWGQGTLVTVSA
                    ^^^^^^^^           ^^^^^^^^                        ^^^^^^^^^^^^^

Best Alignment of Risankizumab light CDRs chain to a sequence from OAS
DIQMTQSPSSLSASVGDRVTITCKASRDVAIAVAWYQQKPGKVPKLLIYWASTRHTGVPSRFSGSGSRTDFTLTISSLQPEDVADYFCHQYSSYPFTFGSGTKLEIK
||.||||....|..|||||.||||||.||.|||||||||||..||||||||||||||||.||.||||.||||||||..|.||.|||||.||||||||||||||||||
DIVMTQSHKFMSTLVGDRVNITCKASQDVNIAVAWYQQKPGQSPKLLIYWASTRHTGVPDRFTGSGSGTDFTLTISNVQSEDLADYFCQQYSSYPFTFGSGTKLEIK
                     ^^^^^^         ^^^                   ^^^^^^^^^

Best Alignment of Risankizumab CDR-H3 chain to a sequence from OAS
QVQLVQSGAEVKKPGSSVKVSCKASGYTFTDQTIHWMRQAPGQGLEWIGYIYPRDDSPKYNENFKGKVTITADKSTSTAYMELSSLRSEDTAVYYCAIPDRSGYAWFIYWGQGTLVTVSS
.|||..|.||..|||.|||.|||.|||||||.|||||.|.|.|||||||||||||.|.||||.||||.|.|||||.|||||.|.||.|||.|||.|||||.||||||.|||||||||||.
-VQLQESDAELVKPGASVKISCKVSGYTFTDHTIHWMKQRPEQGLEWIGYIYPRDGSTKYNEKFKGKATLTADKSSSTAYMQLNSLTSEDSAVYFCAIPDSSGYAWFAYWGQGTLVTVSA
                    ^^^^^^^^           ^^^^^^^^                        ^^^^^^^^^^^^^

Therapeutic : Rituximab

Best Alignment of Rituximab heavy chain to a sequence from OAS
QVQLQQPGAELVKPGASVKMSCKASGYTFTSYNMHWVKQTPGRGLEWIGAIYPGNGDTSYNQKFKGKATLTADKSSSTAYMQLSSLTSEDSAVYYCARSTYYG-GDWYFNVWGAGTTVTVSA
||||||||||||||||||||||||||||||||||||||||||.||||||||||||||||||||||||||||||||||||||||||||||||||||||||.|||...|||.|||||||||||.
QVQLQQPGAELVKPGASVKMSCKASGYTFTSYNMHWVKQTPGQGLEWIGAIYPGNGDTSYNQKFKGKATLTADKSSSTAYMQLSSLTSEDSAVYYCARSHYYGSSGWYFDVWGAGTTVTVSS
                    ^^^^^^^^           ^^^^^^^^                        ^^^^^^^^^^^^^^^

Best Alignment of Rituximab light chain to a sequence from OAS
QIVLSQSPAILSASPGEKVTMTCRASSSVSYIHWFQQKPGSSPKPWIYATSNLASGVPVRFSGSGSGTSYSLTISRVEAEDAATYYCQQWTSNPPTFGGGTKLEIK
||||||||||||||||||||||||||||||||||.|||||||||||||||||||||||.|||||||||||||||||||||||||||||||.|||||||||||||||
QIVLSQSPAILSASPGEKVTMTCRASSSVSYIHWYQQKPGSSPKPWIYATSNLASGVPARFSGSGSGTSYSLTISRVEAEDAATYYCQQWSSNPPTFGGGTKLEIK
                     ^^^^^          ^^^                   ^^^^^^^^^

Best Alignment of Rituximab heavy chain CDRs to a sequence from OAS
QVQLQQPGAELVKPGASVKMSCKASGYTFTSYNMHWVKQTPGRGLEWIGAIYPGNGDTSYNQKFKGKATLTADKSSSTAYMQLSSLTSEDSAVYYCARSTYYGGDWYFNVWGAGTTVTVSA
.|||||.|||||.||||||||||||||||||||||||||||..||||||||||||||||||||||||||||.||||||||||||||||||||||||||..||||||||.|||.|||||||.
-VQLQQSGAELVRPGASVKMSCKASGYTFTSYNMHWVKQTPRQGLEWIGAIYPGNGDTSYNQKFKGKATLTVDKSSSTAYMQLSSLTSEDSAVYYCARDGYYGGDWYFDVWGTGTTVTVSS
                    ^^^^^^^^           ^^^^^^^^                        ^^^^^^^^^^^^^^

Best Alignment of Rituximab light CDRs chain to a sequence from OAS
QIVLSQSPAILSASPGEKVTMTCRASSSVSYIHWFQQKPGSSPKPWIYATSNLASGVPVRFSGSGSGTSYSLTISRVEAEDAATYYCQQWTSNPPTFGGGTKLEIK
||||.|||||.||||||||||||.|||||||.||.|||.|.|.|||.|||||||||||||.|||||||||||||||||.|||||||||||||.|||||||.|||||.|
QIVLTQSPAIMSASPGEKVTMTCSASSSVSYMHWYQQKSGTSPKRWIYATSNLASGVPARFSGSGSGTSYSLTISRMEAEDAATYYCQQWSSNPPTFGAGTKLELK
                     ^^^^^          ^^^                   ^^^^^^^^^

Best Alignment of Rituximab CDR-H3 chain to a sequence from OAS
QVQLQQPGAELVKPGASVKMSCKASGYTFTSYNMHWVKQTPGRGLEWIGAIYPGNGDTSYNQKFKGKATLTADKSSSTAYMQLSSLTSEDSAVYYCARSTYYGGDWYFNVWGAGTTVTVSA
.|||||.||||.|||||||.||||.|||||.|...||||.||.||||||.|.||.|.||.|||||||.|||.||.||||||||||.||||.|||||.||||.||||||||.|||||||.
-VQLQQSGAELMKPGASVKLSCKATGYTFTGYWIEWVKQRPGHGLEWIGEILPGSGSTNYNEKFKGKATFTADTSSNTAYMQLSSLTTEDSAIYYCARGTYYGYDWYFNVWGTGTTVTVSS
                    ^^^^^^^^           ^^^^^^^^                        ^^^^^^^^^^^^^^

Therapeutic : Robatumumab

Best Alignment of Robatumumab heavy chain to a sequence from OAS
EVQLVQSGGGLVKPGGSLRLSCAASGFTFSSFAMHWVRQAPGKGLEWISVIDTR-GATYYADSVKGRFTISRDNAKNSLYLQMNSLRAEDTAVYYCARLGNF------YYGMDVWGQGTTVTVSS
|||||.|||||||||||||||||||||||||.||.||||||||||||.|.|.|....|.|||||||||||||||||||||||||||||||||||||||||.|........|||||||||||||||||
EVQLVESGGGLVKPGGSLRLSCAASGFTFSSYAMSWVRQAPGKGLEWVSAISGSGGSTYYADSVKGRFTISRDNAKNSLYLQMNSLRAEDTAVYYCAREGYYGSGVYYYYGMDVWGQGTTVTVSS
                    ^^^^^^^^           ^^^^^^^^                        ^^^^^^^^^^^^^^^^^^

Best Alignment of Robatumumab light chain to a sequence from OAS
EIVLTQSPGTLSVSPGERATLSCRASQSI-GSSLHWYQQKPGQAPRLLIKYASQSLSGIPDRFSGSGSGTDFTLTISRLEPEDFAVYYCHQSSRLPHTFGQGTKVEIK
||||||||||||.|||||||||||||||..||.|.||||||||||||||..||....||||||||||||||||||||||||||||||||||.||.|.|||||||||||
EIVLTQSPGTLSLSPGERATLSCRASQSVSGSYLAWYQQKPGQAPRLLIYGASSRATGIPDRFSGSGSGTDFTLTISRLEPEDFAVYYCHQYSRSPQTFGQGTKVEIK
                     ^^^^^^^          ^^^                   ^^^^^^^^^

Best Alignment of Robatumumab heavy chain CDRs to a sequence from OAS
EVQLVQSGGGLVKPGGSLRLSCAASGFTFSSFAMHWVRQAPGKGLEWISVIDTRGATYYADSVKGRFTISRDNAKNSLYLQMNSLRAEDTAVYYCARLGNFYYGMDVWGQGTTVTVSS
..............|.||..||||.||||||..|||||||.||||||.|.|.|.|.|||..||||.|||||.||||||||||||||||||||||||||||.||||||||||||||.||
--------------GESLKISCAACGFTFSSYDMHWVRQATGKGLEWVSAIGTAGDTYYPGSVKGQFTISRENAKNSLYLQMNSLRAEDTAVYYCARLGNNYYGMDVWGQGTTVTGSS
                    ^^^^^^^^           ^^^^^^^                        ^^^^^^^^^^^^

Best Alignment of Robatumumab light CDRs chain to a sequence from OAS
EIVLTQSPGTLSVSPGERATLSCRASQSIGSSLHWYQQKPGQAPRLLIKYASQSLSGIPDRFSGSGSGTDFTLTISRLEPEDFAVYYCHQSSRLPHTFGQGTKVEIK
||||||||...||.|.|..|..||||||||||||||||||.|.|.|||||||||.||.|.|.|||||||||||||||..||.||.|.|.||||||||||||||.|||||||
EIVLTQSPDFQSVTPKEKVTITCRASQSIGSSLHWYQQKPDQSPKLLIKYASQSISGVPSRFSGSGSGTDFTLTINSLEAEDAAAYYCHQSSRLPHTFGGGTKVEIK
                     ^^^^^^         ^^^                   ^^^^^^^^^

Best Alignment of Robatumumab CDR-H3 chain to a sequence from OAS
EVQLVQSGGGLVKPGGSLRLSCAASGFTFSSFAMHWVRQAPGKGLEWISVIDTRGATYYADSVKGRFTISRDNAKNSLYLQMNSLRAEDTAVYYCARLGNFYYGMDVWGQGTTVTVSS
.................|||||..||...||....|.||.||||||||..|...|.|.|..|.|.|||.|..||...|...|..|.|||||||||||.||||||||||||||||||
-----------------LRLSCTVSGGSISSYYWSWIRQPPGKGLEWIGYIYYSGSTNYNPSLKSRVTISVDTSKNQFSLKLSSVTAADTAVYYCARLGVFYYGMDVWGQGTTVTVSS
                    ^^^^^^^^           ^^^^^^^                        ^^^^^^^^^^^^

Therapeutic : Roledumab

Best Alignment of Roledumab heavy chain to a sequence from OAS
QVQLVESGGGVVQPGRSLRLSCTASGFTFKNYAMHWVRQAPAKGLEWVATISYDGRNIQYADSVKGRFTFSRDNSQDTLYLQLNSLRPEDTAVYYCARPVRSRWLQ---LGLEDAFHIWGQGTMVTVSS
||||||||||||||||||||||.||||||..||||||||||.|||||||.|||||.|..||||||||||.|||||..|||||.||||.||||||||||...|.||....|...|||.||||||||||||
QVQLVESGGGVVQPGRSLRLSCAASGFTFSSYAMHWVRQAPGKGLEWVAVISYDGSNKYYADSVKGRFTISRDNSKNTLYLQMNSLRAEDTAVYYCARERFSQWLVIPQLPGFDAFDIWGQGTMVTVSS
                    ^^^^^^^^           ^^^^^^^^                        ^^^^^^^^^^^^^^^^^^^^^

Best Alignment of Roledumab light chain to a sequence from OAS
AIRMTQSPSSFSASTGDRVTITCRASQDIRNYVAWYQQKSGKAPKFLIYAASTLQSGVPSRFSGSGSGTDFTLTINSLQSEDFATYYCQQYYNSPPTFGQGTRVEIT
|||||||||||||||||||||||||||.|..|.||||||.|||||||||||||||||||||||||||||||||||..|||||||||||||||..||||||||.|||.
AIRMTQSPSSFSASTGDRVTITCRASQGIGSYLAWYQQKPGKAPKFLIYAASTLQSGVPSRFSGSGSGTDFTLTISCLQSEDFATYYCQQYYSYPPTFGQGTKVEIK
^^^^^^ ^^^ ^^^^^^^^^

Best Alignment of Roledumab heavy chain CDRs to a sequence from OAS
QVQLVESGGGVVQPGRSLRLSCTASGFTFKNYAMHWVRQAPAKGLEWVATISYDGRNIQYADSVKGRFTFSRDNSQDTLYLQLNSLRPEDTAVYYCARPVRSRWLQLGLEDAFHIWGQGTMVTVSS
||||||||||||||||||||||.||||||..|.||||||||.|||||||.|||||.|..||||||.|||.||||....||||.|..|.|||.||.|||.|||.|.|||..|||.|||||.||||||
QVQLVESGGGVVQPGRSLRLSCAASGFTFSSYGMHWVRQAPGKGLEWVAVISYDGSNKYYADSVKDRFTISRDNFKNSLYLQMNNRRAEDTDVYFCARVVRSGWHQLGGYDAFDIWGQGRMVTVSS
^^^^^^^^ ^^^^^^^^ ^^^^^^^^^^^^^^^^^^

Best Alignment of Roledumab light CDRs chain to a sequence from OAS
AIRMTQSPSSFSASTGDRVTITCRASQDIRNYVAWYQQKSGKAPKFLIYAASTLQSGVPSRFSGSGSGTDFTLTINSLQSEDFATYYCQQYYNSPPTFGQGTRVEIT
.|.|||||||.|||.||||||||||||||||...|.|||.|.|||.|.||||.|.|||||||||||||||.||.|.|||..|||||.||||||||||||.||.|.|.
-IQMTQSPSSLSASVGDRVTITCRASQDIRNDLGWFQQKPGRAPKLLVYAASRLESGVPSRFSGSGSGTDYTLAISSLQPDDFATYFCQQYYNSPPTFGPGTKVDIK
^^^^^^ ^^^ ^^^^^^^^^

Best Alignment of Roledumab CDR-H3 chain to a sequence from OAS
QVQLVESGGGVVQPGRSLRLSCTASGFTFKNYAMHWVRQAPAKGLEWVATISYDGRNIQYADSVKGRFTFSRDNSQDTLYLQLNSLRPEDTAVYYCARPVRSRWLQLGLEDAFHIWGQGTMVTVSS
.............|...|.|.|...|..|..|...|.||.|.|||||...|........|..|.|.|.|.|.|.|.....|.|.|....|||||||||.|..||||||.||||.||||||||||||
-------------PSETLSLTCAVYGGSFSGYYWSWIRQPPGKGLEWIGEINHS-GSTNYNPSLKSRVTISVDTSKNQFSLKLSSVTAADTAVYYCARFVTQRWLQLGSEDAFDIWGQGTMVTVSS
^^^^^^^^ ^^^^^^^^ ^^^^^^^^^^^^^^^^^^

Therapeutic : Romosozumab

Best Alignment of Romosozumab heavy chain to a sequence from OAS
EVQLVQSGAEVKKPGASVKVSCKASGYTFTDYNMHWVRQAPGQGLEWMGEINPNSGGAGYNQKFKGRVTMTTDTSTSTAYMELRSLRSDDTAVYYCARLGYDDI---YDDWYFDVWGQGTTVTVSS
.|||||||||||||||||||||||||||||.|.||||||||||||||||.|||||||..|.||..||||||||||||||||||||||||||||||||||.||.....|...|||.|||||.|||||
QVQLVQSGAEVKKPGASVKVSCKASGYTFTGYYMHWVRQAPGQGLEWMGWINPNSGGTNYAQKLQGRVTMTTDTSTSTAYMELRSLRSDDTAVYYCARLDYDYVWGSYRKYYFDYWGQGTLVTVSS
^^^^^^^^ ^^^^^^^^ ^^^^^^^^^^^^^^^^^^

Best Alignment of Romosozumab light chain to a sequence from OAS
DIQMTQSPSSLSASVGDRVTITCRASQDISNYLNWYQQKPGKAPKLLIYYTSRLLSGVPSRFSGSGSGTDFTLTISSLQPEDFATYYCQQGDTLPYTFGGGTKVEIK
|||||||||||||||||||||||||||.|||||||||||||||||||||..|||.|||||||||||||||||||||||||||||||||||.||.|.|||||||||||
DIQMTQSPSSLSASVGDRVTITCRASQSISNYLNWYQQKPGKAPKLLIYAASRLQSGVPSRFSGSGSGTDFTLTISSLQPEDFATYYCQQRDTTPLTFGGGTKVEIK
^^^^^^ ^^^ ^^^^^^^^^

Best Alignment of Romosozumab heavy chain CDRs to a sequence from OAS
EVQLVQSGAEVKKPGASVKVSCKASGYTFTDYNMHWVRQAPGQGLEWMGEINPNSGGAGYNQKFKGRVTMTTDTSTSTAYMELRSLRSDDTAVYYCARLGYDDIYDDWYFDVWGQGTTVTVSS
.|||.|||.|..|||||||..|||||||||||||.||.|..|..|||.|.||||.||..|||||||..|.|.|.|.||||||||||.|.||||||.||||||..|.||||||||.||||||||
-VQLQQSGPELVKPGASVKIPCKASGYTFTDYNMDWVKQSHGKSLEWIGDINPNNGGTIYNQKFKGKATLTVDKSSSTAYMELRSLTSEDTAVYYYARLGYDSSYEDWYFDVWGTGTTVTVSS
^^^^^^^^ ^^^^^^^^ ^^^^^^^^^^^^^^^^

Best Alignment of Romosozumab light CDRs chain to a sequence from OAS
DIQMTQSPSSLSASVGDRVTITCRASQDISNYLNWYQQKPGKAPKLLIYYTSRLLSGVPSRFSGSGSGTDFTLTISSLQPEDFATYYCQQGDTLPYTFGGGTKVEIK
|||.||..||||||.||||||.||||||||||||||||||....||||||||||.|||||||||||||||..||||.|..||.|||.||||||||||||||||.|||
DIQLTQTTSSLSASLGDRVTISCRASQDISNYLNWYQQKPDGTVKLLIYYTSRLHSGVPSRFSGSGSGTDYSLTISNLEQEDIATYFCQQGDTLPYTFGGGTKLEIK
^^^^^^ ^^^ ^^^^^^^^^

Best Alignment of Romosozumab CDR-H3 chain to a sequence from OAS
EVQLVQSGAEVKKPGASVKVSCKASGYTFTDYNMHWVRQAPGQGLEWMGEINPNSGGAGYNQKFKGRVTMTTDTSTSTAYMELRSLRSDDTAVYYCARLGYDDIYDDWYFDVWGQGTTVTVSS
||.||.||....|||.|.|.||.|||.||.||.||||||||..||||...|...|....|....|||.|...|....|......||||.||||||.||.|.||.||||||||||.||||||||
EVHLVESGGGLVKPGGSLKLSCAASGFTFSDYGMHWVRQAPEKGLEWVAYISSGSSTIYYADTVKGRFTISRDNAKNTLFLQMTSLRSEDTAVYYWARKGEDDGYDDWYFDVWGTGTTVTVSS
^^^^^^^^ ^^^^^^^^ ^^^^^^^^^^^^^^^^

Therapeutic : Rontalizumab

Best Alignment of Rontalizumab heavy chain to a sequence from OAS
EVQLVESGGGLVQPGGSLRLSCATSGYTFTEYIIHWVRQAPGKGLEWVASINPDYDITNYNQRFKGRFTISLDKSKRTAYLQMNSLRAEDTAVYYCASWIS----DFFDYWGQGTLVTVSS
|||||||||||||||||||||||.||.||..|..|||||||||||.|||.||.|..||.|....|||||||.|..|.|.||||||||||||||||||.|.......|||||||||||||||
EVQLVESGGGLVQPGGSLRLSCAASGFTFSSYWMHWVRQAPGKGLVWVARINSDGSITTYADSVKGRFTISRDNAKNTLYLQMNSLRAEDTAVYYCARWDYSNFDGFFDYWGQGTLVTVSS
^^^^^^^^ ^^^^^^^^ ^^^^^^^^^^^^^^

Best Alignment of Rontalizumab light chain to a sequence from OAS
DIQMTQSPSSLSASVGDRVTITCRASQSVSTSSYSYMHWYQQKPGKAPKVLISYASNLESGVPSRFSGSGSGTDFTLTISSLQPEDFATYYCQHSWGIPRTFGQGTKVEIK
||||||||||||||.||||||||||||||.....||..|||||||||||||||.||.||||||||||||||||||||||||||||||||||||||.|.|...|||||||||||||
DIQMTQSPSSLSASIGDRVTITCRASQSV----SSYLNWYQQKPGKAPKVLISDASSLESGVPSRFSGSGSGTDFTLTISSLQPEDFATYYCQQSYSSPRTFGQGTKVEIK
^^^^^^^^^^ ^^^ ^^^^^^^^^

Best Alignment of Rontalizumab heavy chain CDRs to a sequence from OAS
EVQLVESGGGLVQPGGSLRLSCATSGYTFTEYIIHWVRQAPGKGLEWVASINPDYDITNYNQRFKGRFTISLDKSKRTAYLQMNSLRAEDTAVYYCASWISDFFDYWGQGTLVTVSS
................|...||.|||||||.|.|||||||||.||||...|||..|||||.|||.||.|...|.|..|.|.|....||..|||||||||||.|.|||||||||||||||
---------------ASVKVSCKTSGYTFTKYLIHWVRQAPGQGLEWMGTINPQGDITNYAQRFQGRITLTEDTSTSTVYMELSSLTSEDTAVYYCASWDSSFFDYWGQGTLVTVSS
^^^^^^^^ ^^^^^^^^ ^^^^^^^^^^

Best Alignment of Rontalizumab light CDRs chain to a sequence from OAS
DIQMTQSPSSLSASVGDRVTITCRASQSVSTSSYSYMHWYQQKPGKAPKVLISYASNLESGVPSRFSGSGSGTDFTLTISSLQPEDFATYYCQHSWGIPRTFGQGTKVEIK
||..||||.||..|.|.|.||.|||||||||||||||||||||||..||.||.||||||||||.|||||||||||||.|.....||.|||||||||.||||||.|||.|||
DIVLTQSPASLAVSLGQRATISCRASQSVSTSSYSYMHWYQQKPGQPPKLLIKYASNLESGVPARFSGSGSGTDFTLNIQPVEEEDTATYYCQHSWEIPRTFGGGTKLEIK
^^^^^^^^^^ ^^^ ^^^^^^^^^

Best Alignment of Rontalizumab CDR-H3 chain to a sequence from OAS
EVQLVESGGGLVQPGGSLRLSCATSGYTFTEYIIHWVRQAPGKGLEWVASINPDYDITNYNQRFKGRFTISLDKSKRTAYLQMNSLRAEDTAVYYCASWISDFFDYWGQGTLVTVSS
||||.||||.|||||||||||||.||..|..|...||||.|||||||...|......|.|.....||||||.|.||.|.|||||||||.|||.|||||||||.||||||||||||||
EVQLLESGGDLVQPGGSLRLSCAASGFIFRNYAMMWVRQSPGKGLEWISTIDGPTTNTHYADSVEGRFTISRDNSKNTVYLQMNSLRAGDTAIYYCASWISDHFDYWGQGTLVTVSS
^^^^^^^^ ^^^^^^^^ ^^^^^^^^^^

Therapeutic : Rovalpituzumab

Best Alignment of Rovalpituzumab heavy chain to a sequence from OAS
QVQLVQSGAEVKKPGASVKVSCKASGYTFTNYGMNWVRQAPGQGLEWMGWINTYTGEPTYADDFKGRVTMTTDTSTSTAYMELRSLRSDDTAVYYCARIGDS--SPSDYWGQGTLVTVSS
|||||||||||||||||||||||||||||||||.|||||||||||||||||.||.|...||....||||.||||||||||||||||||||||||||||.|....||.|||||||||||||
QVQLVQSGAEVKKPGASVKVSCKASGYTFTNYGINWVRQAPGQGLEWMGWISTYNGNTNYAQKLQGRVTVTTDTSTSTAYMELRSLRSDDTAVYYCARDGWDLLSPLDYWGQGTLVTVSS
^^^^^^^^ ^^^^^^^^ ^^^^^^^^^^^^^

Best Alignment of Rovalpituzumab light chain to a sequence from OAS
EIVMTQSPATLSVSPGERATLSCKASQSVSNDVVWYQQKPGQAPRLLIYYASNRYTGIPARFSGSGSGTEFTLTISSLQSEDFAVYYCQQDYTSPWTFGQGTKLEIK
|||||||||||||||||||||||.||||||....|||||||||||||||.||.|.||||||||||||||||||||||||||||||||||||.|||.|||||||||||
EIVMTQSPATLSVSPGERATLSCRASQSVSSNLAWYQQKPGQAPRLLIYGASTRATGIPARFSGSGSGTEFTLTISSLQSEDFAVYYCQQDDTSPYTFGQGTKLEIK
^^^^^^ ^^^ ^^^^^^^^^

Best Alignment of Rovalpituzumab heavy chain CDRs to a sequence from OAS
QVQLVQSGAEVKKPGASVKVSCKASGYTFTNYGMNWVRQAPGQGLEWMGWINTYTGEPTYADDFKGRVTMTTDTSTSTAYMELRSLRSDDTAVYYCARIGDSSPSDYWGQGTLVTVSS
..|.||.|.|.||||..||.|||||||||||||||||.||||.||.||||||||||||||.||||||......||.||||.....|...|||....||.||||..||||.||..|||.

```
-IQVVQAGPELKKPGETVKISCKASGYTFTNYGMNWVKQAPGKGLKWMGWINTYTGEPTYTDDFKGRFAFSLETSASTAYLQIINLKNEDTATDCYARAGDSSDFDYWGRGTTLTVSA
                  ^^^^^^^^            ^^^^^^^^                        ^^^^^^^^^^^
```

Best Alignment of Rovalpituzumab light CDRs chain to a sequence from OAS

```
EIVMTQSPATLSVSPGERATLSCKASQSVSNDVVWYQQKPGQAPRLLIYYASNRYTGIPARFSGSGSGTEFTLTISSLQSEDFAVYYCQQDYTSPWTFGQGTKLEIK
.||.||.|..|.||.|.|.|..|||||||||||.||||||.|.|.|||||||||||||.|.||.|||.||.||.|||..|.||.|||.|||||.||||.|.|||||||
SIVITQTPQFLLVSAGARVTITCKASQSVSNDVAWYQQKPRQSPKLLIYYASNRYTGVPDRFTGSGYGTDFTFTISTGQAEDLAVYFCQQDYSSPWTVGGGTKLEIK
                   ^^^^^^          ^^^                      ^^^^^^^^^
```

Best Alignment of Rovalpituzumab CDR-H3 chain to a sequence from OAS

```
QVQLVQSGAEVKKPGASVKVSCKASGYTFTNYGMNWVRQAPGQGLEWMGWINTYTGEPTYADDFKGRVTMTTDTSTSTAYMELRSLRSDDTAVYYCARIGDSSPSDYWGQGTLVTVSS
.............|.|.|.||.|||.||..|.|.|||||||.||||...|.....|..|.|..|||.|...|......|....|||..|||||||||.||||||||||||||||||
--------------GESLKISCAASGFTFSSYWMSWVRQAPGKGLEWVANIKQDGSEKYYVDSVKGRFTISRDNAKNSLYLQMNSLRAEDTAVYYCAREGDSSPSDYWGQGTLVTVSS
                  ^^^^^^^^            ^^^^^^^^                        ^^^^^^^^^^^
```

Therapeutic : Rozanolixizumab

Best Alignment of Rozanolixizumab heavy chain to a sequence from OAS

```
EVPLVESGGGLVQPGGSLRLSCAVSGFTFSNYGMVWVRQAPGKGLEWVAYIDSDGDNTYYRDSVKGRFTISRDNAKSSLYLQMNSLRAEDTAVYYCTTGIV---RPFLYWGQGTLVTVSS
||.||||||||||||||||||||.|||||.||.|.||||||.|||||||||...||||||||||||||||||||||||||||||.|||.||||.||||||.....|.|.||||||||||||
EVQLVESGGGLVQPGGSLRLSCAASGFTFTNYYMAWVRQAPTKGLEWVAYISTGGDNTYYRDSVKGRFTISRDNAKSSLYLQMDSLRSEDTATYYCTTGDYGYTRLFAYWGQGTLVTVSS
                  ^^^^^^^^            ^^^^^^^^                        ^^^^^^^^^^^^^
```

Best Alignment of Rozanolixizumab light chain to a sequence from OAS

```
DIQMTQSPSSLSASVGDRVTITCKSSQSLVGASGKTYLYWLFQKPGKAPKRLIYLVSTLDSGIPSRFSGSGSGTEFTLTISSLQPEDFATYYCLQGTHFPHTFGQGTKLEIK
|||||||||||||||||||||||..|||........||.|..||||||||||||..|.|.||.|||||||||||||||||||||||||||||||||....|||||||||||||
DIQMTQSPSSLSASVGDRVTITCRASQSI-----SNYLNWYQQKPGKAPKRLIYAASSLQSGVPSRFSGSGSGTEFTLTISSLQPEDFATYYCLQHNNYPHTFGQGTKLEIK
                   ^^^^^^^^^^^          ^^^                     ^^^^^^^^^
```

Best Alignment of Rozanolixizumab heavy chain CDRs to a sequence from OAS

```
EVPLVESGGGLVQPGGSLRLSCAVSGFTFSNYGMVWVRQAPGKGLEWVAYIDSDGDNTYYRDSVKGRFTISRDNAKSSLYLQMNSLRAEDTAVYYCTTGIVRPFLYWGQGTLVTVSS
................|||||||.||||||||.|.||||.|||||.||..|.|||..|.|.||||||||.||||||.||||||||||||||||||.||||.||.||||||||||||
----------------SLRLSCAASGFTFSNYWMHWVRQVPGKGLVWVSRINSDGSSTTYADSVKGRFTVSRDNAKNSLYLQMNSLRAEDTAVYYCATGIVGPFDYWGQGTLVTVSS
                  ^^^^^^^^            ^^^^^^^^                        ^^^^^^^^^^
```

Best Alignment of Rozanolixizumab light CDRs chain to a sequence from OAS

```
DIQMTQSPSSLSASVGDRVTITCKSSQSLVGASGKTYLYWLFQKPGKAPKRLIYLVSTLDSGIPSRFSGSGSGTEFTLTISSLQPEDFATYYCLQGTHFPHTFGQGTKLEIK
|..|||.|..||...|....|.|||||||....|||||.||.|.||..|||||||||.||||.|.||.||||||.|||.||....||...||||||||||||||.|||||||
DVVMTQTPLTLSVTIGQPASISCKSSQSLLDSDGKTYLNWLLQRPGQSPKRLIYLVSKLDSGVPDRFTGSGSGTDFTLKISRVEAEDLGVYYCLQGTHFPHTFGGGTKLEIK
                   ^^^^^^^^^^^          ^^^                     ^^^^^^^^^
```

Best Alignment of Rozanolixizumab CDR-H3 chain to a sequence from OAS

```
EVPLVESGGGLVQPGGSLRLSCAVSGFTFSNYGMVWVRQAPGKGLEWVAYIDSDGDNTYYRDSVKGRFTISRDNAKSSLYLQMNSLRAEDTAVYYCTTGIVRPFLYWGQGTLVTVSS
||.|..||..||.||....|||..|||......|.||.|.|..||||...||.....|.|.....|..||..|......|||..||..||.||.|||||.||||.|||||||||||.
EVQLQQSGAELVRPGALVKLSCTASGFNIKDDYMHWVKQRPEPGLEWIGWIDPENGDTEYASKFQGKATITADTSSNTAYLQLSSLTSEDNAVDYCTTGGVRPFAYWGQGTLVTVSA
                  ^^^^^^^^            ^^^^^^^^                        ^^^^^^^^^^
```

Therapeutic : Sacituzumab

Best Alignment of Sacituzumab heavy chain to a sequence from OAS

```
QVQLQQSGSELKKPGASVKVSCKASGYTFTNYGMNWVKQAPGQGLKWMGWINTYTGEPTYTDDFKGRFAFSLDTSVSTAYLQISSLKADDTAVYFCARGGFGSSYWYFDVWGQGSLVTVSS
|.||.|||.||||||..||.||||||||||||||||||||||||||||||||||||||||||||.|||||||||||.||.|||||||..||..|||.|||||.|.||||||||||||.|..|||||
QIQLVQSGPELKKPGETVKISCKASGYTFTNYGMNWVKQAPGQGLKWMGWINTYTGEPTYADDFKGRFAFSLETSASTAYLQINNLKNEDTATYFCARRGYGSSYWYFDVWGAGTTVTVSS
                  ^^^^^^^^            ^^^^^^^^                        ^^^^^^^^^^^^^^
```

Best Alignment of Sacituzumab light chain to a sequence from OAS

```
DIQLTQSPSSLSASVGDRVSITCKASQDVSIAVAWYQQKPGKAPKLLIYSASYRYTGVPDRFSGSGSGTDFTLTISSLQPEDFAVYYCQQHYITPLTFGAGTKVEIK
||||||||||||||||||||.|||.|||..|..|||||||||||||||||.||....|||.|||||||||||||||||||||||||||||||||||...|||||.|||||||
DIQLTQSPSSLSASVGDRVTITCRASQGISSYVAWYQQKPGKAPKLLIYDASNLQSGVPSRFSGSGSGTDFTLTISSLQPEDFAVYYCQQHNSSPLTFGGGTKVEIK
                   ^^^^^^          ^^^                      ^^^^^^^^^
```

Best Alignment of Sacituzumab heavy chain CDRs to a sequence from OAS

```
QVQLQQSGSELKKPGASVKVSCKASGYTFTNYGMNWVKQAPGQGLKWMGWINTYTGEPTYTDDFKGRFAFSLDTSVSTAYLQISSLKADDTAVYFCARGGFGSSYWYFDVWGQGSLVTVSS
|.||.|||.|.||||..||.||||||||||||||||||||||.||||||||||||||||||.||||||||||||.||.|||||||..||..|.|.||||||||.||||||||||||.|..|||||
QIQLVQSGPERKKPGETVKISCKASGYTFTNYGMNWVKQAPGKGLKWMGWINTYTGEPTYADDFKGRFAFSLETSASTAYLQINNLKNADMATYFCARGGYGSSYWYFDVWGAGTTVTVSS
                  ^^^^^^^^            ^^^^^^^^                        ^^^^^^^^^^^^^^
```

Best Alignment of Sacituzumab light CDRs chain to a sequence from OAS

```
DIQLTQSPSSLSASVGDRVSITCKASQDVSIAVAWYQQKPGKAPKLLIYSASYRYTGVPDRFSGSGSGTDFTLTISSLQPEDFAVYYCQQHYITPLTFGAGTKVEIK
||..|||....|.||||||||||||||||||||||||||||..|||||||||||||||||||||||||||||.||||.|..|.|||||||||||||.|||.|||.|||
DIVMTQSHKFMSTSVGDRVSITCKASQDVSIAVAWYQQKPGQSPKLLIYSASYRYTGVPDRFSGSGSGTDFTFTISSVQADDLAVYYCQQHYITPWTFGGGTKLEIK
                   ^^^^^^          ^^^                      ^^^^^^^^^
```

Best Alignment of Sacituzumab CDR-H3 chain to a sequence from OAS

```
QVQLQQSGSELKKPGASVKVSCKASGYTFTNYGMNWVKQAPGQGLKWMGWINTYTGEPTYTDDFKGRFAFSLDTSVSTAYLQISSLKADDTAVYFCARGGFGSSYWYFDVWGQGSLVTVSS
.||||.||.||.|||||||.|||||||.|..|.||||||.||.||.|.|.|....|...|||......|.|.||||.|.|||...|.||||||||||||||||||||.|.|..|||||
-VQLQESGAELVKPGASVKISCKASGYAFSSYWMNWVKQRPGKGLEWIGQIYPGDGDTNYNGKFKGKAKVTADKSSSTAYMQLSSLTSEDSAVYFCARGGFGSSYWYFDVRGTGTTVTVSS
                  ^^^^^^^^            ^^^^^^^^                        ^^^^^^^^^^^^^^
```

Therapeutic : Sarilumab

Best Alignment of Sarilumab heavy chain to a sequence from OAS

```
EVQLVESGGGLVQPGRSLRLSCAASRFTFDDYAMHWVRQAPGKGLEWVSGISWNSGRIGYADSVKGRFTISRDNAENSLFLQMNGLRAEDTALYYCAKGRD-----------SFDIWGQGTMVTVSS
|||||||||||||||||||||||||.||||||||||||||||||||||||||||||.||||||||||||||||||.|||.|||||||||||||||||||.|............||||||||||||||
EVQLVESGGGLVQPGRSLRLSCAASGFTFDDYAMHWVRQAPGKGLEWVSGISWNSGSIGYADSVKGRFTISRDNAKNSLYLQMNGLRAEDTALYYCAKGIDCSSTSCYSPWNAFDIWGQGTMVTVSS
                  ^^^^^^^^            ^^^^^^^^                        ^^^^^^^^^^^^^^^^^^^^
```

Best Alignment of Sarilumab light chain to a sequence from OAS

```
DIQMTQSPSSVSASVGDRVTITCRASQGISSWLAWYQQKPGKAPKLLIYGASSLESGVPSRFSGSGSGTDFTLTISSLQPEDFASYYCQQANSFPYTFGQGTKLEIK
||||||||||||||||||||||||||||||||||||||||||||||||||||||||||||||||||||||||||||||||||||||.|||||||||||||||||||||
DIQMTQSPSSVSASVGDRVTITCRASQGISSWLAWYQQKPGKAPKLLIYGASSLESGVPSRFSGSGSGTDFTLTISSLQPEDFATYYCQQANSFPYTFGQGTKLEIK
                   ^^^^^^          ^^^                      ^^^^^^^^^
```

Best Alignment of Sarilumab heavy chain CDRs to a sequence from OAS

```
EVQLVESGGGLVQPGRSLRLSCAASRFTFDDYAMHWVRQAPGKGLEWVSGISWNSGRIGYADSVKGRFTISRDNAENSLFLQMNGLRAEDTALYYCAKGRDSFDIWGQGTMVTVSS
................|||||||||.||||||||||||||||||||||||||||||.|||||||||||||||||||.|||.||||.|||||||||||||||||.||||||||.||||||
----------------SLRLSCAASGFTFDDYAMHWVRQAPGKGLEWVSGISWNSGSIGYADSVKGRFTISRDNAKNSLYLQMNSLRAEDTALYYCAKGRDAFDIWGQGAMVTVSS
                  ^^^^^^^^            ^^^^^^^^                        ^^^^^^^^^
```

Best Alignment of Sarilumab light CDRs chain to a sequence from OAS

```
DIQMTQSPSSVSASVGDRVTITCRASQGISSWLAWYQQKPGKAPKLLIYGASSLESGVPSRFSGSGSGTDFTLTISSLQPEDFASYYCQQANSFPYTFGQGTKLEIK
|||||||||||||||||||.||||||||||||||||||||||||||||||||.||||.|||||.||||||||||||||||||||.||||||||||||||||||..||
DIQMTQSPSSVSASVGDRVTFTCRASQGISSWLAWYQQKPGKAPKLLIYGASSLQSGVPLRFSGSRSGTDFTLTISSLQPEDFATYYCQQANSFPYTFGQGTKVDIK
                   ^^^^^^          ^^^                      ^^^^^^^^^
```

Best Alignment of Sarilumab CDR-H3 chain to a sequence from OAS

```
EVQLVESGGGLVQPGRSLRLSCAASRFTFDDYAMHWVRQAPGKGLEWVSGISWNSGRIGYADSVKGRFTISRDNAENSLFLQMNGLRAEDTALYYCAKGRDSFDIWGQGTMVTVSS
.......|.|||.|...|.|||..|......|...|.||.|||||||...|....|...|..|.|.|.|||.|...|...|......|.|||.|||||||||||||||||||||||
```

-------GPGLVKPSETLSLSCIVSGDSISSYYWSWIRQSPGKGLEWIGYIYYS-GSTNYNPSLKSRVTISVDTSNNQFSLKLTSVTAADTAVYYCAKGRDSFDIWGQGTMVTVSS
                    ^^^^^^^^               ^^^^^^^^                                   ^^^^^^^^^

Therapeutic : Satralizumab

Best Alignment of Satralizumab heavy chain to a sequence from OAS
QVQLQESGPGLVKPSETLSLTCAVSGHSISHDHAWSWVRQPPGEGLEWIGFISYSGITNYNPSLQGRVTISRDNSKNTLYLQMNSLRAEDTAVYYCARSLAR----TTAMDYWGEGTLVTVSS
|||||||||||||||.||||||||||.|||....|||||||||.||||||.|..||.|||||||..|||||||||||||||||||||||||||||||.|...........||||.||||||||
QVQLQESGPGLVKPSGTLSLTCAVSGGSISSSNWWSWVRQPPGKGLEWIGEIYHSGSTNYNPSLKSRVTISRDNSKNTLYLQMNSLRAEDTAVYYCAKSKEPGIAAAGEFDYWGQGTLVTVSS
                    ^^^^^^^^^               ^^^^^^^                                   ^^^^^^^^^^^^^^^^

Best Alignment of Satralizumab light chain to a sequence from OAS
DIQMTQSPSSLSASVGDSVTITCQASTDISSHLNWYQQKPGKAPELLIYYGSHLLSGVPSRFSGSGSGTDFTFTISSLEAEDAATYYCGQGNRLPYTFGQGTKVEIE
|||||||||||||||||.||||||||.|||..|||||||||||||||||..|.|..||||||||||||||||||||||..||.|||||.|...|||||||||||||.
DIQMTQSPSSLSASVGDRVTITCQASQDISNYLNWYQQKPGKAPELLIYDASNLQTGVPSRFSGSGSGTDFTFTISSLQPEDIATYYCQQYEDLPYTFGQGTKVEIK
                     ^^^^^^               ^^^                                ^^^^^^^^^

Best Alignment of Satralizumab heavy chain CDRs to a sequence from OAS
QVQLQESGPGLVKPSETLSLTCAVSGHSISHDHAWSWVRQPPGEGLEWIGFISYSGITNYNPSLQGRVTISRDNSKNTLYLQMNSLRAEDTAVYYCARSLARTTAMDYWGEGTLVTVSS
.||||||||||||||..|||||.|.|.||..|.||.|.||.||..|||.|.|||||.|.|||||..|..|.||.|||...||.||...||||.|||||||.|..||||||.||.|||||
DVQLQESGPGLVKPSQSLSLTCTVTGYSITSDYAWNWIRQFPGNKLEWMGYISYSGSTSYNPSLKSRISITRDTSKNQFFLQLNSATTEDTATYYCARSLLRLLAMDYWGQGTSVTVSS
                    ^^^^^^^^^               ^^^^^^^                                   ^^^^^^^^^^^^

Best Alignment of Satralizumab light CDRs chain to a sequence from OAS
DIQMTQSPSSLSASVGDSVTITCQASTDISSHLNWYQQKPGKAPELLIYYGSHLLSGVPSRFSGSGSGTDFTFTISSLEAEDAATYYCGQGNRLPYTFGQGTKVEIE
||||||..||||||.||.|||.|.||.|||.|||||||||.....|||||.|.|.|||||||||||||||...|||.||.||.|||.|.|||.||||||.|||.||.
DIQMTQTTSSLSASLGDRVTISCRASQDISNHLNWYQQKPDGTVKLLIYYTSRLHSGVPSRFSGSGSGTDYSLTISNLEQEDIATYFCQQGNTLPYTFGGGTKLEIK
                     ^^^^^^               ^^^                                ^^^^^^^^^

Best Alignment of Satralizumab CDR-H3 chain to a sequence from OAS
QVQLQESGPGLVKPSETLSLTCAVSGHSISHDHAWSWVRQPPGEGLEWIGFISYS-GITNYNPSLQGRVTISRDNSKNTLYLQMNSLRAEDTAVYYCARSLARTTAMDYWGEGTLVTVSS
||||..||..||.|.....|.|..||.....|....||.|.||.|||||..|......|.||....|..|.....|..|.|.|..||..||.|||.||||||.|.||||.|.||.|||||
QVQLKQSGAELVRPGASVKLSCKASGYTF-TDYYINWVKQRPGQGLEWIARIYPGRCNTYYNEKVKGKATLTAEKSSITAYMQLRSLTSEDSAVYCCARSLAGTRAMDYGGQGTSVTVSS
                    ^^^^^^^^^               ^^^^^^^^                                   ^^^^^^^^^^^^

Therapeutic : Secukinumab

Best Alignment of Secukinumab heavy chain to a sequence from OAS
EVQLVESGGGLVQPGGSLRLSCAASGFTFSNYWMNWVRQAPGKGLEWVAAINQDGSEKYYVGSVKGRFTISRDNAKNSLYLQMNSLRVEDTAVYYCVRDYYDILTDYYIHYWYFDLWGRGTLVTVSS
||||||||||||||||||||||||||||||.|||.||||||||||||||.|.|||||||||.|||||||||||||||||||||||||.||||||||.|.|||....|...|||||||||||||||||
EVQLVESGGGLVQPGGSLRLSCAASGFTFSSYWMSWVRQAPGKGLEWVANIKQDGSEKYYVDSVKGRFTISRDNAKNSLYLQMNSLRAEDTAVYYCARSYYDFWSGYRTSYWYFDLWGRGTLVTVSS
                    ^^^^^^^^               ^^^^^^^^                                   ^^^^^^^^^^^^^^^^^^^

Best Alignment of Secukinumab light chain to a sequence from OAS
EIVLTQSPGTLSLSPGERATLSCRASQSVSSSYLAWYQQKPGQAPRLLIYGASSRATGIPDRFSGSGSGTDFTLTISRLEPEDFAVYYCQQYGSSPCTFGQGTRLEIK
||||||||||||||||||||||||||||||||||||||||||||||||||||||||||||||||||||||||||||||||||||||||||||||||||||||||||||
EIVLTQSPGTLSLSPGERATLSCRASQSVSSSYLAWYQQKPGQAPRLLIYGASSRATGIPDRFSGSGSGTDFTLTISRLEPEDFAVYYCQQYGSSPCTFGQGTRLEIK
                     ^^^^^^^               ^^^                                ^^^^^^^^^

Best Alignment of Secukinumab heavy chain CDRs to a sequence from OAS
EVQLVESGGGLVQPGGSLRLSCAASGFTFSNYWMNWVRQAPGKGLEWVAAINQDGSEKYYVGSVKGRFTISRDNAKNSLYLQMNSLRVEDTAVYYCVRDYYDILTDYYIHYWYFDLWGRGTLVTVSS
.............|||||||||||||||||.|||.||||||||||||||.|||||||||||.|||||||||||||||||.||||||||||||||||.|..|||||.||..|||||||||||||||||
-------------PGGSLRLSCAASGFTFSSYWMSWVRQAPGKGLEWVANINQDGSEKYYVDSVKGRFTISRDNAKNSLCLQMNSLRVEDTAVYYCARANYDILTGYYTGYWYFDLWGRGTLVTVSS
                    ^^^^^^^^               ^^^^^^^^                                   ^^^^^^^^^^^^^^^^^^^

Best Alignment of Secukinumab light CDRs chain to a sequence from OAS
EIVLTQSPGTLSLSPGERATLSCRASQSVSSSYLAWYQQKPGQAPRLLIYGASSRATGIPDRFSGSGSGTDFTLTISRLEPEDFAVYYCQQYGSSPCTFGQGTRLEIK
|.|||||||||||||||||||.|||||||||||||||||||||||||||||||||||||||||||||||||||||||||||||||||||||||||||||||.||||
ETVLTQSPGTLSLSPGERATLPCRASQSVSSSYLAWYQQKPGQAPRLLIYGASSRATGIPDRFSGSGSGTDFTLTISRLEPEDFAVYYCQQYGSSPCTFGQGTKLEIK
                     ^^^^^^^               ^^^                                ^^^^^^^^^

Best Alignment of Secukinumab CDR-H3 chain to a sequence from OAS
EVQLVESGGGLVQPGGSLRLSCAASGFTFSNYWMNWVRQAPGKGLEWVAAINQDGSEKYYVGSVKGRFTISRDNAKNSLYLQMNSLRVEDTAVYYCVRDYYDILTDYYIHYWYFDLWGRGTLVTVSS
......||.....||.||..||..||.....||..||||.|||||||...|....|...|..|..|..|||.|......|||..||...|||.|||.|.||||||.|||.|||||||||||||||||
-----QSGAEVKKPGESLKISCKGSGYSLTSYWIGWVRQMPGKGLEWMGIIYPGDSDTRYSPSFQGQVTISADKSISTAYLQWSSLKASDTAMYYCARLYYDILTGYYINYWYFDLWGRGTLVTVSS
                    ^^^^^^^^               ^^^^^^^^                                   ^^^^^^^^^^^^^^^^^^^

Therapeutic : Seribantumab

Best Alignment of Seribantumab heavy chain to a sequence from OAS
EVQLLESGGGLVQPGGSLRLSCAASGFTFSHYVMAWVRQAPGKGLEWVSSISSSGGWTLYADSVKGRFTISRDNSKNTLYLQMNSLRAEDTAVYYCTRGLKMATIFDYWGQGTLVTVSS
||||||||||||||||||||||||||||||.|.|.||||||||||||||.||.|||.|.||||||||||||||||||||||||||||||||||||||||..|||.||||||||||||||
EVQLLESGGGLVQPGGSLRLSCAASGFTFSSYAMSWVRQAPGKGLEWVSAISGSGGSTYYADSVKGRFTISRDNSKNTLYLQMNSLRAEDTAVYYCTRGWEMATHFDYWGQGTLVTVSS
                    ^^^^^^^^               ^^^^^^^^                                   ^^^^^^^^^^^^

Best Alignment of Seribantumab light chain to a sequence from OAS
QSALTQPASVSGSPGQSITISCTGTSSDVGSYNVVSWYQQHPGKAPKLIIYEVSQRPSGVSNRFSGSKSGNTASLTISGLQTEDEADYYCCSYAGSSIFVIFGGGTKVTVL
|||||||||||||||||||||||||||||||||.||||||||||||||.||||||||||||||||||||||||||||||||.|||||||||||||||.||.||||||||||
QSALTQPASVSGSPGQSITISCTGTSSDVGSYNLVSWYQQHPGKAPKLMIYEVSQRPSGVSNRFSGSKSGNTASLTISGLQAEDEADYYCCSYAGSSTFVVFGGGTKVTVL
                    ^^^^^^^^^               ^^^                                ^^^^^^^^^^^

Best Alignment of Seribantumab heavy chain CDRs to a sequence from OAS
EVQLLESGGGLVQPGGSLRLSCAASGFTFSHYVMAWVRQAPGKGLEWVSSISSSGGWTLYADSVKGRFTISRDNSKNTLYLQMNSLRAEDTAVYYCTRGLKMATIFDYWGQGTLVTVSS
..............|||||||.||||||||.|.|.||||||||||||||.||.|||.|.|||||||||||||||.||.||||||||||||||||||.|||||||.||||||||||||||
--------------GGSLRLSYAASGFTFSSYAMSWVRQAPGKGLEWVSAISGSGGSTYYADSVKGRFTISRDNAKNSLYLQMNSLRAEDTAVYYCARGLKMATPFDYWGQGTLVTVSS
                    ^^^^^^^^               ^^^^^^^^                                   ^^^^^^^^^^^^

Best Alignment of Seribantumab light CDRs chain to a sequence from OAS
QSALTQPASVSGSPGQSITISCTGTSSDVGSYNVVSWYQQHPGKAPKLIIYEVSQRPSGVSNRFSGSKSGNTASLTISGLQTEDEADYYCCSYAGSSIFVIFGGGTKVTVL
|||||||||||||||||.||||||||||||||||||||||||||||||.|||||.|||||.||||||||||||||||||||.||||||||||||||||||.||||||.|||
QSALTQPASVSGSPGQSLTISCTGTSSDVGSYNVVSWYQQHPGKAPKLMIYEVSKRPSGVANRFSGSKSGNTASLTISGLQAEDEADYYCCSYAGSSIFVVFGGGTKLTVL
                    ^^^^^^^^^               ^^^                                ^^^^^^^^^^^

Best Alignment of Seribantumab CDR-H3 chain to a sequence from OAS
EVQLLESGGGLVQPGGSLRLSCAASGFTFSHYVMAWVRQAPGKGLEWVSSISSSGGWTLYADSVKGRFTISRDNSKNTLYLQMNSLRAEDTAVYYCTRGLKMATIFDYWGQGTLVTVSS
.............|||||||||||||||||.|.|.||||||||||||||.||||.....|||||||||||||||.||.||||||||||||||||||.|||.||||||||||||||.|||
-------------PGGSLRLSCAASGFTFSSYSMNWVRQAPGKGLEWVSYISSSSSTIYYADSVKGRFTISRDNAKNSLYLQMNSLRAEDTAVYYCARGLEMATIFDYWGQGTLVNVSS
                    ^^^^^^^^               ^^^^^^^^                                   ^^^^^^^^^^^^

Therapeutic : Setoxaximab

Best Alignment of Setoxaximab heavy chain to a sequence from OAS
EVQLQQPGPELEKPGASVKLSCKASGYSFTDYNMNWVKQNNGESLEWIGKIDPYYGGPSYNQKFKDKATLTVDKSSSTAYMQLKSLTSEDSAVYYCTRGGNRDWYFDVWGAGTTLTVSA
||||||.||||||||||||.||||||||||.||||||||.||.||||||.|||||||.|||||||.||||||||||||||||||||||||||||||.|.||.||||||||||||.|||.
EVQLQQSGPELEKPGASVKISCKASGYSFTGYNMNWVKQSNGKSLEWIGNIDPYYGGTSYNQKFKGKATLTVDKSSSTAYMQLKSLTSEDSAVYYCARDGNYDWYFDVWGAGTTVTVSS
                    ^^^^^^^^               ^^^^^^^^                                   ^^^^^^^^^^^^

Best Alignment of Setoxaximab light chain to a sequence from OAS
DIVLSQSPSSLVVSVGEKVTMSCKSSQSLLYSRNQKNYLAWYQQKPGQSPKVLIYWASTRESGVPDRLTGSGSGTDFTLTISSVKAEDLAVYYCQQYYSYPLTFGAGTKLELK
|||.|||||||.|||||||||||||||||||||||||||||||||||||||.|||||||||||||||.|||||||||||||||||||||||||||||||||||||||||||||
DIVMSQSPSSLAVSVGEKVTMSCKSSQSLLYSRNQKNYLAWYQQKPGQSPKLLIYWASTRESGVPDRFTGSGSGTDFTLTISSVKAEDLAVYYCQQYYSYPLTFGAGTKLELK
^^^^^^^^^^^^ ^^^ ^^^^^^^^^

Best Alignment of Setoxaximab heavy chain CDRs to a sequence from OAS
EVQLQQPGPELEKPGASVKLSCKASGYSFTDYNMNWVKQNNGESLEWIGKIDPYYGGPSYNQKFKDKATLTVDKSSSTAYMQLKSLTSEDSAVYYCTRGGNRDWYFDVWGAGTTLTVSA
||.|...||||||||||||.||||||||||.||||||||.||.||||||.|||||||.|||||||.||||||||||||||||||||||||||||||.|||||.|||||||||||.|||.
EVKLVESGPELEKPGASVKISCKASGYSFTGYNMNWVKQSNGKSLEWIGNIDPYYGGTSYNQKFKGKATLTVDKSSSTAYMQLKSLTSEDSAVYYCARGGNRYWYFDVWGAGTTVTVSS
^^^^^^^^ ^^^^^^^^ ^^^^^^^^^^^^

Best Alignment of Setoxaximab light CDRs chain to a sequence from OAS
DIVLSQSPSSLVVSVGEKVTMSCKSSQSLLYSRNQKNYLAWYQQKPGQSPKVLIYWASTRESGVPDRLTGSGSGTDFTLTISSVKAEDLAVYYCQQYYSYPLTFGAGTKLELK
|||.|||||||.|||||||||||||||||||||||||||||||||||||||.|||||||||||||||.|||||||||||||||||||||||||||||||||||||||||||||
DIVMSQSPSSLAVSVGEKVTMSCKSSQSLLYSRNQKNYLAWYQQKPGQSPKLLIYWASTRESGVPDRFTGSGSGTDFTLTISSVKAEDLAVYYCQQYYSYPLTFGAGTKLELK
^^^^^^^^^^^^ ^^^ ^^^^^^^^^

Best Alignment of Setoxaximab CDR-H3 chain to a sequence from OAS
EVQLQQPGPELEKPGASVKLSCKASGYSFTDYNMNWVKQNNGESLEWIGKIDPYYGGPSYNQKFKDKATLTVDKSSSTAYMQLKSLTSEDSAVYYCTRGGNRDWYFDVWGAGTTLTVSA
.|||||.|.||..|||||||||||.||.||||.|.||||.....|||||.|.|..||...|||||.|||||.|||||||||.|.|||.||||||||||||.|||||||||||||.|||.
QVQLQQSGAELVRPGASVKLSCKALGYTFTDYEMHWVKQTSVHGLEWIGAIHPGSGGTAYNQKFKGKATLTADKSSSTAYMELSSLTAEDSAVYYCTRGGLRDWYFDVWGAGTTVTVSS
^^^^^^^^ ^^^^^^^^ ^^^^^^^^^^^^

Therapeutic : Setrusumab

Best Alignment of Setrusumab heavy chain to a sequence from OAS
QVQLVESGGGLVQPGGSLRLSCAASGFTFRSHWLSWVRQAPGKGLEWVSNINYDGSSTYYADSVKGRFTISRDNSKNTLYLQMNSLRAEDTAVYYCARDTY--LHFDYWGQGTLVTVSS
.||||||||||||||||||||||||||||.|.||.||||||||||.|||.||.|||||.|||||||||||||||||||||||||.||||||||||||||.|..||||||||||||||||
EVQLVESGGGLVQPGGSLRLSCAASGFTFSSYWLHWVRQAPGKGLVWVSRINSDGSSTSYADSVKGRFTISRDNAKNTLYLQMNSLRAEDTAVYYCARDPYQRLHFDYWGQGTLVTVSS
^^^^^^^^ ^^^^^^^^ ^^^^^^^^^^^^

Best Alignment of Setrusumab light chain to a sequence from OAS
DIALTQPASVSGSPGQSITISCTGTSSDVGDINDVSWYQQHPGKAPKLMIYDVNNRPSGVSNRFSGSKSGNTASLTISGLQAEDEADYYCQSYAGSYLSEVFGGGTKLTVL
..||||||||||||||||||||||||||||..|.|||||||||||||||||||.||||||||||||||||||||||||||||||||||||.||||||.|.|||||||||||
QSALTQPASVSGSPGQSITISCTGTSSDVGGYNYVSWYQQHPGKAPKLMIYDVTNRPSGVSNRFSGSKSGNTASLTISGLQAEDEADYYCSSYAGSYTSWVFGGGTKLTVL
^^^^^^^^^ ^^^ ^^^^^^^^^^^

Best Alignment of Setrusumab heavy chain CDRs to a sequence from OAS
QVQLVESGGGLVQPGGSLRLSCAASGFTFRSHWLSWVRQAPGKGLEWVSNINYDGSSTYYADSVKGRFTISRDNSKNTLYLQMNSLRAEDTAVYYCARDTYLHFDYWGQGTLVTVSS
.............||||||||||||||||.|||..||||||||||.|||.||.|||.|.||||||||||||.||.||.||||||||||||||||||||||.||||||||||||||||
-------------PGGSLRLSCAASGFTFSSHWMHWVRQAPGKGLVWVSRINSDGSRTTYADSVKGRFTISKDNAKNSLYLQMNSLRAEDTAVYYCARDTQLHFDYWGQGTLVTVSS
^^^^^^^^ ^^^^^^^^ ^^^^^^^^^^

Best Alignment of Setrusumab light CDRs chain to a sequence from OAS
DIALTQPASVSGSPGQSITISCTGTSSDVGDINDVSWYQQHPGKAPKLMIYDVNNRPSGVSNRFSGSKSGNTASLTISGLQAEDEADYYCQSYAGSYLSEVFGGGTKLTVL
..|||||.|||||||||.||||||||||||..|.|.||||||||||||||||||.|||||..|||||||||||||.||||||.|||||||.||||||||.|||.||..|||
QSALTQPRSVSGSPGQSVTISCTGTSSDVGGYNSVFWYQQHPGKAPKLMIYDVNKRPSGVPARFSGSKSGNTASLSISGLQADDEADYYCCSYAGSYLSRVFGSGTEVTVL
^^^^^^^^^ ^^^ ^^^^^^^^^^^

Best Alignment of Setrusumab CDR-H3 chain to a sequence from OAS
QVQLVESGGGLVQPGGSLRLSCAASGFTFRSHWLSWVRQAPGKGLEWVSNINYDGSSTYYADSVKGRFTISRDNSKNTLYLQMNSLRAEDTAVYYCARDTYLHFDYWGQGTLVTVSS
.|||...|..||.||.|..|||.||||........||.|.|.|..||||...|......|.|.....|..||..|.|.||.|||..||..|||||||||||.|||||||||||..||||
-VQLQQPGAELVKPGASVKLSCTASGFNIKDTYMHWVKQRPEQGLEWIGRIDPANGNTKYDPKFQGKATITADTSSNTAYLQLSSLTSEDTAVYYCARDGYLHFDYWGQGTTLTVSS
^^^^^^^^ ^^^^^^^^ ^^^^^^^^^^

Therapeutic : Sifalimumab

Best Alignment of Sifalimumab heavy chain to a sequence from OAS
QVQLVQSGAEVKKPGASVKVSCKASGYTFTSYSISWVRQAPGQGLEWMGWISVYNGNTNYAQKFQGRVTMTTDTSTSTAYLELRSLRSDDTAVYYCARDPI----AAGYWGQGTLVTVSS
||||||||||||||||||||||||||||||||.|||||||||||||||||||.|||||||||||||||||||||||||||.||||||||||||||||||.|....|.|||||||||||||
QVQLVQSGAEVKKPGASVKVSCKASGYTFTSYDISWVRQAPGQGLEWMGWISAYNGNTNYAQKFQGRVTMTTDTSTSTAYMELRSLRSDDTAVYYCARDGIVGVPAMGYWGQGTLVTVSS
^^^^^^^^ ^^^^^^^^ ^^^^^^^^^^^^^

Best Alignment of Sifalimumab light chain to a sequence from OAS
EIVLTQSPGTLSLSPGERATLSCRASQSVSSTYLAWYQQKPGQAPRLLIYGASSRATGIPDRFSGSGSGTDFTLTISRLEPEDFAVYYCQQYGSSPRTFGQGTKVEIK
||||||||||||||||||||||||||||||||||||||||||||||||||||||||||||||||||||||||||||||||||||||||||||||||||||||||||||
EIVLTQSPGTLSLSPGERATLSCRASQSVSSTYLAWYQQKPGQAPRLLIYGASSRATGIPDRFSGSGSGTDFTLTISRLEPEDFAVYYCQQYGSSPRTFGQGTKVEIK
^^^^^^^ ^^^ ^^^^^^^^^

Best Alignment of Sifalimumab heavy chain CDRs to a sequence from OAS
QVQLVQSGAEVKKPGASVKVSCKASGYTFTSYSISWVRQAPGQGLEWMGWISVYNGNTNYAQKFQGRVTMTTDTSTSTAYLELRSLRSDDTAVYYCARDPIAAGYWGQGTLVTVSS
................||||||||||||||||.||||||||||||||||||||||||||||||.|||||||.||||.|||.|||||||||.|||||||.||||.||||||||||||
----------------SVKVSCKASGYTFTSYGISWVRQAPGQGLEWMGWISVYNGNTNYAQKLQGRVTMTADTSTNTAYMELRSLRSDDSAVYYCARTPIAADYWGQGTLVTVSS
^^^^^^^^ ^^^^^^^^ ^^^^^^^^^

Best Alignment of Sifalimumab light CDRs chain to a sequence from OAS
EIVLTQSPGTLSLSPGERATLSCRASQSVSSTYLAWYQQKPGQAPRLLIYGASSRATGIPDRFSGSGSGTDFTLTISRLEPEDFAVYYCQQYGSSPRTFGQGTKVEIK
||||||||.|||||||||||||||||||||||||||||||||||||||||||||||||||||||||||||||||||||||||||||||||||||||||||.||||.||
EIVLTQSPDTLSLSPGERATLSCRASQSVSSTYLAWYQQKPGQAPRLLIYGASSRATGIPDRFSGSGSGTDFTLTISRLEPEDFAVYYCQQYGSSPRTFGPGTKVDIK
^^^^^^^ ^^^ ^^^^^^^^^

Best Alignment of Sifalimumab CDR-H3 chain to a sequence from OAS
QVQLVQSGAEVKKPGASVKVSCKASGYTFTSYSISWVRQAPGQGLEWMGWISVYNGNTNYAQKFQGRVTMTTDTSTSTAYLELRSLRSDDTAVYYCARDPIAAGYWGQGTLVTVSS
..............|.|...||.|||.|..|...||||||||.||.|...|....|.|.||....||.|...|.|..|.||...|||..|||||||||||||||||||||||||||
--------------GGSLRLSCAASGFTVSSNYMSWVRQAPGKGLGWVSVIYSG-GSTYYADSVKGRFTISRDNSKNTLYLQMNSLRAEDTAVYYCARDPIAAGYWGQGTLVTVSS
^^^^^^^^ ^^^^^^^^ ^^^^^^^^^

Therapeutic : Siltuximab

Best Alignment of Siltuximab heavy chain to a sequence from OAS
EVQLVESGGKLLKPGGSLKLSCAASGFTFSSFAMSWFRQSPEKRLEWVAEISSGGSYTYYPDTVTGRFTISRDNAKNTLYLEMSSLRSEDTAMYYCARGLWG---YYALDYWGQGTSVTVSS
|||||||||.|.|||||||||||||||||||.||||.||||||||||||||||||||||||||||||||||||||||||||||||||||||||||||||..|...|||.|||||||||||||
EVQLVESGGGLVKPGGSLKLSCAASGFTFSSYAMSWVRQSPEKRLEWVAEISSGGSYTYYPDTVTGRFTISRDNAKNTLYLEMSSLRSEDTAMYYCARGGSGYDGYYAMDYWGQGTSVTVSS
^^^^^^^^ ^^^^^^^^ ^^^^^^^^^^^^^^^

Best Alignment of Siltuximab light chain to a sequence from OAS
QIVLIQSPAIMSASPGEKVTMTCSASSSVSYMYWYQQKPGSSPRLLIYDTSNLASGVPVRFSGSGSGTSYSLTISRMEAEDAATYYCQQWSGYPYTFGGGTKLEIK
||||.|||||||||||||||||||||||||||||||||||||||||||||||||||||||||||||||||||||||||||||||||||||||||||||||||||||
QIVLTQSPAIMSASPGEKVTMTCSASSSVSYMYWYQQKPGSSPRLLIYDTSNLASGVPVRFSGSGSGTSYSLTISRMEAEDAATYYCQQWSGYPYTFGGGTKLEIK
^^^^^ ^^^ ^^^^^^^^^

Best Alignment of Siltuximab heavy chain CDRs to a sequence from OAS
EVQLVESGGKLLKPGGSLKLSCAASGFTFSSFAMSWFRQSPEKRLEWVAEISSGGSYTYYPDTVTGRFTISRDNAKNTLYLEMSSLRSEDTAMYYCARGLWGYYALDYWGQGTSVTVSS
|||.|||||.|.|||||||||||||||||||.||||.|||||||||||||||||||||||||.|||||||.||||||||||||||||||||||||||||||.||||.||||||||||||

EVQRVESGGGLVKPGGSLKLSCAASGFTFSSYAMSWVRQSPEKRLEWVAEISSGGSYTYYPDNVTGRFTIARDNAKNTLYLEMSSLRSEDTAMYYCARGLDGYYAMDYWGQGTSVTVSS
^^^^^^^^ ^^^^^^^^ ^^^^^^^^^^^^

Best Alignment of Siltuximab light CDRs chain to a sequence from OAS
QIVLIQSPAIMSASPGEKVTMTCSASSSVSYMYWYQQKPGSSPRLLIYDTSNLASGVPVRFSGSGSGTSYSLTISRMEAEDAATYYCQQWSGYPYTFGGGTKLEIK
||||.|||||||||||||||||||||||||||||||||||||||||||||||||||||||||||||||||||||||||||||||||||||||||||||||||||||
QIVLTQSPAIMSASPGEKVTMTCSASSSVSYMYWYQQKPGSSPRLLIYDTSNLASGVPVRFSGSGSGTSYSLTISRMEAEDAATYYCQQWSGYPYTFGGGTKLEIK
^^^^^ ^^^ ^^^^^^^^^

Best Alignment of Siltuximab CDR-H3 chain to a sequence from OAS
EVQLVESGGKLLKPGGSLKLSCAASGFTFSSFAMSWFRQSPEKRLEWVAEISSGGSYTYYPDTVTGRFTISRDNAKNTLYLEMSSLRSEDTAMYYCARGLWGYYALDYWGQGTSVTVSS
||||..||..|.|||.|.|.|.||...|.||.|....|..|.|...|||...|......|.|.....|..|...|....|.|.|.|||.|||.|.||||||||||||.|||||||||||||
EVQLQQSGPELVKPGASGKMSCKPPGYTFTSYIIHWGKQEPGQGLEWIGYINPYNDGTKYNEKFKGKATLTSDKSSGTAYMELSSLTSEDSAVYYCARGLWGYYAMDYWGQGTSVTVSS
^^^^^^^^ ^^^^^^^^ ^^^^^^^^^^^^

Therapeutic : Simtuzumab

Best Alignment of Simtuzumab heavy chain to a sequence from OAS
QVQLVQSGAEVKKPGASVKVSCKASGYAFTYYLIEWVRQAPGQGLEWIGVINPGSGGTNYNEKFKGRATITADKSTSTAYMELSSLRSEDTAVYFCARNWM--------NFDYWGQGTTVTVSS
||||.|||||..||||||||||||||||||.||||||.|.||||||||||||||||||||||||||.||.|||||.|||||.||||.|||.|||||||||..........|||||||||.||||
QVQLQQSGAELVKPGASVKVSCKASGYAFTNYLIEWVKQRPGQGLEWIGVINPGSGGTNYNEKFKGKATLTADKSSSTAYMQLSSLTSEDSAVYFCARNWDYYGSSYLYYFDYWGQGTTLTVSS
^^^^^^^^ ^^^^^^^^ ^^^^^^^^^^^^^^^^^

Best Alignment of Simtuzumab light chain to a sequence from OAS
DIVMTQTPLSLSVTPGQPASISCRSSKSLLHSNGNTYLYWFLQKPGQSPQFLIYRMSNLASGVPDRFSGSGSGTDFTLKISRVEAEDVGVYYCMQHLEYPYTFGGGTKVEIK
||||||...|..|||||..||||||||||||||||||||||||.||||||.|||||||||||||||||||||||||||.|||||||||||||||||||||||||||||.|||
DIVMTQAAPSVPVTPGQSVSISCRSSKSLLHSNGNTYLYWFLQRPGQSPQLLIYRMSNLASGVPDRFSGSGSGTDFTLRISRVEAEDVGVYYCMQHLEYPYTFGGGTKLEIK
^^^^^^^^^^^ ^^^ ^^^^^^^^^

Best Alignment of Simtuzumab heavy chain CDRs to a sequence from OAS
QVQLVQSGAEVKKPGASVKVSCKASGYAFTYYLIEWVRQAPGQGLEWIGVINPGSGGTNYNEKFKGRATITADKSTSTAYMELSSLRSEDTAVYFCARNWMNFDYWGQGTTVTVSS
||||.|||||...||.||||||||||||||.||||||.|.||||||||||||||||||||||||||.||.|||||.|||||.||||.|||.||.||||||.||||||||||.||||
QVQLQQSGAELVRPGTSVKVSCKASGYAFTNYLIEWVKQRPGQGLEWIGVINPGSGGTNYNEKFKGKATLTADKSSSTAYMQLSSLTSEDSAVFFCARNWENFDYWGQGTTLTVSS
^^^^^^^^ ^^^^^^^^ ^^^^^^^^^

Best Alignment of Simtuzumab light CDRs chain to a sequence from OAS
DIVMTQTPLSLSVTPGQPASISCRSSKSLLHSNGNTYLYWFLQKPGQSPQFLIYRMSNLASGVPDRFSGSGSGTDFTLKISRVEAEDVGVYYCMQHLEYPYTFGGGTKVEIK
.|||.|...|..||||...||||||||||||||||||||||||.||||||.|||||||||||||||||||||||.|||.|||||||||||||||||||||||||||||.|||
-IVMMQAAPSVPVTPGESVSISCRSSKSLLHSNGNTYLYWFLQRPGQSPQLLIYRMSNLASGVPDRFSGSGSGTAFTLRISRVEAEDVGVYYCMQHLEYPYTFGGGTKLEIK
^^^^^^^^^^^ ^^^ ^^^^^^^^^

Best Alignment of Simtuzumab CDR-H3 chain to a sequence from OAS
QVQLVQSGAEVKKPGASVKVSCKASGYAFTYYLIEWVRQAPGQGLEWIGVINPGSGGTNYNEKFKGRATITADKSTSTAYMELSSLRSEDTAVYFCARNWMNFDYWGQGTTVTVSS
..............|.|...||.|||..|..|...|||.|||.||||...|...|....|....|||.||..|......|....|||.||||||.|||||||||||||||.|||||
--------------GGSLRLSCAASGFTFSSYSLNWVRLAPGKGLEWVSSISRDSDYIYYADSVKGRFTISRDNARNSLYLQMNSLRVEDTAVYYCARNWMNFDYWGQGTLVTVSS
^^^^^^^^ ^^^^^^^^ ^^^^^^^^^

Therapeutic : Sirukumab

Best Alignment of Sirukumab heavy chain to a sequence from OAS
EVQLVESGGGLVQPGGSLRLSCAASGFTFSPFAMSWVRQAPGKGLEWVAKISPGGSWTYYSDTVTGRFTISRDNAKNSLYLQMNSLRAEDTAVYYCARQLWG------------YYALDIWGQGTTVTVSS
||||||||||||||||||||||||||||||...||||||||||||||||.|...||.|||.|.|.|||||||||||||||||||||||||||||||||..||............||..|.|||||||||||
EVQLVESGGGLVQPGGSLRLSCAASGFTFSSYWMSWVRQAPGKGLEWVANIKQDGSETYYVDSVKGRFTISRDNAKNSLYLQMNSLRAEDTAVYYCAREGWGCSGGSCYGTYYYYYGMDVWGQGTTVTVSS
^^^^^^^^ ^^^^^^^^ ^^^^^^^^^^^^^^^^^^^^^^^^

Best Alignment of Sirukumab light chain to a sequence from OAS
EIVLTQSPATLSLSPGERATLSCSASISV-SYMYWYQQKPGQAPRLLIYDMSNLASGIPARFSGSGSGTDFTLTISSLEPEDFAVYYCMQWSGYPYTFGGGTKVEIK
|||||||||||||||||||||||.||.||.||..||||||||||||||||.||.||||||||||||||||||||||||||||||||||.|.|..|.|||||||||||
EIVLTQSPATLSLSPGERATLSCRASQSVNSYLAWYQQKPGQAPRLLIYDASNRASGIPARFSGSGSGTDFTLTISSLEPEDFAVYYCQQRSDWPLTFGGGTKVEIK
^^^^^^ ^^^ ^^^^^^^^^

Best Alignment of Sirukumab heavy chain CDRs to a sequence from OAS
EVQLVESGGGLVQPGGSLRLSCAASGFTFSPFAMSWVRQAPGKGLEWVAKISPGGSWTYYSDTVTGRFTISRDNAKNSLYLQMNSLRAEDTAVYYCARQLWGYYALDIWGQGTTVTVSS
||..||||||||.|||||.|||||||||||..|||||||.|...|||||.||||||.|||.|.|.||.|||||||||.|||||..|.|||.|||||||.|..|||||.|||||.|||||
EVHVVESGGGLVKPGGSLKLSCAASGFTFSSYAMSWVRQTPEQRLEWVATISPGGSYTYYPDNVKGRVTISRDNAKNNLYLQMSHLKSEDSAVYYCARGLGIYYALDYWGQGTSVTVSS
^^^^^^^^ ^^^^^^^^ ^^^^^^^^^^^^

Best Alignment of Sirukumab light CDRs chain to a sequence from OAS
EIVLTQSPATLSLSPGERATLSCSASISVSYMYWYQQKPGQAPRLLIYDMSNLASGIPARFSGSGSGTDFTLTISSLEPEDFAVYYCMQWSGYPYTFGGGTKVEIK
.||||||||..|.||||..|..||||.|||||||||||||..|||||||.||||||.|.|||||||||...||||..|.||.|.|||.||||||||||||||.|||
QIVLTQSPAIMSASPGEKVTMTCSASSSVSYMYWYQQKPGSSPRLLIYDTSNLASGVPVRFSGSGSGTSYSLTISRMEAEDAATYYCQQWSGYPYTFGGGTKLEIK
^^^^^ ^^^ ^^^^^^^^^

Best Alignment of Sirukumab CDR-H3 chain to a sequence from OAS
EVQLVESGGGLVQPGGSLRLSCAASGFTF-SPFAMSWVRQAPGKGLEWVAKISPGGSWTYYSDTVTGRFTISRDNAKNSLYLQMNSLRAEDTAVYYCARQLWGYYALDIWGQGTTVTVSS
.|||.|||..||.|.|.|.|.|..||.........|||||.|||||||...|...............|.|||..|.||...|.|.|..|.|||||||||.|.||||||||||||.|||||
QVQLQESGPRLVKPSGTLSLTCSVSGDPISQGHWWSWVRQPPGKGLEWIGAIFHI-GSINSNPPLKSRVTISVANSKNQFSLEMTSVTAADTAVYYCARYLSGYYALDIWGQGTLVTVSS
^^^^^^^^^ ^^^^^^^^ ^^^^^^^^^^^^

Therapeutic : Solanezumab

Best Alignment of Solanezumab heavy chain to a sequence from OAS
EVQLVESGGGLVQPGGSLRLSCAASGFTFSRYSMSWVRQAPGKGLELVAQINSVGNSTYYPDTVKGRFTISRDNAKNTLYLQMNSLRAEDTAVYYCASG----DYWGQGTLVTVSS
||||||||||||||||||||||||||||||.|||.|||||||||||.|..|||.|.||.|.|.|||||||||||||||||||||||||||||||||||||....|||||||||||||
EVQLVESGGGLVQPGGSLRLSCAASGFTFSSYSMNWVRQAPGKGLEWVSRINSDGSSTSYADSVKGRFTISRDNAKNTLYLQMNSLRAEDTAVYYCASGGVVVDYWGQGTLVTVSS
^^^^^^^^ ^^^^^^^^ ^^^^^^^^^

Best Alignment of Solanezumab light chain to a sequence from OAS
DVVMTQSPLSLPVTLGQPASISCRSSQSLIYSDGNAYLHWFLQKPGQSPRLLIYKVSNRFSGVPDRFSGSGSGTDFTLKISRVEAEDVGVYYCSQSTH-VPWTFGQGTKVEIK
|||||||||||||||||||||||||||||.|||||.|||||.|.||||||.||||||||.|||||||||||||||||||||||||||||||||.||||..|||||||||||||
DVVMTQSPLSLPVTLGQPASISCRSSQSLVYSDGNIYLHWFQQRPGQSPRRLIYKVSNRDSGVPDRFSGSGSGTDFTLKISRVEAEDVGVYYCMQSTHWPPWTFGQGTKVEIK
^^^^^^^^^^^ ^^^ ^^^^^^^^^^

Best Alignment of Solanezumab heavy chain CDRs to a sequence from OAS
EVQLVESGGGLVQPGGSLRLSCAASGFTFSRYSMSWVRQAPGKGLELVAQINSVGNSTYYPDTVKGRFTISRDNAKNTLYLQMNSLRAEDTAVYYCASGDYWGQGTLVTVSS
..............||||||||||||||||.|.|.||||||||||..|..|||||.||.|.|.||||||||||||||||||||||||||||||||||.||||||||||||||
--------------GGSLRLSCAASGFTFSSYWMHWVRQAPGKGLVWVSRINSVGSSTSYADSVKGRFTISRDNAKNTLYLQMNSLRAEDTAVYYCARGDYWGQGTLVTVSS
^^^^^^^^ ^^^^^^^^ ^^^^^

Best Alignment of Solanezumab light CDRs chain to a sequence from OAS
DVVMTQSPLSLPVTLGQPASISCRSSQSLIYSDGNAYLHWFLQKPGQSPRLLIYKVSNRFSGVPDRFSGSGSGTDFTLKISRVEAEDVGVYYCSQSTHVPWTFGQGTKVEIK
..|||||||||||.||...||||||||||.||||||||||.||||||||.||||.||||||||||||||||||||||||||||||||.|||.||||||||||||.|||.|||
--VMTQSPLSLPVSLGDQTSISCRSSQSLVYSDGNAYLHWYLQKPGQSPELLIYTVSNRFSGVPDRFSGSGSGTDFTLKISRVEAEDLGVYFCSQSTHVPWTFGGGTKLEIK
^^^^^^^^^^^ ^^^ ^^^^^^^^^

Best Alignment of Solanezumab CDR-H3 chain to a sequence from OAS
EVQLVESGGGLVQPGGSLRLSCAASGFTFSRYSMSWVRQAPGKGLELVAQINSVGNSTYYPDTVKGRFTISRDNAKNTLYLQMNSLRAEDTAVYYCASGDYWGQGTLVTVSS
||||..|...||.||.|....|.|||.||..|.|.||.|..||.||....||.....|.|....||..|...|....|.|....||..||||||||||||||||||.|||||

```
EVQLQQSRPELVKPGASVKIPCKASGYTFTDYNMDWVNQSHGKSLEWIGDINPNNGGTIYNQKFKGKATLTVDKSSSTAYMELRSLTSEDTAVYYCASGDYWGQGTSVTVSS
                  ^^^^^^^^              ^^^^^^^^                              ^^^^^
```

Therapeutic : Spartalizumab

Best Alignment of Spartalizumab heavy chain to a sequence from OAS

```
EVQLVQSGAEVKKPGESLRISCKGSGYTFTTYWMHWVRQATGQGLEWMGNIYPGTGGSNFDEKFKNRVTITADKSTSTAYMELSSLRSEDTAVYYCTRWTT----GTGAYWGQGTTVTVSS
..|||||||||||||||.||||||||||||||..|||||.||||||||.|.|..|..|...||..||||||||||||||||||||||||||||||.|.......|....|||||||||||
--QLVQSGAEVKKPGESLKISCKGSGYTFTTYWIAWVRQAPGQGLEWMGRIIPILGIANYAQKFQGRVTITADKSTSTAYMELSSLRSEDTAVYYCARGSSTWFLGAFDIWGQGTTVTVSS
                  ^^^^^^^^              ^^^^^^^^                              ^^^^^^^^^^^^^^
```

Best Alignment of Spartalizumab light chain to a sequence from OAS

```
EIVLTQSPATLSLSPGERATLSCKSSQSLLDSGNQKNFLTWYQQKPGQAPRLLIYWASTRESGVPSRFSGSGSGTDFTFTISSLEAEDAATYYCQNDYSYPYTFGQGTKVEIK
|||.||||||||.|||||||||||||||.|.|.|.||.|.||||||||.|.|||||||||||||||.||||||||||||.|||||.|||.|.||||..||.|.|||||||||||
EIVMTQSPATLSVSPGERATLSCKSSQSVLYSSNNKNYLAWYQQKPGQPPKLLIYWASTRESGVPDRFSGSGSGTDFTLTISSLQAEDVAVYYCQQYYSTPWTFGQGTKVEIK
                   ^^^^^^^^^^^^             ^^^                         ^^^^^^^^^
```

Best Alignment of Spartalizumab heavy chain CDRs to a sequence from OAS

```
EVQLVQSGAEVKKPGESLRISCKGSGYTFTTYWMHWVRQATGQGLEWMGNIYPGTGGSNFDEKFKNRVTITADKSTSTAYMELSSLRSEDTAVYYCTRWTTGTGAYWGQGTTVTVSS
.|||.|.|||..|||.|...|||.||||||.||||||.|..||||||.||||||.|..|.|||||...|.|.|.|.|||||.||||.|||.||||||||.||.||||||||||.||||.
QVQLQQPGAELVKPGASVKLSCKASGYTFTSYWMHWVKQRPGQGLEWIGNIYPGSGSTNYDEKFKSKATLTVDTSSSTAYMQLSSLTSEDSAVYYCTRGTTATGAYWGQGTLVTVSA
                  ^^^^^^^^              ^^^^^^^^                              ^^^^^^^^^^
```

Best Alignment of Spartalizumab light CDRs chain to a sequence from OAS

```
EIVLTQSPATLSLSPGERATLSCKSSQSLLDSGNQKNFLTWYQQKPGQAPRLLIYWASTRESGVPSRFSGSGSGTDFTFTISSLEAEDAATYYCQNDYSYPYTFGQGTKVEIK
.||.||||..|....||..|.|||||||||.||||||.||||||||||.|.||||||||||||||.||.|||||||||.||||..|||.|.||||||||||||||.|||.|||
DIVMTQSPSSLTVTAGEKVTMSCKSSQSLLNSGNQKNYLTWYQQKPGQPPKLLIYWASTRESGVPDRFTGSGSGTDFTLTISSVQAEDLAVYYCQNDYSYPYTFGSGTKLEIK
                   ^^^^^^^^^^^^             ^^^                         ^^^^^^^^^
```

Best Alignment of Spartalizumab CDR-H3 chain to a sequence from OAS

```
EVQLVQSGAEVKKPGESLRISCKGSGYTFTTYWMHWVRQATGQGLEWMGNIYPGTGGSNFDEKFKNRVTITADKSTSTAYMELSSLRSEDTAVYYCTRWTTGTGAYWGQGTTVTVSS
.|||.|||||...||.|...|||.||||||.||..||.|..||||||.|||||.....|...|||...|.|.|||.|||||.|||..|||.||||||||.|||||||||||.||||.
QVQLQQSGAELVRPGASVKLSCKASGYTFTSYWINWVKQRPGQGLEWIGNIYPSDSYTNYNQKFKDKATLTVDKSSSTAYMQLSSPTSEDSAVYYCTRWGTGTGAYWGQGTLVTVSA
                  ^^^^^^^^              ^^^^^^^^                              ^^^^^^^^^^
```

Therapeutic : Suvratoxumab

Best Alignment of Suvratoxumab heavy chain to a sequence from OAS

```
EVQLVESGGGLVQPGGSLRLSCAASGFTFSSHDMHWVRQATGKGLEWVSGIGTAGDTYYPDSVKGRFTISRENAKNSLYLQMNSLRAGDTAVYYCARDRYSPT--GHYYGMDVWGQGTTVTVSS
|||||||||||||||||||||||||||||||.||||||||||||||||||.||||||||||.||||||||||||||||||||||||||||||||||||||||...||..|.||||||||||||||||
EVQLVESGGGLVQPGGSLRLSCAASGFTFSSYDMHWVRQATGKGLEWVSAIGTAGDTYYPGSVKGRFTISRENAKNSLYLQMNSLRAGDTAVYYCARDVRMPTKHGNYYGMDVWGQGTTVTVSS
                  ^^^^^^^^              ^^^^^^^                               ^^^^^^^^^^^^^^^^^^
```

Best Alignment of Suvratoxumab light chain to a sequence from OAS

```
DIQMTQSPSTLSASVGDRVTITCRASQSISSWLAWYQQKPGKAPKLLIYKASSLESGVPSRFSGSGSGTEFTLTISSLQPDDFATYYCKQYADYWTFGQGTKVEIK
|||||||||||||||||||||||||||||||||||||||||||||||||||||||||||||||||||||||||||||||||||||||.||.||||||||||||||
DIQMTQSPSTLSASVGDRVTITCRASQSISSWLAWYQQKPGKAPKLLIYKASSLESGVPSRFSGSGSGTEFTLTISSLQPDDFATYYCQQYNDYWTFGQGTKVEIK
                   ^^^^^^             ^^^                         ^^^^^^^^
```

Best Alignment of Suvratoxumab heavy chain CDRs to a sequence from OAS

```
EVQLVESGGGLVQPGGSLRLSCAASGFTFSSHDMHWVRQATGKGLEWVSGIGTAGDTYYPDSVKGRFTISRENAKNSLYLQMNSLRAGDTAVYYCARDRYSPTGHYYGMDVWGQGTTVTVSS
.............|||||||||||||||||.|||||||||||||||||.||||||||||||||||||||.|.||.|||.||||||.|||||||||||.||..||||||||||||||||||||
-------------GGSLRLSCAASGFTFSSYDMHWVRQATGKGLEWVSAIGTAGDTYYPDSVKGRFTISRDNSKNTLYLLMNSLRAEDTAVYYCARDDYSIMGHYYGMDVWGQGTTVTVSS
                  ^^^^^^^^              ^^^^^^^                               ^^^^^^^^^^^^^^^^
```

Best Alignment of Suvratoxumab light CDRs chain to a sequence from OAS

```
DIQMTQSPSTLSASVGDRVTITCRASQSISSWLAWYQQKPGKAPKLLIYKASSLESGVPSRFSGSGSGTEFTLTISSLQPDDFATYYCKQYADYWTFGQGTKVEIK
||||||||||||||||||||||||||||||||||||||||||||||.|.|||||.|||||||||||||||||||||||||||||||||||||.|||||||||||||||||
DIQMTQSPSTLSASVGDRVTITCRASQSISSWLAWYQQKPGKAPELVIYKASRLESGVPSRFSGSGSGTEFTLTISSLQPDDFATYYCQQYADYWTFGQGTKVEIK
                   ^^^^^^             ^^^                         ^^^^^^^^
```

Best Alignment of Suvratoxumab CDR-H3 chain to a sequence from OAS

```
EVQLVESGGGLVQPGGSLRLSCAASGFTF--SSHDMHWVRQATGKGLEWVSGIGTAGDTYYPDSVKGRFTISRENAKNSLYLQMNSLRAGDTAVYYCARDRYSPTGHYYGMDVWGQGTTVTVSS
................|.|.|..||...........|.||..||||||.......|.|||..|.|.|.|||....||...|...|..|.||||||||||||||..||||||||||||||||||.
-------------SETLSLTCTVSGGSISSGDYYWSWIRQPPGKGLEWIGYFYYSGSTYYNPSLKSRVTISVDTSKNQFSLKLSSVTAADTAVYYCARDRYSPPIHYYGMDVWGQGTTVTVS-
                  ^^^^^^^^^^             ^^^^^^^                               ^^^^^^^^^^^^^^^^
```

Therapeutic : Tabalumab

Best Alignment of Tabalumab heavy chain to a sequence from OAS

```
QVQLQQWGAGLLKPSETLSLTCAVYGGSFSGYYWSWIRQPPGKGLEWIGEINHSGSTNYNPSLKSRVTISVDTSKNQFSLKLSSVTAADTAVYYCARGYYDILT--GYYYYFDYWGQGTLVTVSS
||||||||||||||||||||||||||||||||||||||||||||||||||||||||||||||||||||||||||||||||||||||||||||||||||||||||....||||||||||||||||||
QVQLQQWGAGLLKPSETLSLTCAVYGGSFSGYYWSWIRQPPGKGLEWIGEINHSGSTNYNPSLKSRVTISVDTSKNQFSLKLSSVTAADTAVYYCARGYYDILTGYSGYYYFDYWGQGTLVTVSS
                  ^^^^^^^^              ^^^^^^^                              ^^^^^^^^^^^^^^^^^^^
```

Best Alignment of Tabalumab light chain to a sequence from OAS

```
EIVLTQSPATLSLSPGERATLSCRASQSVSRYLAWYQQKPGQAPRLLIYDASNRATGIPARFSGSGSGTDSTLTISSLEPEDFAVYYCQQRSNWPRTFGQGTKVEIK
||||||||||||||||||||||||||||||||||||||||||||||||||||||||||||||||||||||.|||||||||||||||||||||||||||||||||||||
EIVLTQSPATLSLSPGERATLSCRASQSVSRYLAWYQQKPGQAPRLLIYDASNRATGIPARFSGSGSGTDFTLTISSLEPEDFAVYYCQQRSNWPRTFGQGTKVEIK
                   ^^^^^^             ^^^                         ^^^^^^^^^
```

Best Alignment of Tabalumab heavy chain CDRs to a sequence from OAS

```
QVQLQQWGAGLLKPSETLSLTCAVYGGSFSGYYWSWIRQPPGKGLEWIGEINHSGSTNYNPSLKSRVTISVDTSKNQFSLKLSSVTAADTAVYYCARGYYDILTGYYYYFDYWGQGTLVTVSS
.............|||||||||||||||||||||||||||||||||||||||||||||||||||||.|||||||||||||||||||.|||||||||||.||||||||||||||||
-------------SETLSLTCAVYGGSFSGYYWSWIRQPPGKGLEWIGEINHSGSTNYNPSLKSRVTISVDTSENQFSLKLSSVTAADTAVYYRARGYYDILTGYLYYFDYWGQGTLVTVSS
                  ^^^^^^^^              ^^^^^^^                              ^^^^^^^^^^^^^^^^^
```

Best Alignment of Tabalumab light CDRs chain to a sequence from OAS

```
EIVLTQSPATLSLSPGERATLSCRASQSVSRYLAWYQQKPGQAPRLLIYDASNRATGIPARFSGSGSGTDSTLTISSLEPEDFAVYYCQQRSNWPRTFGQGTKVEIK
||||||||||||||||||||||||||||||||||||||||||||||||||||||||||||||||||||||.|||||||||||||||||||||||||||||||||||||
EIVLTQSPATLSLSPGERATLSCRASQSVSRYLAWYQQKPGQAPRLLIYDASNRATGIPARFSGSGSGTDFTLTISSLEPEDFAVYYCQQRSNWPRTFGQGTKVEIK
                   ^^^^^^             ^^^                         ^^^^^^^^^
```

Best Alignment of Tabalumab CDR-H3 chain to a sequence from OAS

```
QVQLQQWGAGLLKPSETLSLTCAVYGGSFSGYYWSWIRQPPGKGLEWIGEINHS-GSTNYNPSLKSRVTISVDTSKNQFSLKLSSVTAADTAVYYCARGYYDILTGYYYYFDYWGQGTLVTVSS
............|.|.|...|...|.||..|...|.||.|||||||.|.|......|.|.||....||||.|.|.....|..||..|.|||.||||||||||||||||||||||||||||||
------------PGESLKISCKGSGYSFTSYWIGWVRQMPGKGLEWMGIIYPGDSDTRYSPSFQGQVTISADKSISTAYLQWSSLKASDTAMYYCARGYYDILTGYYYYFDYWGQGTLVTVSS
                  ^^^^^^^^              ^^^^^^^^                              ^^^^^^^^^^^^^^^^
```

Therapeutic : Tanezumab

Best Alignment of Tanezumab heavy chain to a sequence from OAS

```
QVQLQESGPGLVKPSETLSLTCTVSGFSLIGYDLNWIRQPPGKGLEWIGIIWGD-GTTDYNSAVKSRVTISKDTSKNQFSLKLSSVTAADTAVYYCARGGYWYA--TSYYFDYWGQGTLVTVSS
|||||||||||||||||||||.|||.|..||...||||||||||||||.|.|..|.||||...|||||||.||||||||||||||||||||||||||.||.|....||||||||||.||||||
QVQLQESGPGLVKPSETLSLTCAVSGGSFSGYYWGWIRQPPGKGLEWIGYIRGSRGSTDYNPSLKSRVTISTDTSKNQFSLKLSSVTAADTAVYYCARKGYYYSGSPSYYFDYWGQGVLVTVSS
                  ^^^^^^^^              ^^^^^^^^                              ^^^^^^^^^^^^^^^^
```

Best Alignment of Tanezumab light chain to a sequence from OAS
```
DIQMTQSPSSLSASVGDRVTITCRASQSISNNLNWYQQKPGKAPKLLIYYTSRFHSGVPSRFSGSGSGTDFTFTISSLQPEDIATYYCQQEHTLPYTFGQGTKLEIK
||||||||||||||||||||||||||||||..|||||||||||||||||.||...|||||||||||||||||||||||||||||||||||...||||||||||||||
DIQMTQSPSSLSASVGDRVTITCRASQSISTYLNWYQQKPGKAPKLLIYATSSLQSGVPSRFSGSGSGTDFTFTISSLQPEDIATYYCQQYDNLPYTFGQGTKLEIK
                        ^^^^^^          ^^^                       ^^^^^^^^^
```

Best Alignment of Tanezumab heavy chain CDRs to a sequence from OAS
```
QVQLQESGPGLVKPSETLSLTCTVSGFSLIGYDLNWIRQPPGKGLEWIGIIWGDGTTDYNSAVKSRVTISKDTSKNQFSLKLSSVTAADTAVYYCARGGYWYATSYYFDYWGQGTLVTVSS
.|||.|||||||.||..||.|||||||||.||..||.||||||||||.|.|||||.||||||.|||..||||.||.|..||..|....|||.||||||||.|..|||||||||||..||||
-VQLKESGPGLVAPSQSLSITCTVSGFSLTGYGVNWVRQPPGKGLEWLGMIWGDGSTDYNSALKSRLSISKDNSKSQVFLKMNSLQTDDTARYYCARGGYYYGSSYYFDYWGQGTTLTVSS
                        ^^^^^^^^          ^^^^^^^                        ^^^^^^^^^^^^^^^
```

Best Alignment of Tanezumab light CDRs chain to a sequence from OAS
```
DIQMTQSPSSLSASVGDRVTITCRASQSISNNLNWYQQKPGKAPKLLIYYTSRFHSGVPSRFSGSGSGTDFTFTISSLQPEDIATYYCQQEHTLPYTFGQGTKLEIK
||||||..||||||.||||||.|||||.||||||||||||....|||||||||.||||||||||||||||...|||.|..||||||.|||..||||||||.|||||||
DIQMTQTTSSLSASLGDRVTISCRASQDISNNLNWYQQKPDGTVKLLIYYTSRLHSGVPSRFSGSGSGTDYSLTISNLEQEDIATYFCQQGNTLPYTFGGGTKLEIK
                        ^^^^^^          ^^^                       ^^^^^^^^^
```

Best Alignment of Tanezumab CDR-H3 chain to a sequence from OAS
```
QVQLQESGPGLVKPSETLSLTCTVSGFSLIGYDLNWIRQPPGKGLEWIGIIWGD-GTTDYNSAVKSRVTISKDTSKNQFSLKLSSVTAADTAVYYCARGGYWYATSYYFDYWGQGTLVTVSS
.||||..|..||.|.....|.|..||.....|...|..|.||.||||||.|.......|.||...|...|...|.|.......|||.|..|.|||||||||||.||.|||||||||||..||||
-VQLQQPGAELVRPGSSVKLSCKASGYTFTSYWMDWVKQRPGQGLEWIGNIYPSDSETHYNQKFKDKATLTVDKSSSTAYMQLSSLTSEDSAVYYCARGGYYYAISYYFDYWGQGTTLTVSS
                        ^^^^^^^^          ^^^^^^^^                        ^^^^^^^^^^^^^^^
```

Therapeutic : Tarextumab

Best Alignment of Tarextumab heavy chain to a sequence from OAS
```
EVQLVESGGGLVQPGGSLRLSCAASGFTFSSSGMSWVRQAPGKGLEWVSVIASSGSNTYYADSVKGRFTISRDNSKNTLYLQMNSLRAEDTAVYYCARSI-----FYTTWGQGTLVTVSS
||||||||||||||||||||||||||||||||..|||||||||||||||||.|..||..|||||||||||||||||||||||||||||||||||||||||.......||.|||||||||||
EVQLVESGGGLVQPGGSLRLSCAASGFTFSSYAMSWVRQAPGKGLEWVSAISGSGGSTYYADSVKGRFTISRDNSKNTLYLQMNSLRAEDTAVYYCARSPGDSSGYYTYWGQGTLVTVSS
                        ^^^^^^^^          ^^^^^^^^                        ^^^^^^^^^^^^^
```

Best Alignment of Tarextumab light chain to a sequence from OAS
```
DIVLTQSPATLSLSPGERATLSCRASQSVRSNYLAWYQQKPGQAPRLLIYGASSRATGVPARFSGSGSGTDFTLTISSLEPEDFAVYYCQQYSNFPITFGQGTKVEIK
||||||||||||||||||||||||||||||.|||||||||||||||||||||.||.||||.|||||||||||||||||||||||||||||||||||||||||.|.|||||||||||
DIVLTQSPATLSLSPGERATLSCRASQSV-SNYLAWYQQKPGQAPRLLIYDASNRATGIPARFSGSGSGTDFTLTISSLEPEDFAVYYCQQYSNSPRTFGQGTKVEIK
                        ^^^^^^^          ^^^                       ^^^^^^^^^
```

Best Alignment of Tarextumab heavy chain CDRs to a sequence from OAS
```
EVQLVESGGGLVQPGGSLRLSCAASGFTFSSSGMSWVRQAPGKGLEWVSVIASSGSNTYYADSVKGRFTISRDNSKNTLYLQMNSLRAEDTAVYYCARSIFYTTWGQGTLVTVSS
..............|||||||||||||||||.|||||||||||||||||.|||||.||||||||||||||||||||||..||.||||.||||.|||||.|...|||.||||||||
--------------GGSLRLSCAASGFTFSSSAMSWVRQAPGKGLEWVSLIASSGLNTYYADSVKGRFTISRDNSKNTVSLQINSLRVEDTAIYYCARDIELSTWGLGTLVTVSS
                        ^^^^^^^^          ^^^^^^^^                        ^^^^^^^^
```

Best Alignment of Tarextumab light CDRs chain to a sequence from OAS
```
DIVLTQSPATLSLSPGERATLSCRASQSVRSNYLAWYQQKPGQAPRLLIYGASSRATGVPARFSGSGSGTDFTLTISSLEPEDFAVYYCQQYSNFPITFGQGTKVEIK
.|||||||.|||||||||||.||||.|||||||.|||||.||||||.||||||.||||.|.|.||||||.|||||||.|||||||||||||||.|||.||||||||.|||
EIVLTQSPGTLSLSPGERATVSCRATQSVRSNYVAWYQQRPGQAPRFLIYGASNRATGIPDRLSGSGSGADFTLTISRLEPEDFAVYYCQQYGNFPYTFGQGTKLEIK
                        ^^^^^^^          ^^^                       ^^^^^^^^^
```

Best Alignment of Tarextumab CDR-H3 chain to a sequence from OAS
```
EVQLVESGGGLVQPGGSLRLSCAASGFTFSSSGMSWVRQAPGKGLEWVSVIASSGSNTYYADSVKGRFTISRDNSKNTLYLQMNSLRAEDTAVYYCARSIFYTTWGQGTLVTVSS
..............|.||..|||||||||||..|.||||||||||.|||.|.|.||.|.|||||||||||||||.||||||||||||||||||||||||..|||||||||||||
--------------GESLKISCAASGFTFSSYWMHWVRQAPGKGLVWVSRINSDGSSTSYADSVKGRFTISRDNAKNTLYLQMNSLRAEDTAVYYCARSITMTTWGQGTLVTVSS
                        ^^^^^^^^          ^^^^^^^^                        ^^^^^^^^
```

Therapeutic : Teplizumab

Best Alignment of Teplizumab heavy chain to a sequence from OAS
```
QVQLVQSGGGVVQPGRSLRLSCKASGYTFTRYTMHWVRQAPGKGLEWIGYINPSRGYTNYNQKVKDRFTISRDNSKNTAFLQMDSLRPEDTGVYFCARYYDD-HYCLDYWGQGTPVTVSS
|||||.||||||||||||||||.|||.||..|.|.|.||||||||||..||..|..||||...||.|||||||||||||..|||.|||.|||.||.|||.|.|..|..||||||||.|||||
QVQLVESGGGVVQPGRSLRLSCAASGFTFSDYYMSWIRQAPGKGLEWVSYISSSSSYTNYADSVKGRFTISRDNSKNTLYLQMNSLRAEDTAVYYCARNYGDYGYYFDYWGQGTLVTVSS
                        ^^^^^^^^          ^^^^^^^^                        ^^^^^^^^^^^^^
```

Best Alignment of Teplizumab light chain to a sequence from OAS
```
DIQMTQSPSSLSASVGDRVTITCSASSSV-SYMNWYQQTPGKAPKRWIYDTSKLASGVPSRFSGSGSGTDYTFTISSLQPEDIATYYCQQWSSNPFTFGQGTKLQIT
|||||||||||||||||||||||.||..|.||.|||||.||||||..||..|.|.||||||||||||||||||||||||||||||.|||||.|.||.|||||.||||.|.
DIQMTQSPSSLSASVGDRVTITCRASENVNSYLNWYQQKPGKAPKLLIYEASTLRSGVPSRFSGSGSGTDYTFTISSLQPEDLATYYCLQYSSSPFTFGPGTKLDIK
                        ^^^^^^          ^^^                       ^^^^^^^^^
```

Best Alignment of Teplizumab heavy chain CDRs to a sequence from OAS
```
QVQLVQSGGGVVQPGRSLRLSCKASGYTFTRYTMHWVRQAPGKGLEWIGYINPSRGYTNYNQKVKDRFTISRDNSKNTAFLQMDSLRPEDTGVYFCARYYDDHYCLDYWGQGTPVTVSS
|.||..|......||.|...|||||||||||||||||.|.||.||||||||||||.|||.||||.||..|...|.|..||..|..||..||..||.||||||..|..||||||||..|..|
QLQLQHSASELARPGASVQMSCKASGYTFTRYTMHWVQQRPGQGLEWIGYINPSSGYTEYNQKLKDKTTLTADKSSSTAYMQLSSLTSEDYAVYYCARYYDYDYAMDYWGQGTTDTDTS
                        ^^^^^^^^          ^^^^^^^^                        ^^^^^^^^^^^^
```

Best Alignment of Teplizumab light CDRs chain to a sequence from OAS
```
DIQMTQSPSSLSASVGDRVTITCSASSSVSYMNWYQQTPGKAPKRWIYDTSKLASGVPSRFSGSGSGTDYTFTISSLQPEDIATYYCQQWSSNPFTFGQGTKLQIT
.|..||||...|||.|..||.||||||||||||.||||..|..||||||||||||||||.||||||||||.|..||||...||.||||||||||||||||||.||||...
QIVLTQSPAIMSASPGEKVTMTCSASSSVSYMHWYQQKSGTSPKRWIYDTSKLASGVPARFSGSGSGTSYSLTISSMEAEDAATYYCQQWSSNPFTFGAGTKLELK
                        ^^^^^          ^^^                       ^^^^^^^^^
```

Best Alignment of Teplizumab CDR-H3 chain to a sequence from OAS
```
QVQLVQSGGGVVQPGRSLRLSCKASGYTFTRYTMHWVRQAPGKGLEWIGYINPS--RGYTNYNQKVKDRFTISRDNSKNTAFLQMDSLRPEDTGVYFCARYYDDHYCLDYWGQGTPVTVSS
.|..|.||||.||||.||.|||.|||.|||.|.|.||||.|||.|||.|.|........|.|...||.||||||||||.....|||..||.||...|.||||.||.||||||||||||..||||
EVKVVESGGGLVQPGGSLSLSCAASGFTFTDYYMSWVRQPPGKALEWLGFIRNKANGYTTEYSASVKGRFTISRDNSQSILYLQMNALRAEDSATYYCARYLDDGYCLDYWGQGTTLTVSS
                        ^^^^^^^^          ^^^^^^^^^^                         ^^^^^^^^^^^^
```

Therapeutic : Teprotumumab

Best Alignment of Teprotumumab heavy chain to a sequence from OAS
```
QVELVESGGGVVQPGRSQRLSCAASGFTFSSYGMHWVRQAPGKGLEWVAIIWFDGSSTYYADSVRGRFTISRDNSKNTLYLQMNSLRAEDTAVYFCARELGR----------RYFDLWGRGTLVSVSS
||.||||||||||||||.|||||||||||||||||||||||||||||||||||.||.|||..||||||.||||||||||||||||||||||||||||||||.||||||............||||||||||||.|||
QVQLVESGGGVVQPGRSLRLSCAASGFTFSSYGMHWVRQAPGKGLEWVAVIWYDGSNKYYADSVKGRFTISRDNSKNTLYLQMNSLRAEDTAVYYCARELGSWGYGSKPGKDWYFDLWGRGTLVTVSS
                        ^^^^^^^^          ^^^^^^^^                        ^^^^^^^^^^^^^^^^^^^^^
```

Best Alignment of Teprotumumab light chain to a sequence from OAS
```
EIVLTQSPATLSLSPGERATLSCRASQSVSSYLAWYQQKPGQAPRLLIYDASKRATGIPARFSGSGSGTDFTLTISSLEPEDFAVYYCQQRSKWPPWTFGQGTKVESK
||||||||||||||||||||||||||||||||||||||||||||||||||||||.|||||||||||||||||||||||||||||||||||||||||||||.||||||||||||||||
EIVLTQSPATLSLSPGERATLSCRASQSVSSYLAWYQQKPGQAPRLLIYDASNRATGIPARFSGSGSGTDFTLTISSLEPEDFAVYYCQQRSNWPPWTFGQGTKVESK
                        ^^^^^^          ^^^                       ^^^^^^^^^^
```

Best Alignment of Teprotumumab heavy chain CDRs to a sequence from OAS
```
QVELVESGGGVVQPGRSQRLSCAASGFTFSSYGMHWVRQAPGKGLEWVAIIWFDGSSTYYADSVRGRFTISRDNSKNTLYLQMNSLRAEDTAVYFCARELGRRYFDLWGRGTLVSVSS
................|.|||||||||||||||||||||||||||||||||.||||||||.|||||||.||||||||||||.||||||||.||||.|||||.|.||||.||..||||||||||||.|||
```

```
----------------SLRLSCAASGFTFSSYGMHWVRQAPGKGLEWVAAIWFDGSSKYYADSVKGRFTISRDNSKSTLYLQMSSLRVEDTAIYYCARGLGHWYFDLWGRGTLVTVSS
                          ^^^^^^^^                ^^^^^^^^                                ^^^^^^^^^^^

Best Alignment of Teprotumumab light CDRs chain to a sequence from OAS
EIVLTQSPATLSLSPGERATLSCRASQSVSSYLAWYQQKPGQAPRLLIYDASKRATGIPARFSGSGSGTDFTLTISSLEPEDFAVYYCQQRSKWPPWTFGQGTKVESK
||||||||||||||||||||||||||||||||||||||||||||||||||||.|||||||||||||||||||.|||||||||||||||||||||||||||||||||.|
EIVLTQSPATLSLSPGERATLSCRASQSVSSYLAWYQQKPGQAPRLLIYDASNRATGIPARFSGSGSGTDFTHTISSLEPEDFAVYYCQQRSKWPPWTFGQGTKVEIK
                           ^^^^^^                ^^^                             ^^^^^^^^^^

Best Alignment of Teprotumumab CDR-H3 chain to a sequence from OAS
QVELVESGGGVVQPGRSQRLSCAASGFTFSSYGMHWVRQAPGKGLEWVAIIWFDGSSTYYADSVRGRFTISRDNSKNTLYLQMNSLRAEDTAVYFCARELGRRYFDLWGRGTLVSVSS
..............|.|.|||||.|||||....||||||||||||||||.|||||...|||||..|||||||||||||||||||.||..|||||.|||.|||||||||||||||.|||
--------------GGSLRLSCAPSGFTFRNSAMHWVRQAPGKGLEWVALIWFDGTKEYYADSLKGRFTISRDNSKNTLYLQMNRLRVDDTAVYYCARDLGRRYFDLWGRGTLVTVSS
                          ^^^^^^^^                ^^^^^^^^                                ^^^^^^^^^^^

Therapeutic : Tesidolumab

Best Alignment of Tesidolumab heavy chain to a sequence from OAS
EVQLVQSGAEVKKPGSSVKVSCKASGGTFSSYAISWVRQAPGQGLEWMGGIGPFFGTANYAQKFQGRVTITADESTSTAYMELSSLRSEDTAVYYCARDTP-------------YFDYWGQGTLVTVSS
.||||||||||||||||||||||||||||||||||||||||||||||||||.|.|||||||||||||||||||||||||||||||||||||||||||||||.............|||||||||||||||
QVQLVQSGAEVKKPGSSVKVSCKASGGTFSSYAISWVRQAPGQGLEWMGGIIPIFGTANYAQKFQGRVTITADESTSTAYMELSSLRSEDTAVYYCARDTPRDYHDSSGYYSPGYFDYWGQGTLVTVSS
                          ^^^^^^^^                ^^^^^^^^                                ^^^^^^^^^^^^^^^^^^^^^

Best Alignment of Tesidolumab light chain to a sequence from OAS
SYELTQPLSVSVALGQTARITCSGDSIPNYYVYWYQQKPGQAPVLVIYDDSNRPSGIPERFSGSNSGNTATLTISRAQAGDEADYYCQSFDSSLNAEVFGGGTKLTVL
||||||||||||||||||||||.|..|....|.|||||||||||||||.|||||||||||||||||||||||||||||||||||||||..||...|||||||||||||
SYELTQPLSVSVALGQTARITCGGNNIGSKNVHWYQQKPGQAPVLVIYRDSNRPSGIPERFSGSNSGNTATLTISRAQAGDEADYYCQVWDS-STAEVFGGGTKLTVL
                          ^^^^^^             ^^^                              ^^^^^^^^^^^

Best Alignment of Tesidolumab heavy chain CDRs to a sequence from OAS
EVQLVQSGAEVKKPGSSVKVSCKASGGTFSSYAISWVRQAPGQGLEWMGGIGPFFGTANYAQKFQGRVTITADESTSTAYMELSSLRSEDTAVYYCARDTPYFDYWGQGTLVTVSS
................|||||||||||||||||||||||||||||||||||.|.||||||||||||||||||||||||||||||||||||||||||||||||||||||||||||||
----------------SVKVSCKASGGTFSSYAISWVRQAPGQGLEWMGGIIPIFGTANYAQKFQGRVTITADESTSTAYMELSSLRSEDTAVYYCARDTPYFDYWGQGTLVTVSS
                          ^^^^^^^^                ^^^^^^^^                                ^^^^^^^^^

Best Alignment of Tesidolumab light CDRs chain to a sequence from OAS
SYELTQPLSVSVALGQTARITCSGDSIPNYYVYWYQQKPGQAPVLVIYDDSNRPSGIPERFSGSNSGNTATLTISRAQAGDEADYYCQSFDSSLNAEVFGGGTKLTVL
|.|.||...||||||||.||||.|||..|||..|||||||||||.|||.||.||||.|.|||||.||..|.|.|...||.|||.|||||.||||...||||||||||.
SSEVTQDPAVSVALGQTVRITCQGDSLRNYYANWYQQKPGQAPVVVIYYDSDRPSGVPDRFSGSKSGTSASLAITGLQAEDEAAYYCQSYDSSLSGSVFGGGTKLTVV
                          ^^^^^^             ^^^                              ^^^^^^^^^^^

Best Alignment of Tesidolumab CDR-H3 chain to a sequence from OAS
EVQLVQSGAEVKKPGSSVKVSCKASGGTFSSYAISWVRQAPGQGLEWMGGIGPFFGTANYAQKFQGRVTITADESTSTAYMELSSLRSEDTAVYYCARDTPYFDYWGQGTLVTVSS
................|...||.|||.|||||...|||||||.||||...|........||....||.||..|.|..|.|....|||.||||||||||||||||||||||||||||
----------------SLRLSCAASGFTFSSYGMHWVRQAPGKGLEWVAVIWYDGSNKYYADSVKGRFTISRDNSKNTLYLQMNSLRAEDTAVYYCARDTPYFDYWGQGTLVTVSS
                          ^^^^^^^^                ^^^^^^^^                                ^^^^^^^^^

Therapeutic : Tezepelumab

Best Alignment of Tezepelumab heavy chain to a sequence from OAS
QMQLVESGGGVVQPGRSLRLSCAASGFTFRTYGMHWVRQAPGKGLEWVAVIWYDGSNKHYADSVKGRFTITRDNSKNTLNLQMNSLRAEDTAVYYCARAPQWELVHEAFDIWGQGTMVTVSS
|.|||||||||||||||||||||||||||..|||||||||||||||||||||||||||.|||||||||||.||||||||.||||||||||||||||||..|..||..|||||||||||||||
QVQLVESGGGVVQPGRSLRLSCAASGFTFSSYGMHWVRQAPGKGLEWVAVIWYDGSNKYYADSVKGRFTISRDNSKNTLYLQMNSLRAEDTAVYYCARDSQQWLVRSAFDIWGQGTMVTVSS
                          ^^^^^^^^                ^^^^^^^^                                ^^^^^^^^^^^^^^^

Best Alignment of Tezepelumab light chain to a sequence from OAS
SYVLTQPPSVSVAPGQTARITCGGNNLGSKSVHWYQQKPGQAPVLVVYDDSDRPSWIPERFSGSNSGNTATLTISRGEAGDEADYYCQVWDSSSDHVVFGGGTKLTVL
||||||||||||||||||||||||||.||||||||||||||||||||||||||||.||||||||||||||||||||||||||||||||||||||||||||||||||||
SYVLTQPPSVSVAPGQTARITCGGNNIGSKSVHWYQQKPGQAPVLVVYDDSDRPSGIPERFSGSNSGNTATLTISRGEAGDEADYYCQVWDSSSDHVVFGGGTKLTVL
                          ^^^^^^             ^^^                              ^^^^^^^^^^^

Best Alignment of Tezepelumab heavy chain CDRs to a sequence from OAS
QMQLVESGGGVVQPGRSLRLSCAASGFTFRTYGMHWVRQAPGKGLEWVAVIWYDGSNKHYADSVKGRFTITRDNSKNTLNLQMNSLRAEDTAVYYCARAPQWELVHEAFDIWGQGTMVTVSS
................|||||||||||||.|||.||||||||||||||||||||||||.|||||||||||.||||||||.||||||||||||||||||...|||.|.|||||||||||||||
----------------SLRLSCAASGFTFSTYGIHWVRQAPGKGLEWVAVIWYDGSNKYYADSVKGRFTISRDNSKNTLYLQMNSLRAEDTAVYYCARGGWWELPHDAFDIWGQGTMVTVSS
                          ^^^^^^^^                ^^^^^^^^                                ^^^^^^^^^^^^^^^

Best Alignment of Tezepelumab light CDRs chain to a sequence from OAS
SYVLTQPPSVSVAPGQTARITCGGNNLGSKSVHWYQQKPGQAPVLVVYDDSDRPSWIPERFSGSNSGNTATLTISRGEAGDEADYYCQVWDSSSDHVVFGGGTKLTVL
|||||||||||.||||||||||.|.||||||||||||||||||||||||||||||.||||||||||||||||||||.|.|||||||||||||||||||||||||||||
SYVLTQPPSVSLAPGQTARITCEGDNLGSKSVHWYQQKPGQAPVLVVYDDSDRPSGIPERFSGSNSGNTATLTISRVEGGDEADYYCQVWDSSSDHVVFGGGTKLTVL
                          ^^^^^^             ^^^                              ^^^^^^^^^^^

Best Alignment of Tezepelumab CDR-H3 chain to a sequence from OAS
QMQLVESGGGVVQPGRSLRLSCAASGFTFRTYGMHWVRQAPGKGLEWVAVIWYDGSNKHYADSVKGRFTITRDNSKNTLNLQMNSLRAEDTAVYYCARAPQWELVHEAFDIWGQGTMVTVSS
|.||.|||.|.|.|...|.|.|..||.....|...|.||.|||||||...|.|......|..|.|.|.||..|.|||...|...||.|.|||||||||||||||...|||||||||||||.||
QVQLQESGPGLVKPSETLSLTCTVSGGSISSYYWSWIRQPPGKGLEWIGYIYYS-GGTNYNPSLKSRVTISVDTSKNQFSLKLSSLTAADTAVYYCARAPQWELLPDAFDIWGQGTMVTASS
                          ^^^^^^^^                ^^^^^^^^                                ^^^^^^^^^^^^^^^

Therapeutic : Tigatuzumab

Best Alignment of Tigatuzumab heavy chain to a sequence from OAS
EVQLVESGGGLVQPGGSLRLSCAASGFTFSSYVMSWVRQAPGKGLEWVATISSGGSYTYYPDSVKGRFTISRDNAKNTLYLQMNSLRAEDTAVYYCARRGDSMITTDYWGQGTLVTVSS
||||||||||||||||||||||||||||||||.||||||||||||||||.|...||..||.||||||||||||||||.||||||||||||||||||||||.|...||||||||||||||
EVQLVESGGGLVQPGGSLRLSCAASGFTFSSYWMSWVRQAPGKGLEWVANIKQDGSEKYYVDSVKGRFTISRDNAKNSLYLQMNSLRAEDTAVYYCARRGYS-SHTDYWGQGTLVTVSS
                          ^^^^^^^^                ^^^^^^^^                                ^^^^^^^^^^^^

Best Alignment of Tigatuzumab light chain to a sequence from OAS
DIQMTQSPSSLSASVGDRVTITCKASQDVGTAVAWYQQKPGKAPKLLIYWASTRHTGVPSRFSGSGSGTDFTLTISSLQPEDFATYYCQQYS-SYRTFGQGTKVEIK
|||||||||||||||||||||||.||||.||..||||||||||||||||.|||.|.|||.||||||||||||||||||||||||||||||.....||||||||||||
DIQMTQSPSSLSASVGDRVTITCRASQDSGTWLAWYQQKPGKAPKLLIYKASTLHSGVPARFSGSGSGTDFTLTISSLQPEDFATYYCQQHHTTPRTFGQGTKVEIK
                           ^^^^^^             ^^^                             ^^^^^^^^^

Best Alignment of Tigatuzumab heavy chain CDRs to a sequence from OAS
EVQLVESGGGLVQPGGSLRLSCAASGFTFSSYVMSWVRQAPGKGLEWVATISSGGSYTYYPDSVKGRFTISRDNAKNTLYLQMNSLRAEDTAVYYCARRGDSMITTDYWGQGTLVTVSS
|||.|||||.||.|||||.|||||||||||||.||||||.|...||||||||||||||||||||.||||||||||||||||||.||..||||...||||||.|||.||||.||..||.|
EVQGVESGGDLVKPGGSLKLSCAASGFTFSSYGMSWVRQTPDQRLEWVATISSGGSYTYYPDSVQGRFTISRDNAKNTLYLQMSSLKSEDTAMDGCARRGDPMITVDYWGEGTTLTVAS
                          ^^^^^^^^                ^^^^^^^^                                ^^^^^^^^^^^^

Best Alignment of Tigatuzumab light CDRs chain to a sequence from OAS
DIQMTQSPSSLSASVGDRVTITCKASQDVGTAVAWYQQKPGKAPKLLIYWASTRHTGVPSRFSGSGSGTDFTLTISSLQPEDFATYYCQQYSSYRTFGQGTKVEIK
||.||||....|.|||||..|||||||||||||||.|||||..||||||||||||||||.||.|||||||||||.|..|.||.|.|.|||||||||||.|||.|.|
DIVMTQSHTFMSTSVGDRGSITCKASQDVGTAVAWHQQKPGQSPKLLIYWASTRHTGVPDRFTGSGSGTDFTLTSSNVQSEDLADYFCQQYSSYRTFGAGTKLELK
                           ^^^^^^             ^^^                             ^^^^^^^^
Best Alignment of Tigatuzumab CDR-H3 chain to a sequence from OAS
EVQLVESGGGLVQPGGSLRLSCAASGFTFSSYVMSWVRQAPGKGLEWVATISSGGSYTYYPDSVKGRFTISRDNAKNTLYLQMNSLRAEDTAVYYCARRGDSMITTDYWGQGTLVTVSS
|||.|||||.||.|||||.|||||||||||||.||||||.|...||||||||||||||||||||.||||||||||||||||||.||..||||...||||||.|||.||||.||..||.|
```

EVQGVESGGDLVKPGGSLKLSCAASGFTFSSYGMSWVRQTPDQRLEWVATISSGGSYTYYPDSVQGRFTISRDNAKNTLYLQMSSLKSEDTAMDGCARRGDPMITVDYWGEGTTLTVAS
          ^^^^^^^^          ^^^^^^^^                    ^^^^^^^^^^^^

Therapeutic : Tildrakizumab

Best Alignment of Tildrakizumab heavy chain to a sequence from OAS
QVQLVQSGAEVKKPGASVKVSCKASGYIFITYWMTWVRQAPGQGLEWMGQIFPASGSADYNEKFEGRVTMTTDTSTSTAYMELRSLRSDDTAVYYCARGGG-----GFAYWGQGTLVTVSS
|||||||||||||||||||||||||||.|..|.|.||||||||||||||.|.|..||..|..||.||||||||||||||||||||||||||||||||||||......|.|||||||||||||
QVQLVQSGAEVKKPGASVKVSCKASGYTFTSYYMHWVRQAPGQGLEWMGIINPSGGSTSYAQKFQGRVTMTTDTSTSTAYMELRSLRSDDTAVYYCARGGGSYGQDYFDYWGQGTLVTVSS
          ^^^^^^^^          ^^^^^^^^                    ^^^^^^^^^^^^^^

Best Alignment of Tildrakizumab light chain to a sequence from OAS
DIQMTQSPSSLSASVGDRVTITCRTSENIYSYLAWYQQKPGKAPKLLIYNAKTLAEGVPSRFSGSGSGTDFTLTISSLQPEDFATYYCQHHYGIPFTFGQGTKVEIK
||||||||||||||||||||||||.|.||.||||||||||||.||||||.|.||..|||||||||||||||||||||||||||||||||.|...|.|||||||||||
DIQMTQSPSSLSASVGDRVTITCRASQNISSYLAWYQQKPGKVPKLLIYAASTLQSGVPSRFSGSGSGTDFTLTISSLQPEDFATYYCQQHNSHPRTFGQGTKVEIK
           ^^^^^^          ^^^                    ^^^^^^^^^

Best Alignment of Tildrakizumab heavy chain CDRs to a sequence from OAS
QVQLVQSGAEVKKPGASVKVSCKASGYIFITYWMTWVRQAPGQGLEWMGQIFPASGSADYNEKFEGRVTMTTDTSTSTAYMELRSLRSDDTAVYYCARGGGGFAYWGQGTLVTVSS
||||.|||.|...||||||.||||||||||||||.||.|.|||||||.|||||||||..|||.|||..|.|.|||.|||||.|.||.|.|.|.||||||||||||.||||||..||||
QVQLQQSGPELVRPGASVKLSCKASGYIFITYWMNWVKQRPGQGLEWIGQIFPASGSTNYNEMFEGKATLTVDTSSSTAYMQLSSLTSEDSAVYYCARGGGGFDYWGQGTTLTVSS
          ^^^^^^^^          ^^^^^^^^                    ^^^^^^^^^

Best Alignment of Tildrakizumab light CDRs chain to a sequence from OAS
DIQMTQSPSSLSASVGDRVTITCRTSENIYSYLAWYQQKPGKAPKLLIYNAKTLAEGVPSRFSGSGSGTDFTLTISSLQPEDFATYYCQHHYGIPFTFGQGTKVEIK
||||||||.||||.||..||||||.||||||||.|||||.||.|..|.|||||||||||||||||||||.|.|.|.|||||||..||||||||||||||.|||.|||
DIQMTQSPASLSAFVGETVTITCRASENIYSYLVWYQQKQGKSPPFLVYNAKTLAEGVPSRFSGSGSGTQFSLKINSLQPEDFGSYYCQHHYGIPFTFGSGTKLEIK
           ^^^^^^          ^^^                    ^^^^^^^^^

Best Alignment of Tildrakizumab CDR-H3 chain to a sequence from OAS
QVQLVQSGAEVKKPGASVKVSCKASGYIFITYWMTWVRQAPGQGLEWMGQIFPASGSADYNEKFEGRVTMTTDTSTSTAYMELRSLRSDDTAVYYCARGGGGFAYWGQGTLVTVSS
.....|||.|..|||||||.|||||||.|..|.|.||.|..|..|||.|.|.|..|...||.||.|..|.|.|.|.||||||||||.|.|.||||.|||||||||||||||||||.
----QQSGPELVKPGASVKMSCKASGYTFTDYYMHWVKQSHGKSLEWIGYIYPNNGGNGYNQKFKGKATLTVDKSSSTAYMELRSLTSEDSAVYYWARGGGGFAYWGQGTLVTVSA
          ^^^^^^^^          ^^^^^^^^                    ^^^^^^^^^

Therapeutic : Timolumab

Best Alignment of Timolumab heavy chain to a sequence from OAS
QVQLVESGGGVVQPGRSLRLSCAASGFTFFSYAMHWVRQTPGKGLEWVAVIWFDGSNENYVDSVKGRFTISRDNSKNTLYLQMNTLRAEDTAVYYCARDAW-----SYFDYWGQGTLVTVSS
|||||||||||||||||||||||||||||.|||||||||.||||||||||||.||||..|.|||||||||||||||||||||||.||||||||||||.|.|.....||||||||||||||||
QVQLVESGGGVVQPGRSLRLSCAASGFTFSSYAMHWVRQAPGKGLEWVAVIWYDGSNKYYADSVKGRFTISRDNSKNTLYLQMNSLRAEDTAVYYCAKDLWSGSYYSYFDYWGQGTLVTVSS
          ^^^^^^^^          ^^^^^^^^                    ^^^^^^^^^^^^^^^

Best Alignment of Timolumab light chain to a sequence from OAS
VIQLTQSPSSLSASVGDRVTITCRASQGISRALAWYQQKPGKGPKLLIYDASSLESGVPSRFSGSGSGTDFTLTISSLQPEDFATYYCQQFNSYPLTFGGGTKVEIK
.|||||||||||||||||||||||||||||.|||||||||||.||||||||||||||||||||||||||||||||||||||||||||||||||||||||||||||||
AIQLTQSPSSLSASVGDRVTITCRASQGISSALAWYQQKPGKAPKLLIYDASSLESGVPSRFSGSGSGTDFTLTISSLQPEDFATYYCQQFNSYPLTFGGGTKVEIK
           ^^^^^^          ^^^                    ^^^^^^^^^

Best Alignment of Timolumab heavy chain CDRs to a sequence from OAS
QVQLVESGGGVVQPGRSLRLSCAASGFTFFSYAMHWVRQTPGKGLEWVAVIWFDGSNENYVDSVKGRFTISRDNSKNTLYLQMNTLRAEDTAVYYCARDAWSYFDYWGQGTLVTVSS
..............|.||..|||||||||.||.||||||.|||||||||.||||||||.|.|||||||||||||.|||||||||.||||||||||||||...|||||||||||||||
--------------GESLKISCAASGFTFTSYGMHWVRQAPGKGLEWVAFIWFDGSNEFYADSVKGRFTISRDNAKNTLYLQMNSLRAEDTAVYYCARDKFYYFDYWGQGTLVTVSS
          ^^^^^^^^          ^^^^^^^^                    ^^^^^^^^^^

Best Alignment of Timolumab light CDRs chain to a sequence from OAS
VIQLTQSPSSLSASVGDRVTITCRASQGISRALAWYQQKPGKGPKLLIYDASSLESGVPSRFSGSGSGTDFTLTISSLQPEDFATYYCQQFNSYPLTFGGGTKVEIK
.|||||||||||||||||||||||||||||||||||||||||.|||||||||||.||||||||||||||||||||.||||||||.|||||||||||||.||||||||
AIQLTQSPSSLSASVGDRVTITCRASQGISRALAWYQQKPGKLPKLLIYDASSLQSGVPSRFSGSGSGTDFTLTINSLQPEDFAAYYCQQFNSYPLTFAGGTKVEIK
           ^^^^^^          ^^^                    ^^^^^^^^^

Best Alignment of Timolumab CDR-H3 chain to a sequence from OAS
QVQLVESGGGVVQPGRSLRLSCAASGFTFFSYAMHWVRQTPGKGLEWVAVIWFDGSNENYVDSVKGRFTISRDNSKNTLYLQMNTLRAEDTAVYYCARDAWSYFDYWGQGTLVTVSS
.................|.|.||..|..|..|...|.||..||||||...|.......||..|.|.|.|||.|.|||...|......|.||||||||||||||||||||||||||||
-------------TSETLSLTCAVYGGSFSGYYWSWIRQPAGKGLEWIGRIYTS-GSTNYNPSLKSRVTISVDTSKNQFSLKLSSVTAADTAVYYCARDAWSYFDYWGQGTLVTVSS
          ^^^^^^^^          ^^^^^^^^                    ^^^^^^^^^^

Therapeutic : Tislelizumab

Best Alignment of Tislelizumab heavy chain to a sequence from OAS
QVQLQESGPGLVKPSETLSLTCTVSGFSLTSYGVHWIRQPPGKGLEWIGVIYADGSTNYNPSLKSRVTISKDTSKNQVSLKLSSVTAADTAVYYCARAYGN----YWYIDVWGQGTTVTVSS
||||||||||||||||||||||||||.|..||...||||||||||||||.||..||||||||||||||||.||||||.||||||||||||||||||||.|.....|.|||||||||||||||
QVQLQESGPGLVKPSETLSLTCTVSGGSISSYYWSWIRQPPGKGLEWIGYIYYSGSTNYNPSLKSRVTISVDTSKNQFSLKLSSVTAADTAVYYCARAGGYGSWYYYYIDVWGQGTTVTVSS
          ^^^^^^^^          ^^^^^^^                    ^^^^^^^^^^^^^^^^

Best Alignment of Tislelizumab light chain to a sequence from OAS
DIVMTQSPDSLAVSLGERATINCKSSESV-----SNDVAWYQQKPGQPPKLLINYAFHRFTGVPDRFSGSGYGTDFTLTISSLQAEDVAVYYCHQAYSSPYTFGQGTKLEIK
||||||||||||||||||||||||||.||......|.||||||||||||||||..|..|..||||||||||.|||||||||||||||||||||.|.||||||||||||||||
DIVMTQSPDSLAVSLGERATINCKSSQSVLYSNNKNYVAWYQQKPGQPPKLLIYWASTRESGVPDRFSGSGSGTDFTLTISSLQAEDVAVYYCQQYYSSPYTFGQGTKLEIK
           ^^^^^^^^^^^          ^^^                    ^^^^^^^^^

Best Alignment of Tislelizumab heavy chain CDRs to a sequence from OAS
QVQLQESGPGLVKPSETLSLTCTVSGFSLTSYGVHWIRQPPGKGLEWIGVIYADGSTNYNPSLKSRVTISKDTSKNQVSLKLSSVTAADTAVYYCARAYGNYWYIDVWGQGTTVTVSS
.||||.||||||.||..||.||||||||||||||||.||||||||||.|||.|.||||||..|.||..||||.||.||.||..|....|||.||||||||||||.||||.||||||||
-VQLQQSGPGLVAPSQSLSITCTVSGFSLTSYGVHWVRQPPGKGLEWLGVIWAGGSTNYNSALMSRLSISKDNSKSQVFLKMNSLQTDDTAMYYCARAYGNYWYFDVWGAGTTVTVSS
          ^^^^^^^^          ^^^^^^^                    ^^^^^^^^^^^^

Best Alignment of Tislelizumab light CDRs chain to a sequence from OAS
DIVMTQSPDSLAVSLGERATINCKSSESVSNDVAWYQQKPGQPPKLLINYAFHRFTGVPDRFSGSGYGTDFTLTISSLQAEDVAVYYCHQAYSSPYTFGQGTKLEIK
.|||||.|..|.||.|.|.||.||.|.||||||||||||||..|||||.||..|.|||||||.|||||||||.|||..||||.|||.|||.||||||||.|||||||
-IVMTQTPTFLLVSAGDRVTITCKASQSVSNDVAWYQQKPGKSPKLLIYYASNRYTGVPDRFTGSGYGTDFTFTISTVQAEDLAVYFCHQDYSSPYTFGGGTKLEIK
           ^^^^^^          ^^^                    ^^^^^^^^^

Best Alignment of Tislelizumab CDR-H3 chain to a sequence from OAS
QVQLQESGPGLVKPSETLSLTCTVSGFSLTSYGVHWIRQPPGKGLEWIGVIYAD-GSTNYNPSLKSRVTISKDTSKNQVSLKLSSVTAADTAVYYCARAYGNYWYIDVWGQGTTVTVSS
.|.|.|||..||||...|.|.|..|||...|||..|.||.|.|.|||...|......|.|..|.|.|.|||.|.|..||...|..||....|||.||||||||||||.||||.||||||||
EVNLVESGGDLVKPGGSLKLSCAASGFTFSSYGMSWVRQTPDKRLEWVATISSGGSYTYYPDSVKGRFTISRDNAKNTLYLQMSSLKSEDTAMYYCARAYGNYWYFDVWGAGTTVTVSS
          ^^^^^^^^          ^^^^^^^^                    ^^^^^^^^^^^^

Therapeutic : Tisotumab

Best Alignment of Tisotumab heavy chain to a sequence from OAS
EVQLLESGGGLVQPGGSLRLSCAASGFTFSNYAMSWVRQAPGKGLEWVSSISGSGDYTYYTDSVKGRFTISRDNSKNTLYLQMNSLRAEDTAVYYCARSPWG-YYLDSWGQGTLVTVSS
||||||||||||||||||||||||||||||.||||||||||||||||||.|||||..|||.||||||||||||||||||||||||||||||||||||||||..||.|.|||||||||||
EVQLLESGGGLVQPGGSLRLSCAASGFTFSSYAMSWVRQAPGKGLEWVSAISGSGGSTYYADSVKGRFTISRDNSKNTLYLQMNSLRAEDTAVYYCARSPWAPYYFDYWGQGTLVTVSS
          ^^^^^^^^          ^^^^^^^^                    ^^^^^^^^^^^^

Best Alignment of Tisotumab light chain to a sequence from OAS
DIQMTQSPPSLSASAGDRVTITCRASQGISSRLAWYQQKPEKAPKSLIYAASSLQSGVPSRFSGSGSGTDFTLTISSLQPEDFATYYCQQYNSYPYTFGQGTKLEIK
||||||||.|||||.||||||||||||||||.|||||||||||||||||||||||||||||||||||||||||||||||||||||||||||||||||||||||||||
DIQMTQSPSSLSASVGDRVTITCRASQGISSWLAWYQQKPEKAPKSLIYAASSLQSGVPSRFSGSGSGTDFTLTISSLQPEDFATYYCQQYNSYPYTFGQGTKLEIK
^^^^^^ ^^^ ^^^^^^^^^

Best Alignment of Tisotumab heavy chain CDRs to a sequence from OAS
EVQLLESGGGLVQPGGSLRLSCAASGFTFSNYAMSWVRQAPGKGLEWVSSISGSGDYTYYTDSVKGRFTISRDNSKNTLYLQMNSLRAEDTAVYYCARSPWGYYLDSWGQGTLVTVSS
.............|||||||||||||||||.|||.||||||||||||||.||||||.|||.||||||||||||.|||||||||.||||||||.||||||.|.||.|||||||||||||
-------------PGGSLRLSCAASGFTFSSYAMNWVRQAPGKGLEWVSFISGSGDTTYYADSVKGRFTISRDSSKNTLYLQMDSLRAEDTAAYYCARSSWPYYFDSWGQGTLVTVSS
^^^^^^^^ ^^^^^^^^ ^^^^^^^^^^^

Best Alignment of Tisotumab light CDRs chain to a sequence from OAS
DIQMTQSPPSLSASAGDRVTITCRASQGISSRLAWYQQKPEKAPKSLIYAASSLQSGVPSRFSGSGSGTDFTLTISSLQPEDFATYYCQQYNSYPYTFGQGTKLEIK
||||||||.|||||.||||||||||||||||||||||||||.|||||||||||.|.|||||||||||||||||||||||||||||||||||||||||||||||||||
DIQMTQSPSSLSASVGDRVTITCRASQGISSRLAWYQQKPEQAPKSLIYAASSWQRGVPSRFSGSGSGTDFTLTISSLQPEDFATYYCQQYNSYPYTFGQGTKLEIK
^^^^^^ ^^^ ^^^^^^^^^

Best Alignment of Tisotumab CDR-H3 chain to a sequence from OAS
EVQLLESGGGLVQPGGSLRLSCAASGFTFSNYAMSWVRQAPGKGLEWVSSISGSGDYTYYTDSVKGRFTISRDNSKNTLYLQMNSLRAEDTAVYYCARSPWGYYLDSWGQGTLVTVSS
.................|.|.|..||...|||...|.||.|||||||...||.|...|.|..|.|.|.|.|.|||...|...|..|.|||||.|||||.|||||.|||||||||||
--------------SETLSLTCTVSGGSVSNYHWGWLRQSPGKGLEWIGDISYS-RITNYNPSLKSRVTMSVDTSKNQFSLRLSSVTAADTAVYFCARSPKGYYLDYWGQGTLVTVSS
^^^^^^^^ ^^^^^^^^ ^^^^^^^^^^^

Therapeutic : Tocilizumab

Best Alignment of Tocilizumab heavy chain to a sequence from OAS
QVQLQESGPGLVRPSQTLSLTCTVSGYSIT-SDHAWSWVRQPPGRGLEWIGYISYSGITTYNPSLKSRVTMLRDTSKNQFSLRLSSVTAADTAVYYCARSLAR-------TTAMDYWGQGSLVTVSS
||||||||||||.||||||||||||||.||...|..|||.||||.||||||||.|||.|.||||||||||...|||||||||.||||||||||||||||||||..........|||||||.||||||
QVQLQESGPGLVKPSQTLSLTCTVSGGSISSGDYYWSWIRQPPGKGLEWIGYIYYSGSTYYNPSLKSRVTISVDTSKNQFSLKLSSVTAADTAVYYCARSLARYYYDSSGYWGMDYWGQGTLVTVSS
^^^^^^^^^^ ^^^^^^^ ^^^^^^^^^^^^^^^^^^^

Best Alignment of Tocilizumab light chain to a sequence from OAS
DIQMTQSPSSLSASVGDRVTITCRASQDISSYLNWYQQKPGKAPKLLIYYTSRLHSGVPSRFSGSGSGTDFTFTISSLQPEDIATYYCQQGNTLPYTFGQGTKVEIK
|||||||||||||||||||||||||||.|||||||||||||||||||||..|.|.|||||||||||||||||||||||||||||||||||...||||||||||||||
DIQMTQSPSSLSASVGDRVTITCRASQSISSYLNWYQQKPGKAPKLLIYAASSLQSGVPSRFSGSGSGTDFTFTISSLQPEDIATYYCQQYDNLPYTFGQGTKVEIK
^^^^^^ ^^^ ^^^^^^^^^

Best Alignment of Tocilizumab heavy chain CDRs to a sequence from OAS
QVQLQESGPGLVRPSQTLSLTCTVSGYSITSDHAWSWVRQPPGRGLEWIGYISYSGITTYNPSLKSRVTMLRDTSKNQFSLRLSSVTAADTAVYYCARSLARTTAMDYWGQGSLVTVSS
.|||||||||||.|||.|||||||.|||||||.||.|.||.||..|||.|||||||.|.||||||||....||||||||.|.|.|.|..|||.|||||||.|..||||||||..|||||
DVQLQESGPGLVKPSQSLSLTCTVTGYSITSDYAWNWIRQFPGNKLEWMGYISYSGSTSYNPSLKSRISITRDTSKNQFFLQLNSATTEDTATYYCARSLLRLLAMDYWGQGTSVTVSS
^^^^^^^^^ ^^^^^^^ ^^^^^^^^^^^^

Best Alignment of Tocilizumab light CDRs chain to a sequence from OAS
DIQMTQSPSSLSASVGDRVTITCRASQDISSYLNWYQQKPGKAPKLLIYYTSRLHSGVPSRFSGSGSGTDFTFTISSLQPEDIATYYCQQGNTLPYTFGQGTKVEIK
||||||..||||||.||||||.||||||||||||||||||....||||||||||||||||||||||||||...|||.|..||||||.||||||||||||.|||.|||
DIQMTQTTSSLSASLGDRVTISCRASQDISSYLNWYQQKPDGTVKLLIYYTSRLHSGVPSRFSGSGSGTDYSLTISNLEQEDIATYFCQQGNTLPYTFGGGTKLEIK
^^^^^^ ^^^ ^^^^^^^^^

Best Alignment of Tocilizumab CDR-H3 chain to a sequence from OAS
QVQLQESGPGLVRPSQTLSLTCTVSGYSITSDHAWSWVRQPPGRGLEWIGYISYS-GITTYNPSLKSRVTMLRDTSKNQFSLRLSSVTAADTAVYYCARSLARTTAMDYWGQGSLVTVSS
||||..||..||||.....|.|..|||....|....||.|.||.|||||..|......|.||...|...|.....|.......|.|.|..|.|||.||||||.|.||||.|||..|||||
QVQLKQSGAELVRPGASVKLSCKASGYTF-TDYYINWVKQRPGQGLEWIARIYPGRCNTYYNEKVKGKATLTAEKSSITAYMQLRSLTSEDSAVYCCARSLAGTRAMDYGGQGTSVTVSS
^^^^^^^^^ ^^^^^^^^ ^^^^^^^^^^^^

Therapeutic : Tomuzotuximab

Best Alignment of Tomuzotuximab heavy chain to a sequence from OAS
QVQLKQSGPGLVQPSQSLSITCTVSGFSLTNYGVHWVRQSPGKGLEWLGVIWSGGNTDYNTPFTSRLSINKDNSKSQVFFKMNSLQSNDTAIYYCARALTYYDYEFAYWGQGTLVTVST
||||||||||||||||||||||||||||||.||||||||||||||||||||||||.||||..|.|||||.||||||||||||||||.||||||||||.|.|.||||||||||||||||.
QVQLKQSGPGLVQPSQSLSITCTVSGFSLTSYGVHWVRQSPGKGLEWLGVIWSGGSTDYNAAFISRLSISKDNSKSQVFFKMNSLQANDTAIYYCARGLYY-DYEFAYWGQGTLVTVSA
^^^^^^^^ ^^^^^^^ ^^^^^^^^^^^^^

Best Alignment of Tomuzotuximab light chain to a sequence from OAS
DILLTQSPVILSVSPGERVSFSCRASQSIGTNIHWYQQRTNGSPRLLIKYASESISGIPSRFSGSGSGTDFTLSINSVESEDIADYYCQQNNNWPTTFGAGTKLELK
||||||||.||||||||||||||||||||||.|||||||||||||||||||||||||||||||||||||||||||||||||||||||||||||||.|||||||||||
DILLTQSPAILSVSPGERVSFSCRASQSIGTSIHWYQQRTNGSPRLLIKYASESISGIPSRFSGSGSGTDFTLSINSVESEDIADYYCQQNNNWPLTFGAGTKLELK
^^^^^^ ^^^ ^^^^^^^^^

Best Alignment of Tomuzotuximab heavy chain CDRs to a sequence from OAS
QVQLKQSGPGLVQPSQSLSITCTVSGFSLTNYGVHWVRQSPGKGLEWLGVIWSGGNTDYNTPFTSRLSINKDNSKSQVFFKMNSLQSNDTAIYYCARALTYYDYEFAYWGQGTLVTVST
.|.|..||||||||||||||||||||||||.||||||||||||||||||||||||.||||..|.|||||.||||||||||||||||..|||||||||.|.||||.|||||||||||||.
-VMLVESGPGLVQPSQSLSITCTVSGFSLTSYGVHWVRQSPGKGLEWLGVIWSGGSTDYNAAFISRLSISKDNSKSQVFFKMNSLQADDTAIYYCARNLDYYDYGFAYWGQGTLVTVSA
^^^^^^^^ ^^^^^^^ ^^^^^^^^^^^^^

Best Alignment of Tomuzotuximab light CDRs chain to a sequence from OAS
DILLTQSPVILSVSPGERVSFSCRASQSIGTNIHWYQQRTNGSPRLLIKYASESISGIPSRFSGSGSGTDFTLSINSVESEDIADYYCQQNNNWPTTFGAGTKLELK
||||||||.||||||||||||||||||||||.|||||||||||||||||||||||||||||||||||||.|||.|||||||||||||||||||||||||.|||||.|
DILLTQSPAILSVSPGERVSFSCRASQSIGTSIHWYQQRTNGSPRLLIKYASESISGIPSRFSGSGSGTYFTLTINSVESEDIADYYCQQNNNWPTTFGGGTKLEIK
^^^^^^ ^^^ ^^^^^^^^^

Best Alignment of Tomuzotuximab CDR-H3 chain to a sequence from OAS
QVQLKQSGPGLVQPSQSLSITCTVSGFSLTNYGVHWVRQSPGKGLEWLGVIWSG-GNTDYNTPFTSRLSINKDNSKSQVFFKMNSLQSNDTAIYYCARALTYYDYEFAYWGQGTLVTVST
.|||.|||..||.|..|....|..||...|.|...|..|..||.|||.|.......|.||..|......|....|.|.|......|.|.|.|.|||||||.||||||||||||||||||.
-VQLQQSGAELVKPGASVKMSCKASGYTFTTYPIEWMKQNHGKSLEWIGNFHPYNDDTKYNEKFKGKATLTVEKSSSTVYLELSRLTSDDSAVYYCARALIYYDYEFAYWGQGTLVTVSA
^^^^^^^^ ^^^^^^^^ ^^^^^^^^^^^^^

Therapeutic : Tovetumab

Best Alignment of Tovetumab heavy chain to a sequence from OAS
QVQLVESGGGLVKPGGSLRLSCAASGFTFSDYYMNWIRQAPGKGLEWVSYISSSGSIIYYADSVKGRFTISRDNAKNSLYLQMNSLRAEDTAVYYCAREGRIA-------ARGMDVWGQGTTVTVSS
||||||||||||||||||||||||||||||||||.|||||||||||||||||||||.|||||||||||||||||||||||||||||||||||||||||.||.|.........|||||||||||||||
QVQLVESGGGLVKPGGSLRLSCAASGFTFSDYYMSWIRQAPGKGLEWVSYISSSGSTIYYADSVKGRFTISRDNAKNSLYLQMNSLRAEDTAVYYCARVGRAAAGNQRTDYYGMDVWGQGTTVTVSS
^^^^^^^^ ^^^^^^^^ ^^^^^^^^^^^^^^^^^^^

Best Alignment of Tovetumab light chain to a sequence from OAS
DIQMTQSPSSLSASVGDRVSITCRPSQSFSRYINWYQQKPGKAPKLLIHAASSLVGGVPSRFSGSGSGTDFTLTISSLQPEDFATYYCQQTYSNPPITFGQGTRLEMK
|||||||||||||||||||.||||.|||.|.|.|||||||||||||||||||||..||||||||||||||||||||||||||||||||||.||.||||||||||||.|
DIQMTQSPSSLSASVGDRVTITCRASQSISSYLNWYQQKPGKAPKLLIHAASSLQSGVPSRFSGSGSGTDFTLTISSLQPEDFATYYCQQSYSTPPITFGQGTRLEIK
^^^^^^ ^^^ ^^^^^^^^^^

Best Alignment of Tovetumab heavy chain CDRs to a sequence from OAS
QVQLVESGGGLVKPGGSLRLSCAASGFTFSDYYMNWIRQAPGKGLEWVSYISSSGSIIYYADSVKGRFTISRDNAKNSLYLQMNSLRAEDTAVYYCAREGRIAARGMDVWGQGTTVTVSS
................||||||||||||||||||.|||||||||||||||||||.|...|||||||||||||||||||||||||||||||||||||.|||||||||||||||||||

----------------SLRLSCAASGFTFSDYYMSWIRQAPGKGLEWVSYISSSSSHTNYADSVKGRFTISRDNAKNSLYLQMNSLRAEDTAVYYCAREPRIAARGMDVWGQGTTVTVSS
^^^^^^^^ ^^^^^^^^ ^^^^^^^^^^^^^

Best Alignment of Tovetumab light CDRs chain to a sequence from OAS
DIQMTQSPSSLSASVGDRVSITCRPSQSFSRYINWYQQKPGKAPKLLIHAASSLVGGVPSRFSGSGSGTDFTLTISSLQPEDFATYYCQQTYSNPPITFGQGTRLEMK
||||||.||.|||||||||.||||.|||.|||.||||||||||||.||.|||.|..|||||||||||||||||||||||||||||.|||||||||.||||||||||.|
DIQMTQTPSPLSASVGDRVTITCRASQSISRYLNWYQQKPGKAPKPLIYAASNLQSGVPSRFSGSGSGTDFTLTISSLQPEDFATDYCQQTYSNPFITFGQGTRLEIK
^^^^^^ ^^^ ^^^^^^^^^^

Best Alignment of Tovetumab CDR-H3 chain to a sequence from OAS
QVQLVESGGGLVKPGGSLRLSCAASGFTFSDYYMNWIRQAPGKGLEWVSYISSSGSIIYYADSVKGRFTISRDNAKNSLYLQMNSLRAEDTAVYYCAREGRIAARGMDVWGQGTTVTVSS
................||||||||||||||||||.|||||||||||||||||||||.|...||||||||||||||||||||||||||||||||||||||||.||||||||||||||||||||
----------------SLRLSCAASGFTFSDYYMSWIRQAPGKGLEWVSYISSSSSHTNYADSVKGRFTISRDNAKNSLYLQMNSLRAEDTAVYYCAREPRIAARGMDVWGQGTTVTVSS
^^^^^^^^ ^^^^^^^^ ^^^^^^^^^^^^^

Therapeutic : Tralokinumab

Best Alignment of Tralokinumab heavy chain to a sequence from OAS
QVQLVQSGAEVKKPGASVKVSCKASGYTFTNYGLSWVRQAPGQGLEWMGWISANNGDTNYGQEFQGRVTMTTDTSTSTAYMELRSLRSDDTAVYYCARDSSSSWARWFFDLWGRGTLVTVSS
||||||||||||||||||||||||||||||.||.|||||||||||||||||||.||.|||.|..|||||||||||||||||||||||||||||||||||..|||..|.||||||||||||||
QVQLVQSGAEVKKPGASVKVSCKASGYTFTSYGISWVRQAPGQGLEWMGWISAYNGNTNYAQKLQGRVTMTTDTSTSTAYMELRSLRSDDTAVYYCARDKGSSWIYWYFDLWGRGTLVTVSS
^^^^^^^^ ^^^^^^^^ ^^^^^^^^^^^^^^^

Best Alignment of Tralokinumab light chain to a sequence from OAS
SYVLTQPPSVSVAPGKTARITCGGNIIGSKLVHWYQQKPGQAPVLVIYDDGDRPSGIPERFSGSNSGNTATLTISRVEAGDEADYYCQVWDTGSDPVVFGGGTKLTVL
||.||||||||||||||||||||||.||||.|||||||||||||||.||||||||||||||||||||||||||||||||||||||||||||.||||||||||||||||
SYELTQPPSVSVAPGKTARITCGGNNIGSKSVHWYQQKPGQAPVLVVYDDGDRPSGIPERFSGSNSGNTATLTISRVEAGDEADYYCQVWDSGSDPVVFGGGTKLTVL
^^^^^^ ^^^ ^^^^^^^^^^^

Best Alignment of Tralokinumab heavy chain CDRs to a sequence from OAS
QVQLVQSGAEVKKPGASVKVSCKASGYTFTNYGLSWVRQAPGQGLEWMGWISANNGDTNYGQEFQGRVTMTTDTSTSTAYMELRSLRSDDTAVYYCARDSSSSWARWFFDLWGRGTLVTVSS
.....|||||||||||||||||.||||||||||.|||||||||||||||||||.||||.|.|..||||||||||.|||||||||||.|||||||.|||||..|..|.|||||||||||||||
-----QSGAEVKKPGASVKVSCTASGYTFTNYGISWVRQAPGQGLEWMGWISAYNGDTDYAQKLQGRVTMTTDTLTSTAYMELRSLTSDDTAVYHCARDSQLSPTRGFFDLWGRGTLVTVSS
^^^^^^^^ ^^^^^^^^ ^^^^^^^^^^^^^^^

Best Alignment of Tralokinumab light CDRs chain to a sequence from OAS
SYVLTQPPSVSVAPGKTARITCGGNIIGSKLVHWYQQKPGQAPVLVIYDDGDRPSGIPERFSGSNSGNTATLTISRVEAGDEADYYCQVWDTGSDPVVFGGGTKLTVL
|||||||||||||||.|||||||||.||||.|||||||||||||||.|||||||||||||||||||||||||||.|||||||||.|||||.|||||||||||||||||
SYVLTQPPSVSVAPGQTARITCGGNNIGSKSVHWYQQKPGQAPVLVVYDDGDRPSGIPERFSGSNSGNTATLTITRVEAGDEADFYCQVWETGSDPVVFGGGTKLTVL
^^^^^^ ^^^ ^^^^^^^^^^^

Best Alignment of Tralokinumab CDR-H3 chain to a sequence from OAS
QVQLVQSGAEVKKPGASVKVSCKASGYTFTNYGLSWVRQAPGQGLEWMGWISANNGDTNYGQEFQGRVTMTTDTSTSTAYMELRSLRSDDTAVYYCARDSSSSWARWFFDLWGRGTLVTVSS
..............|.|.|.||.|||.||..||..|||||||.||||...|........|.....||.|...|.|..|.|....|||..|||||||||||||||..|.||||||||||||||
--------------GESLKISCAASGFTFSSYGMHWVRQAPGKGLEWVAVIWYDGSNKYYADSVKGRFTISRDNSKNTLYLQMNSLRAEDTAVYYCARDSSSSWSNWYFDLWGRGTLVTVSS
^^^^^^^^ ^^^^^^^^ ^^^^^^^^^^^^^^^

Therapeutic : Trastuzumab

Best Alignment of Trastuzumab heavy chain to a sequence from OAS
EVQLVESGGGLVQPGGSLRLSCAASGFNIKDTYIHWVRQAPGKGLEWVARIYPTNGYTRYADSVKGRFTISADTSKNTAYLQMNSLRAEDTAVYYCSRWGGDG-FYAMDYWGQGTLVTVSS
.|||.||||||||||||||||||||||||...|||||||||||||||||.||.....|.|||||||||||||||||||||||||||||||||||||.|....|...|||||||||||||||
QVQLQESGGGLVQPGGSLRLSCAASGFNISSYYIHWVRQAPGKGLEWVASIYSSSSSTYYADSVKGRFTISADTSKNTAYLQMNSLRAEDTAVYYCARGPVFGPYAAMDYWGQGTLVTVSS
^^^^^^^^ ^^^^^^^^ ^^^^^^^^^^^^^^

Best Alignment of Trastuzumab light chain to a sequence from OAS
DIQMTQSPSSLSASVGDRVTITCRASQDVNTAVAWYQQKPGKAPKLLIYSASFLYSGVPSRFSGSRSGTDFTLTISSLQPEDFATYYCQQHYTTPPTFGQGTKVEIK
||||||||||||||||||||||||||||..|..||||||||||||||||.||.|.||||.|||||.|||||||||||||||||||||||||.|||.|||||||||||
DIQMTQSPSSLSASVGDRVTITCRASQDSGTWLAWYQQKPGKAPKLLIYKASTLHSGVPARFSGSGSGTDFTLTISSLQPEDFATYYCQQHHTTPRTFGQGTKVEIK
^^^^^^ ^^^ ^^^^^^^^^

Best Alignment of Trastuzumab heavy chain CDRs to a sequence from OAS
EVQLVESGGGLVQPGGSLRLSCAASGFNIKDTYIHWVRQAPGKGLEWVARIYPTNGYTRYADSVKGRFTISADTSKNTAYLQMNSLRAEDTAVYYCSRWGGDGFYAMDYWGQGTLVTVSS
||.|||||..||.||.|..|||.||||||||||.|||.|.|..||||..||.|.||.|.|.....|..||.|||.||||||..||..|||||||..||||||.|||||||||..|||||
EVMLVESGAELVKPGASVKLSCTASGFNIKDTYMHWVKQRPEQGLEWIGRIDPANGNTKYDPKFQGKATITADTSSNTAYLQLSSLTSEDTAVYYGARWGGDGVYAMDYWGQGPSVTVSS
^^^^^^^^ ^^^^^^^^ ^^^^^^^^^^^^^

Best Alignment of Trastuzumab light CDRs chain to a sequence from OAS
DIQMTQSPSSLSASVGDRVTITCRASQDVNTAVAWYQQKPGKAPKLLIYSASFLYSGVPSRFSGSRSGTDFTLTISSLQPEDFATYYCQQHYTTPPTFGQGTKVEIK
||.||||....|.||||||.|||.|||||||||||||||||..||||||||||...|||.||.||.||||||.||.|.|.||.|.|||||||.||||||.|||.|||
DIMMTQSHKFMSTSVGDRVSITCKASQDVNTAVAWYQQKPGHSPKLLIYSASFRCTGVPDRFTGSGSGTDFTFTINSVQAEDLAVYYCQQHYSTPPTFGGGTKLEIK
^^^^^^ ^^^ ^^^^^^^^^

Best Alignment of Trastuzumab CDR-H3 chain to a sequence from OAS
EVQLVESGGGLVQPGGSLRLSCAASGFNIKDTYIHWVRQAPGKGLEWVARIYPTNGYTRYADSVKGRFTISADTSKNTAYLQMNSLRAEDTAVYYCSRWGGDGFYAMDYWGQGTLVTVSS
.|||...|..||.||.|..|||.|||.........||.|.|..||||..||.|.....||....|.......|.|..|||.|..||..||.||||||||||..|||||||||||.|||||
QVQLQQPGAELVRPGASVKLSCKASGYTFTNYWMNWVKQRPEQGLEWIGRIDPYDSEIRYNQKFKDKAILTVDKSSRTAYMQLSSLTSEDSAVYYCSRWGGITFYAMDYWGQGTSVTVSS
^^^^^^^^ ^^^^^^^^ ^^^^^^^^^^^^^

Therapeutic : Tregalizumab

Best Alignment of Tregalizumab heavy chain to a sequence from OAS
EEQLVESGGGLVKPGGSLRLSCAASGFSFSDCRMYWLRQAPGKGLEWIGVISVKSENYGANYAESVRGRFTISRDDSKNTVYLQMNSLKTEDTAVYYCSASYYRYD-VGAWFAYWGQGTLVTVSS
|.|||||||||||||||||||||||||.||...|.|.||||||||||.|.|..|...|...||.||.|||||||||||||.|||||||||||||||||..|||.......||.||||||||||||
EVQLVESGGGLVKPGGSLRLSCAASGFTFSNAWMSWVRQAPGKGLEWVGRIRSKANSYATAYAASVKGRFTISRDDSKNTAYLQMNSLKTEDTAVYYCTSSYYDFWSGSLWFDYWGQGTLVTVSS
^^^^^^^^ ^^^^^^^^^^ ^^^^^^^^^^^^^^^^

Best Alignment of Tregalizumab light chain to a sequence from OAS
DIVMTQSPDSLAVSLGERATINCRASKSVST--SGYSYIYWYQQKPGQPPKLLIYLASILESGVPDRFSGSGSGTDFTLTISSLQAEDVAVYYCQHSRELPWTFGQGTKVEIK
||||||||||||||||||||||||||.||....|......|||.|||||||||||.|||.||||||||||||||||||||||||||||||||||||....|||||||||||||
DIVMTQSPDSLAVSLGERATINCRASQSVLHSSSNKNFLAWYQHKPGQPPKLLIYWASIRESGVPDRFSGSGSGTDFTLTISSLQAEDVAVYYCQHYYSTPWTFGQGTKVEIK
^^^^^^^^^^^^ ^^^ ^^^^^^^^^

Best Alignment of Tregalizumab heavy chain CDRs to a sequence from OAS
EEQLVESGGGLVKPGGSLRLSCAASGFSFSDCRMYWLRQAPGKGLEWIGVISVKSENYGANYAESVRGRFTISRDDSKNTVYLQMNSLKTEDTAVYYCSASYYRYDVGAWFAYWGQGTLVTVSS
..||||.|||||.||.||.|||..|||.||..||.||||.|||.||||.||.|||.||||||||||.|||.|||||||..|||.||....||||.|.||.|.|.||.||||||||||||||||.
QVQLVETGGGLVRPGNSLKLSCVTSGFTFSNYRMHWLRQPPGKRLEWIAVITVKSDNYGANYAESVKGRFAISRDDSKSSVYLEMNRVREEDTATYFCSRSGYDYDEGAWFAYWGQGTLVTVSA
^^^^^^^^ ^^^^^^^^^^ ^^^^^^^^^^^^^^^

Best Alignment of Tregalizumab light CDRs chain to a sequence from OAS
DIVMTQSPDSLAVSLGERATINCRASKSVSTSGYSYIYWYQQKPGQPPKLLIYLASILESGVPDRFSGSGSGTDFTLTISSLQAEDVAVYYCQHSRELPWTFGQGTKVEIK
|||.||||.|||||||.||||.||||||||||||||..||||||||||||.|||||.||||||.|||||||||||||.|.....||.|.||||||||||||||.|||.|||
DIVLTQSPASLAVSLGQRATISCRASKSVSTSGYSYMHWYQQKPGQPPKLIIYLASNLESGVPARFSGSGSGTDFTLNIHPVEEEDAATYYCQHSRELPWTFGGGTKLEIK
^^^^^^^^^^ ^^^ ^^^^^^^^^

Best Alignment of Tregalizumab CDR-H3 chain to a sequence from OAS
EEQLVESGGGLVKPGGSLRLSCAASGFSFSDCRMYWLRQAPGKGLEWIGVISVKSENYGANYAESVRGRFTISRDDSKNTVYLQMNSLKTEDTAVYYCSASYYRYDVGAWFAYWGQGTLVTVSS
|.||..||..|..||.|...||.|||..|....|.|..|.||.||||||.|.....|....|.....|........|..|.|....||..||.|||||.||||||||||||||||||||||||.

```
EVQLQQSGTVLARPGASVKMSCKASGYTFTSYWMHWVKQRPGQGLEWIGAIYPG--NSDTSYNQKFKGKAKLTAVTSTSTAYMELSSLTNEDSAVYYCTASYYRYDVGAWFAYWGQGTLVTVSA
                   ^^^^^^^^          ^^^^^^^^^^                      ^^^^^^^^^^^^^^^
```

Therapeutic : Tremelimumab

Best Alignment of Tremelimumab heavy chain to a sequence from OAS

```
QVQLVESGGGVVQPGRSLRLSCAASGFTFSSYGMHWVRQAPGKGLEWVAVIWYDGSNKYYADSVKGRFTISRDNSKNTLYLQMNSLRAEDTAVYYCARDPRGATL-YYYYYGMDVWGQGTTVTVSS
|||||||||||||||||||||||||||||||||||||||||||||||||||||||||||||||||||||||||||||||||||||||||||||||||||..||||.||||||||||||||||||||
QVQLVESGGGVVQPGRSLRLSCAASGFTFSSYGMHWVRQAPGKGLEWVAVIWYDGSNKYYADSVKGRFTISRDNSKNTLYLQMNSLRAEDTAVYYCARDRVGATLYYYYYYGMDVWGQGTTVTVSS
                   ^^^^^^^^          ^^^^^^^^                        ^^^^^^^^^^^^^^^^^^
```

Best Alignment of Tremelimumab light chain to a sequence from OAS

```
DIQMTQSPSSLSASVGDRVTITCRASQSINSYLDWYQQKPGKAPKLLIYAASSLQSGVPSRFSGSGSGTDFTLTISSLQPEDFATYYCQQYYSTPFTFGPGTKVEIK
|||||||||||||||||||||||||||||||||.||||||||||||||||||||||||||||||||||||||||||||||||||||||||.||||||||.|||||||
DIQMTQSPSSLSASVGDRVTITCRASQSINSYLNWYQQKPGKAPKLLIYAASSLQSGVPSRFSGSGSGTDFTLTISSLQPEDFATYYCQQSYSTPFTFGGGTKVEIK
                    ^^^^^^          ^^^                     ^^^^^^^^^
```

Best Alignment of Tremelimumab heavy chain CDRs to a sequence from OAS

```
QVQLVESGGGVVQPGRSLRLSCAASGFTFSSYGMHWVRQAPGKGLEWVAVIWYDGSNKYYADSVKGRFTISRDNSKNTLYLQMNSLRAEDTAVYYCARDPRGATLYYYYYGMDVWGQGTTVTVSS
..............|.||||||||||||||||||||||||||||||||||||||||||||||||||||||||||||||||||||||.|||.|||||||||||||||||||.
--------------GGSLRLSCAASGFTFSSYGMHWVRQAPGKGLEWVAVIWYDGSNKYYADSVKGRFTISRDNSKNTLYLQMNSLRAEDTAVYYCARDPLGATVYYYYYGMDVWGQGTTVTVS-
                   ^^^^^^^^          ^^^^^^^^                        ^^^^^^^^^^^^^^^^^
```

Best Alignment of Tremelimumab light CDRs chain to a sequence from OAS

```
DIQMTQSPSSLSASVGDRVTITCRASQSINSYLDWYQQKPGKAPKLLIYAASSLQSGVPSRFSGSGSGTDFTLTISSLQPEDFATYYCQQYYSTPFTFGPGTKVEIK
|||||||||||||||||||||||||||||||||.||||.|||||||||||||.||||||.|||||||.||||||||||||||||||||||.|||||||||||||.||
DIQMTQSPSSLSASVGDRVTITCRASQSINSYLNWYQQRPGKAPKLLIYAASGLQSGVPARFSGSGSETDFTLTISSLQPEDFATYYCQQSYSTPFTFGPGTKVDIK
                    ^^^^^^          ^^^                     ^^^^^^^^^
```

Best Alignment of Tremelimumab CDR-H3 chain to a sequence from OAS

```
QVQLVESGGGVVQPGRSLRLSCAASGFTFSSYGMHWVRQAPGKGLEWVAVIWYDGSNKYYADSVKGRFTISRDNSKNTLYLQMNSLRAEDTAVYYCARDPRGATLYYYYYGMDVWGQGTTVTVSS
..............|.||||||||||||||||.|.||||||||||||||.|..|||.|||.|||||||||||||.||.||||||||||||||||||||||.|||.||||||||||||||||||||
--------------GGSLRLSCAASGFTFSSYWMSWVRQAPGKGLEWVANIKQDGSEKYYVDSVKGRFTISRDNAKNSLYLQMNSLRAEDTAVYYCARDPVGATFYYYYYGMDVWGQGTTVTVSS
                   ^^^^^^^^          ^^^^^^^^                        ^^^^^^^^^^^^^^^^^
```

Therapeutic : Trevogrumab

Best Alignment of Trevogrumab heavy chain to a sequence from OAS

```
EVQVLESGGDLVQPGGSLRLSCAASGFTFSAYAMTWVRQAPGKGLEWVSAISGSGGSAYYADSVKGRFTISRDNSKNTVYLQMNSLRAEDTAVYYCAKDGAWK------MSGLDVWGQGTTVIVSS
|||.|||||.||||||||||||||||||||.|||.||||||||||||||||||||||.||||||||||||||||||||.|||||||||||||||||||||.|.........|.|||||||||.|||
EVQLLESGGGLVQPGGSLRLSCAASGFTFSSYAMSWVRQAPGKGLEWVSAISGSGGSTYYADSVKGRFTISRDNSKNTLYLQMNSLRAEDTAVYYCAKDGKWRHNLSYYYYGMDVWGQGTTVTVSS
                   ^^^^^^^^          ^^^^^^^^                        ^^^^^^^^^^^^^^^^^^
```

Best Alignment of Trevogrumab light chain to a sequence from OAS

```
DIQMTQSPASLSASVGDRVTITCRASQDISDYLAWYQQKPGKIPRLLIYTTSTLQSGVPSRFRGSGSGTDFTLTISSLQPEDVATYYCQKYDSAPLTFGGGTKVEIK
||||||||.|||||||||||||||||||||.|||||||||||.|.|||||||||||||||||.||||||||||||||||||||||||||||.|||||||||||||||
DIQMTQSPSSLSASVGDRVTITCRASQDISNYLAWYQQKPGKVPKLLIYTTSTLQSGVPSRFSGSGSGTDFTLTISSLQPEDVATYYCQKYSSAPLTFGGGTKVEIK
                    ^^^^^^          ^^^                     ^^^^^^^^^
```

Best Alignment of Trevogrumab heavy chain CDRs to a sequence from OAS

```
EVQVLESGGDLVQPGGSLRLSCAASGFTFSAYAMTWVRQAPGKGLEWVSAISGSGGSAYYADSVKGRFTISRDNSKNTVYLQMNSLRAEDTAVYYCAKDGAWKMSGLDVWGQGTTVIVSS
..............||||||||||||||||.|||.||||||||||||||||||||||.||||||||||||||||||||.||||||||||||||||||||.||..||.|||||||||.||.
--------------GGSLRLSCAASGFTFSSYAMSWVRQAPGKGLEWVSAISGSGGSTYYADSVKGRFTISRDNSKNTLYLQMNSLRAEDTAVYYCAKDRAWLPSGMDVWGQGTTVTVS-
                   ^^^^^^^^          ^^^^^^^^                        ^^^^^^^^^^^^^
```

Best Alignment of Trevogrumab light CDRs chain to a sequence from OAS

```
DIQMTQSPASLSASVGDRVTITCRASQDISDYLAWYQQKPGKIPRLLIYTTSTLQSGVPSRFRGSGSGTDFTLTISSLQPEDVATYYCQKYDSAPLTFGGGTKVEIK
||||||||.||||||||||||||.||||||.|||||||||||.|.|||||||||||||||||.||||||||||||||||||||||||||||.||||||||||.||||
DIQMTQSPSSLSASVGDRVTITCQASQDISNYLAWYQQKPGKVPKLLIYTTSTLQSGVPSRFSGSGSGTDFTLTISSLQPEDVATYYCQKYSSAPLTFGGGAKVEIK
                    ^^^^^^          ^^^                     ^^^^^^^^^
```

Best Alignment of Trevogrumab CDR-H3 chain to a sequence from OAS

```
EVQVLESGGDLVQPGGSLRLSCAASGFTFSAYAMTWVRQAPGKGLEWVSAISGSGGSAYYADSVKGRFTISRDNSKNTVYLQMNSLRAEDTAVYYCAKDGAWKMSGLDVWGQGTTVIVSS
................||||||||||||||.|.|.|||||||||||||..|...|...||||||||||||||||||||.||||||||||||||||||||||.|..|||.|||||.|.|||
----------------SLRLSCAASGFTFSSYGMHWVRQAPGKGLEWVAVIWYDGSNKYYADSVKGRFTISRDNSKNTLYLQMNSLRAEDTAVYYCAKDGATKIMGLDYWGQGTLVTVSS
                   ^^^^^^^^          ^^^^^^^^                        ^^^^^^^^^^^^^
```

Therapeutic : Ublituximab

Best Alignment of Ublituximab heavy chain to a sequence from OAS

```
QAYLQQSGAELVRPGASVKMSCKASGYTFTSYNMHWVKQTPRQGLEWIGGIYPGNGDTSYNQKFKGKATLTVGKSSSTAYMQLSSLTSEDSAVYFCARYDN----YAMDYWGQGTSVTVSS
|||||||||||||||||||||||||||||||||||||||||||||||||.|||||||||||||||||||||.||||||||||||||||||||||||||.||.....||||||||||||||||
QAYLQQSGAELVRPGASVKMSCKASGYTFTSYNMHWVKQTPRQGLEWIGAIYPGNGDTSYNQKFKGKATLTVDKSSSTAYMQLSSLTSEDSAVYFCARGDYGSSYNYAMDYWGQGTSVTVSS
                   ^^^^^^^^          ^^^^^^^^                        ^^^^^^^^^^^^^^^
```

Best Alignment of Ublituximab light chain to a sequence from OAS

```
QIVLSQSPAILSASPGEKVTMTCRASSSVSYMHWYQQKPGSSPKPWIYATSNLASGVPARFSGSGSGTSYSFTISRVEAEDAATYYCQQWTFNPPTFGGGTRLEIK
|||||||||||||||||||||||||||||||||||||||||||||||||||||||||||||||||||||||.||||||||||||||||||..|||||||||.||||
QIVLSQSPAILSASPGEKVTMTCRASSSVSYMHWYQQKPGSSPKPWIYATSNLASGVPARFSGSGSGTSYSLTISRVEAEDAATYYCQQWSSNPPTFGGGTKLEIK
                    ^^^^^          ^^^                     ^^^^^^^^^
```

Best Alignment of Ublituximab heavy chain CDRs to a sequence from OAS

```
QAYLQQSGAELVRPGASVKMSCKASGYTFTSYNMHWVKQTPRQGLEWIGGIYPGNGDTSYNQKFKGKATLTVGKSSSTAYMQLSSLTSEDSAVYFCARYDYNYAMDYWGQGTSVTVSS
|..||||||||||||||||||||||||||||||||||||||||||||||.|||||||||||||||||||||.|||||||||||||||||||||||||||||.||||||||||||||||
QVQLQQSGAELVRPGASVKMSCKASGYTFTSYNMHWVKQTPRQGLEWIGAIYPGNGDTSYNQKFKGKATLTVDKSSSTAYMQLSSLTSEDSAVYFCARYDYDYAMDYWGQGTSVTVSS
                   ^^^^^^^^          ^^^^^^^^                        ^^^^^^^^^^^
```

Best Alignment of Ublituximab light CDRs chain to a sequence from OAS

```
QIVLSQSPAILSASPGEKVTMTCRASSSVSYMHWYQQKPGSSPKPWIYATSNLASGVPARFSGSGSGTSYSFTISRVEAEDAATYYCQQWTFNPPTFGGGTRLEIK
||||.|||||.||||||||||||.||||||||||||||.|.|||.||||||||||||||||||||||||||.||||.|||||||||||||..||||||.||.||.|
QIVLTQSPAIMSASPGEKVTMTCSASSSVSYMHWYQQKSGTSPKRWIYATSNLASGVPARFSGSGSGTSYSLTISRMEAEDAATYYCQQWSSNPPTFGAGTKLELK
                    ^^^^^          ^^^                     ^^^^^^^^^
```

Best Alignment of Ublituximab CDR-H3 chain to a sequence from OAS

```
QAYLQQSGAELVRPGASVKMSCKASGYTFTSYNMHWVKQTPRQGLEWIGGIYPGNGDTSYNQKFKGKATLTVGKSSSTAYMQLSSLTSEDSAVYFCARYDYNYAMDYWGQGTSVTVSS
|..|||.|||||.||||||.|||.||||||||.||||||.|..||||||.|.|..|.|.||.|||.||||||.|.|||||||||||||||||||.|||||||||||||||||||||||
QVQLQQPGAELVKPGASVKLSCKSSGYTFTSYWMHWVKQRPGRGLEWIGRIDPNSGGTKYNEKFKSKATLTVDKPSSTAYMQLSSLTSEDSAVYYCARYDYNYAMDYWGQGTSVTVSS
                   ^^^^^^^^          ^^^^^^^^                        ^^^^^^^^^^^
```

Therapeutic : Urelumab

Best Alignment of Urelumab heavy chain to a sequence from OAS

```
QVQLQQWGAGLLKPSETLSLTCAVYGGSFSGYYWSWIRQSPEKGLEWIGEINHGGYVTYNPSLESRVTISVDTSKNQFSLKLSSVTAADTAVYYCARDYGPGN----YDWYFDLWGRGTLVTVSS
|||||||||||||||||||||||||||||||||||||||.|.|||||||||||.|...|||||.|||||||||||||||||||||||||||||||||.||.|.......||||||||||||||||
QVQLQQWGAGLLKPSETLSLTCAVYGGSFSGYYWSWIRQPPGKGLEWIGEINHSGSTNYNPSLKSRVTISVDTSKNQFSLKLSSVTAADTAVYYCARSYGSGSYYNRRYWYFDLWGRGTLVTVSS
                   ^^^^^^^^          ^^^^^^^                       ^^^^^^^^^^^^^^^^^^
```

Best Alignment of Urelumab light chain to a sequence from OAS

```
EIVLTQSPATLSLSPGERATLSCRASQSVSSYLAWYQQKPGQAPRLLIYDASNRATGIPARFSGSGSGTDFTLTISSLEPEDFAVYYCQQRSNWPPALTFGGGTKVEIK
|||||||||||||||||||||||||||||||||||||||||||||||||||||||||||||||||||||||||||||||||||||||||||||||||||||||||||||
EIVLTQSPATLSLSPGERATLSCRASQSVSSYLAWYQQKPGQAPRLLIYDASNRATGIPARFSGSGSGTDFTLTISSLEPEDFAVYYCQQRSNWPPALTFGGGTKVEIK
                         ^^^^^^                 ^^^                                   ^^^^^^^^^^^
```

Best Alignment of Urelumab heavy chain CDRs to a sequence from OAS

```
QVQLQQWGAGLLKPSETLSLTCAVYGGSFSGYYWSWIRQSPEKGLEWIGEINHGGYVTYNPSLESRVTISVDTSKNQFSLKLSSVTAADTAVYYCARDYGPGNYDWYFDLWGRGTLVTVSS
.............||||||||||||||||||||||||||.|.|||||||||||.|...|||||.||||||||||||||||||||||||||||||||||||.||..||||||||||||||||
-------------PSETLSLTCAVYGGSFSGYYWSWIRQPPGKGLEWIGEINHSGSTNYNPSLKSRVTISVDTSKNQFSLKLSSVTAADTAVYYCARDYGSGNTQWYFDLWGRGTLVTVSS
                         ^^^^^^^^                 ^^^^^^^                                    ^^^^^^^^^^^^^^^
```

Best Alignment of Urelumab light CDRs chain to a sequence from OAS

```
EIVLTQSPATLSLSPGERATLSCRASQSVSSYLAWYQQKPGQAPRLLIYDASNRATGIPARFSGSGSGTDFTLTISSLEPEDFAVYYCQQRSNWPPALTFGGGTKVEIK
|||||||||||||||||||||||||||||||||||||||||||||||||||||||||||||||||||||||||||||||||||||||||||||||||||||||||||||
EIVLTQSPATLSLSPGERATLSCRASQSVSSYLAWYQQKPGQAPRLLIYDASNRATGIPARFSGSGSGTDFTLTISSLEPEDFAVYYCQQRSNWPPALTFGGGTKVEIK
                         ^^^^^^                 ^^^                                   ^^^^^^^^^^^
```

Best Alignment of Urelumab CDR-H3 chain to a sequence from OAS

```
QVQLQQWGAGLLKPSETLSLTCAVYGGSFSGYYWSWIRQSPEKGLEWIGEINHG-GYVTYNPSLESRVTISVDTSKNQFSLKLSSVTAADTAVYYCARDYGPGNYDWYFDLWGRGTLVTVSS
.............|...|.|.||..|..||.|...|.||.|.|||||...|........|..|...|.|||.|.|||...|...|..|.||||||||||.||.|||||||||||||||||||
-------------PGGSLRLSCAASGFTFSSYWMHWVRQAPGKGLEWVAVISYDGSNKYYADSVKGRFTISRDNSKNTLYLQMNSLRAEDTAVYYCARDGGPWNYDWYFDLWGRGTLVTVSS
                         ^^^^^^^^                 ^^^^^^^^                                    ^^^^^^^^^^^^^^^
```

Therapeutic : Ustekinumab

Best Alignment of Ustekinumab heavy chain to a sequence from OAS

```
EVQLVQSGAEVKKPGESLKISCKGSGYSFTTYWLGWVRQMPGKGLDWIGIMSPVDSDIRYSPSFQGQVTMSVDKSITTAYLQWNSLKASDTAMYYCARRRPG---QGYFDFWGQGTLVTVSS
||||||||||||||||||||||||||||||.||||||||||||||.|.||..|.|||.||||||||||||.|.||||.||||||.|||||||||||||||..|....||||.|||||||||||
EVQLVQSGAEVKKPGESLKISCKGSGYSFTSYWLGWVRQMPGKGLEWMGIIYPGDSDTRYSPSFQGQVTISADKSISTAYLQWSSLKASDTAMYYCARRKVGAISWGYFDYWGQGTLVTVSS
                         ^^^^^^^^                 ^^^^^^^^                                    ^^^^^^^^^^^^^^^
```

Best Alignment of Ustekinumab light chain to a sequence from OAS

```
DIQMTQSPSSLSASVGDRVTITCRASQGISSWLAWYQQKPEKAPKSLIYAASSLQSGVPSRFSGSGSGTDFTLTISSLQPEDFATYYCQQYNIYPYTFGQGTKLEIK
||||||||||||||||||||||||||||||||||||||||||||||||||||||||||||||||||||||||||||||||||||||||||||.||||||||||||||
DIQMTQSPSSLSASVGDRVTITCRASQGISSWLAWYQQKPEKAPKSLIYAASSLQSGVPSRFSGSGSGTDFTLTISSLQPEDFATYYCQQYNSYPYTFGQGTKLEIK
                         ^^^^^^                 ^^^                                   ^^^^^^^^^
```

Best Alignment of Ustekinumab heavy chain CDRs to a sequence from OAS

```
EVQLVQSGAEVKKPGESLKISCKGSGYSFTTYWLGWVRQMPGKGLDWIGIMSPVDSDIRYSPSFQGQVTMSVDKSITTAYLQWNSLKASDTAMYYCARRRPGQGYFDFWGQGTLVTVSS
..............||||||||||||||||.||.|||||||||||.|.||.||.|||||||||||||||.|.||||.||||||.||||||||||||||.|..|||||||||||||||||
--------------GESLKISCKGSGYSFTRYWIGWVRQMPGKGLEWMGIISPGDSDIRYSPSFQGQVTISADKSISTAYLQWSSLKASDTAMYYCARERSSQGYFDFWGQGTLVTVSS
                         ^^^^^^^^                 ^^^^^^^^                                    ^^^^^^^^^^^^
```

Best Alignment of Ustekinumab light CDRs chain to a sequence from OAS

```
DIQMTQSPSSLSASVGDRVTITCRASQGISSWLAWYQQKPEKAPKSLIYAASSLQSGVPSRFSGSGSGTDFTLTISSLQPEDFATYYCQQYNIYPYTFGQGTKLEIK
||||||||||||||||||||||||||||||||||||||.|||||||||||||.||||||||||||||||||||||||||||||||||||||||||||||||||||||
DIQMTQSPSSLSASVGDRVTITCRASQGISSWLAWYQQRPEKAPKSLIYAASTLQSGVPSRFSGSGSGTDFTLTISSLQPEDFATYYCQQYNIYPYTFGQGTKLEIK
                         ^^^^^^                 ^^^                                   ^^^^^^^^^
```

Best Alignment of Ustekinumab CDR-H3 chain to a sequence from OAS

```
EVQLVQSGAEVKKPGESLKISCKGSGYSFTTYWLGWVRQMPGKGLDWIGIMSPVDSDIRYSPSFQGQVTMSVDKSITTAYLQWNSLKASDTAMYYCARRRPGQGYFDFWGQGTLVTVSS
...|.|||.||||.|.|.|.|||.|.|.||.|...||||.||.||.|.|..|.......|...|||.|||..|.|.||.|....||...|||.|||||.|||.|||||||||.||||||
---LGQSGGEVKKSGASVKVSCKASHYTFTDYHINWVRQAPGQGLEWMGCISTYNGGTKYAQKFQGRVTMTTDTSTTTVYMELRSLRSDDTAVYYCARDRPGYGYFDFWGQGSLVTVSS
                         ^^^^^^^^                 ^^^^^^^^                                    ^^^^^^^^^^^^
```

Therapeutic : Utomilumab

Best Alignment of Utomilumab heavy chain to a sequence from OAS

```
EVQLVQSGAEVKKPGESLRISCKGSGYSFSTYWISWVRQMPGKGLEWMGKIYPGDSYTNYSPSFQGQVTISADKSISTAYLQWSSLKASDTAMYYCARGYG-IFDYWGQGTLVTVSS
||||||||||||||||||.|||||||||||||||.||||||||||||||.|||.||.|.|||||||||||||||||||||||||||||||||||||||.||..||||||||||||||
EVQLVQSGAEVKKPGESLKISCKGSGYSFSTYWIGWVRQMPGKGLEWMGIIYPRDSDTRYSPSFQGQVTISADKSISTAYLQWSSLKASDTAMYYCARVYGYEFDYWGQGTLVTVSS
                         ^^^^^^^^                 ^^^^^^^^                                    ^^^^^^^^^^
```

Best Alignment of Utomilumab light chain to a sequence from OAS

```
SYELTQPPSVSVSPGQTASITCSGDNIGDQYAHWYQQKPGQSPVLVIYQDKNRPSGIPERFSGSNSGNTATLTISGTQAMDEADYYCATYTGFGSLAVFGGGTKLTVL
|||||||||||||||||||||||||..||.||.|||||||||||||||||..|||||||||||||||||||||||||||||||||||..|.|...|||||||||||||
SYELTQPPSVSVSPGQTASITCSGDKLGDKYACWYQQKPGQSPVLVIYQDSKRPSGIPERFSGSNSGNTATLTISGTQAMDEADYYCCSYAG--LLAVFGGGTKLTVL
                         ^^^^^^                 ^^^                                   ^^^^^^^^^^^
```

Best Alignment of Utomilumab heavy chain CDRs to a sequence from OAS

```
EVQLVQSGAEVKKPGESLRISCKGSGYSFSTYWISWVRQMPGKGLEWMGKIYPGDSYTNYSPSFQGQVTISADKSISTAYLQWSSLKASDTAMYYCARGYGIFDYWGQGTLVTVSS
..............||||.|||.|||||||||||.||||||||||||||.||||||.|.||||||||||||||.||||||||||||||||||.||||||.|.||||||||||||||
--------------GESLKISCQGSGYSFSTYWIGWVRQMPGKGLEWMGIIYPGDSDTRYSPSFQGQVTISADRSISTAYLQWSSLKASDTAIYYCARGSGNFDYWGQGTLVTVSS
                         ^^^^^^^^                 ^^^^^^^^                                    ^^^^^^^^^
```

Best Alignment of Utomilumab light CDRs chain to a sequence from OAS

```
SYELTQPPSVSVSPGQTASITCSGDNIGDQYAHWYQQKPGQSPVLVIYQDKNRPSGIPERFSGSNSGNTATLTISGTQAMDEADYYCATYTGFGSLAVFGGGTKLTVL
.|||||||||||||||||||||||||.||||..|||||||||||||||||..|||||||||||||||||||||||||||||||||||..||...|..||||||.||||
-YELTQPPSVSVSPGQTASITCSGDNLGDQYVCWYQQKPGQSPVLVIYQDSKRPSGIPERFSGSNSGNTATLTISGTQAMDEADYYCSSYTTTTSAGVFGGGTTLTVL
                         ^^^^^^                 ^^^                                   ^^^^^^^^^^^
```

Best Alignment of Utomilumab CDR-H3 chain to a sequence from OAS

```
EVQLVQSGAEVKKPGESLRISCKGSGYSFSTYWISWVRQMPGKGLEWMGKIYPGDSYTNYSPSFQGQVTISADKSISTAYLQWSSLKASDTAMYYCARGYGIFDYWGQGTLVTVSS
.|||..||.....|..||.|.|..||.|...||||.|||||||.|.|......|.|.........||.|.|.|...|...||...|||.|||||||||||||||||..||||
QVQLKESGPGLVAPSQSLSITCTVSGFSLTGYGVNWVRQPPGKGLEWLGMIWGD-GSTDYNSALKSRLSISKDNSKSQVFLKMNSLQTDDTARYYCARGYGIFDYWGQGTTLTVSS
                         ^^^^^^^^                 ^^^^^^^^                                    ^^^^^^^^^
```

Therapeutic : Vadastuximab

Best Alignment of Vadastuximab heavy chain to a sequence from OAS

```
QVQLVQSGAEVKKPGASVKVSCKASGYTFTNYDINWVRQAPGQGLEWIGWIYPGDGSTKYNEKFKAKATLTADTSTSTAYMELRSLRSDDTAVYYCASGYE------DAMDYWGQGTTVTVSS
||||.|||.|..|||||||.||||||||||.||||||.|.|||||||||||||.|||||||||||.|||||.|||.|||||||.||.|.|.||||||.||........|||||||||||||||
QVQLQQSGPELVKPGASVKLSCKASGYTFTSYDINWVKQRPGQGLEWIGWIYPRDGSTKYNEKFKGKATLTVDTSSSTAYMELQSLTSEDSAVYYCARGYYGSSYEGNAMDYWGQGTTVTVSS
                         ^^^^^^^^                 ^^^^^^^^                                    ^^^^^^^^^^^^^^^^
```

Best Alignment of Vadastuximab light chain to a sequence from OAS

```
DIQMTQSPSSLSASVGDRVTINCKASQDINSYLSWFQQKPGKAPKTLIYRANRLVDGVPSRFSGSGSGQDYTLTISSLQPEDFATYYCLQYDEFPLTFGGGTKVEIK
||||||||||||||.|.||||.||||||||||||||||||||.||||||||||||||||||||||||||||.||||||..||...||||||||||.|||||||.|||
DIQMTQSPSSLSASLGERVTITCKASQDINSYLSWFQQKPGKSPKTLIYRANRLVDGVPSRFSGSGSGQDYSLTISSLEYEDMGIYYCLQYDEFPYTFGGGTKLEIK
                         ^^^^^^                 ^^^                                   ^^^^^^^^^
```

Best Alignment of Vadastuximab heavy chain CDRs to a sequence from OAS

```
QVQLVQSGAEVKKPGASVKVSCKASGYTFTNYDINWVRQAPGQGLEWIGWIYPGDGSTKYNEKFKAKATLTADTSTSTAYMELRSLRSDDTAVYYCASGYEDAMDYWGQGTTVTVSS
.|||.|||.|..||||.||.||||||||||.||||||.|.|||||||||||||||||||||||||.|||||||.|.|||||.|.||.|...|||.|||.|.|||||||||||.|||||
```

```
-VQLQQSGPELVKPGALVKISCKASGYTFTSYDINWVKQRPGQGLEWIGWIYPGDGSTKYNEKFKGKATLTADKSSSTAYMQLSSLTSENSAVYFCASNYADAMDYWGQGTSVTVSS
                         ^^^^^^^^                 ^^^^^^^^                                      ^^^^^^^^^^
Best Alignment of Vadastuximab light CDRs chain to a sequence from OAS
DIQMTQSPSSLSASVGDRVTINCKASQDINSYLSWFQQKPGKAPKTLIYRANRLVDGVPSRFSGSGSGQDYTLTISSLQPEDFATYYCLQYDEFPLTFGGGTKVEIK
||.||||||||..||.|.||||.|||||||||||||||||||.|||||||||||||||||||||||||||||.||||||..||...|||||||||||||||||||.|||
DIKMTQSPSSMYASLGERVTITCKASQDINSYLSWFQQKPGKSPKTLIYRANRLVDGVPSRFSGSGSGQDYSLTISSLEYEDMGIYYCLQYDEFPLTFGGGTKLEIK
                          ^^^^^^               ^^^                                    ^^^^^^^^^

Best Alignment of Vadastuximab CDR-H3 chain to a sequence from OAS
QVQLVQSGAEVKKPGASVKVSCKASGYTFTNYDINWVRQAPGQGLEWIGWIYPGDGSTKYNEKFKAKATLTADTSTSTAYMELRSLRSDDTAVYYCASGYEDAMDYWGQGTTVTVSS
.|||.|.|||..|||||||.||||||||||.|...||.|.||.||||||.|.|..|.||||||.|||||.|.|...|||||.|.||.|..|||||||||||||||||||||.|||||
-VQLQQPGAELVKPGASVKLSCKASGYTFTSYWMHWVKQRPGRGLEWIGRIDPNSGGTKYNEKFKSKATLTVDKPSSTAYMQLSSLTSEDSAVYYCASGYEDAMDYWGQGTSVTVSS
                         ^^^^^^^^                 ^^^^^^^^                                      ^^^^^^^^^^

Therapeutic : Varlilumab

Best Alignment of Varlilumab heavy chain to a sequence from OAS
QVQLVESGGGVVQPGRSLRLSCAASGFTFSSYDMHWVRQAPGKGLEWVAVIWYDGSNKYYADSVKGRFTISRDNSKNTLYLQMNSLRAEDTAVYYCARGSGN---WGFFDYWGQGTLVTVSS
||||||||||||||||||||||||||||||||.|||||||||||||||||||||||||||||||||||||||||||||||||||||||||||||||||||||....|..||||||||||||||
QVQLVESGGGVVQPGRSLRLSCAASGFTFSSYGMHWVRQAPGKGLEWVAVIWYDGSNKYYADSVKGRFTISRDNSKNTLYLQMNSLRAEDTAVYYCARGSGYSSSWYYFDYWGQGTLVTVSS
                         ^^^^^^^^                 ^^^^^^^^                                      ^^^^^^^^^^^^^^^

Best Alignment of Varlilumab light chain to a sequence from OAS
DIQMTQSPSSLSASVGDRVTITCRASQGISRWLAWYQQKPEKAPKSLIYAASSLQSGVPSRFSGSGSGTDFTLTISSLQPEDFATYYCQQYNTYPRTFGQGTKVEIK
||||||||||||||||||||||||||||||.|||||||||||||||||||||||||||||||||||||||||||||||||||||||||||||.||||||||||||||
DIQMTQSPSSLSASVGDRVTITCRASQGISSWLAWYQQKPEKAPKSLIYAASSLQSGVPSRFSGSGSGTDFTLTISSLQPEDFATYYCQQYNSYPRTFGQGTKVEIK
                          ^^^^^^               ^^^                                    ^^^^^^^^^

Best Alignment of Varlilumab heavy chain CDRs to a sequence from OAS
QVQLVESGGGVVQPGRSLRLSCAASGFTFSSYDMHWVRQAPGKGLEWVAVIWYDGSNKYYADSVKGRFTISRDNSKNTLYLQMNSLRAEDTAVYYCARGSGNWGFFDYWGQGTLVTVSS
||||||||||||||||||||||||||||||||.|||||||||||||||||||||||||||.||||||||||||||||||||||||||||||||||||||||.|.|||||||||||||||
QVQLVESGGGVVQPGRSLRLSCAASGFTFSSYGMHWVRQAPGKGLEWVAVIWYDGSNKYYGDSVKGRFTISRDNSKNTLYLQMNSLRAEDTAVYYCARGSGSWRFFDYWGQGTLVTVSS
                         ^^^^^^^^                 ^^^^^^^^                                      ^^^^^^^^^^^^

Best Alignment of Varlilumab light CDRs chain to a sequence from OAS
DIQMTQSPSSLSASVGDRVTITCRASQGISRWLAWYQQKPEKAPKSLIYAASSLQSGVPSRFSGSGSGTDFTLTISSLQPEDFATYYCQQYNTYPRTFGQGTKVEIK
|||||||||||||||||||||||||||||||||.||||||..|||||.||||.|||||||||||||||||||||||||||||||||.|||||||||||||||||..|||
DIQMTQSPSSLSASVGDRVTITCRASQGISRWLVWYQQKPGEAPKSLSYAASRLQSGVPSRFSGSGSGTDFTLTISSLQPEDFAIYYCQQYNTYPRTFGQGTRLEIK
                          ^^^^^^               ^^^                                    ^^^^^^^^^

Best Alignment of Varlilumab CDR-H3 chain to a sequence from OAS
QVQLVESGGGVVQPGRSLRLSCAASGFTFSSYDMHWVRQAPGKGLEWVAVIWYDGSNKYYADSVKGRFTISRDNSKNTLYLQMNSLRAEDTAVYYCARGSGNWGFFDYWGQGTLVTVSS
.............||.||||||||||||||||.||||||||||||||||||.||||||||||||||||||||||||||.||||||||||||||||||||||.|||||||||||||||||
-------------PGGSLRLSCAASGFTFSSYAMHWVRQAPGKGLEWVAVISYDGSNKYYADSVKGRFTISRDNSKNTVYLQMNSLRAEDTAVYYCARGSGYWGFFDYWGQGTLVTVSS
                         ^^^^^^^^                 ^^^^^^^^                                      ^^^^^^^^^^^^

Therapeutic : Vatelizumab

Best Alignment of Vatelizumab heavy chain to a sequence from OAS
QVQLQESGPGLVKPSETLSLTCTVSGFSLTNYGIHWIRQPPGKGLEWLGVIWARGFTNYNSALMSRLTISKDNSKNQVSLKLSSVTAADTAVYYCARANDGV---YYAMDYWGQGTLVTVSS
|||||.||||||.||..||.||||||||||.||.||.|||||||||||||||.|.|||||||||.||||||.||.||..|....|||.|||||||||....|||||||||||.|||||
QVQLQQSGPGLVAPSQSLSITCTVSGFSLTSYGVHWVRQPPGKGLEWLGVIWAGGSTNYNSALMSRLSISKDNSKSQVFLKMNSLQTDDTAMYYCARANDGYYADYYAMDYWGQGTSVTVSS
                         ^^^^^^^^                 ^^^^^^^                                      ^^^^^^^^^^^^^^^^

Best Alignment of Vatelizumab light chain to a sequence from OAS
DFVMTQSPAFLSVTPGEKVTITCSAQSSV-NYIHWYQQKPDQAPKKLIYDTSKLASGVPSRFSGSGSGTDYTFTISSLEAEDAATYYCQQWTTNPLTFGQGTKVEIK
..|||||||||||||||||||||.|.....||..|||||||||||.||...|...|||||||||||||||.|||||||||||||||||||....|||||.|||||||
--VMTQSPAFLSVTPGEKVTITCQASEGIGNYLYWYQQKPDQAPKLLIKYASQSISGVPSRFSGSGSGTDFTFTISSLEAEDAATYYCQQGNKHPLTFGGGTKVEIK
                          ^^^^^^               ^^^                                    ^^^^^^^^^

Best Alignment of Vatelizumab heavy chain CDRs to a sequence from OAS
QVQLQESGPGLVKPSETLSLTCTVSGFSLTNYGIHWIRQPPGKGLEWLGVIWARGFTNYNSALMSRLTISKDNSKNQVSLKLSSVTAADTAVYYCARANDGVYYAMDYWGQGTLVTVSS
.|.|.|||||||..|..||.||||||||||.||.||.|||||||||||||||.|.||||.||||||.|||||||.||.||..|....|||.||||||.||.|||||||||||.|||||
-VKLVESGPGLVATSQSLSITCTVSGFSLTSYGVHWVRQPPGKGLEWLGVIWAGGSTNYNLALMSRLSISKDNSKSQVFLKMNSLQTDDTAMYYCARAYDGYYYAMDYWGQGTSVTVSS
                         ^^^^^^^^                 ^^^^^^^                                      ^^^^^^^^^^^^^

Best Alignment of Vatelizumab light CDRs chain to a sequence from OAS
DFVMTQSPAFLSVTPGEKVTITCSAQSSVNYIHWYQQKPDQAPKKLIYDTSKLASGVPSRFSGSGSGTDYTFTISSLEAEDAATYYCQQWTTNPLTFGQGTKVEIK
..|||||||..|...||.||..|.|.|||||..|||||....||..||||||||||||.|||||||||.|..||||.|||||||||||||..||||||.|||.|||
-IVMTQSPAIMSASLGERVTMSCRASSSVNYMYWYQQKSGTSPKRWIYDTSKLASGVPARFSGSGSGTSYSLTISSMEAEDAATYYCQQWSSNPLTFGGGTKLEIK
                          ^^^^^               ^^^                                    ^^^^^^^^^

Best Alignment of Vatelizumab CDR-H3 chain to a sequence from OAS
QVQLQESGPGLVKPSETLSLTCTVSGFSLTNYGIHWIRQPPGKGLEWLGVIWAR-GFTNYNSALMSRLTISKDNSKNQVSLKLSSVTAADTAVYYCARANDGVYYAMDYWGQGTLVTVSS
.||||.||..||||.....|.||.|||....|..||..|....||||.|.|....|.|.|........||..|.|.|...|.|||.|..|||||||||||||.|||||||||||.|||||
-VQLQQSGAELVKPGASVKLSCTASGFNIKDYYMHWVKQRTEQGLEWIGRIDPEDGETKYAPKFQGKATITADTSSNTAYLQLSSLTSEDTAVYYCARANDGEYYAMDYWGQGTSVTVSS
                         ^^^^^^^^                 ^^^^^^^^                                      ^^^^^^^^^^^^^

Therapeutic : Vedolizumab

Best Alignment of Vedolizumab heavy chain to a sequence from OAS
QVQLVQSGAEVKKPGASVKVSCKGSGYTFTSYWMHWVRQAPGQRLEWIGEIDPSESNTNYNQKFKGRVTLTVDISASTAYMELSSLRSEDTAVYYCARGGYDG------WDYAIDYWGQGTLVTVSS
|||||||||||||||||||||||.||||||||.|||||||||||||||.|.|.....||.|.|||.||||.|.|.||||||||||||||||||||||||||||||.........|..|||||||||||||
QVQLVQSGAEVKKPGASVKVSCKASGYTFTSYAMHWVRQAPGQRLEWMGWINAGNGNTKYSQKFQGRVTITRDTSASTAYMELSSLRSEDTAVYYCARGGYDYVWGSYITQYYFDYWGQGTLVTVSS
                         ^^^^^^^^                 ^^^^^^^^                                      ^^^^^^^^^^^^^^^^^^^^

Best Alignment of Vedolizumab light chain to a sequence from OAS
DVVMTQSPLSLPVTPGEPASISCRSSQSLAKSYGNTYLSWYLQKPGQSPQLLIYGISNRFSGVPDRFSGSGSGTDFTLKISRVEAEDVGVYYCLQGTH-QPYTFGQGTKVEIK
|||||||||||||||||||||||||||||..|.|..||.||||||||||||||..|||.|||||||||||||||||||||||||||||||||.||||..|||||||||.|||
DVVMTQSPLSLPVTPGEPASISCRSSQSLLHSNGYNYLDWYLQKPGQSPQLLIYLGSNRASGVPDRFSGSGSGTDFTLKISRVEAEDVGVYYCMQGTHWPPYTFGQGTKLEIK
                          ^^^^^^^^^^^               ^^^                                    ^^^^^^^^^^

Best Alignment of Vedolizumab heavy chain CDRs to a sequence from OAS
QVQLVQSGAEVKKPGASVKVSCKGSGYTFTSYWMHWVRQAPGQRLEWIGEIDPSESNTNYNQKFKGRVTLTVDISASTAYMELSSLRSEDTAVYYCARGGYDGWDYAIDYWGQGTLVTVSS
.|||.|.|||..|||||||.|||.||||||||||.||.|.|||.||||||||||.|.|||||||||..|||||.|.|||||.||||.|||.||||||||||||.|||.|||||||.|||||
-VQLQQPGAELVKPGASVKLSCKASGYTFTSYWMQWVKQRPGQGLEWIGEIDPSDSYTNYNQKFKGKATLTVDTSSSTAYMQLSSLTSEDSAVYYCARGGYDGEDYAMDYWGQGTSVTVSS
                         ^^^^^^^^                 ^^^^^^^^                                      ^^^^^^^^^^^^^^

Best Alignment of Vedolizumab light CDRs chain to a sequence from OAS
DVVMTQSPLSLPVTPGEPASISCRSSQSLAKSYGNTYLSWYLQKPGQSPQLLIYGISNRFSGVPDRFSGSGSGTDFTLKISRVEAEDVGVYYCLQGTHQPYTFGQGTKVEIK
.||.||.||||||..|....|||||||||||.|||||||||||.|||||||||||||||||||||||||||||||||||||||||....||.|.|||||||||||||||.|||.|||
-VVVTQTPLSLPVSFGAQVPISCRSSQSLANSYGNTYLSWYLHKPGQSPQLLIYGISNRFSGVPDRFSGSGSGTDFTLKISTIKPEDLGMYYCLQGTHQPYTFGGGTKLEIK
                          ^^^^^^^^^^^               ^^^                                    ^^^^^^^^^
Best Alignment of Vedolizumab CDR-H3 chain to a sequence from OAS
QVQLVQSGAEVKKPGASVKVSCKGSGYTFTSYWMHWVRQAPGQRLEWIGEIDPSESNTNYNQKFKGRVTLTVDISASTAYMELSSLRSEDTAVYYCARGGYDGWDYAIDYWGQGTLVTVSS
.|.|..||....|||.|.|.||..||.||.|.||||.|..||||...|......|.|....|||.|...|......|...|||||||||.||||||||||.|||.|||||||.|||||
```

```
EVKLMESGGGLVKPGGSLKLSCAASGFTFSSYAMSWVRQTPEKRLEWVASISSG-GSTYYPDSVKGRFTISRDNARNILYLQMSSLRSEDTAMYYCARGGYDGADYAMDYWGQGTSVTVSS
                         ^^^^^^^^        ^^^^^^^^                    ^^^^^^^^^^^^^^
```

Therapeutic : Veltuzumab

Best Alignment of Veltuzumab heavy chain to a sequence from OAS

```
QVQLQQSGAEVKKPGSSVKVSCKASGYTFTSYNMHWVKQAPGQGLEWIGAIYPGMGDTSYNQKFKGKATLTADESTNTAYMELSSLRSEDTAFYYCARSTYYG-GDWYFDVWGQGTTVTVSS
||||||||||..|||.|||.|||||||||||||||||||.||||||||||||||.||||||||||||||||||.|..||||.||||.|||.|.||||||.|||...||||||||.||||||||
QVQLQQSGAELVKPGASVKMSCKASGYTFTSYNMHWVKQTPGQGLEWIGAIYPGNGDTSYNQKFKGKATLTADKSSSTAYMQLSSLTSEDSAVYYCARSHYYGSSYWYFDVWGAGTTVTVSS
                         ^^^^^^^^        ^^^^^^^^                    ^^^^^^^^^^^^^^^
```

Best Alignment of Veltuzumab light chain to a sequence from OAS

```
DIQLTQSPSSLSASVGDRVTMTCRASSSV-SYIHWFQQKPGKAPKPWIYATSNLASGVPVRFSGSGSGTDYTFTISSLQPEDIATYYCQQWTSNPPTFGGGTKLEIK
|||.||||||||||||||||.|||||.||..|.||||||||||||..|||.|||.||||.||||||||||.|.|||||||||.|||||||..|.|.|||.|||||||
DIQMTQSPSSLSASVGDRVTITCRASQSVSTYLHWFQQKPGKAPKLLIYAASNLQSGVPSRFSGSGSGTDFTLTISSLQPEDFATYYCQQSYSTPGTFGQGTKLEIK
                          ^^^^^^       ^^^                 ^^^^^^^^^
```

Best Alignment of Veltuzumab heavy chain CDRs to a sequence from OAS

```
QVQLQQSGAEVKKPGSSVKVSCKASGYTFTSYNMHWVKQAPGQGLEWIGAIYPGMGDTSYNQKFKGKATLTADESTNTAYMELSSLRSEDTAFYYCARSTYYGGDWYFDVWGQGTTVTVSS
.|||||||||...||.|||.||||||||||||||||||||.|.|||||||||||||.||||||||||||||||.|.|..||||.||||.|||.|.|||||..|||||||||||||.||||||||
-VQLQQSGAELVRPGASVKMSCKASGYTFTSYNMHWVKQTPRQGLEWIGAIYPGNGDTSYNQKFKGKATLTVDKSSSTAYMQLSSLTSEDSAVYYCARDGYYGGDWYFDVWGTGTTVTVSS
                         ^^^^^^^^        ^^^^^^^^                    ^^^^^^^^^^^^^^
```

Best Alignment of Veltuzumab light CDRs chain to a sequence from OAS

```
DIQLTQSPSSLSASVGDRVTMTCRASSSVSYIHWFQQKPGKAPKPWIYATSNLASGVPVRFSGSGSGTDYTFTISSLQPEDIATYYCQQWTSNPPTFGGGTKLEIK
.|.|||||...|||.|..|||||.|||||||.||.|||.|..||.|||||||||||||.|||||||||.|..|||....||.||||||||.|||||||.|||||.|
QIVLTQSPAIMSASPGEKVTMTCSASSSVSYMHWYQQKSGTSPKRWIYATSNLASGVPARFSGSGSGTSYSLTISRMEAEDAATYYCQQWSSNPPTFGAGTKLELK
                          ^^^^^       ^^^                 ^^^^^^^^^
```

Best Alignment of Veltuzumab CDR-H3 chain to a sequence from OAS

```
QVQLQQSGAEVKKPGSSVKVSCKASGYTFTSYNMHWVKQAPGQGLEWIGAIYPGMGDTSYNQKFKGKATLTADESTNTAYMELSSLRSEDTAFYYCARSTYYGGDWYFDVWGQGTTVTVSS
.|.|..||....|||.|.|.||.|||.||.||.|.||.|.|...|||...|..|...|.|....||..|...|...|..|...|.|.|||||.|||||||||||.||||||||.||||||||
EVKLMESGGGLVKPGGSLKLSCAASGFTFSSYAMSWVRQTPEKRLEWVATISDGGSYTYYPDNVKGRFTISRDNAKNNLYLQMSHLKSEDTAMYYCARSTYYGYDWYFDVWGTGTTVTVSS
                         ^^^^^^^^        ^^^^^^^^                    ^^^^^^^^^^^^^^
```

Therapeutic : Visilizumab

Best Alignment of Visilizumab heavy chain to a sequence from OAS

```
QVQLVQSGAEVKKPGASVKVSCKASGYTFISYTMHWVRQAPGQGLEWMGYINPRSGYTHYNQKLKDKATLTADKSASTAYMELSSLRSEDTAVYYCARSAYYDYDGFAYWGQGTLVTVSS
||||.|||||...||||||.|||||||||.||||||||.|.||||||||.|||||.||||.||||.||||||||||||.||||||.||||.|||.|||||||||.|||||||||||||||||||||.
QVQLQQSGAELARPGASVKMSCKASGYTFTSYTMHWVKQRPGQGLEWIGYINPSSGYTKYNQKFKDKATLTADKSSSTAYMQLSSLTSEDSAVYYCARSVYYDYDGFAYWGQGTLVTVSA
                         ^^^^^^^^        ^^^^^^^^                    ^^^^^^^^^^^^^
```

Best Alignment of Visilizumab light chain to a sequence from OAS

```
DIQMTQSPSSLSASVGDRVTITCSASSSV-SYMNWYQQKPGKAPKRLIYDTSKLASGVPSRFSGSGSGTDFTLTISSLQPEDFATYYCQQWSSNPPTFGGGTKVEIK
||||||||||||||||||||||||.||.|..||.|||||||||||||.|||.||.|.||||||||||||||||||||||||||||||||||||..|.||||||||||||||
DIQMTQSPSSLSASVGDRVTITCRASQSISSYLNWYQQKPGKAPKLLIYATSSLQSGVPSRFSGSGSGTDFTLTISSLQPEDFATYYCQQSYSTPPTFGGGTKVEIK
                          ^^^^^^       ^^^                 ^^^^^^^^^
```

Best Alignment of Visilizumab heavy chain CDRs to a sequence from OAS

```
QVQLVQSGAEVKKPGASVKVSCKASGYTFISYTMHWVRQAPGQGLEWMGYINPRSGYTHYNQKLKDKATLTADKSASTAYMELSSLRSEDTAVYYCARSAYYDYDGFAYWGQGTLVTVSS
.|||.||||||...||||||.||||||||||.||||||||.|.||||||||.|||||.||||.||||.||||||||||||.||||||.||||.|||.|.|||||||.|||||||||||||||||||||.
-VQLQQSGAELARPGASVKMSCKASGYTFTSYTMHWVKQRPGQGLEWIGYINPSSGYTKYNQKFKDKATLTADKSSSTAYMQLSSLTSEDSAGYYCARSVYYDYDGFAYWGQGTLVTVSA
                         ^^^^^^^^        ^^^^^^^^                    ^^^^^^^^^^^^^
```

Best Alignment of Visilizumab light CDRs chain to a sequence from OAS

```
DIQMTQSPSSLSASVGDRVTITCSASSSVSYMNWYQQKPGKAPKRLIYDTSKLASGVPSRFSGSGSGTDFTLTISSLQPEDFATYYCQQWSSNPPTFGGGTKVEIK
.|..||||...|||.|..||.||||||||||||.||||||||..|..||||||.||||||.||||||||||...||||||...||.||||||||||||||||||||||.|||
QIVLTQSPAIMSASPGEKVTMTCSASSSVSYMYWYQQKPGSSPRLLIYDTSNLASGVPVRFSGSGSGTSYSLTISSMEAEDAATYYCQQWSSNPPTFGGGTKLEIK
                          ^^^^^       ^^^                 ^^^^^^^^^
```

Best Alignment of Visilizumab CDR-H3 chain to a sequence from OAS

```
QVQLVQSGAEVKKPGASVKVSCKASGYTFISYTMHWVRQAPGQGLEWMGYINPRSGYTHYNQKLKDKATLTADKSASTAYMELSSLRSEDTAVYYCARSAYYDYDGFAYWGQGTLVTVSS
.|||.||||||..||||||||.||.|||.....|.||||.|...|||||.|.|.|..|.|.|..|....||.|.|.|..|||..||.|.||||||||||||||||||||||||||||||||||.
EVQLQQSGAELVKPGASVKLSCTASGFNINDYYMHWVKQRTEQGLEWIGRIDPEEGETKYAPKFQGNATITSDTSSNTAYLQLSRLTSEDTAVYYCARSAYYDYDGFAYWGQGTLVTVSA
                         ^^^^^^^^        ^^^^^^^^                    ^^^^^^^^^^^^^
```

Therapeutic : Vonlerolizumab

Best Alignment of Vonlerolizumab heavy chain to a sequence from OAS

```
EVQLVQSGAEVKKPGASVKVSCKASGYTFTDSYMSWVRQAPGQGLEWIGDMYPDNGDSSYNQKFRERVTITRDTSTSTAYLELSSLRSEDTAVYYCVLAPR-------WYFSVWGQGTLVTVSS
.||||||||||||||||||||||||||||||..||.|||||||||||||||...|..|..||.|||..|||.|||||||||.|.||||||||||||||||||..|||.........|..||||||||||||
QVQLVQSGAEVKKPGASVKVSCKASGYTFTSYYMHWVRQAPGQGLEWIGIINPSGGSTSYAQKFQGRVTMTRDTSTSTVYMELSSLRSEDTAVYYCARAPRITMIVVVLLFDYWGQGTLVTVSS
                         ^^^^^^^^        ^^^^^^^^                    ^^^^^^^^^^^^^^^^^
```

Best Alignment of Vonlerolizumab light chain to a sequence from OAS

```
DIQMTQSPSSLSASVGDRVTITCRASQDISNYLNWYQQKPGKAPKLLIYYTSRLRSGVPSRFSGSGSGTDFTLTISSLQPEDFATYYCQQGHTLPPTFGQGTKVEIK
|||||||||||||||||||||||||||.||||||||||||||||||.||||.||.|.|||||||||||||||||||||||||||||||||||||||..|||.||||||||||||
DIQMTQSPSSLSASVGDRVTITCRASQSISNYLNWYQQKPGKAPNLLIYATSSLQSGVPSRFSGSGSGTDFTLTISSLQPEDFATYYCQQCFTLPWTFGQGTKVEIK
                          ^^^^^^       ^^^                 ^^^^^^^^^
```

Best Alignment of Vonlerolizumab heavy chain CDRs to a sequence from OAS

```
EVQLVQSGAEVKKPGASVKVSCKASGYTFTDSYMSWVRQAPGQGLEWIGDMYPDNGDSSYNQKFRERVTITRDTSTSTAYLELSSLRSEDTAVYYCVLAPRWYFSVWGQGTLVTVSS
.|||.|||.|..||||||||.||||||||||||...|||.|..|||||||||..||..|...||.||....|.|.|.|..|||..||||.|||.|||.|||.||||||.|||.||.||||||
QVQLQQSGPELVKPGASVKMSCKASGYTFTDYVISWVKQRTGQGLEWIGEIYPGSGSTYYNEKFKGKATLTADKSSNTAYMQLSSLTSEDSAVYFCVLRPRWYFDVWGAGTTVTVSS
                         ^^^^^^^^        ^^^^^^^^                    ^^^^^^^^^^
```

Best Alignment of Vonlerolizumab light CDRs chain to a sequence from OAS

```
DIQMTQSPSSLSASVGDRVTITCRASQDISNYLNWYQQKPGKAPKLLIYYTSRLRSGVPSRFSGSGSGTDFTLTISSLQPEDFATYYCQQGHTLPPTFGQGTKVEIK
||||.|...|||||.|.|.||.||||||||||||||||||||||....|.|||||||||.|||.|.||||||||||||..|||||||..||.|||.|||||.||||||||..||...|
DIQMIQTTYSLSASFGGRATISCRASQDISNYLNWYQQKPDGTVKHLIYYTSRLHSGVTSKFSGSGSGTDYSLTISSLEQEDIATYFCQQGNTLPPTFGARTKLGLK
                          ^^^^^^       ^^^                 ^^^^^^^^^
```

Best Alignment of Vonlerolizumab CDR-H3 chain to a sequence from OAS

```
EVQLVQSGAEVKKPGASVKVSCKASGYTFTDSYMSWVRQAPGQGLEWIGDMYPDNGDSSYNQKFRERVTITRDTSTSTAYLELSSLRSEDTAVYYCVLAPRWYFSVWGQGTLVTVSS
.|||.|||.|..||||||||.||||||||||||...|||.|..|||||||||..||..|...||.||....|.|.|.|..|||..||||.|||.|||.|||.||||||.|||.||.||||||
QVQLQQSGPELVKPGASVKMSCKASGYTFTDYVISWVKQRTGQGLEWIGEIYPGSGSTYYNEKFKGKATLTADKSSNTAYMQLSSLTSEDSAVYFCVLRPRWYFDVWGAGTTVTVSS
                         ^^^^^^^^        ^^^^^^^^                    ^^^^^^^^^^
```

Therapeutic : Zalutumumab

Best Alignment of Zalutumumab heavy chain to a sequence from OAS

```
QVQLVESGGGVVQPGRSLRLSCAASGFTFSTYGMHWVRQAPGKGLEWVAVIWDDGSYKYYGDSVKGRFTISRDNSKNTLYLQMNSLRAEDTAVYYCARDGITMVRGVMKDYFDYWGQGTLVTVSS
||||||||||||||||||||||||||||||.||||||||||||||||||||||||.|||.|||.||||||||||||||||||||||||||||||||||||||||||||||.||||||||...||||||||||||||||
QVQLVESGGGVVQPGRSLRLSCAASGFTFSSYGMHWVRQAPGKGLEWVAVIWYDGSNKYYADSVKGRFTISRDNSKNTLYLQMNSLRAEDTAVYYCARDPITMVRGVITTYFDYWGQGTLVTVSS
                         ^^^^^^^^        ^^^^^^^^                    ^^^^^^^^^^^^^^^^^
```

Best Alignment of Zalutumumab light chain to a sequence from OAS

```
AIQLTQSPSSLSASVGDRVTITCRASQDISSALVWYQQKPGKAPKLLIYDASSLESGVPSRFSGSESGTDFTLTISSLQPEDFATYYCQQFNSYPLTFGGGTKVEIK
|||||||||||||||||||||||||||.|||||||||||||||||||||||||||||||||||||.|||||||||||||||||||||||||||||||||||||||||
AIQLTQSPSSLSASVGDRVTITCRASQGISSALVWYQQKPGKAPKLLIYDASSLESGVPSRFSGSGSGTDFTLTISSLQPEDFATYYCQQFNSYPLTFGGGTKVEIK
                          ^^^^^^                 ^^^                                    ^^^^^^^^^
```

Best Alignment of Zalutumumab heavy chain CDRs to a sequence from OAS

```
QVQLVESGGGVVQPGRSLRLSCAASGFTFSTYGMHWVRQAPGKGLEWVAVIWDDGSYKYYGDSVKGRFTISRDNSKNTLYLQMNSLRAEDTAVYYCARDGITMVRGVMKDYFDYWGQGTLVTVSS
.|.|||||||||||.|||||||||||||||||||||||.||||||||||||.|||.|||.|||||||||||||||||||||||||||||||||||||||||||||||.|.||.|||||.|||||
-VRLVESGGGVVQPGGSLRLSCAASGFTFSTYGMHWVRQSPGKGLEWVAVIWYDGSNKYYADSVKGRFTISRDNSKNTLYLQMNSLRAEDTAVYYCARDGITMVRGVMADAFDIWGQGTMVTVSS
                         ^^^^^^^^                 ^^^^^^^^                                      ^^^^^^^^^^^^^^^^^
```

Best Alignment of Zalutumumab light CDRs chain to a sequence from OAS

```
AIQLTQSPSSLSASVGDRVTITCRASQDISSALVWYQQKPGKAPKLLIYDASSLESGVPSRFSGSESGTDFTLTISSLQPEDFATYYCQQFNSYPLTFGGGTKVEIK
||||||||||||||||||||||||.||||||||.|||||||||||||||||||||||||||||||.|||||||||||||||||||||||||||||||||||||||||
AIQLTQSPSSLSASVGDRVTITCRTSQDISSALAWYQQKPGKAPKLLIYDASSLESGVPSRFSGSGSGTDFTLTISSLQPEDFATYYCQQFNSYPLTFGGGTKVEIK
                          ^^^^^^                 ^^^                                    ^^^^^^^^^
```

Best Alignment of Zalutumumab CDR-H3 chain to a sequence from OAS

```
QVQLVESGGGVVQPGRSLRLSCAASGFTFSTYGMHWVRQAPGKGLEWVAVIWDDGSYKYYGDSVKGRFTISRDNSKNTLYLQMNSLRAEDTAVYYCARDGITMVRGVMKDYFDYWGQGTLVTVSS
..............|.||||||||||||||.|.|.|||||||||||||.||.......||.||||||||||||||||||||||||||||||||||||||||||||||..||||||||||||||||
--------------GGSLRLSCAASGFTFSSYAMSWVRQAPGKGLEWVSVISSG-GSTYYADSVKGRFTISRDNSKNTLYLQMNSLRAEDTAVYYCARDGITMVRGVISDYFDYWGQGTLVTVSS
                         ^^^^^^^^                 ^^^^^^^^                                      ^^^^^^^^^^^^^^^^^
```

Therapeutic : Zanolimumab

Best Alignment of Zanolimumab heavy chain to a sequence from OAS

```
QVQLQQWGAGLLKPSETLSLTCAVYGGSFSGYYWSWIRQPPGKGLEWIGEINHSGSTNYNPSLKSRVTISVDTSKNQFSLKLSSVTAADTAVYYCARVIN-------------WFDPWGQGTLVTVSS
||||||||||||||||||||||||||||||||||||||||||||||||||||||||||||||||||||||||||||||||||||||||||||||||||.|.............|||||||||||||||
QVQLQQWGAGLLKPSETLSLTCAVYGGSFSGYYWSWIRQPPGKGLEWIGEINHSGSTNYNPSLKSRVTISVDTSKNQFSLKLSSVTAADTAVYYCARVANVVVPAARRKMGWGWFDPWGQGTLVTVSS
                         ^^^^^^^^                 ^^^^^^^                                      ^^^^^^^^^^^^^^^^^^^^^
```

Best Alignment of Zanolimumab light chain to a sequence from OAS

```
DIQMTQSPSSVSASVGDRVTITCRASQDISSWLAWYQHKPGKAPKLLIYAASSLQSGVPSRFSGSGSGTDFTLTISSLQPEDFATYYCQQANSFPYTFGQGTKLEIK
|||||||||||||||||||||||||||||||||||||.|||||||||||||||||||||||||||||||||||||||||||||||||||||||||||||||||||||
DIQMTQSPSSVSASVGDRVTITCRASQDISSWLAWYQQKPGKAPKLLIYAASSLQSGVPSRFSGSGSGTDFTLTISSLQPEDFATYYCQQANSFPYTFGQGTKLEIK
                          ^^^^^^                 ^^^                                    ^^^^^^^^^
```

Best Alignment of Zanolimumab heavy chain CDRs to a sequence from OAS

```
QVQLQQWGAGLLKPSETLSLTCAVYGGSFSGYYWSWIRQPPGKGLEWIGEINHSGSTNYNPSLKSRVTISVDTSKNQFSLKLSSVTAADTAVYYCARVINWFDPWGQGTLVTVSS
............|||||||||||||||||||||||||||||||||||||||||||||||||||||||||||||||||||||||||||||||||||||||||||||||||||||||
------------KPSETLSLTCAVYGGSFSGYYWSWIRQPPGKGLEWIGEINHSGSTNYNPSLKSRVTISVDTSKNQFSLKLSSVTAADTAVYYCARVINWFDPWGQGTLVTVSS
                         ^^^^^^^^                 ^^^^^^^                                      ^^^^^^^^^
```

Best Alignment of Zanolimumab light CDRs chain to a sequence from OAS

```
DIQMTQSPSSVSASVGDRVTITCRASQDISSWLAWYQHKPGKAPKLLIYAASSLQSGVPSRFSGSGSGTDFTLTISSLQPEDFATYYCQQANSFPYTFGQGTKLEIK
||||||||||||||.|||.||||||||||||||||||.|||.||||||||||.|..|||.||||.|||||||||||||||||||||||||||||||||||||..||
DIQMTQSPSSVSASIGDRLTITCRASQDISSWLAWYQQKPGQAPKLLIYAASRLRTGVPTRFSGGGSGTDFTLTISSLQPEDFATYYCQQANSFPYTFGQGTKVDIK
                          ^^^^^^                 ^^^                                    ^^^^^^^^^
```

Best Alignment of Zanolimumab CDR-H3 chain to a sequence from OAS

```
QVQLQQWGAGLLKPSETLSLTCAVYGGSFSGYYWSWIRQPPGKGLEWIGEINHSGSTNYNPSLKSRVTISVDTSKNQFSLKLSSVTAADTAVYYCARVINWFDPWGQGTLVTVSS
..............||||||||.|.|||.|.||||||||||||||||||.|..|||||||||||||||||||||||||||||.||||||||||||||||||||||||||||||||
--------------SETLSLTCTVSGGSISSYYWSWIRQPPGKGLEWIGYIYYSGSTNYNPSLKSRVTISVDTSKNQFSLKLNSVTAADTAVYYCARVINWFDPWGQGTLVTVSS
                         ^^^^^^^^                 ^^^^^^^                                      ^^^^^^^^^
```

Therapeutic : Zolbetuximab

Best Alignment of Zolbetuximab heavy chain to a sequence from OAS

```
QVQLQQPGAELVRPGASVKLSCKASGYTFTSYWINWVKQRPGQGLEWIGNIYPSDSYTNYNQKFKDKATLTVDKSSSTAYMQLSSPTSEDSAVYYCTRSWRG--NSFDYWGQGTTLTVSS
||||||||||||||||||||||||||||||||||||||||||||||||||||||||||||||||||||||||||||||||||||||||||||||||||||....|.||||||||||||||
QVQLQQPGAELVRPGASVKLSCKASGYTFTSYWINWVKQRPGQGLEWIGNIYPSDSYTNYNQKFKDKATLTVDKSSSTAYMQLSSPTSEDSAVYYCTRSWGNYENYFDYWGQGTTLTVSS
                         ^^^^^^^^                 ^^^^^^^^                                      ^^^^^^^^^^^^^
```

Best Alignment of Zolbetuximab light chain to a sequence from OAS

```
DIVMTQSPSSLTVTAGEKVTMSCKSSQSLLNSGNQKNYLTWYQQKPGQPPKLLIYWASTRESGVPDRFTGSGSGTDFTLTISSVQAEDLAVYYCQNDYSYPFTFGSGTKLEIK
|||||||||||||||||||||||||||||||||||||||||||||||||||||||||||||||||||||||||||||||||||||||||||||||||||||||||||||||||
DIVMTQSPSSLTVTAGEKVTMSCKSSQSLLNSGNQKNYLTWYQQKPGQPPKLLIYWASTRESGVPDRFTGSGSGTDFTLTISSVQAEDLAVYYCQNDYSYPFTFGSGTKLEIK
                          ^^^^^^^^^^^^                 ^^^                                    ^^^^^^^^^
```

Best Alignment of Zolbetuximab heavy chain CDRs to a sequence from OAS

```
QVQLQQPGAELVRPGASVKLSCKASGYTFTSYWINWVKQRPGQGLEWIGNIYPSDSYTNYNQKFKDKATLTVDKSSSTAYMQLSSPTSEDSAVYYCTRSWRGNSFDYWGQGTTLTVSS
.|.|...|||||||||||||||||||||||||||||||||||||||||||||||||||||||||||||||||||||||||||||||||||||||||...|||||||||||||||||
-VHLVESGAELVRPGASVKLSCKASGYTFTSYWINWVKQRPGQGLEWIGNIYPSDSYTNYNQKFKDKATLTVDKSSSTAYMQLSSPTSEDSAVYYCTRRTYGNSFDYWGQGTTLTVSS
                         ^^^^^^^^                 ^^^^^^^^                                      ^^^^^^^^^^^
```

Best Alignment of Zolbetuximab light CDRs chain to a sequence from OAS

```
DIVMTQSPSSLTVTAGEKVTMSCKSSQSLLNSGNQKNYLTWYQQKPGQPPKLLIYWASTRESGVPDRFTGSGSGTDFTLTISSVQAEDLAVYYCQNDYSYPFTFGSGTKLEIK
|||||||||||||||||||||||||||||||||||||||||||||||||||||||||||||||||||||||||||||||||||||||||||||||||||||||||.|||||.|
DIVMTQSPSSLTVTAGEKVTMSCKSSQSLLNSGNQKNYLTWYQQKPGQPPKLLIYWASTRESGVPDRFTGSGSGTDFTLTISSVQAEDLAVYYCQNDYSYPFTFGAGTKLELK
                          ^^^^^^^^^^^^                 ^^^                                    ^^^^^^^^^
```

Best Alignment of Zolbetuximab CDR-H3 chain to a sequence from OAS

```
QVQLQQPGAELVRPGASVKLSCKASGYTFTSYWINWVKQRPGQGLEWIGNIYPSDSYTNYNQKFKDKATLTVDKSSSTAYMQLSSPTSEDSAVYYCTRSWRGNSFDYWGQGTTLTVSS
..................|.|..||....||...|..|.||.||||||.||.|...||||...|...|..||.|.......|||.|..|.|||||.||.||||||||||||..||||
--------------SETLSLTCTVSGGSISSYYWSWIRQPPGKGLEWIGYIYYS-GSTNYNPSLKSRVTISVDTSKNQFSLKLSSVTAADTAVYYCARSSRGNSFDYWGQGTLVTVSS
                         ^^^^^^^^                 ^^^^^^^^                                      ^^^^^^^^^^^
```

### References


1. GaBI Journal Editor. 2017. Patent expiry dates for biologicals: 2016 update. *Generics Biosimilars Initiat. J.* .
